# Discovery of Novel Targets for Important Human and Plant Fungal Pathogens via Automated Computational Pipeline HitList

**DOI:** 10.1101/2024.10.22.619685

**Authors:** David E. Condon, Brenda K. Schroeder, Paul A. Rowley, F. Marty Ytreberg

## Abstract

Fungi are a major threat to human health and agricultural productivity, causing 1.7 million human deaths and billions of dollars in crop losses and spoilage annually. While various antifungal compounds have been developed to combat these fungi in medical and agricultural settings, but there are concerns that treatment effectiveness is waning due to the emergence of acquired drug resistance and to novel pathogens. Effectiveness is further hampered due to the limited number of modes of action for available antifungal compounds.To develop new strategies for the control and mitigation of fungal disease and spoilage, new antifungals are needed with novel fungal-specific protein targets that can overcome resistance and prevent host toxicity. New antifungals can add new methods of targeting fungi that have no effective control measures. The increasing availability of complete genomes of pathogenic and spoilage fungi has enabled identification of novel protein and RNA targets essential for viability and not found in host plants or humans. In this study, an automated bioinformatics pipeline utilizing BLAST, ClustalΩ, and subtractive genomics was created and used to identify potential new targets for any system of hosts and pathogens with available genomic or proteomic data. This pipeline called HitList allows *in silico* screening of thousands of possible targets. HitList was then used to generate a list of potential antifungal targets for the World Health Organization fungal priority pathogens list and the top 10 agricultural fungal pathogens. Known antifungal targets were found, validating the approach, and an additional eight novel protein targets were discovered that could be used for the rational design of antifungal compounds.

## 1 Introduction

Fungal infections in humans can range from mild subcutaneous and mucosal infections[1] to life-threatening systemic disease. In particular, the rise of sophisticated medical devices, interventions, and therapeutics has increased vulnerable and immunocompromised patient populations that tend to be more susceptible to mycoses.[2, 3] As a result, fungal diseases are the 5th largest cause of human mortality.[4]. Populations that are especially vulnerable to fungal infections include prematurely born children[5] and HIV,[6] leukemia,[7] and organ transplant patients.[8] Fungal infections can be a heavy cost for the health care system.[9, 10] In 2018, 666,235 fungal infections were diagnosed in the United States alone.[11] Invasive fungal diseases can have high mortality rates, depending on the etiological agent. Specifically, *Candida* yeasts are a leading cause of nosocomial fungal infections, particularly *Candida albicans*, which is a common fungal species associated with human mucosal surfaces.[12] Recent outbreaks of mucomycoses associated with the COVID-19 pandemic have also highlighted the lack of effective fungal treatments, with mortality rates *>*40%.[13] Non-life-threatening infections include mucosal infections by *Candida* yeasts. Specifically, vulvovaginal candidiasis affects 75% of women at least once in their lifetime and is a major cause of morbidity, especially due to recurrent infections caused by drug-resistant species of yeasts.[14] The World Health Organization (WHO) recently published a list of the most concerning fungal pathogens,[15] highlighting the global concern over fungal infection.

Agriculture incurs substantial losses from many different fungal pathogens in all regions of the world.[16] For example *Pyricularia oryzae* destroying 10-35% of rice, *Ustilago maydis*[17] destroying 2-20% of corn.[18] Fungal pathogens of agriculture such as *Puccina*,[19] *Botrytis*,[20] and others[21] can be devastating to yields of many different crops[22] and endanger food security among a growing population.[23] Fungal pathogens such as *Blumeria graminis* are obligate plant pathogens, even having several specialized forms adapted to parasitize specific plants.[24] Globalization can also introduce new fungal pathogens to agriculture from different regions of the world, potentially introducing new fungi for which plants and farmers have no experience,[25] such as *Mycosphaerella graminicola*, which spread to the Americas and Australia over the last 500 years.[26] *Colletotrichum truncatum* has a very wide host range, including soybeans,[27] pepper, eggplant, muskmelon, chickpea, grapes, and tomatoes.[28] In addition, the production of mycotoxins by fungi can contaminate agricultural products,[29] and threaten human and animal health if consumed.[30, 31]Infection by pathogens such such as *Pyricularia oryzae* and *Fusarium* species, that can cause devastating economic losses[32] costing farmers billions of dollars annually[33] in antifungal, i.e., fungicide, treatments. In addition to acting as direct pathogens, some fungi, such as saprophytes, can spoil food stores.[34] Use of fungicides in agriculture also creates resistance in *Aspergillus* to azole drugs,[35] used for both agricultural and human medicinal applications,[36] linking agricultural and human pathogens. The repeated use of fungicides with limited number of modes of action results in an increase in resistant strains within fungal populations that are pathogens of humans or agriculture.[37, 38] It is widely understood that there is a critical need for new classes of antifungal compounds.[39, 40] Developing antifungal compounds with new modes of action could provide more options and extend the window of efficacy for current treatments, leading to new uses in the case of any contraindication to a particular drug. One key limiting factor in the development of new antifungal drugs is the lack of novel protein targets.[41] While screening methods can identify new antimicrobial compounds without knowledge of the molecular target,[42] an ideal compound should maximize harm against the pathogen and minimize off-target effects for the host human or plant. This can be accomplished by designing compounds to bind targets that are absent in the host, which can be difficult due to the shared eukaryotic ancestry between fungi, plants and humans, and shared biochemical pathways between humans, plants, and fungi.[43] However, differences in biochemistry between fungi, humans, and plants present opportunities for drug targeting. For example, within fungal systems, ergosterol serves many of the same cellular functions that cholesterol serves in humans, but it is not present in animals, hence proteins in the ergosterol synthesis pathway have been targeted by many different antifungals (Table S1). Similarly, azoles targeting 14α-demethylase[44, 45] and morpholines targeting Erg2 and Erg24 in the ergosterol synthesis pathway are commonly used antifungals.[46] (Table S1). Likewise, the echinocandins target the pathway of cell wall β-(1,3) glucan synthesis,[47]but are exclusively used against human and animal infections.[48, 49] As the cell wall is a unique structure to fungi, this pathway is absent in humans and leads to fungal osmotic instability and death.[50] Unfortunately, many of these antifungal compounds also have undesirable properties (Table S1). Azoles are used against both human and agricultural pathogens,[51] and must be used for extended periods to be effective. Despite their potent activity against fungal pathways of ergosterol synthesis, they have been associated with hepatotoxicity and numerous hormone-related effects.[52] In addition, polyenes such as amphotericin B have severe renal toxicity.[53, 54] Proliferation of resistance to current antifungal compounds further highlights the need for new modes of action.[51] Examples include *Candida*’s resistance to echinocandins[55] and flucytosine,[56] azole resistance in *Aspergillus*[57] and *Candida*[58, 59, 60] and fungal species that are resistant to multiple types of antifungal drugs.[61]Some other minor antifungals exist in addition to the major classes mentioned above.[62]

The most common method for developing antifungal compounds involves high-throughput screening of large chemical libraries to identify possible antifungal candidates[63]. This process is expensive and time-consuming, with the added challenge that the cellular target of the compound is unknown. Computer-aided drug design, or CADD[64] has also been used in the design and development of antifungals to treat human disease and it is significantly cheaper and faster for screening large libraries of compounds. Virtual screening is performed by using docking strategies to simulate the binding of a ligand with a given protein target[65], then applying various scoring functions to estimate the protein-ligand binding strength.[66] CADD has been used to identify further inhibitors of known drug targets, but has yet to produce a viable drug.[67] At least 70 human drugs have been approved that used virtual screening as part of the drug discovery process, including Captopril, Norfloxacin, and Imatinib, but anti-fungals were not among them.[68].

In this study, a bioinformatics pipeline, HitList, was developed and used to identify possible targets of antimicrobials by applying it to determine antifungal protein targets. The purpose was to discover novel antifungal targets for the World Health Organization critical fungal pathogen list and the top 10 agricultural fungal pathogens. Recent advances in the computational determination of protein structure are also accelerating the pace of drug discovery due to the increased availability of protein structures. The current study started with a list of essential genes from the model fungus *Saccharomyces cerevisiae*. Each of 201 essential genes in *S. cerevisiae* was used to identify orthologous proteins in humans, plants, and fungal pathogens. Subtractive genomics[69] was then used to identify protein regions that could serve as good targets.HitList surveyed more than a thousand proteins. HitList was validated by identifying proteins that already have known antifungal inhibitors, but we also found eight proteins that had not been previously considered as antifungal targets.

## 2 Methods

### 2.1 Source Data and Resources

Protein sequences encoded by essential genes from the Database for Essential Genes (DEG)[70] for *S. cerevisiae* (yeast) were downloaded from http://essentialgene.org. The pathogen targets were identified by utilizing the World Health Organization critical pathogen list[71] (Table S3) and the top 10 agricultural fungal pathogen list (Table S4)[32]. Proteome data was downloaded for common agricultural organisms that can be hosts for fungi *Homo sapiens* (human), *Glycine max* (soy), *Oryza sativa* (rice), *Solanum tuberosum* (potato), and *Zea mays* (corn) from https://ftp.ncbi.nlm.nih.gov/genomes/refseq/ (Table S2). Finally, pathogen proteomes were downloaded from from https://ftp.ncbi.nlm.nih.gov and fungidb.org (Tables S3 and S4).

### 2.2 Bioinformatics Pipeline

The pipeline starts with all genes from *S. cerevisiae* as obtained from the DEG. Of the 1,110 that were considered essential, 909 genes that have human orthologs according to the Saccharomyces Genome Database (SGD)[72] were eliminated from consideration (Figure 1). The remaining 201 genes were used as queries in a BLASTP analysis using default parameters[73] (version 2.14) against all hosts and pathogens proteomes. Proteins with homology to query proteins having an expectation value above 0.1 were excluded. For each query protein, the top resulting protein from each pathogen and each host BLAST analysis was placed together and aligned with Clustal 1.2.4[74] to obtain multiple sequence alignments (MSAs). MSA regions that show alignment with pathogens and not with hosts are considered potentially targetable, using an approach that has previously been called “subtractive genomics.”[69]. This approach identifies amino acid sequences that are likely only present in the pathogens, and absent in the hosts, avoiding effects on the host. Analgous to a study showing that amino acid sequence could determine whether or not an immunoglobulin E protein would bind to casein for milk-allergic patients,[75] we sought to identify amino acid regions of proteins that would be susceptible to any sort of potential inhibitor or antibody.

**Figure 1:**
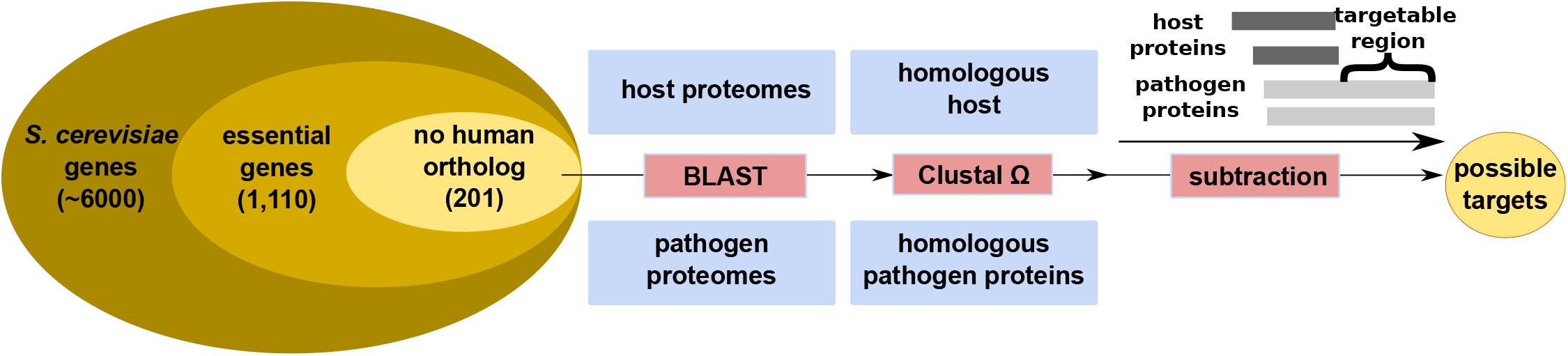
Graphical representation of the bioinformatics pipeline that identifies potentially targetable proteins. Considering only genes that have no human ortholog according to the Saccharomyces Genome Database (SGD)[72], the field of essential genes is narrowed to 201. These genes are then used as BLAST[77] queries against all pathogen and host proteomes (Section S1). ClustalΩ was then used to create MSAs to identify targetable regions according via subtractive genomics.[69]

Fig 2 shows an example comparing two proteins, one with a poor targetability, and another with excellent targetability. The targetability of a region is quantified by the number of pathogens present at each residue in the MSA, the more pathogen proteins are present within a given region, the better that region is for a target. Proteins with known experimental, e.g., NMR or X-ray, structures are preferred, as eventual structure-based drug design[76] would have a more reliable starting point for design of anti-fungal compounds. Additionally, where the protein is located within the cell is also important, as drugs must be able to penetrate the cell membrane.

**Figure 2:**
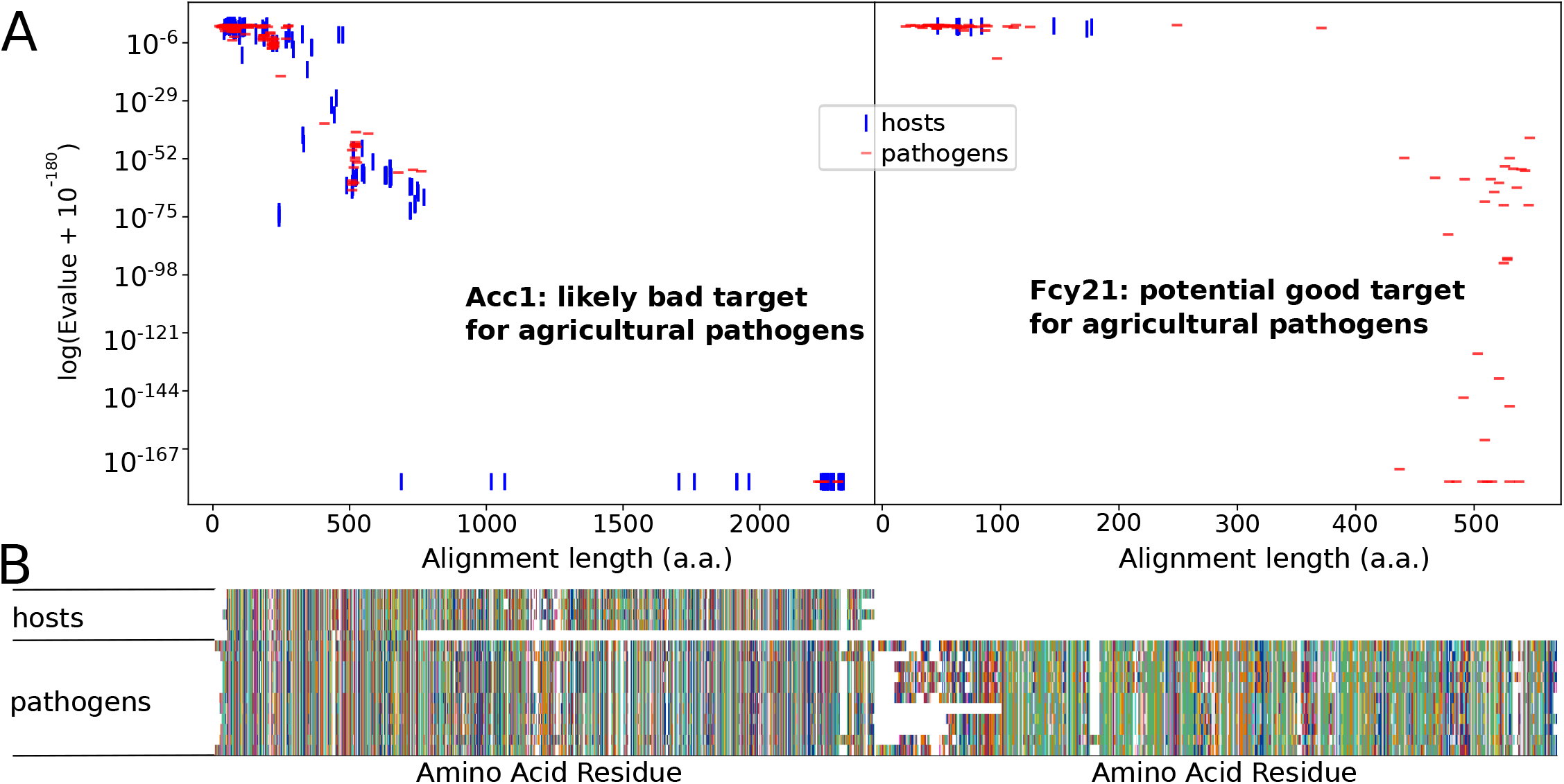
An example of two essential yeast proteins analyzed for target suitability among the top 10 agricultural fungal pathogens. The figure shows results from BLAST searches performed for each essential gene with alignment statistics shown. **(A)** Acc1 is an example of a bad target that shows high similarity between homologous proteins of hosts and pathogens, while on the right Fcy21 is an example of a potentially good target that shows neither strong expectation values nor long alignment lengths with host proteins. **(B)** The strong alignment between host and pathogen proteins for Acc1, and the lack of host proteins for Fcy21 is visible in multiple sequence alignments, which are visualized using a modified version of CIAlign[78].

The similarity of the amino acids within each MSA was quantified by Sneath’s similarity index[79] (Table S6, section S3).Sneath’s φ was chosen because of its intuitive nature, in that it ranks within (0,1], and an amino acid compared with itself can be exactly 1. The Sneath index is calculated at each position within an MSA by comparing every amino acid in a group, against every other amino acid in that group:

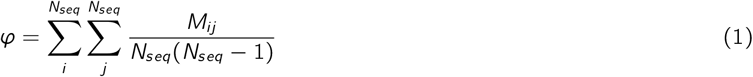

where *M* values for two amino acids *i* and *j* are given by the Sneath similarity in Table S6, and *N*_*seq*_ is the number of sequences at that position. For example, consider two peptides, each containing nine alanine residues. In this case, all Sneath similarity values would be 1.0. Comparing a 9-mer of alanine against a 9-mer of cysteine would give a Sneath similarity of 0.87 across the 9 residues. Uncertain amino acids, e.g. “X” are set to the minimum Sneath value of 0.003 to avoid type II errors. If a known amino acid is compared with an ambiguous amino acid, e.g. A with B, where B could be either D or N, then the Sneath index of A and B is set at the mean of *M*_*A/D*_ and *M*_*A/N*_.

## 3 Results & Discussion

In this study, our newly developed pipeline HitList was used to identify target proteins against two groups of pathogens: WHO fungal pathogens (Table S3), and agricultural pathogens (Table S4). Hosts of the WHO fungal pathogens are humans only. By contrast, hosts for the agricultural pathogens includes the plants and the humans that consume the plants (Table S2). Using *S. cerevisiae* as a model fungus, we considered 201 essential genes that do not have human orthologs according to the SGD[72, 95] (Fig. 1). No similar database exists for plants, so plant orthologs were not considered. BLASTP[73] searches were conducted of 20 fungal pathogen proteomes in Section S1 using 201 protein queries of essential yeast proteins with no annotated human homologs (Fig. 1). Top hits from each pathogen and host were combined into a single MSA for each protein using ClustalΩ.[74] Regions of pathogen proteins that do not match any host were then selected as possible targets.

Pathogen proteins that have relatively strong bit scores and alignment length to an essential protein for *S. cerevisiae* and not to any host protein, while having strong homogeneity as measured by φ, i.e. approximately within the range 0.7 ≤ *φ* ≤ 1, are considered excellent possible targets (e.g. “Good Target” in Fig. 2) Conversely, proteins that have good alignment between both host and pathogen proteins and to the *S. cerevisiae* essential protein, especially over the entire length, are considered less desirable targets (e.g. “Bad Target” in Fig. 2). The longer the alignment length, and the more pathogens show strong alignment and φ, the more potential the protein could have as a target for antifungal compounds as more distinct surfaces and structural features will be present that can be targeted by CADD. That is, proteins in the top right corners of Fig. 3 represent the best preliminary targets.

**Figure 3:**
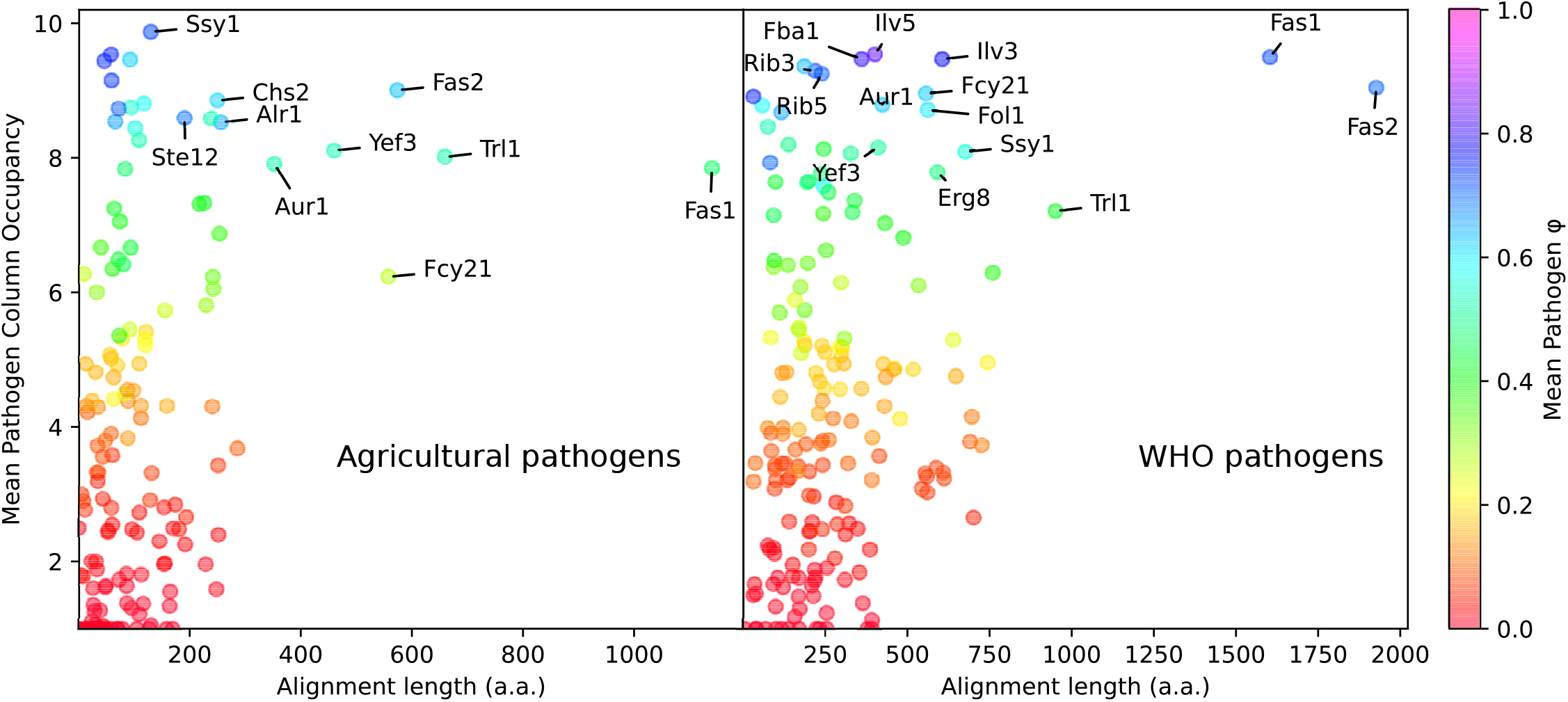
The output of the pipeline can be visualized by a scatterplot of alignment length, mean pathogen column occupancy, and φ. All proteins tested for targeting are plotted by targetable length and number of pathogens. The best protein targets are labeled.

Table 2 summarizes our list of most promising potential protein targets. Of the 16 proteins in the table, seven are in common between WHO and agricultural fungal pathogens and thus could be targets that lead to the development of broad spectrum antifungal compounds. “Broad-spectrum” here refers to a drug effectively targeting multiple genera of pathogen organisms that are not closely related to one another. While the potential target list is greater than these 16 proteins, these are the most visually conspicuous in Fig 3) and hence further analysis will focus on the proteins listed in Table 2. A larger number of possible protein targets were found for the WHO pathogens than for agricultural pathogens, likely because agricultural hosts include human and also a wide phylogenetic range of plants, as compared to WHO pathogens that only include human as a host.

**Table 1:** The host and pathogen species used in this study are listed above, selecting economically important plants and their pathogens. Hosts used for agricultural pathogens are indicated by ^A, W^ indicates WHO pathogens, and human is considered for both groups. Source data for each species is indicated in Table S1.

| Hosts | Human WHO Pathogens | Ag. Pathogens |
| --- | --- | --- |
| *Glycine max* (soy)<sup>A</sup><br>*Homo sapiens* (human)<sup>AW</sup><br>*Oryza sativa* (rice)<sup>A</sup><br>*Solanum tuberosum* (potato)<sup>A</sup><br>*Zea mays* (maize, or corn)<sup>A</sup> | *Aspergillus fumigatus*<br>*Candida albicans*<br>*Candida auris*<br>*Candida parapsilosis*<br>*Candida tropicalis*<br>*Cryptococcus neoformans* JEC21<br>*Cryptococcus neoformans* B 3501A<br>*Cryptococcus neoformans* grubii H99<br>*Histoplasma capsulatum*<br>*Nakaseomyces glabratus* | *Blumeria graminis*<br>*Botrytis cinerea*<br>*Colletotrichum truncatum*<br>*Fusarium graminearum*<br>*Mycosphaerella graminicola*<br>*Puccinia graminis*<br>*Puccinia striiformis*<br>*Puccinia triticina*<br>*Pyricularia oryzae*<br>*Ustilago maydis* |

**Table 2:** *S. cerevisiae* proteins that show high potential for antifungal targeting for pathogens in the agricultural and WHO pathogen lists (labeled in Figure 3). Superscript ^W^ indicates that the protein is only a good target with the WHO pathogens, superscript ^A^ indicates that the protein is only a good target for agricultural pathogens and no superscript means it is a good target for both lists. The UniProt IDs are provided in the second column. The third column contains information summarized from the SGD and UniProt.[72, 94] More detailed information is available in Section S4.

| Protein | UniProt ID | Length (a.a.) | Summary | Lit. |
| --- | --- | --- | --- | --- |
| Alr1<sup>AW</sup> | Q08269 | 859 | Transmembrane protein involved in $Mg^{2+}$ transport | [80] |
| Aur1<sup>AW</sup> | P36107 | 401 | Inositol phosphoceramide synthase involved in sphingolipid biosynthesis | [81] |
| Chs2<sup>A</sup> | P14180 | 963 | Chitin synthase II; catalyzes transfer of N-acetylglucosamine to chitin | [82, 83, 84] |
| Erg8<sup>W</sup> | P24521 | 451 | Phosphomevalonate kinase; contributes to ergosterol biosynthesis | none |
| Fas1<sup>AW</sup> | P07149 | 2051 | $\beta$ Subunit of cytoplasmic fatty acid synthase; complex catalyzes the synthesis of long-chain saturated fatty acids | [85, 86, 87] |
| Fas2<sup>AW</sup> | P19097 | 1887 | $\alpha$ Subunit of cytoplasmic fatty acid synthase; complex catalyzes the synthesis of long-chain saturated fatty acids | [85, 86, 87] |
| Fba1<sup>W</sup> | P14540 | 359 | Fructose 1,6-bisphosphate aldolase; required for gluconeogenesis and glycolysis | [88] |
| Fcy21 | P40039 | 528 | Nucleobase transporter; moves nucleobases through plasma membrane | none |
| Fol1<sup>W</sup> | P53848 | 824 | Multifunctional enzyme of the folic acid biosynthesis pathway | [89] |
| Ilv3<sup>W</sup> | P39522 | 585 | Mitochondrial dihydroxy-acid dehydratase for branched-chain amino acid biosynthesis | none |
| Ilv5<sup>W</sup> | P06168 | 395 | Mitochondrial ketol-acid reductoisomerase for branched-chain amino acid biosynthesis | none |
| Rib3<sup>W</sup> | Q99258 | 208 | Cytosolic synthase involved in riboflavin biosynthesis | none |
| Rib5<sup>W</sup> | P38145 | 238 | Riboflavin synthase; catalyzes the last step of the riboflavin biosynthesis pathway | none |
| Ssy1<sup>AW</sup> | Q03770 | 852 | senses external amino acid concentration; transmits intracellular signals that regulate expression of a.a. permease genes | none |
| Ste12<sup>A</sup> | P13574 | 688 | A sequence-specific DNA binding transcription factor for genes involved in mating and filamentous growth | none |
| Trl1<sup>AW</sup> | P09880 | 827 | RNA ligase (ATP) and GTP-dependent RNA kinase; role in mediating nonsense-mediated decay | [90, 91, 92] |
| Yef3<sup>AW</sup> | P16521 | 1044 | Translation elongation factor and ATPase involved in translational elongation and termination | [41, 93] |

### Identification of Previously Identified Targets Validates the Pipeline

Figure 3 shows our pipeline is working, as it identifies previously known targets that already have known inhibitors. Our method picked out Fas1 and Fas2 (Sections S4.5 × S4.6); these subunits of fatty acid synthase[96] have been previously identified as viable targets against fungi, and inhibitors have already been found,[85, 86, 87] serving as a validation of our pipeline. Fas1 is the specific target of the antifungal compound NPD6433.[86, 97] Our analysis suggests Fas2 is a better target for WHO pathogens compared to agricultural pathogens, because host plant proteins show weakly aligned sequences, and fungal sequences also align more weakly (Figs S93 and S97). Indeed, the natural products CT2108A and CT2108B are effective antifungals against *Candida, S. cerevisiae*, and *Cryptococcus*, yet both natural products are made by the fungus *Penicillium solitum*. One protein of *P. solitum* also has an evalue of 0 against S. cerevisiae Fas1 and Fas2, and yet *P. solitum* isn’t affected by its own natural product, so good alignment alone is not an indicator that an inhibitor will have the same activity. Fas1 BLAST hits for the plant pathogens *P. graminis, P. triticina*, and *P. striiformis* are much weaker than for other agricultural pathogens, as demonstrated by the lower φ value in Fig. 3. BLAST searches against the non-redundant protein database show that Fas1 is broadly conserved across many different genera of fungi from Asomycota, Basidomycota, and Zoopagomycota (Fig. S92) and Fas2 (Fig. S106). Table 2 also shows Chitin synthase 2 (Chs2) as a possible agricultural target, and this protein already has known inhibitors including nikkomycins and polyoxins.[82, 83, 84] However, as the human protein hyaluronan synthase has a strong BLAST alignment, Chs2 is not an optimum choice for drug targeting. Nonetheless, chitin synthase (cf. Section S4.3) has no equivalent in humans or plants according to the SGD, agreeing with a search against the NCBI’s non-redundant database[98] (NR) (cf. Figs. S60 × S65). Even if Chs2 were successfully targeted, a Chs2-knockout strain of the human pathogen *C. albicans* did not have attenuated virulence,[99] so targeting Chs2 with potential therapeutics may not be effective in treatment of infections. Finally, the plasma membrane Mg^2+^ transporter Alr1[100] was identified in the pipeline, which has been shown to be a potential antifungal target and is known to be inhibited by Bovine pancreatic trypsin inhibitor.[80]

HitList also identified some proteins that have been previously suggested as potential antifungal targets in the literature but have no known inhibitors. Proteins that are part of chemical pathways that are absent in hosts are especially attractive, as the chance of side effects is significantly lower. For example, folate synthesis (Fol1)[101], ceramide phosphoinositol transferase, part of sphingolipid synthesis (Aur1),[102] and fructose 1,6-bisphosphate aldolase (Fba1)[88] were identified as potential antifungal targets by this analysis and others, and do not have mammalian equivalents.[81, 102] Folate synthesis is an effective target of antibacterials, such as trimethoprim and sulfamethoxazole, [103] and may be effective in fungi too. The pipeline identified Trl1 responsible for splicing of introns from nascent tRNA as a potential target. Since this is done very differently in fungi compared to metazoa, Trl1 a possible antifungal target.[104, 92, 90, 91] Yeast Elongation factor 3 (Yef3 or eEF3) is very well conserved among fungi [41] and identified as a potential target using this pipeline. Because non-fungi use only two elongation factors when translating mRNA, as opposed to three for fungi, Yef3 has potential as a target[105, 93] Indeed, Figs. S251 × S256 both show strong conservation of Yef3 across the fungal kingdom, however, our study also reveals 960 metazoan species with strong hits (Fig S255). This make intuitive sense because Yef3 contains ATP-binding cassette domains[106] that are very common throughout many domains of life (Table S40), and inhibitors must be carefully designed to avoid unintentional effects on host organisms. Further, results suggest that Trl1 and Yef3 are less likely to be broad spectrum targets as can be seen with the relatively low φ and weaker MSA (Figs 3, S243, S231), compared to other proteins identified in the pipeline.

### Identification of Eight Novel Antifungal Protein Targets

Most importantly, our study has also uncovered a rich variety of novel possible protein targets that have not been previously identified to our knowledge (Table 2): Erg8, Fcy21, Ilv3, Ilv5, Rib3, Rib5, Ssy1, and Ste12. As will be detailed below, some of these targets are particularly well-suited for WHO pathogens and some for agricultural pathogens, while others appear to be candidates for antifungal development against both pathogen lists.

HitList identifies potential broad-spectrum antifungal target proteins that are particularly well-suited for treating pathogens in the WHO pathogen list. The protein pair Ilv3 and Ilv5,[107] involved in branched chain amino acid synthesis,[108, 109] is an especially attractive target because 1) no human proteins have strong alignments via BLAST, 2) there is a high φ value between pathogen hit sequences, and 3) branched chain amino acid synthesis does not occur in metazoa.[110] The last point is critical since inhibition of a protein pathway that is present in a pathogen, but not in a host, is unlikely to cause harm to the host. The dihydroxyacid dehydratase enzyme performs the same chemistry as Ilv3 in cyanobacteria and is inhibited by aspterric acid.[111] However, the broad-spectrum antifungal properties of aspterric acid are in doubt since it is a natural product of the fungus *Aspergillus terreus*.[112, 113] Further experimentation is necessary to determine whether aspterric acid or related derivatives have potential to function as a broad-spectrum antifungals. The pipeline also identified Rib3 and Rib5 as attractive protein targets for the WHO pathogen list, with high φ values. These proteins function in riboflavin synthesis, which does not occur in metazoa. Our study suggests that Ilv3, Ilv5, Rib3 and Rib5 are not as suitable of targets for developing agricultural antifungals use because riboflavin[114] and branched chain amino acid synthesis[109] also occurs in plants (see Figs S185 and S202).

Ssy1 is an amino acid sensor protein that has potential for broad-spectrum use against both agriculture and WHO pathogens, but appears to be better suited for agricultural application. This apparently novel antifungal target shows much stronger alignment with *Candida* pathogens than *Cryptococcus* (Fig S205), which reduces the φ value within WHO pathogens. Furthermore, *Nakaseomyces* shows great differences with *Candida* and *Cryptococcus* sequence hits with Ssy1 (Figs S204 and S205). BLAST hits against the Non-Redundant protein database animal and plant proteins are few and weak (Figs. S212 and S217), so any well-designed drug against Ssy1 would be unlikely to affect animals and plants. However, as a membrane protein, drug design need not be concerned with membrane penetration.

Fcy21, for “flucytosine resistance,” is another potential novel protein target for antifungal compounds, which has potential to be an antifungal target for WHO human pathogens. Fcy21 is a putative purine-cytosine permease.[115, 116] This protein is related to the target of flucytosine (Table 2) Fcy2p, but cannot substitute for its function. No human protein shows significant alignment with Fcy21 (Fig S121), but strong conservation is seen among pathogenic fungi, apart from the first 100 residues in the N-terminal domain. No non-fungal orthologs exist according to the SGD, and searches against the non-redundant database indicate strong conservation among WHO pathogenic fungi, and no hits among metazoa and insignificant hits among plants (Fig S133). Interestingly, Fcy21 does not have homology with proteins from several of the agricultural fungal pathogens of agricultural products (Fig. S125 × Table S22), hence Fcy21 is not likely to be as useful for developing agricultural antifungals. Ste12 was also shown as a possible target for WHO human pathogens, but is not labeled in Figure 3 as it has weak alignment with the model S. cerevisiae (Figs S218 and S222), and is overshadowed by other targets. Ste12, a transcription factor that is important for the virulence of mycoparasites, including *Trichoderma atroviride* which is a parasite of plant pathogens,[117, 118] shows good antifungal potential for the top 10 agricultural fungal pathogens (Section S4.15). Ste12 is one of the two proteins that is a good target against agricultural pathogens, but not against human pathogens (Figs S222 × S218).

Our study shows that agricultural host proteomes have much stronger hits to Erg8 (Fig. S70) than human (Fig. S66), so any antifungal drug targeting of Erg8 would likely be more effective against human pathogens. Erg8 is part of the ergosterol synthesis pathway, enabling phosphomevalonate kinase activity. While the ergosterol synthesis pathway is well-studied and targeted by azole[45, 119] and morpholine antifungals,[120] Erg8 appears to be a novel as a potential target. The difference in primary sequence of the Erg8 homolog from *Cryptococcus neoformans* compared to the homologs from other pathogen species is significant, with large gaps in the multiple sequence alignment because of *C. neoformans* (cf. Fig S66).

### Limitations of the Study

The primary limitation of the pipeline is it relies on the list of essential genes. Here, essential genes for *S. cerevisiae* from DEG that do not have human orthologs according to the SGD are used. If a gene is not on this list, this gene will For example, consider Fks1, the target of echinocandin antifungals. Fks1 does not appear on our list because the β-1,3-glucan synthase proteins (Fks1, Fks2, Fks3) are not considered essential genes according to the DEG.[70] Nonetheless, BLAST analysis of Fks1 against the non-redundant database identifies protein homologs in 1,149 fungal species (Fig. S262), and a multiple sequence alignment of these demonstrates that Fks1 is an ideal target (Fig S261), as there are very large regions of the protein in the MSA that only have alignment to pathogen protein domains. This shows that Fks1 would have appeared on our final list of target proteins if it was in the initial pool of genes considered. Such results should be expected, as fungal cell wall synthesis is unique to fungi.[121]

All species need to have a sequenced genome and proteome in order to be used

HitList is designed to avoid possible effects on the host, and does so at the possible risk of missing otherwise good targets, such as would have been targeted by the azoles. Erg11, the target of azole antifungals (Table S1), shows a human protein having nearly identical alignment as fungal proteins (Fig S257) which can be further seen with alignment bit scores (Fig S41). Indeed, human lanosterol and non-targeted cytochrome P450 are also affected by azole antifungals.[122] Similarly, Erg24 is a target of the morpholine antifungals, and shows a strong sequence similarity with delta(14)-sterol reductase, and has human orthologs according to SGD.[72] However, Fks1, Fks2/Gsc2, and Fks3 do not have any homologous human proteins, and did not show up as hits with this pipeline as they were not included in DEG.

HitList requires a well-characterized organism closely related to the target pathogens such as *S. cerevisiae*, with a sequenced genome, predicted and/or directly sequenced proteins, and a list of essential genes.The genomes of the pathogens must be sequenced and available, of sufficient quality, preferably with a predicted proteome. Our study was possible due to the very high quality genome annotation of *S. cerevisiae*, and the availability of the pathogen data from NCBI.

Identified target proteins should also be well-characterized structurally, e.g. by NMR or X-ray crystallography. Proteins should have known active sites to be used for computational chemistry docking programs, so that potential inhibitors can be screened.[123] Many proteins in Table 2, such as Ssy1 and Fcy21, do not have any experimental structures available as of publication, while proteins such as Yef3 does have an X-ray structure available.[93] Future research of possible protein targets should consider the presence and quality of existing structures.

### Diversity of Pathogens and Hosts Hinders Target Selection

The number of hosts/pathogens and their identities affects the possible protein targets and the length of the targetable regions identified by the pipeline. Consider Ssy1, which has a much longer targetable section for the WHO pathogens than for the top 10 agricultural pathogens (Fig 3 and Section S4.14). An MSA of the Ssy1 homologs from the top 10 agricultural pathogen and respective host proteomes shows large gaps, reducing the potential target length (Fig. S208). Gaps also exist for Ssy1 homologs from the WHO pathogen proteomes (Fig S208) in approximately the same regions as for the Ssy1 homologs from the top 10 agricultural pathogens, but the homologs from the *Candida* species and *N. glabratus* proteomes exhibit homology across these gaps, lengthening the potential region that could be targeted with antifungal compounds. For the agricultural pathogens, the plant proteins within the MSA shorten the targetable region. The larger the number of hosts and pathogens that are included in the analysis, and the more diversity within each group, the fewer and shorter the targets will be for antifungal compounds.

The phylogenetic diversity of the ascomycetes is very wide, [124] challenging identification of targets.Consider the extreme distance between the two ascomycete genera *Saccharomyces* and *Schizosaccharomyces*, which is on the order of 350 million years of separation.[125] However, previous phylogenetic analysis[124] shows that *S. cerevisiae* is a better choice than *Schizosaccharomyces* for the ascomycetes investigated here. Indeed, *Cryptococcus* and *Puccina* are both genera within basidomycota, which is in an entirely different phylum than *S. cerevisiae’*s ascomycota, and at an even greater evolutionary distance than between *Schizosaccharomyces* and *Saccharomyces*. No basidomycete has the quality of annotation that *S. cerevisiae* has and a known essential gene list. Nonetheless, potential protein targets were found that included the basidomycetes. The length of targetable proteins is reduced by increasing diversity of pathogens and hosts.Consider, for example, using only *Candida* pathogens in the WHO list for HitList. When this analysis is performed, Ccc1 is a protein that shows as targetable, but does not show as targetable when the entire WHO list is considered. Ccc1[126] (Section S6.1) is a Mn^2+^/Fe^2+^ transporter that has previously been identified as a potential antifungal target.[127] Indeed, Ccc1 is particularly attractive because no metazoan equivalent exists, and *Candida* is a human pathogen. *Cryptococcus* proteins clearly cluster differently from the other fungal genera (Figs S265 and S267), which is intuitive as *Cryptococcus* is a basidomycete genus, while all other pathogens are ascomycetes. *Candida* pathogen proteins clearly show both strong alignment to one another and to *S. cerevisiae*, while *Cryptococcus* shows its own cluster that doesn’t align strongly to *S. cerevisiae* (Fig. S266). Strong homology exists between the *Candida* proteins and *S. cerevisiae*, making more probable that this protein is essential in *Candida* as well, agreeing with phylogenetic distances in Figure S267. Inspection of Fig. S265 implies that the broad-spectrum targeting region is approximately from residue 110-170 in MSA coordinates, a narrow region of the protein to target with a compound. Given that malfunctioning immune antibodies can target decapeptides,[75] Ccc1 is a possible target for *Cryptococcus*, but development of an anti-fungal drug could be challenging. By contrast, using only *Candida* species as pathogens suggest that residues 100-340 are targetable (Fig. S265), which could make compound design simpler.

## Conclusion

A new, automated bioinformatics pipeline (HitList) that uses subtractive genomics to identify protein targets on a massively parallel scale for any pathogen-host combination was developed. HitList was applied to the discovery of novel antifungal targets for the WHO critical list of fungal pathogens and the top 10 agricultural fungal pathogens, using essential genes from *S. cerevisiae*.

Current anti-fungal targets, such as Fas1 and Fas2, and previously hypothesized anti-fungal targets such as Trl1 and Yef3 were identified as potential targets by HitList. Both lists of pathogens and hosts showed established targets. Many previously identified targets that were identified do not yet have inhibitors, such as Alr1, Aur1, and Trl1.

Most importantly, HitList identified novel protein targets that could be used to develop broad-spectrum antifungals that have not been previously mentioned in the literature, to our knowledge: Erg8, Fcy21, Ilv3, Ilv5, Rib3, Rib5, Ssy1, and Ste12. Based on our analysis, we anticipate that some of these targets will be more well-suited for developing antifungals against the WHO list (Erg8, Ilv3, Ilv5, Rib3, and Rib5), Ste12 against the agriculture list only, and others against both lists (Fcy21 and Ssy1). In addition, stronger potential targets are identified for the WHO/human pathogen list, since it has a single host, as compared to the agricultural top 10 list, that has five hosts across two different domains of life.

While HitList identifies proteins that could potentially be targets of compounds for antifungal purposes (Table 2), more research is clearly needed in order to validate these targets and develop possible inhibitors that could lead to new classes of antifungal drugs.

Finally, we note that HitList could be used to investigate any list of pathogens and hosts for any kingdom of life, provided that genomes and/or proteomes are available. The HitList software to run the bioinformatics pipeline and identify potential protein targets is freely available at https://github.com/YtrebergLab.

