## Supporting Information for "Discovery of Novel Targets for Important Human and Plant Fungal Pathogens via Automated Computational Pipeline HitList"

David E. Condon

October 22, 2024

#### Contents

**S1 Data Sources** **S10**

**S2 BLAST Expectation Value Histograms** **S11**
S2.1 WHO Pathogens . . . S11
S2.2 Agricultural Pathogens . . . S18

**S3 Amino Acid Similarity** **S24**

**S4 Best Targetable Proteins** **S24**
S4.1 Alr1 . . . S24
S4.1.1 WHO Critical Pathogens . . . S24
S4.1.2 Top 10 Agricultural Fungal Pathogens . . . S27
S4.1.3 NR . . . S31
S4.2 Aur1 . . . S35
S4.2.1 WHO Critical Pathogens . . . S35
S4.2.2 Top 10 Agricultural Fungal Pathogens . . . S38
S4.2.3 NR . . . S42
S4.3 Chs2 . . . S46
S4.3.1 WHO Critical Pathogens . . . S46
S4.3.2 Top 10 Agricultural Fungal Pathogens . . . S49
S4.3.3 NR . . . S53
S4.4 Erg8 . . . S59
S4.4.1 WHO Critical Pathogens . . . S59
S4.4.2 Top 10 Agricultural Fungal Pathogens . . . S62
S4.4.3 NR . . . S66
S4.5 Fas1 . . . S72
S4.5.1 WHO Critical Pathogens . . . S72
S4.5.2 Top 10 Agricultural Fungal Pathogens . . . S75
S4.5.3 NR . . . S79
S4.6 Fas2 . . . S83
S4.6.1 WHO Critical Pathogens . . . S83
S4.6.2 Top 10 Agricultural Fungal Pathogens . . . S86
S4.6.3 NR . . . S90
S4.7 Fba1 . . . S96
S4.7.1 WHO Critical Pathogens . . . S96
S4.7.2 Top 10 Agricultural Fungal Pathogens . . . S99
S4.7.3 NR . . . S103
S4.8 Fcy21 . . . S109
S4.8.1 WHO Critical Pathogens . . . S109
S4.8.2 Top 10 Agricultural Fungal Pathogens . . . S112
S4.8.3 NR . . . S116
S4.9 Fol1 . . . S120
S4.9.1 WHO Critical Pathogens . . . S120
S4.9.2 Top 10 Agricultural Fungal Pathogens . . . S123
S4.9.3 NR . . . S127

S4.10 Ilv3 . . . S133
S4.10.1 WHO Critical Pathogens . . . S133
S4.10.2 Top 10 Agricultural Fungal Pathogens . . . S136
S4.10.3 NR . . . S140
S4.11 Ilv5 . . . S146
S4.11.1 WHO Critical Pathogens . . . S146
S4.11.2 Top 10 Agricultural Fungal Pathogens . . . S149
S4.11.3 NR . . . S153
S4.12 Rib3 . . . S159
S4.12.1 WHO Critical Pathogens . . . S159
S4.12.2 Top 10 Agricultural Fungal Pathogens . . . S162
S4.12.3 NR . . . S166
S4.13 Rib5 . . . S172
S4.13.1 WHO Critical Pathogens . . . S172
S4.13.2 Top 10 Agricultural Fungal Pathogens . . . S175
S4.13.3 NR . . . S179
S4.14 Ssy1 . . . S185
S4.14.1 WHO Critical Pathogens . . . S185
S4.14.2 Top 10 Agricultural Fungal Pathogens . . . S188
S4.14.3 NR . . . S192
S4.15 Ste12 . . . S198
S4.15.1 WHO Critical Pathogens . . . S198
S4.15.2 Top 10 Agricultural Fungal Pathogens . . . S201
S4.15.3 NR . . . S205
S4.16 Trl1 . . . S209
S4.16.1 WHO Critical Pathogens . . . S209
S4.16.2 Top 10 Agricultural Fungal Pathogens . . . S212
S4.16.3 NR . . . S216
S4.17 Yef3 . . . S220
S4.17.1 WHO Critical Pathogens . . . S220
S4.17.2 Top 10 Agricultural Fungal Pathogens . . . S223
S4.17.3 NR . . . S227

**S5 Previously identified protein targets S233**
S5.1 Erg11 . . . S233
S5.2 Erg24 . . . S235
S5.3 Erg2 . . . S236
S5.4 Fks1 . . . S237
S5.5 Fks3 . . . S239
S5.6 Gsc2 . . . S240

**S6 Genus-specific Good Targets S241**
S6.1 Ccc1 . . . S241
S6.1.1 WHO Critical Pathogens . . . S241
S6.1.2 Aspergillus . . . S243
S6.1.3 Candida . . . S243
S6.1.4 Cryptococcus . . . S244
S6.1.5 Top 10 Agricultural Fungal Pathogens . . . S244

### List of Tables

S1 Major current human anti-fungal compound classes and their targets, all genes refer to *Saccharomyces cerevisiae*. S10
S2 Species and the data sources used for hosts. The Top 10 agricultural pathogens group used every species here, while the WHO pathogens only used *H. sapiens*. . . . S10
S3 World Health Organization fungal pathogens and the data sources used. . . . S10
S4 Top ten agricultural pathogens and the data sources used. . . . S11
S5 Pairwise alignment info from *S. cerevisiae* query proteins. . . . S24

S6 Similarity between amino acids can be quantified by Sneath's index $\phi$, where identical amino acids have a value of 1, and the greater the index between two amino acids is, the more similar the amino acids are to one another.. S24
S7 Pairwise alignment info from yeast Alr1 (DEG20010935), cf. Figure S26. . . . . S26
S8 Pairwise alignment info from yeast Alr1 (DEG20010935), cf. Figure S30. . . . . S29
S9 Pairwise alignment info from yeast Aur1 (DEG20010598), cf. Figure S39. . . . . S37
S10 Pairwise alignment info from yeast Aur1 (DEG20010598), cf. Figure S43. . . . . S40
S11 Pairwise alignment info from yeast Chs2 (DEG20010039), cf. Figure S52. . . . . S48
S12 Pairwise alignment info from yeast Chs2 (DEG20010039), cf. Figure S56. . . . . S51
S13 Pairwise alignment info from yeast Erg8 (DEG20010822), cf. Figure S66. . . . . S61
S14 Pairwise alignment info from yeast Erg8 (DEG20010822), cf. Figure S70. . . . . S64
S15 Pairwise alignment info from yeast Fas1 (DEG20010641), cf. Figure S80. . . . . S74
S16 Pairwise alignment info from yeast Fas1 (DEG20010641), cf. Figure S84. . . . . S77
S17 Pairwise alignment info from yeast Fas2 (DEG20011054), cf. Figure S93. . . . . S85
S18 Pairwise alignment info from yeast Fas2 (DEG20011054), cf. Figure S97. . . . . S88
S19 Pairwise alignment info from yeast Fba1 (DEG20010617), cf. Figure S107. . . . . S98
S20 Pairwise alignment info from yeast Fba1 (DEG20010617), cf. Figure S111. . . . . S101
S21 Pairwise alignment info from yeast Fcy21 (DEG20010294), cf. Figure S121. . . . . S111
S22 Pairwise alignment info from yeast Fcy21 (DEG20010294), cf. Figure S125. . . . . S114
S23 Pairwise alignment info from yeast Fol1 (DEG20010889), cf. Figure S134. . . . . S122
S24 Pairwise alignment info from yeast Fol1 (DEG20010889), cf. Figure S138. . . . . S125
S25 Pairwise alignment info from yeast Ilv3 (DEG20010579), cf. Figure S148. . . . . S135
S26 Pairwise alignment info from yeast Ilv3 (DEG20010579), cf. Figure S152. . . . . S138
S27 Pairwise alignment info from yeast Ilv5 (DEG20010747), cf. Figure S162. . . . . S148
S28 Pairwise alignment info from yeast Ilv5 (DEG20010747), cf. Figure S166. . . . . S151
S29 Pairwise alignment info from yeast Rib3 (DEG20010261), cf. Figure S176. . . . . S161
S30 Pairwise alignment info from yeast Rib3 (DEG20010261), cf. Figure S180. . . . . S164
S31 Pairwise alignment info from yeast Rib5 (DEG20010082), cf. Figure S190. . . . . S174
S32 Pairwise alignment info from yeast Rib5 (DEG20010082), cf. Figure S194. . . . . S177
S33 Pairwise alignment info from yeast Ssy1 (DEG20010185), cf. Figure S204. . . . . S187
S34 Pairwise alignment info from yeast Ssy1 (DEG20010185), cf. Figure S208. . . . . S190
S35 Pairwise alignment info from yeast Ste12 (DEG20010472), cf. Figure S218. . . . . S200
S36 Pairwise alignment info from yeast Ste12 (DEG20010472), cf. Figure S222. . . . . S203
S37 Pairwise alignment info from yeast Trl1 (DEG20010555), cf. Figure S231. . . . . S211
S38 Pairwise alignment info from yeast Trl1 (DEG20010555), cf. Figure S235. . . . . S214
S39 Pairwise alignment info from yeast Yef3 (DEG20010729), cf. Figure S243. . . . . S222
S40 Pairwise alignment info from yeast Yef3 (DEG20010729), cf. Figure S247. . . . . S225
S41 Pairwise alignment info from yeast Erg11 . . . . . S235
S42 Pairwise alignment info from yeast Erg24 . . . . . S236
S43 Pairwise alignment info from yeast Erg2 . . . . . S237
S44 Pairwise alignment info from yeast Fks1 . . . . . S239
S45 Pairwise alignment info from yeast Fks3 . . . . . S240
S46 Pairwise alignment info from yeast Gsc2 . . . . . S241
S47 Pairwise alignment info from *S. cerevisiae* Ccc1 with all pathogens. . . . . S241
S48 Pairwise alignment info from *S. cerevisiae* Ccc1 with all pathogens. . . . . S245

#### List of Figures

S1 Expectation values for *A.fumigatus*, cf. Table S5. . . . . S11
S2 Expectation values for *C.albicans*, cf. Table S5. . . . . S12
S3 Expectation values for *C.auris*, cf. Table S5. . . . . S12
S4 Expectation values for *C.neoformans.B.3501A*, cf. Table S5. . . . . S13
S5 Expectation values for *C.neoformans.JEC21*, cf. Table S5. . . . . S13
S6 Expectation values for *C.neoformans.grubii.H99*, cf. Table S5. . . . . S14
S7 Expectation values for *C.parapsilosis*, cf. Table S5. . . . . S14
S8 Expectation values for *C.tropicalis*, cf. Table S5. . . . . S15
S9 Expectation values for *G.max*, cf. Table S5. . . . . S15
S10 Expectation values for *H.capsulatum*, cf. Table S5. . . . . S16

S11 Expectation values for *H. sapiens*, cf. Table S5. . . . S16
S12 Expectation values for *N.glabratus*, cf. Table S5. . . . S17
S13 Expectation values for *O.sativa*, cf. Table S5. . . . S17
S14 Expectation values for *S.tuberosum*, cf. Table S5. . . . S18
S15 Expectation values for *Z.mays*, cf. Table S5. . . . S18
S16 Expectation values for *B.cinerea*, cf. Table S5. . . . S19
S17 Expectation values for *B.graminis*, cf. Table S5. . . . S19
S18 Expectation values for *C.truncatum*, cf. Table S5. . . . S20
S19 Expectation values for *F.graminearum*, cf. Table S5. . . . S20
S20 Expectation values for *M.graminicola*, cf. Table S5. . . . S21
S21 Expectation values for *P.graminis*, cf. Table S5. . . . S21
S22 Expectation values for *P.oryzae*, cf. Table S5. . . . S22
S23 Expectation values for *P.striiformis*, cf. Table S5. . . . S22
S24 Expectation values for *P.triticina*, cf. Table S5. . . . S23
S25 Expectation values for *U.maydis*, cf. Table S5. . . . S23
S26 Multiple sequence alignment of yeast Alr1 (WHO Critical Pathogens). Cf. Figure S27 for alignment quality, and Figure S28 for Sneath similarity. Cf. Table S7 for protein names, and pairwise alignment metrics with yeast Alr1. S25
S27 Multiple sequence alignment quality of Alr1 (WHO Critical Pathogens). Cf. Figures S26 and S29. . . . S25
S28 Sneath Similarity of Alr1 for WHO Critical Pathogens, cf. Figure S26 . . . . S26
S29 Phylogeny of hits for Alr1 made using clustal omega and visualized using ggtree, cf. Figures S26 and S27. . . . S27
S30 Multiple sequence alignment of yeast Alr1 (Top 10 Agricultural Fungal Pathogens). Cf. Figure S31 for alignment quality, and Figure S32 for Sneath similarity. Cf. Table S8 for protein names, and pairwise alignment metrics with yeast Alr1. . . . S27
S31 Multiple sequence alignment quality of Alr1 (Top 10 Agricultural Fungal Pathogens). Cf. Figures S30 and S33. S28
S32 Sneath Similarity of Alr1 for Top 10 Agricultural Fungal Pathogens, cf. Figure S30 . . . . S30
S33 Phylogeny of hits for Alr1 made using clustal omega and visualized using ggtree, cf. Figures S30 and S31. . . . S30
S34 Non-redundant (NR) protein hits for DEG20010935/Alr1, with expectation value of no more than 0.1. Green points are medians, and red points are arithmetic means. . . . S31
S35 Non-redundant (NR) protein hits for Alr1 in the kingdom Metazoa. . . . S32
S36 Non-redundant (NR) protein hits for Alr1 in the kingdom Viridiplantae. . . . S33
S37 Non-redundant (NR) protein hits for Alr1 in the kingdom Fungi. . . . S34
S38 Non-redundant (NR) protein hits for DEG20010935/Alr1 at 20 amino acid length queries. . . . S35
S39 Multiple sequence alignment of yeast Aur1 (WHO Critical Pathogens). Cf. Figure S40 for alignment quality, and Figure S41 for Sneath similarity. Cf. Table S9 for protein names, and pairwise alignment metrics with yeast Aur1. S36
S40 Multiple sequence alignment quality of Aur1 (WHO Critical Pathogens). Cf. Figures S39 and S42. . . . S36
S41 Sneath Similarity of Aur1 for WHO Critical Pathogens, cf. Figure S39 . . . . S37
S42 Phylogeny of hits for Aur1 made using clustal omega and visualized using ggtree, cf. Figures S39 and S40. . . . S38
S43 Multiple sequence alignment of yeast Aur1 (Top 10 Agricultural Fungal Pathogens). Cf. Figure S44 for alignment quality, and Figure S45 for Sneath similarity. Cf. Table S10 for protein names, and pairwise alignment metrics with yeast Aur1. . . . S38
S44 Multiple sequence alignment quality of Aur1 (Top 10 Agricultural Fungal Pathogens). Cf. Figures S43 and S46. S39
S45 Sneath Similarity of Aur1 for Top 10 Agricultural Fungal Pathogens, cf. Figure S43 . . . . S41
S46 Phylogeny of hits for Aur1 made using clustal omega and visualized using ggtree, cf. Figures S43 and S44. . . . S41
S47 Non-redundant (NR) protein hits for DEG20010598/Aur1, with expectation value of no more than 0.1. Green points are medians, and red points are arithmetic means. . . . S42
S48 Non-redundant (NR) protein hits for Aur1 in the kingdom Viridiplantae. . . . S43
S49 Non-redundant (NR) protein hits for Aur1 in the kingdom Fungi. . . . S44
S50 Non-redundant (NR) protein hits for Aur1 in the kingdom Metazoa. . . . S45
S51 Non-redundant (NR) protein hits for DEG20010598/Aur1 at 20 amino acid length queries. . . . S46
S52 Multiple sequence alignment of yeast Chs2 (WHO Critical Pathogens). Cf. Figure S53 for alignment quality, and Figure S54 for Sneath similarity. Cf. Table S11 for protein names, and pairwise alignment metrics with yeast Chs2. S47
S53 Multiple sequence alignment quality of Chs2 (WHO Critical Pathogens). Cf. Figures S52 and S55. . . . S47
S54 Sneath Similarity of Chs2 for WHO Critical Pathogens, cf. Figure S52 . . . . S48
S55 Phylogeny of hits for Chs2 made using clustal omega and visualized using ggtree, cf. Figures S52 and S53. . . . S49
S56 Multiple sequence alignment of yeast Chs2 (Top 10 Agricultural Fungal Pathogens). Cf. Figure S57 for alignment quality, and Figure S58 for Sneath similarity. Cf. Table S12 for protein names, and pairwise alignment metrics with yeast Chs2. . . . S49
S57 Multiple sequence alignment quality of Chs2 (Top 10 Agricultural Fungal Pathogens). Cf. Figures S56 and S59. S50

S58 Sneath Similarity of Chs2 for Top 10 Agricultural Fungal Pathogens, cf. Figure S56 . . . S52
S59 Phylogeny of hits for Chs2 made using clustal omega and visualized using ggtree, cf. Figures S56 and S57. . . . S52
S60 Non-redundant (NR) protein hits for DEG20010039/Chs2, with expectation value of no more than 0.1. Green points are medians, and red points are arithmetic means. . . . S53
S61 Non-redundant (NR) protein hits for Chs2 in the kingdom Viridiplantae. . . . S54
S62 Non-redundant (NR) protein hits for Chs2 in the kingdom Fungi. . . . S55
S63 Non-redundant (NR) protein hits for Chs2 in the kingdom SAR. . . . S56
S64 Non-redundant (NR) protein hits for Chs2 in the kingdom Metazoa. . . . S57
S65 Non-redundant (NR) protein hits for DEG20010039/Chs2 at 20 amino acid length queries. . . . S58
S66 Multiple sequence alignment of yeast Erg8 (WHO Critical Pathogens). Cf. Figure S67 for alignment quality, and Figure S68 for Sneath similarity. Cf. Table S13 for protein names, and pairwise alignment metrics with yeast Erg8. S59
S67 Multiple sequence alignment quality of Erg8 (WHO Critical Pathogens). Cf. Figures S66 and S69. . . . S60
S68 Sneath Similarity of Erg8 for WHO Critical Pathogens, cf. Figure S66 . . . S61
S69 Phylogeny of hits for Erg8 made using clustal omega and visualized using ggtree, cf. Figures S66 and S67. . . . S62
S70 Multiple sequence alignment of yeast Erg8 (Top 10 Agricultural Fungal Pathogens). Cf. Figure S71 for alignment quality, and Figure S72 for Sneath similarity. Cf. Table S14 for protein names, and pairwise alignment metrics with yeast Erg8. . . . S62
S71 Multiple sequence alignment quality of Erg8 (Top 10 Agricultural Fungal Pathogens). Cf. Figures S70 and S73. S63
S72 Sneath Similarity of Erg8 for Top 10 Agricultural Fungal Pathogens, cf. Figure S70 . . . S65
S73 Phylogeny of hits for Erg8 made using clustal omega and visualized using ggtree, cf. Figures S70 and S71. . . . S65
S74 Non-redundant (NR) protein hits for DEG20010822/Erg8, with expectation value of no more than 0.1. Green points are medians, and red points are arithmetic means. . . . S66
S75 Non-redundant (NR) protein hits for Erg8 in the kingdom Metazoa. . . . S67
S76 Non-redundant (NR) protein hits for Erg8 in the kingdom Viridiplantae. . . . S68
S77 Non-redundant (NR) protein hits for Erg8 in the kingdom Fungi. . . . S69
S78 Non-redundant (NR) protein hits for Erg8 in the kingdom SAR. . . . S70
S79 Non-redundant (NR) protein hits for DEG20010822/Erg8 at 20 amino acid length queries. . . . S71
S80 Multiple sequence alignment of yeast Fas1 (WHO Critical Pathogens). Cf. Figure S81 for alignment quality, and Figure S82 for Sneath similarity. Cf. Table S15 for protein names, and pairwise alignment metrics with yeast Fas1. S72
S81 Multiple sequence alignment quality of Fas1 (WHO Critical Pathogens). Cf. Figures S80 and S83. . . . S73
S82 Sneath Similarity of Fas1 for WHO Critical Pathogens, cf. Figure S80 . . . S74
S83 Phylogeny of hits for Fas1 made using clustal omega and visualized using ggtree, cf. Figures S80 and S81. . . . S75
S84 Multiple sequence alignment of yeast Fas1 (Top 10 Agricultural Fungal Pathogens). Cf. Figure S85 for alignment quality, and Figure S86 for Sneath similarity. Cf. Table S16 for protein names, and pairwise alignment metrics with yeast Fas1. . . . S75
S85 Multiple sequence alignment quality of Fas1 (Top 10 Agricultural Fungal Pathogens). Cf. Figures S84 and S87. S76
S86 Sneath Similarity of Fas1 for Top 10 Agricultural Fungal Pathogens, cf. Figure S84 . . . S78
S87 Phylogeny of hits for Fas1 made using clustal omega and visualized using ggtree, cf. Figures S84 and S85. . . . S78
S88 Non-redundant (NR) protein hits for DEG20010641/Fas1, with expectation value of no more than 0.1. Green points are medians, and red points are arithmetic means. . . . S79
S89 Non-redundant (NR) protein hits for Fas1 in the kingdom Fungi. . . . S80
S90 Non-redundant (NR) protein hits for Fas1 in the kingdom Viridiplantae. . . . S81
S91 Non-redundant (NR) protein hits for Fas1 in the kingdom SAR. . . . S82
S92 Non-redundant (NR) protein hits for DEG20010641/Fas1 at 20 amino acid length queries. . . . S83
S93 Multiple sequence alignment of yeast Fas2 (WHO Critical Pathogens). Cf. Figure S94 for alignment quality, and Figure S95 for Sneath similarity. Cf. Table S17 for protein names, and pairwise alignment metrics with yeast Fas2. S84
S94 Multiple sequence alignment quality of Fas2 (WHO Critical Pathogens). Cf. Figures S93 and S96. . . . S84
S95 Sneath Similarity of Fas2 for WHO Critical Pathogens, cf. Figure S93 . . . S85
S96 Phylogeny of hits for Fas2 made using clustal omega and visualized using ggtree, cf. Figures S93 and S94. . . . S86
S97 Multiple sequence alignment of yeast Fas2 (Top 10 Agricultural Fungal Pathogens). Cf. Figure S98 for alignment quality, and Figure S99 for Sneath similarity. Cf. Table S18 for protein names, and pairwise alignment metrics with yeast Fas2. . . . S86
S98 Multiple sequence alignment quality of Fas2 (Top 10 Agricultural Fungal Pathogens). Cf. Figures S97 and S100. S87
S99 Sneath Similarity of Fas2 for Top 10 Agricultural Fungal Pathogens, cf. Figure S97 . . . S89
S100 Phylogeny of hits for Fas2 made using clustal omega and visualized using ggtree, cf. Figures S97 and S98. . . . S89
S101 Non-redundant (NR) protein hits for DEG20011054/Fas2, with expectation value of no more than 0.1. Green points are medians, and red points are arithmetic means. . . . S90
S102 Non-redundant (NR) protein hits for Fas2 in the kingdom Metazoa. . . . S91

S103 Non-redundant (NR) protein hits for Fas2 in the kingdom SAR. . . . S92
S104 Non-redundant (NR) protein hits for Fas2 in the kingdom Viridiplantae. . . . S93
S105 Non-redundant (NR) protein hits for Fas2 in the kingdom Fungi. . . . S94
S106 Non-redundant (NR) protein hits for DEG20011054/Fas2 at 20 amino acid length queries. . . . S95
S107 Multiple sequence alignment of yeast Fba1 (WHO Critical Pathogens). Cf. Figure S108 for alignment quality, and Figure S109 for Sneath similarity. Cf. Table S19 for protein names, and pairwise alignment metrics with yeast Fba1. . . . S96
S108 Multiple sequence alignment quality of Fba1 (WHO Critical Pathogens). Cf. Figures S107 and S110. . . . S97
S109 Sneath Similarity of Fba1 for WHO Critical Pathogens, cf. Figure S107 . . . S98
S110 Phylogeny of hits for Fba1 made using clustal omega and visualized using ggtree, cf. Figures S107 and S108. . . S99
S111 Multiple sequence alignment of yeast Fba1 (Top 10 Agricultural Fungal Pathogens). Cf. Figure S112 for alignment quality, and Figure S113 for Sneath similarity. Cf. Table S20 for protein names, and pairwise alignment metrics with yeast Fba1. . . . S99
S112 Multiple sequence alignment quality of Fba1 (Top 10 Agricultural Fungal Pathogens). Cf. Figures S111 and S114. S100
S113 Sneath Similarity of Fba1 for Top 10 Agricultural Fungal Pathogens, cf. Figure S111 . . . S102
S114 Phylogeny of hits for Fba1 made using clustal omega and visualized using ggtree, cf. Figures S111 and S112. . S102
S115 Non-redundant (NR) protein hits for DEG20010617/Fba1, with expectation value of no more than 0.1. Green points are medians, and red points are arithmetic means. . . . S103
S116 Non-redundant (NR) protein hits for Fba1 in the kingdom Viridiplantae. . . . S104
S117 Non-redundant (NR) protein hits for Fba1 in the kingdom SAR. . . . S105
S118 Non-redundant (NR) protein hits for Fba1 in the kingdom Metazoa. . . . S106
S119 Non-redundant (NR) protein hits for Fba1 in the kingdom Fungi. . . . S107
S120 Non-redundant (NR) protein hits for DEG20010617/Fba1 at 20 amino acid length queries. . . . S108
S121 Multiple sequence alignment of yeast Fcy21 (WHO Critical Pathogens). Cf. Figure S122 for alignment quality, and Figure S123 for Sneath similarity. Cf. Table S21 for protein names, and pairwise alignment metrics with yeast Fcy21. . . . S109
S122 Multiple sequence alignment quality of Fcy21 (WHO Critical Pathogens). Cf. Figures S121 and S124. . . . S110
S123 Sneath Similarity of Fcy21 for WHO Critical Pathogens, cf. Figure S121 . . . S111
S124 Phylogeny of hits for Fcy21 made using clustal omega and visualized using ggtree, cf. Figures S121 and S122. . S112
S125 Multiple sequence alignment of yeast Fcy21 (Top 10 Agricultural Fungal Pathogens). Cf. Figure S126 for alignment quality, and Figure S127 for Sneath similarity. Cf. Table S22 for protein names, and pairwise alignment metrics with yeast Fcy21. . . . S112
S126 Multiple sequence alignment quality of Fcy21 (Top 10 Agricultural Fungal Pathogens). Cf. Figures S125 and S128. . . . S113
S127 Sneath Similarity of Fcy21 for Top 10 Agricultural Fungal Pathogens, cf. Figure S125 . . . S115
S128 Phylogeny of hits for Fcy21 made using clustal omega and visualized using ggtree, cf. Figures S125 and S126. . S115
S129 Non-redundant (NR) protein hits for DEG20010294/Fcy21, with expectation value of no more than 0.1. Green points are medians, and red points are arithmetic means. . . . S116
S130 Non-redundant (NR) protein hits for Fcy21 in the kingdom Fungi. . . . S117
S131 Non-redundant (NR) protein hits for Fcy21 in the kingdom SAR. . . . S118
S132 Non-redundant (NR) protein hits for Fcy21 in the kingdom Viridiplantae. . . . S119
S133 Non-redundant (NR) protein hits for DEG20010294/Fcy21 at 20 amino acid length queries. . . . S120
S134 Multiple sequence alignment of yeast Fol1 (WHO Critical Pathogens). Cf. Figure S135 for alignment quality, and Figure S136 for Sneath similarity. Cf. Table S23 for protein names, and pairwise alignment metrics with yeast Fol1. . . . S121
S135 Multiple sequence alignment quality of Fol1 (WHO Critical Pathogens). Cf. Figures S134 and S137. . . . S121
S136 Sneath Similarity of Fol1 for WHO Critical Pathogens, cf. Figure S134 . . . S122
S137 Phylogeny of hits for Fol1 made using clustal omega and visualized using ggtree, cf. Figures S134 and S135. . . S123
S138 Multiple sequence alignment of yeast Fol1 (Top 10 Agricultural Fungal Pathogens). Cf. Figure S139 for alignment quality, and Figure S140 for Sneath similarity. Cf. Table S24 for protein names, and pairwise alignment metrics with yeast Fol1. . . . S123
S139 Multiple sequence alignment quality of Fol1 (Top 10 Agricultural Fungal Pathogens). Cf. Figures S138 and S141. S124
S140 Sneath Similarity of Fol1 for Top 10 Agricultural Fungal Pathogens, cf. Figure S138 . . . S126
S141 Phylogeny of hits for Fol1 made using clustal omega and visualized using ggtree, cf. Figures S138 and S139. . . S126
S142 Non-redundant (NR) protein hits for DEG20010889/Fol1, with expectation value of no more than 0.1. Green points are medians, and red points are arithmetic means. . . . S127
S143 Non-redundant (NR) protein hits for Fol1 in the kingdom SAR. . . . S128
S144 Non-redundant (NR) protein hits for Fol1 in the kingdom Viridiplantae. . . . S129

S145 Non-redundant (NR) protein hits for Fol1 in the kingdom Fungi. . . . . . S130
S146 Non-redundant (NR) protein hits for Fol1 in the kingdom Metazoa. . . . . . S131
S147 Non-redundant (NR) protein hits for DEG20010889/Fol1 at 20 amino acid length queries. . . . . . S132
S148 Multiple sequence alignment of yeast Ilv3 (WHO Critical Pathogens). Cf. Figure S149 for alignment quality, and Figure S150 for Sneath similarity. Cf. Table S25 for protein names, and pairwise alignment metrics with yeast Ilv3. S133
S149 Multiple sequence alignment quality of Ilv3 (WHO Critical Pathogens). Cf. Figures S148 and S151. . . . . . S134
S150 Sneath Similarity of Ilv3 for WHO Critical Pathogens, cf. Figure S148 . . . . . . S135
S151 Phylogeny of hits for Ilv3 made using clustal omega and visualized using ggtree, cf. Figures S148 and S149. . . S136
S152 Multiple sequence alignment of yeast Ilv3 (Top 10 Agricultural Fungal Pathogens). Cf. Figure S153 for alignment quality, and Figure S154 for Sneath similarity. Cf. Table S26 for protein names, and pairwise alignment metrics with yeast Ilv3. . . . . . S136
S153 Multiple sequence alignment quality of Ilv3 (Top 10 Agricultural Fungal Pathogens). Cf. Figures S152 and S155. S137
S154 Sneath Similarity of Ilv3 for Top 10 Agricultural Fungal Pathogens, cf. Figure S152 . . . . . . S139
S155 Phylogeny of hits for Ilv3 made using clustal omega and visualized using ggtree, cf. Figures S152 and S153. . . S139
S156 Non-redundant (NR) protein hits for DEG20010579/Ilv3, with expectation value of no more than 0.1. Green points are medians, and red points are arithmetic means. . . . . . S140
S157 Non-redundant (NR) protein hits for Ilv3 in the kingdom Metazoa. . . . . . S141
S158 Non-redundant (NR) protein hits for Ilv3 in the kingdom SAR. . . . . . S142
S159 Non-redundant (NR) protein hits for Ilv3 in the kingdom Fungi. . . . . . S143
S160 Non-redundant (NR) protein hits for Ilv3 in the kingdom Viridiplantae. . . . . . S144
S161 Non-redundant (NR) protein hits for DEG20010579/Ilv3 at 20 amino acid length queries. . . . . . S145
S162 Multiple sequence alignment of yeast Ilv5 (WHO Critical Pathogens). Cf. Figure S163 for alignment quality, and Figure S164 for Sneath similarity. Cf. Table S27 for protein names, and pairwise alignment metrics with yeast Ilv5. S146
S163 Multiple sequence alignment quality of Ilv5 (WHO Critical Pathogens). Cf. Figures S162 and S165. . . . . . S147
S164 Sneath Similarity of Ilv5 for WHO Critical Pathogens, cf. Figure S162 . . . . . . S148
S165 Phylogeny of hits for Ilv5 made using clustal omega and visualized using ggtree, cf. Figures S162 and S163. . . S149
S166 Multiple sequence alignment of yeast Ilv5 (Top 10 Agricultural Fungal Pathogens). Cf. Figure S167 for alignment quality, and Figure S168 for Sneath similarity. Cf. Table S28 for protein names, and pairwise alignment metrics with yeast Ilv5. . . . . . S149
S167 Multiple sequence alignment quality of Ilv5 (Top 10 Agricultural Fungal Pathogens). Cf. Figures S166 and S169. S150
S168 Sneath Similarity of Ilv5 for Top 10 Agricultural Fungal Pathogens, cf. Figure S166 . . . . . . S152
S169 Phylogeny of hits for Ilv5 made using clustal omega and visualized using ggtree, cf. Figures S166 and S167. . . S152
S170 Non-redundant (NR) protein hits for DEG20010747/Ilv5, with expectation value of no more than 0.1. Green points are medians, and red points are arithmetic means. . . . . . S153
S171 Non-redundant (NR) protein hits for Ilv5 in the kingdom Metazoa. . . . . . S154
S172 Non-redundant (NR) protein hits for Ilv5 in the kingdom SAR. . . . . . S155
S173 Non-redundant (NR) protein hits for Ilv5 in the kingdom Viridiplantae. . . . . . S156
S174 Non-redundant (NR) protein hits for Ilv5 in the kingdom Fungi. . . . . . S157
S175 Non-redundant (NR) protein hits for DEG20010747/Ilv5 at 20 amino acid length queries. . . . . . S158
S176 Multiple sequence alignment of yeast Rib3 (WHO Critical Pathogens). Cf. Figure S177 for alignment quality, and Figure S178 for Sneath similarity. Cf. Table S29 for protein names, and pairwise alignment metrics with yeast Rib3. . . . . . S159
S177 Multiple sequence alignment quality of Rib3 (WHO Critical Pathogens). Cf. Figures S176 and S179. . . . . . S160
S178 Sneath Similarity of Rib3 for WHO Critical Pathogens, cf. Figure S176 . . . . . . S161
S179 Phylogeny of hits for Rib3 made using clustal omega and visualized using ggtree, cf. Figures S176 and S177. . . S162
S180 Multiple sequence alignment of yeast Rib3 (Top 10 Agricultural Fungal Pathogens). Cf. Figure S181 for alignment quality, and Figure S182 for Sneath similarity. Cf. Table S30 for protein names, and pairwise alignment metrics with yeast Rib3. . . . . . S162
S181 Multiple sequence alignment quality of Rib3 (Top 10 Agricultural Fungal Pathogens). Cf. Figures S180 and S183. S163
S182 Sneath Similarity of Rib3 for Top 10 Agricultural Fungal Pathogens, cf. Figure S180 . . . . . . S165
S183 Phylogeny of hits for Rib3 made using clustal omega and visualized using ggtree, cf. Figures S180 and S181. . . S165
S184 Non-redundant (NR) protein hits for DEG20010261/Rib3, with expectation value of no more than 0.1. Green points are medians, and red points are arithmetic means. . . . . . S166
S185 Non-redundant (NR) protein hits for Rib3 in the kingdom Viridiplantae. . . . . . S167
S186 Non-redundant (NR) protein hits for Rib3 in the kingdom SAR. . . . . . S168
S187 Non-redundant (NR) protein hits for Rib3 in the kingdom Fungi. . . . . . S169
S188 Non-redundant (NR) protein hits for Rib3 in the kingdom Metazoa. . . . . . S170
S189 Non-redundant (NR) protein hits for DEG20010261/Rib3 at 20 amino acid length queries. . . . . . S171

S190 Multiple sequence alignment of yeast Rib5 (WHO Critical Pathogens). Cf. Figure S191 for alignment quality, and Figure S192 for Sneath similarity. Cf. Table S31 for protein names, and pairwise alignment metrics with yeast Rib5. . . . . . S172
S191 Multiple sequence alignment quality of Rib5 (WHO Critical Pathogens). Cf. Figures S190 and S193. . . . . . S173
S192 Sneath Similarity of Rib5 for WHO Critical Pathogens, cf. Figure S190 . . . . . S174
S193 Phylogeny of hits for Rib5 made using clustal omega and visualized using ggtree, cf. Figures S190 and S191. . . S175
S194 Multiple sequence alignment of yeast Rib5 (Top 10 Agricultural Fungal Pathogens). Cf. Figure S195 for alignment quality, and Figure S196 for Sneath similarity. Cf. Table S32 for protein names, and pairwise alignment metrics with yeast Rib5. . . . . . S175
S195 Multiple sequence alignment quality of Rib5 (Top 10 Agricultural Fungal Pathogens). Cf. Figures S194 and S197. S176
S196 Sneath Similarity of Rib5 for Top 10 Agricultural Fungal Pathogens, cf. Figure S194 . . . . . S178
S197 Phylogeny of hits for Rib5 made using clustal omega and visualized using ggtree, cf. Figures S194 and S195. . . S178
S198 Non-redundant (NR) protein hits for DEG20010082/Rib5, with expectation value of no more than 0.1. Green points are medians, and red points are arithmetic means. . . . . . S179
S199 Non-redundant (NR) protein hits for Rib5 in the kingdom Fungi. . . . . . S180
S200 Non-redundant (NR) protein hits for Rib5 in the kingdom SAR. . . . . . S181
S201 Non-redundant (NR) protein hits for Rib5 in the kingdom Metazoa. . . . . . S182
S202 Non-redundant (NR) protein hits for Rib5 in the kingdom Viridiplantae. . . . . . S183
S203 Non-redundant (NR) protein hits for DEG20010082/Rib5 at 20 amino acid length queries. . . . . . S184
S204 Multiple sequence alignment of yeast Ssy1 (WHO Critical Pathogens). Cf. Figure S205 for alignment quality, and Figure S206 for Sneath similarity. Cf. Table S33 for protein names, and pairwise alignment metrics with yeast Ssy1. . . . . . S185
S205 Multiple sequence alignment quality of Ssy1 (WHO Critical Pathogens). Cf. Figures S204 and S207. . . . . . S186
S206 Sneath Similarity of Ssy1 for WHO Critical Pathogens, cf. Figure S204 . . . . . S187
S207 Phylogeny of hits for Ssy1 made using clustal omega and visualized using ggtree, cf. Figures S204 and S205. . . S188
S208 Multiple sequence alignment of yeast Ssy1 (Top 10 Agricultural Fungal Pathogens). Cf. Figure S209 for alignment quality, and Figure S210 for Sneath similarity. Cf. Table S34 for protein names, and pairwise alignment metrics with yeast Ssy1. . . . . . S188
S209 Multiple sequence alignment quality of Ssy1 (Top 10 Agricultural Fungal Pathogens). Cf. Figures S208 and S211. S189
S210 Sneath Similarity of Ssy1 for Top 10 Agricultural Fungal Pathogens, cf. Figure S208 . . . . . S191
S211 Phylogeny of hits for Ssy1 made using clustal omega and visualized using ggtree, cf. Figures S208 and S209. . . S191
S212 Non-redundant (NR) protein hits for DEG20010185/Ssy1, with expectation value of no more than 0.1. Green points are medians, and red points are arithmetic means. . . . . . S192
S213 Non-redundant (NR) protein hits for Ssy1 in the kingdom Fungi. . . . . . S193
S214 Non-redundant (NR) protein hits for Ssy1 in the kingdom Metazoa. . . . . . S194
S215 Non-redundant (NR) protein hits for Ssy1 in the kingdom SAR. . . . . . S195
S216 Non-redundant (NR) protein hits for Ssy1 in the kingdom Viridiplantae. . . . . . S196
S217 Non-redundant (NR) protein hits for DEG20010185/Ssy1 at 20 amino acid length queries. . . . . . S197
S218 Multiple sequence alignment of yeast Ste12 (WHO Critical Pathogens). Cf. Figure S219 for alignment quality, and Figure S220 for Sneath similarity. Cf. Table S35 for protein names, and pairwise alignment metrics with yeast Ste12. . . . . . S198
S219 Multiple sequence alignment quality of Ste12 (WHO Critical Pathogens). Cf. Figures S218 and S221. . . . . S199
S220 Sneath Similarity of Ste12 for WHO Critical Pathogens, cf. Figure S218 . . . . . S200
S221 Phylogeny of hits for Ste12 made using clustal omega and visualized using ggtree, cf. Figures S218 and S219. . S201
S222 Multiple sequence alignment of yeast Ste12 (Top 10 Agricultural Fungal Pathogens). Cf. Figure S223 for alignment quality, and Figure S224 for Sneath similarity. Cf. Table S36 for protein names, and pairwise alignment metrics with yeast Ste12. . . . . . S201
S223 Multiple sequence alignment quality of Ste12 (Top 10 Agricultural Fungal Pathogens). Cf. Figures S222 and S225. . . . . . S202
S224 Sneath Similarity of Ste12 for Top 10 Agricultural Fungal Pathogens, cf. Figure S222 . . . . . S204
S225 Phylogeny of hits for Ste12 made using clustal omega and visualized using ggtree, cf. Figures S222 and S223. . S204
S226 Non-redundant (NR) protein hits for DEG20010472/Ste12, with expectation value of no more than 0.1. Green points are medians, and red points are arithmetic means. . . . . . S205
S227 Non-redundant (NR) protein hits for Ste12 in the kingdom Viridiplantae. . . . . . S206
S228 Non-redundant (NR) protein hits for Ste12 in the kingdom Metazoa. . . . . . S207
S229 Non-redundant (NR) protein hits for Ste12 in the kingdom Fungi. . . . . . S208
S230 Non-redundant (NR) protein hits for DEG20010472/Ste12 at 20 amino acid length queries. . . . . . S209

S231 Multiple sequence alignment of yeast Trl1 (WHO Critical Pathogens). Cf. Figure S232 for alignment quality, and Figure S233 for Sneath similarity. Cf. Table S37 for protein names, and pairwise alignment metrics with yeast Trl1. . . . S210
S232 Multiple sequence alignment quality of Trl1 (WHO Critical Pathogens). Cf. Figures S231 and S234. . . . S210
S233 Sneath Similarity of Trl1 for WHO Critical Pathogens, cf. Figure S231 . . . S211
S234 Phylogeny of hits for Trl1 made using clustal omega and visualized using ggtree, cf. Figures S231 and S232. . . S212
S235 Multiple sequence alignment of yeast Trl1 (Top 10 Agricultural Fungal Pathogens). Cf. Figure S236 for alignment quality, and Figure S237 for Sneath similarity. Cf. Table S38 for protein names, and pairwise alignment metrics with yeast Trl1. . . . S212
S236 Multiple sequence alignment quality of Trl1 (Top 10 Agricultural Fungal Pathogens). Cf. Figures S235 and S238.S213
S237 Sneath Similarity of Trl1 for Top 10 Agricultural Fungal Pathogens, cf. Figure S235 . . . S215
S238 Phylogeny of hits for Trl1 made using clustal omega and visualized using ggtree, cf. Figures S235 and S236. . . S215
S239 Non-redundant (NR) protein hits for DEG20010555/Trl1, with expectation value of no more than 0.1. Green points are medians, and red points are arithmetic means. . . . S216
S240 Non-redundant (NR) protein hits for Trl1 in the kingdom Viridiplantae. . . . S217
S241 Non-redundant (NR) protein hits for Trl1 in the kingdom Fungi. . . . S218
S242 Non-redundant (NR) protein hits for DEG20010555/Trl1 at 20 amino acid length queries. . . . S219
S243 Multiple sequence alignment of yeast Yef3 (WHO Critical Pathogens). Cf. Figure S244 for alignment quality, and Figure S245 for Sneath similarity. Cf. Table S39 for protein names, and pairwise alignment metrics with yeast Yef3. . . . S220
S244 Multiple sequence alignment quality of Yef3 (WHO Critical Pathogens). Cf. Figures S243 and S246. . . . S221
S245 Sneath Similarity of Yef3 for WHO Critical Pathogens, cf. Figure S243 . . . S222
S246 Phylogeny of hits for Yef3 made using clustal omega and visualized using ggtree, cf. Figures S243 and S244. . . S223
S247 Multiple sequence alignment of yeast Yef3 (Top 10 Agricultural Fungal Pathogens). Cf. Figure S248 for alignment quality, and Figure S249 for Sneath similarity. Cf. Table S40 for protein names, and pairwise alignment metrics with yeast Yef3. . . . S223
S248 Multiple sequence alignment quality of Yef3 (Top 10 Agricultural Fungal Pathogens). Cf. Figures S247 and S250.S224
S249 Sneath Similarity of Yef3 for Top 10 Agricultural Fungal Pathogens, cf. Figure S247 . . . S226
S250 Phylogeny of hits for Yef3 made using clustal omega and visualized using ggtree, cf. Figures S247 and S248. . . S226
S251 Non-redundant (NR) protein hits for DEG20010729/Yef3, with expectation value of no more than 0.1. Green points are medians, and red points are arithmetic means. . . . S227
S252 Non-redundant (NR) protein hits for Yef3 in the kingdom Fungi. . . . S228
S253 Non-redundant (NR) protein hits for Yef3 in the kingdom SAR. . . . S229
S254 Non-redundant (NR) protein hits for Yef3 in the kingdom Viridiplantae. . . . S230
S255 Non-redundant (NR) protein hits for Yef3 in the kingdom Metazoa. . . . S231
S256 Non-redundant (NR) protein hits for DEG20010729/Yef3 at 20 amino acid length queries. . . . S232
S257 MSA for Erg11; cf. Table S41 for gene names and exact alignment values. . . . S233
S258 Histogram of Erg11 hits against NR in 20-aa windows . . . S234
S259 MSA for Erg24; cf. Table S42 for gene names and exact alignment values. . . . S235
S260 MSA for Erg2; cf. Table S43 for gene names and exact alignment values. . . . S236
S261 MSA for Fks1; cf. Table S44 for gene names and exact alignment values. . . . S237
S262 Histogram of Fks1 hits against NR in 20-aa windows . . . S238
S263 MSA for Fks3; cf. Table S45 for gene names and exact alignment values. . . . S239
S264 MSA for Gsc2; cf. Table S46 for gene names and exact alignment values. . . . S240
S265 MSA of Ccc1, with all pathogens in the WHO Critical Pathogens group. Cf. Table S47 and Fig. S266. . . . S242
S266 2-D MSA of Ccc1, with all pathogens in the group. Cf. Table S47. . . . S242
S267 Phylogeny of Ccc1, with all pathogens in the group. Cf. Table S47, Figures S265 and S266. Plot was made using ggtree and ggplot2. . . . S243
S268 MSA of Ccc1 with only *Aspergillus*, with all genes in the group. Cf. Table S47 and Fig. S265. . . . S243
S269 MSA of Ccc1 with only *Candida*, with all genes in the group. Cf. Table S47 and Fig. S265. . . . S244
S270 MSA of Ccc1 with only *Cryptococcus*, with all genes in the group. Cf. Table S47 and Fig. S265. . . . S244
S271 MSA of Ccc1, with all pathogens in the Top 10 Agricultural Fungal Pathogens group. Cf. Table S48 and Fig. S272. . . . S246
S272 2-D MSA of Ccc1, with all pathogens in the group. Cf. Table S48. . . . S247

| Drug Class | Mechanism | Drawbacks |
|---|---|---|
| echinocandins | β-(1,3)-glucan synthesis[1] (Fks1, Fks2/Gsc2, Fks3) | poor bioavailability, IV dosing |
| azoles | ergosterol synthesis (Erg11)inhibition of CYP-dependent 14α-demethylase | hepatotoxicity, weeks-months treatment |
| pyrimidine analogs | disrupts RNA & DNA synthesis | ineffective against *Aspergillus*; hepatotoxic |
| allylamines/thiocarbamates | squalene epoxidase/Erg1 (e.g. Terbinafine) | CYP2D6 drug-drug interactions |
| morpholines | Erg2/Erg24 in ergosterol biosynthesis: δ-14 reductase and δ-7–δ-8 isomerase | |
| polyenes | ergosterol in fungal cell membrane | severe renal toxicity |

Table S1: Major current human anti-fungal compound classes and their targets, all genes refer to *Saccharomyces cerevisiae*.

### S1 Data Sources

| Species | Source |
|---|---|
| *Glycine max* | GCF_000004515.6_Glycine_max_v4.0_protein |
| *Homo sapiens* | GCF_000001405 40_GRCh38.p14_protein |
| *Oryza sativa* | GCF_001433935.1_IRGSP-1.0_protein |
| *Solanum tuberosum* | GCF_000226075.1_SolTub_3.0_protein |
| *Zea mays* | GCF_902167145.1_Zm-B73-REFERENCE-NAM-5.0_protein |

Table S2: Species and the data sources used for hosts. The Top 10 agricultural pathogens group used every species here, while the WHO pathogens only used *H. sapiens*.

| Species | Source |
|---|---|
| *Aspergillus fumigatus* | GCF_000002655 1_ASM265v1_protein |
| *Candida albicans* | GCF_000182965 3_ASM18296v3_protein |
| *Candida auris* | GCF_003013715 1_ASM301371v2_protein |
| *Candida parapsilosis* | GCF_000182765 1_ASM18276v2_protein |
| *Candida tropicalis* | GCF_000006335 3_ASM633v3_protein |
| *Cryptococcus.neoformans JEC21* | GCF_000091045 1_ASM9104v1_protein |
| *Cryptococcus.neoformans.B 3501A* | GCF_000149385 1_ASM14938v1_protein |
| *Cryptococcus.neoformans.grubii H99* | GCF_000149245 1_CNA3_protein |
| *Histoplasma capsulatum* | GCF_000150115 1_ASM15011v1_protein |
| *Nakaseomyces glabratus* | GCF_000002545 3_ASM254v2_protein |

Table S3: World Health Organization fungal pathogens and the data sources used.

| Species | Source |
|---|---|
| *Blumeria graminis* | FungiDB-65_BgraminisTritici96224_AnnotatedProteins |
| *Botrytis cinerea* | GCF_000143535 2_ASM14353v4_protein |
| *Colletotrichum truncatum* | GCF_014235925 1_CTRU02_protein |
| *Fusarium graminearum* | GCF_000240135 3_ASM24013v3_protein |
| *Mycosphaerella graminicola* | FungiDB-65_ZtriticiIPO323_AnnotatedProteins |
| *Puccinia graminis* | GCF_000149925 1_ASM14992v1_protein |
| *Puccinia striiformis* | GCF_021901695 1_Pst134E36_v1_pri_protein |
| *Puccinia triticina* | GCF_026914185 1_ASM2691418v1_protein |
| *Pyricularia oryzae* | FungiDB-65_PoryzaeBR32_AnnotatedProteins fasta |
| *Ustilago maydis* | GCF_000328475 2_Umaydis521_2.0_protein |

Table S4: Top ten agricultural pathogens and the data sources used.

### S2 BLAST Expectation Value Histograms

#### S2.1 WHO Pathogens

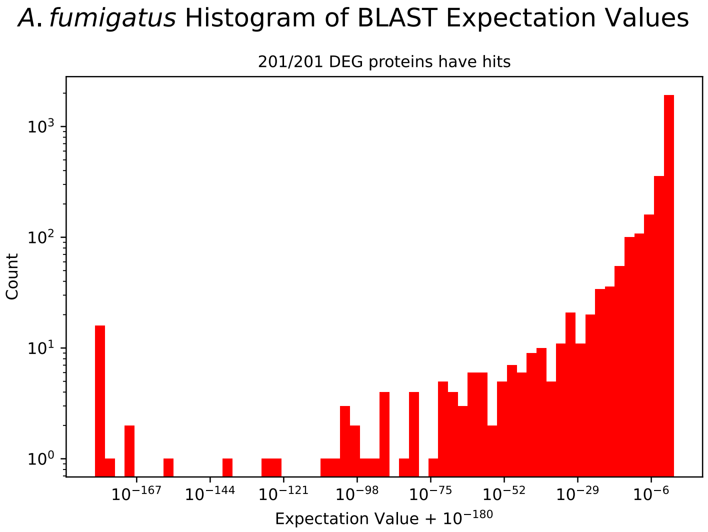


Figure S1: Expectation values for *A.fumigatus*, cf. Table S5.

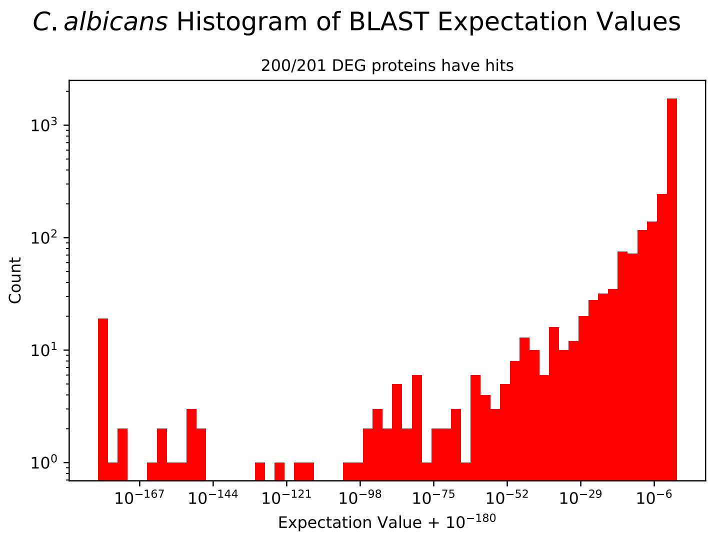


Figure S2: Expectation values for *C.albicans*, cf. Table S5.

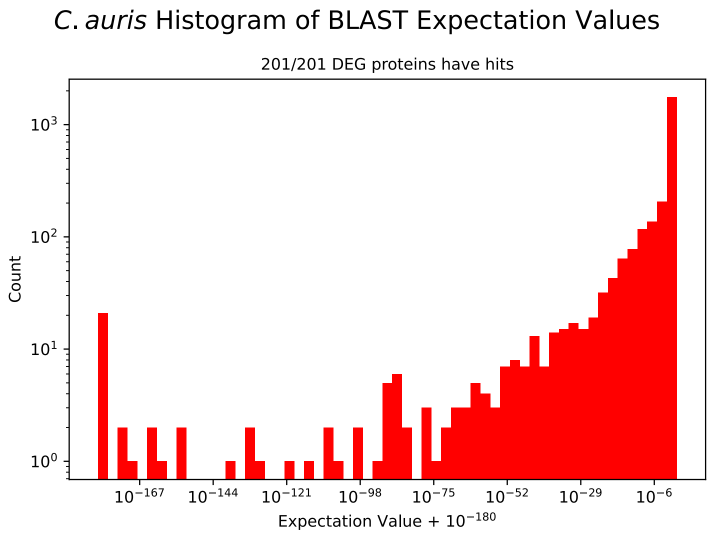


Figure S3: Expectation values for *C.auris*, cf. Table S5.

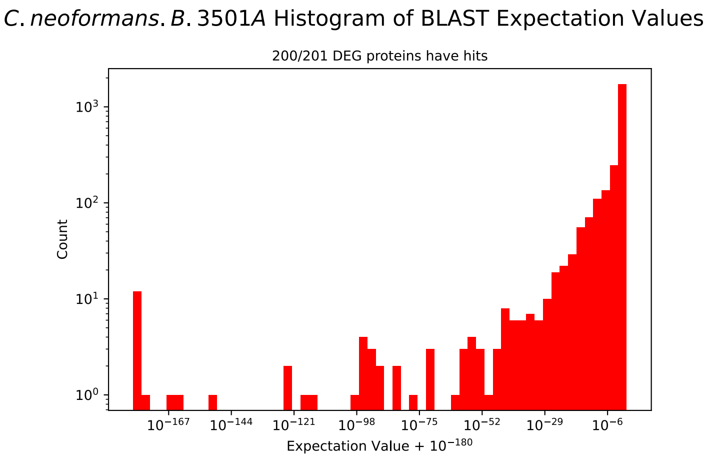


Figure S4: Expectation values for *C.neoformans.B.3501A*, cf. Table S5.

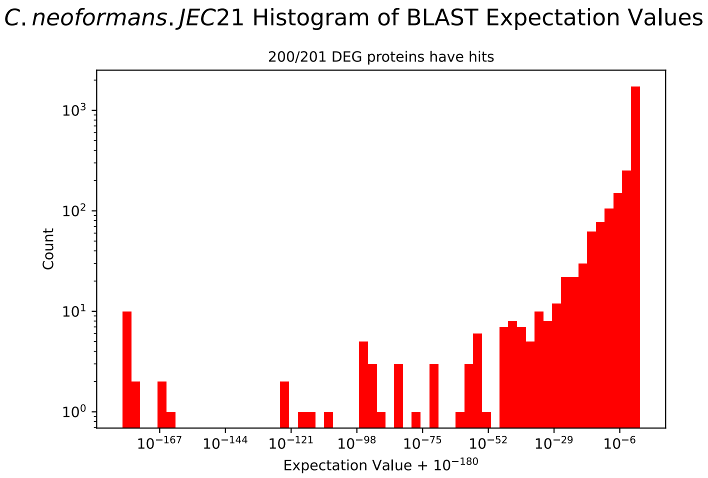


Figure S5: Expectation values for *C.neoformans.JEC21*, cf. Table S5.

#### *C. neoformans. grubii. H*99 Histogram of BLAST Expectation Values

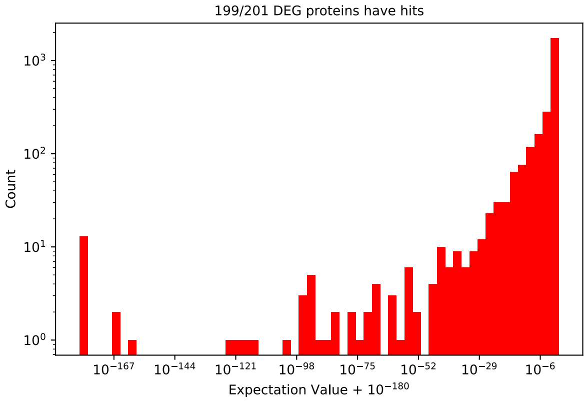


Figure S6: Expectation values for *C.neoformans.grubii.H99*, cf. Table S5.

#### *C. parapsilosis* Histogram of BLAST Expectation Values

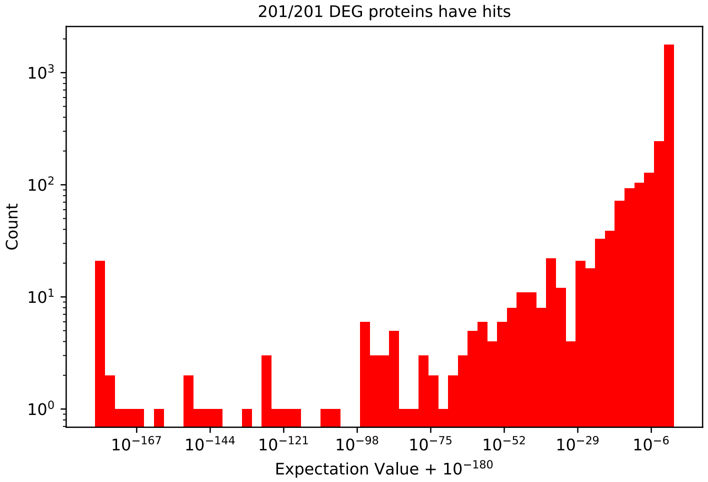


Figure S7: Expectation values for *C.parapsilosis*, cf. Table S5.

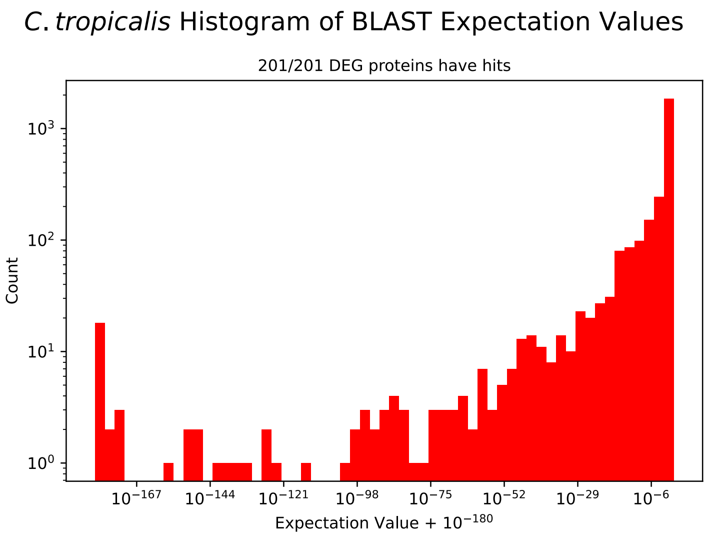


Figure S8: Expectation values for *C.tropicalis*, cf. Table S5.

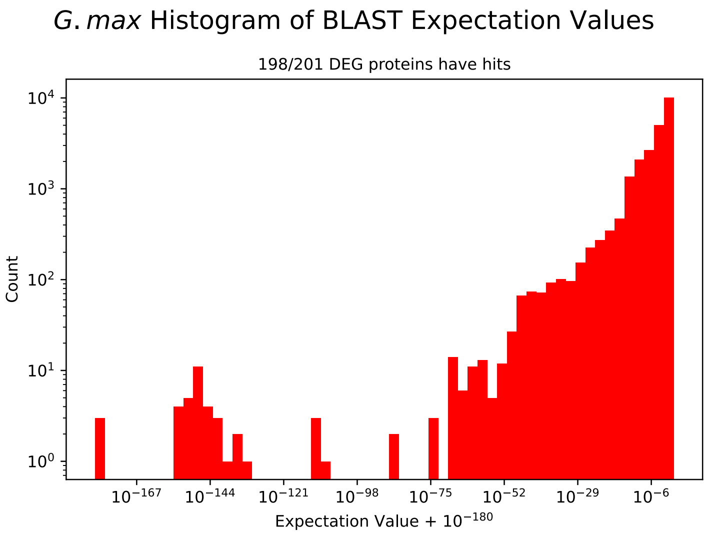


Figure S9: Expectation values for *G.max*, cf. Table S5.

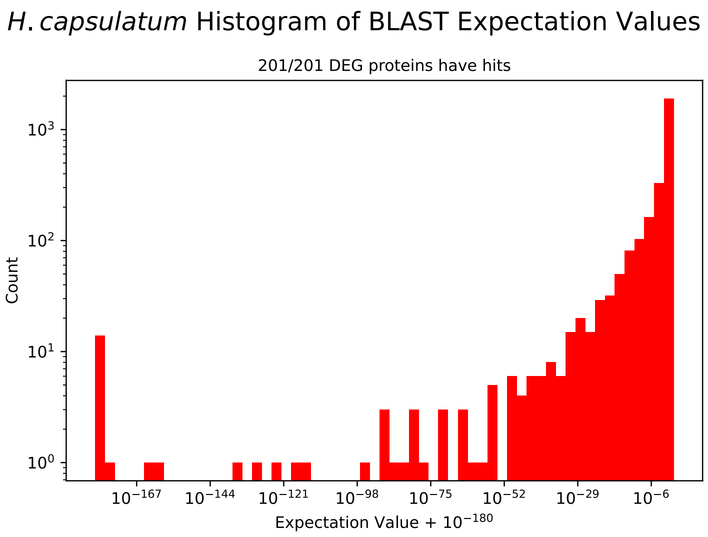


Figure S10: Expectation values for *H.capsulatum*, cf. Table S5.

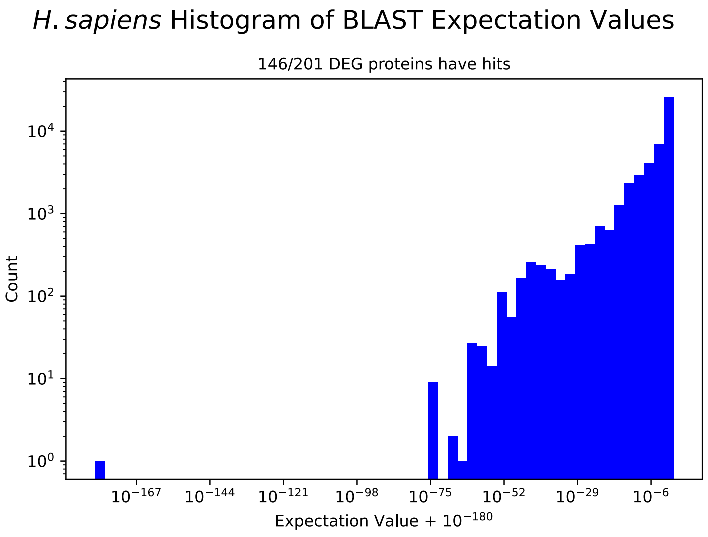


Figure S11: Expectation values for *H. sapiens*, cf. Table S5.

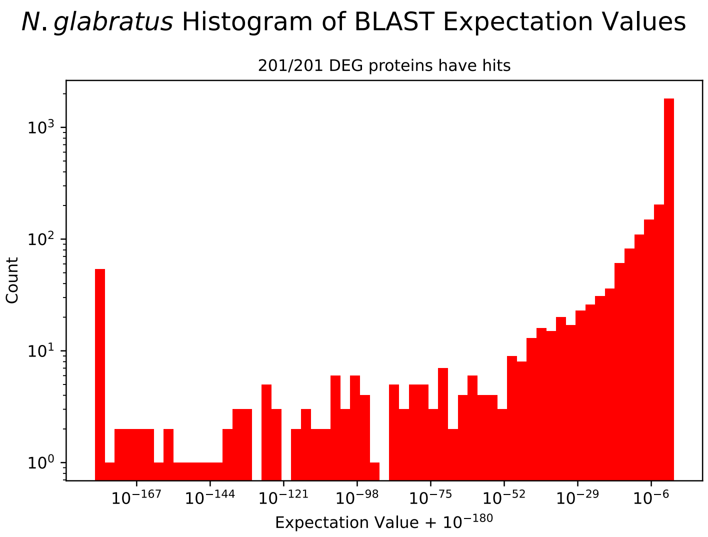


Figure S12: Expectation values for *N.glabratus*, cf. Table S5.

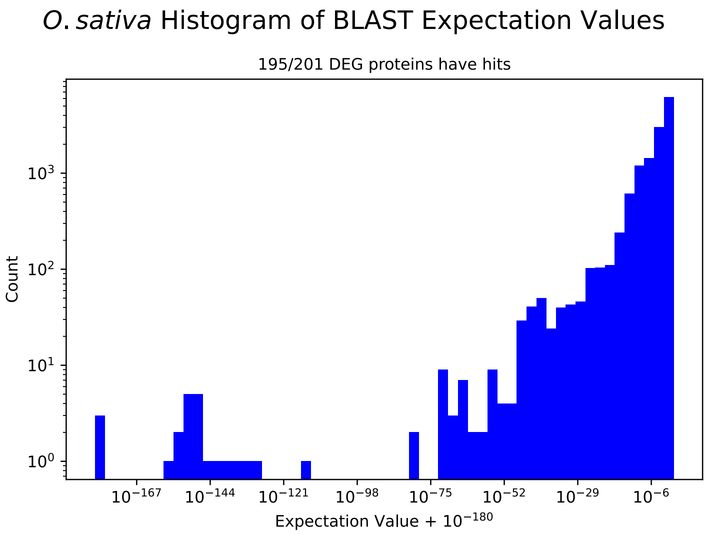


Figure S13: Expectation values for *O.sativa*, cf. Table S5.

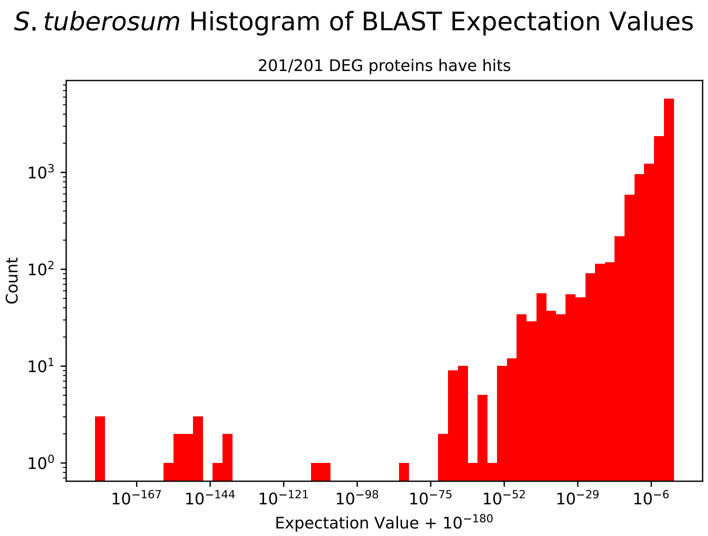


Figure S14: Expectation values for *S.tuberosum*, cf. Table S5.

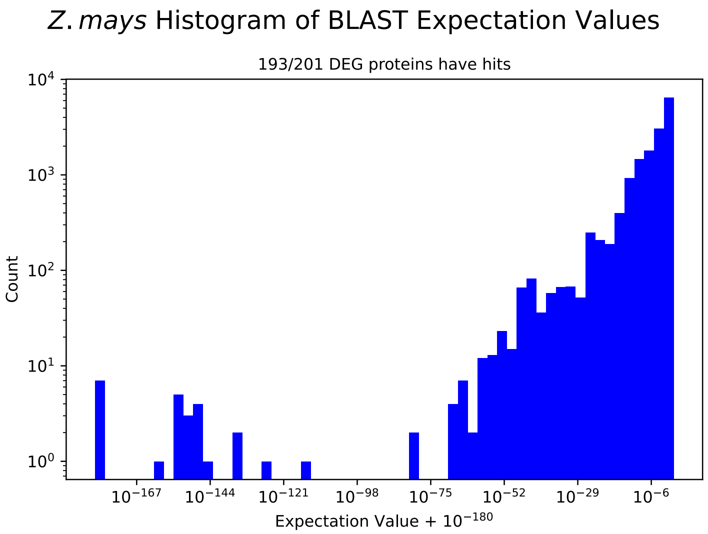


Figure S15: Expectation values for *Z.mays*, cf. Table S5.

#### S2.2 Agricultural Pathogens

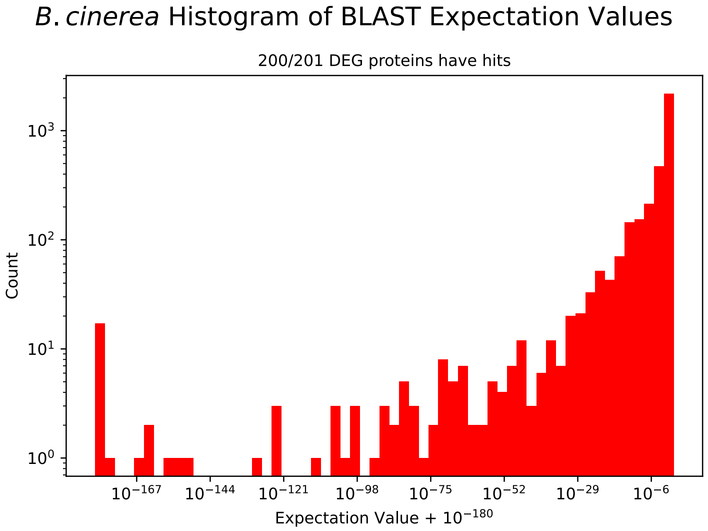


Figure S16: Expectation values for *B.cinerea*, cf. Table S5.

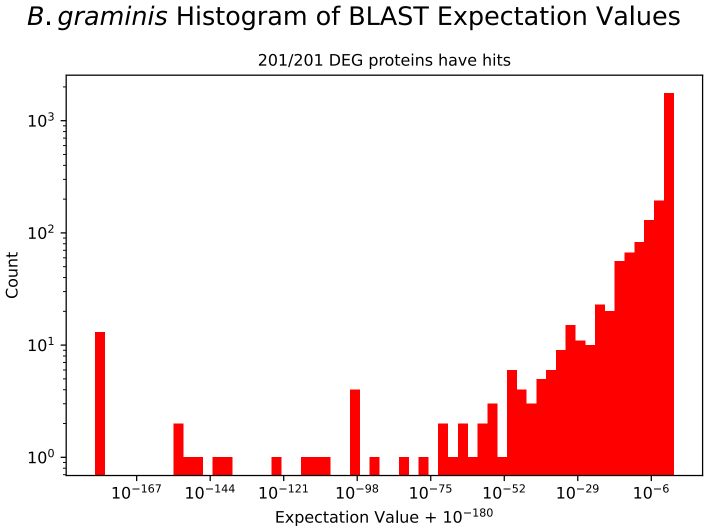


Figure S17: Expectation values for *B.graminis*, cf. Table S5.

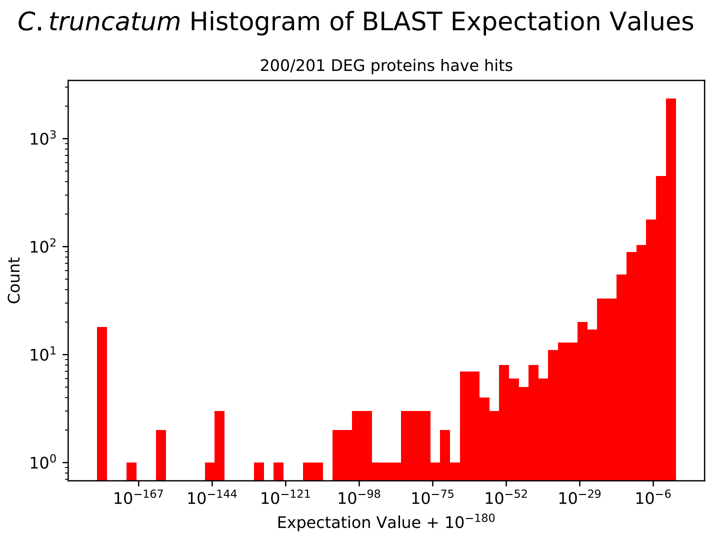


Figure S18: Expectation values for *C.truncatum*, cf. Table S5.

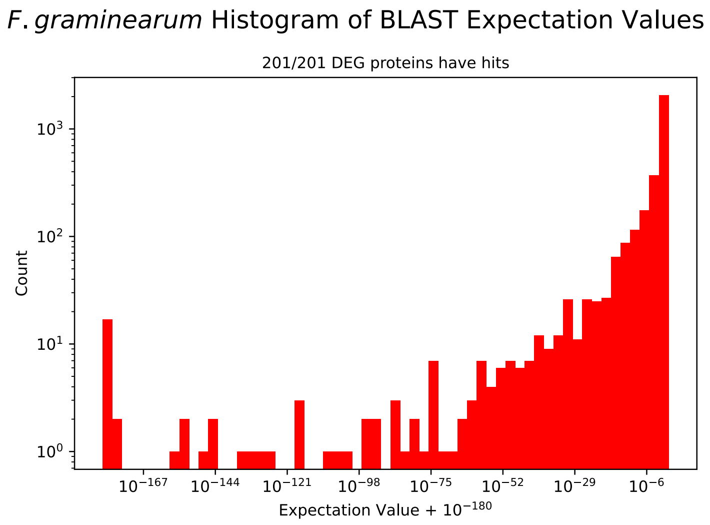


Figure S19: Expectation values for *F.graminearum*, cf. Table S5.

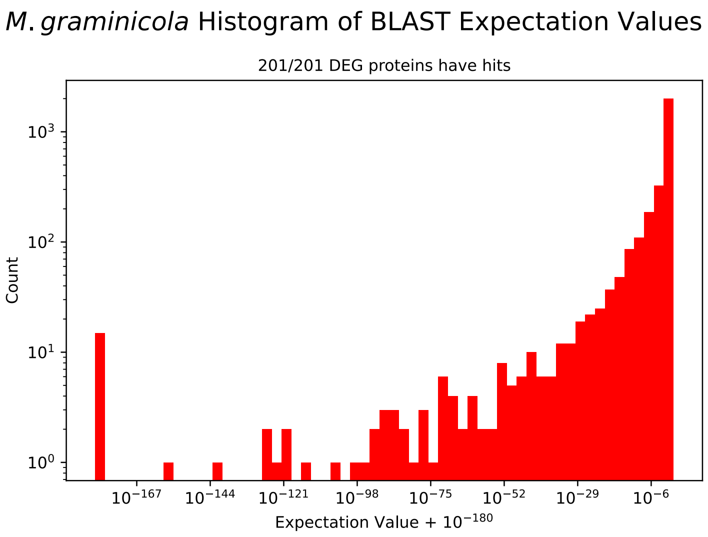


Figure S20: Expectation values for *M.graminicola*, cf. Table S5.

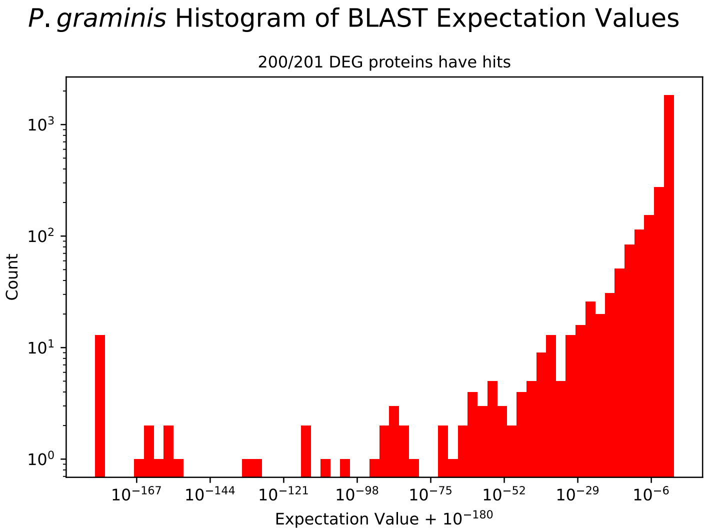


Figure S21: Expectation values for *P.graminis*, cf. Table S5.

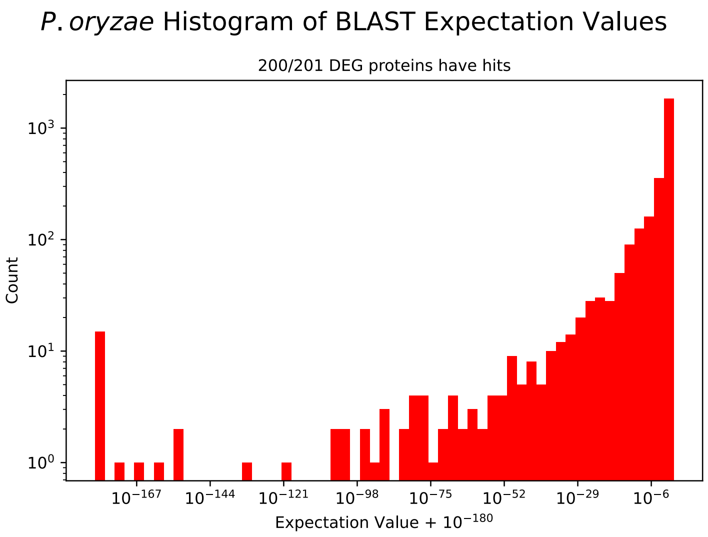


Figure S22: Expectation values for *P.oryzae*, cf. Table S5.

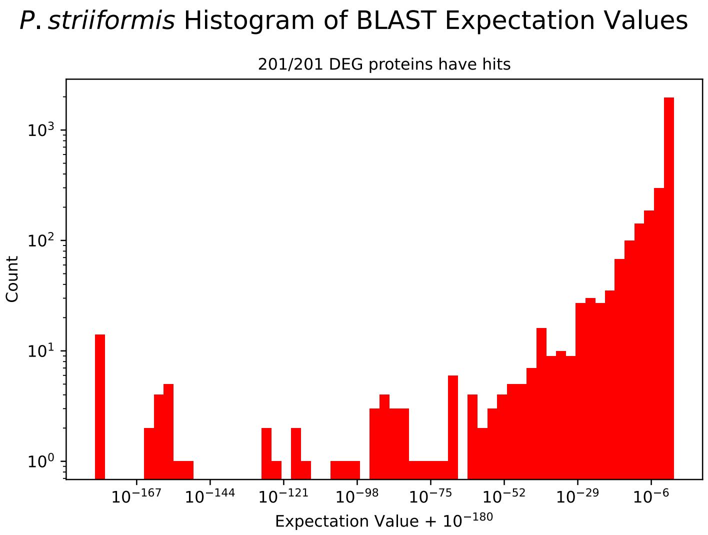


Figure S23: Expectation values for *P.striiformis*, cf. Table S5.

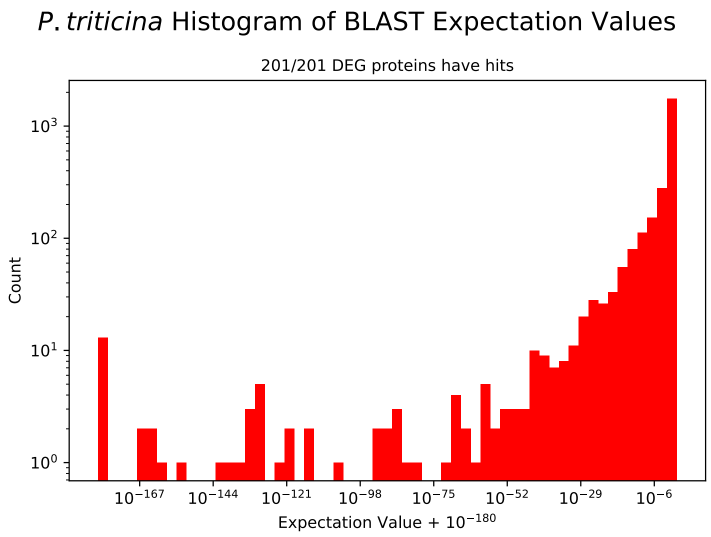


Figure S24: Expectation values for *P.triticina*, cf. Table S5.

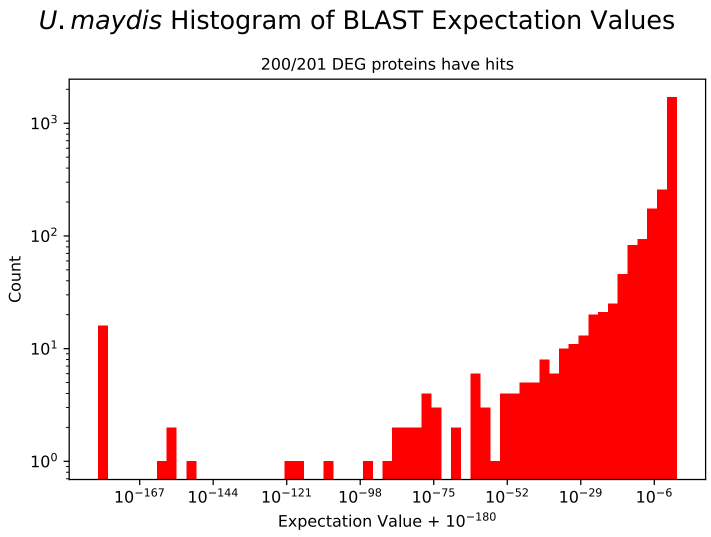


Figure S25: Expectation values for *U.maydis*, cf. Table S5.

| Species | $\leq 10^{-180}$ | $\leq 10^{-167}$ | $\leq 10^{-144}$ | $\leq 10^{-121}$ | $\leq 10^{-98}$ | $\leq 10^{-75}$ | $\leq 10^{-52}$ | $\leq 10^{-29}$ | $\leq 10^{-6}$ | $\leq 10^{-1}$ |
|---|---|---|---|---|---|---|---|---|---|---|
| *A.fumigatus* | 15 | 19 | 20 | 23 | 29 | 41 | 72 | 143 | 613 | 1,244 |
| *C.albicans* | 19 | 22 | 32 | 34 | 38 | 60 | 83 | 164 | 636 | 1,064 |
| *C.auris* | 21 | 24 | 29 | 33 | 40 | 57 | 83 | 172 | 636 | 1,013 |
| *C.neoformans.B.3501A* | 11 | 13 | 16 | 18 | 20 | 33 | 46 | 85 | 482 | 939 |
| *C.neoformans.JEC21* | 10 | 12 | 15 | 17 | 20 | 33 | 46 | 92 | 509 | 983 |
| *C.neoformans.grubii.H99* | 11 | 13 | 16 | 17 | 21 | 36 | 53 | 99 | 552 | 1,057 |
| *C.parapsilosis* | 21 | 25 | 31 | 37 | 41 | 63 | 92 | 170 | 632 | 1,074 |
| *C.tropicalis* | 17 | 23 | 28 | 35 | 39 | 57 | 85 | 168 | 627 | 1,071 |
| *G.max* | 3 | 3 | 26 | 34 | 38 | 40 | 100 | 676 | 7,405 | 15,703 |
| *H.capsulatum* | 14 | 15 | 17 | 20 | 22 | 32 | 45 | 103 | 533 | 1,117 |
| *H. sapiens* | 1 | 1 | 1 | 1 | 1 | 1 | 165 | 1,538 | 12,762 | 25,391 |
| *N.glabratus* | 52 | 60 | 70 | 87 | 110 | 134 | 166 | 268 | 733 | 1,122 |
| *O.sativa* | 3 | 3 | 17 | 22 | 23 | 25 | 58 | 307 | 3,691 | 8,756 |
| *S.tuberosum* | 3 | 3 | 11 | 14 | 16 | 17 | 50 | 321 | 3,265 | 7,327 |
| *Z.mays* | 7 | 7 | 21 | 24 | 25 | 27 | 87 | 492 | 5,188 | 10,501 |
| *B.cinerea* | 17 | 18 | 24 | 28 | 34 | 52 | 83 | 154 | 806 | 1,650 |
| *B.graminis* | 12 | 13 | 17 | 20 | 26 | 30 | 42 | 92 | 448 | 824 |
| *C.truncatum* | 18 | 19 | 22 | 27 | 35 | 52 | 80 | 150 | 604 | 1,471 |
| *F.graminearum* | 16 | 19 | 24 | 29 | 35 | 47 | 76 | 159 | 625 | 1,316 |

Table S5: Pairwise alignment info from *S. cerevisiae* query proteins.

### S3 Amino Acid Similarity

| | A | C | D | E | F | G | H | I | K | L | M | N | P | Q | R | S | T | V | W | Y |
|---|---|---|---|---|---|---|---|---|---|---|---|---|---|---|---|---|---|---|---|---|
| A | 1 | 0.578 | 0.156 | 0.159 | 0.356 | 0.659 | 0.32 | 0.588 | 0.426 | 0.643 | 0.421 | 0.28 | 0.553 | 0.356 | 0.315 | 0.477 | 0.417 | 0.675 | 0.224 | 0.214 |
| C | 0.578 | 1 | 0.243 | 0.221 | 0.289 | 0.29 | 0.288 | 0.348 | 0.269 | 0.398 | 0.612 | 0.485 | 0.318 | 0.482 | 0.324 | 0.613 | 0.485 | 0.43 | 0.188 | 0.223 |
| D | 0.156 | 0.243 | 1 | 0.84 | 0.172 | 0.015 | 0.2 | 0.338 | 0.248 | 0.387 | 0.292 | 0.637 | 0.015 | 0.492 | 0.236 | 0.33 | 0.254 | 0.279 | 0.028 | 0.287 |
| E | 0.159 | 0.221 | 0.84 | 1 | 0.219 | 0.049 | 0.201 | 0.366 | 0.435 | 0.333 | 0.402 | 0.578 | 0.003 | 0.685 | 0.382 | 0.312 | 0.218 | 0.239 | 0.086 | 0.261 |
| F | 0.356 | 0.289 | 0.172 | 0.219 | 1 | 0.259 | 0.605 | 0.487 | 0.381 | 0.57 | 0.465 | 0.34 | 0.282 | 0.459 | 0.339 | 0.38 | 0.254 | 0.38 | 0.741 | 0.729 |
| G | 0.659 | 0.29 | 0.015 | 0.049 | 0.259 | 1 | 0.183 | 0.371 | 0.309 | 0.38 | 0.149 | 0.19 | 0.499 | 0.163 | 0.149 | 0.323 | 0.396 | 0.437 | 0.138 | 0.163 |
| H | 0.32 | 0.288 | 0.2 | 0.201 | 0.605 | 0.183 | 1 | 0.368 | 0.421 | 0.45 | 0.345 | 0.459 | 0.172 | 0.406 | 0.396 | 0.342 | 0.208 | 0.3 | 0.484 | 0.504 |
| I | 0.588 | 0.348 | 0.338 | 0.366 | 0.487 | 0.371 | 0.368 | 1 | 0.477 | 0.889 | 0.494 | 0.456 | 0.419 | 0.453 | 0.342 | 0.36 | 0.493 | 0.843 | 0.287 | 0.266 |
| K | 0.426 | 0.269 | 0.248 | 0.435 | 0.381 | 0.309 | 0.421 | 0.477 | 1 | 0.492 | 0.482 | 0.401 | 0.295 | 0.545 | 0.733 | 0.285 | 0.224 | 0.419 | 0.297 | 0.285 |
| L | 0.643 | 0.398 | 0.387 | 0.333 | 0.57 | 0.38 | 0.45 | 0.889 | 0.492 | 1 | 0.515 | 0.506 | 0.432 | 0.501 | 0.36 | 0.411 | 0.432 | 0.785 | 0.368 | 0.347 |
| M | 0.421 | 0.612 | 0.292 | 0.402 | 0.465 | 0.149 | 0.345 | 0.494 | 0.482 | 0.515 | 1 | 0.518 | 0.265 | 0.699 | 0.522 | 0.48 | 0.409 | 0.465 | 0.355 | 0.307 |
| N | 0.28 | 0.485 | 0.637 | 0.578 | 0.34 | 0.19 | 0.459 | 0.456 | 0.401 | 0.506 | 0.518 | 1 | 0.172 | 0.589 | 0.427 | 0.581 | 0.488 | 0.395 | 0.316 | 0.391 |
| P | 0.553 | 0.318 | 0.015 | 0.003 | 0.282 | 0.499 | 0.172 | 0.419 | 0.295 | 0.432 | 0.265 | 0.172 | 1 | 0.168 | 0.155 | 0.321 | 0.33 | 0.473 | 0.211 | 0.179 |
| Q | 0.356 | 0.482 | 0.492 | 0.685 | 0.459 | 0.163 | 0.406 | 0.453 | 0.545 | 0.501 | 0.699 | 0.589 | 0.168 | 1 | 0.561 | 0.501 | 0.34 | 0.416 | 0.353 | 0.368 |
| R | 0.315 | 0.324 | 0.236 | 0.382 | 0.339 | 0.149 | 0.396 | 0.342 | 0.733 | 0.36 | 0.522 | 0.427 | 0.155 | 0.561 | 1 | 0.317 | 0.258 | 0.307 | 0.288 | 0.407 |
| S | 0.477 | 0.613 | 0.33 | 0.312 | 0.38 | 0.323 | 0.342 | 0.36 | 0.285 | 0.411 | 0.48 | 0.581 | 0.321 | 0.501 | 0.317 | 1 | 0.668 | 0.439 | 0.243 | 0.354 |
| T | 0.417 | 0.485 | 0.254 | 0.218 | 0.254 | 0.396 | 0.208 | 0.493 | 0.224 | 0.432 | 0.409 | 0.488 | 0.33 | 0.34 | 0.258 | 0.668 | 1 | 0.551 | 0.176 | 0.285 |
| V | 0.675 | 0.43 | 0.279 | 0.239 | 0.38 | 0.437 | 0.3 | 0.843 | 0.419 | 0.785 | 0.465 | 0.395 | 0.473 | 0.416 | 0.307 | 0.439 | 0.551 | 1 | 0.195 | 0.199 |
| W | 0.224 | 0.188 | 0.028 | 0.086 | 0.741 | 0.138 | 0.484 | 0.287 | 0.297 | 0.368 | 0.355 | 0.316 | 0.211 | 0.353 | 0.288 | 0.243 | 0.176 | 0.195 | 1 | 0.565 |
| Y | 0.214 | 0.223 | 0.287 | 0.261 | 0.729 | 0.163 | 0.504 | 0.266 | 0.285 | 0.347 | 0.307 | 0.391 | 0.179 | 0.368 | 0.407 | 0.354 | 0.285 | 0.199 | 0.565 | 1 |

Table S6: Similarity between amino acids can be quantified by Sneath's index $\phi$, where identical amino acids have a value of 1, and the greater the index between two amino acids is, the more similar the amino acids are to one another..

### S4 Best Targetable Proteins

#### S4.1 Alr1

##### S4.1.1 WHO Critical Pathogens

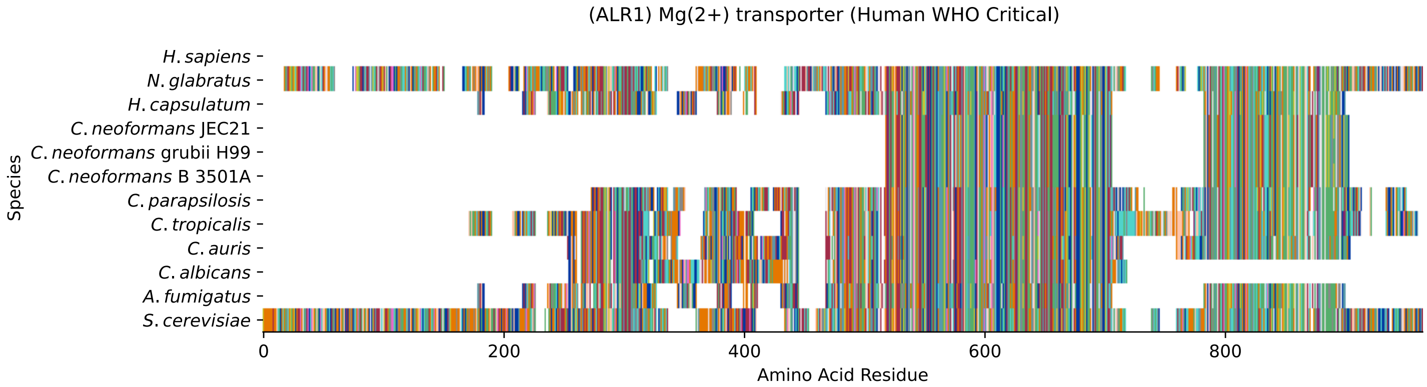


Figure S26: Multiple sequence alignment of yeast Alr1 (WHO Critical Pathogens). Cf. Figure S27 for alignment quality, and Figure S28 for Sneath similarity. Cf. Table S7 for protein names, and pairwise alignment metrics with yeast Alr1.

#### Alr1 MSA Quality

| | *A. f.* | *C. a.* | *C. a.* | *C. p.* | *C. t.* | *C. n.* | *C. n.* | *C. n.* | *H. c.* | *N. g.* | *S. c.* |
|---|---|---|---|---|---|---|---|---|---|---|---|
| *A. fumigatus* | 1062.75 | 285.419 | 398.282 | 378.252 | 294.278 | 371.318 | 372.089 | 371.318 | 735.717 | 406.371 | 393.275 |
| *C. albicans* | 269.626 | 903.279 | 526.554 | 503.442 | 650.203 | 254.603 | 254.603 | 254.603 | 276.559 | 446.047 | 454.907 |
| *C. auris* | 379.022 | 520.776 | 1152.5 | 581.637 | 662.914 | 366.311 | 366.696 | 366.311 | 409.453 | 584.719 | 579.326 |
| *C. parapsilosis* | 372.474 | 495.352 | 598.201 | 1176.39 | 503.056 | 360.533 | 360.533 | 360.533 | 401.749 | 582.793 | 579.711 |
| *C. tropicalis* | 275.404 | 647.506 | 686.411 | 676.781 | 1389.02 | 254.603 | 254.603 | 254.603 | 296.975 | 614.379 | 637.876 |
| *C. neoformans* B 3501A | 356.295 | 253.832 | 366.311 | 350.903 | 254.603 | 643.269 | 641.728 | 643.269 | 363.999 | 357.066 | 338.961 |
| *C. neoformans* grubii H99 | 357.066 | 253.832 | 366.696 | 350.903 | 254.603 | 641.728 | 642.114 | 641.728 | 362.073 | 357.066 | 339.347 |
| *C. neoformans* JEC21 | 356.295 | 253.832 | 366.311 | 350.903 | 254.603 | 643.269 | 641.728 | 643.269 | 363.999 | 357.066 | 338.961 |
| *H. capsulatum* | 709.523 | 275.789 | 409.453 | 385.956 | 296.975 | 363.999 | 362.073 | 363.999 | 1058.9 | 406.371 | 404.445 |
| *N. glabratus* | 387.111 | 453.751 | 602.823 | 589.726 | 613.994 | 357.836 | 357.836 | 357.836 | 411.764 | 1640.55 | 922.924 |
| *S. cerevisiae* | 385.571 | 456.833 | 600.127 | 582.408 | 637.491 | 351.673 | 352.829 | 351.673 | 414.846 | 926.776 | 1796.17 |

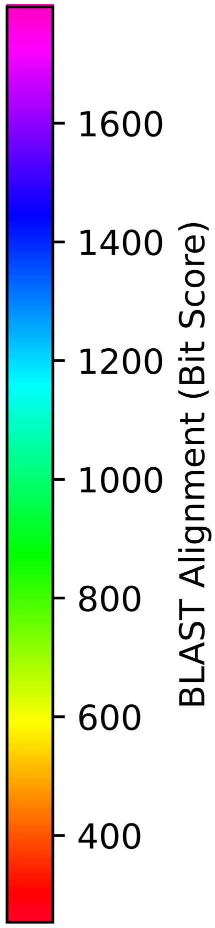


Figure S27: Multiple sequence alignment quality of Alr1 (WHO Critical Pathogens). Cf. Figures S26 and S29.

| Species | Hit Protein | Hit Length (a.a.) | evalue | align_len | bit_score | identity | positive | score | gaps | % identity | % positive |
|---|---|---|---|---|---|---|---|---|---|---|---|
| H.sapiens | - | - | - | - | - | - | - | - | - | - | - |
| N.glabratus | XP_445759.1 uncharacterized p-rotein CAGL0E01617g Nakaseomyc-es glabratus | 856 | 0 | 856 | 927.161 | 513 | 605 | 2395 | 85 | 59.7 | 70.4 |
| H.capsulatum | XP_045290975.1 magnesium tran-sporter ALR1 Histoplasma capsu-latum G186AR | 601 | 1.6e-135 | 601 | 417.157 | 242 | 337 | 1071 | 98 | 28.2 | 39.2 |
| C.neoformans.JEC21 | XP_571831.1 hypothetical prot-ein CNG00910 Cryptococcus neof-ormans var. neoformans JEC21 | 350 | 1.6e-114 | 350 | 366.311 | 169 | 234 | 939 | 44 | 19.7 | 27.2 |
| C.neoformans.grubii.H99 | XP_012051300.1 magnesium tran-sporter Cryptococcus neoforman-s var. grubii H99 | 350 | 9e-115 | 350 | 367.081 | 169 | 234 | 941 | 44 | 19.7 | 27.2 |
| C.neoformans.B.3501A | XP_774407.1 hypothetical prot-ein CNBG3880 Cryptococcus neof-ormans var. neoformans B-3501A | 350 | 1.5e-114 | 350 | 366.311 | 169 | 234 | 939 | 44 | 19.7 | 27.2 |
| C.parapsilosis | XP_036663332.1 uncharacterize-d protein CPAR2 101870 Candida parapsilosis | 591 | 0 | 591 | 585.874 | 320 | 394 | 1509 | 48 | 37.3 | 45.9 |
| C.tropicalis | XP_002548119.1 magnesium tran-sporter ALR1 Candida tropicali-s MYA-3404 | 727 | 0 | 727 | 639.032 | 368 | 467 | 1647 | 120 | 42.8 | 54.4 |
| C.auris | XP_028892080.2 hypothetical p-rotein Candida auris | 600 | 0 | 600 | 605.905 | 328 | 409 | 1561 | 91 | 38.2 | 47.6 |
| C.albicans | XP_721686.1 Mg(2+) transporte-r Candida albicans SC5314 | 443 | 2.2e-149 | 443 | 461.455 | 244 | 293 | 1186 | 49 | 28.4 | 34.1 |
| A.fumigatus | XP_754049.1 CorA family metal ion transporter, putative Asp-ergillus fumigatus Af293 | 601 | 6.9e-124 | 601 | 387.111 | 241 | 331 | 993 | 94 | 28.1 | 38.5 |

Table S7: Pairwise alignment info from yeast Alr1 (DEG20010935), cf. Figure S26.

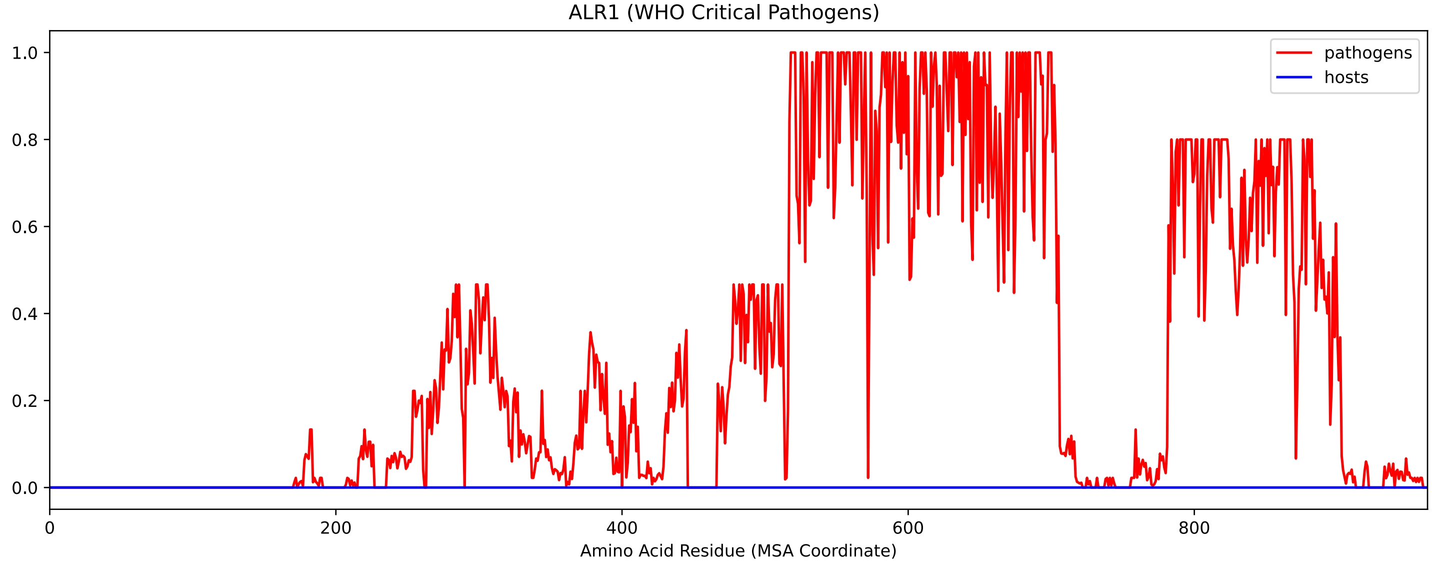


Figure S28: Sneath Similarity of Alr1 for WHO Critical Pathogens, cf. Figure S26

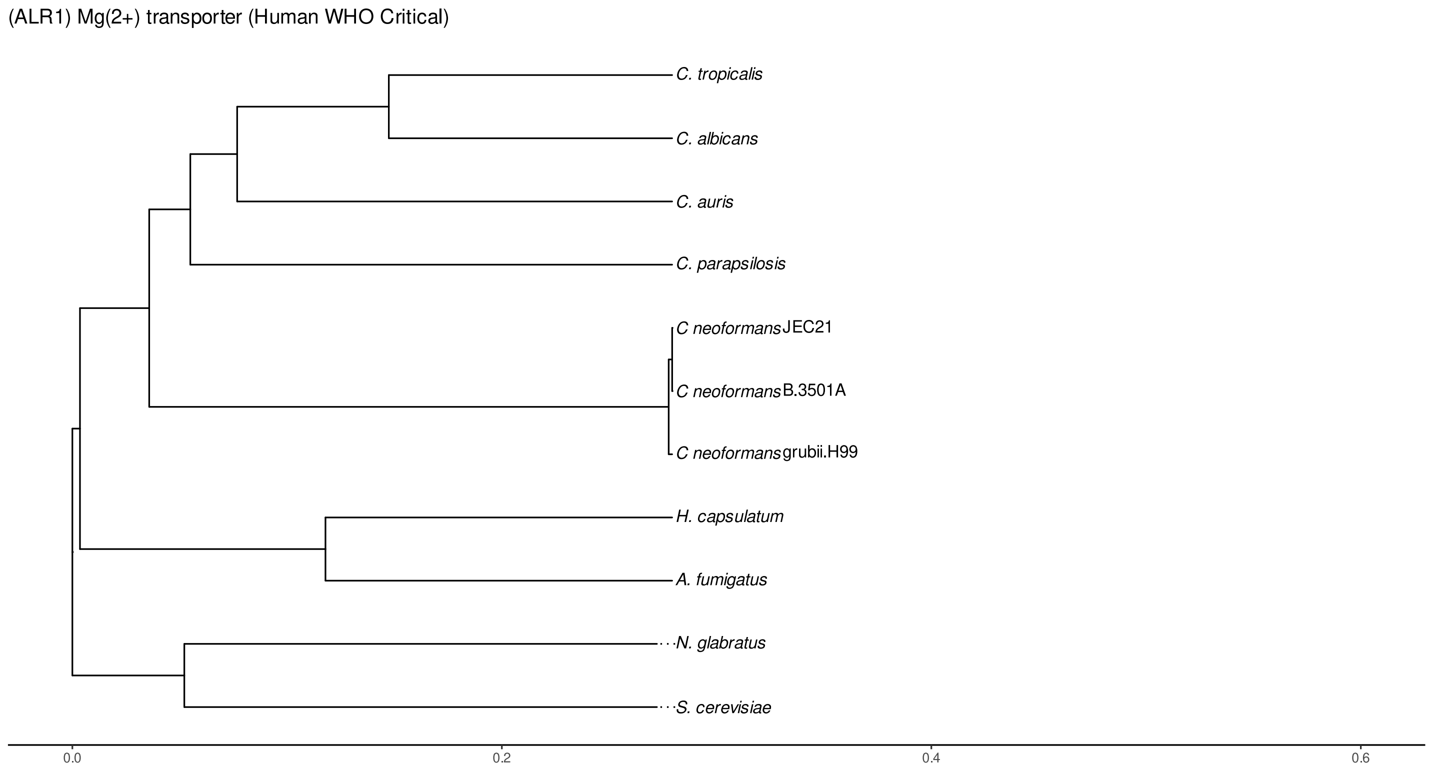


Figure S29: Phylogeny of hits for Alr1 made using clustal omega and visualized using ggtree, cf. Figures S26 and S27.

##### S4.1.2 Top 10 Agricultural Fungal Pathogens

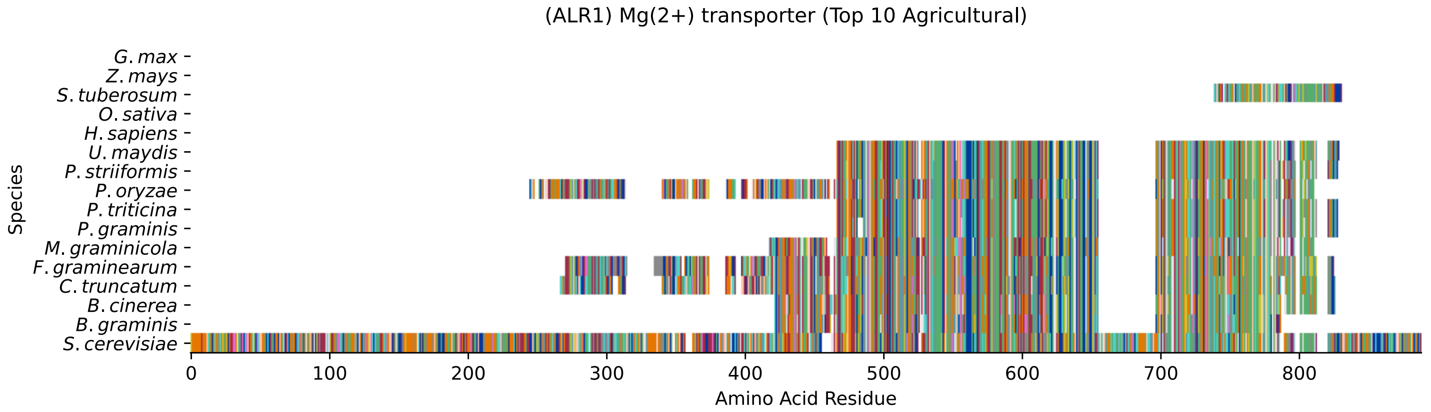


Figure S30: Multiple sequence alignment of yeast Alr1 (Top 10 Agricultural Fungal Pathogens). Cf. Figure S31 for alignment quality, and Figure S32 for Sneath similarity. Cf. Table S8 for protein names, and pairwise alignment metrics with yeast Alr1.

### Alr1 MSA Quality

| | *B. g.* | *B. c.* | *C. t.* | *F. g.* | *M. g.* | *P. g.* | *P. t.* | *P. o.* | *P. s.* | *U. m.* | *S. c.* |
|---|---|---|---|---|---|---|---|---|---|---|---|
| *B. graminis* | 670.233 | 546.199 | 518.079 | 505.368 | 409.453 | 330.872 | 340.887 | 513.457 | 336.265 | 329.717 | 334.724 |
| *B. cinerea* | 546.199 | 726.857 | 527.709 | 501.516 | 405.216 | 326.635 | 333.183 | 493.812 | 330.872 | 329.331 | 325.094 |
| *C. truncatum* | 518.079 | 527.709 | 942.184 | 649.818 | 417.157 | 329.717 | 338.961 | 615.15 | 335.109 | 328.946 | 344.354 |
| *F. graminearum* | 507.679 | 503.827 | 659.833 | 959.518 | 404.831 | 325.479 | 336.265 | 569.311 | 331.643 | 333.183 | 337.806 |
| *M. graminicola* | 409.453 | 405.216 | 417.157 | 402.905 | 707.597 | 348.206 | 354.755 | 419.468 | 352.443 | 354.369 | 375.17 |
| *P. graminis* | 330.872 | 326.635 | 329.717 | 323.939 | 348.206 | 629.787 | 614.379 | 335.495 | 620.542 | 549.28 | 337.806 |
| *P. triticina* | 340.887 | 333.183 | 338.961 | 335.109 | 354.755 | 614.379 | 637.876 | 342.428 | 623.239 | 556.984 | 345.125 |
| *P. oryzae* | 513.457 | 493.812 | 615.15 | 557.37 | 419.468 | 335.495 | 342.428 | 965.296 | 340.887 | 337.421 | 328.176 |
| *P. striiformis* | 336.265 | 330.872 | 335.109 | 329.717 | 352.443 | 620.542 | 623.239 | 340.887 | 637.876 | 560.836 | 350.903 |
| *U. maydis* | 329.717 | 329.331 | 328.946 | 332.028 | 354.369 | 549.28 | 556.984 | 337.421 | 560.836 | 637.106 | 347.436 |
| *S. cerevisiae* | 335.88 | 332.413 | 353.599 | 340.887 | 387.497 | 343.584 | 352.058 | 338.191 | 357.066 | 359.377 | 1796.17 |

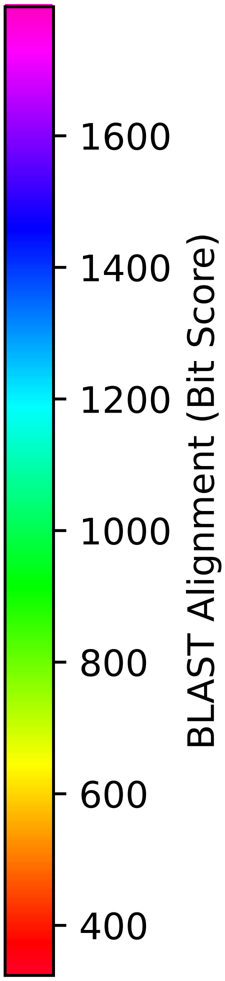


Figure S31: Multiple sequence alignment quality of Alr1 (Top 10 Agricultural Fungal Pathogens). Cf. Figures S30 and S33.

| Species | Hit Protein | Hit Length (a.a.) | evalue | align_len | bit_score | identity | positive | score | gaps | % identity | % positive |
|---|---|---|---|---|---|---|---|---|---|---|---|
| G.max | - | - | - | - | - | - | - | - | - | - | - |
| Z.mays | - | - | - | - | - | - | - | - | - | - | - |
| S.tuberosum | XP_006363963.1 PREDICTED: uncharacterized protein LOC102592-202 Solanum tuberosum | 93 | 0.016 | 93 | 38.891 | 31 | 48 | 89 | 15 | 3.6 | 5.6 |
| O.sativa | - | - | - | - | - | - | - | - | - | - | - |
| H.sapiens | - | - | - | - | - | - | - | - | - | - | - |
| U.maydis | XP_011386252.1 uncharacterized protein UMAG 00361 Ustilago maydis 521 | 348 | 9.2e-117 | 348 | 374.785 | 170 | 237 | 961 | 42 | 19.8 | 27.6 |
| P.striiformis | XP_047797628.1 hypothetical protein Pst134EA 030450 Puccinia striiformis f. sp. tritici | 348 | 3.5e-115 | 348 | 369.777 | 169 | 234 | 948 | 43 | 19.7 | 27.2 |
| P.oryzae | mRNA M BR32 EuGene 00004751-p1 — transcript=mRNA M BR32 EuGene 00004751 — gene=M BR32 EuGene 00004751 — organism=Pyricularia oryzae BR32 — gene product=unspecified product — transcript product=unspecified product — location=BR32 scaffold00001:1398493-1400442(-) — protein length=621 — sequence SO=supercontig — SO=protein coding gene — is pseudo=false | 560 | 1e-106 | 560 | 340.887 | 219 | 285 | 873 | 110 | 25.5 | 33.2 |
| P.triticina | XP_053028148.1 uncharacterized protein PtA15 17A74 Puccinia triticina | 348 | 9.9e-115 | 348 | 364.385 | 166 | 231 | 934 | 43 | 19.3 | 26.9 |
| P.graminis | XP_003336742.2 hypothetical protein PGTG 17997 Puccinia graminis f. sp. tritici CRL 75-36-700-3 | 348 | 2.9e-109 | 348 | 359.762 | 167 | 229 | 922 | 47 | 19.4 | 26.7 |
| M.graminicola | ZTRI 8.421.mRNA-p1 — transcript=ZTRI 8.421.mRNA — gene=ZTRI 8.421 — organism=Zymoseptoria tritici IPO323 — gene product=similar to cora family metal ion transporter — transcript product=similar to cora family metal ion transporter — location=Ztri chr 8:1277069-1279072(−) — protein length=667 — sequence SO=chromosome — SO=protein coding gene — is pseudo=false | 384 | 2.1e-127 | 384 | 396.741 | 196 | 252 | 1018 | 49 | 22.8 | 29.3 |
| F.graminearum | XP_011318426.1 hypothetical protein FGSG 02495 Fusarium graminearum PH-1 | 536 | 3.8e-109 | 536 | 347.821 | 202 | 286 | 891 | 84 | 23.5 | 33.3 |
| C.truncatum | XP_036584879.1 CorA-like Mg2+ transporter Colletotrichum truncatum | 538 | 8.3e-113 | 538 | 357.451 | 215 | 287 | 916 | 88 | 25.0 | 33.4 |
| B.cinerea | XP_024547928.1 hypothetical protein BCIN 03g07480 Botrytis cinerea B05.10 | 392 | 4.5e-107 | 392 | 342.813 | 171 | 237 | 878 | 52 | 19.9 | 27.6 |
| B.graminis | VDB86198.1 — transcript=BGT96224V316 LOCUS3808 t1 — gene=BGT96224V316 LOCUS3808 — organism=Blumeria graminis f. sp. tritici 96224 — gene product=unspecified product — transcript product=unspecified product — location=LR026988:17506578-17508517(+) — protein length=630 — sequence SO=chromosome — SO=protein coding gene — is pseudo=false | 362 | 3.8e-107 | 362 | 341.658 | 171 | 223 | 875 | 47 | 19.9 | 26.0 |

Table S8: Pairwise alignment info from yeast Alr1 (DEG20010935), cf. Figure S30.

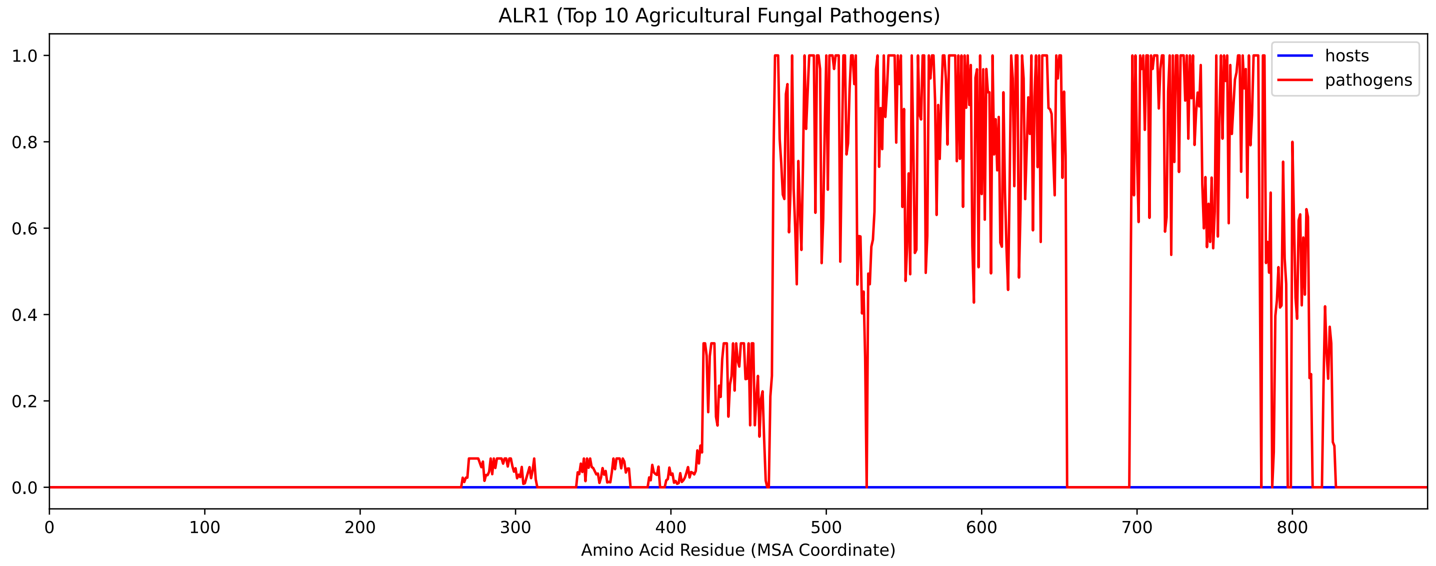


Figure S32: Sneath Similarity of Alr1 for Top 10 Agricultural Fungal Pathogens, cf. Figure S30

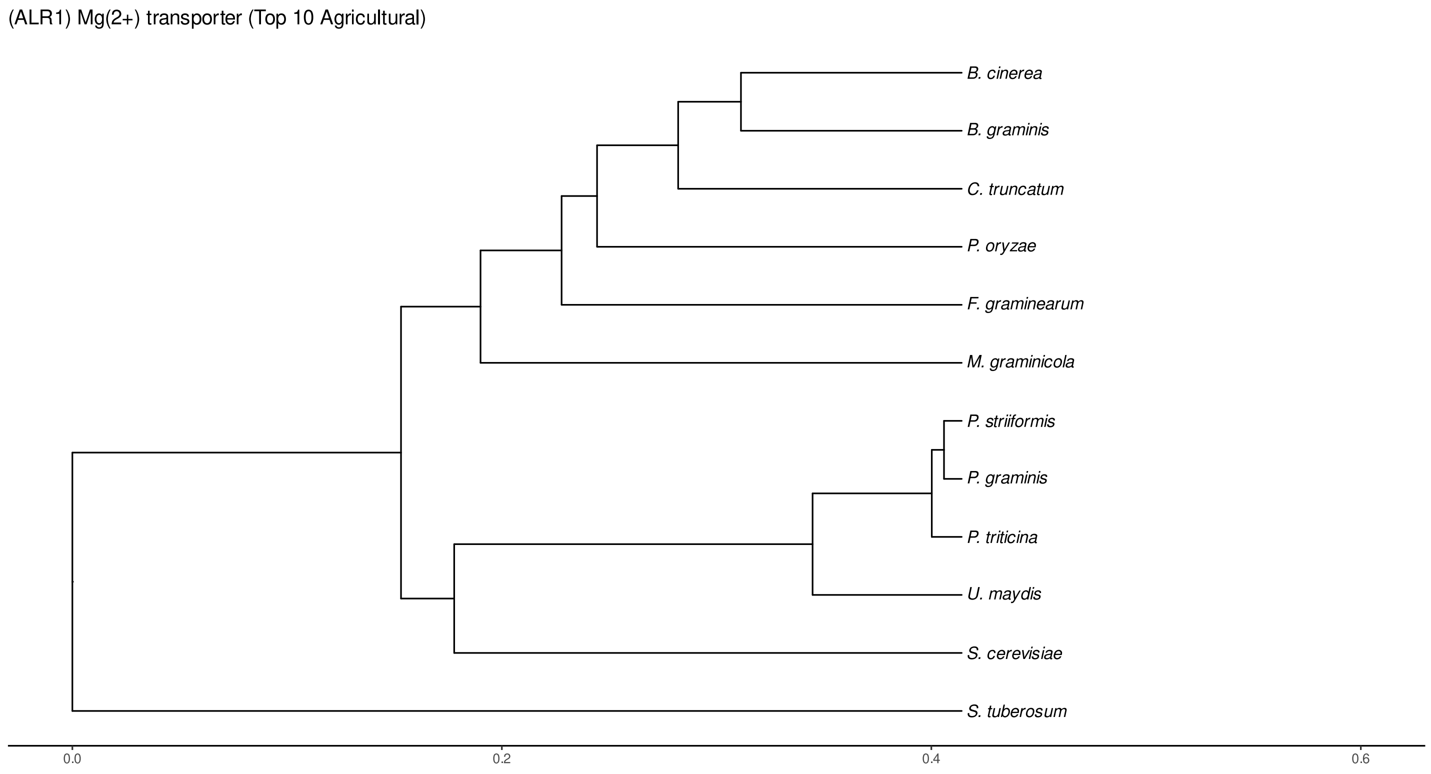


Figure S33: Phylogeny of hits for Alr1 made using clustal omega and visualized using ggtree, cf. Figures S30 and S31.

### S4.1.3 NR

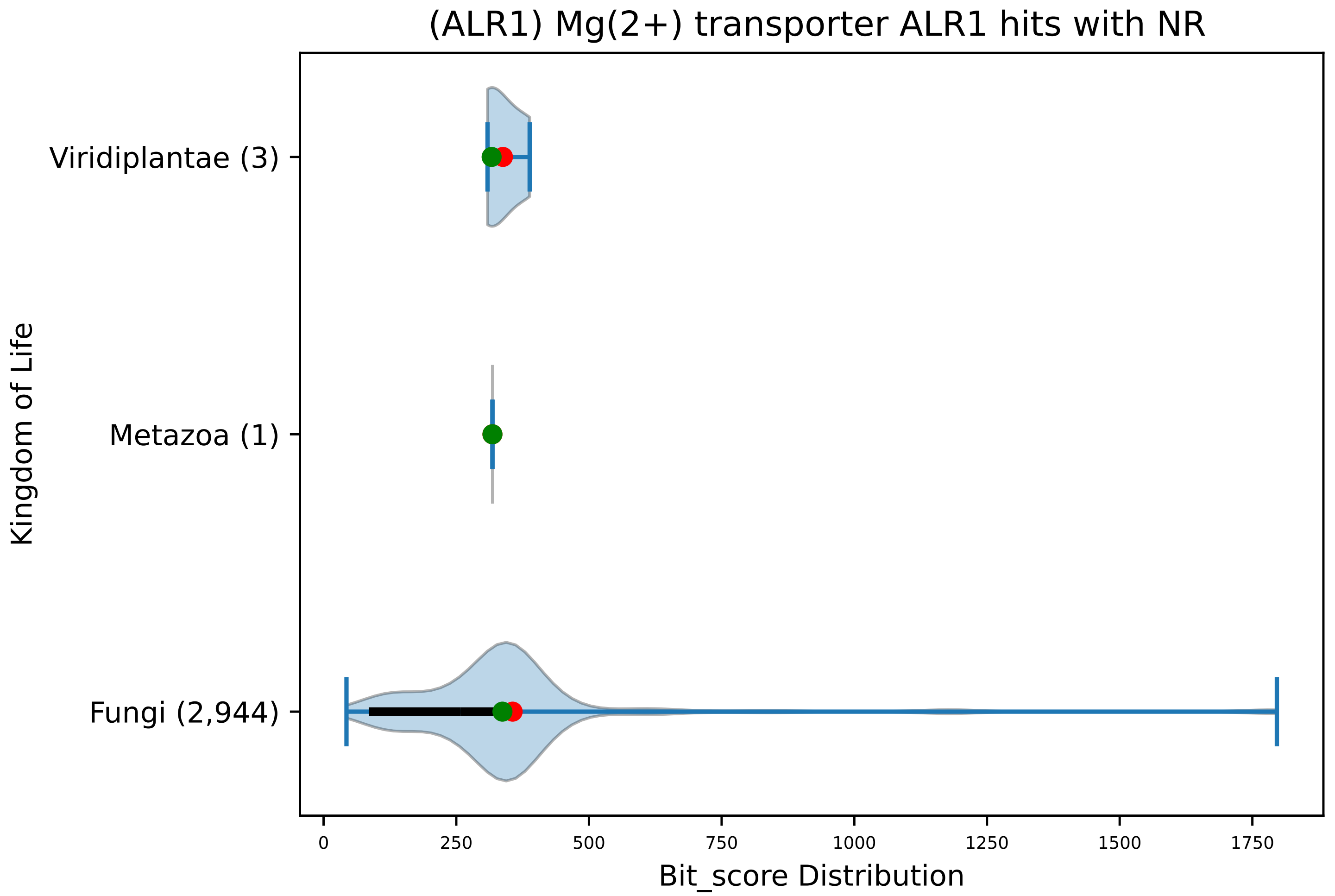


Figure S34: Non-redundant (NR) protein hits for DEG20010935/Alr1, with expectation value of no more than 0.1. Green points are medians, and red points are arithmetic means.

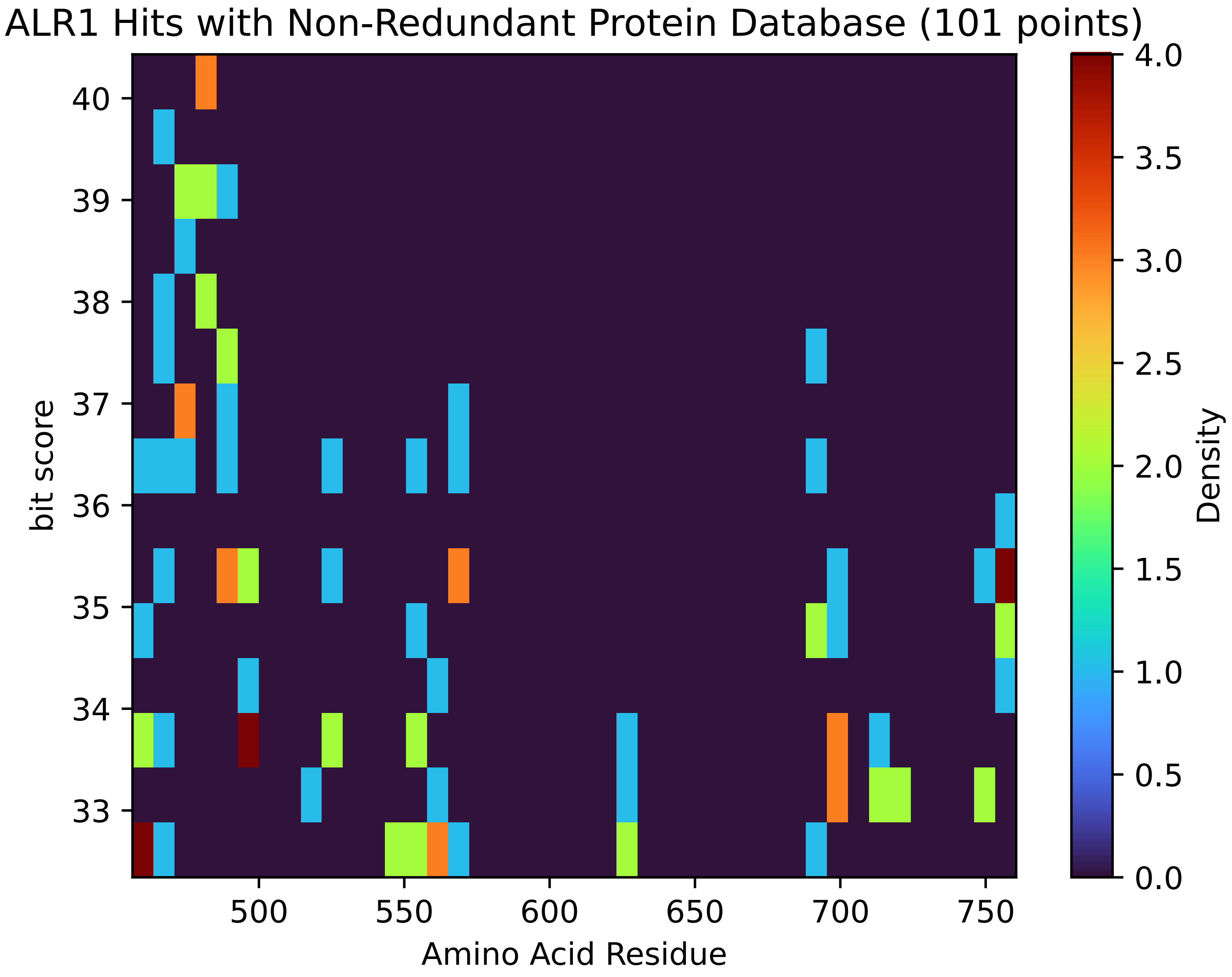


Figure S35: Non-redundant (NR) protein hits for Alr1 in the kingdom Metazoa.

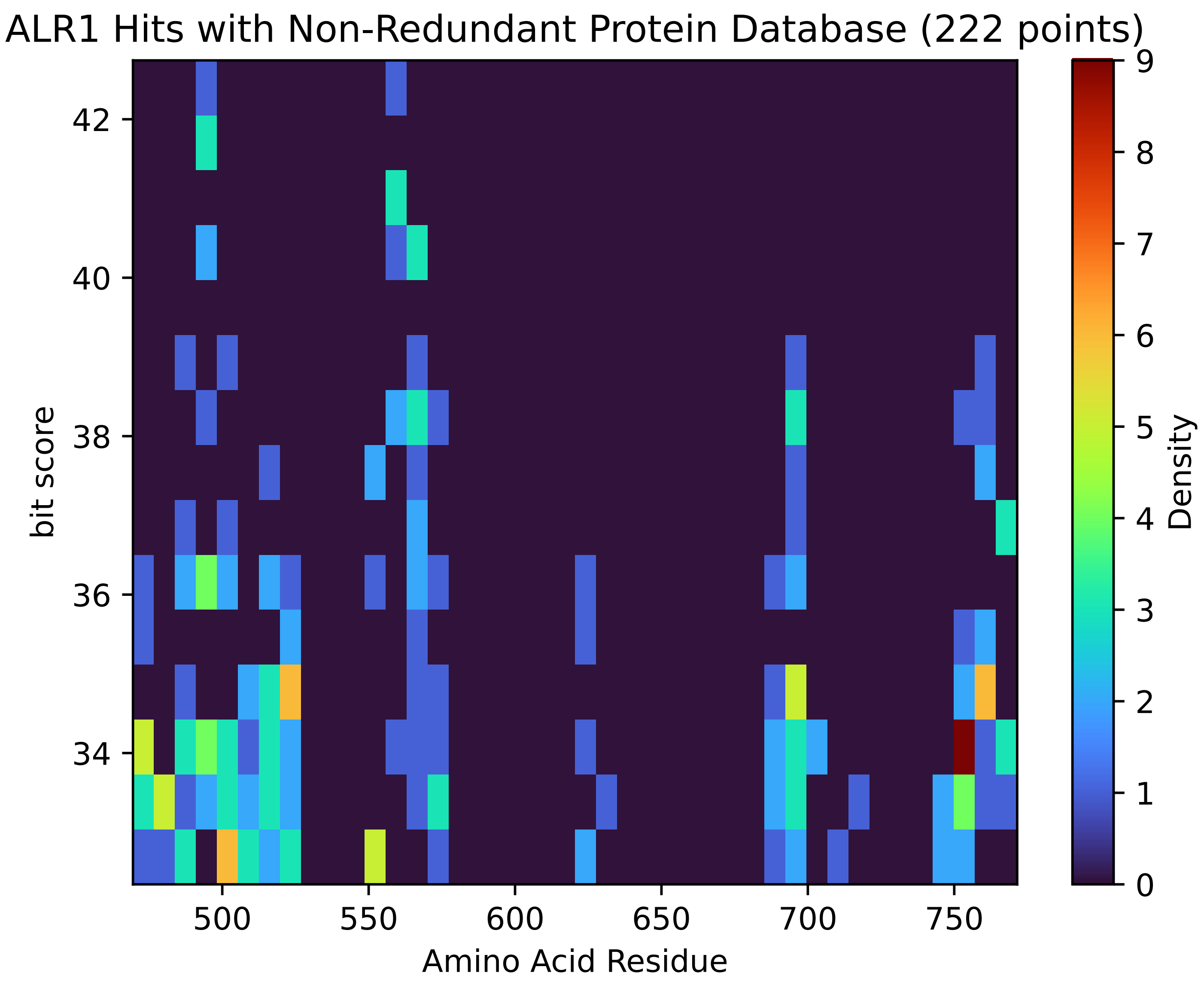


Figure S36: Non-redundant (NR) protein hits for Alr1 in the kingdom Viridiplantae.

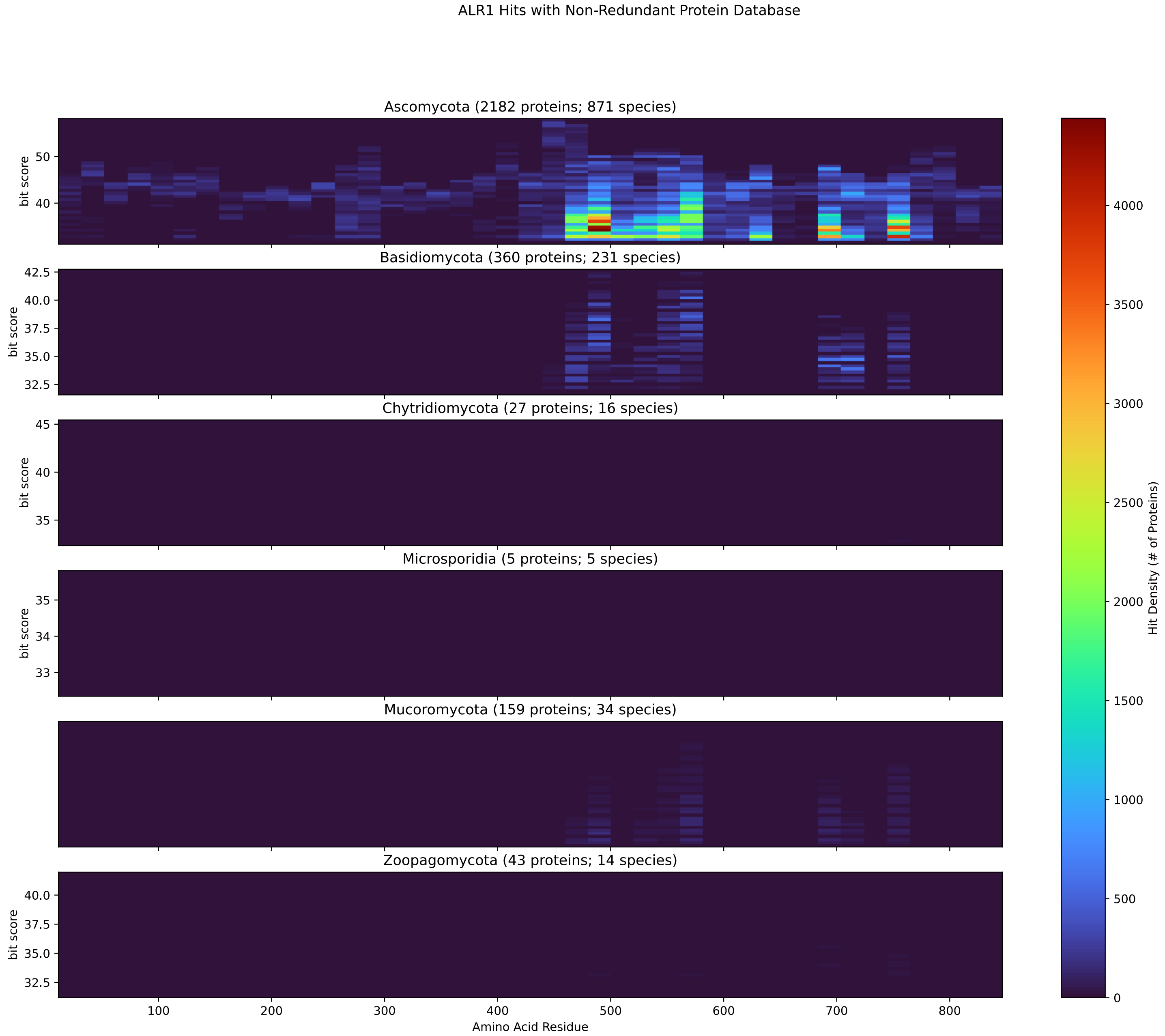


Figure S37: Non-redundant (NR) protein hits for Alr1 in the kingdom Fungi.

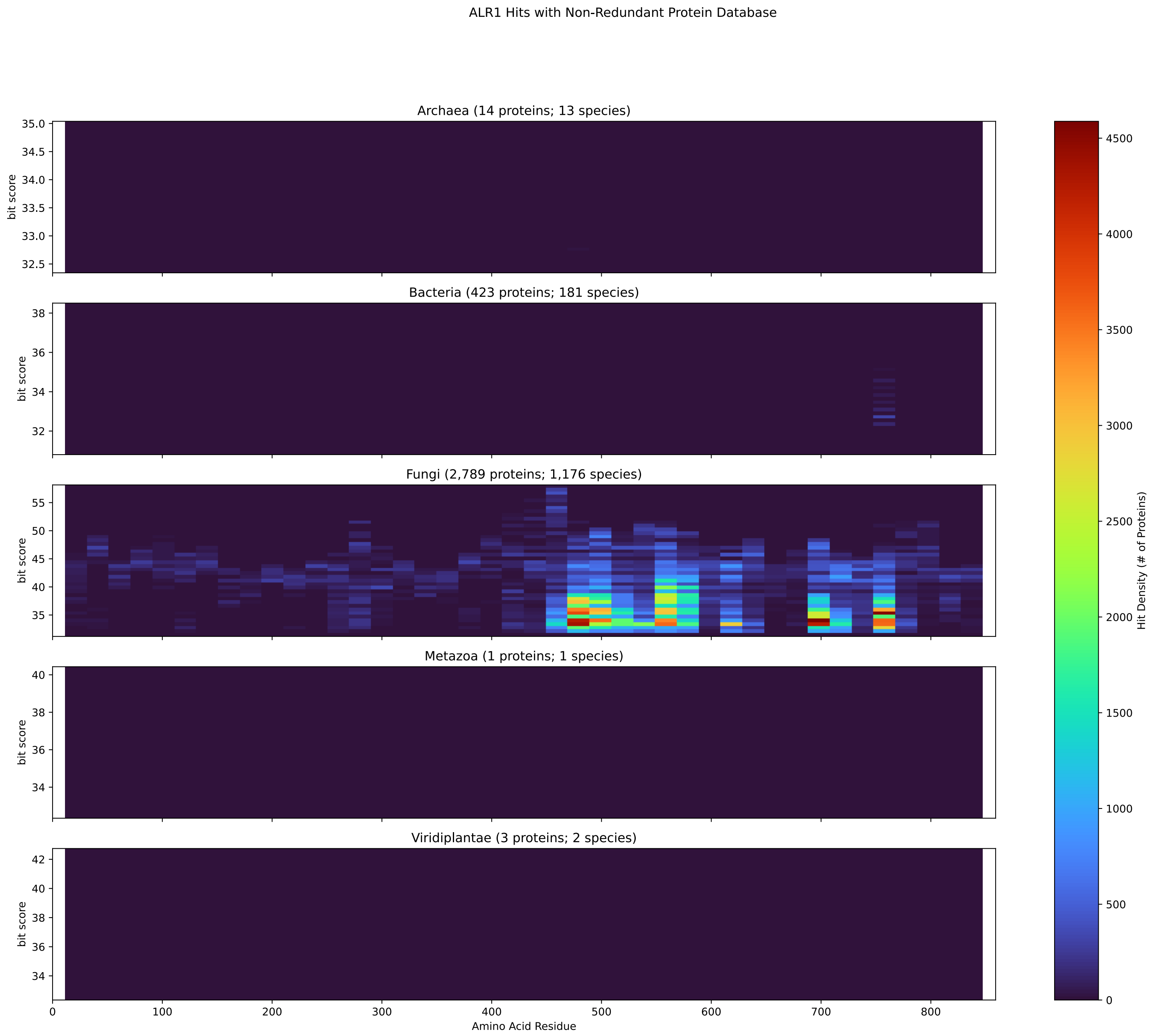


Figure S38: Non-redundant (NR) protein hits for DEG20010935/Alr1 at 20 amino acid length queries.

#### S4.2 Aur1

##### S4.2.1 WHO Critical Pathogens

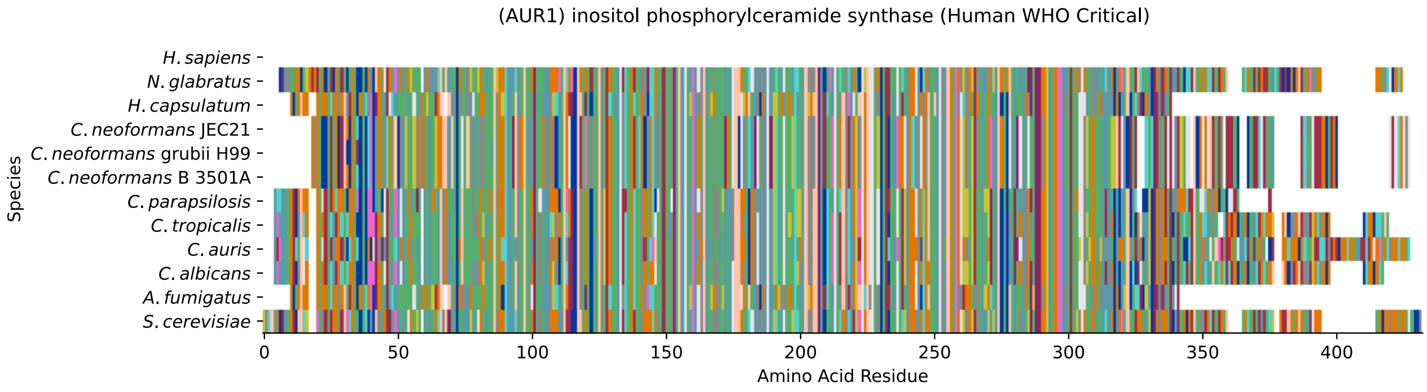


Figure S39: Multiple sequence alignment of yeast Aur1 (WHO Critical Pathogens). Cf. Figure S40 for alignment quality, and Figure S41 for Sneath similarity. Cf. Table S9 for protein names, and pairwise alignment metrics with yeast Aur1.

#### Aur1 MSA Quality

| | *A. f.* | *C. a.* | *C. a.* | *C. p.* | *C. t.* | *C. n.* | *C. n.* | *C. n.* | *H. c.* | *N. g.* | *S. c.* |
|---|---|---|---|---|---|---|---|---|---|---|---|
| *A. fumigatus* | 662.529 | 313.538 | 266.929 | 318.546 | 325.479 | 268.47 | 266.544 | 268.47 | 509.22 | 305.834 | 296.59 |
| *C. albicans* | 310.457 | 810.446 | 451.825 | 514.227 | 711.064 | 285.419 | 283.493 | 285.419 | 307.76 | 377.867 | 432.18 |
| *C. auris* | 272.322 | 466.848 | 843.958 | 443.351 | 465.307 | 280.411 | 281.182 | 280.411 | 271.937 | 362.459 | 390.578 |
| *C. parapsilosis* | 314.694 | 514.227 | 428.713 | 722.62 | 517.309 | 267.7 | 270.396 | 267.7 | 303.523 | 388.267 | 432.565 |
| *C. tropicalis* | 321.627 | 711.064 | 449.129 | 517.309 | 810.831 | 293.893 | 291.197 | 293.893 | 309.686 | 371.703 | 434.876 |
| *C. neoformans* B 3501A | 265.003 | 285.419 | 271.166 | 267.7 | 293.893 | 760.37 | 747.273 | 760.37 | 258.07 | 282.337 | 284.648 |
| *C. neoformans* grubii H99 | 263.462 | 283.493 | 271.937 | 270.396 | 291.197 | 747.273 | 758.829 | 747.273 | 257.299 | 283.878 | 283.108 |
| *C. neoformans* JEC21 | 265.003 | 285.419 | 271.166 | 267.7 | 293.893 | 760.37 | 747.273 | 760.37 | 258.07 | 282.337 | 284.648 |
| *H. capsulatum* | 499.975 | 307.76 | 262.692 | 303.523 | 309.686 | 258.07 | 257.299 | 258.07 | 653.284 | 294.664 | 296.204 |
| *N. glabratus* | 301.212 | 377.867 | 347.821 | 388.267 | 371.703 | 282.337 | 283.878 | 282.337 | 294.664 | 799.275 | 515.383 |
| *S. cerevisiae* | 293.123 | 432.18 | 376.326 | 432.565 | 434.876 | 284.648 | 283.108 | 284.648 | 296.204 | 515.383 | 817.379 |

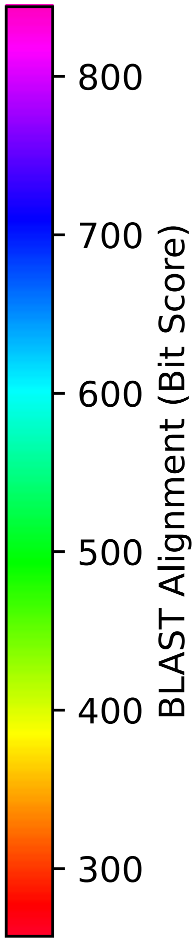


Figure S40: Multiple sequence alignment quality of Aur1 (WHO Critical Pathogens). Cf. Figures S39 and S42.

| Species | Hit Protein | Hit Length (a.a.) | evalue | align_len | bit_score | identity | positive | score | gaps | % identity | % positive |
|---|---|---|---|---|---|---|---|---|---|---|---|
| H.sapiens | - | - | - | - | - | - | - | - | - | - | - |
| N.glabratus | XP_448347.1 uncharacterized p-rotein CAGL0K02805g Nakaseomyc-es glabratus | 393 | 0 | 393 | 516.153 | 252 | 302 | 1328 | 5 | 62.8 | 75.3 |
| H.capsulatum | XP_045289424.1 aureobasidin r-esistance protein Aur1 Histopl-asma capsulatum G186AR | 326 | 7.5e-98 | 326 | 297.36 | 148 | 211 | 760 | 2 | 36.9 | 52.6 |
| C.neoformans.JEC21 | XP_572823.1 inositolphosphory-lceramide synthase, putative C-ryptococcus neoformans var. ne-oformans JEC21 | 379 | 4.2e-93 | 379 | 285.804 | 154 | 225 | 730 | 14 | 38.4 | 56.1 |
| C.neoformans.grubii.H99 | XP_012051809.1 inositolphosph-orylceramide synthase Cryptoco-ccus neoformans var. grubii H9-9 | 379 | 1.4e-92 | 379 | 284.648 | 151 | 224 | 727 | 14 | 37.7 | 55.9 |
| C.neoformans.B.3501A | XP_774053.1 hypothetical prot-ein CNBH0990 Cryptococcus neof-ormans var. neoformans B-3501A | 379 | 4.1e-93 | 379 | 285.804 | 154 | 225 | 730 | 14 | 38.4 | 56.1 |
| C.parapsilosis | XP_036664447.1 uncharacterize-d protein CPAR2 302580 Candida parapsilosis | 357 | 1e-151 | 357 | 436.802 | 216 | 261 | 1122 | 2 | 53.9 | 65.1 |
| C.tropicalis | XP_002551078.1 aureobasidin A resistance protein Candida tr-opicalis MYA-3404 | 397 | 8.4e-152 | 397 | 435.647 | 215 | 282 | 1119 | 5 | 53.6 | 70.3 |
| C.auris | XP_028888388.2 inositol phosp-horylceramide synthase Candida auris | 416 | 7.9e-129 | 416 | 376.326 | 208 | 271 | 965 | 23 | 51.9 | 67.6 |
| C.albicans | XP_715708.1 inositol phosphor-ylceramide synthase Candida al-bicans SC5314 | 399 | 8.4e-151 | 399 | 433.335 | 219 | 279 | 1113 | 11 | 54.6 | 69.6 |
| A.fumigatus | XP_754623.1 aureobasidin resi-stance protein Aur1 Aspergillu-s fumigatus Af293 | 329 | 1.3e-96 | 329 | 294.278 | 148 | 207 | 752 | 2 | 36.9 | 51.6 |

Table S9: Pairwise alignment info from yeast Aur1 (DEG20010598), cf. Figure S39.

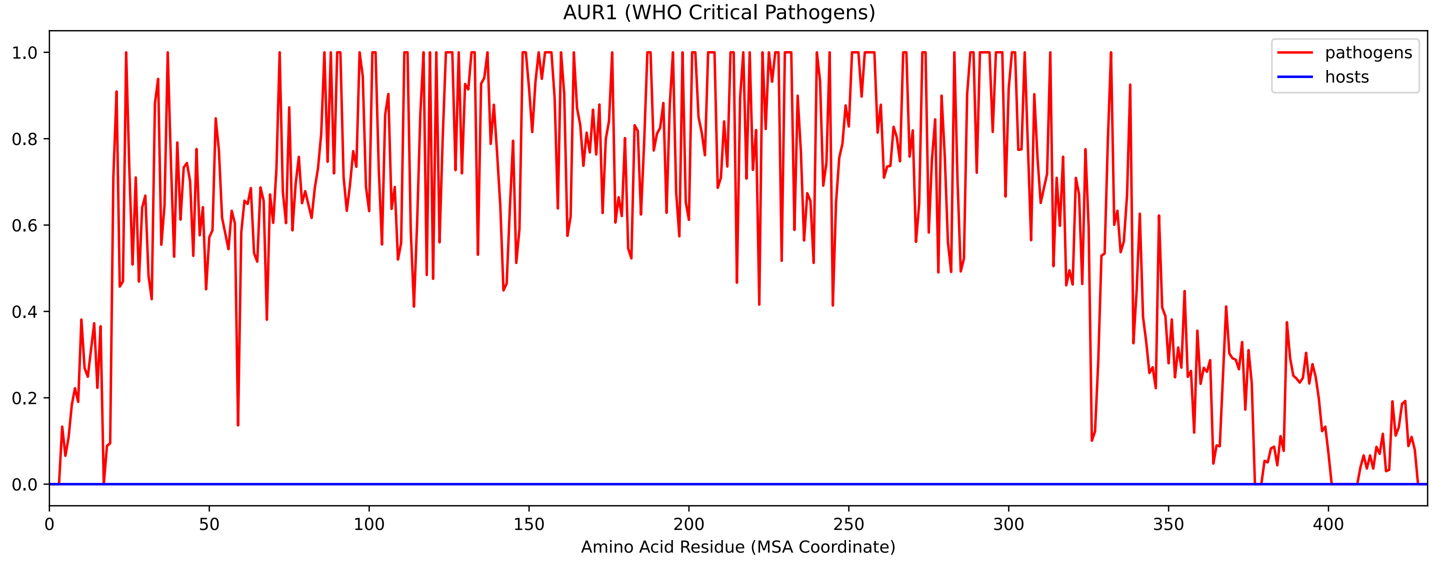


Figure S41: Sneath Similarity of Aur1 for WHO Critical Pathogens, cf. Figure S39

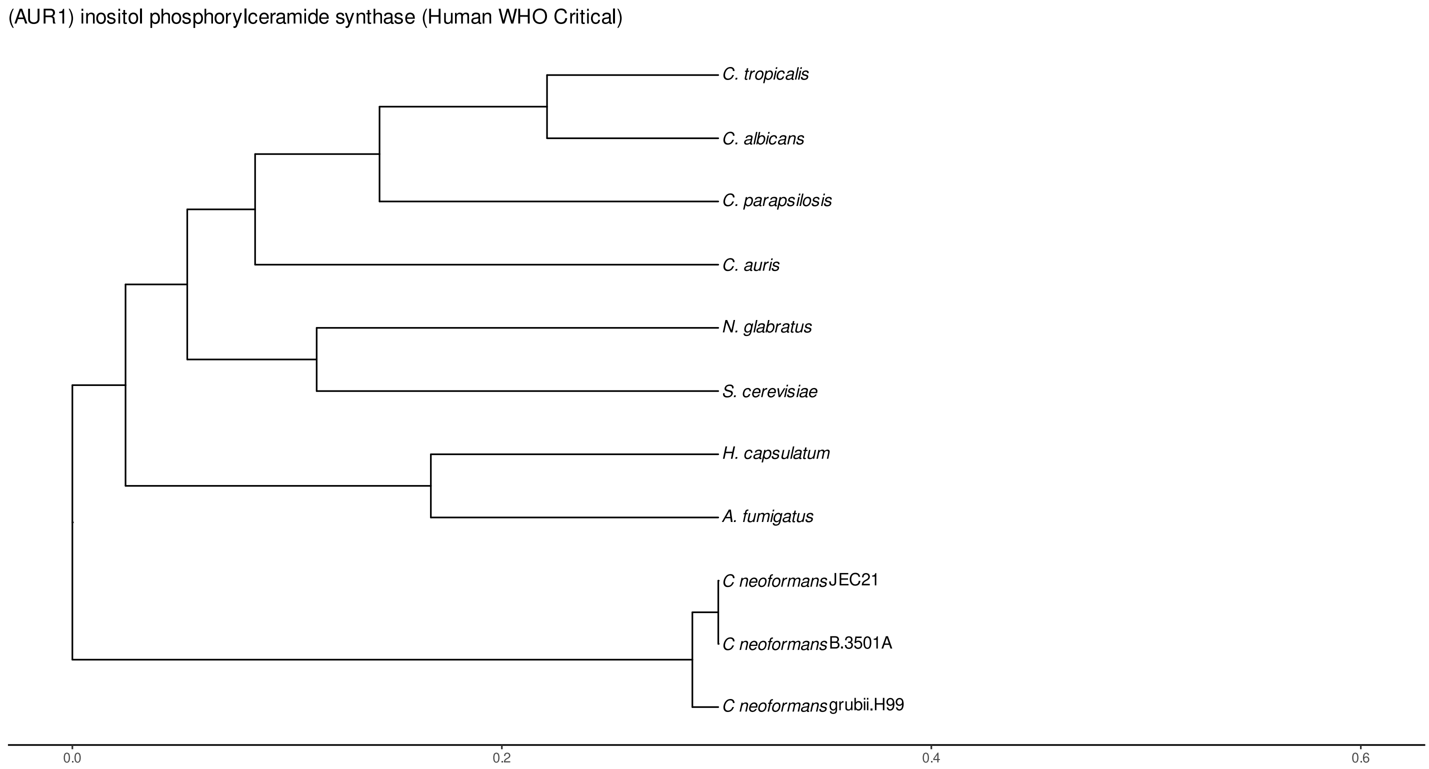


Figure S42: Phylogeny of hits for Aur1 made using clustal omega and visualized using ggtree, cf. Figures S39 and S40.

##### S4.2.2 Top 10 Agricultural Fungal Pathogens

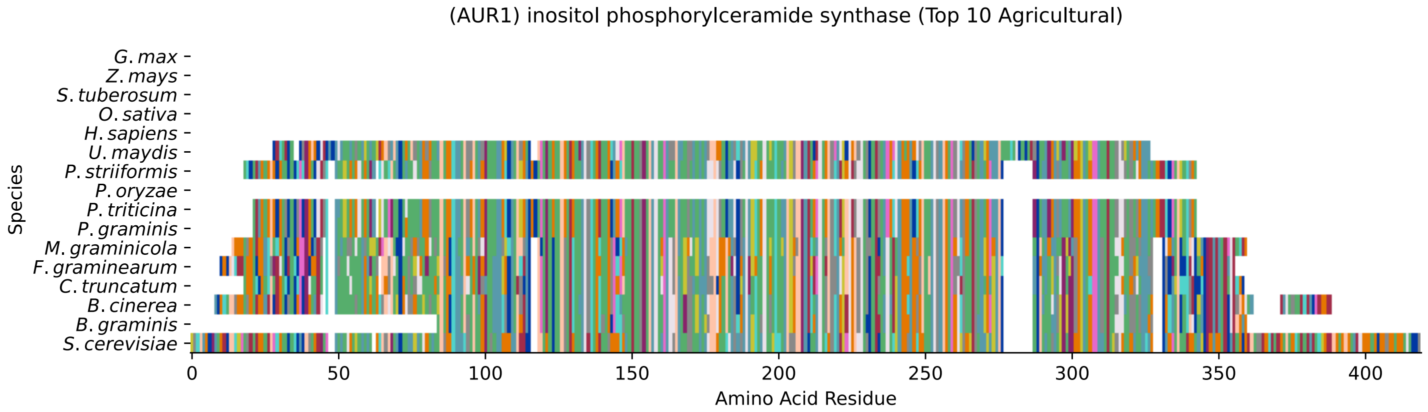


Figure S43: Multiple sequence alignment of yeast Aur1 (Top 10 Agricultural Fungal Pathogens). Cf. Figure S44 for alignment quality, and Figure S45 for Sneath similarity. Cf. Table S10 for protein names, and pairwise alignment metrics with yeast Aur1.

#### Aur1 MSA Quality

| | B. g. | B. c. | C. t. | F. g. | M. g. | P. g. | P. t. | P. o. | P. s. | U. m. | S. c. |
|---|---|---|---|---|---|---|---|---|---|---|---|
| *B. graminis* | 532.332 | 456.062 | 343.969 | 368.622 | 402.134 | 275.404 | 273.863 | 0 | 273.478 | 251.906 | 298.13 |
| *B. cinerea* | 457.603 | 717.998 | 397.512 | 409.838 | 472.241 | 308.531 | 307.375 | 0 | 306.22 | 278.485 | 322.398 |
| *C. truncatum* | 352.829 | 404.831 | 656.751 | 444.121 | 384.415 | 288.5 | 285.419 | 0 | 288.886 | 247.284 | 303.908 |
| *F. graminearum* | 368.622 | 402.134 | 422.935 | 671.389 | 379.407 | 280.796 | 280.026 | 0 | 275.404 | 244.588 | 285.804 |
| *M. graminicola* | 402.134 | 461.84 | 372.089 | 379.407 | 665.611 | 277.715 | 277.33 | 0 | 280.411 | 251.521 | 314.309 |
| *P. graminis* | 275.404 | 301.597 | 277.715 | 280.796 | 277.715 | 620.157 | 613.224 | 0 | 578.941 | 303.138 | 269.626 |
| *P. triticina* | 273.863 | 300.056 | 274.248 | 280.026 | 277.33 | 613.224 | 620.157 | 0 | 578.17 | 300.827 | 267.314 |
| *P. oryzae* | 0 | 0 | 0 | 0 | 0 | 0 | 0 | 0 | 0 | 0 | 0 |
| *P. striiformis* | 273.478 | 298.516 | 275.404 | 275.404 | 280.411 | 578.941 | 578.17 | 0 | 627.476 | 301.212 | 273.478 |
| *U. maydis* | 270.396 | 291.967 | 257.684 | 264.618 | 271.937 | 326.635 | 323.939 | 0 | 324.324 | 593.964 | 245.743 |
| *S. cerevisiae* | 298.13 | 319.316 | 291.582 | 285.804 | 314.309 | 269.626 | 267.314 | 0 | 273.478 | 220.705 | 817.379 |

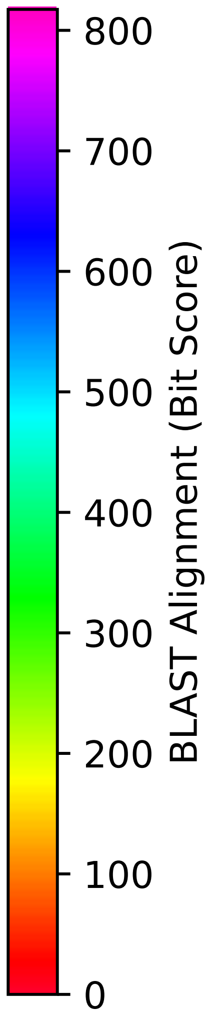


Figure S44: Multiple sequence alignment quality of Aur1 (Top 10 Agricultural Fungal Pathogens). Cf. Figures S43 and S46.

| Species | Hit Protein | Hit Length (a.a.) | evalue | align_len | bit_score | identity | positive | score | gaps | % identity | % positive |
|---|---|---|---|---|---|---|---|---|---|---|---|
| G.max | - | - | - | - | - | - | - | - | - | - | - |
| Z.mays | - | - | - | - | - | - | - | - | - | - | - |
| S.tuberosum | - | - | - | - | - | - | - | - | - | - | - |
| O.sativa | - | - | - | - | - | - | - | - | - | - | - |
| H.sapiens | - | - | - | - | - | - | - | - | - | - | - |
| U.maydis | XP_011387609.1 uncharacterized protein UMAG 01613 Ustilago maydis 521 | 295 | 2.2e-69 | 295 | 225.713 | 133 | 182 | 574 | 11 | 33.2 | 45.4 |
| P.striiformis | XP_047811613.1 hypothetical protein Pst134EA 002784 Puccinia striiformis f. sp. tritici | 312 | 1.6e-89 | 312 | 275.404 | 144 | 196 | 703 | 5 | 35.9 | 48.9 |
| P.oryzae | - | - | - | - | - | - | - | - | - | - | - |
| P.triticina | XP_053018037.1 uncharacterized protein PtA15 2A799 Puccinia triticina | 309 | 1.1e-87 | 309 | 271.552 | 143 | 193 | 693 | 5 | 35.7 | 48.1 |
| P.graminis | XP_003333402.2 hypothetical protein PGTG 15186 Puccinia graminis f. sp. tritici CRL 75-36-700-3 | 309 | 4.6e-88 | 309 | 273.092 | 144 | 194 | 697 | 5 | 35.9 | 48.4 |
| M.graminicola | ZTRI 6.447.mRNA-p1 — transcript=ZTRI 6.447.mRNA — gene=ZTRI 6.447 — organism=Zymoseptoria tritici IPO323 — gene product=similar to aureobasidin resistance protein aur1 — transcript product=similar to aureobasidin resistance protein aur1 — location=Ztri chr 6:1662238-1663581(+) — protein length=447 — sequence SO=chromosome — SO=protein coding gene — is pseudo=false | 330 | 2.1e-105 | 330 | 317.39 | 163 | 219 | 812 | 4 | 40.6 | 54.6 |
| F.graminearum | XP_011315988.1 hypothetical protein FGSG 00338 Fusarium graminearum PH-1 | 331 | 7.1e-94 | 331 | 288.115 | 149 | 211 | 736 | 7 | 37.2 | 52.6 |
| C.truncatum | XP_036579559.1 PAP2 superfamily protein Colletotrichum truncatum | 323 | 3.8e-96 | 323 | 294.278 | 162 | 210 | 752 | 1 | 40.4 | 52.4 |
| B.cinerea | XP_001546315.2 Bcaur1 Botrytis cinerea B05.10 | 364 | 2.2e-106 | 364 | 320.087 | 166 | 223 | 819 | 11 | 41.4 | 55.6 |
| B.graminis | VDB89011.1 — transcript=BGT96224V316 LOCUS4784 t1 — gene=BGT96224V316 LOCUS4784 — organism=Blumeria graminis f. sp. tritici 96224 — gene product=unspecified product — transcript product=unspecified product — location=LR026990:2935260-2936323(-) — protein length=337 — sequence SO=chromosome — SO=protein coding gene — is pseudo=false | 261 | 3.1e-100 | 261 | 299.671 | 143 | 182 | 766 | 0 | 35.7 | 45.4 |

Table S10: Pairwise alignment info from yeast Aur1 (DEG20010598), cf. Figure S43.

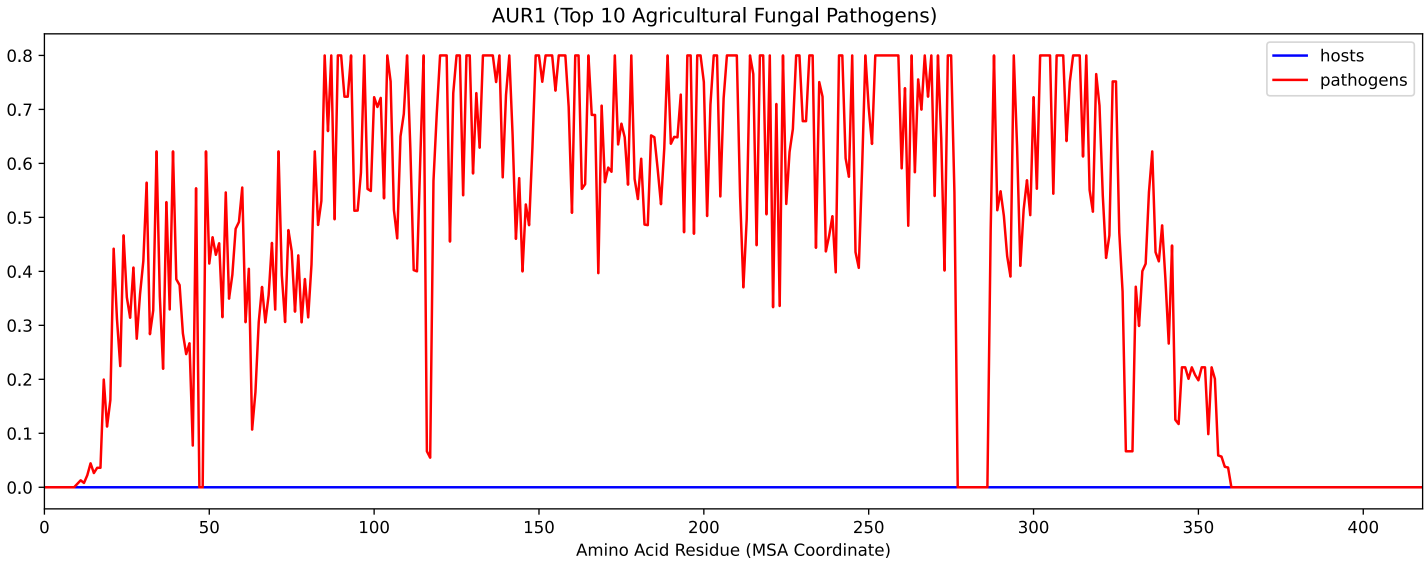


Figure S45: Sneath Similarity of Aur1 for Top 10 Agricultural Fungal Pathogens, cf. Figure S43

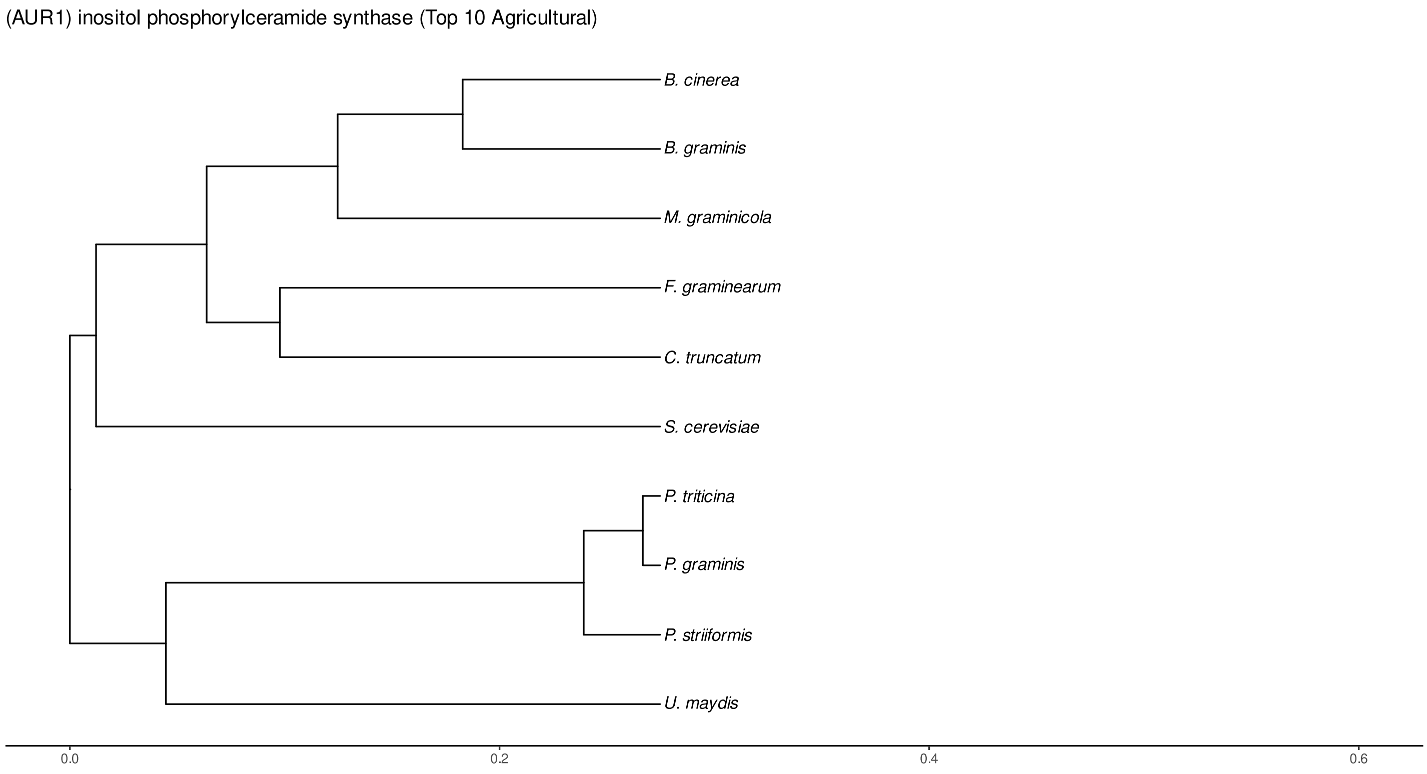


Figure S46: Phylogeny of hits for Aur1 made using clustal omega and visualized using ggtree, cf. Figures S43 and S44.

### S4.2.3 NR

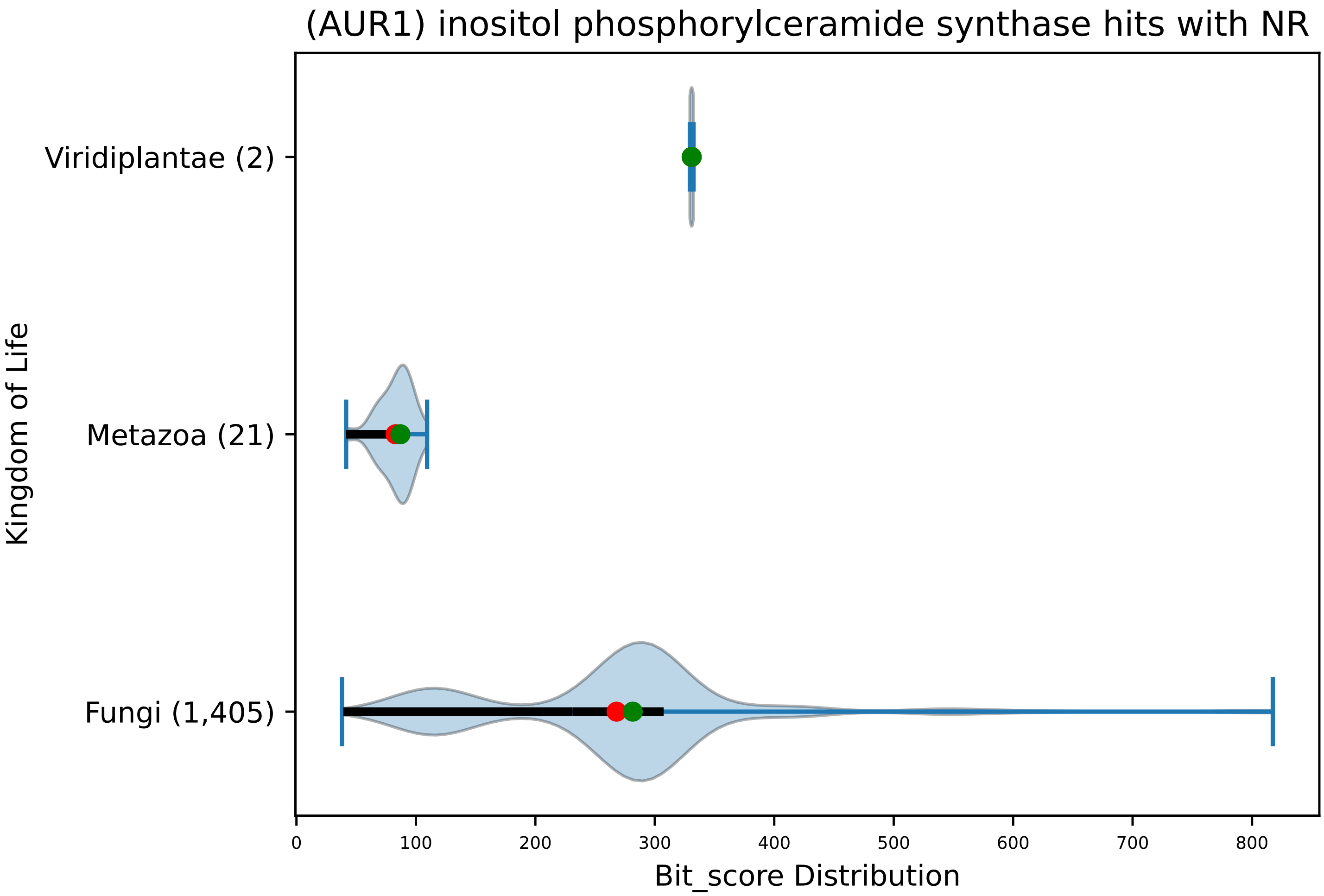


Figure S47: Non-redundant (NR) protein hits for DEG20010598/Aur1, with expectation value of no more than 0.1. Green points are medians, and red points are arithmetic means.

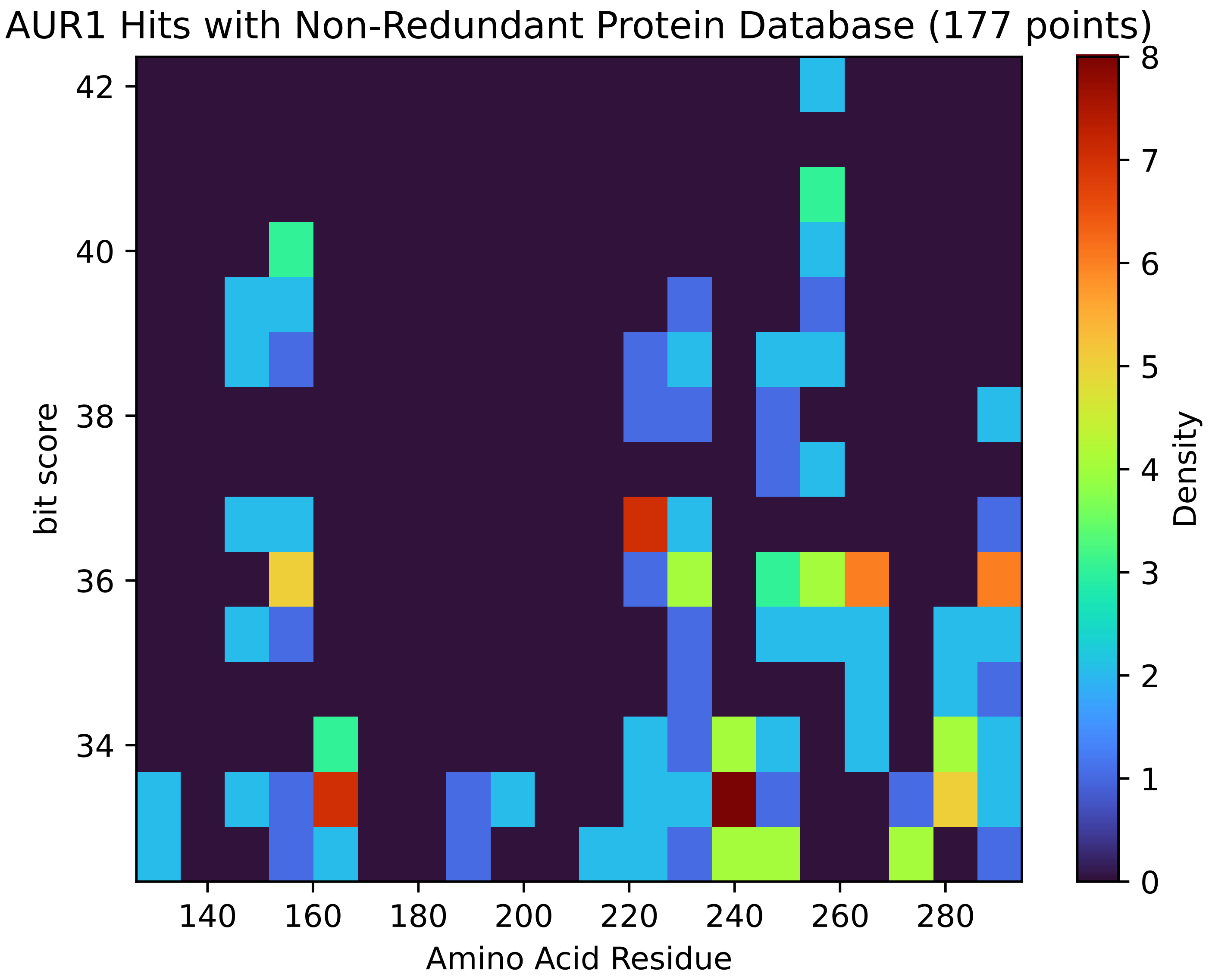


Figure S48: Non-redundant (NR) protein hits for Aur1 in the kingdom Viridiplantae.

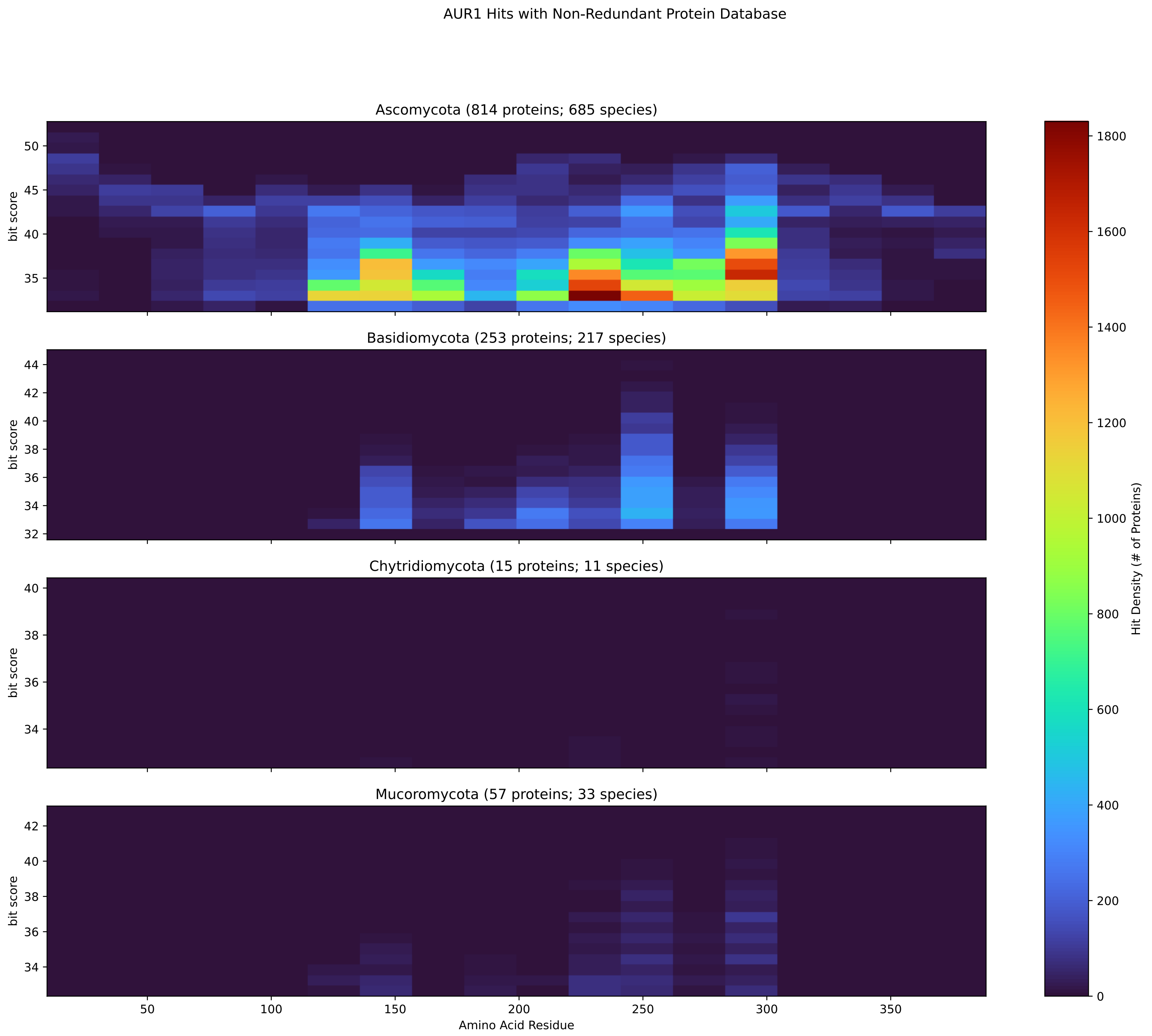


Figure S49: Non-redundant (NR) protein hits for Aur1 in the kingdom Fungi.

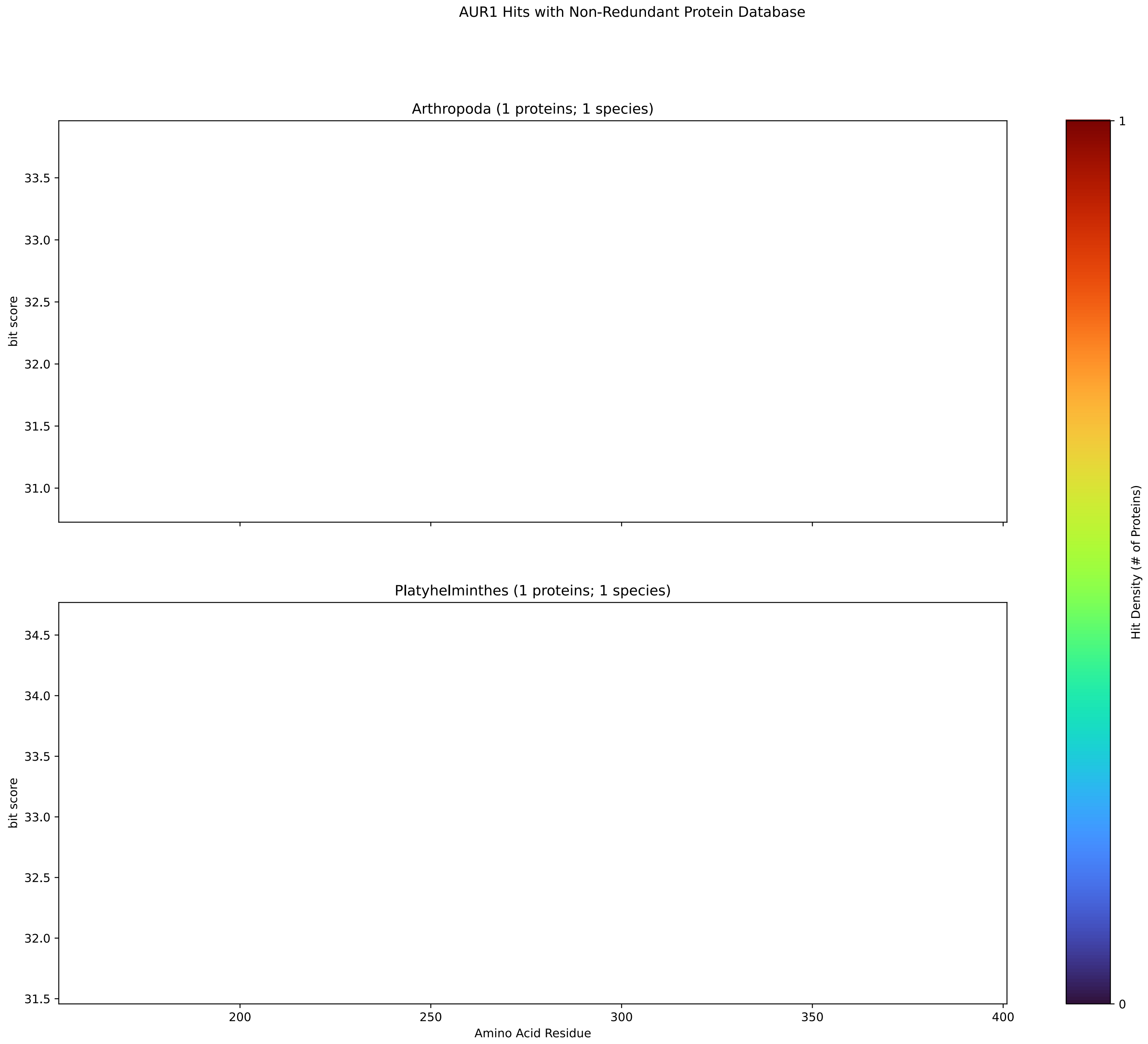


Figure S50: Non-redundant (NR) protein hits for Aur1 in the kingdom Metazoa.

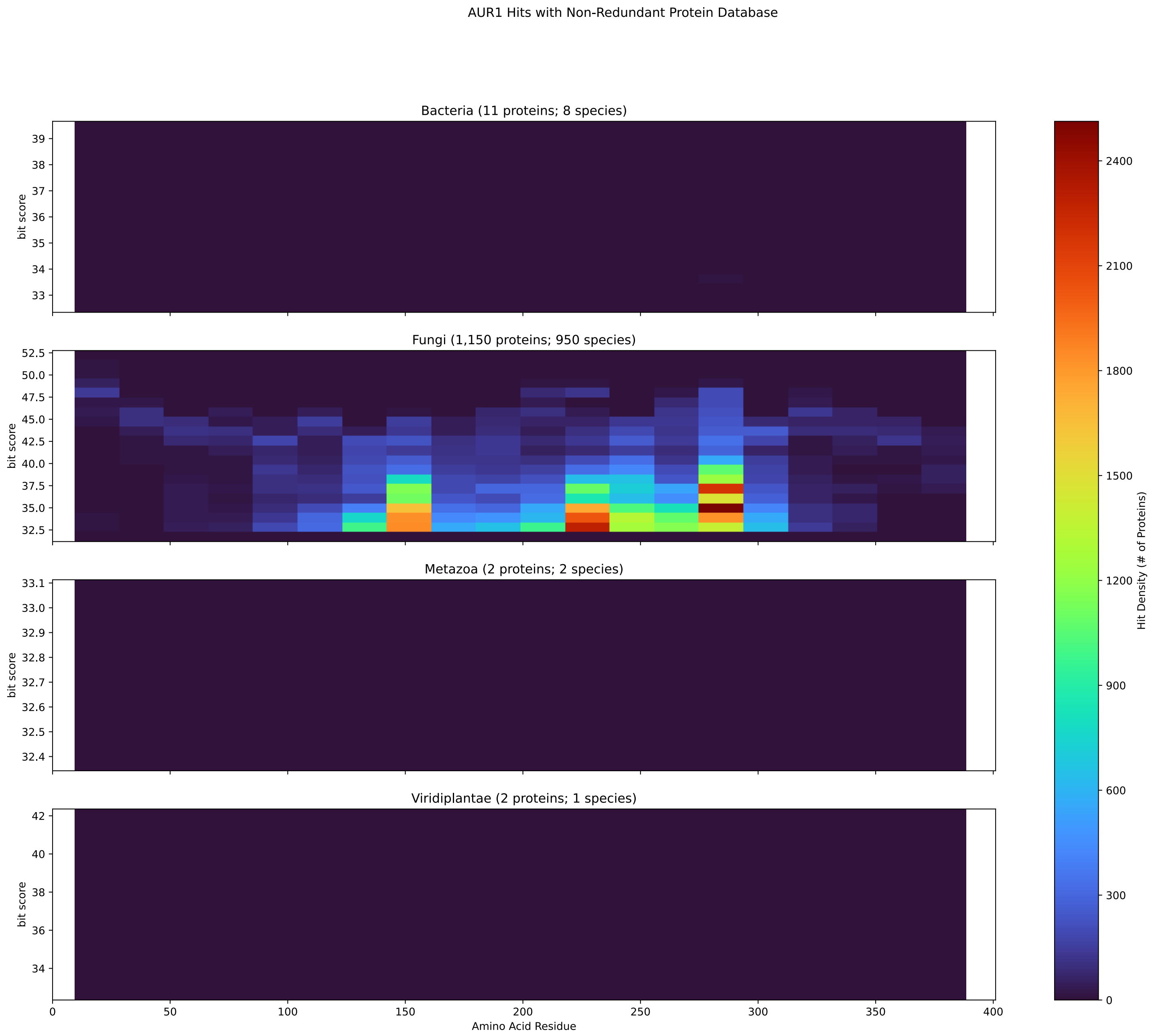


Figure S51: Non-redundant (NR) protein hits for DEG20010598/Aur1 at 20 amino acid length queries.

#### S4.3 Chs2

##### S4.3.1 WHO Critical Pathogens

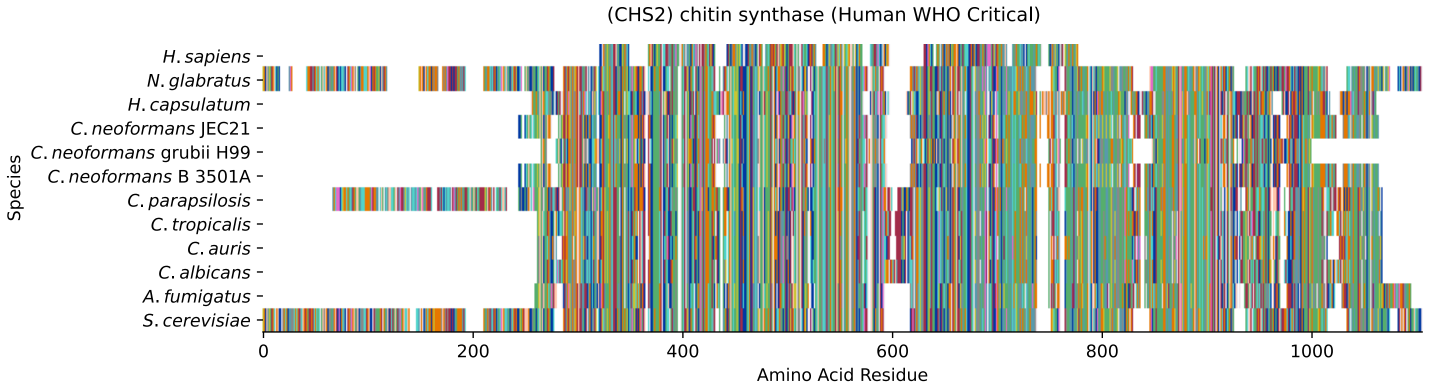


Figure S52: Multiple sequence alignment of yeast Chs2 (WHO Critical Pathogens). Cf. Figure S53 for alignment quality, and Figure S54 for Sneath similarity. Cf. Table S11 for protein names, and pairwise alignment metrics with yeast Chs2.

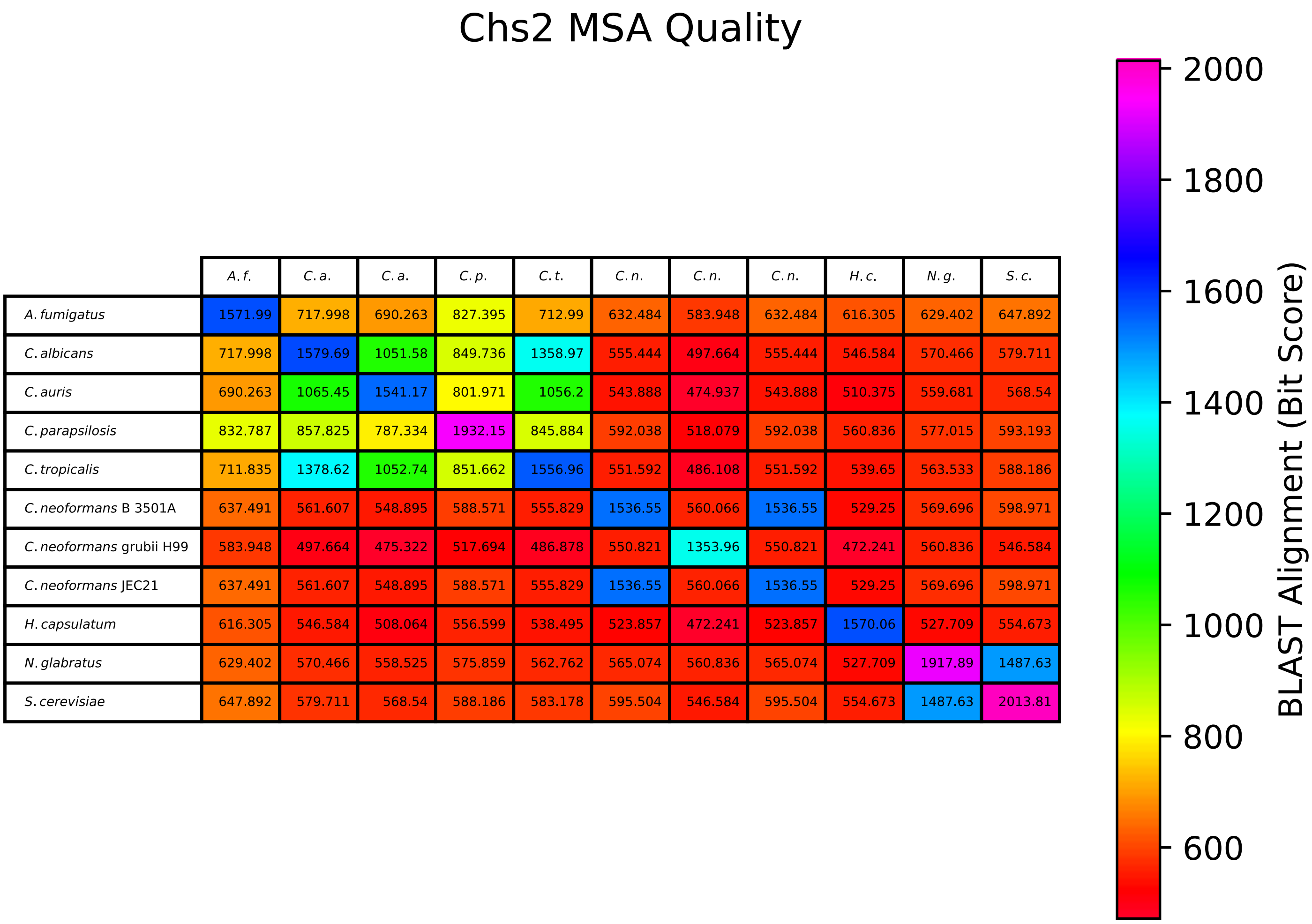


| | *A. f.* | *C. a.* | *C. a.* | *C. p.* | *C. t.* | *C. n.* | *C. n.* | *C. n.* | *H. c.* | *N. g.* | *S. c.* |
|---|---|---|---|---|---|---|---|---|---|---|---|
| *A. fumigatus* | 1571.99 | 717.998 | 690.263 | 827.395 | 712.99 | 632.484 | 583.948 | 632.484 | 616.305 | 629.402 | 647.892 |
| *C. albicans* | 717.998 | 1579.69 | 1051.58 | 849.736 | 1358.97 | 555.444 | 497.664 | 555.444 | 546.584 | 570.466 | 579.711 |
| *C. auris* | 690.263 | 1065.45 | 1541.17 | 801.971 | 1056.2 | 543.888 | 474.937 | 543.888 | 510.375 | 559.681 | 568.54 |
| *C. parapsilosis* | 832.787 | 857.825 | 787.334 | 1932.15 | 845.884 | 592.038 | 518.079 | 592.038 | 560.836 | 577.015 | 593.193 |
| *C. tropicalis* | 711.835 | 1378.62 | 1052.74 | 851.662 | 1556.96 | 551.592 | 486.108 | 551.592 | 539.65 | 563.533 | 588.186 |
| *C. neoformans* B 3501A | 637.491 | 561.607 | 548.895 | 588.571 | 555.829 | 1536.55 | 560.066 | 1536.55 | 529.25 | 569.696 | 598.971 |
| *C. neoformans* grubii H99 | 583.948 | 497.664 | 475.322 | 517.694 | 486.878 | 550.821 | 1353.96 | 550.821 | 472.241 | 560.836 | 546.584 |
| *C. neoformans* JEC21 | 637.491 | 561.607 | 548.895 | 588.571 | 555.829 | 1536.55 | 560.066 | 1536.55 | 529.25 | 569.696 | 598.971 |
| *H. capsulatum* | 616.305 | 546.584 | 508.064 | 556.599 | 538.495 | 523.857 | 472.241 | 523.857 | 1570.06 | 527.709 | 554.673 |
| *N. glabratus* | 629.402 | 570.466 | 558.525 | 575.859 | 562.762 | 565.074 | 560.836 | 565.074 | 527.709 | 1917.89 | 1487.63 |
| *S. cerevisiae* | 647.892 | 579.711 | 568.54 | 588.186 | 583.178 | 595.504 | 546.584 | 595.504 | 554.673 | 1487.63 | 2013.81 |

Figure S53: Multiple sequence alignment quality of Chs2 (WHO Critical Pathogens). Cf. Figures S52 and S55.

| Species | Hit Protein | Hit Length (a.a.) | evalue | align_len | bit_score | identity | positive | score | gaps | % identity | % positive |
|---|---|---|---|---|---|---|---|---|---|---|---|
| H.sapiens | NP_001186209.1 hyaluronan synthase 3 isoform a Homo sapiens | 422 | 1.9e-05 | 422 | 51.2174 | 95 | 165 | 121 | 68 | 9.9 | 17.1 |
| N.glabratus | XP_447459.1 uncharacterized protein CAGL0I04818g Nakaseomyces glabratus | 964 | 0 | 964 | 1487.24 | 734 | 806 | 3849 | 53 | 76.2 | 83.7 |
| H.capsulatum | XP_045283732.1 chitin synthase Histoplasma capsulatum G186AR | 761 | 0 | 761 | 557.755 | 310 | 441 | 1436 | 73 | 32.2 | 45.8 |
| C.neoformans.JEC21 | XP_571995.1 chitin synthase 1-, putative Cryptococcus neoformans var. neoformans JEC21 | 744 | 0 | 744 | 598.586 | 310 | 450 | 1542 | 38 | 32.2 | 46.7 |
| C.neoformans.grubii.H99 | XP_012053352.1 chitin synthase Cryptococcus neoformans var. grubii H99 | 655 | 0 | 655 | 551.206 | 294 | 396 | 1419 | 20 | 30.5 | 41.1 |
| C.neoformans.B.3501A | XP_774244.1 hypothetical protein CNBG2250 Cryptococcus neoformans var. neoformans B-3501A | 744 | 0 | 744 | 598.586 | 310 | 450 | 1542 | 38 | 32.2 | 46.7 |
| C.parapsilosis | XP_036668304.1 uncharacterized protein CPAR2 701490 Candida parapsilosis | 958 | 0 | 958 | 589.726 | 357 | 501 | 1519 | 122 | 37.1 | 52.0 |
| C.tropicalis | XP_002551423.1 chitin synthase 2 Candida tropicalis MYA-340-4 | 759 | 0 | 759 | 588.186 | 315 | 452 | 1515 | 68 | 32.7 | 46.9 |
| C.auris | XP_028890166.1 hypothetical protein Candida auris | 747 | 0 | 747 | 572.392 | 309 | 445 | 1474 | 57 | 32.1 | 46.2 |
| C.albicans | XP_717760.2 Chs8p Candida albicans SC5314 | 767 | 0 | 767 | 583.948 | 319 | 456 | 1504 | 77 | 33.1 | 47.4 |
| A.fumigatus | XP_749322.1 chitin synthase A Aspergillus fumigatus Af293 | 761 | 0 | 761 | 649.818 | 343 | 472 | 1675 | 51 | 35.6 | 49.0 |

Table S11: Pairwise alignment info from yeast Chs2 (DEG20010039), cf. Figure S52.

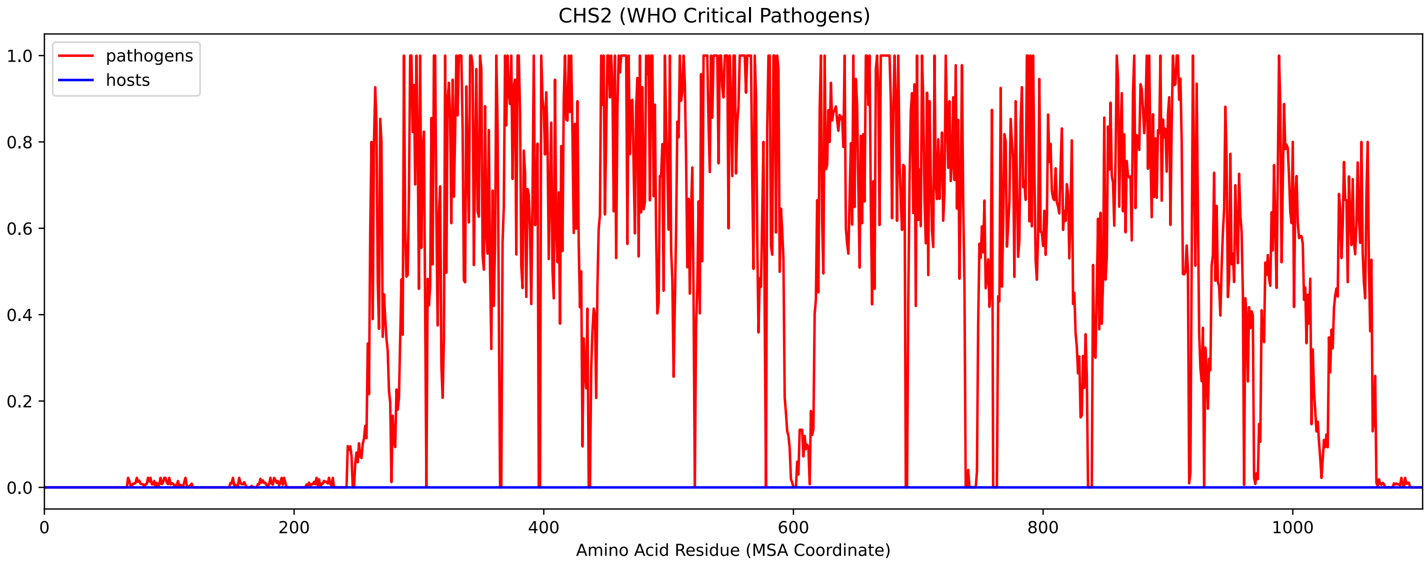


Figure S54: Sneath Similarity of Chs2 for WHO Critical Pathogens, cf. Figure S52

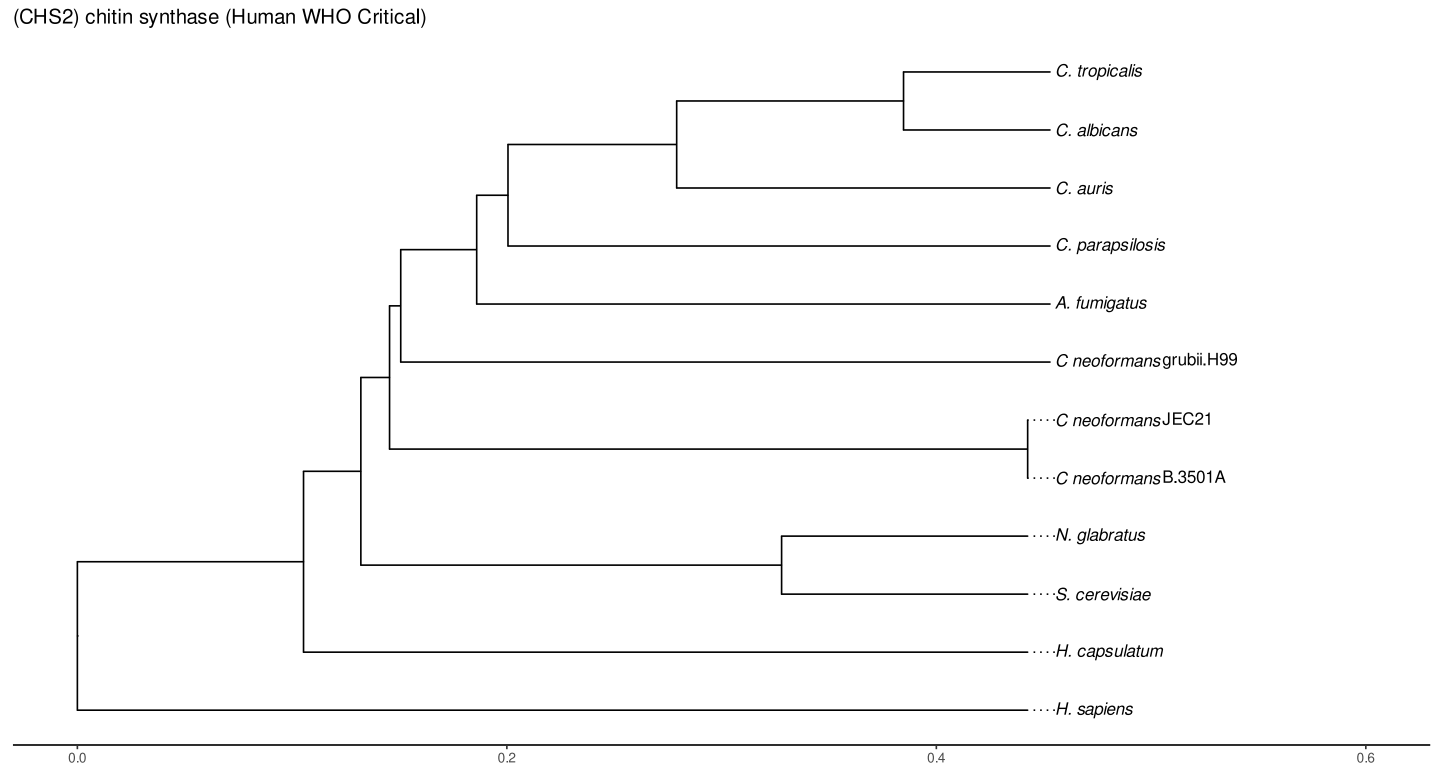


Figure S55: Phylogeny of hits for Chs2 made using clustal omega and visualized using ggtree, cf. Figures S52 and S53.

##### S4.3.2 Top 10 Agricultural Fungal Pathogens

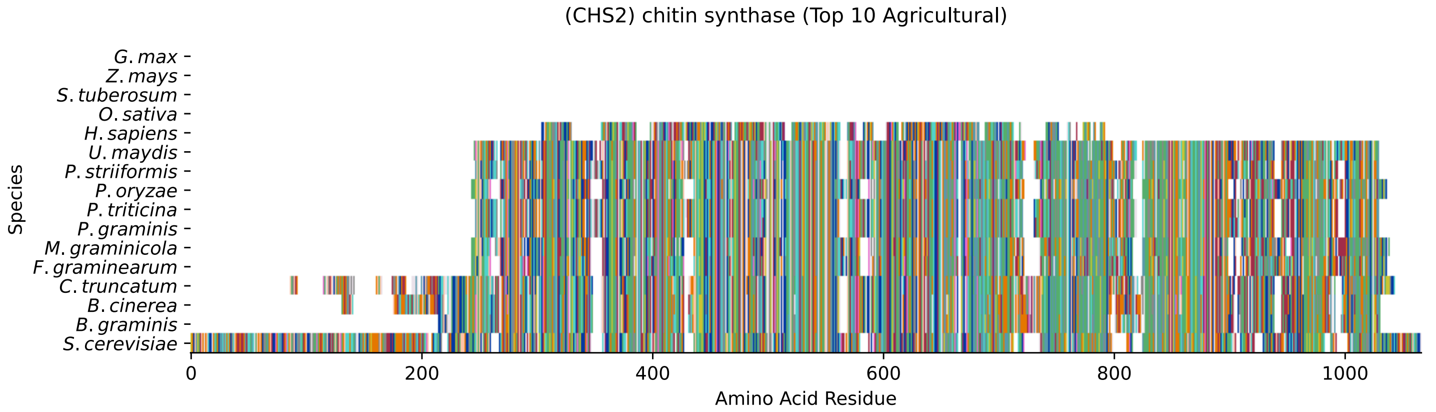


Figure S56: Multiple sequence alignment of yeast Chs2 (Top 10 Agricultural Fungal Pathogens). Cf. Figure S57 for alignment quality, and Figure S58 for Sneath similarity. Cf. Table S12 for protein names, and pairwise alignment metrics with yeast Chs2.

#### Chs2 MSA Quality

| | *B. g.* | *B. c.* | *C. t.* | *F. g.* | *M. g.* | *P. g.* | *P. t.* | *P. o.* | *P. s.* | *U. m.* | *S. c.* |
|---|---|---|---|---|---|---|---|---|---|---|---|
| *B. graminis* | 1612.43 | 1402.11 | 1264.98 | 602.823 | 608.601 | 869.766 | 871.307 | 598.971 | 873.618 | 941.799 | 558.91 |
| *B. cinerea* | 1425.22 | 1721.44 | 1267.29 | 578.17 | 595.89 | 865.144 | 871.692 | 578.17 | 872.848 | 923.309 | 555.829 |
| *C. truncatum* | 1264.98 | 1250.73 | 1788.08 | 602.823 | 608.216 | 860.136 | 857.825 | 587.03 | 859.751 | 894.034 | 562.762 |
| *F. graminearum* | 602.823 | 576.63 | 602.823 | 1519.6 | 1026.16 | 628.632 | 629.402 | 1237.63 | 618.616 | 625.165 | 628.632 |
| *M. graminicola* | 608.601 | 594.349 | 608.216 | 1026.16 | 1524.22 | 604.364 | 602.438 | 1067.37 | 598.201 | 619.772 | 632.484 |
| *P. graminis* | 869.766 | 858.21 | 860.136 | 628.632 | 604.364 | 1538.47 | 1508.43 | 618.616 | 1479.92 | 1047.73 | 545.428 |
| *P. triticina* | 871.307 | 863.218 | 857.825 | 629.402 | 602.438 | 1508.43 | 1536.93 | 620.157 | 1472.6 | 1053.51 | 544.658 |
| *P. oryzae* | 598.971 | 577.015 | 587.03 | 1237.63 | 1067.37 | 618.616 | 620.157 | 1509.97 | 620.157 | 617.461 | 640.573 |
| *P. striiformis* | 873.618 | 864.374 | 859.751 | 618.616 | 598.201 | 1479.92 | 1472.6 | 620.157 | 1541.17 | 1046.96 | 543.502 |
| *U. maydis* | 941.799 | 914.45 | 894.034 | 625.165 | 619.772 | 1047.73 | 1053.51 | 617.461 | 1046.96 | 1539.24 | 551.977 |
| *S. cerevisiae* | 558.91 | 551.592 | 562.762 | 628.632 | 632.484 | 545.428 | 544.658 | 640.573 | 543.502 | 551.977 | 2013.81 |

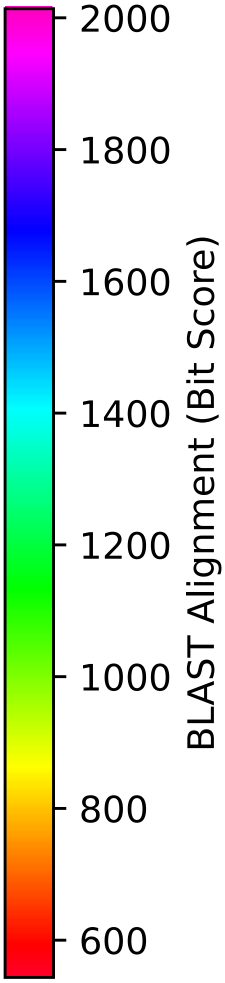


Figure S57: Multiple sequence alignment quality of Chs2 (Top 10 Agricultural Fungal Pathogens). Cf. Figures S56 and S59.

| Species | Hit Protein | Hit Length (a.a.) | evalue | align_len | bit_score | identity | positive | score | gaps | % identity | % positive |
|---|---|---|---|---|---|---|---|---|---|---|---|
| G.max | - | - | - | - | - | - | - | - | - | - | - |
| Z.mays | - | - | - | - | - | - | - | - | - | - | - |
| S.tuberosum | - | - | - | - | - | - | - | - | - | - | - |
| O.sativa | - | - | - | - | - | - | - | - | - | - | - |
| H.sapiens | NP_001186209.1 hyaluronan synthase 3 isoform a Homo sapiens | 422 | 1.9e-05 | 422 | 51.2174 | 95 | 165 | 121 | 68 | 9.9 | 17.1 |
| U.maydis | XP_011391524.1 chitin synthase 1 Ustilago maydis 521 | 761 | 0 | 761 | 555.444 | 307 | 447 | 1430 | 94 | 31.9 | 46.4 |
| P.striiformis | XP_047799288.1 hypothetical protein Pst134EA 026738 Puccinia striiformis f. sp. tritici | 753 | 0 | 753 | 546.584 | 306 | 437 | 1407 | 78 | 31.8 | 45.4 |
| P.oryzae | mRNA M BR32 EuGene 00045151-p1 — transcript=mRNA M BR32 EuGene 00045151 — gene=M BR32 EuGene 00045151 — organism=Pyricularia oryzae BR32 — gene product=unspecified product — transcript product=unspecified product — location=BR32 scaffold00004:267300-270156(-) — protein length=886 — sequence SO=supercontig — SO=protein coding gene — is pseudo=false | 738 | 0 | 738 | 643.269 | 329 | 449 | 1658 | 54 | 34.2 | 46.6 |
| P.triticina | XP_053028600.1 uncharacterized protein PtA15 18A101 Puccinia triticina | 754 | 0 | 754 | 547.74 | 302 | 435 | 1410 | 80 | 31.4 | 45.2 |
| P.graminis | XP_003336234.2 chitin synthase Puccinia graminis f. sp. tritici CRL 75-36-700-3 | 754 | 0 | 754 | 548.895 | 303 | 434 | 1413 | 80 | 31.5 | 45.1 |
| M.graminicola | ZTRI 1.444.mRNA-p1 — transcript=ZTRI 1.444.mRNA — gene=ZTRI 1.444 — organism=Zymoseptoria tritici IPO323 — gene product=similar to chitin synthase 2 — transcript product=similar to chitin synthase 2 — location=Ztri chr 1:1478238-1481305(-) — protein length=946 — sequence SO=chromosome — SO=protein coding gene — is pseudo=false | 744 | 0 | 744 | 634.41 | 331 | 457 | 1635 | 57 | 34.4 | 47.5 |
| F.graminearum | XP_011319287.1 chitin synthase 3 Fusarium graminearum PH-1 | 742 | 0 | 742 | 630.558 | 329 | 452 | 1625 | 58 | 34.2 | 46.9 |
| C.truncatum | XP_036585044.1 chitin synthase Colletotrichum truncatum | 888 | 0 | 888 | 562.762 | 338 | 486 | 1449 | 97 | 35.1 | 50.5 |
| B.cinerea | XP_001557191.1 BcCHSIIIa Botrytis cinerea B05.10 | 835 | 0 | 835 | 553.132 | 326 | 467 | 1424 | 92 | 33.9 | 48.5 |
| B.graminis | VCU40918.1 — transcript=BGT96224V316 LOCUS2169 t1 — gene=BGT96224V316 LOCUS2169 — organism=Blumeria graminis f. sp. tritici 96224 — gene product=unspecified product — transcript product=unspecified product — location=LR026987:2394926-2397807(+) — protein length=900 — sequence SO=chromosome — SO=protein coding gene — is pseudo=false | 792 | 0 | 792 | 559.296 | 314 | 451 | 1440 | 87 | 32.6 | 46.8 |

Table S12: Pairwise alignment info from yeast Chs2 (DEG20010039), cf. Figure S56.

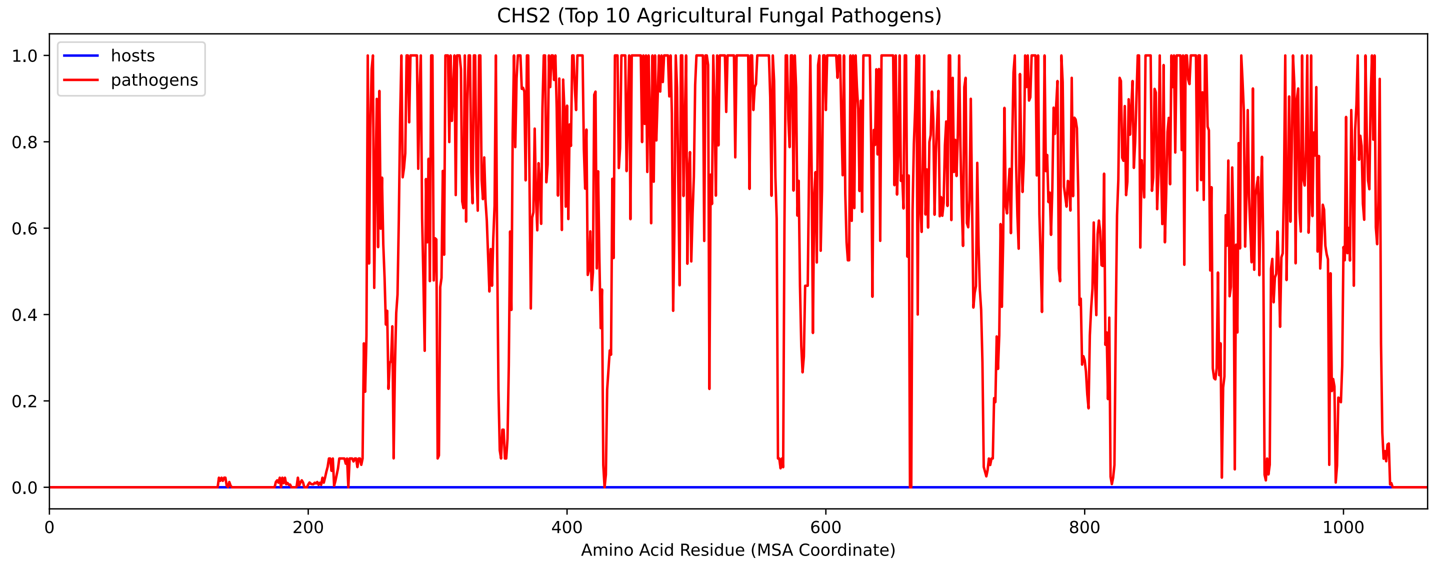


Figure S58: Sneath Similarity of Chs2 for Top 10 Agricultural Fungal Pathogens, cf. Figure S56

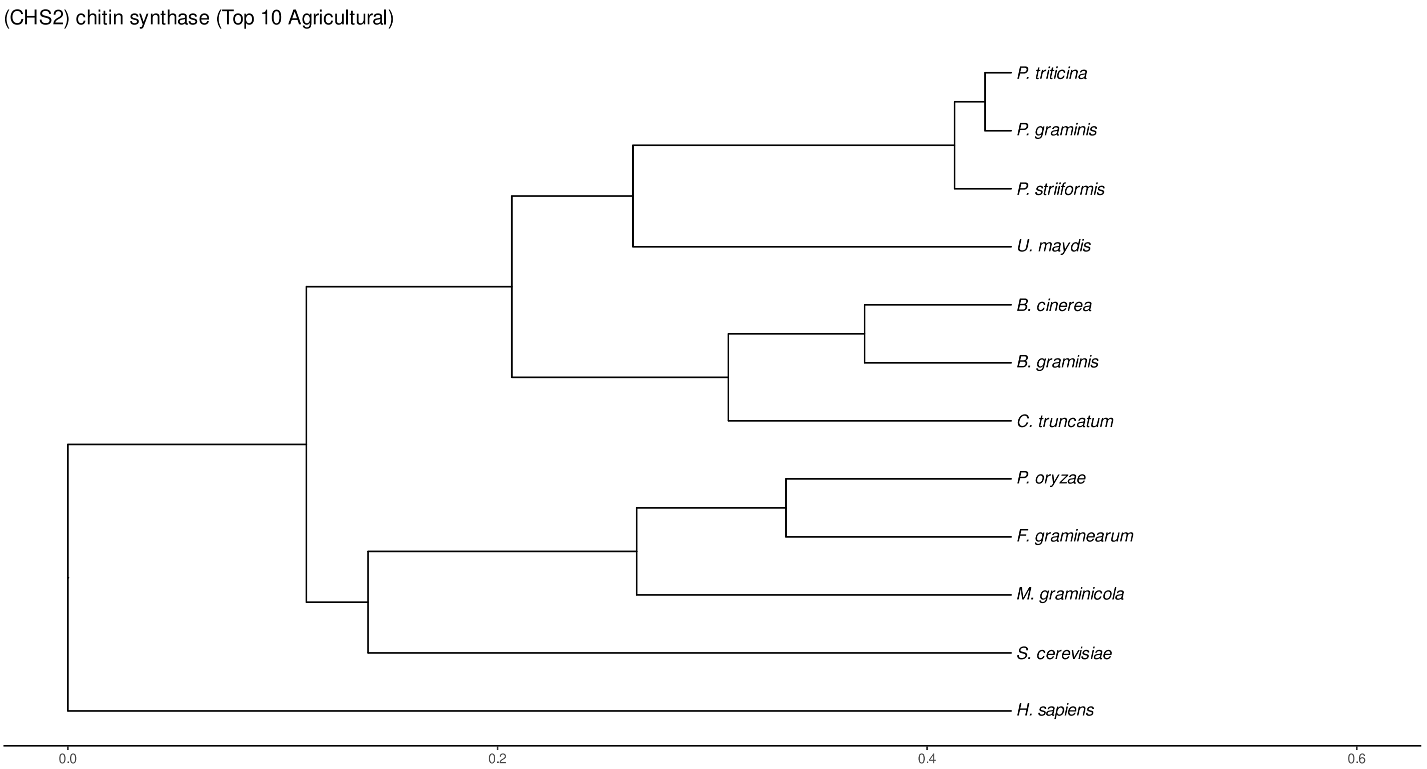


Figure S59: Phylogeny of hits for Chs2 made using clustal omega and visualized using ggtree, cf. Figures S56 and S57.

### S4.3.3 NR

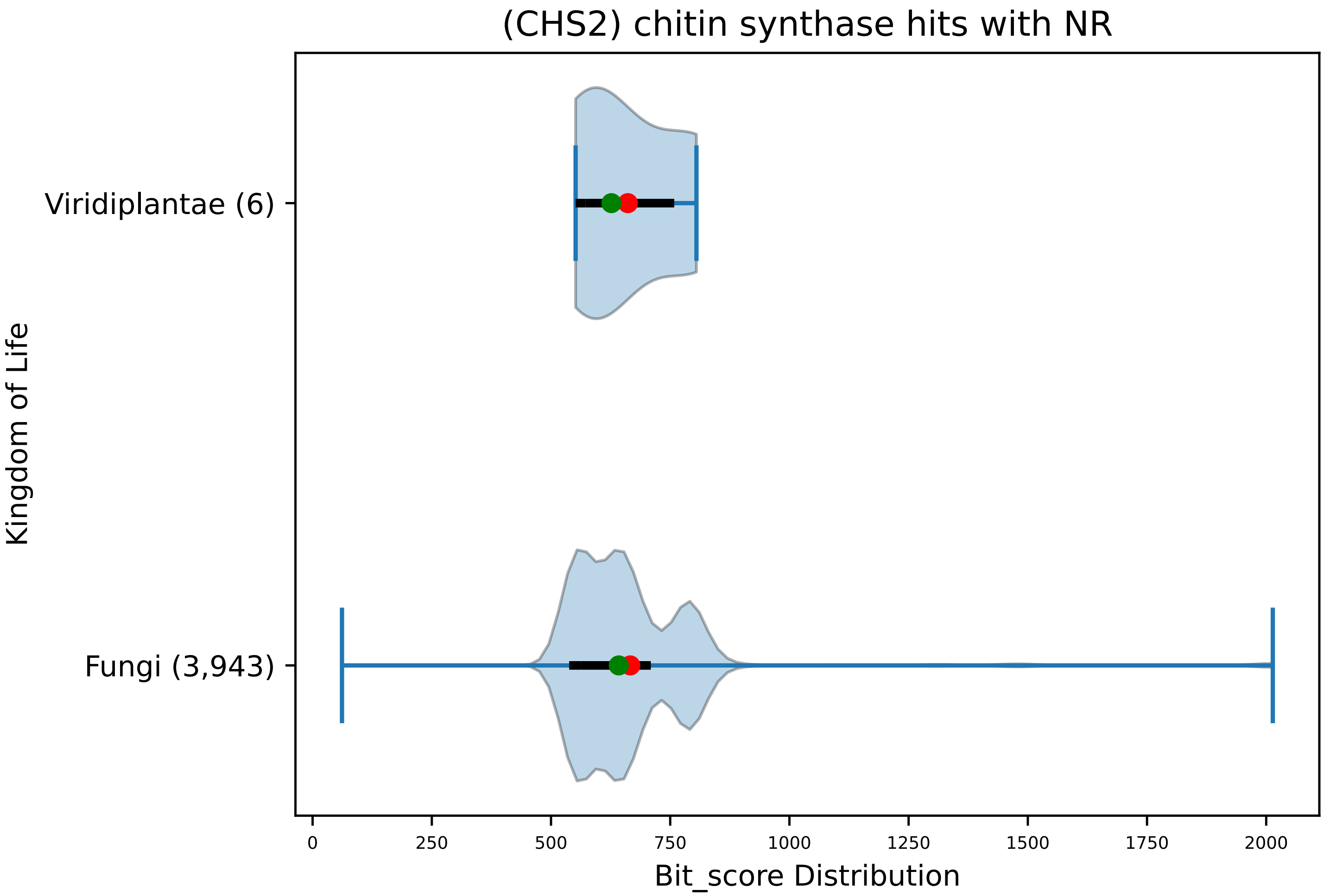


Figure S60: Non-redundant (NR) protein hits for DEG20010039/Chs2, with expectation value of no more than 0.1. Green points are medians, and red points are arithmetic means.

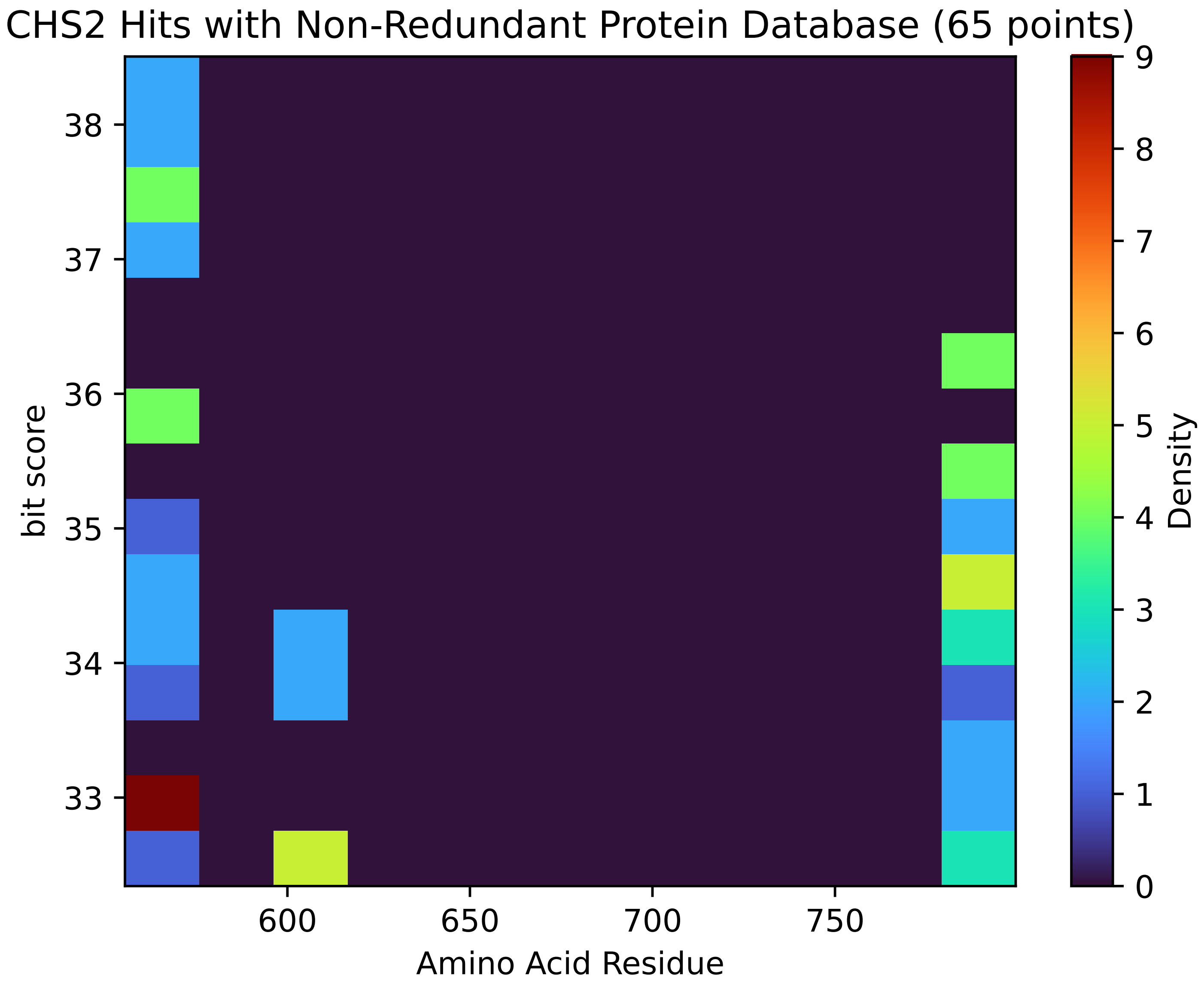


Figure S61: Non-redundant (NR) protein hits for Chs2 in the kingdom Viridiplantae.

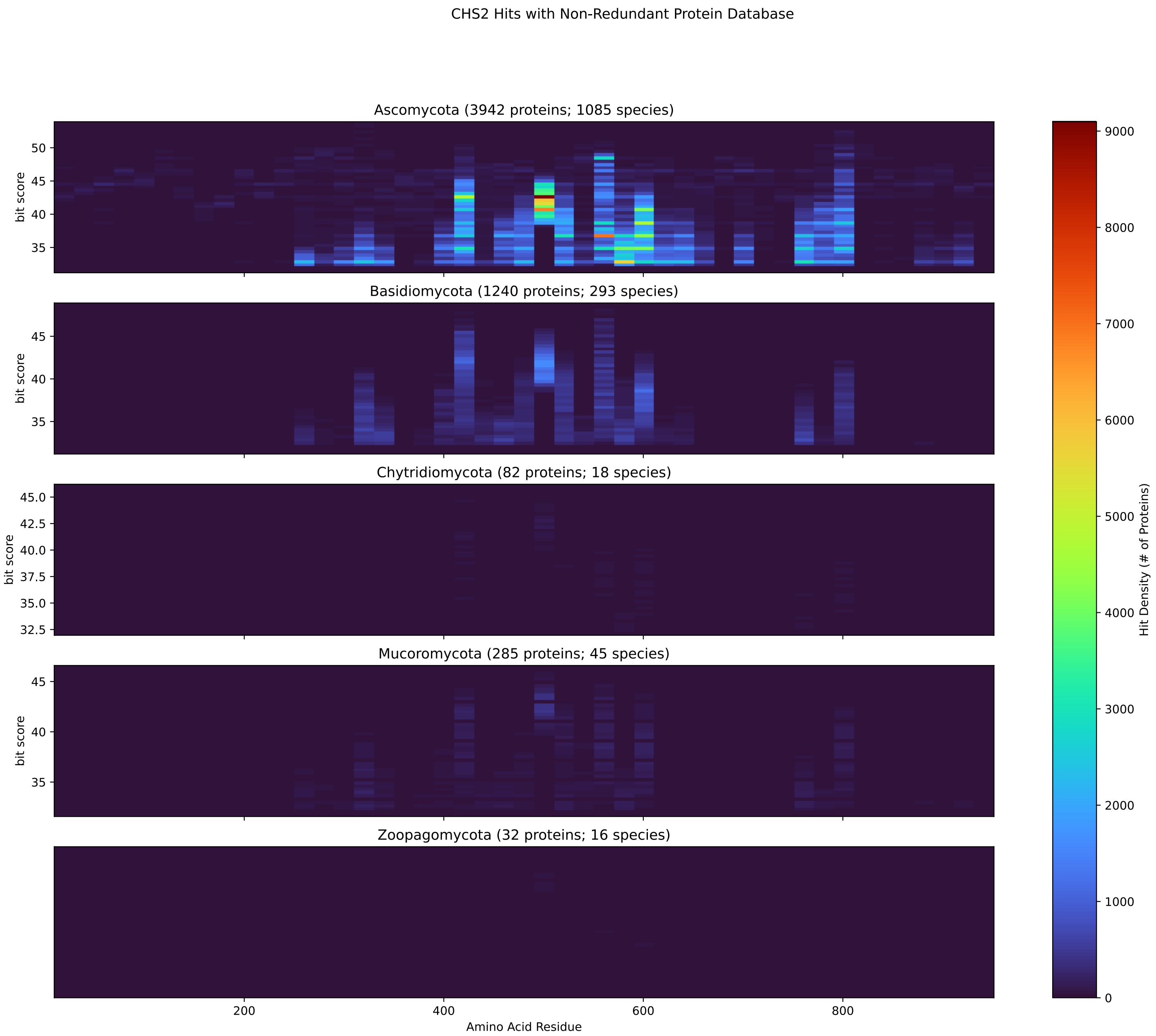


Figure S62: Non-redundant (NR) protein hits for Chs2 in the kingdom Fungi.

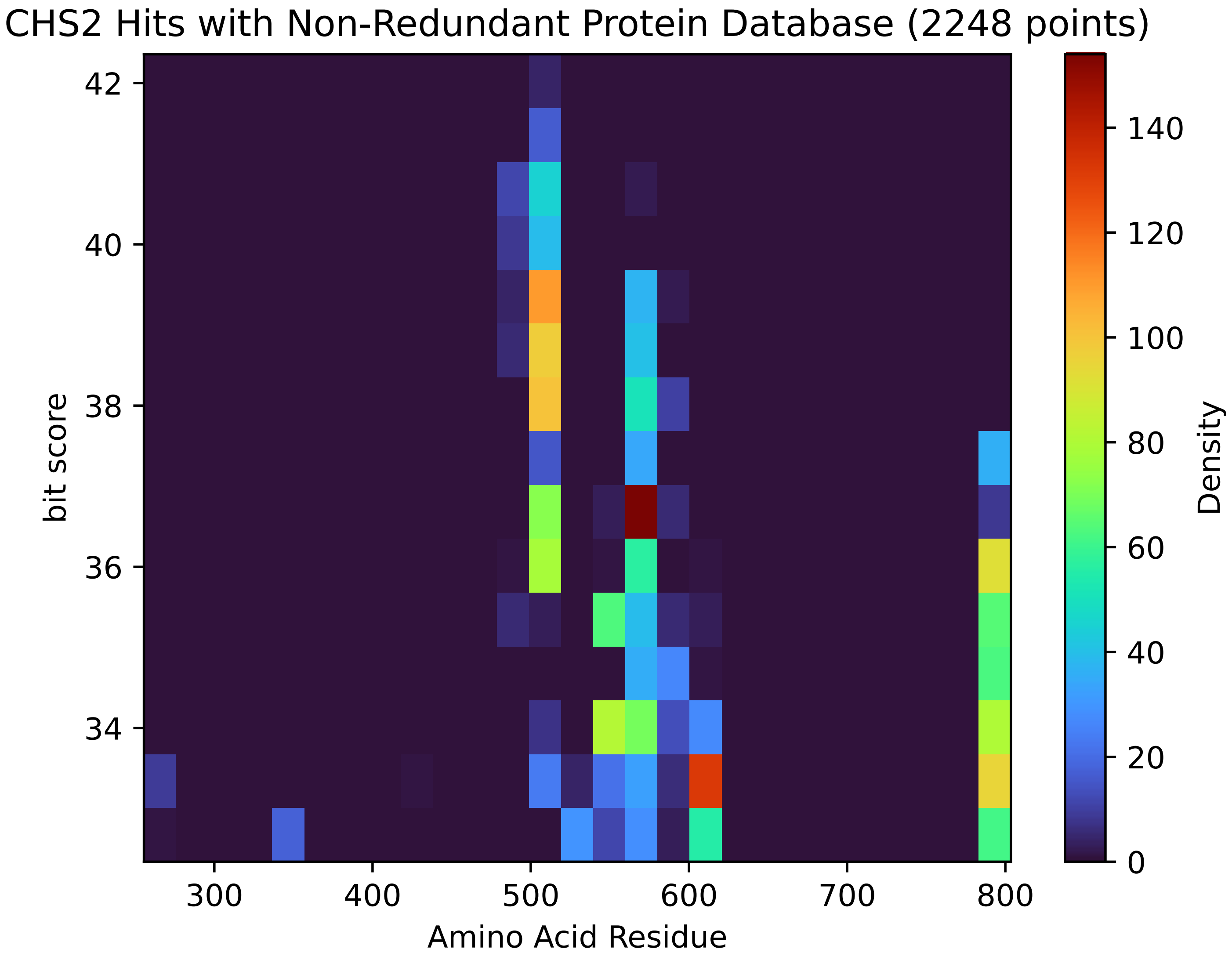


Figure S63: Non-redundant (NR) protein hits for Chs2 in the kingdom SAR.

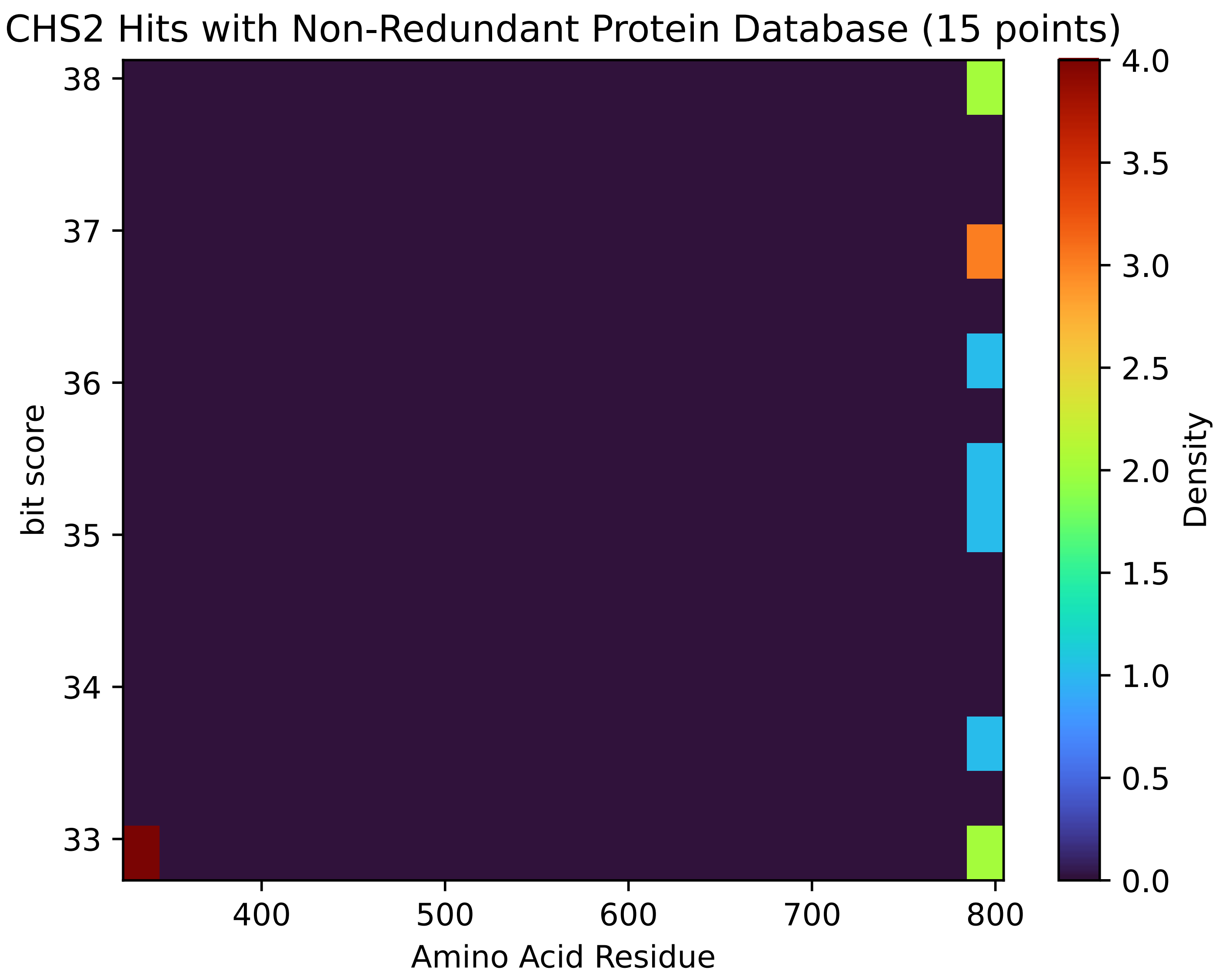


Figure S64: Non-redundant (NR) protein hits for Chs2 in the kingdom Metazoa.

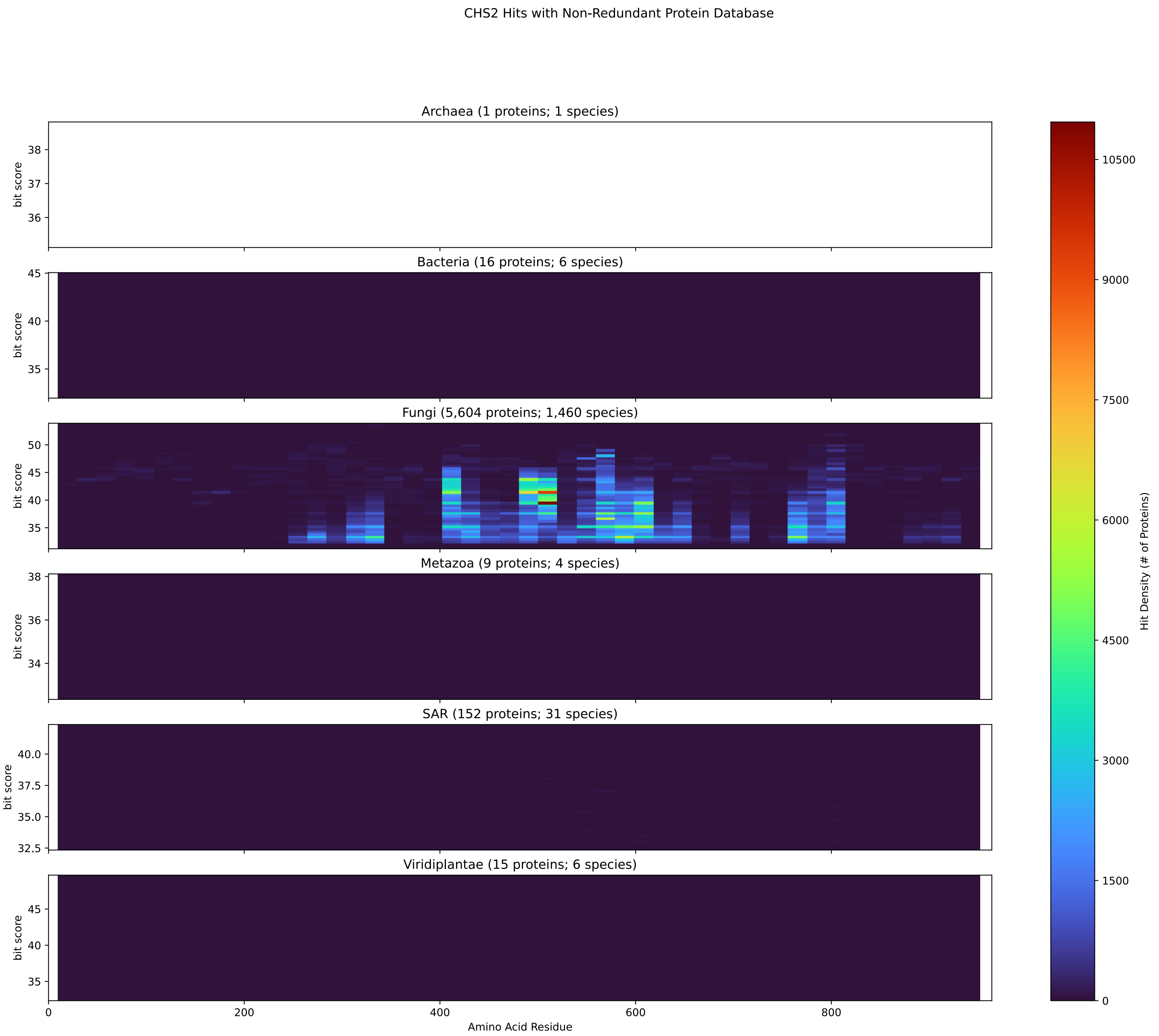


Figure S65: Non-redundant (NR) protein hits for DEG20010039/Chs2 at 20 amino acid length queries.

#### S4.4 Erg8

##### S4.4.1 WHO Critical Pathogens

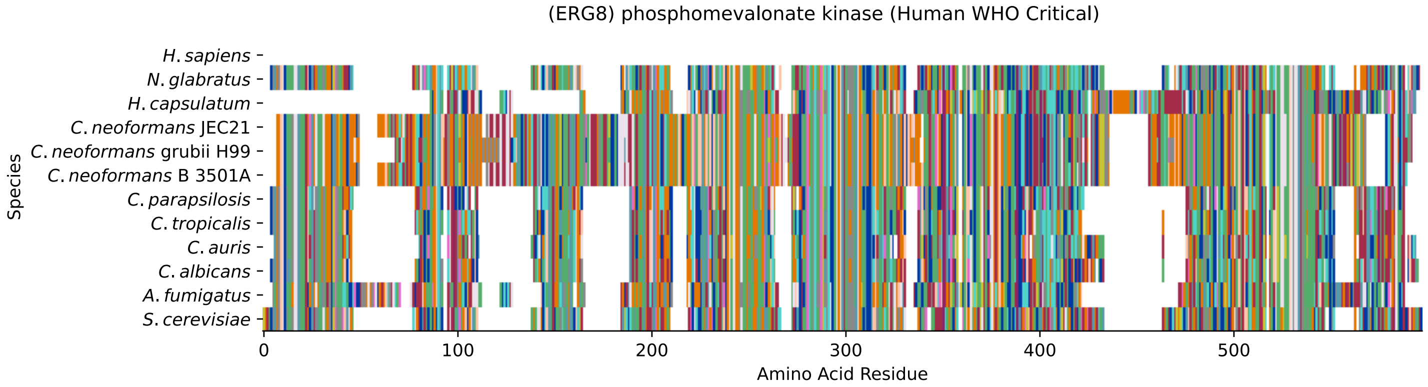


Figure S66: Multiple sequence alignment of yeast Erg8 (WHO Critical Pathogens). Cf. Figure S67 for alignment quality, and Figure S68 for Sneath similarity. Cf. Table S13 for protein names, and pairwise alignment metrics with yeast Erg8.

#### Erg8 MSA Quality

| | *A. f.* | *C. a.* | *C. a.* | *C. p.* | *C. t.* | *C. n.* | *C. n.* | *C. n.* | *H. c.* | *N. g.* | *S. c.* |
|---|---|---|---|---|---|---|---|---|---|---|---|
| *A. fumigatus* | 961.444 | 220.32 | 218.394 | 248.44 | 231.876 | 196.438 | 202.216 | 200.29 | 350.132 | 217.238 | 214.927 |
| *C. albicans* | 238.424 | 883.248 | 478.019 | 461.84 | 641.728 | 162.925 | 179.874 | 171.785 | 132.109 | 365.925 | 315.849 |
| *C. auris* | 214.927 | 444.121 | 876.7 | 415.231 | 439.113 | 158.303 | 176.022 | 165.622 | 126.331 | 370.548 | 327.02 |
| *C. parapsilosis* | 241.891 | 431.409 | 415.231 | 847.81 | 479.174 | 155.221 | 161.384 | 156.377 | 172.17 | 344.354 | 297.745 |
| *C. tropicalis* | 228.409 | 607.06 | 439.113 | 479.174 | 838.951 | 153.68 | 157.147 | 151.754 | 164.466 | 356.681 | 307.76 |
| *C. neoformans* B 3501A | 229.18 | 174.866 | 191.43 | 185.652 | 186.422 | 1092.41 | 887.871 | 1061.98 | 171.4 | 166.392 | 177.178 |
| *C. neoformans* grubii H99 | 225.713 | 183.726 | 202.986 | 187.193 | 184.496 | 855.129 | 1081.24 | 878.241 | 173.326 | 193.356 | 178.718 |
| *C. neoformans* JEC21 | 228.409 | 175.637 | 195.667 | 184.496 | 177.948 | 1061.98 | 897.116 | 1097.03 | 176.792 | 172.17 | 179.874 |
| *H. capsulatum* | 427.557 | 165.622 | 190.66 | 193.356 | 180.644 | 167.933 | 170.629 | 172.555 | 872.848 | 164.851 | 167.933 |
| *N. glabratus* | 213.772 | 337.806 | 370.548 | 344.354 | 356.681 | 134.806 | 160.229 | 135.961 | 150.984 | 909.442 | 409.068 |
| *S. cerevisiae* | 204.912 | 296.59 | 327.02 | 297.745 | 307.76 | 145.976 | 154.066 | 150.599 | 142.51 | 409.068 | 941.028 |

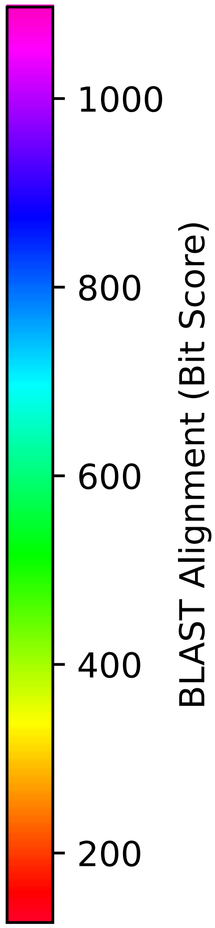


Figure S67: Multiple sequence alignment quality of Erg8 (WHO Critical Pathogens). Cf. Figures S66 and S69.

| Species | Hit Protein | Hit Length (a.a.) | evalue | align_len | bit_score | identity | positive | score | gaps | % identity | % positive |
|---|---|---|---|---|---|---|---|---|---|---|---|
| H.sapiens | - | - | - | - | - | - | - | - | - | - | - |
| N.glabratus | XP_446144.1 uncharacterized protein CAGL0F03993g Nakaseomyces glabratus | 449 | 4.9e-141 | 449 | 409.453 | 210 | 298 | 1051 | 11 | 46.6 | 66.1 |
| H.capsulatum | XP_045287201.1 phosphomevalonate kinase Histoplasma capsulatum G186AR | 446 | 1.6e-37 | 446 | 142.51 | 132 | 202 | 358 | 72 | 29.3 | 44.8 |
| C.neoformans.JEC21 | XP_568385.1 expressed protein Cryptococcus neoformans var. neoformans JEC21 | 546 | 1.9e-40 | 546 | 150.599 | 159 | 240 | 379 | 115 | 35.3 | 53.2 |
| C.neoformans.grubii.H99 | XP_012053077.1 phosphomevalonate kinase Cryptococcus neoformans var. grubii H99 | 536 | 1.6e-41 | 536 | 153.68 | 156 | 244 | 387 | 102 | 34.6 | 54.1 |
| C.neoformans.B.3501A | XP_772275.1 hypothetical protein CNBM0120 Cryptococcus neoformans var. neoformans B-3501A | 540 | 6.5e-39 | 540 | 146.362 | 152 | 237 | 368 | 108 | 33.7 | 52.5 |
| C.parapsilosis | XP_036667432.1 uncharacterized protein CPAR2 400710 Candida parapsilosis | 446 | 1.1e-97 | 446 | 297.745 | 179 | 256 | 761 | 44 | 39.7 | 56.8 |
| C.tropicalis | XP_002545449.1 hypothetical protein CTRG 00230 Candida tropicalis MYA-3404 | 448 | 3.5e-101 | 448 | 307.375 | 182 | 265 | 786 | 46 | 40.4 | 58.8 |
| C.auris | XP_028891858.2 phosphomevalonate kinase Candida auris | 448 | 6.1e-109 | 448 | 327.02 | 197 | 255 | 837 | 33 | 43.7 | 56.5 |
| C.albicans | XP_722678.1 phosphomevalonate kinase Candida albicans SC5314 | 454 | 4.7e-97 | 454 | 296.59 | 191 | 269 | 758 | 35 | 42.4 | 59.6 |
| A.fumigatus | XP_753576.1 phosphomevalonate kinase Aspergillus fumigatus Af293 | 487 | 5.3e-61 | 487 | 204.912 | 157 | 233 | 520 | 71 | 34.8 | 51.7 |

Table S13: Pairwise alignment info from yeast Erg8 (DEG20010822), cf. Figure S66.

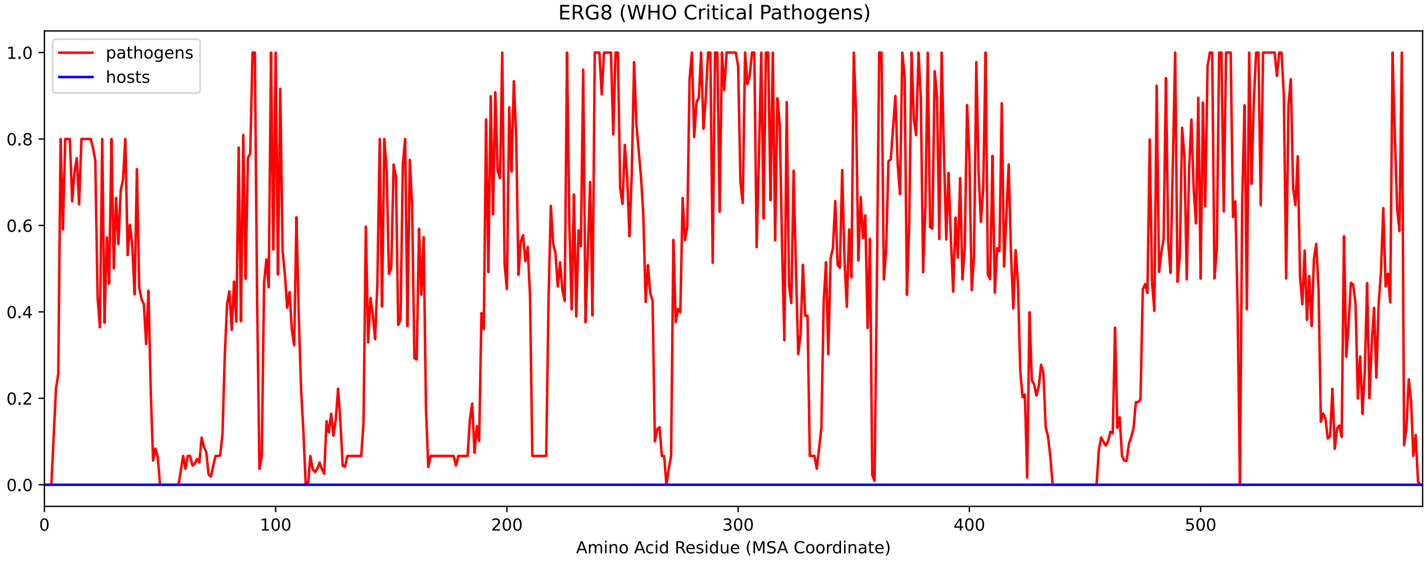


Figure S68: Sneath Similarity of Erg8 for WHO Critical Pathogens, cf. Figure S66

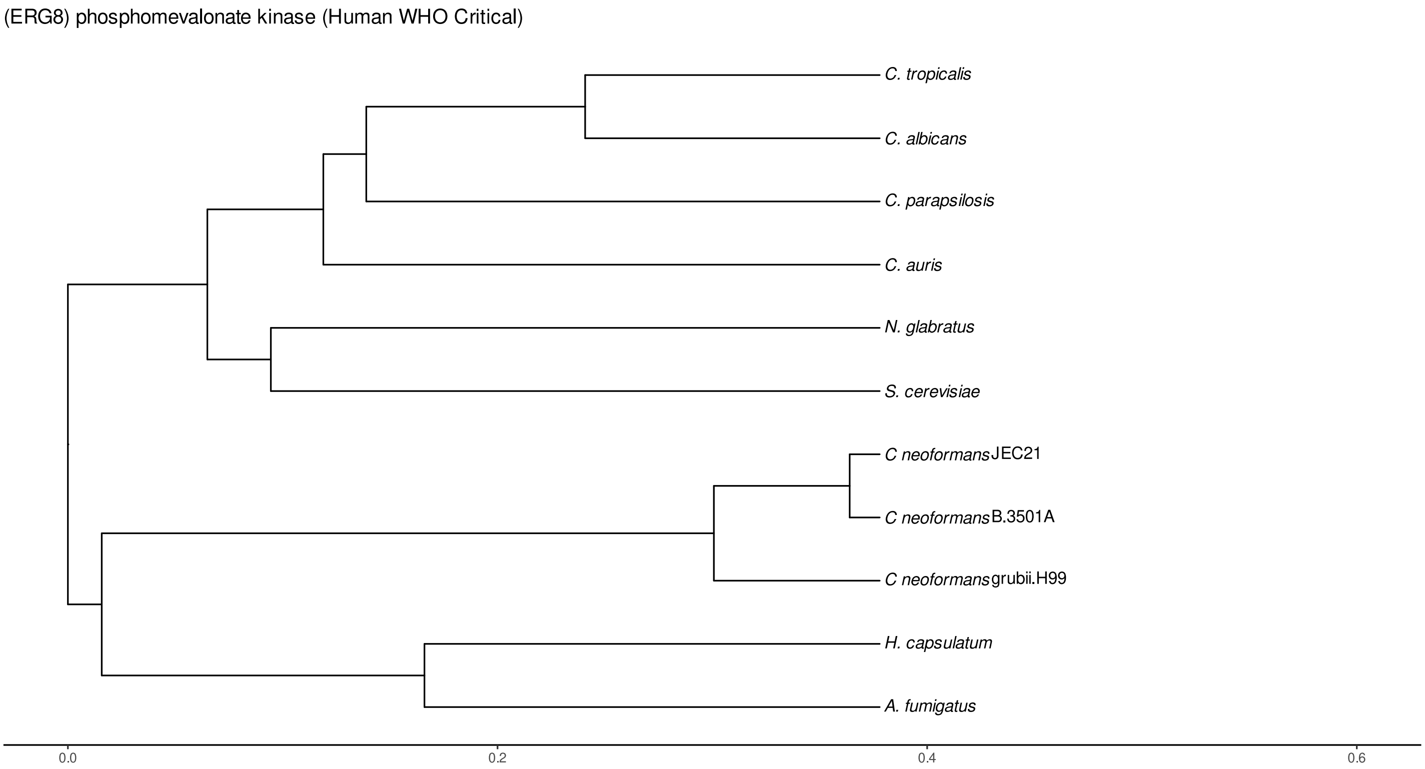


Figure S69: Phylogeny of hits for Erg8 made using clustal omega and visualized using ggtree, cf. Figures S66 and S67.

##### S4.4.2 Top 10 Agricultural Fungal Pathogens

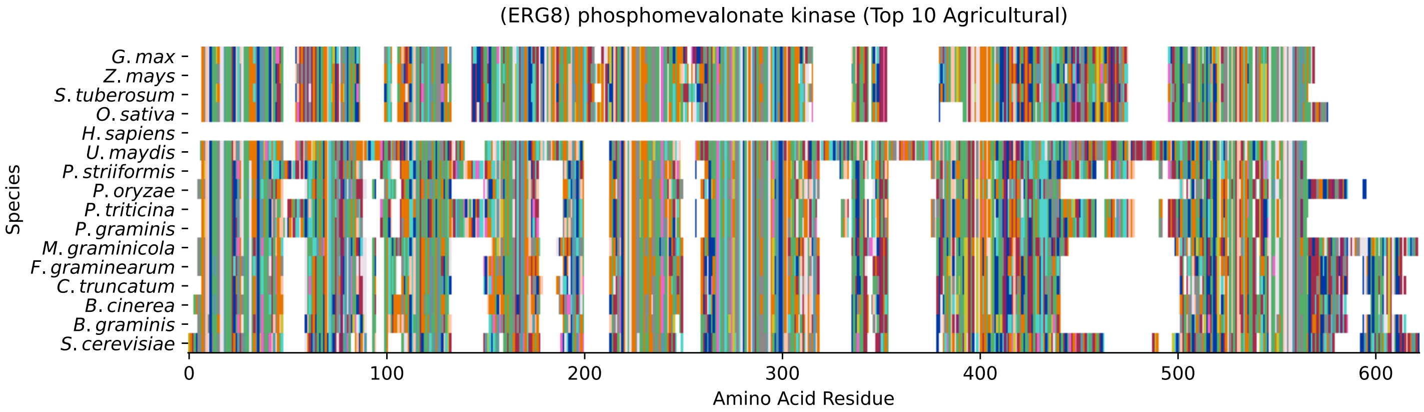


Figure S70: Multiple sequence alignment of yeast Erg8 (Top 10 Agricultural Fungal Pathogens). Cf. Figure S71 for alignment quality, and Figure S72 for Sneath similarity. Cf. Table S14 for protein names, and pairwise alignment metrics with yeast Erg8.

#### Erg8 MSA Quality

| | *B. g.* | *B. c.* | *C. t.* | *F. g.* | *M. g.* | *P. g.* | *P. t.* | *P. o.* | *P. s.* | *U. m.* | *S. c.* |
|---|---|---|---|---|---|---|---|---|---|---|---|
| *B. graminis* | 882.093 | 407.142 | 379.407 | 392.119 | 332.798 | 209.149 | 210.305 | 369.777 | 216.853 | 187.578 | 213.772 |
| *B. cinerea* | 391.734 | 863.218 | 483.411 | 483.026 | 401.749 | 222.246 | 222.246 | 454.136 | 226.098 | 180.259 | 228.794 |
| *C. truncatum* | 355.14 | 476.478 | 876.7 | 600.127 | 378.252 | 236.113 | 237.654 | 581.252 | 232.261 | 184.882 | 256.529 |
| *F. graminearum* | 370.933 | 477.633 | 600.127 | 874.774 | 390.578 | 225.328 | 229.565 | 540.421 | 216.468 | 174.096 | 251.521 |
| *M. graminicola* | 360.918 | 450.284 | 428.713 | 442.195 | 927.546 | 224.942 | 228.409 | 412.535 | 241.891 | 187.963 | 242.662 |
| *P. graminis* | 191.43 | 217.624 | 236.113 | 225.328 | 184.882 | 929.858 | 746.503 | 243.432 | 745.732 | 241.891 | 198.749 |
| *P. triticina* | 192.971 | 219.935 | 237.654 | 229.565 | 187.963 | 746.503 | 927.161 | 247.669 | 719.924 | 244.588 | 181.8 |
| *P. oryzae* | 347.051 | 450.669 | 581.252 | 540.421 | 365.155 | 243.432 | 247.669 | 822.772 | 241.121 | 181.415 | 245.358 |
| *P. striiformis* | 199.904 | 223.402 | 232.261 | 216.468 | 201.83 | 745.732 | 719.924 | 241.121 | 937.562 | 243.817 | 176.022 |
| *U. maydis* | 173.326 | 177.563 | 184.496 | 173.711 | 150.984 | 244.588 | 244.588 | 191.43 | 245.358 | 1056.59 | 155.992 |
| *S. cerevisiae* | 196.052 | 226.868 | 256.529 | 251.521 | 205.297 | 198.749 | 181.8 | 245.358 | 176.022 | 155.606 | 941.028 |

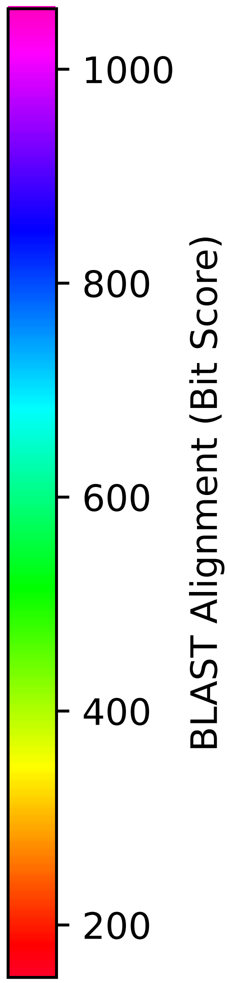


Figure S71: Multiple sequence alignment quality of Erg8 (Top 10 Agricultural Fungal Pathogens). Cf. Figures S70 and S73.

| Species | Hit Protein | Hit Length (a.a.) | evalue | align_len | bit_score | identity | positive | score | gaps | % identity | % positive |
|---|---|---|---|---|---|---|---|---|---|---|---|
| G.max | XP_003526704.1 phosphomevalonate kinase, peroxisomal isoform X3 Glycine max | 462 | 2.3e-42 | 462 | 157.918 | 143 | 217 | 398 | 66 | 31.7 | 48.1 |
| Z.mays | NP_001355169.1 phosphomevalonate kinase Zea mays | 470 | 2.2e-51 | 470 | 182.185 | 151 | 217 | 461 | 78 | 33.5 | 48.1 |
| S.tuberosum | XP_006352929.1 PREDICTED: phosphomevalonate kinase-like Solanum tuberosum | 455 | 3.3e-53 | 455 | 186.422 | 149 | 222 | 472 | 56 | 33.0 | 49.2 |
| O.sativa | XP_015632523.1 phosphomevalonate kinase, peroxisomal isoform X2 Oryza sativa Japonica Group | 473 | 3.6e-48 | 473 | 172.94 | 148 | 212 | 437 | 81 | 32.8 | 47.0 |
| H.sapiens | - | - | - | - | - | - | - | - | - | - | - |
| U.maydis | XP_011386538.1 phosphomevalonate kinase Ustilago maydis 521 | 526 | 9.7e-42 | 526 | 155.221 | 148 | 214 | 391 | 131 | 32.8 | 47.5 |
| P.striiformis | XP_047806802.1 hypothetical protein Pst134EA 013927 Puccinia striiformis f. sp. tritici | 459 | 2.6e-49 | 459 | 175.637 | 141 | 217 | 444 | 63 | 31.3 | 48.1 |
| P.oryzae | mRNA M BR32 EuGene 00127411-p1 — transcript=mRNA M BR32 EuGene 00127411 — gene=M BR32 EuGene 00127411 — organism=Pyricularia oryzae BR32 — gene product=unspecified product — transcript product=unspecified product — location=BR32 scaffold00023:241792-243188(+) — protein length=445 — sequence SO=supercontig — SO=protein coding gene — is pseudo=false | 451 | 9.1e-77 | 451 | 245.358 | 164 | 242 | 625 | 58 | 36.4 | 53.7 |
| P.triticina | XP_053022619.1 uncharacterized protein PtA15 7A793 Puccinia triticina | 458 | 1.1e-51 | 458 | 182.185 | 144 | 222 | 461 | 63 | 31.9 | 49.2 |
| P.graminis | XP_003888790.1 hypothetical protein PGTG 22505 Puccinia graminis f. sp. tritici CRL 75-36-700-3 | 460 | 7.7e-58 | 460 | 198.364 | 145 | 222 | 503 | 67 | 32.2 | 49.2 |
| M.graminicola | ZTRI 9.521.mRNA-p1 — transcript=ZTRI 9.521.mRNA — gene=ZTRI 9.521 — organism=Zymoseptoria tritici IPO323 — gene product=similar to phosphomevalonate kinase — transcript product=similar to phosphomevalonate kinase — location=Ztri chr 9:1739392-1740886(-) — protein length=464 — sequence SO=chromosome — SO=protein coding gene — is pseudo=false | 478 | 4.3e-61 | 478 | 204.912 | 166 | 244 | 520 | 59 | 36.8 | 54.1 |
| F.graminearum | XP_011327956.1 hypothetical protein FGSG 09764 Fusarium graminearum PH-1 | 461 | 4.6e-79 | 461 | 251.521 | 164 | 244 | 641 | 59 | 36.4 | 54.1 |
| C.truncatum | XP_036588544.1 phosphomevalonate kinase Colletotrichum truncatum | 467 | 1.5e-80 | 467 | 255.758 | 173 | 248 | 652 | 68 | 38.4 | 55.0 |
| B.cinerea | XP_001553931.1 Bcerg8 Botrytis cinerea B05.10 | 460 | 1.9e-68 | 460 | 226.098 | 155 | 233 | 575 | 55 | 34.4 | 51.7 |
| B.graminis | VDB92769.1 — transcript=BGT96224V316 LOCUS6538 t1 — gene=BGT96224V316 LOCUS6538 — organism=Blumeria graminis f. sp. tritici 96224 — gene product=unspecified product — transcript product=unspecified product — location=LR026992:3440777-3442168(-) — protein length=438 — sequence SO=chromosome — SO=protein coding gene — is pseudo=false | 462 | 3.7e-58 | 462 | 195.667 | 159 | 220 | 496 | 60 | 35.3 | 48.8 |

Table S14: Pairwise alignment info from yeast Erg8 (DEG20010822), cf. Figure S70.

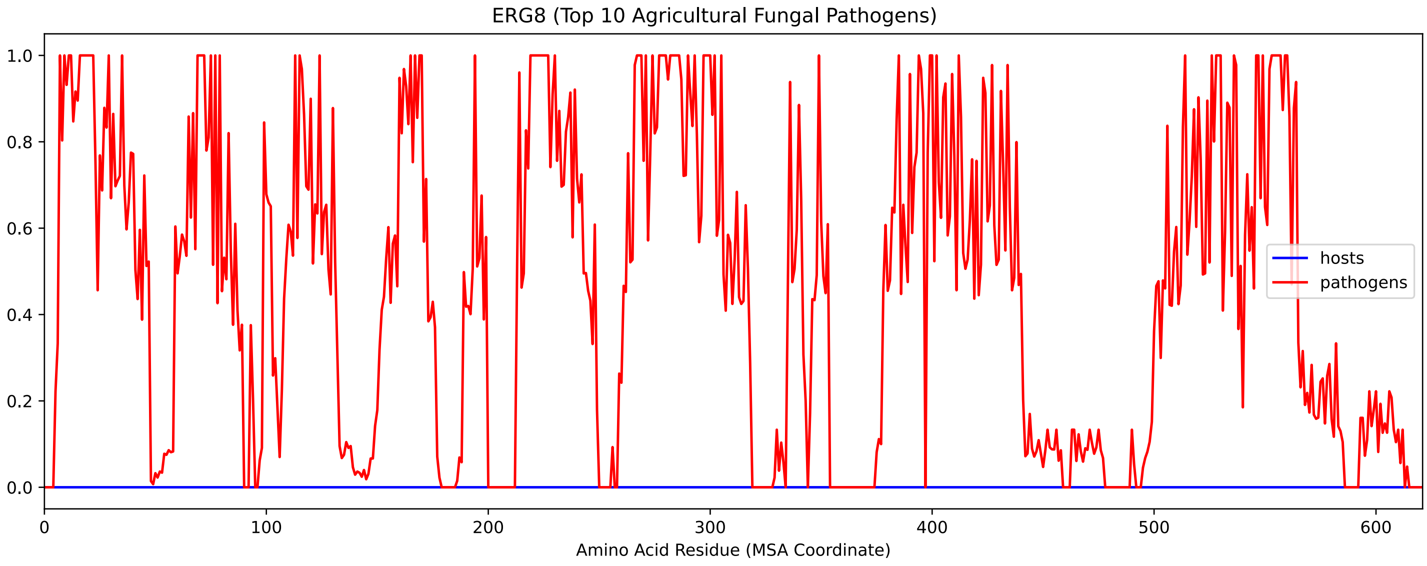


Figure S72: Sneath Similarity of Erg8 for Top 10 Agricultural Fungal Pathogens, cf. Figure S70

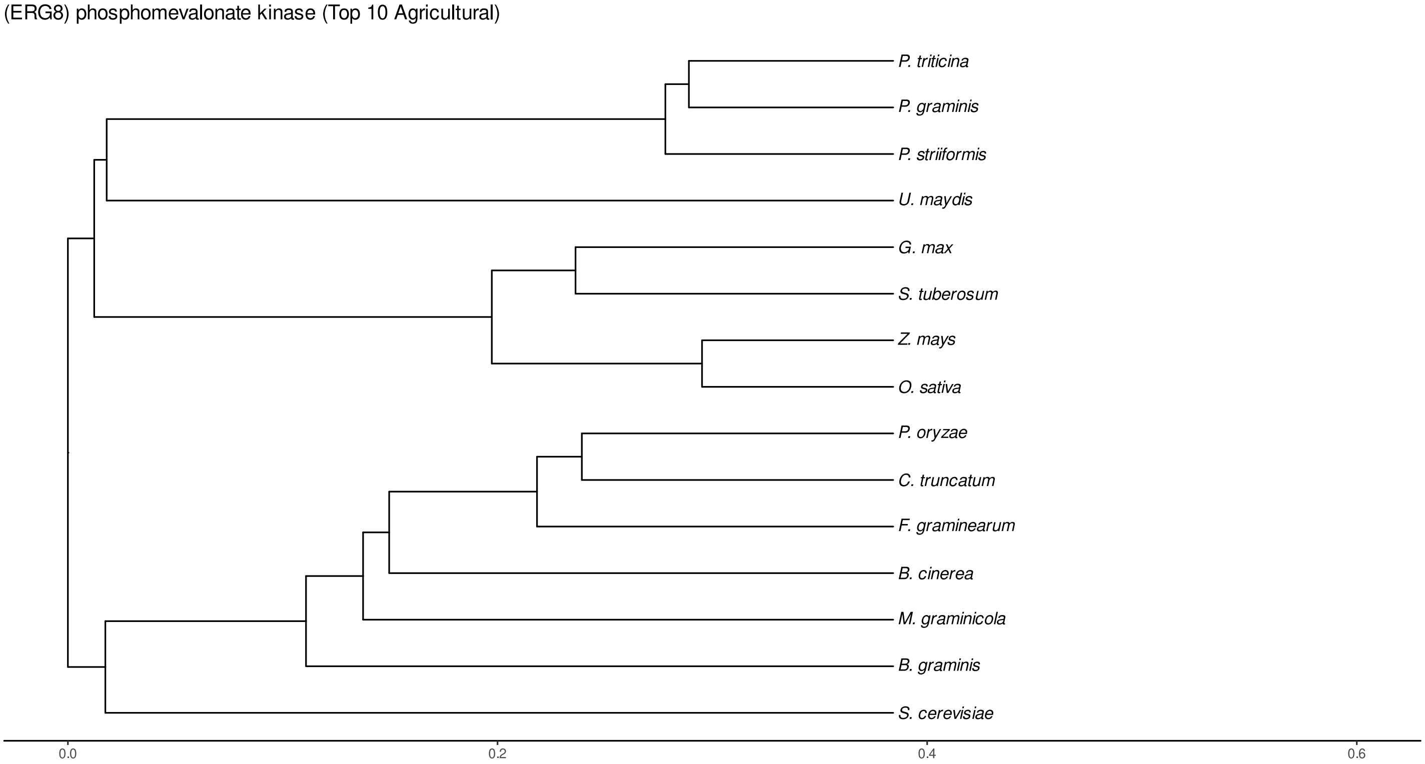


Figure S73: Phylogeny of hits for Erg8 made using clustal omega and visualized using ggtree, cf. Figures S70 and S71.

#### S4.4.3 NR

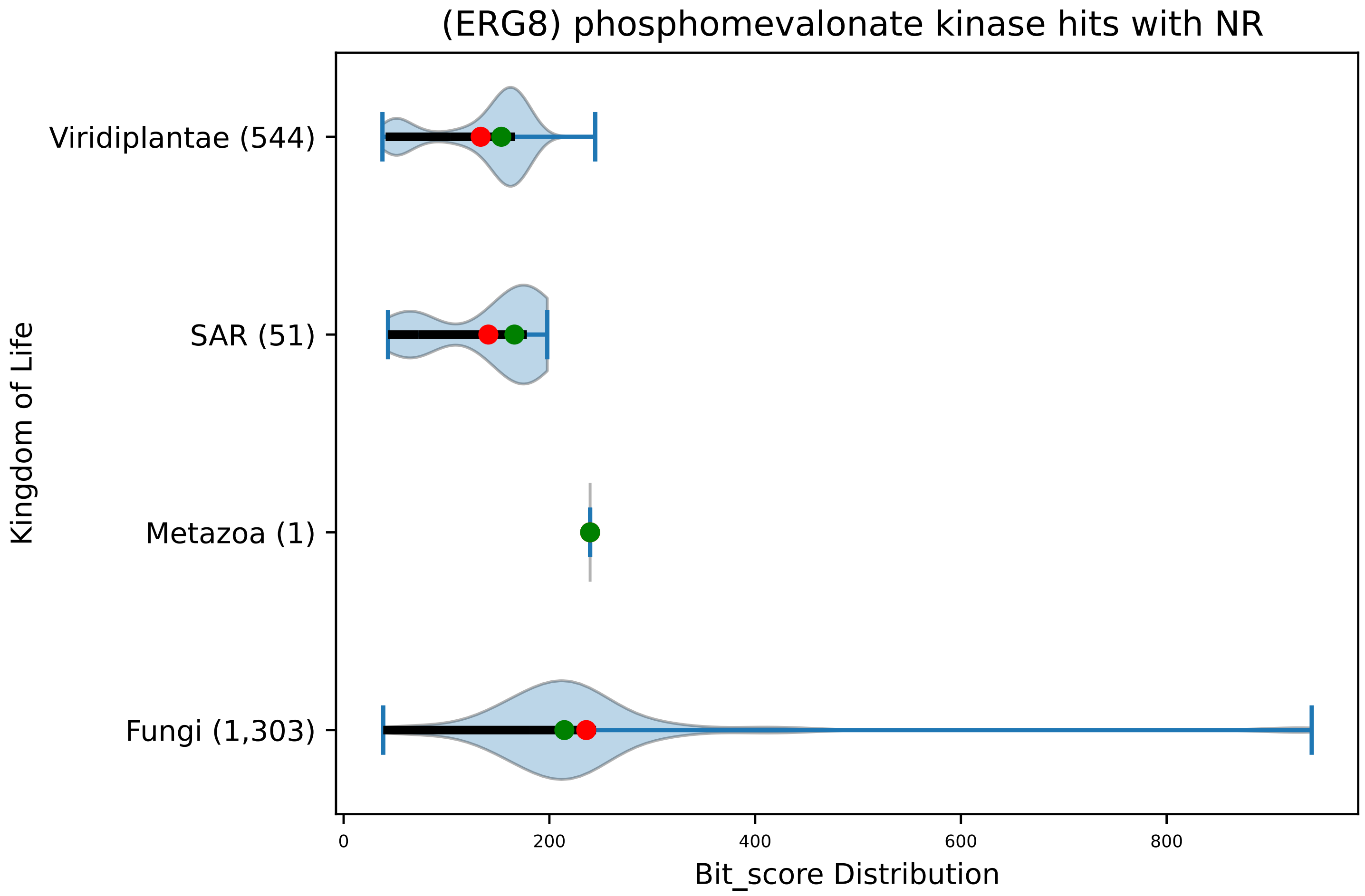


Figure S74: Non-redundant (NR) protein hits for DEG20010822/Erg8, with expectation value of no more than 0.1. Green points are medians, and red points are arithmetic means.

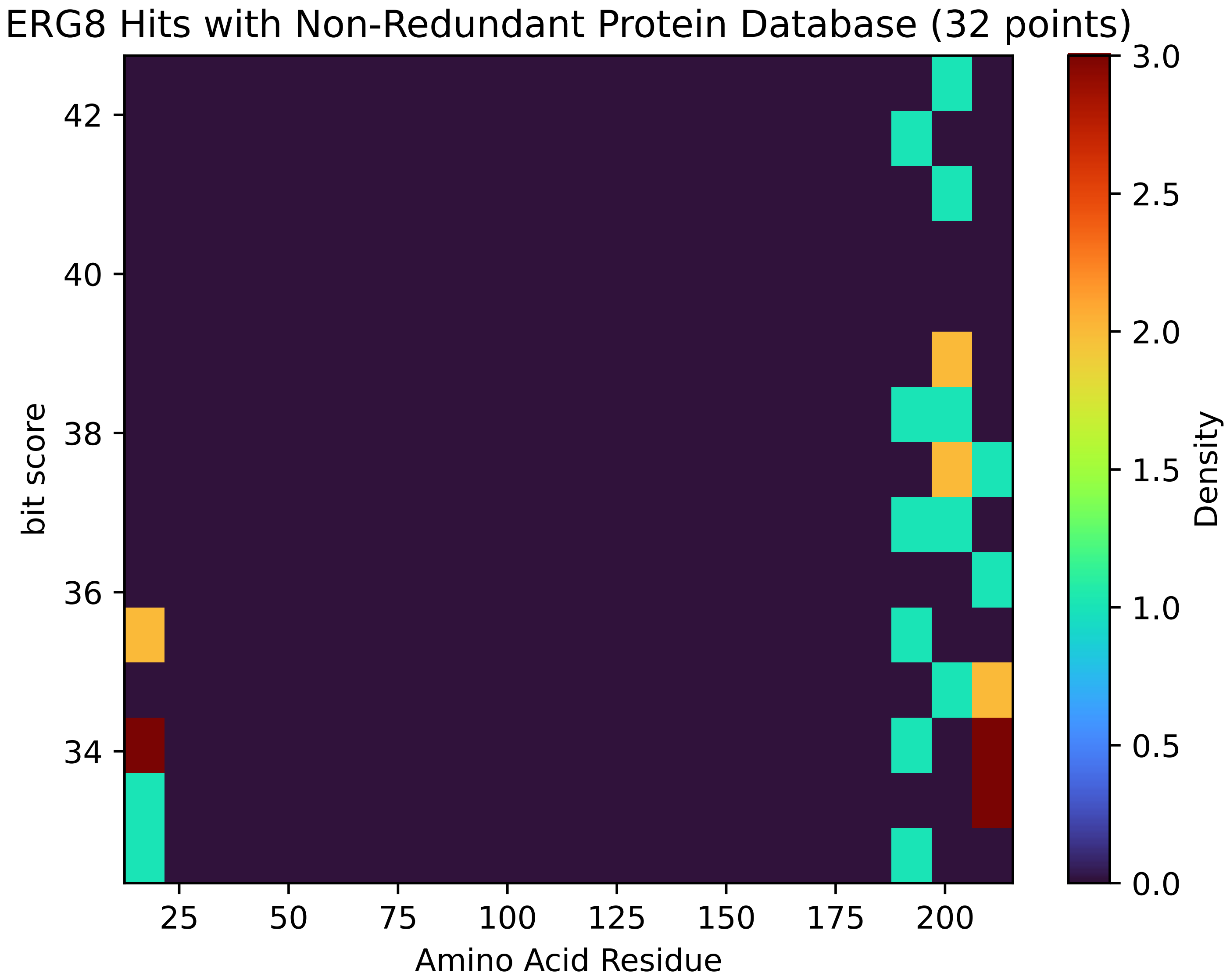


Figure S75: Non-redundant (NR) protein hits for Erg8 in the kingdom Metazoa.

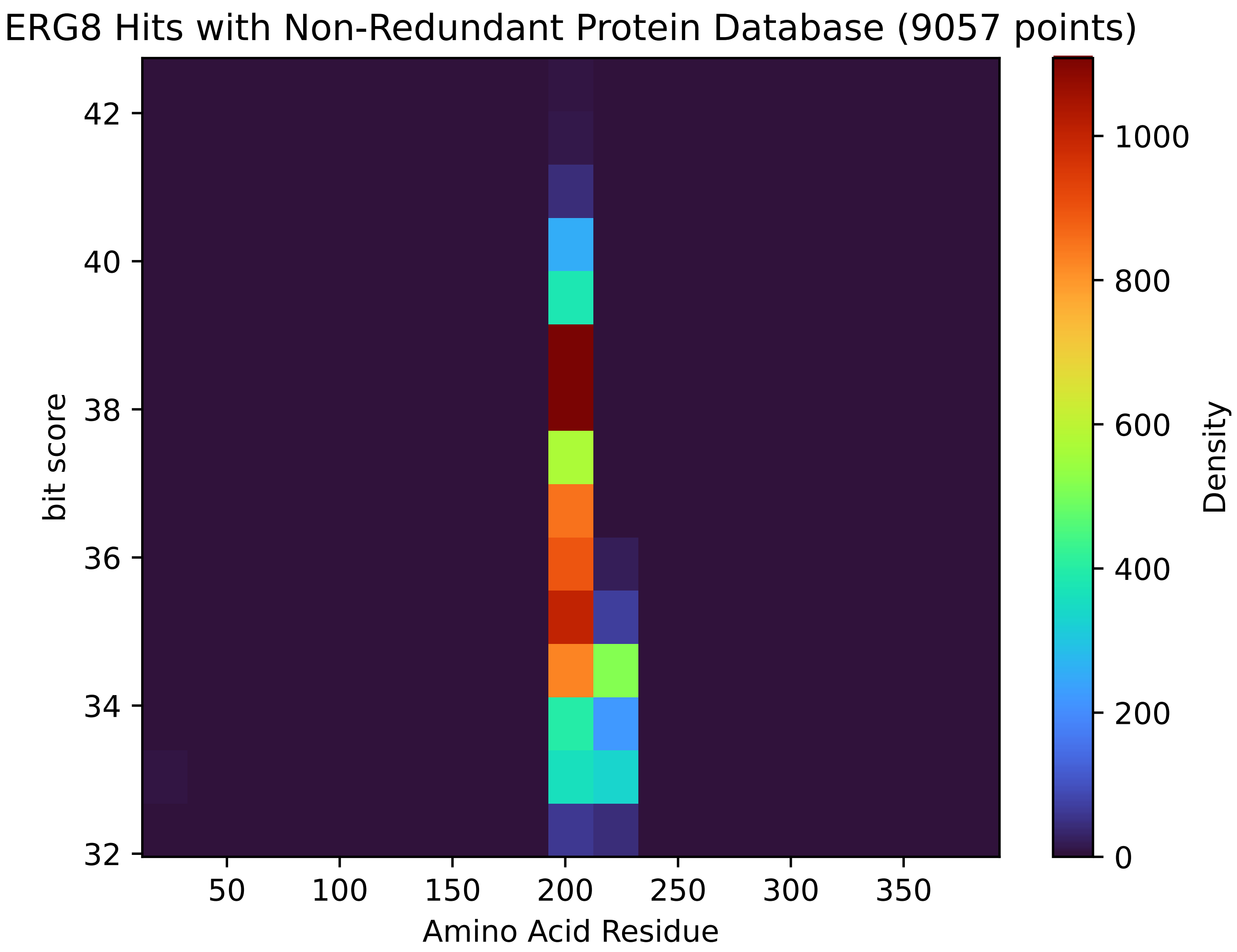


Figure S76: Non-redundant (NR) protein hits for Erg8 in the kingdom Viridiplantae.

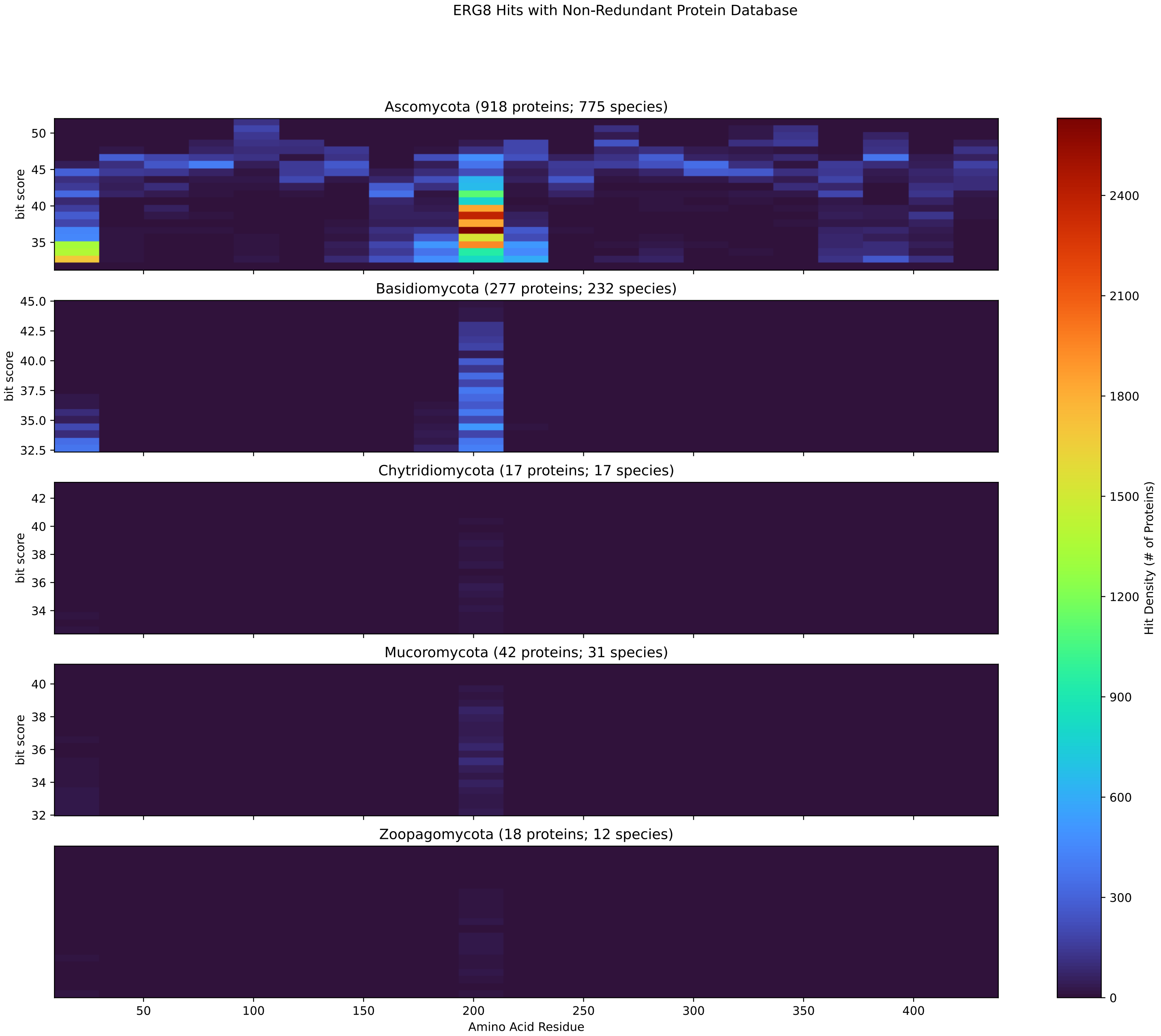


Figure S77: Non-redundant (NR) protein hits for Erg8 in the kingdom Fungi.

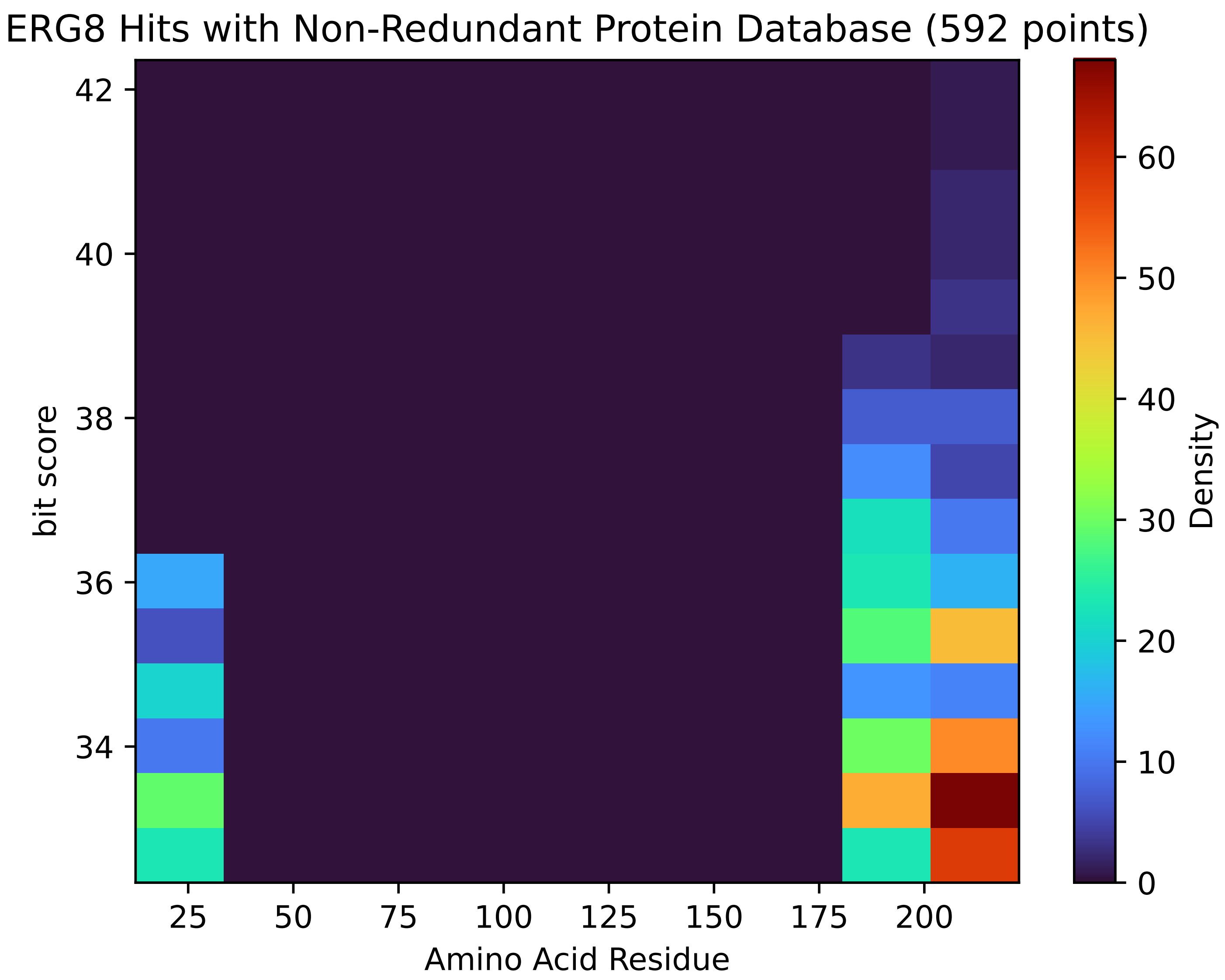


Figure S78: Non-redundant (NR) protein hits for Erg8 in the kingdom SAR.

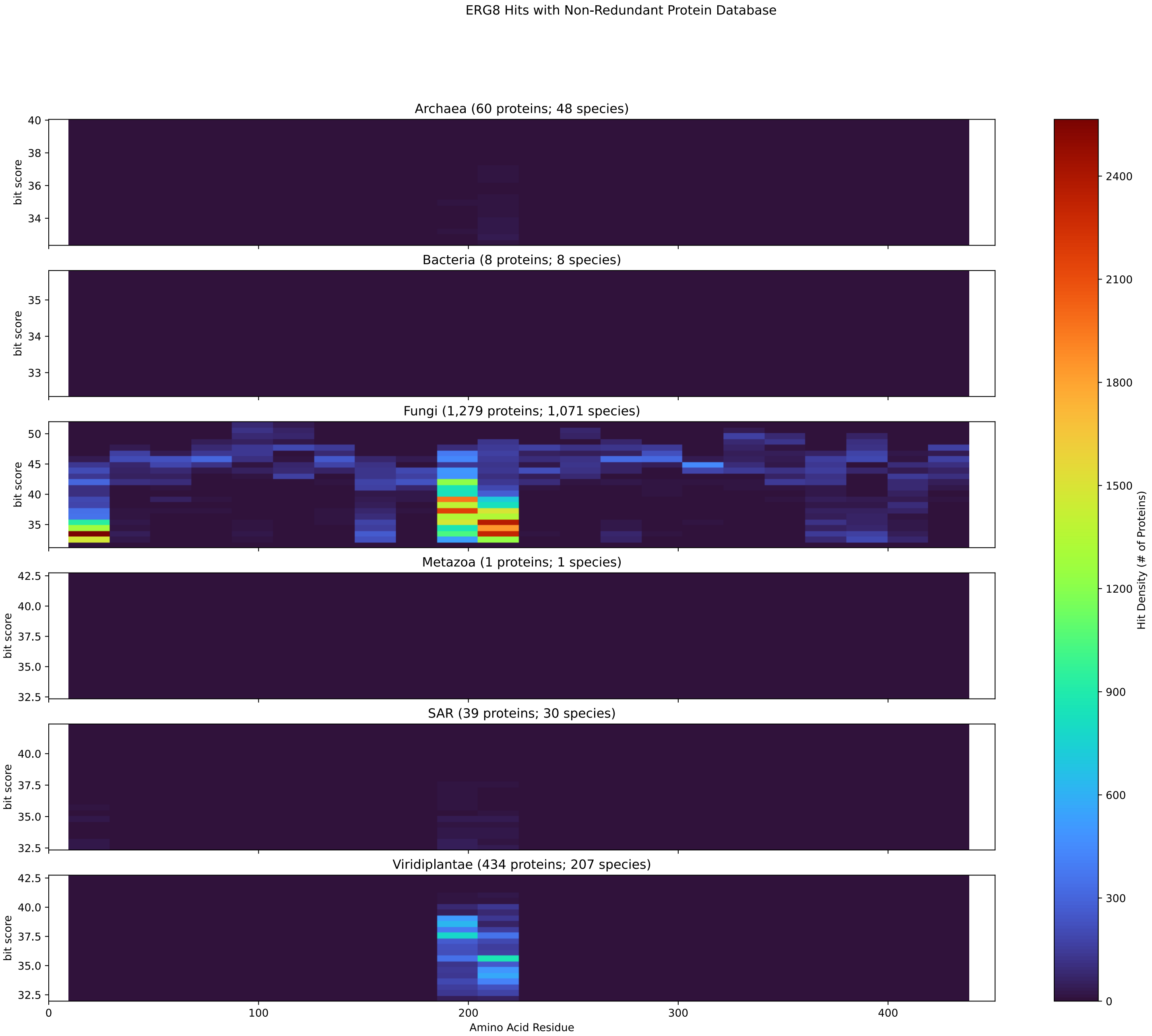


Figure S79: Non-redundant (NR) protein hits for DEG20010822/Erg8 at 20 amino acid length queries.

#### S4.5 Fas1

##### S4.5.1 WHO Critical Pathogens

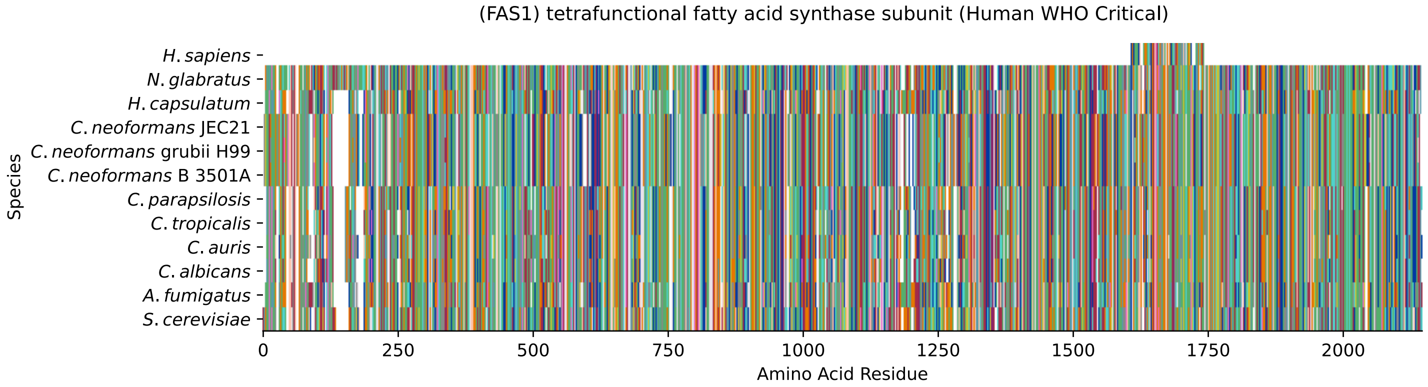


Figure S80: Multiple sequence alignment of yeast Fas1 (WHO Critical Pathogens). Cf. Figure S81 for alignment quality, and Figure S82 for Sneath similarity. Cf. Table S15 for protein names, and pairwise alignment metrics with yeast Fas1.

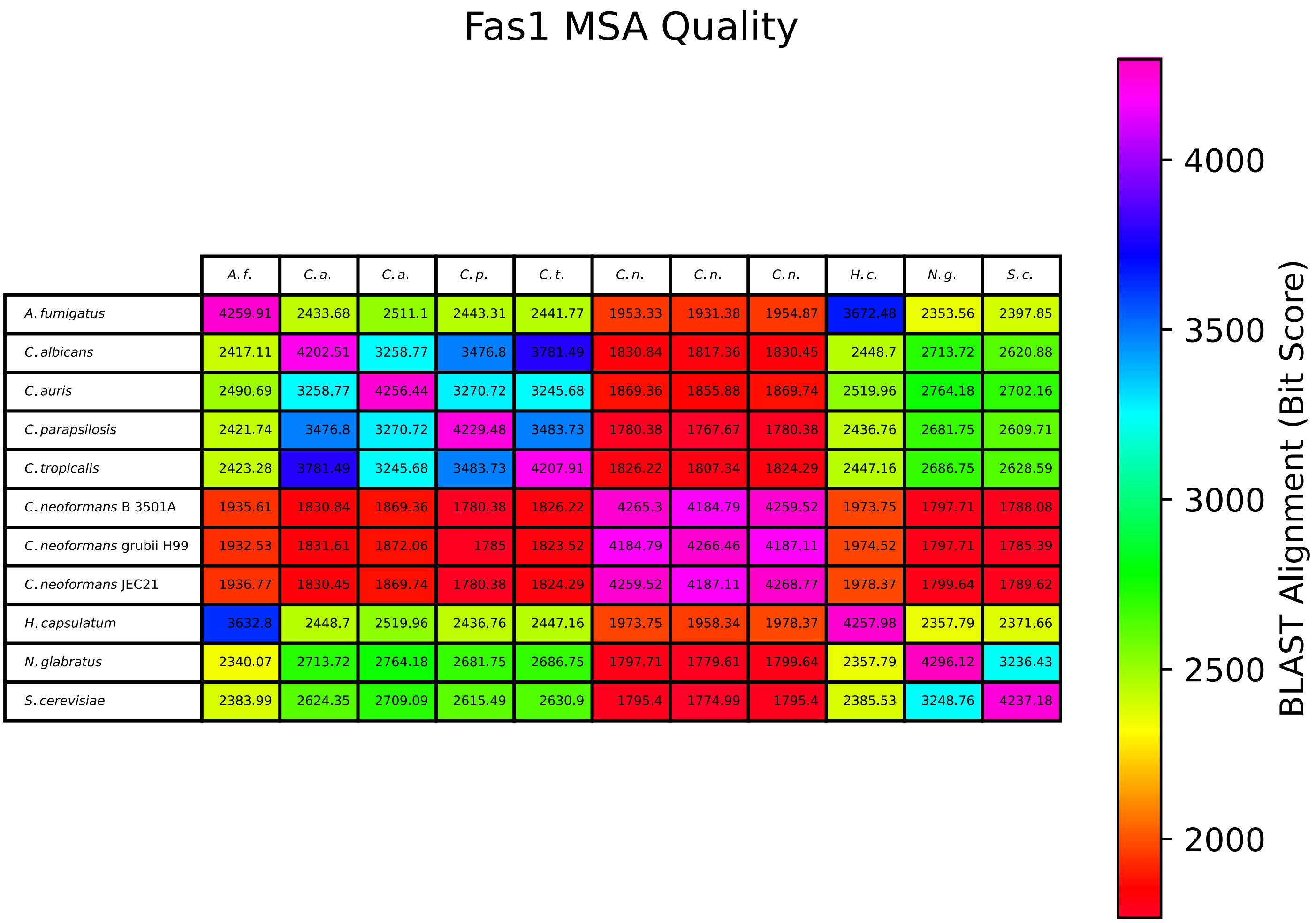


| | *A. f.* | *C. a.* | *C. a.* | *C. p.* | *C. t.* | *C. n.* | *C. n.* | *C. n.* | *H. c.* | *N. g.* | *S. c.* |
|---|---|---|---|---|---|---|---|---|---|---|---|
| *A. fumigatus* | 4259.91 | 2433.68 | 2511.1 | 2443.31 | 2441.77 | 1953.33 | 1931.38 | 1954.87 | 3672.48 | 2353.56 | 2397.85 |
| *C. albicans* | 2417.11 | 4202.51 | 3258.77 | 3476.8 | 3781.49 | 1830.84 | 1817.36 | 1830.45 | 2448.7 | 2713.72 | 2620.88 |
| *C. auris* | 2490.69 | 3258.77 | 4256.44 | 3270.72 | 3245.68 | 1869.36 | 1855.88 | 1869.74 | 2519.96 | 2764.18 | 2702.16 |
| *C. parapsilosis* | 2421.74 | 3476.8 | 3270.72 | 4229.48 | 3483.73 | 1780.38 | 1767.67 | 1780.38 | 2436.76 | 2681.75 | 2609.71 |
| *C. tropicalis* | 2423.28 | 3781.49 | 3245.68 | 3483.73 | 4207.91 | 1826.22 | 1807.34 | 1824.29 | 2447.16 | 2686.75 | 2628.59 |
| *C. neoformans* B 3501A | 1935.61 | 1830.84 | 1869.36 | 1780.38 | 1826.22 | 4265.3 | 4184.79 | 4259.52 | 1973.75 | 1797.71 | 1788.08 |
| *C. neoformans* grubii H99 | 1932.53 | 1831.61 | 1872.06 | 1785 | 1823.52 | 4184.79 | 4266.46 | 4187.11 | 1974.52 | 1797.71 | 1785.39 |
| *C. neoformans* JEC21 | 1936.77 | 1830.45 | 1869.74 | 1780.38 | 1824.29 | 4259.52 | 4187.11 | 4268.77 | 1978.37 | 1799.64 | 1789.62 |
| *H. capsulatum* | 3632.8 | 2448.7 | 2519.96 | 2436.76 | 2447.16 | 1973.75 | 1958.34 | 1978.37 | 4257.98 | 2357.79 | 2371.66 |
| *N. glabratus* | 2340.07 | 2713.72 | 2764.18 | 2681.75 | 2686.75 | 1797.71 | 1779.61 | 1799.64 | 2357.79 | 4296.12 | 3236.43 |
| *S. cerevisiae* | 2383.99 | 2624.35 | 2709.09 | 2615.49 | 2630.9 | 1795.4 | 1774.99 | 1795.4 | 2385.53 | 3248.76 | 4237.18 |

Figure S81: Multiple sequence alignment quality of Fas1 (WHO Critical Pathogens). Cf. Figures S80 and S83.

| Species | Hit Protein | Hit Length (a.a.) | evalue | align_len | bit_score | identity | positive | score | gaps | % identity | % positive |
|---|---|---|---|---|---|---|---|---|---|---|---|
| H.sapiens | NP_001278957.1 peroxisomal mu-ltifunctional enzyme type 2 is-oform 5 Homo sapiens | 133 | 2.1e-06 | 133 | 55.4546 | 40 | 64 | 132 | 9 | 2.0 | 3.1 |
| N.glabratus | XP_445436.1 uncharacterized p-rotein CAGL0D00528g Nakaseomyc-es glabratus | 2083 | 0 | 2083 | 3247.6 | 1545 | 1782 | 8419 | 44 | 75.3 | 86.9 |
| H.capsulatum | XP_045283903.1 fatty acid syn-thase beta subunit dehydratase Histoplasma capsulatum G186AR | 2069 | 0 | 2069 | 2385.14 | 1151 | 1510 | 6180 | 34 | 56.1 | 73.6 |
| C.neoformans.JEC21 | XP_571100.1 fatty-acid syntha-se complex protein, putative C-ryptococcus neoformans var. ne-oformans JEC21 | 2090 | 0 | 2090 | 1798.1 | 933 | 1327 | 4656 | 66 | 45.5 | 64.7 |
| C.neoformans.grubii.H99 | XP_012049942.1 fatty acid syn-thase subunit beta, fungi type Cryptococcus neoformans var. grubii H99 | 2089 | 0 | 2089 | 1774.6 | 924 | 1316 | 4595 | 64 | 45.1 | 64.2 |
| C.neoformans.B.3501A | XP_775164.1 hypothetical prot-ein CNBE4370 Cryptococcus neof-ormans var. neoformans B-3501A | 2090 | 0 | 2090 | 1795.4 | 933 | 1327 | 4649 | 66 | 45.5 | 64.7 |
| C.parapsilosis | XP_036664454.1 uncharacterize-d protein CPAR2 302650 Candida parapsilosis | 2064 | 0 | 2064 | 2615.49 | 1280 | 1576 | 6778 | 42 | 62.4 | 76.8 |
| C.tropicalis | XP_002550943.1 fatty acid syn-thase beta subunit dehydratase Candida tropicalis MYA-3404 | 2059 | 0 | 2059 | 2630.51 | 1279 | 1576 | 6817 | 40 | 62.4 | 76.8 |
| C.auris | XP_028891947.2 fatty acid syn-thase subunit beta Candida aur-is | 2058 | 0 | 2058 | 2711.02 | 1291 | 1601 | 7026 | 16 | 62.9 | 78.1 |
| C.albicans | XP_716817.1 tetrafunctional f-atty acid synthase subunit Can-dida albicans SC5314 | 2061 | 0 | 2061 | 2624.74 | 1278 | 1586 | 6802 | 46 | 62.3 | 77.3 |
| A.fumigatus | XP_748739.1 fatty acid syntha-se beta subunit, putative Aspe-rgillus fumigatus Af293 | 2071 | 0 | 2071 | 2387.07 | 1173 | 1510 | 6185 | 37 | 57.2 | 73.6 |

Table S15: Pairwise alignment info from yeast Fas1 (DEG20010641), cf. Figure S80.

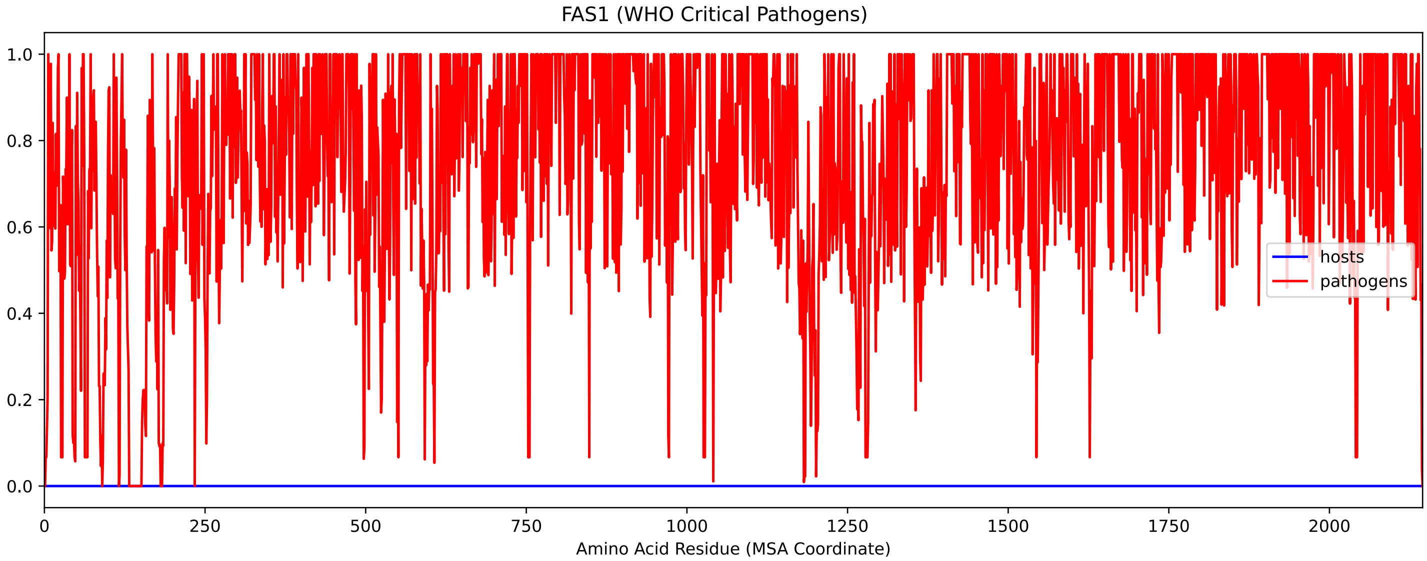


Figure S82: Sneath Similarity of Fas1 for WHO Critical Pathogens, cf. Figure S80

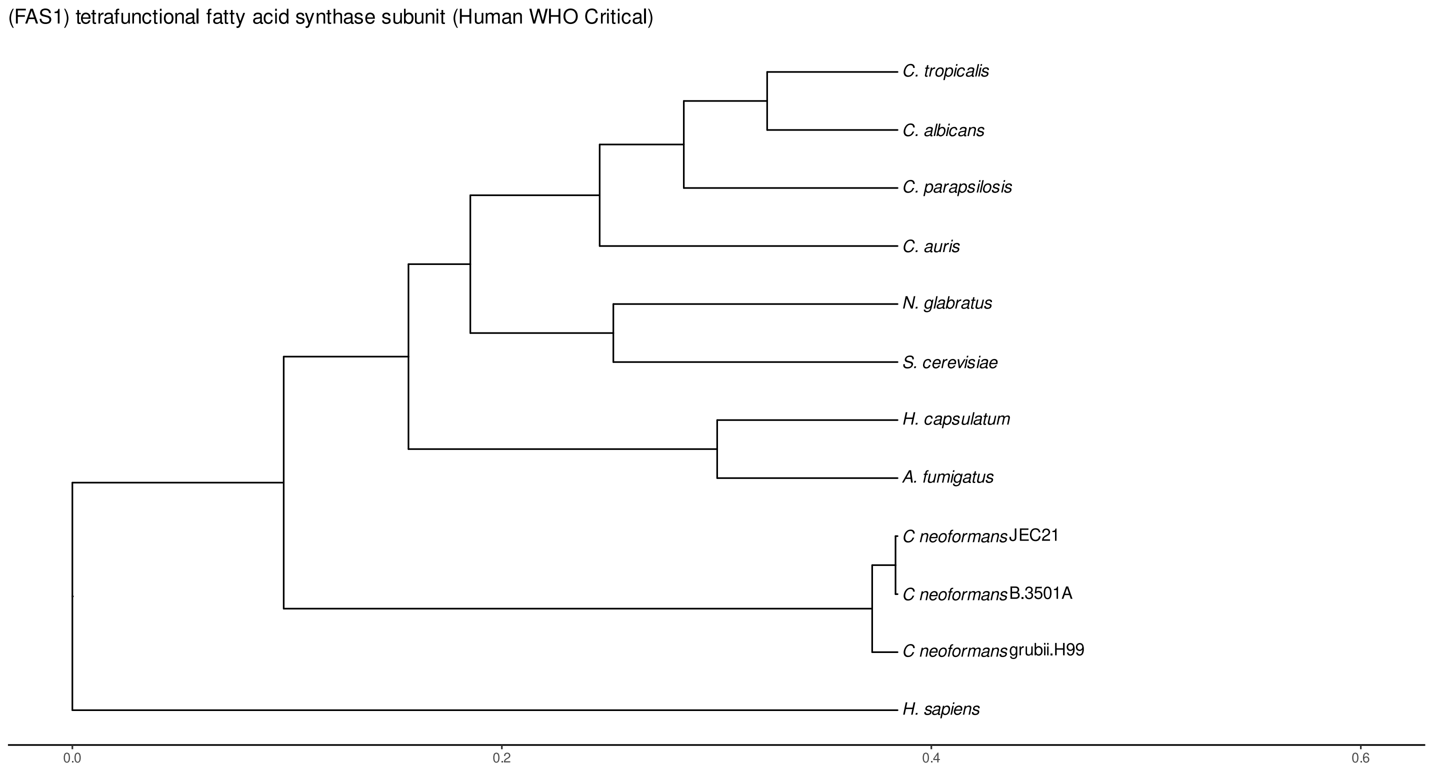


Figure S83: Phylogeny of hits for Fas1 made using clustal omega and visualized using ggtree, cf. Figures S80 and S81.

##### S4.5.2 Top 10 Agricultural Fungal Pathogens

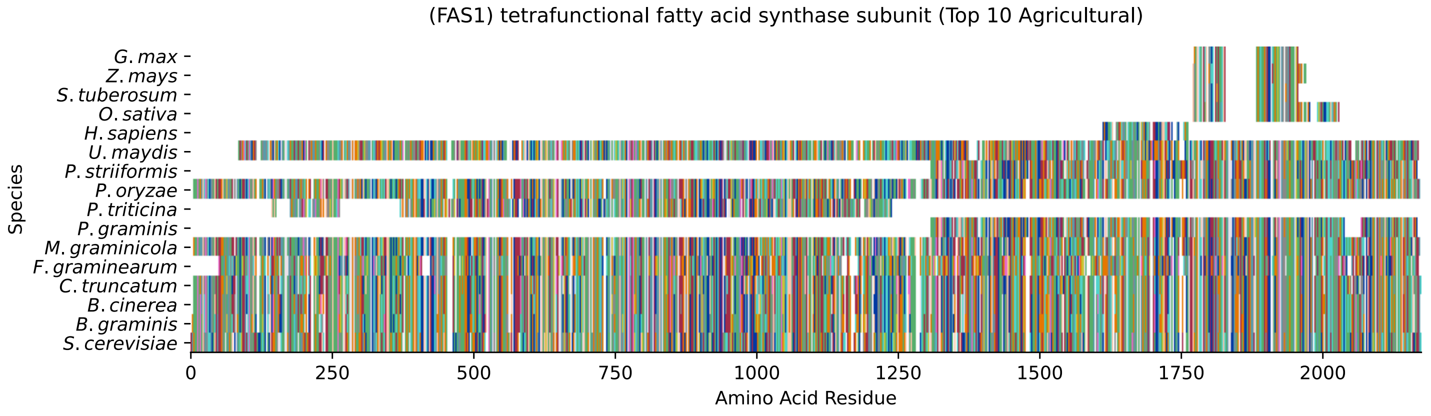


Figure S84: Multiple sequence alignment of yeast Fas1 (Top 10 Agricultural Fungal Pathogens). Cf. Figure S85 for alignment quality, and Figure S86 for Sneath similarity. Cf. Table S16 for protein names, and pairwise alignment metrics with yeast Fas1.

#### Fas1 MSA Quality

| | B.g. | B.c. | C.t. | F.g. | M.g. | P.g. | P.t. | P.o. | P.s. | U.m. | S.c. |
|---|---|---|---|---|---|---|---|---|---|---|---|
| *B. graminis* | 4277.63 | 3644.74 | 3292.67 | 1448.72 | 3231.81 | 729.939 | 728.783 | 3298.45 | 779.63 | 1353.96 | 0 |
| *B. cinerea* | 3644.74 | 4262.6 | 3456.77 | 1434.08 | 3377.8 | 739.569 | 737.258 | 3434.81 | 790.415 | 1324.3 | 0 |
| *C. truncatum* | 3303.84 | 3472.17 | 4263.76 | 1430.62 | 3163.24 | 734.561 | 726.472 | 3437.89 | 768.459 | 1333.16 | 0 |
| *F. graminearum* | 1461.43 | 1444.1 | 1433.31 | 4100.05 | 1413.67 | 433.335 | 535.798 | 1403.65 | 434.491 | 968.763 | 0 |
| *M. graminicola* | 3231.81 | 3377.8 | 3155.93 | 1400.19 | 4274.55 | 711.449 | 739.954 | 3121.26 | 776.548 | 1322.38 | 0 |
| *P. graminis* | 729.939 | 739.569 | 734.176 | 432.95 | 711.449 | 1688.32 | 0 | 721.85 | 1579.69 | 517.694 | 0 |
| *P. triticina* | 736.102 | 745.347 | 737.643 | 536.954 | 752.281 | 0 | 1930.22 | 730.324 | 0 | 711.064 | 0 |
| *P. oryzae* | 3307.69 | 3446.37 | 3437.51 | 1402.5 | 3131.66 | 721.85 | 720.694 | 4271.46 | 770.385 | 1309.28 | 0 |
| *P. striiformis* | 779.63 | 790.415 | 768.074 | 434.876 | 776.548 | 1579.69 | 0 | 770.385 | 1746.87 | 532.717 | 0 |
| *U. maydis* | 1353.96 | 1324.3 | 1327.38 | 959.133 | 1322.38 | 517.694 | 699.893 | 1302.35 | 532.717 | 4087.34 | 0 |
| *S. cerevisiae* | 0 | 0 | 0 | 0 | 0 | 0 | 0 | 0 | 0 | 0 | 0 |

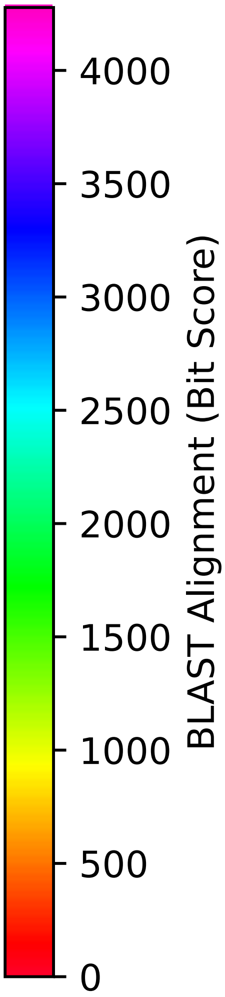


Figure S85: Multiple sequence alignment quality of Fas1 (Top 10 Agricultural Fungal Pathogens). Cf. Figures S84 and S87.

| Species | Hit Protein | Hit Length (a.a.) | evalue | align_len | bit_score | identity | positive | score | gaps | % identity | % positive |
|---|---|---|---|---|---|---|---|---|---|---|---|
| G.max | NP_001238312.2 malonyltransfe- rase Glycine max | 184 | 2.3e-05 | 184 | 50.0618 | 52 | 73 | 118 | 63 | 2.5 | 3.6 |
| Z.mays | NP_001150334.1 uncharacterize- d protein LOC100283964 Zea may- s | 203 | 0.00035 | 203 | 46.2098 | 52 | 82 | 108 | 68 | 2.5 | 4.0 |
| S.tuberosum | XP_006348452.1 PREDICTED: mal- onyl CoA-acyl carrier protein transacylase Solanum tuberosum | 187 | 3.6e-05 | 187 | 48.521 | 52 | 74 | 114 | 63 | 2.5 | 3.6 |
| O.sativa | XP_015628213.1 uncharacterize- d protein LOC4332547 Oryza sat- iva Japonica Group | 256 | 0.00017 | 256 | 46.595 | 63 | 101 | 109 | 73 | 3.1 | 4.9 |
| H.sapiens | NP_001278957.1 peroxisomal mu- ltifunctional enzyme type 2 is- oform 5 Homo sapiens | 133 | 2.1e-06 | 133 | 55.4546 | 40 | 64 | 132 | 9 | 2.0 | 3.1 |
| U.maydis | XP_011392728.1 fatty acid syn- thase FAS2 Ustilago maydis 521 | 2052 | 0 | 2052 | 1249.57 | 742 | 1122 | 3232 | 159 | 36.2 | 54.7 |
| P.striiformis | XP_047808761.1 hypothetical p- rotein Pst134EA 009828 Puccini- a striiformis f. sp. tritici | 845 | 0 | 845 | 725.702 | 394 | 543 | 1872 | 32 | 19.2 | 26.5 |
| P.oryzae | mRNA M BR32 EuGene 00072671-p1 — transcript=mRNA M BR32 EuGe- ne 00072671 — gene=M BR32 EuGe- ne 00072671 — organism=Pyricul- aria oryzae BR32 — gene produc- t=unspecified product — transc- ript product=unspecified produ- ct — location=BR32 scaffold000- 07:1168071-1174501(+) — protei- n length=2120 — sequence SO=su- percontig — SO=protein coding gene — is pseudo=false | 2074 | 0 | 2074 | 2357.79 | 1160 | 1502 | 6109 | 40 | 56.6 | 73.2 |
| P.triticina | XP_053022848.1 uncharacterize- d protein PtA15 8A197 Puccinia triticina | 979 | 0 | 979 | 722.235 | 406 | 556 | 1863 | 75 | 19.8 | 27.1 |
| P.graminis | XP_003325251.2 fatty acid syn- thase subunit beta Puccinia gr- aminis f. sp. tritici CRL 75-3- 6-700-3 | 846 | 0 | 846 | 671.774 | 375 | 518 | 1732 | 62 | 18.3 | 25.3 |
| M.graminicola | ZTRI 3.48.mRNA-p1 — transcript- =ZTRI 3.48.mRNA — gene=ZTRI 3.- 48 — organism=Zymoseptoria tri- tici IPO323 — gene product=sim- ilar to fatty acid synthase su- bunit beta dehydratase — trans- cript product=similar to fatty acid synthase subunit beta de- hydratase — location=Ztri chr 3:163778-170201(+) — protein l- ength=2121 — sequence SO=chrom- osome — SO=protein coding gene — is pseudo=false | 2068 | 0 | 2068 | 2360.49 | 1142 | 1498 | 6116 | 33 | 55.7 | 73.0 |
| F.graminearum | XP_011315624.1 hypothetical p- rotein FGSG 11656 Fusarium gra- minearum PH-1 | 2049 | 0 | 2049 | 1257.28 | 742 | 1121 | 3252 | 123 | 36.2 | 54.7 |
| C.truncatum | XP_036581307.1 fatty acid syn- thase beta subunit dehydratase Colletotrichum truncatum | 2074 | 0 | 2074 | 2330.06 | 1147 | 1490 | 6037 | 40 | 55.9 | 72.6 |
| B.cinerea | XP_024545929.1 hypothetical p- rotein BCIN 01g00450 Botrytis cinerea B05.10 | 2069 | 0 | 2069 | 2420.58 | 1174 | 1520 | 6272 | 34 | 57.2 | 74.1 |
| B.graminis | VCU38897.1 — transcript=BGT962- 24V316 LOCUS150 t1 — gene=BGT9- 6224V316 LOCUS150 — organism=B- lumeria graminis f. sp. tritic- i 96224 — gene product=unspeci- fied product — transcript prod- uct=unspecified product — loca- tion=LR026984:2368020-2374354(- -) — protein length=2096 — seq- uence SO=chromosome — SO=prote- in coding gene — is pseudo=fal- se | 2072 | 0 | 2072 | 2378.21 | 1162 | 1508 | 6162 | 36 | 56.7 | 73.5 |

Table S16: Pairwise alignment info from yeast Fas1 (DEG20010641), cf. Figure S84.

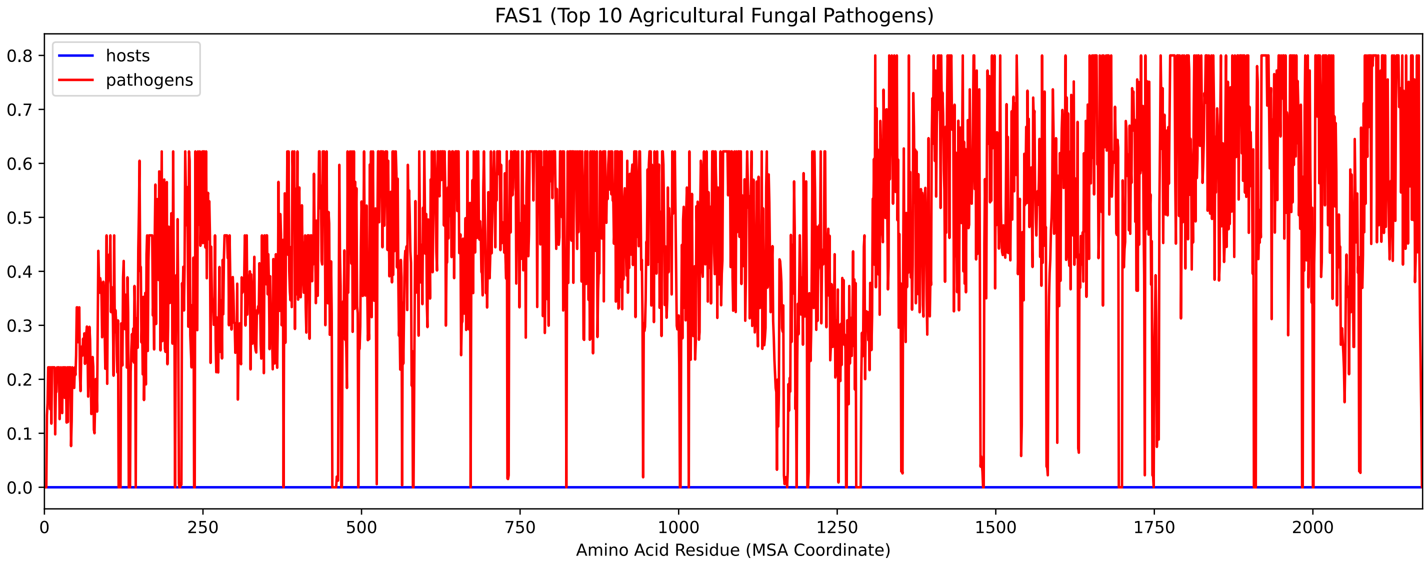


Figure S86: Sneath Similarity of Fas1 for Top 10 Agricultural Fungal Pathogens, cf. Figure S84

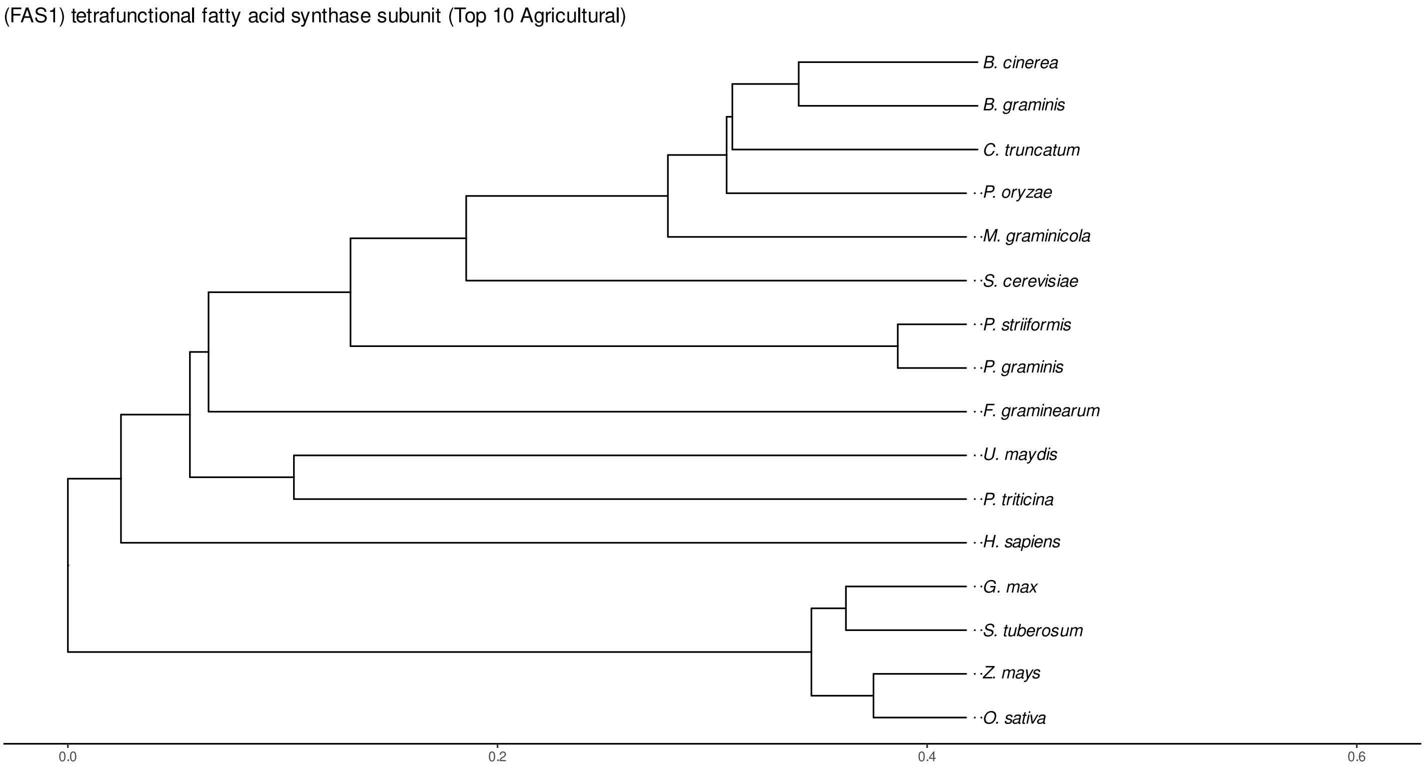


Figure S87: Phylogeny of hits for Fas1 made using clustal omega and visualized using ggtree, cf. Figures S84 and S85.

### S4.5.3 NR

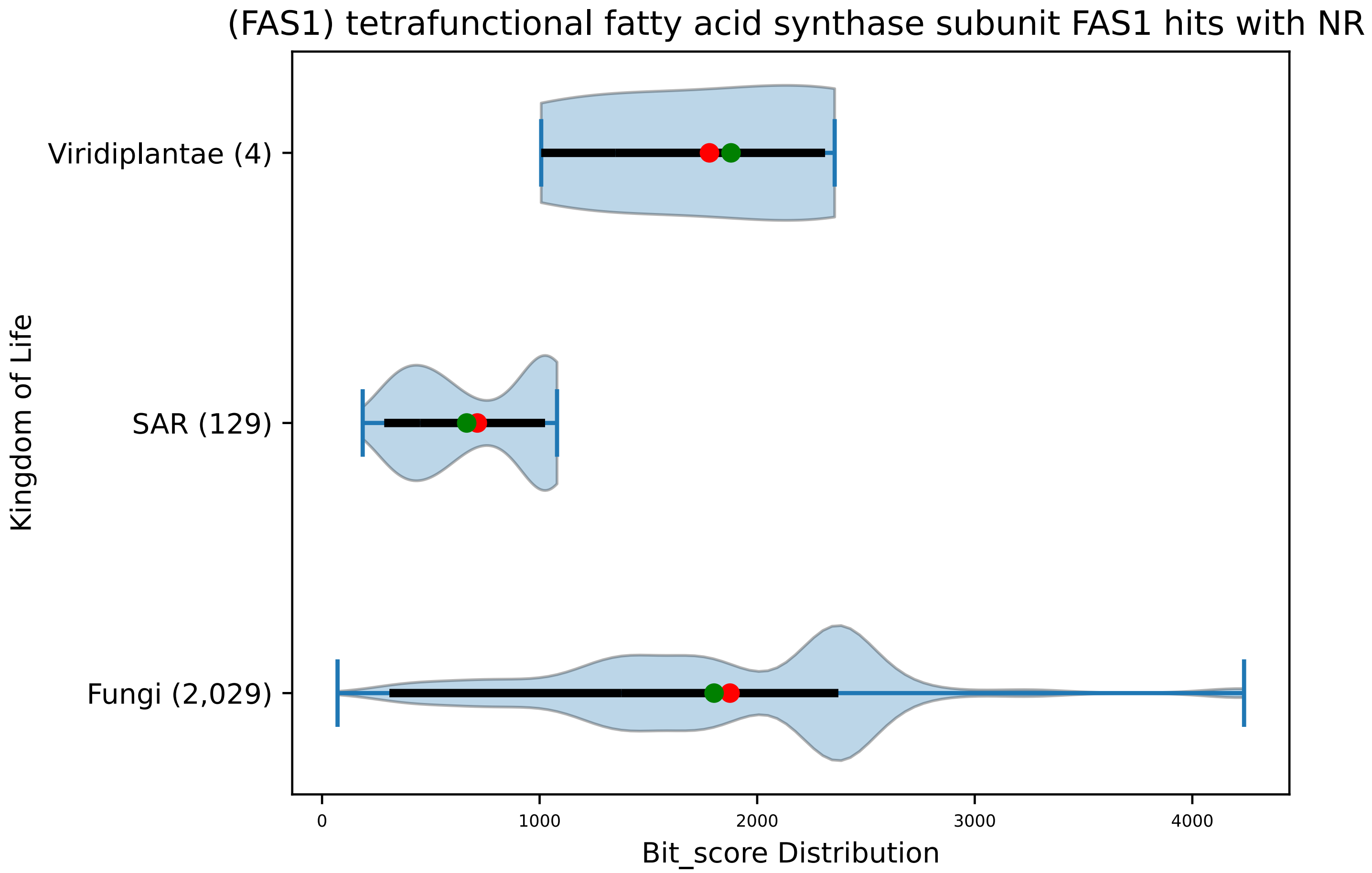


Figure S88: Non-redundant (NR) protein hits for DEG20010641/Fas1, with expectation value of no more than 0.1. Green points are medians, and red points are arithmetic means.

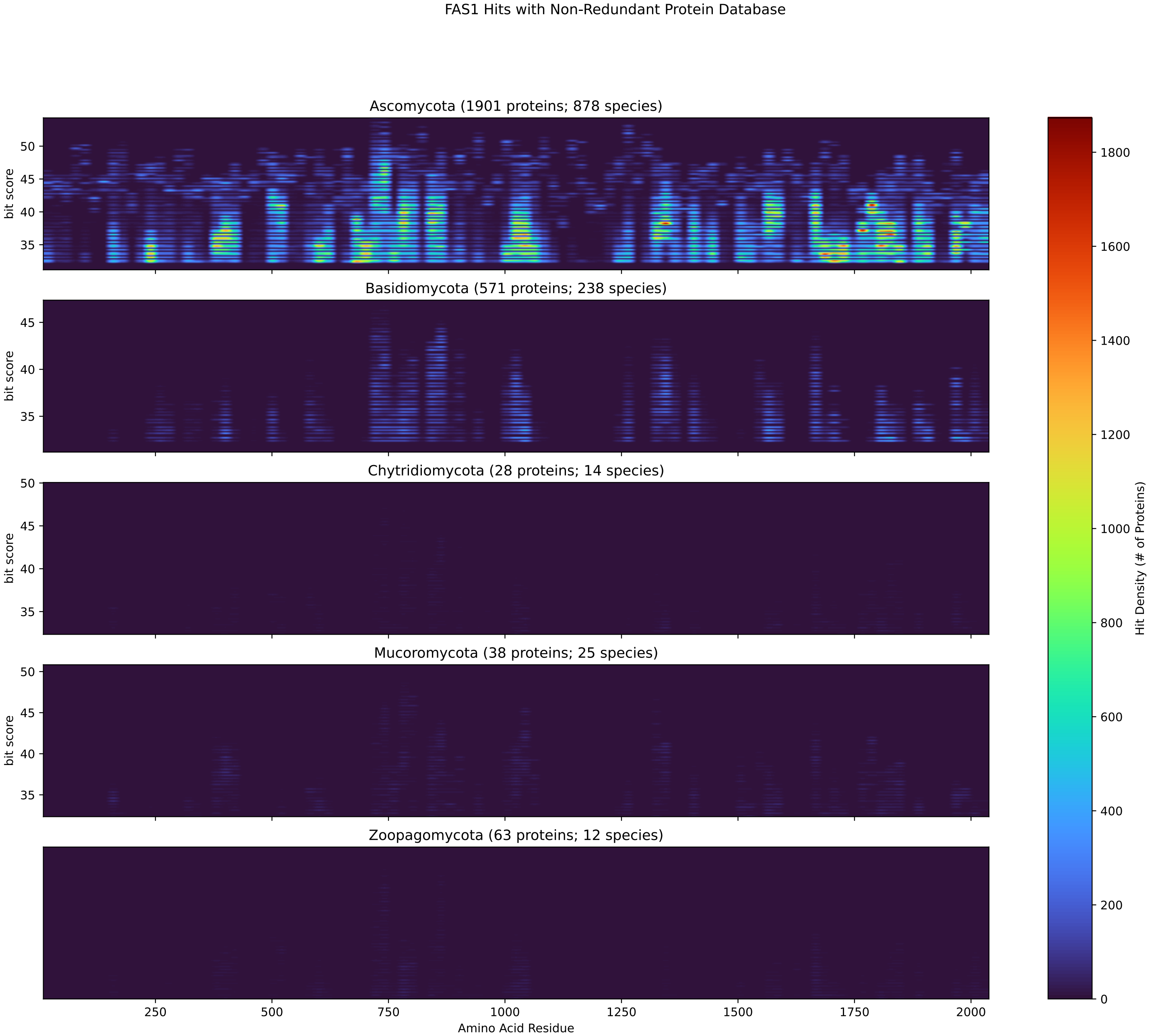


Figure S89: Non-redundant (NR) protein hits for Fas1 in the kingdom Fungi.

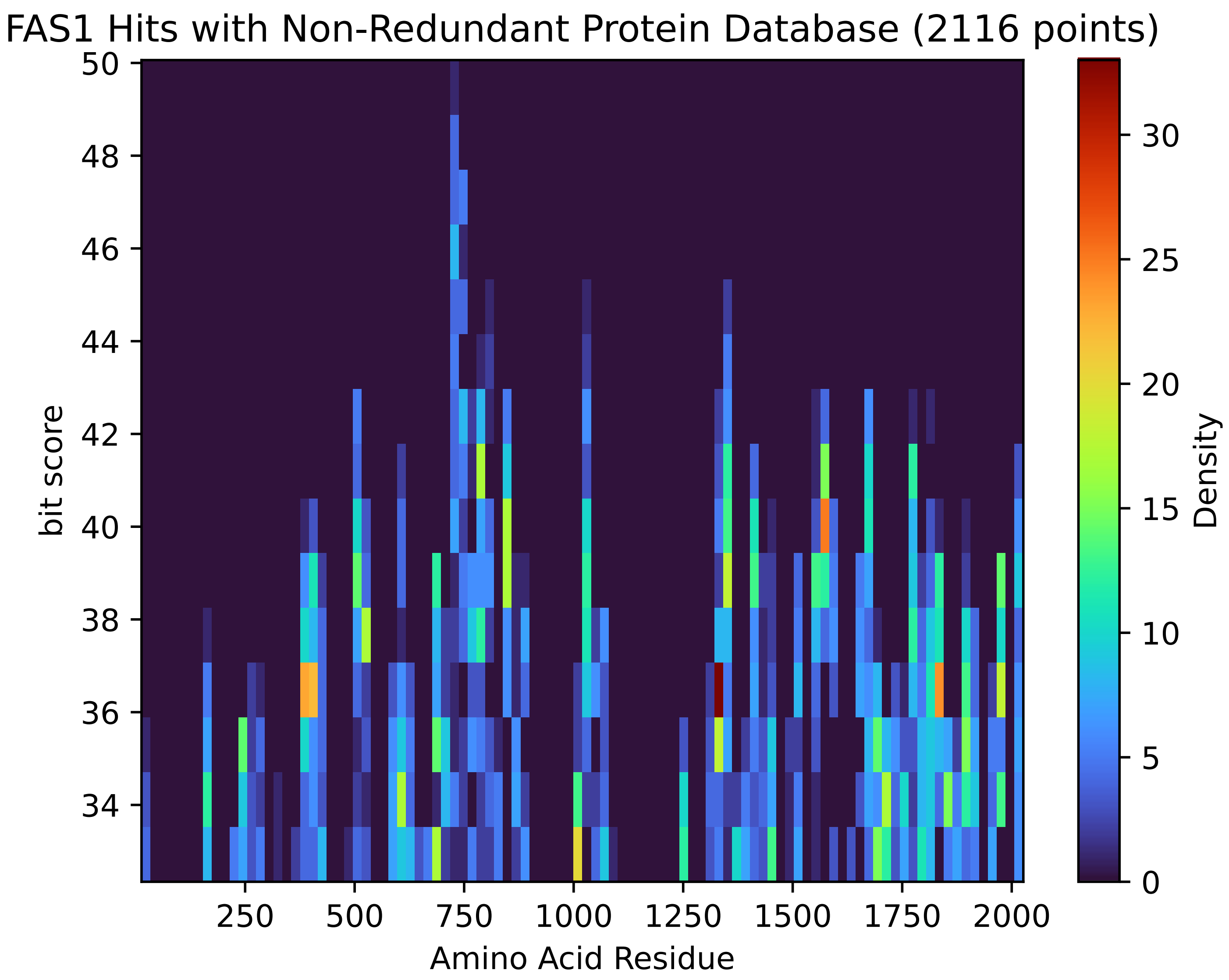


Figure S90: Non-redundant (NR) protein hits for Fas1 in the kingdom Viridiplantae.

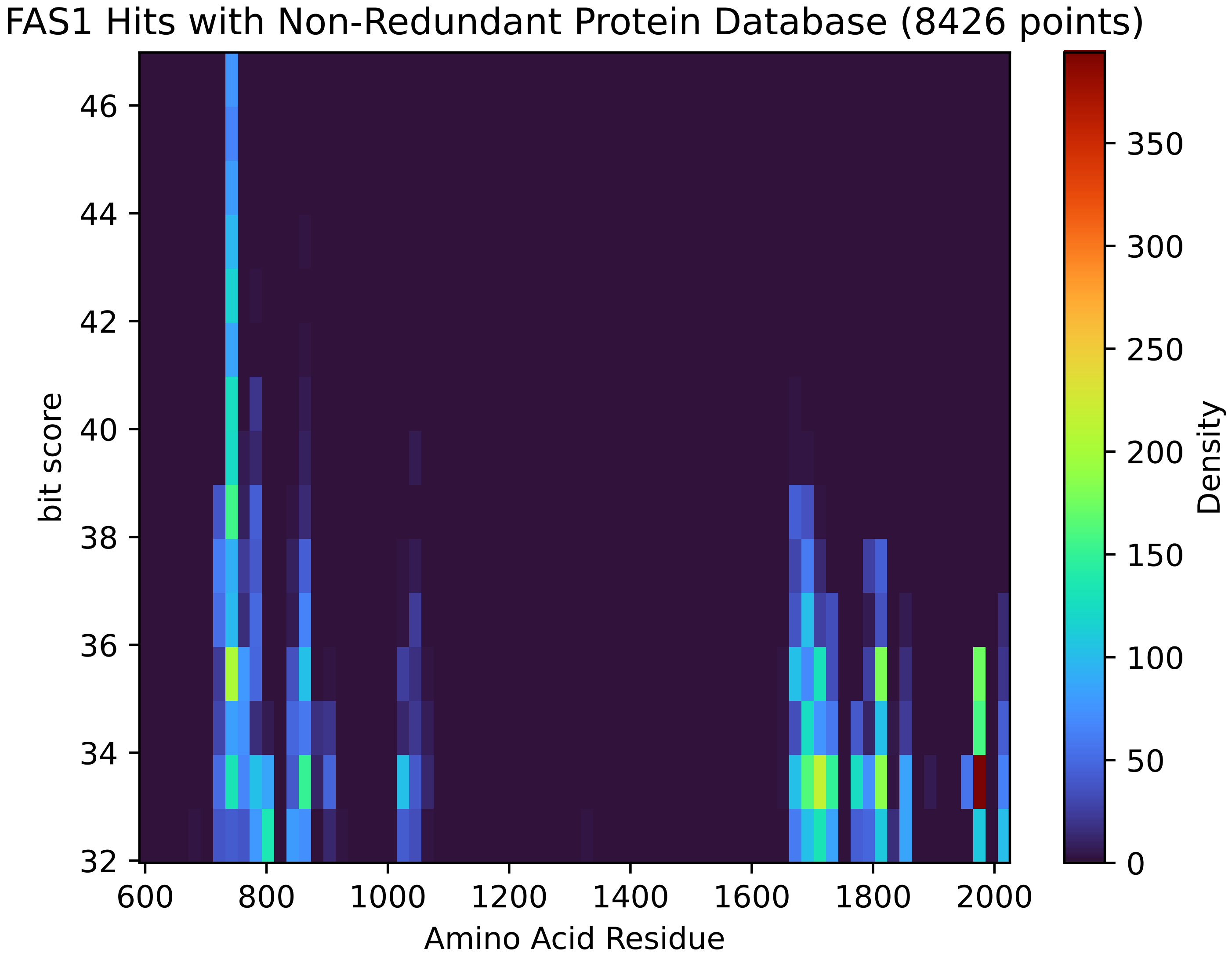


Figure S91: Non-redundant (NR) protein hits for Fas1 in the kingdom SAR.

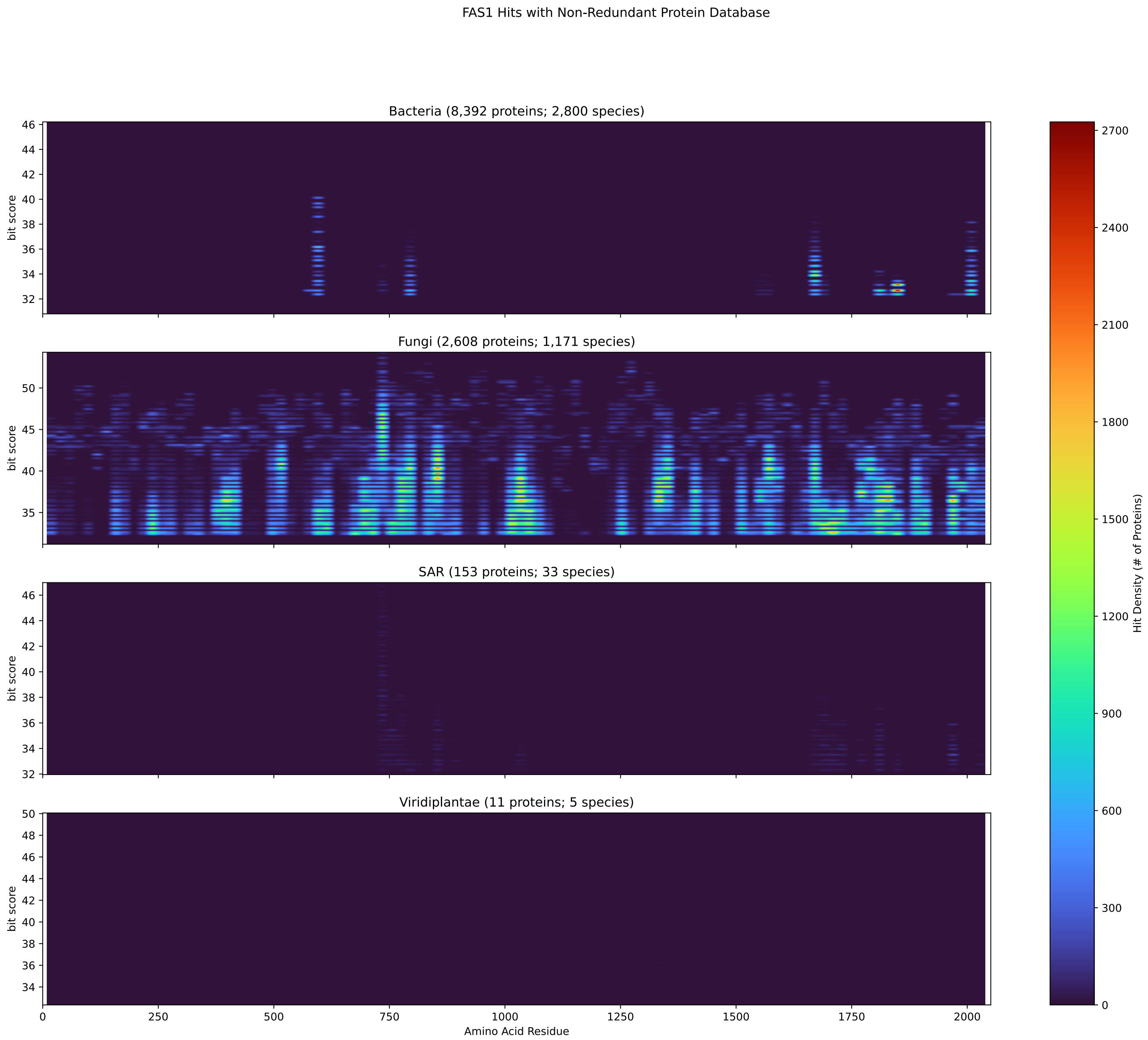


Figure S92: Non-redundant (NR) protein hits for DEG20010641/Fas1 at 20 amino acid length queries.

#### S4.6 Fas2

##### S4.6.1 WHO Critical Pathogens

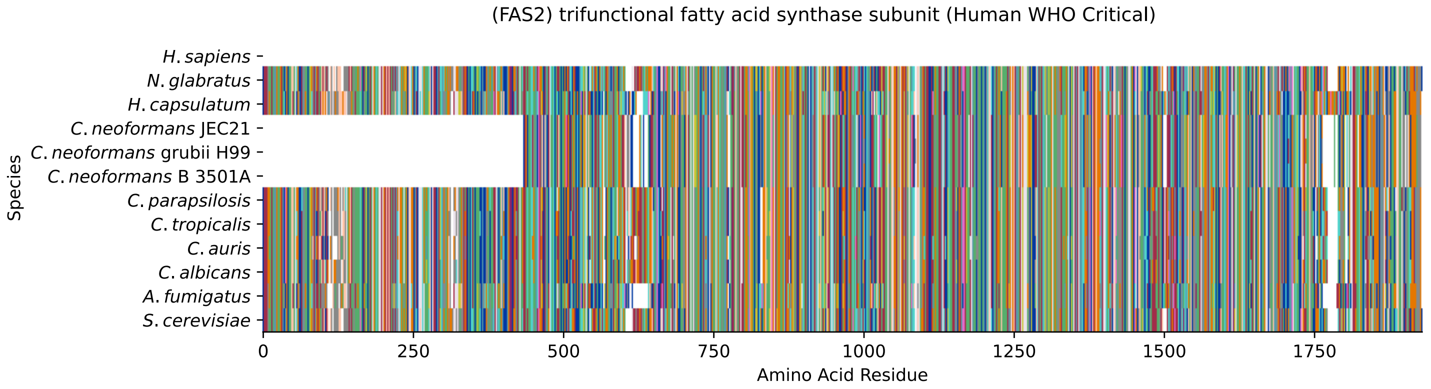


Figure S93: Multiple sequence alignment of yeast Fas2 (WHO Critical Pathogens). Cf. Figure S94 for alignment quality, and Figure S95 for Sneath similarity. Cf. Table S17 for protein names, and pairwise alignment metrics with yeast Fas2.

#### Fas2 MSA Quality

| | *A. f.* | *C. a.* | *C. a.* | *C. p.* | *C. t.* | *C. n.* | *C. n.* | *C. n.* | *H. c.* | *N. g.* | *S. c.* |
|---|---|---|---|---|---|---|---|---|---|---|---|
| *A. fumigatus* | 3842.74 | 2488.38 | 2437.53 | 2421.35 | 2487.22 | 1565.05 | 1565.44 | 1565.05 | 3117.41 | 2419.81 | 2377.44 |
| *C. albicans* | 2484.91 | 3889.73 | 3194.06 | 3346.98 | 3537.66 | 1434.47 | 1436.01 | 1434.47 | 2402.09 | 2714.49 | 2709.48 |
| *C. auris* | 2439.07 | 3210.62 | 3875.48 | 3163.24 | 3219.48 | 1464.9 | 1463.74 | 1464.9 | 2374.36 | 2674.04 | 2682.13 |
| *C. parapsilosis* | 2442.54 | 3374.72 | 3179.81 | 3895.89 | 3382.42 | 1432.54 | 1436.4 | 1432.54 | 2360.1 | 2663.64 | 2672.5 |
| *C. tropicalis* | 2479.52 | 3533.81 | 3200.99 | 3345.83 | 3892.81 | 1441.02 | 1442.17 | 1441.02 | 2407.48 | 2725.27 | 2714.87 |
| *C. neoformans* B 3501A | 1582.77 | 1451.42 | 1476.07 | 1437.17 | 1459.51 | 2954.08 | 2914.41 | 2954.08 | 1488.78 | 1442.17 | 1457.97 |
| *C. neoformans* grubii H99 | 1582.39 | 1452.19 | 1472.6 | 1437.94 | 1459.12 | 2914.41 | 2949.84 | 2914.41 | 1490.32 | 1447.57 | 1455.66 |
| *C. neoformans* JEC21 | 1582.77 | 1451.42 | 1476.07 | 1437.17 | 1459.51 | 2954.08 | 2914.41 | 2954.08 | 1488.78 | 1442.17 | 1457.97 |
| *H. capsulatum* | 3137.05 | 2430.21 | 2394.39 | 2360.1 | 2439.45 | 1504.58 | 1503.8 | 1504.58 | 3944.81 | 2335.84 | 2339.69 |
| *N. glabratus* | 2420.19 | 2728.35 | 2668.65 | 2645.15 | 2743.76 | 1422.14 | 1427.92 | 1422.14 | 2313.88 | 3887.42 | 3219.1 |
| *S. cerevisiae* | 2385.91 | 2712.95 | 2671.73 | 2649.77 | 2722.58 | 1440.25 | 1436.78 | 1440.25 | 2321.2 | 3225.65 | 3890.5 |

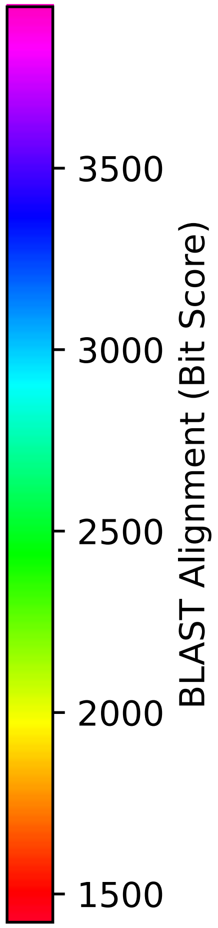


Figure S94: Multiple sequence alignment quality of Fas2 (WHO Critical Pathogens). Cf. Figures S93 and S96.

| Species | Hit Protein | Hit Length (a.a.) | evalue | align_len | bit_score | identity | positive | score | gaps | % identity | % positive |
|---|---|---|---|---|---|---|---|---|---|---|---|
| H.sapiens | - | - | - | - | - | - | - | - | - | - | - |
| N.glabratus | XP_445956.1 uncharacterized p-rotein CAGL0E06138g Nakaseomyc-es glabratus | 1886 | 0 | 1886 | 3224.49 | 1554 | 1723 | 8359 | 5 | 82.4 | 91.3 |
| H.capsulatum | XP_045283904.1 fatty acid syn-thase subunit alpha Histoplasm-a capsulatum G186AR | 1916 | 0 | 1916 | 2321.2 | 1158 | 1460 | 6014 | 48 | 61.4 | 77.4 |
| C.neoformans.JEC21 | XP_571099.1 fatty-acid syntha-se complex protein, putative C-ryptococcus neoformans var. ne-oformans JEC21 | 1469 | 0 | 1469 | 1441.02 | 736 | 989 | 3729 | 55 | 39.0 | 52.4 |
| C.neoformans.grubii.H99 | XP_012049943.1 fatty acid syn-thase subunit alpha, fungi typ-e Cryptococcus neoformans var. grubii H99 | 1463 | 0 | 1463 | 1436.78 | 736 | 985 | 3718 | 43 | 39.0 | 52.2 |
| C.neoformans.B.3501A | XP_775163.1 hypothetical prot-ein CNBE4360 Cryptococcus neof-ormans var. neoformans B-3501A | 1469 | 0 | 1469 | 1441.02 | 736 | 989 | 3729 | 55 | 39.0 | 52.4 |
| C.parapsilosis | XP_036665363.1 uncharacterize-d protein CPAR2 807400 Candida parapsilosis | 1900 | 0 | 1900 | 2651.31 | 1294 | 1565 | 6871 | 34 | 68.6 | 82.9 |
| C.tropicalis | XP_002548204.1 fatty acid syn-thase alpha subunit Candida tr-opicalis MYA-3404 | 1899 | 0 | 1899 | 2721.42 | 1319 | 1584 | 7053 | 33 | 69.9 | 83.9 |
| C.auris | XP_028891558.2 fatty acid syn-thase subunit alpha Candida au-ris | 1898 | 0 | 1898 | 2671.73 | 1311 | 1567 | 6924 | 38 | 69.5 | 83.0 |
| C.albicans | XP_723161.2 trifunctional fat-ty acid synthase subunit Candi-da albicans SC5314 | 1900 | 0 | 1900 | 2712.95 | 1313 | 1588 | 7031 | 34 | 69.6 | 84.2 |
| A.fumigatus | XP_748738.1 fatty acid syntha-se alpha subunit FasA Aspergil-lus fumigatus Af293 | 1894 | 0 | 1894 | 2387.07 | 1167 | 1470 | 6185 | 49 | 61.8 | 77.9 |

Table S17: Pairwise alignment info from yeast Fas2 (DEG20011054), cf. Figure S93.

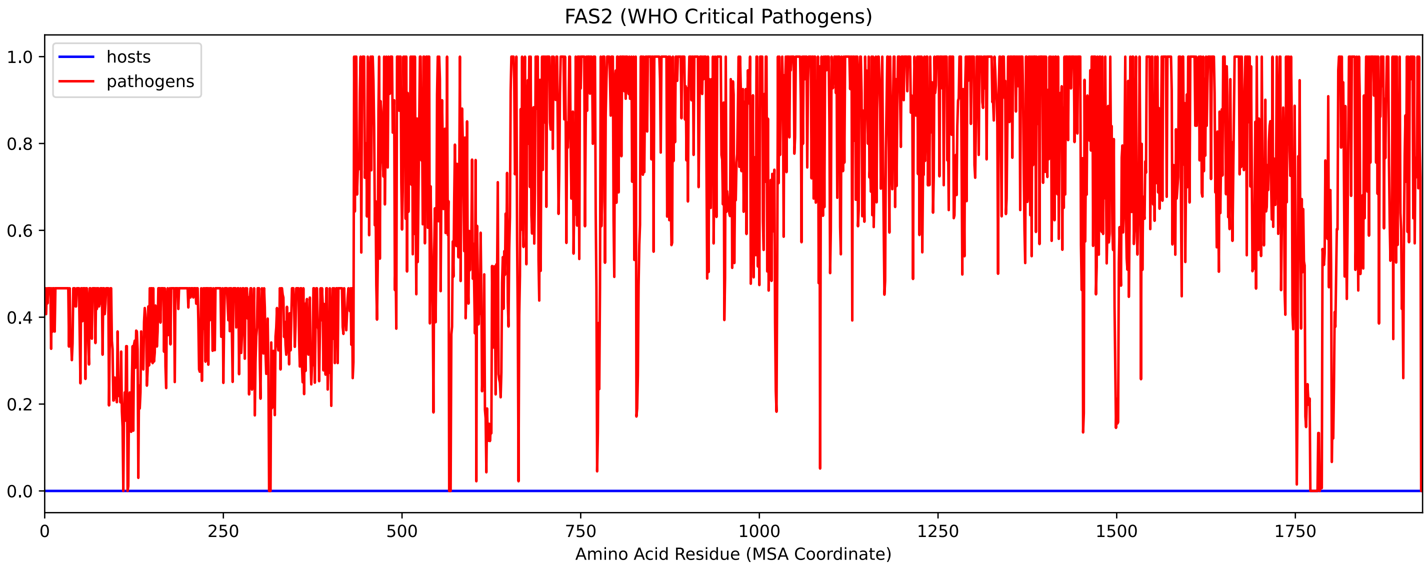


Figure S95: Sneath Similarity of Fas2 for WHO Critical Pathogens, cf. Figure S93

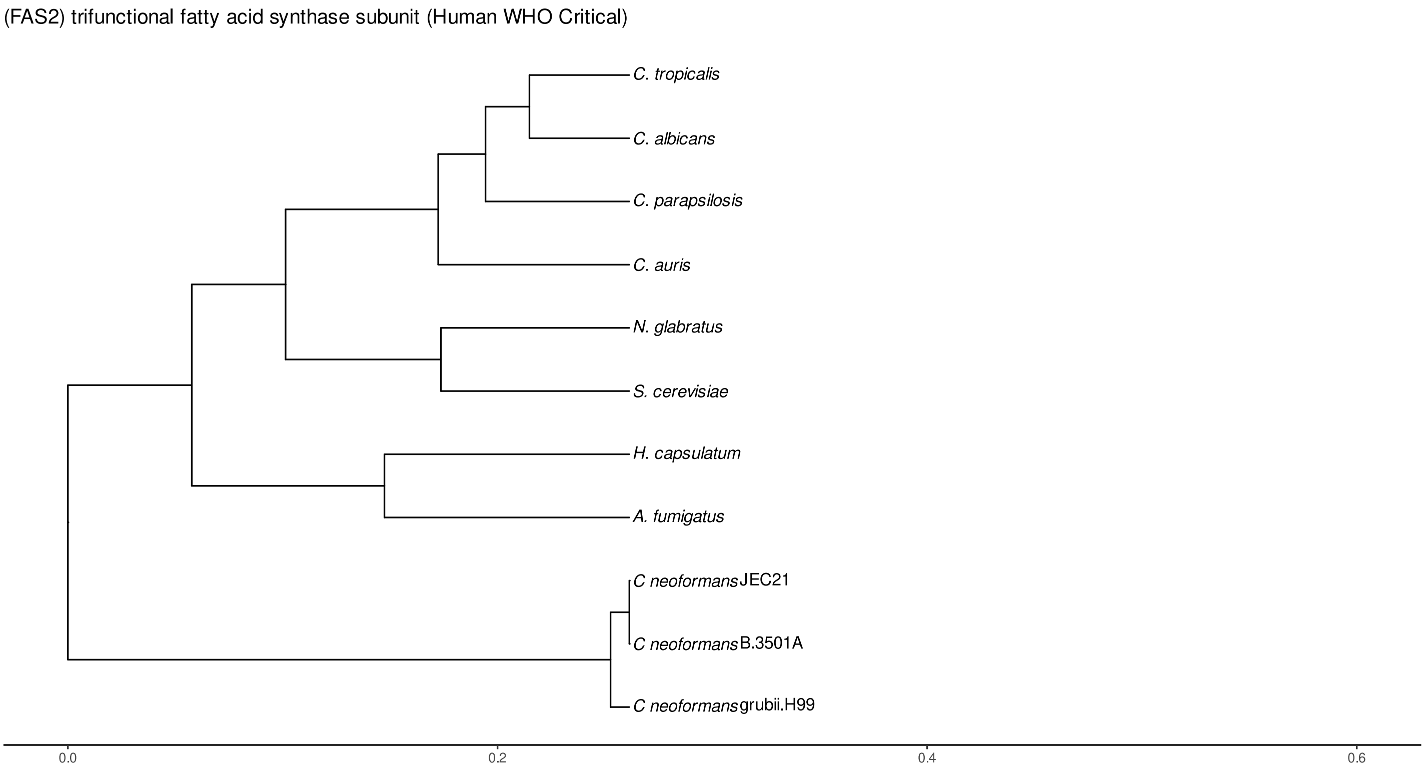


Figure S96: Phylogeny of hits for Fas2 made using clustal omega and visualized using ggtree, cf. Figures S93 and S94.

##### S4.6.2 Top 10 Agricultural Fungal Pathogens

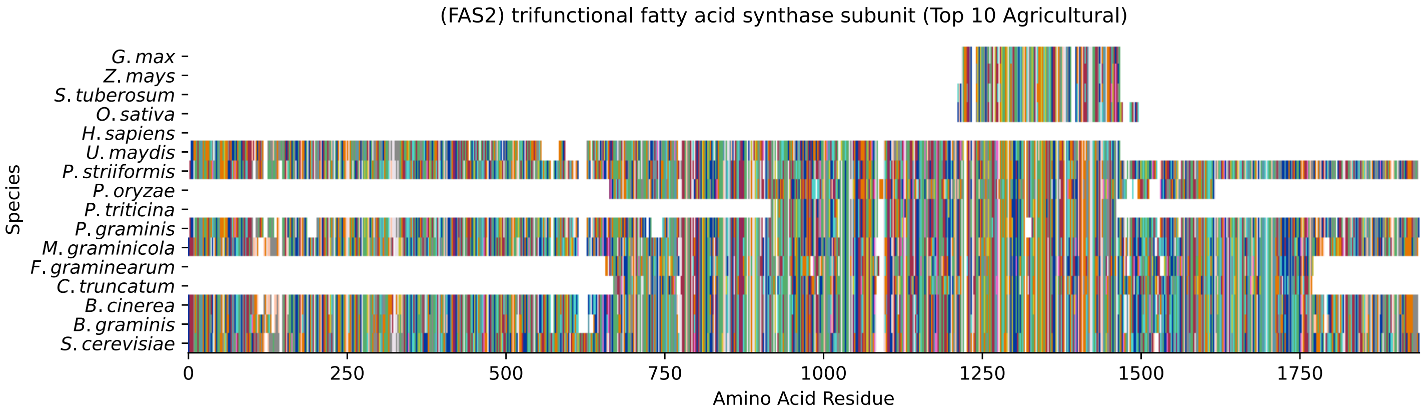


Figure S97: Multiple sequence alignment of yeast Fas2 (Top 10 Agricultural Fungal Pathogens). Cf. Figure S98 for alignment quality, and Figure S99 for Sneath similarity. Cf. Table S18 for protein names, and pairwise alignment metrics with yeast Fas2.

#### Fas2 MSA Quality

| | *B. g.* | *B. c.* | *C. t.* | *F. g.* | *M. g.* | *P. g.* | *P. t.* | *P. o.* | *P. s.* | *U. m.* | *S. c.* |
|---|---|---|---|---|---|---|---|---|---|---|---|
| *B. graminis* | 3860.07 | 3311.16 | 1193.33 | 1192.18 | 3006.08 | 1768.05 | 615.92 | 863.603 | 1937.15 | 1090.1 | 0 |
| *B. cinerea* | 3341.21 | 3853.52 | 1193.33 | 1191.79 | 3124.72 | 1769.59 | 613.224 | 878.241 | 1947.55 | 1067.37 | 0 |
| *C. truncatum* | 1216.45 | 1215.29 | 2209.88 | 1026.16 | 1218.37 | 903.664 | 483.411 | 800.431 | 983.4 | 703.36 | 0 |
| *F. graminearum* | 1202.19 | 1201.04 | 1010.75 | 2237.61 | 1181.01 | 895.575 | 495.352 | 769.615 | 980.704 | 687.182 | 0 |
| *M. graminicola* | 3057.31 | 3151.69 | 1206.43 | 1177.93 | 3860.07 | 1794.63 | 613.224 | 872.848 | 1964.5 | 1070.07 | 0 |
| *P. graminis* | 1794.63 | 1776.91 | 883.634 | 887.486 | 1793.09 | 3749.52 | 988.408 | 650.203 | 3353.53 | 1013.45 | 0 |
| *P. triticina* | 615.92 | 613.609 | 483.026 | 495.738 | 613.224 | 988.023 | 1081.24 | 392.504 | 1030.01 | 554.288 | 0 |
| *P. oryzae* | 887.1 | 900.968 | 801.971 | 788.875 | 895.96 | 670.233 | 412.535 | 1855.11 | 761.911 | 639.802 | 0 |
| *P. striiformis* | 1976.44 | 1962.96 | 987.252 | 993.03 | 1970.67 | 3369.33 | 1030.01 | 758.444 | 3826.56 | 1149.42 | 0 |
| *U. maydis* | 1109.75 | 1074.31 | 684.871 | 687.567 | 1074.69 | 1019.99 | 555.058 | 619.387 | 1129.78 | 2762.64 | 0 |
| *S. cerevisiae* | 0 | 0 | 0 | 0 | 0 | 0 | 0 | 0 | 0 | 0 | 0 |

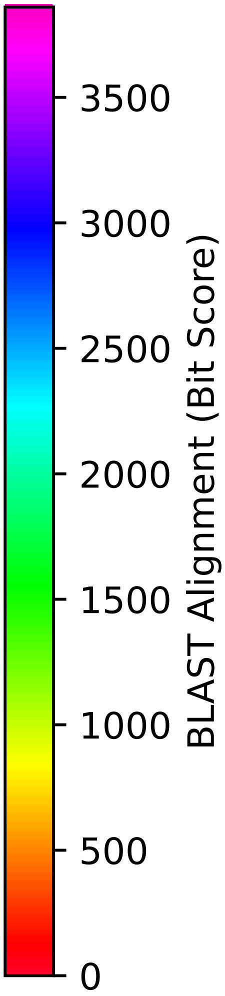


Figure S98: Multiple sequence alignment quality of Fas2 (Top 10 Agricultural Fungal Pathogens). Cf. Figures S97 and S100.

| Species | Hit Protein | Hit Length (a.a.) | evalue | align_len | bit_score | identity | positive | score | gaps | % identity | % positive |
|---|---|---|---|---|---|---|---|---|---|---|---|
| G.max | NP_001236069.1 plastid 3-keto--acyl-ACP synthase II-B precursor Glycine max | 255 | 1.8e-06 | 255 | 53.9138 | 67 | 111 | 128 | 34 | 3.6 | 5.9 |
| Z.mays | NP_001169617.1 3-oxoacyl-acyl--carrier-protein synthase II, chloroplastic-like precursor Zea mays | 255 | 1e-05 | 255 | 51.2174 | 61 | 106 | 121 | 34 | 3.2 | 5.6 |
| S.tuberosum | XP_006345248.1 PREDICTED: 3-oxoacyl-acyl-carrier-protein synthase II, chloroplastic-like Solanum tuberosum | 254 | 4.2e-07 | 254 | 55.0694 | 65 | 112 | 131 | 22 | 3.4 | 5.9 |
| O.sativa | XP_015630630.1 3-oxoacyl-acyl--carrier-protein synthase II, chloroplastic-like Oryza sativa Japonica Group | 278 | 1.1e-05 | 278 | 50.8322 | 63 | 112 | 120 | 34 | 3.3 | 5.9 |
| H.sapiens | - | - | - | - | - | - | - | - | - | - | - |
| U.maydis | XP_011392728.1 fatty acid synthase FAS2 Ustilago maydis 521 | 1432 | 0 | 1432 | 1070.07 | 601 | 851 | 2766 | 91 | 31.8 | 45.1 |
| P.striiformis | XP_047808761.1 hypothetical protein Pst134EA 009828 Puccinia striiformis f. sp. tritici | 1894 | 0 | 1894 | 1812.74 | 947 | 1279 | 4694 | 54 | 50.2 | 67.8 |
| P.oryzae | mRNA M BR32 EuGene 00106401-p1 — transcript=mRNA M BR32 EuGene 00106401 — gene=M BR32 EuGene 00106401 — organism=Pyricularia oryzae BR32 — gene product=unspecified product — transcript product=unspecified product — location=BR32 scaffold000-14:1001674-1005232(-) — protein length=1167 — sequence SO=supercontig — SO=protein coding gene — is pseudo=false | 937 | 0 | 937 | 833.558 | 426 | 617 | 2152 | 50 | 22.6 | 32.7 |
| P.triticina | XP_053019002.1 uncharacterized protein PtA15 3A818 Puccinia triticina | 524 | 0 | 524 | 601.282 | 289 | 388 | 1549 | 7 | 15.3 | 20.6 |
| P.graminis | XP_003325251.2 fatty acid synthase subunit beta Puccinia graminis f. sp. tritici CRL 75-3-6-700-3 | 1899 | 0 | 1899 | 1659.43 | 902 | 1225 | 4296 | 105 | 47.8 | 64.9 |
| M.graminicola | ZTRI 3.47.mRNA-p1 — transcript=ZTRI 3.47.mRNA — gene=ZTRI 3.-47 — organism=Zymoseptoria tritici IPO323 — gene product=similar to fatty acid synthase subunit alpha — transcript product=similar to fatty acid synthase subunit alpha — location=Ztri chr 3:156668-162355(-) — protein length=1859 — sequence SO=chromosome — SO=protein coding gene — is pseudo=false | 1902 | 0 | 1902 | 2352.01 | 1149 | 1459 | 6094 | 63 | 60.9 | 77.3 |
| F.graminearum | XP_011315622.1 hypothetical protein FGSG 00036 Fusarium graminearum PH-1 | 1091 | 0 | 1091 | 1127.46 | 548 | 761 | 2915 | 22 | 29.0 | 40.3 |
| C.truncatum | XP_036575084.1 fatty acid synthase subunit alpha reductase Colletotrichum truncatum | 1080 | 0 | 1080 | 1102.04 | 540 | 762 | 2849 | 25 | 28.6 | 40.4 |
| B.cinerea | XP_024545928.1 Bcfas2 Botrytis cinerea B05.10 | 1899 | 0 | 1899 | 2455.63 | 1188 | 1483 | 6363 | 56 | 63.0 | 78.6 |
| B.graminis | VCU38898.1 — transcript=BGT962-24V316 LOCUS151 t1 — gene=BGT9-6224V316 LOCUS151 — organism=Blumeria graminis f. sp. tritici 96224 — gene product=unspecified product — transcript product=unspecified product — location=LR026984:2375831-2381518(+) — protein length=1864 — sequence SO=chromosome — SO=protein coding gene — is pseudo=false | 1904 | 0 | 1904 | 2449.86 | 1186 | 1482 | 6348 | 60 | 62.9 | 78.5 |

Table S18: Pairwise alignment info from yeast Fas2 (DEG20011054), cf. Figure S97.

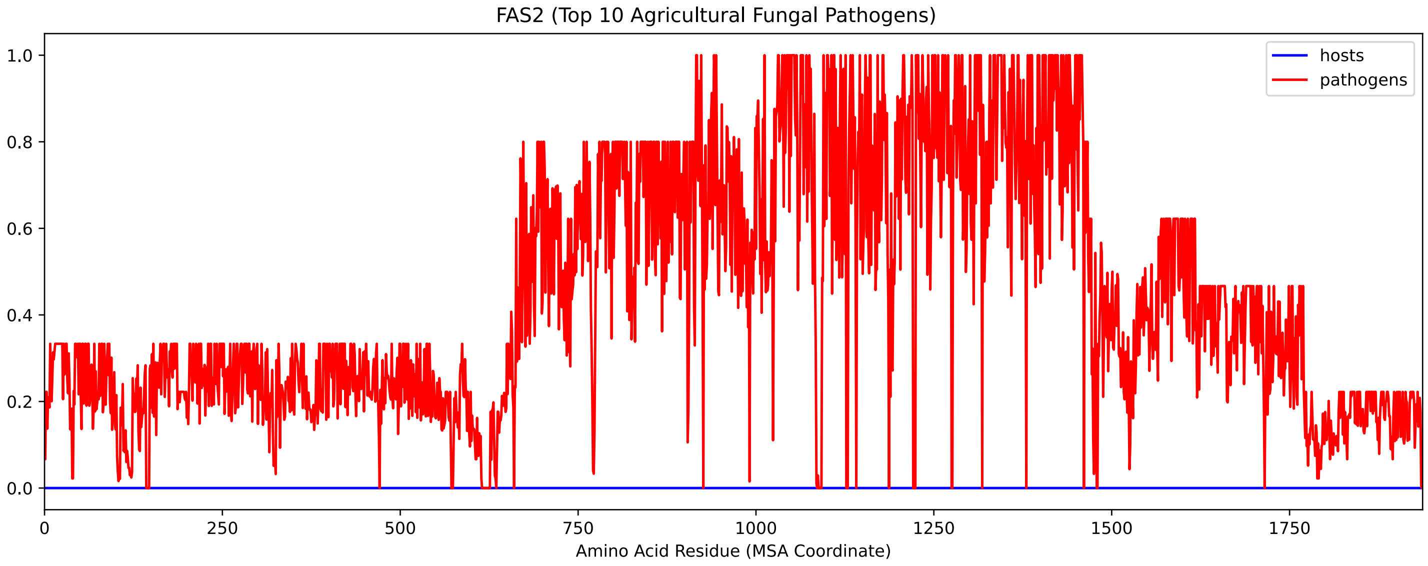


Figure S99: Sneath Similarity of Fas2 for Top 10 Agricultural Fungal Pathogens, cf. Figure S97

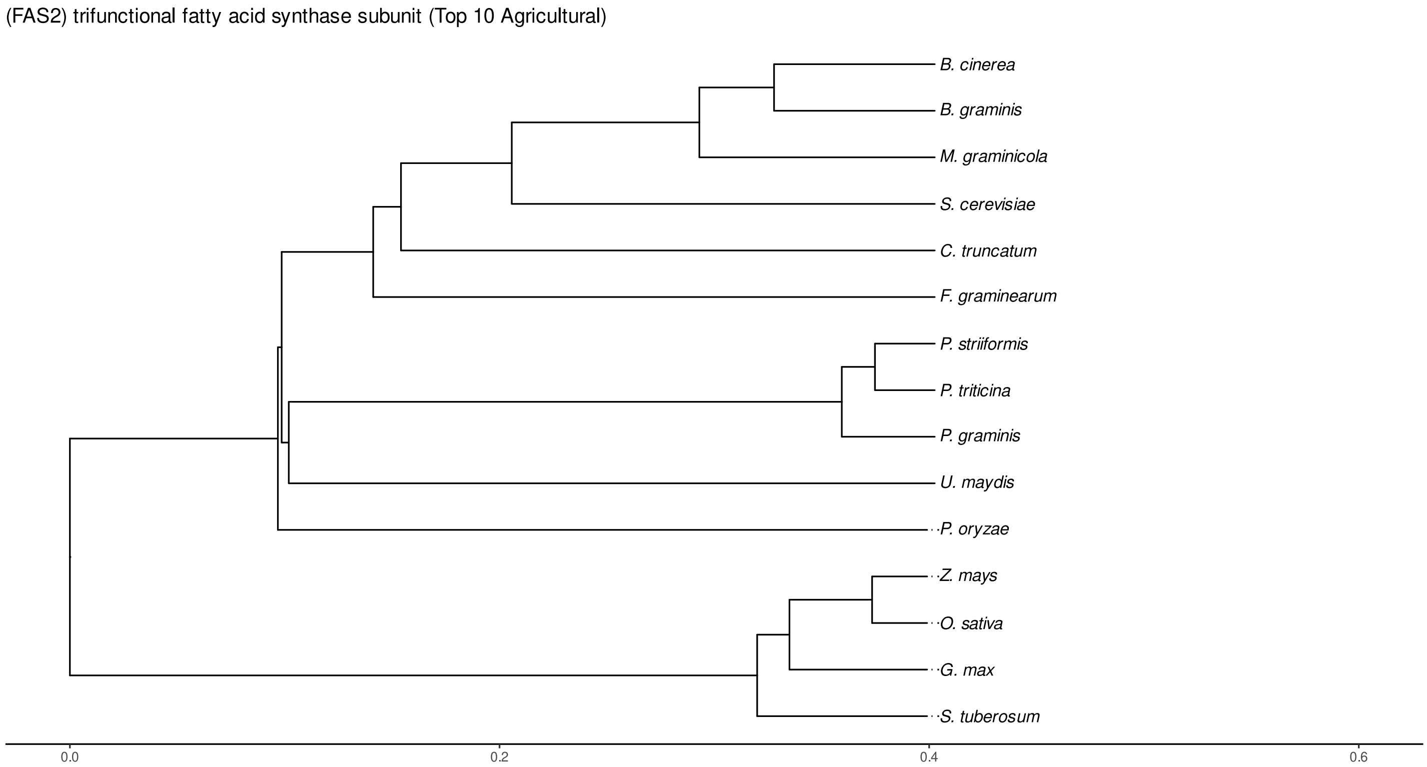


Figure S100: Phylogeny of hits for Fas2 made using clustal omega and visualized using ggtree, cf. Figures S97 and S98.

### S4.6.3 NR

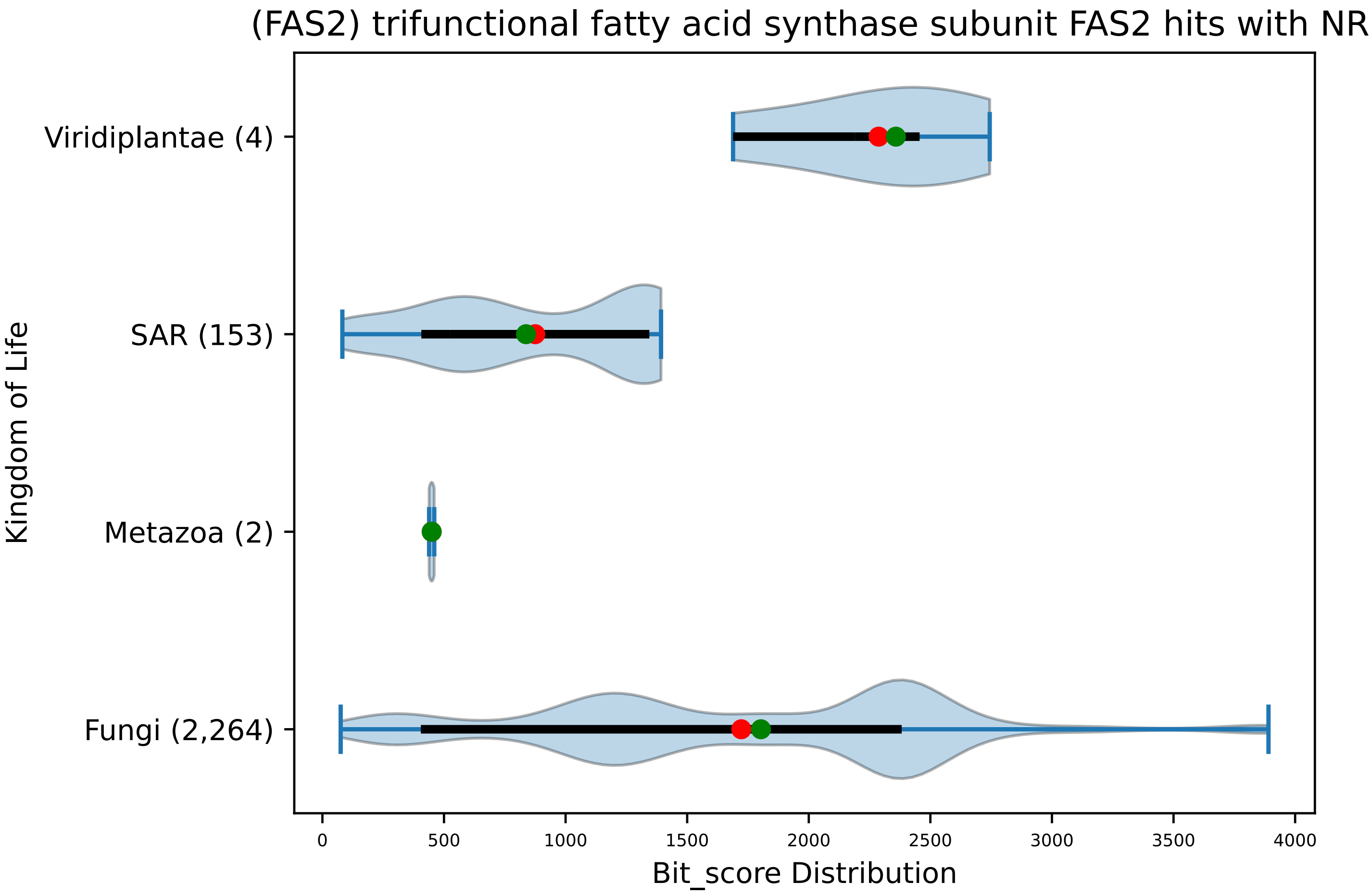


Figure S101: Non-redundant (NR) protein hits for DEG20011054/Fas2, with expectation value of no more than 0.1. Green points are medians, and red points are arithmetic means.

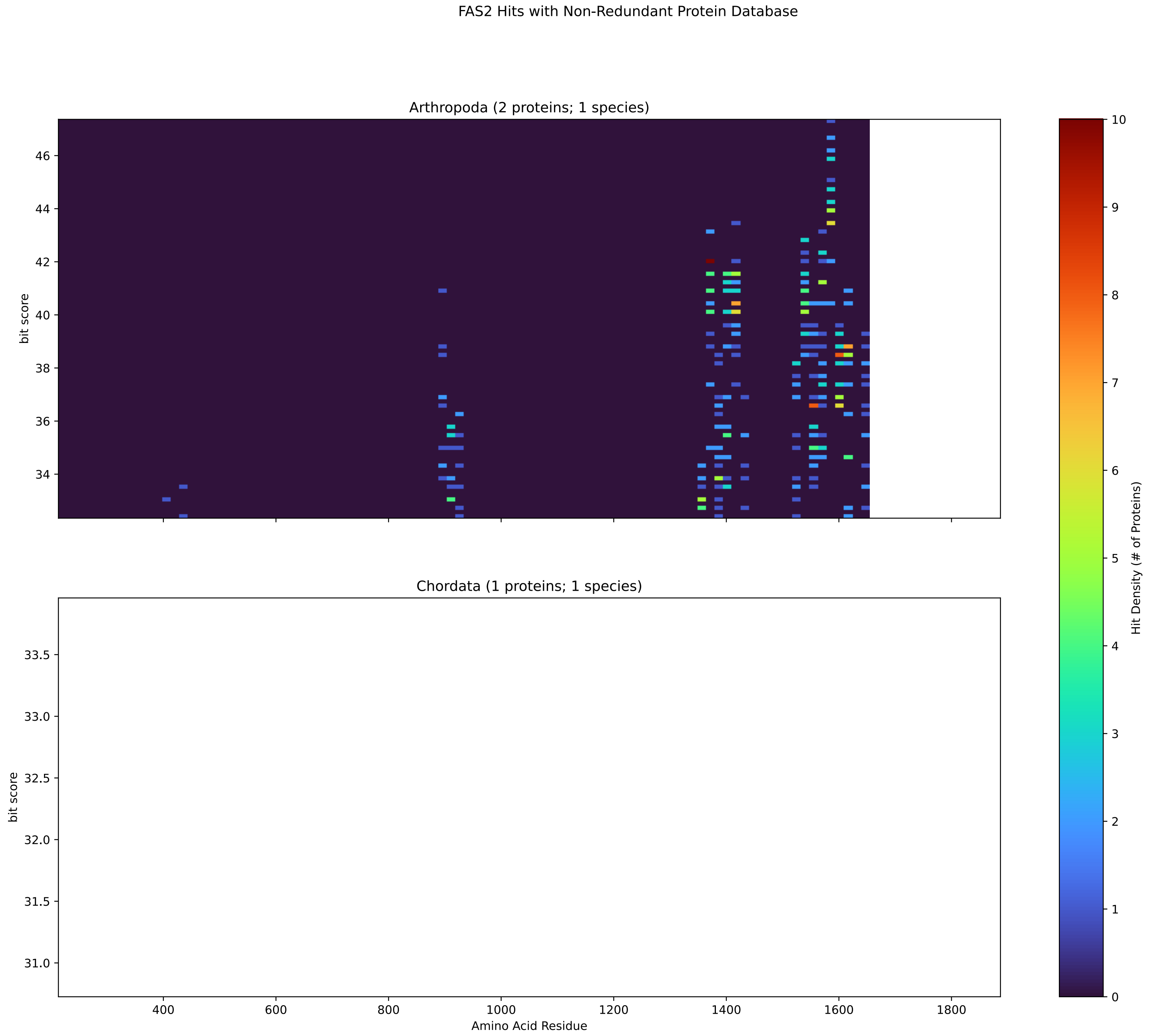


Figure S102: Non-redundant (NR) protein hits for Fas2 in the kingdom Metazoa.

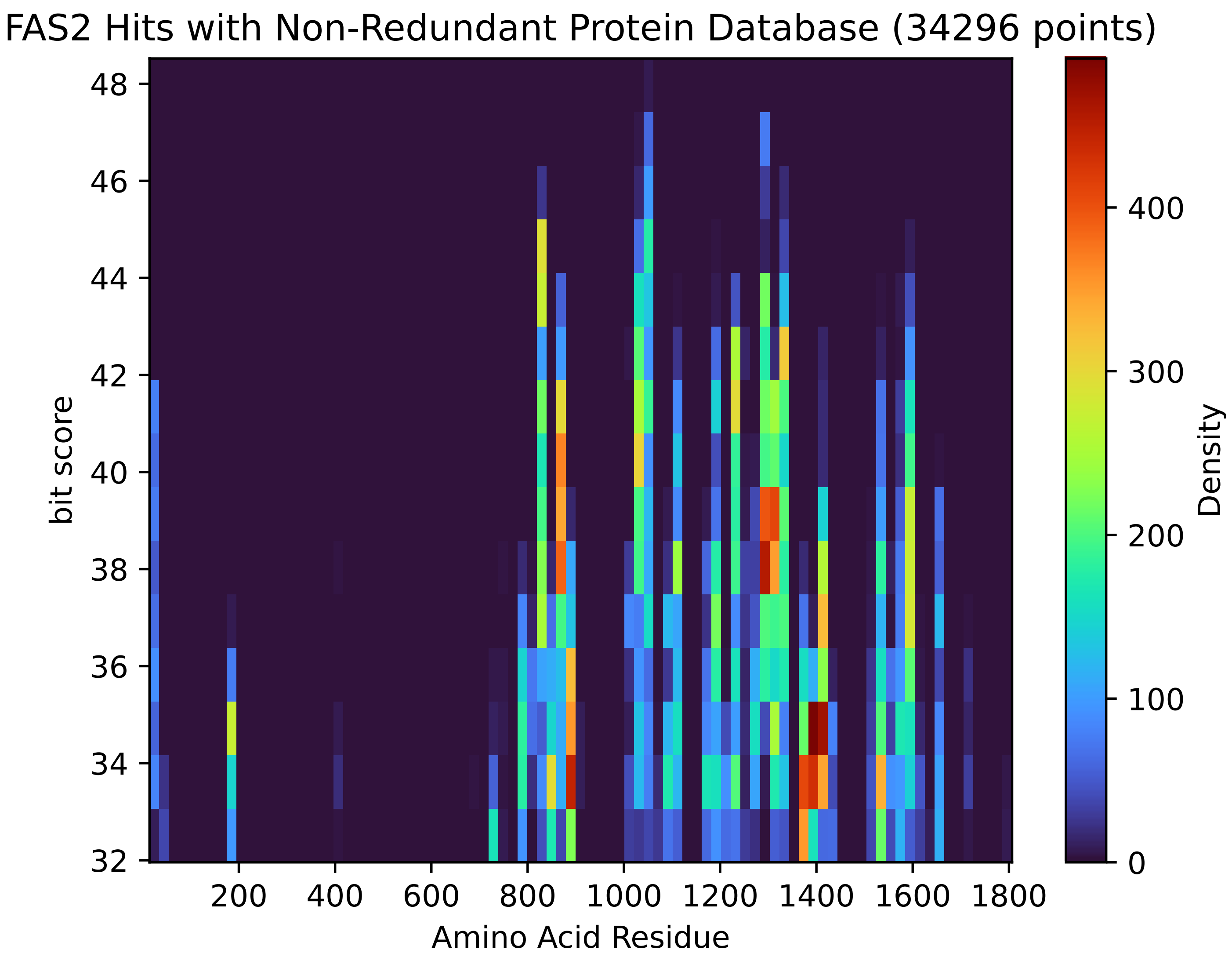


Figure S103: Non-redundant (NR) protein hits for Fas2 in the kingdom SAR.

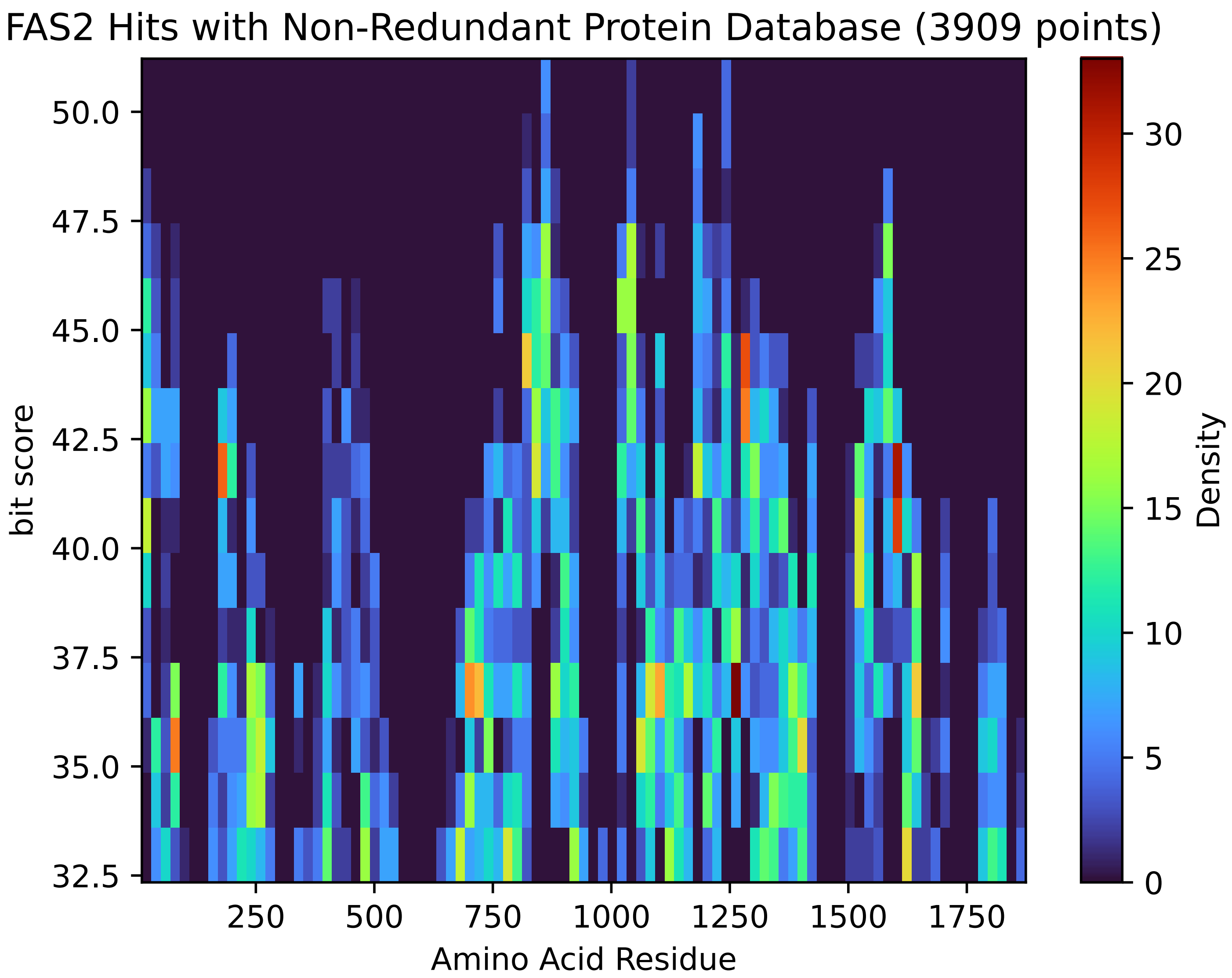


Figure S104: Non-redundant (NR) protein hits for Fas2 in the kingdom Viridiplantae.

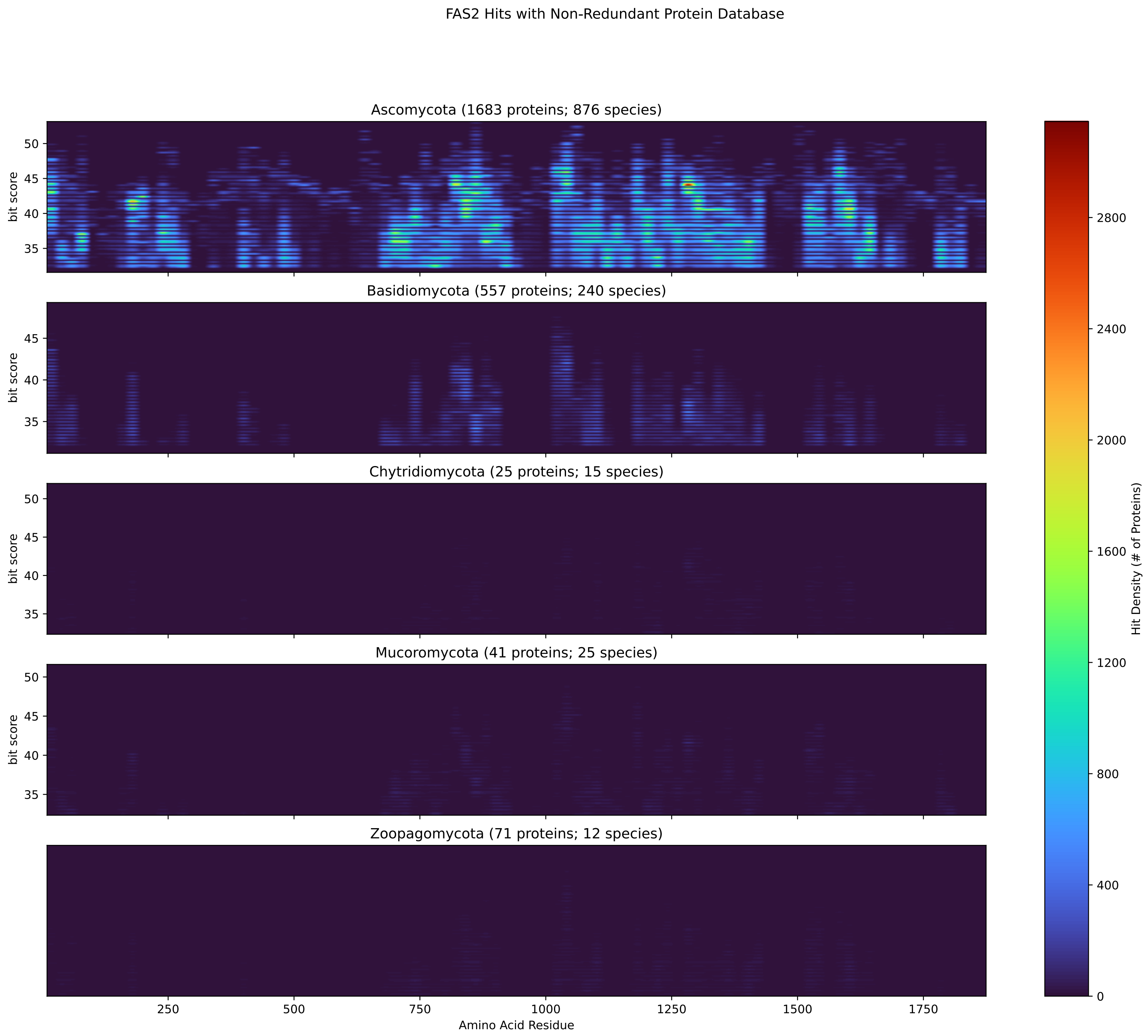


Figure S105: Non-redundant (NR) protein hits for Fas2 in the kingdom Fungi.

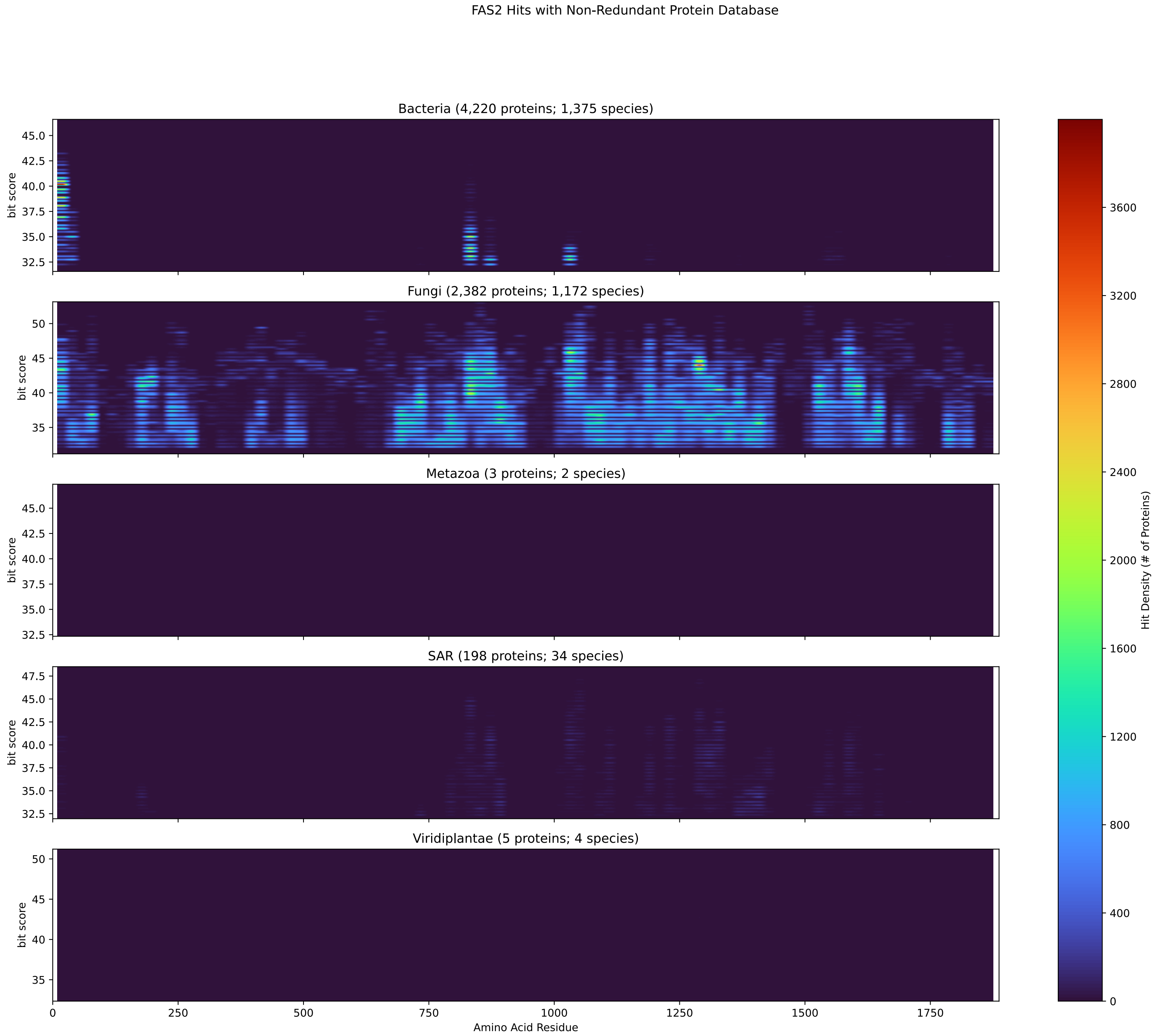


Figure S106: Non-redundant (NR) protein hits for DEG20011054/Fas2 at 20 amino acid length queries.

#### S4.7 Fba1

##### S4.7.1 WHO Critical Pathogens

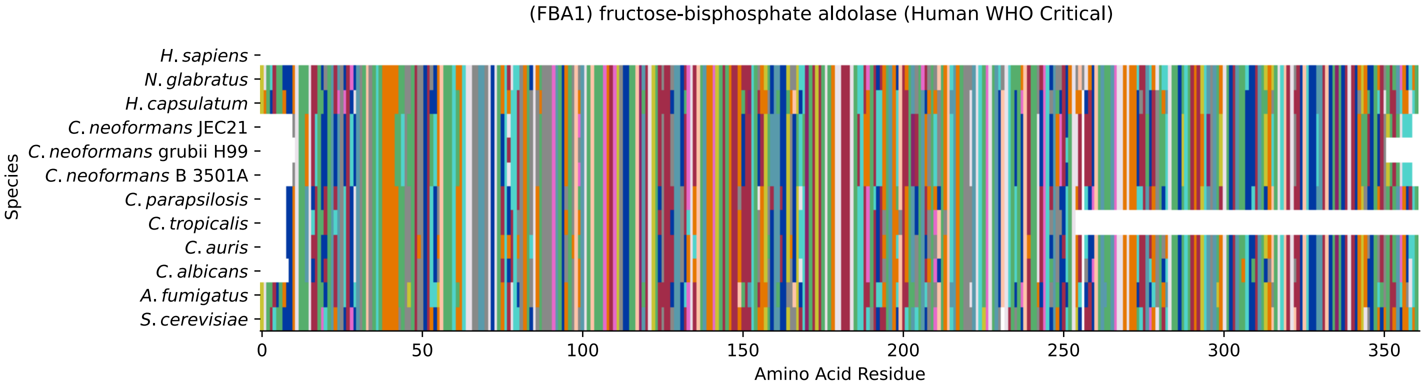


Figure S107: Multiple sequence alignment of yeast Fba1 (WHO Critical Pathogens). Cf. Figure S108 for alignment quality, and Figure S109 for Sneath similarity. Cf. Table S19 for protein names, and pairwise alignment metrics with yeast Fba1.

### Fba1 MSA Quality

| | A. f. | C. a. | C. a. | C. p. | C. t. | C. n. | C. n. | C. n. | H. c. | N. g. | S. c. |
|---|---|---|---|---|---|---|---|---|---|---|---|
| A. fumigatus | 746.503 | 515.383 | 555.444 | 512.686 | 363.999 | 498.819 | 502.286 | 498.819 | 628.246 | 557.37 | 530.02 |
| C. albicans | 516.153 | 728.013 | 650.588 | 642.499 | 450.284 | 481.1 | 482.256 | 481.1 | 506.908 | 590.497 | 548.51 |
| C. auris | 536.569 | 631.328 | 733.791 | 624.78 | 427.172 | 502.671 | 505.753 | 502.671 | 526.939 | 609.372 | 572.007 |
| C. parapsilosis | 512.686 | 641.343 | 643.654 | 730.324 | 434.106 | 493.812 | 494.967 | 493.812 | 512.301 | 579.326 | 552.747 |
| C. tropicalis | 365.54 | 451.44 | 446.047 | 435.647 | 508.449 | 348.591 | 351.673 | 348.591 | 357.451 | 413.69 | 394.045 |
| C. neoformans B 3501A | 499.204 | 480.33 | 519.235 | 493.812 | 347.051 | 719.153 | 687.952 | 719.153 | 504.212 | 520.39 | 508.834 |
| C. neoformans grubii H99 | 502.671 | 481.485 | 522.316 | 494.197 | 350.132 | 687.952 | 701.434 | 687.952 | 499.975 | 523.472 | 514.227 |
| C. neoformans JEC21 | 499.204 | 480.33 | 519.235 | 493.812 | 347.051 | 719.153 | 687.952 | 719.153 | 504.212 | 520.39 | 508.834 |
| H. capsulatum | 629.017 | 506.908 | 542.347 | 512.686 | 356.681 | 504.982 | 500.745 | 504.982 | 742.265 | 543.117 | 526.168 |
| N. glabratus | 540.036 | 572.392 | 609.372 | 561.992 | 395.971 | 504.212 | 507.679 | 504.212 | 528.094 | 749.969 | 631.328 |
| S. cerevisiae | 512.686 | 531.176 | 572.007 | 535.413 | 376.711 | 491.115 | 496.123 | 491.115 | 509.22 | 631.328 | 749.584 |

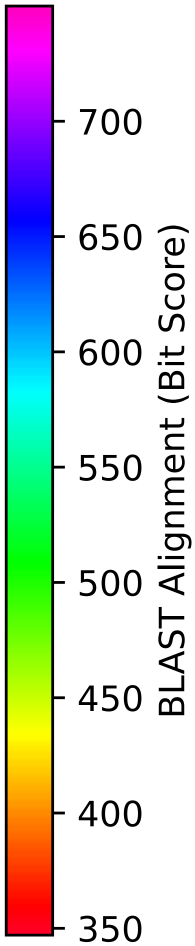


Figure S108: Multiple sequence alignment quality of Fba1 (WHO Critical Pathogens). Cf. Figures S107 and S110.

| Species | Hit Protein | Hit Length (a.a.) | evalue | align_len | bit_score | identity | positive | score | gaps | % identity | % positive |
|---|---|---|---|---|---|---|---|---|---|---|---|
| H.sapiens | - | - | - | - | - | - | - | - | - | - | - |
| N.glabratus | XP_448879.1 uncharacterized p-rotein CAGL0L02497g Nakaseomyc-es glabratus | 361 | 0 | 361 | 631.328 | 300 | 332 | 1627 | 2 | 83.6 | 92.5 |
| H.capsulatum | XP_045285275.1 fructose 1,6-b-iphosphate aldolase Histoplasm-a capsulatum G186AR | 360 | 0 | 360 | 509.22 | 246 | 288 | 1310 | 1 | 68.5 | 80.2 |
| C.neoformans.JEC21 | XP_568771.1 fructose-bisphosp-hate aldolase, putative Crypto-coccus neoformans var. neoform-ans JEC21 | 348 | 5e-176 | 348 | 491.5 | 237 | 284 | 1264 | 1 | 66.0 | 79.1 |
| C.neoformans.grubii.H99 | XP_012047721.1 fructose-bisph-osphate aldolase 1 Cryptococcu-s neoformans var. grubii H99 | 339 | 8.6e-178 | 339 | 496.123 | 238 | 284 | 1276 | 1 | 66.3 | 79.1 |
| C.neoformans.B.3501A | XP_777012.1 hypothetical prot-ein CNBB5380 Cryptococcus neof-ormans var. neoformans B-3501A | 348 | 4.8e-176 | 348 | 491.5 | 237 | 284 | 1264 | 1 | 66.0 | 79.1 |
| C.parapsilosis | XP_036667483.1 uncharacterize-d protein CPAR2 401230 Candida parapsilosis | 352 | 0 | 352 | 535.413 | 257 | 299 | 1378 | 1 | 71.6 | 83.3 |
| C.tropicalis | XP_002545430.1 fructose-bisph-osphate aldolase Candida tropi-calis MYA-3404 | 246 | 9.7e-133 | 246 | 379.793 | 185 | 215 | 974 | 1 | 51.5 | 59.9 |
| C.auris | XP_028888316.1 fructose-bisph-osphate aldolase Candida auris | 352 | 0 | 352 | 572.007 | 263 | 310 | 1473 | 1 | 73.3 | 86.4 |
| C.albicans | XP_722690.1 fructose-bisphosp-hate aldolase Candida albicans SC5314 | 351 | 0 | 351 | 531.176 | 257 | 300 | 1367 | 1 | 71.6 | 83.6 |
| A.fumigatus | XP_754452.1 fructose-bisphosp-hate aldolase, class II Asperg-illus fumigatus Af293 | 360 | 0 | 360 | 512.686 | 246 | 290 | 1319 | 1 | 68.5 | 80.8 |

Table S19: Pairwise alignment info from yeast Fba1 (DEG20010617), cf. Figure S107.

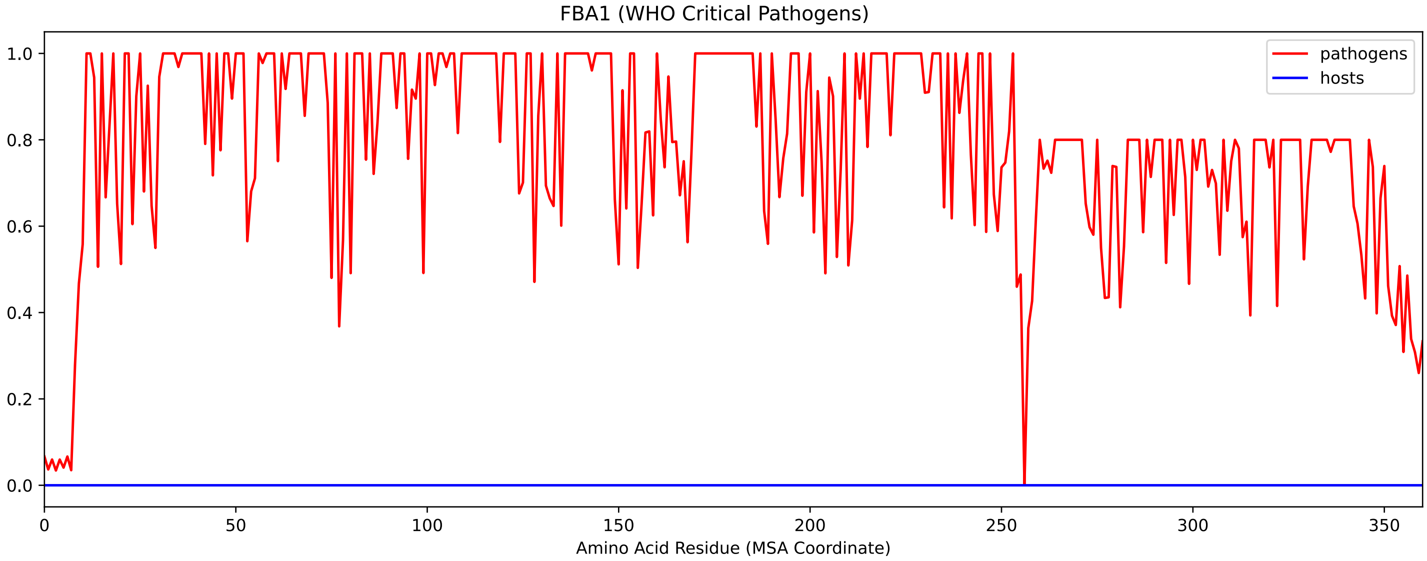


Figure S109: Sneath Similarity of Fba1 for WHO Critical Pathogens, cf. Figure S107

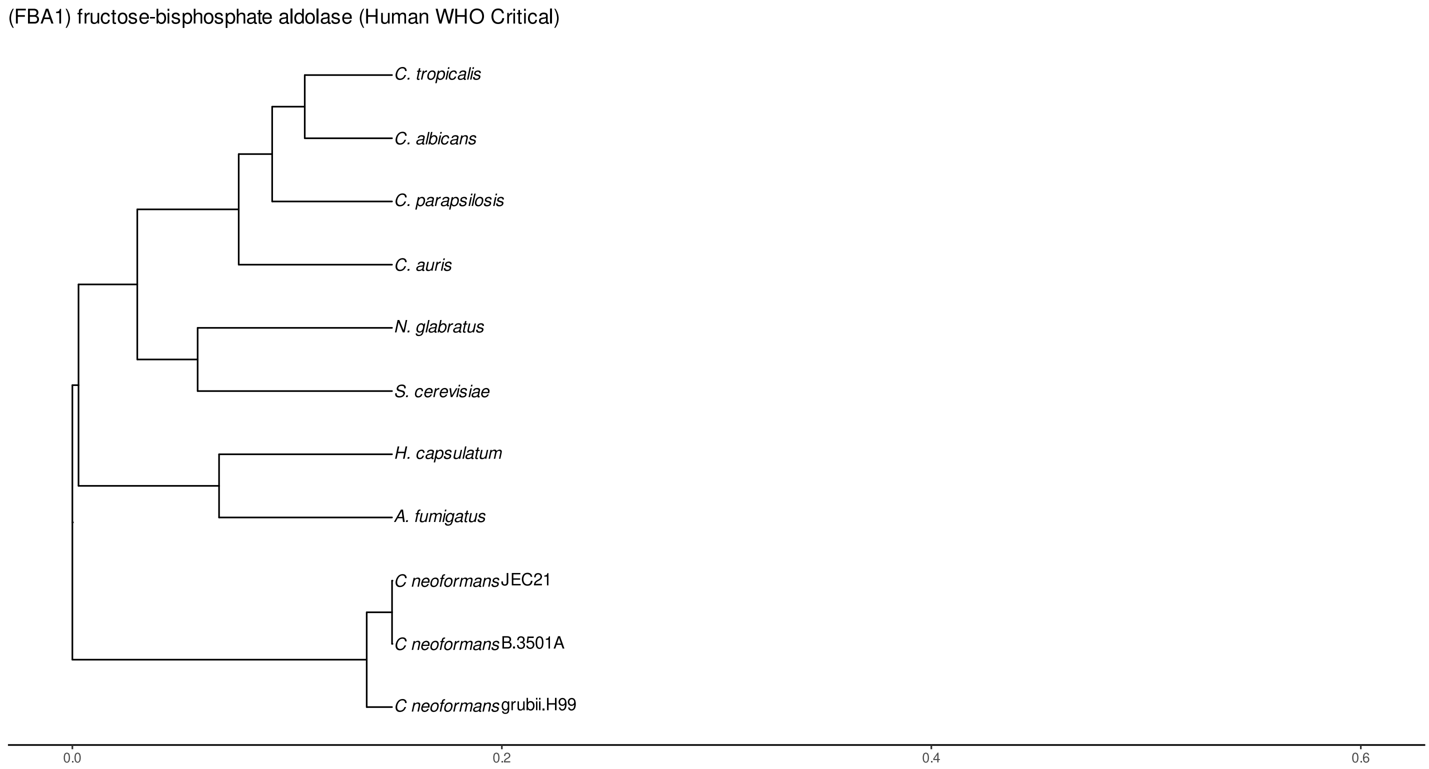


Figure S110: Phylogeny of hits for Fba1 made using clustal omega and visualized using ggtree, cf. Figures S107 and S108.

##### S4.7.2 Top 10 Agricultural Fungal Pathogens

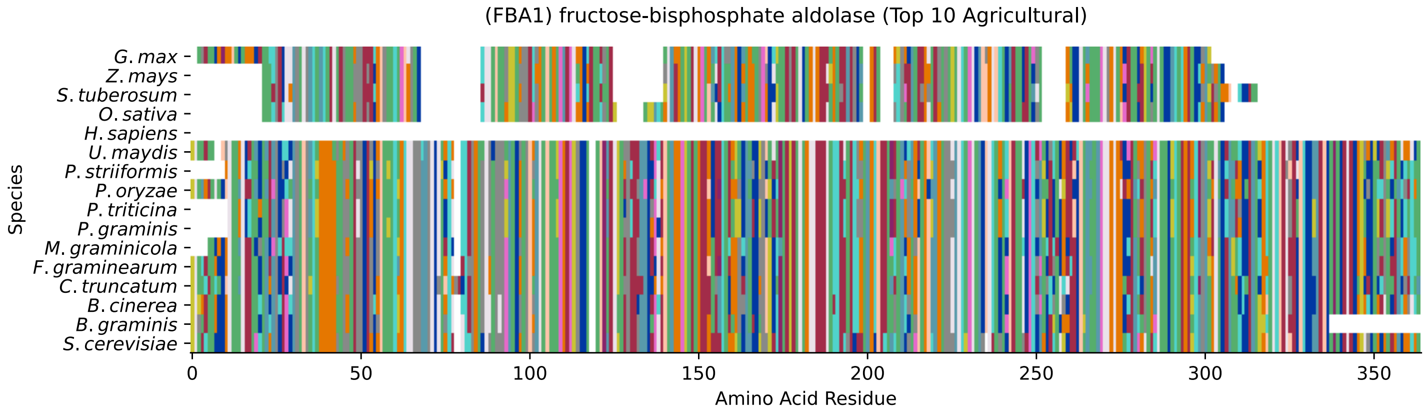


Figure S111: Multiple sequence alignment of yeast Fba1 (Top 10 Agricultural Fungal Pathogens). Cf. Figure S112 for alignment quality, and Figure S113 for Sneath similarity. Cf. Table S20 for protein names, and pairwise alignment metrics with yeast Fba1.

### Fba1 MSA Quality

| | B. g. | B. c. | C. t. | F. g. | M. g. | P. g. | P. t. | P. o. | P. s. | U. m. | S. c. |
|---|---|---|---|---|---|---|---|---|---|---|---|
| B. graminis | 685.256 | 585.489 | 499.59 | 579.326 | 536.954 | 453.366 | 454.907 | 516.538 | 449.129 | 494.582 | 500.745 |
| B. cinerea | 585.489 | 744.191 | 543.888 | 628.246 | 579.711 | 471.855 | 473.781 | 540.806 | 464.922 | 520.776 | 534.258 |
| C. truncatum | 516.153 | 560.066 | 745.347 | 556.214 | 519.235 | 466.463 | 465.692 | 557.37 | 463.381 | 495.738 | 486.108 |
| F. graminearum | 579.326 | 628.246 | 536.954 | 746.117 | 596.275 | 486.108 | 489.574 | 556.984 | 484.182 | 521.161 | 524.628 |
| M. graminicola | 554.673 | 596.66 | 518.85 | 615.15 | 734.561 | 466.077 | 470.7 | 558.91 | 470.315 | 509.605 | 525.783 |
| P. graminis | 470.315 | 488.804 | 466.463 | 504.982 | 466.077 | 729.554 | 713.761 | 458.373 | 671.003 | 547.354 | 478.789 |
| P. triticina | 472.626 | 490.73 | 465.692 | 508.449 | 471.085 | 713.761 | 728.013 | 458.373 | 663.685 | 540.421 | 484.182 |
| P. oryzae | 533.487 | 558.14 | 556.984 | 575.859 | 558.91 | 458.373 | 457.988 | 746.503 | 454.521 | 501.13 | 503.827 |
| P. striiformis | 466.077 | 482.256 | 462.611 | 503.827 | 470.7 | 671.003 | 663.685 | 454.907 | 729.939 | 544.658 | 483.026 |
| U. maydis | 494.582 | 520.776 | 481.485 | 521.161 | 494.582 | 531.561 | 525.013 | 485.337 | 529.635 | 744.577 | 501.13 |
| S. cerevisiae | 500.745 | 534.258 | 469.929 | 524.628 | 508.064 | 461.455 | 466.077 | 485.723 | 464.537 | 501.13 | 749.584 |

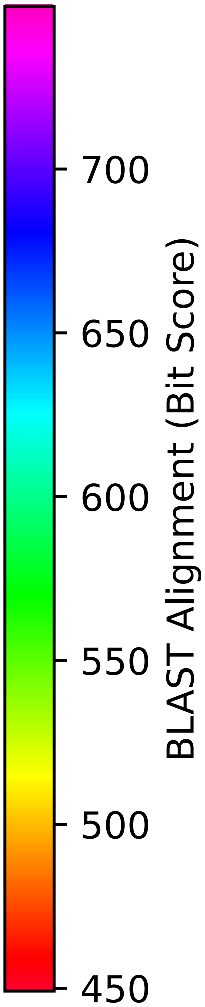


Figure S112: Multiple sequence alignment quality of Fba1 (Top 10 Agricultural Fungal Pathogens). Cf. Figures S111 and S114.

| Species | Hit Protein | Hit Length (a.a.) | evalue | align_len | bit_score | identity | positive | score | gaps | % identity | % positive |
|---|---|---|---|---|---|---|---|---|---|---|---|
| G.max | XP_003530061.1 uncharacterized protein LOC100779987 isoform X1 Glycine max | 297 | 3.2e-09 | 297 | 60.077 | 76 | 123 | 144 | 45 | 21.2 | 34.3 |
| Z.mays | NP_001333690.1 uncharacterized protein LOC100280420 Zea mays | 281 | 4e-06 | 281 | 49.6766 | 74 | 113 | 117 | 43 | 20.6 | 31.5 |
| S.tuberosum | XP_006341517.1 PREDICTED: uncharacterized protein LOC102593631 Solanum tuberosum | 292 | 6.5e-07 | 292 | 51.6026 | 80 | 118 | 122 | 47 | 22.3 | 32.9 |
| O.sativa | XP_015642840.1 uncharacterized protein LOC4340684 isoform X1 Oryza sativa Japonica Group | 285 | 2.5e-07 | 285 | 53.1434 | 77 | 116 | 126 | 44 | 21.4 | 32.3 |
| H.sapiens | - | - | - | - | - | - | - | - | - | - | - |
| U.maydis | XP_011386477.1 putative fructose-bisphosphate aldolase FBA1 Ustilago maydis 521 | 360 | 8.2e-180 | 360 | 501.13 | 231 | 287 | 1289 | 3 | 64.3 | 79.9 |
| P.striiformis | XP_047806328.1 hypothetical protein Pst134EA 013487 Puccinia striiformis f. sp. tritici | 350 | 2.2e-164 | 350 | 464.151 | 218 | 279 | 1193 | 1 | 60.7 | 77.7 |
| P.oryzae | mRNA M BR32 EuGene 00018841-p1 — transcript=mRNA M BR32 EuGene 00018841 — gene=M BR32 EuGene 00018841 — organism=Pyricularia oryzae BR32 — gene product=unspecified product — transcript product=unspecified product — location=BR32 scaffold000-02:792025-793370(-) — protein length=360 — sequence SO=supercontig — SO=protein coding gene — is pseudo=false | 360 | 1.5e-173 | 360 | 485.723 | 238 | 290 | 1249 | 1 | 66.3 | 80.8 |
| P.triticina | XP_053028624.1 uncharacterized protein PtA15 18A126 Puccinia triticina | 349 | 1.2e-165 | 349 | 467.233 | 221 | 277 | 1201 | 1 | 61.6 | 77.2 |
| P.graminis | XP_003320224.1 fructose-bisphosphate aldolase, class II Puccinia graminis f. sp. tritici CRL 75-36-700-3 | 349 | 6.8e-164 | 349 | 461.455 | 218 | 276 | 1186 | 1 | 60.7 | 76.9 |
| M.graminicola | ZTRI 2.441.mRNA-p1 — transcript=ZTRI 2.441.mRNA — gene=ZTRI 2.441 — organism=Zymoseptoria tritici IPO323 — gene product=similar to fructose-bisphosphate aldolase — transcript product=similar to fructose-bisphosphate aldolase — location=Ztri chr 2:1394657-1395855(-) — protein length=359 — sequence SO=chromosome — SO=protein coding gene — is pseudo=false | 355 | 0 | 355 | 508.834 | 243 | 295 | 1309 | 1 | 67.7 | 82.2 |
| F.graminearum | XP_011318737.1 fructose-bisphosphate aldolase Fusarium graminearum PH-1 | 360 | 0 | 360 | 524.628 | 243 | 293 | 1350 | 1 | 67.7 | 81.6 |
| C.truncatum | XP_036586175.1 fructose-bisphosphate aldolase Colletotrichum truncatum | 362 | 8.2e-161 | 362 | 468.774 | 232 | 278 | 1205 | 3 | 64.6 | 77.4 |
| B.cinerea | XP_001556818.1 Bcfba1 Botrytis cinerea B05.10 | 360 | 0 | 360 | 534.258 | 250 | 291 | 1375 | 1 | 69.6 | 81.1 |
| B.graminis | VDB89263.1 — transcript=BGT962-24V316 LOCUS4968 t1 — gene=BGT-96224V316 LOCUS4968 — organism=Blumeria graminis f. sp. tritici 96224 — gene product=unspecified product — transcript product=unspecified product — location=LR026990:8163448-8164635(+) — protein length=344 — sequence SO=chromosome — SO=protein coding gene — is pseudo=false | 333 | 1e-179 | 333 | 500.36 | 235 | 273 | 1287 | 1 | 65.5 | 76.0 |

Table S20: Pairwise alignment info from yeast Fba1 (DEG20010617), cf. Figure S111.

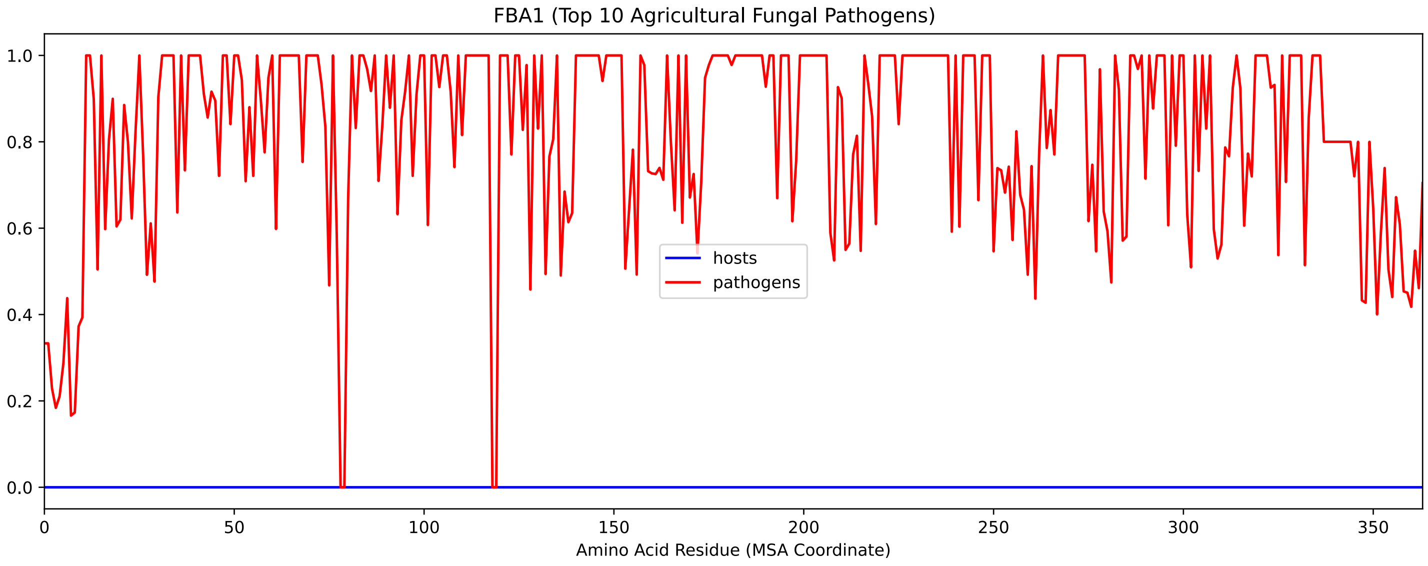


Figure S113: Sneath Similarity of Fba1 for Top 10 Agricultural Fungal Pathogens, cf. Figure S111

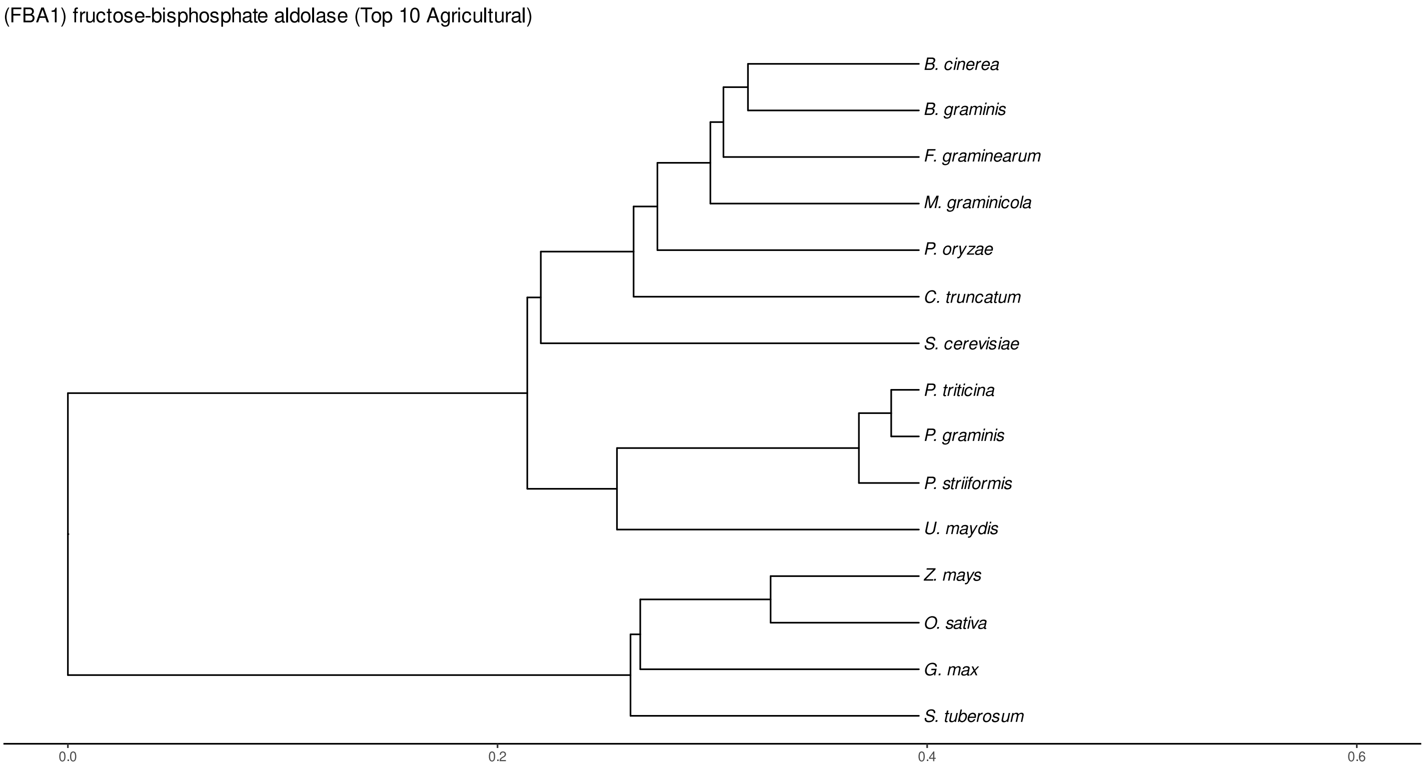


Figure S114: Phylogeny of hits for Fba1 made using clustal omega and visualized using ggtree, cf. Figures S111 and S112.

### S4.7.3 NR

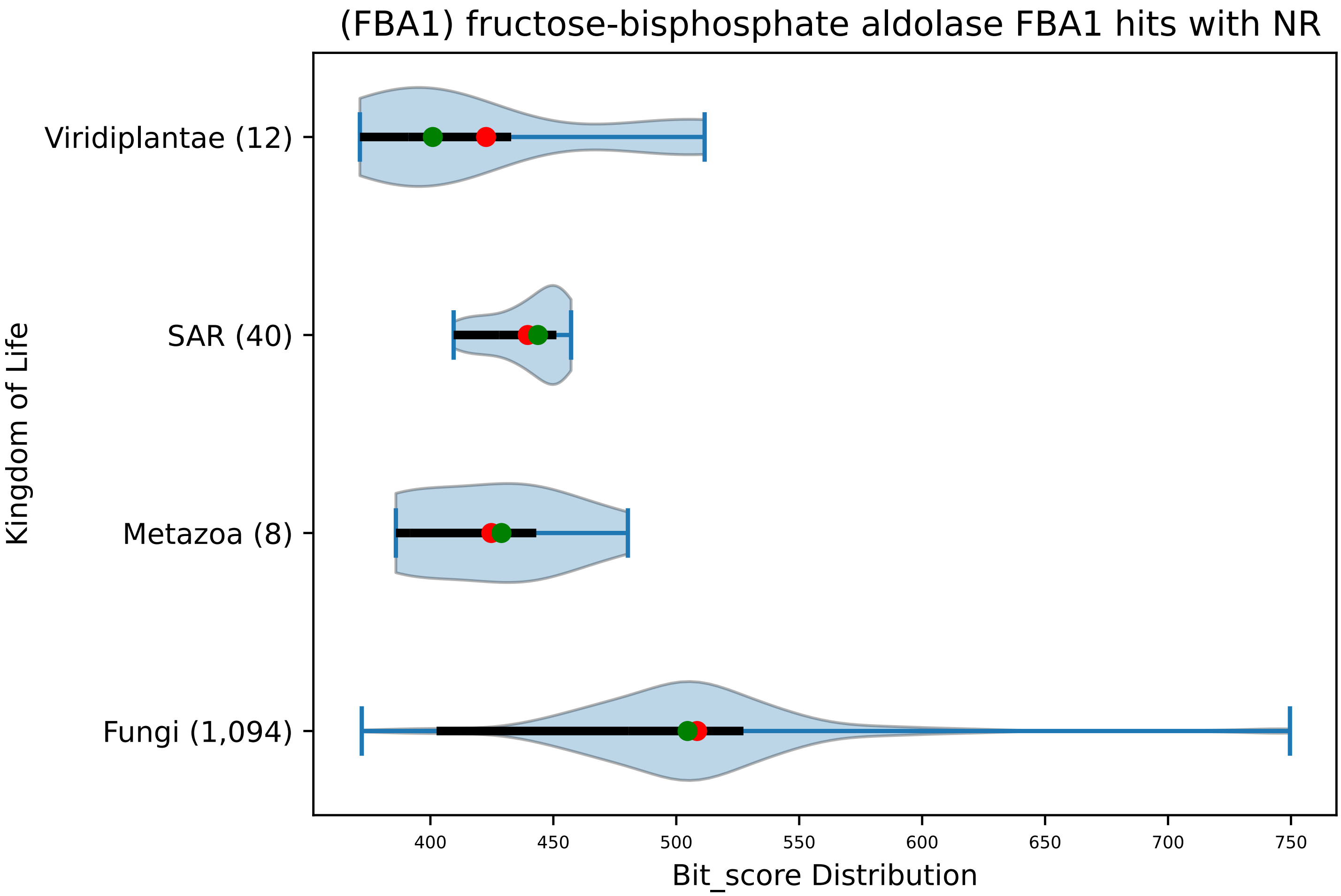


Figure S115: Non-redundant (NR) protein hits for DEG20010617/Fba1, with expectation value of no more than 0.1. Green points are medians, and red points are arithmetic means.

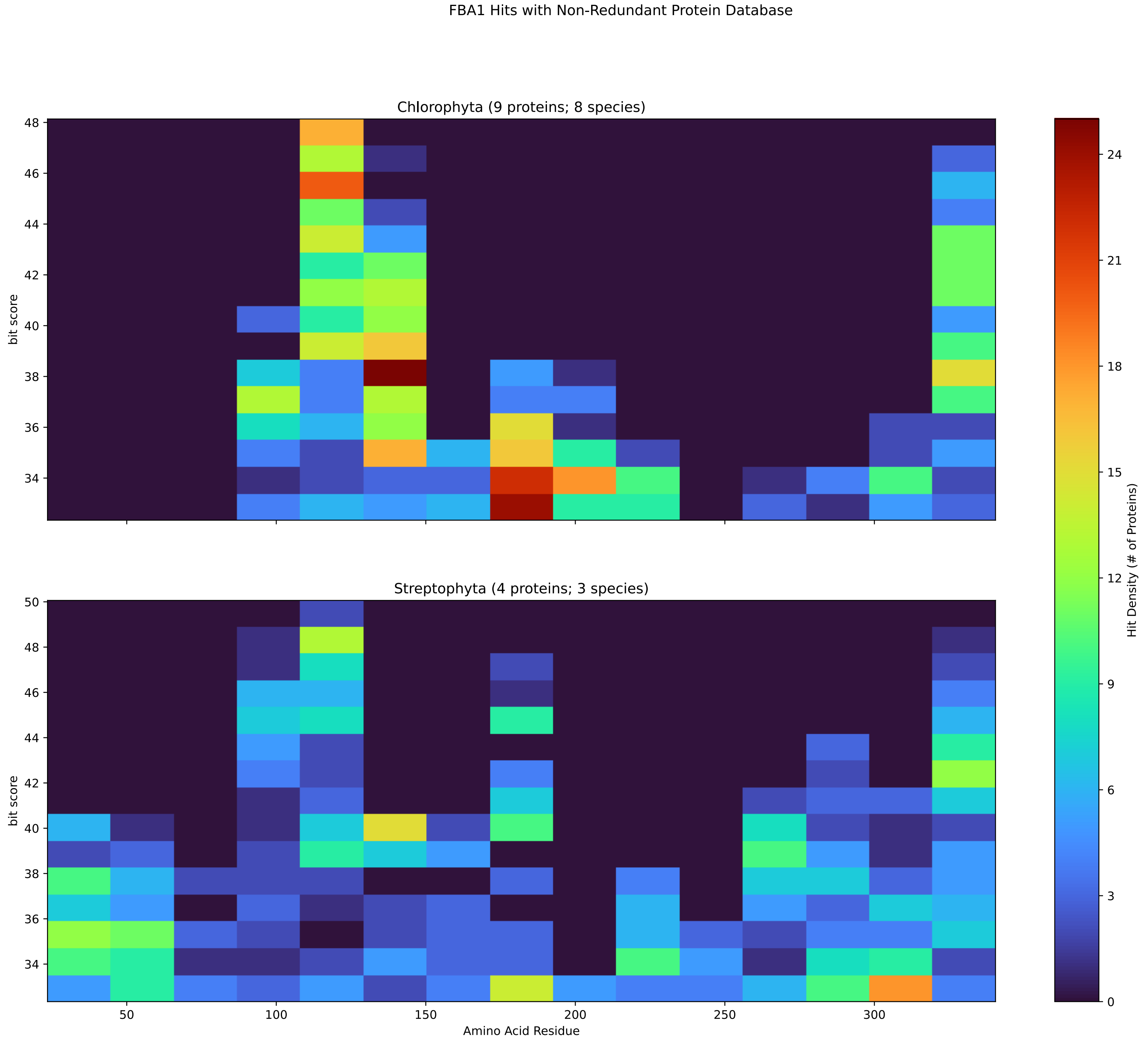


Figure S116: Non-redundant (NR) protein hits for Fba1 in the kingdom Viridiplantae.

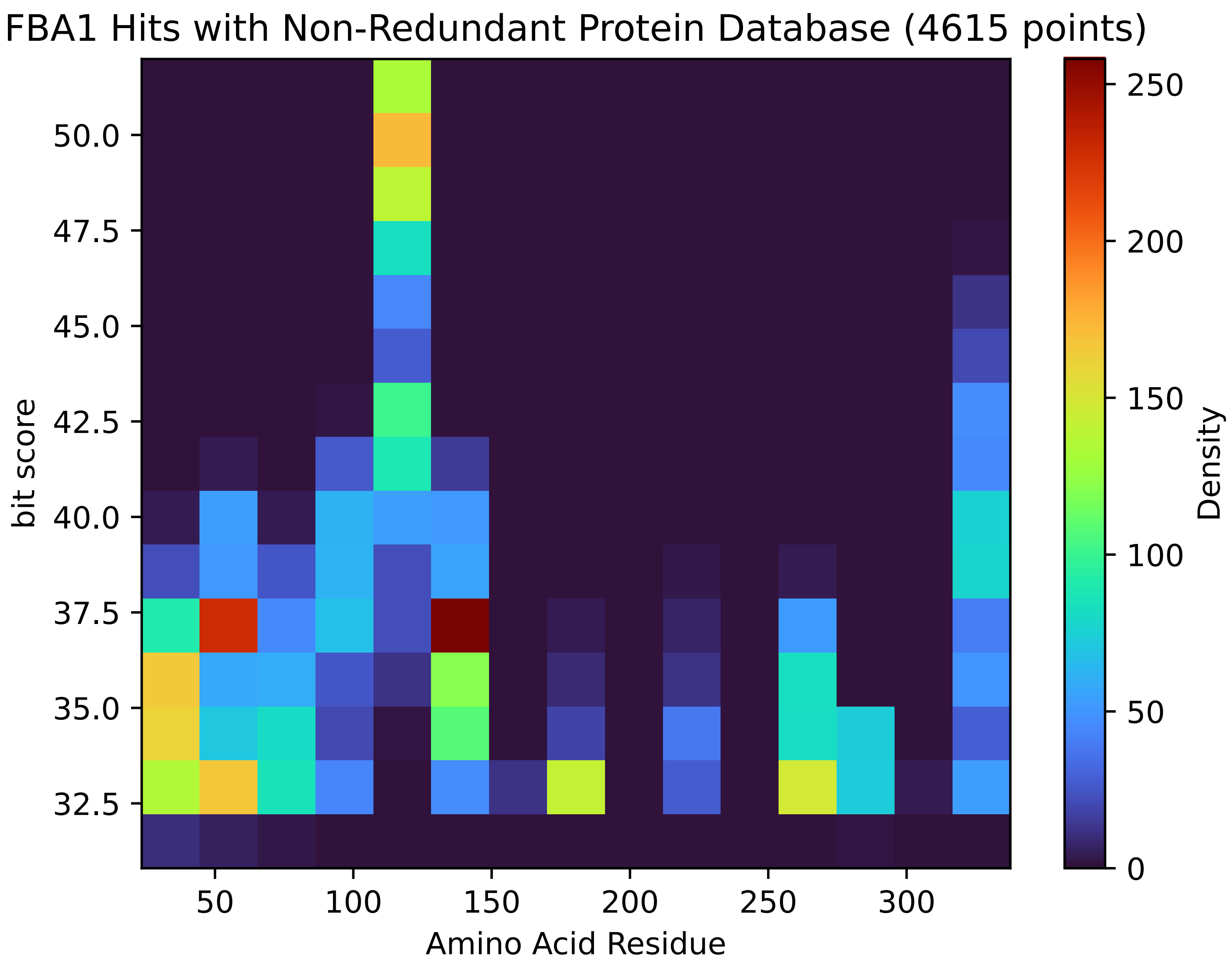


Figure S117: Non-redundant (NR) protein hits for Fba1 in the kingdom SAR.

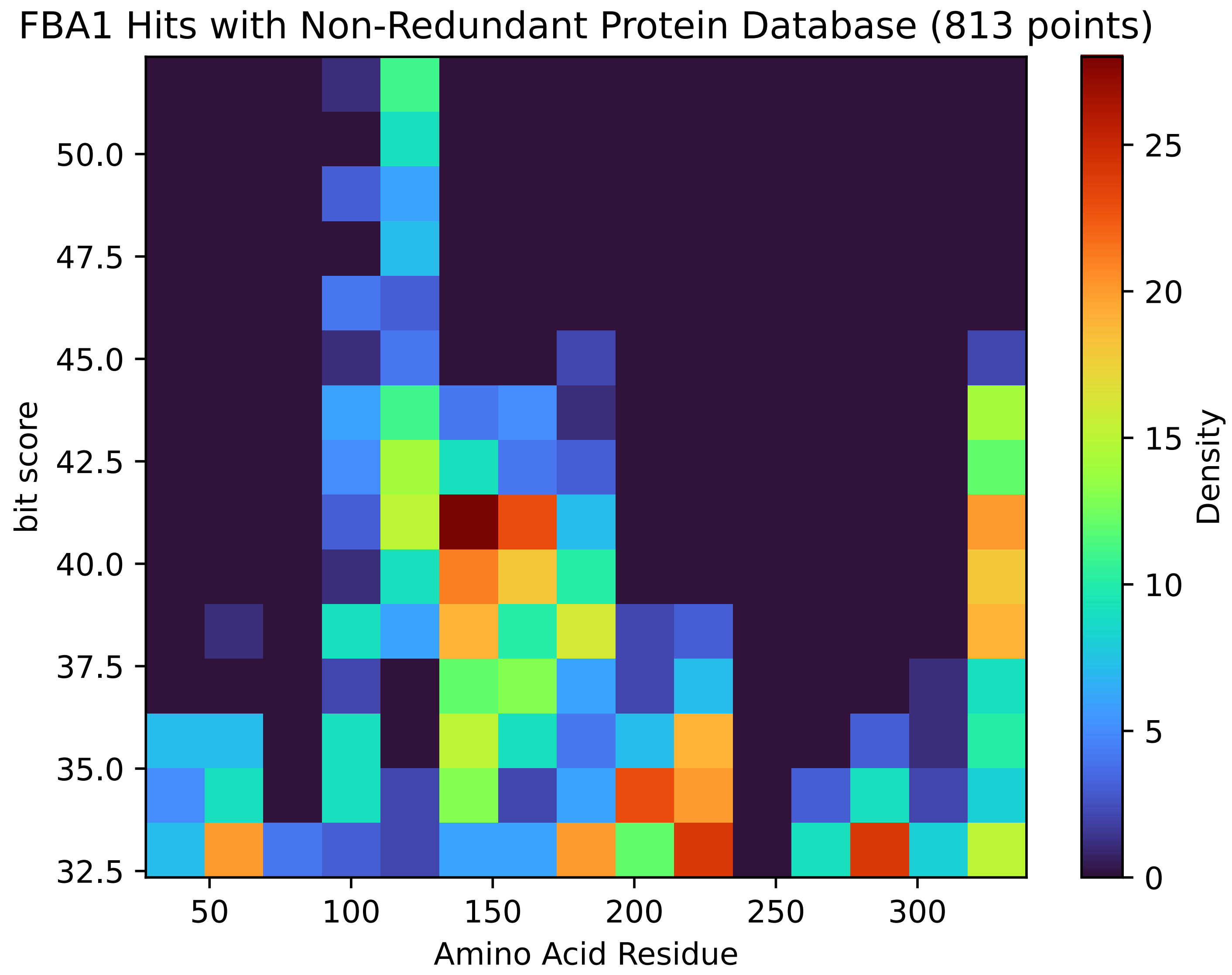


Figure S118: Non-redundant (NR) protein hits for Fba1 in the kingdom Metazoa.

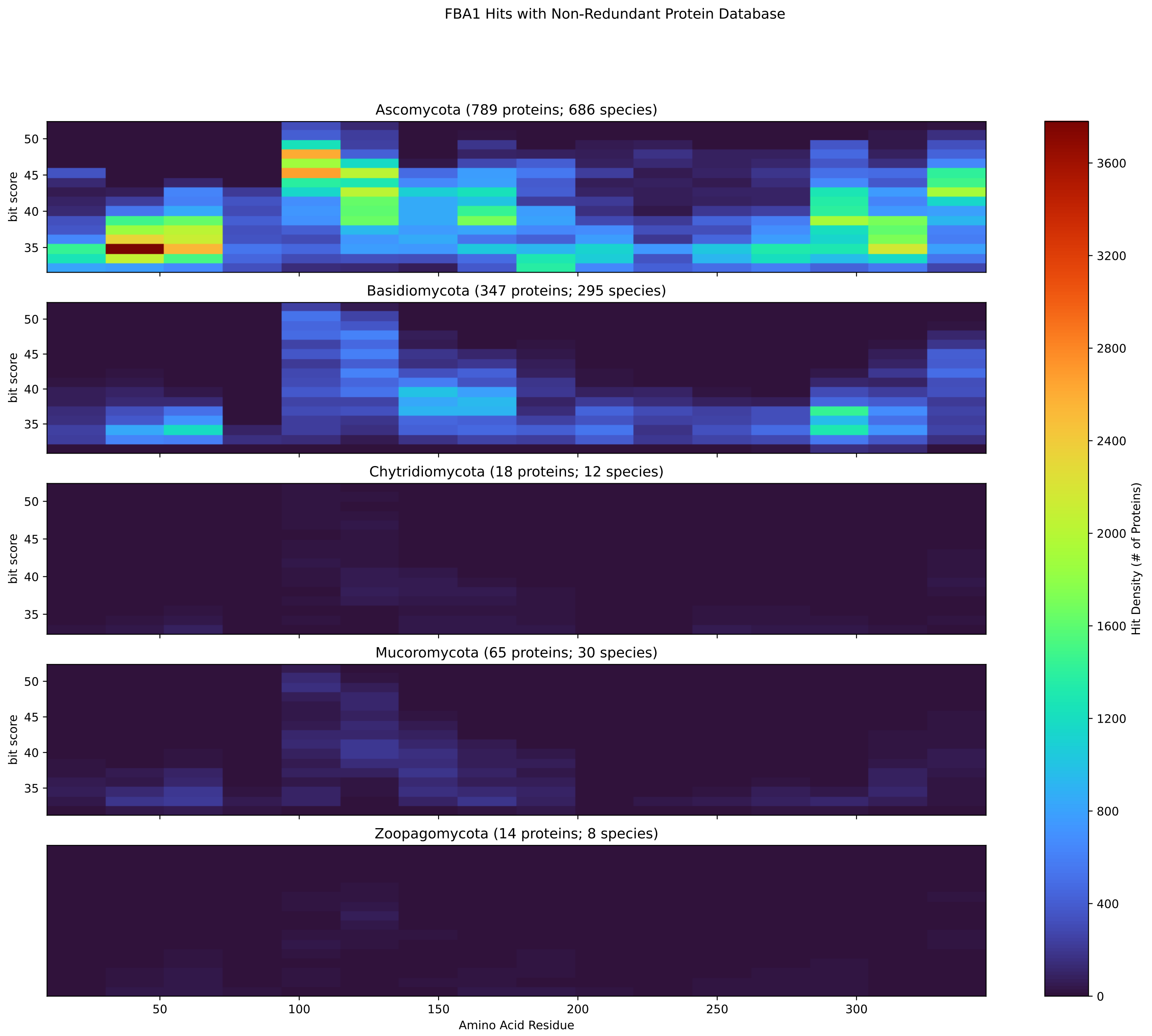


Figure S119: Non-redundant (NR) protein hits for Fba1 in the kingdom Fungi.

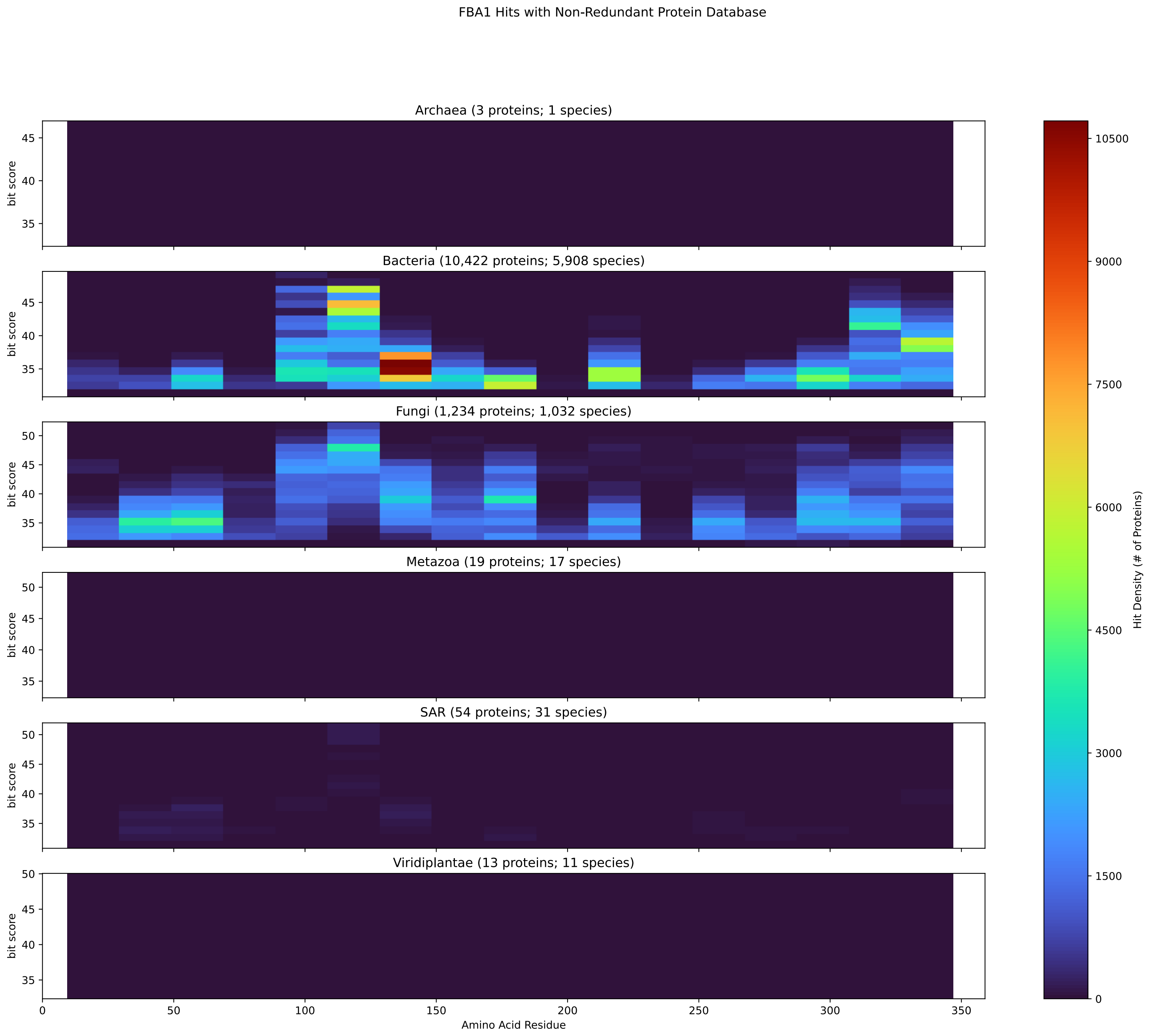


Figure S120: Non-redundant (NR) protein hits for DEG20010617/Fba1 at 20 amino acid length queries.

#### S4.8 Fcy21

##### S4.8.1 WHO Critical Pathogens

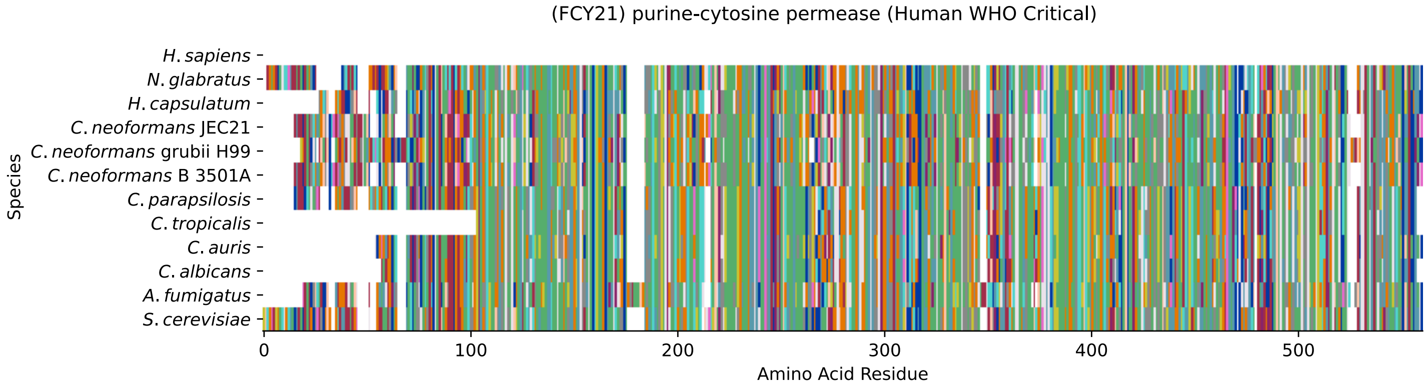


Figure S121: Multiple sequence alignment of yeast Fcy21 (WHO Critical Pathogens). Cf. Figure S122 for alignment quality, and Figure S123 for Sneath similarity. Cf. Table S21 for protein names, and pairwise alignment metrics with yeast Fcy21.

#### Fcy21 MSA Quality

| | *A. f.* | *C. a.* | *C. a.* | *C. p.* | *C. t.* | *C. n.* | *C. n.* | *C. n.* | *H. c.* | *N. g.* | *S. c.* |
|---|---|---|---|---|---|---|---|---|---|---|---|
| *A. fumigatus* | 1047.73 | 413.305 | 379.022 | 408.683 | 371.703 | 327.405 | 322.783 | 327.405 | 501.13 | 352.443 | 412.535 |
| *C. albicans* | 424.476 | 969.148 | 682.559 | 786.563 | 748.814 | 290.426 | 293.508 | 290.426 | 394.045 | 443.736 | 552.362 |
| *C. auris* | 379.022 | 672.929 | 990.334 | 658.677 | 608.986 | 266.929 | 301.982 | 266.929 | 382.489 | 443.351 | 533.872 |
| *C. parapsilosis* | 419.083 | 786.949 | 668.307 | 1042.72 | 723.391 | 298.13 | 285.419 | 298.13 | 388.267 | 454.907 | 544.273 |
| *C. tropicalis* | 382.489 | 748.814 | 618.231 | 723.005 | 878.241 | 261.922 | 261.922 | 261.922 | 374.015 | 421.009 | 509.99 |
| *C. neoformans* B 3501A | 338.961 | 295.049 | 278.485 | 301.212 | 265.774 | 1047.73 | 306.22 | 1047.73 | 319.316 | 293.508 | 305.064 |
| *C. neoformans* grubii H99 | 348.206 | 318.546 | 333.569 | 310.457 | 286.189 | 323.168 | 1057.36 | 323.168 | 364.77 | 314.694 | 323.168 |
| *C. neoformans* JEC21 | 338.961 | 295.049 | 278.485 | 301.212 | 265.774 | 1047.73 | 306.22 | 1047.73 | 319.316 | 293.508 | 305.064 |
| *H. capsulatum* | 501.13 | 382.104 | 382.489 | 376.711 | 363.229 | 305.834 | 330.102 | 305.834 | 1005.36 | 363.999 | 387.111 |
| *N. glabratus* | 358.607 | 445.277 | 454.136 | 456.447 | 422.165 | 285.419 | 296.204 | 285.419 | 369.777 | 1053.51 | 633.639 |
| *S. cerevisiae* | 412.535 | 543.117 | 533.872 | 534.643 | 500.745 | 295.049 | 291.197 | 295.049 | 387.111 | 620.542 | 1078.16 |

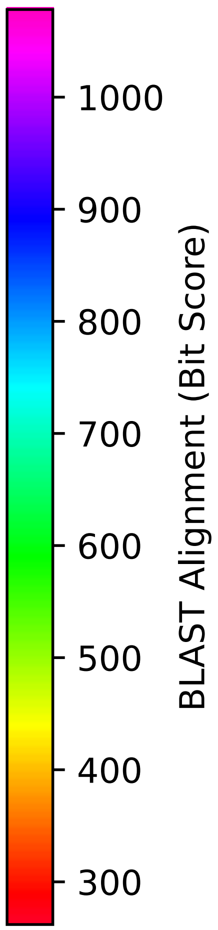


Figure S122: Multiple sequence alignment quality of Fcy21 (WHO Critical Pathogens). Cf. Figures S121 and S124.

| Species | Hit Protein | Hit Length (a.a.) | evalue | align_len | bit_score | identity | positive | score | gaps | % identity | % positive |
|---|---|---|---|---|---|---|---|---|---|---|---|
| H.sapiens | - | - | - | - | - | - | - | - | - | - | - |
| N.glabratus | XP_445191.1 uncharacterized p-rotein CAGL0C00231g Nakaseomyc-es glabratus | 530 | 0 | 530 | 621.313 | 303 | 394 | 1601 | 14 | 57.4 | 74.6 |
| H.capsulatum | XP_045291446.1 purine cytosin-e permease Fcy2 Histoplasma ca-psulatum G186AR | 503 | 5.4e-130 | 503 | 387.111 | 198 | 293 | 993 | 6 | 37.5 | 55.5 |
| C.neoformans.JEC21 | XP_572600.1 cytosine-purine p-ermease, putative Cryptococcus neoformans var. neoformans JE-C21 | 525 | 4.4e-94 | 525 | 294.664 | 184 | 278 | 753 | 20 | 34.8 | 52.7 |
| C.neoformans.grubii.H99 | XP_012052683.1 cytosine perme-ase Cryptococcus neoformans va-r. grubii H99 | 528 | 1.1e-92 | 528 | 291.197 | 191 | 288 | 744 | 23 | 36.2 | 54.5 |
| C.neoformans.B.3501A | XP_773950.1 hypothetical prot-ein CNBH4020 Cryptococcus neof-ormans var. neoformans B-3501A | 525 | 4.3e-94 | 525 | 294.664 | 184 | 278 | 753 | 20 | 34.8 | 52.7 |
| C.parapsilosis | XP_036665281.1 uncharacterize-d protein CPAR2 806580 Candida parapsilosis | 511 | 0 | 511 | 535.413 | 264 | 358 | 1378 | 3 | 50.0 | 67.8 |
| C.tropicalis | XP_002547752.1 purine-cytosin-e permease FCY2 Candida tropic-alis MYA-3404 | 437 | 9.4e-176 | 437 | 500.745 | 247 | 322 | 1288 | 2 | 46.8 | 61.0 |
| C.auris | XP_028891542.1 hypothetical p-rotein Candida auris | 485 | 0 | 485 | 535.028 | 258 | 349 | 1377 | 2 | 48.9 | 66.1 |
| C.albicans | XP_714531.2 purine-cytosine p-ermease Candida albicans SC531-4 | 479 | 0 | 479 | 542.732 | 269 | 349 | 1397 | 2 | 50.9 | 66.1 |
| A.fumigatus | XP_755319.1 purine-cytosine p-ermease Aspergillus fumigatus Af293 | 521 | 8.4e-140 | 521 | 412.92 | 222 | 319 | 1060 | 23 | 42.0 | 60.4 |

Table S21: Pairwise alignment info from yeast Fcy21 (DEG20010294), cf. Figure S121.

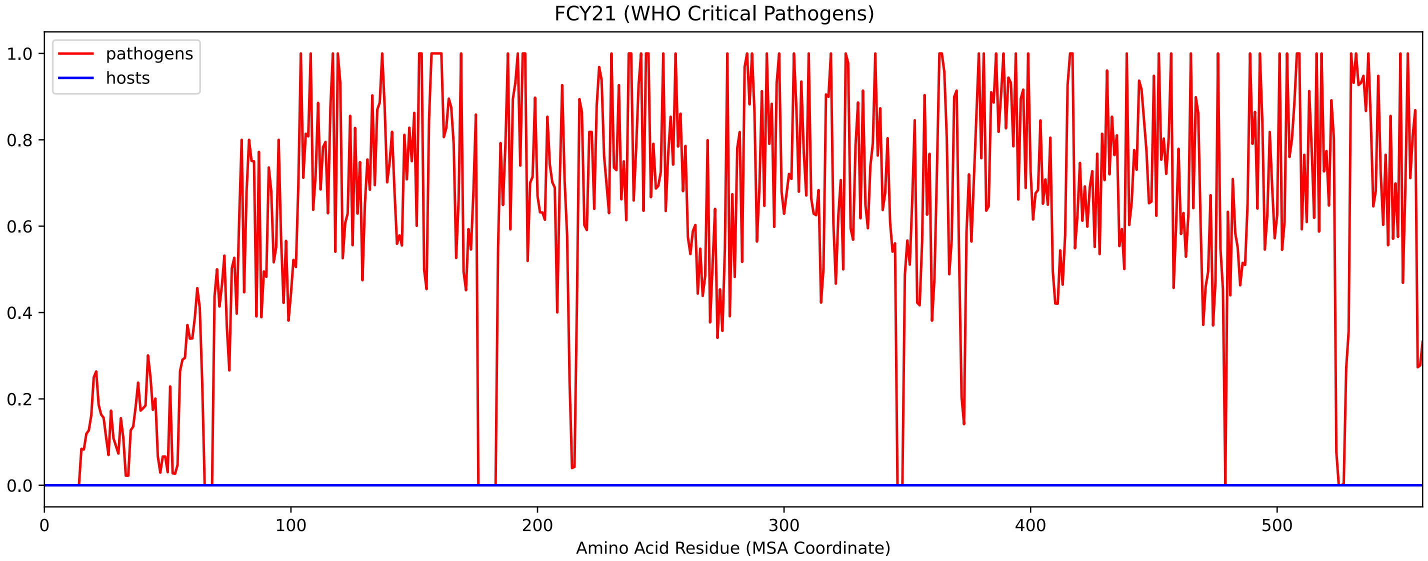


Figure S123: Sneath Similarity of Fcy21 for WHO Critical Pathogens, cf. Figure S121

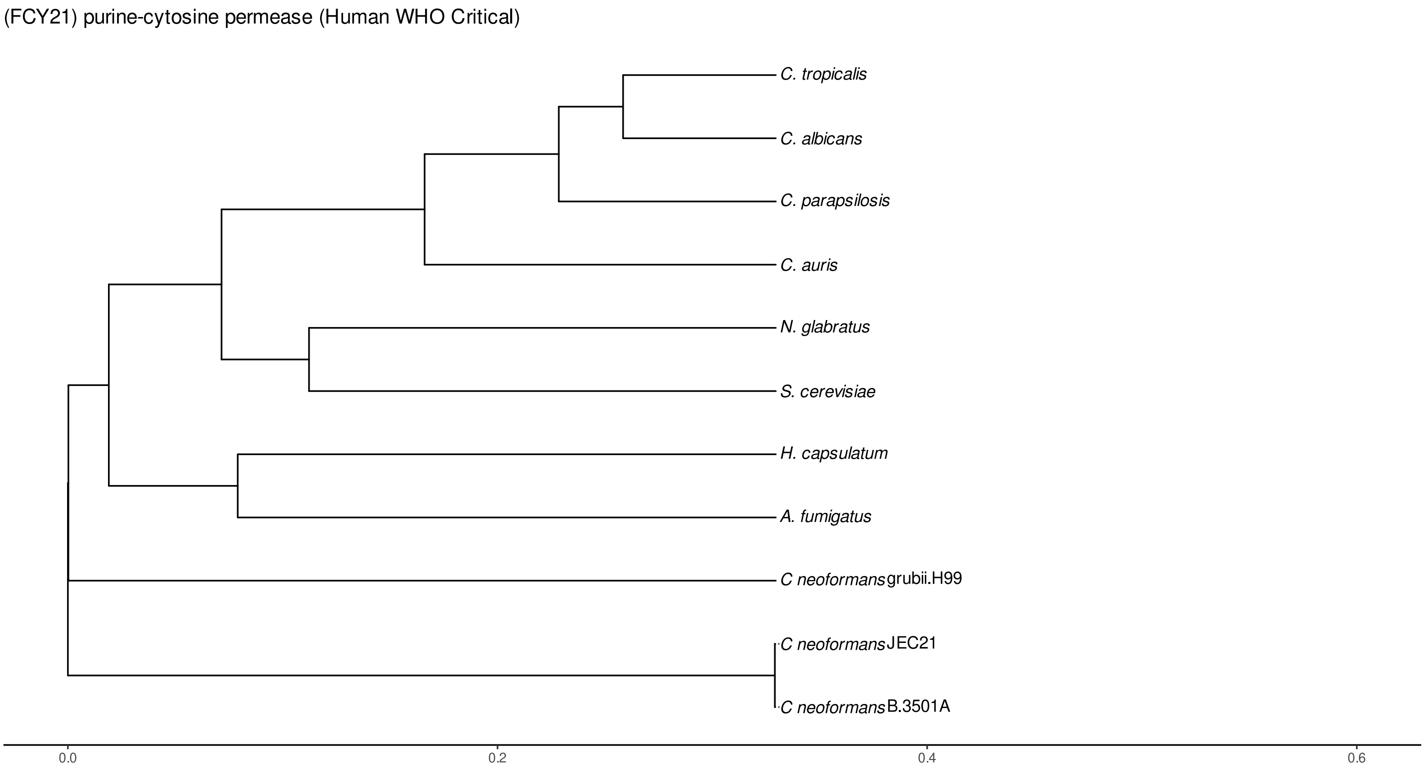


Figure S124: Phylogeny of hits for Fcy21 made using clustal omega and visualized using ggtree, cf. Figures S121 and S122.

##### S4.8.2 Top 10 Agricultural Fungal Pathogens

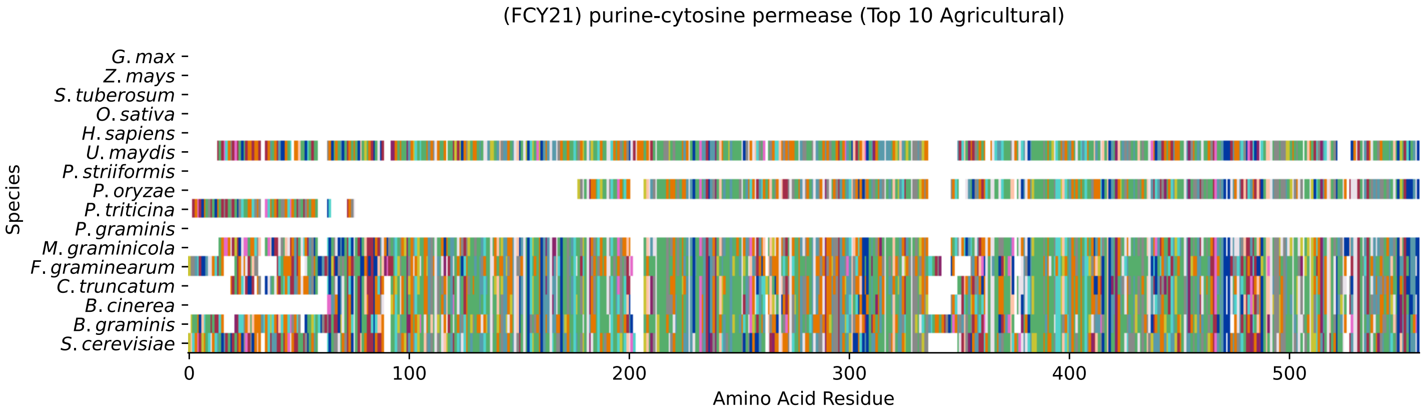


Figure S125: Multiple sequence alignment of yeast Fcy21 (Top 10 Agricultural Fungal Pathogens). Cf. Figure S126 for alignment quality, and Figure S127 for Sneath similarity. Cf. Table S22 for protein names, and pairwise alignment metrics with yeast Fcy21.

### Fcy21 MSA Quality

| | *B. g.* | *B. c.* | *C. t.* | *F. g.* | *M. g.* | *P. g.* | *P. t.* | *P. o.* | *P. s.* | *U. m.* | *S. c.* |
|---|---|---|---|---|---|---|---|---|---|---|---|
| *B. graminis* | 1088.17 | 248.054 | 262.307 | 335.88 | 223.402 | 0 | 0 | 198.749 | 0 | 179.489 | 0 |
| *B. cinerea* | 237.654 | 946.421 | 634.41 | 252.677 | 570.852 | 0 | 0 | 451.825 | 0 | 232.646 | 0 |
| *C. truncatum* | 250.751 | 634.41 | 1028.47 | 250.366 | 601.668 | 0 | 0 | 474.167 | 0 | 241.506 | 0 |
| *F. graminearum* | 323.168 | 252.677 | 250.366 | 1056.59 | 226.483 | 0 | 0 | 210.305 | 0 | 201.06 | 0 |
| *M. graminicola* | 224.942 | 585.104 | 617.461 | 239.58 | 1045.03 | 0 | 0 | 422.935 | 0 | 236.113 | 0 |
| *P. graminis* | 0 | 0 | 0 | 0 | 0 | 0 | 0 | 0 | 0 | 0 | 0 |
| *P. triticina* | 0 | 0 | 0 | 0 | 0 | 0 | 111.309 | 0 | 0 | 0 | 0 |
| *P. oryzae* | 186.037 | 451.825 | 474.167 | 210.305 | 411.379 | 0 | 0 | 720.309 | 0 | 182.57 | 0 |
| *P. striiformis* | 0 | 0 | 0 | 0 | 0 | 0 | 0 | 0 | 0 | 0 | 0 |
| *U. maydis* | 169.859 | 232.646 | 241.506 | 201.06 | 232.646 | 0 | 0 | 182.57 | 0 | 1034.25 | 0 |
| *S. cerevisiae* | 0 | 0 | 0 | 0 | 0 | 0 | 0 | 0 | 0 | 0 | 0 |

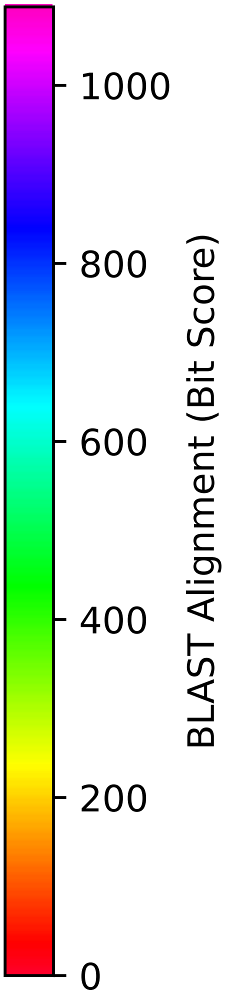


Figure S126: Multiple sequence alignment quality of Fcy21 (Top 10 Agricultural Fungal Pathogens). Cf. Figures S125 and S128.

| Species | Hit Protein | Hit Length (a.a.) | evalue | align_len | bit_score | identity | positive | score | gaps | % identity | % positive |
|---|---|---|---|---|---|---|---|---|---|---|---|
| G.max | - | - | - | - | - | - | - | - | - | - | - |
| Z.mays | - | - | - | - | - | - | - | - | - | - | - |
| S.tuberosum | - | - | - | - | - | - | - | - | - | - | - |
| O.sativa | - | - | - | - | - | - | - | - | - | - | - |
| H.sapiens | - | - | - | - | - | - | - | - | - | - | - |
| U.maydis | XP_011390683.1 uncharacterize-d protein UMAG 04197 Ustilago maydis 521 | 520 | 2.8e-61 | 520 | 208.764 | 144 | 260 | 530 | 20 | 27.3 | 49.2 |
| P.striiformis | - | - | - | - | - | - | - | - | - | - | - |
| P.oryzae | mRNA M BR32 EuGene 00017951-p1 — transcript=mRNA M BR32 EuGe-ne 00017951 — gene=M BR32 EuGe-ne 00017951 — organism=Pyricul-aria oryzae BR32 — gene produc-t=unspecified product — transc-ript product=unspecified produ-ct — location=BR32 scaffold000-02:511879-513124(-) — protein length=359 — sequence SO=super-contig — SO=protein coding gen-e — is pseudo=false | 362 | 1.2e-90 | 362 | 280.796 | 152 | 222 | 717 | 5 | 28.8 | 42.0 |
| P.triticina | XP_053023736.1 uncharacterize-d protein PtA15 9A306 Puccinia triticina | 61 | 0.085 | 61 | 34.2686 | 20 | 33 | 77 | 0 | 3.8 | 6.2 |
| P.graminis | - | - | - | - | - | - | - | - | - | - | - |
| M.graminicola | ZTRI 1.37.mRNA-p1 — transcript-=ZTRI 1.37.mRNA — gene=ZTRI 1.-37 — organism=Zymoseptoria tri-tici IPO323 — gene product=hyp-othetical protein — transcript product=hypothetical protein — location=Ztri chr 1:223670-2-26762(-) — protein length=626 — sequence SO=chromosome — SO=-protein coding gene — is pseud-o=false | 522 | 9.4e-125 | 522 | 377.867 | 209 | 309 | 969 | 11 | 39.6 | 58.5 |
| F.graminearum | XP_011320615.1 hypothetical p-rotein FGSG 13426 Fusarium gra-minearum PH-1 | 535 | 2e-82 | 535 | 265.003 | 183 | 272 | 676 | 22 | 34.7 | 51.5 |
| C.truncatum | XP_036587609.1 purine-cytosin-e permease (NCS1 nucleoside tr-ansporter) Colletotrichum trun-catum | 513 | 6.7e-141 | 513 | 415.616 | 215 | 315 | 1067 | 10 | 40.7 | 59.7 |
| B.cinerea | XP_001559974.1 hypothetical p-rotein BCIN 05g03050 Botrytis cinerea B05.10 | 475 | 3.8e-130 | 475 | 388.267 | 205 | 289 | 996 | 7 | 38.8 | 54.7 |
| B.graminis | VCU39960.1 — transcript=BGT962-24V316 LOCUS1207 t1 — gene=BGT-96224V316 LOCUS1207 — organism-=Blumeria graminis f. sp. trit-ici 96224 — gene product=unspe-cified product — transcript pr-oduct=unspecified product — lo-cation=LR026985:8766531-876835-2(+) — protein length=534 — se-quence SO=chromosome — SO=prot-ein coding gene — is pseudo=fa-lse | 543 | 1.1e-62 | 543 | 212.616 | 165 | 263 | 540 | 30 | 31.2 | 49.8 |

Table S22: Pairwise alignment info from yeast Fcy21 (DEG20010294), cf. Figure S125.

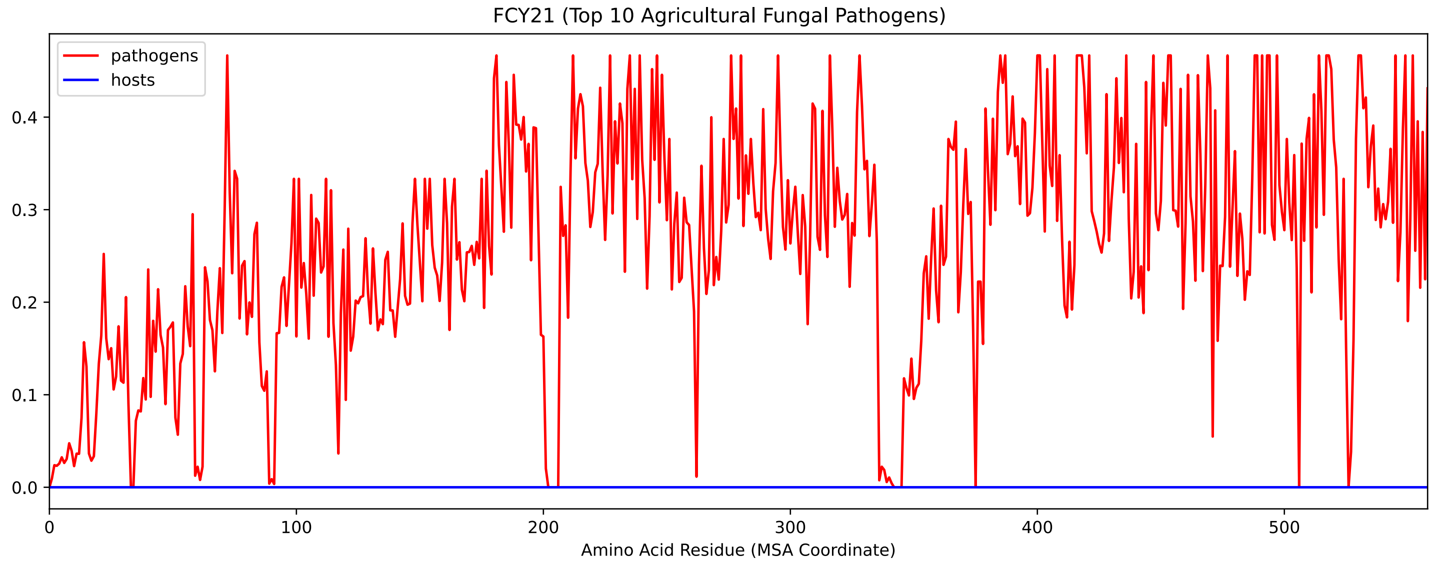


Figure S127: Sneath Similarity of Fcy21 for Top 10 Agricultural Fungal Pathogens, cf. Figure S125

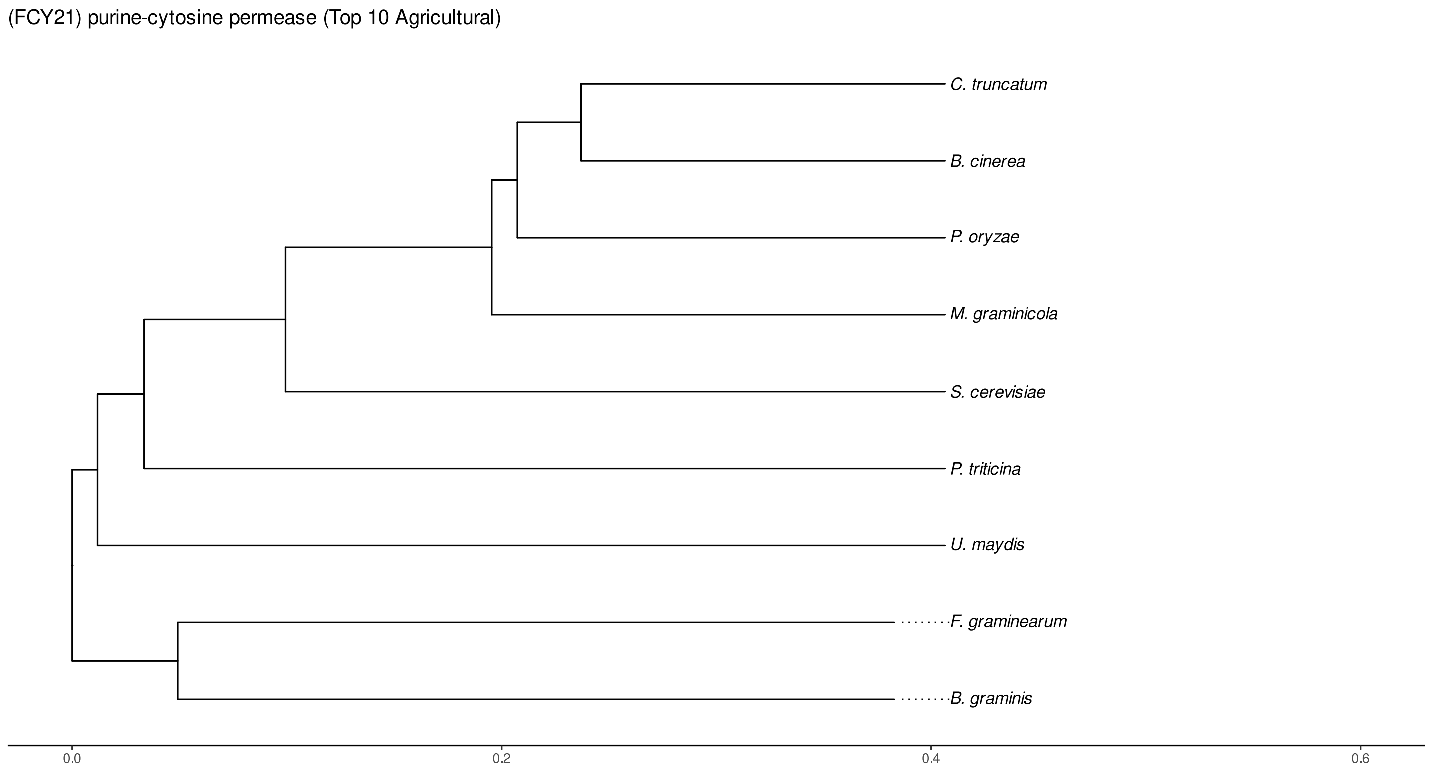


Figure S128: Phylogeny of hits for Fcy21 made using clustal omega and visualized using ggtree, cf. Figures S125 and S126.

#### S4.8.3 NR

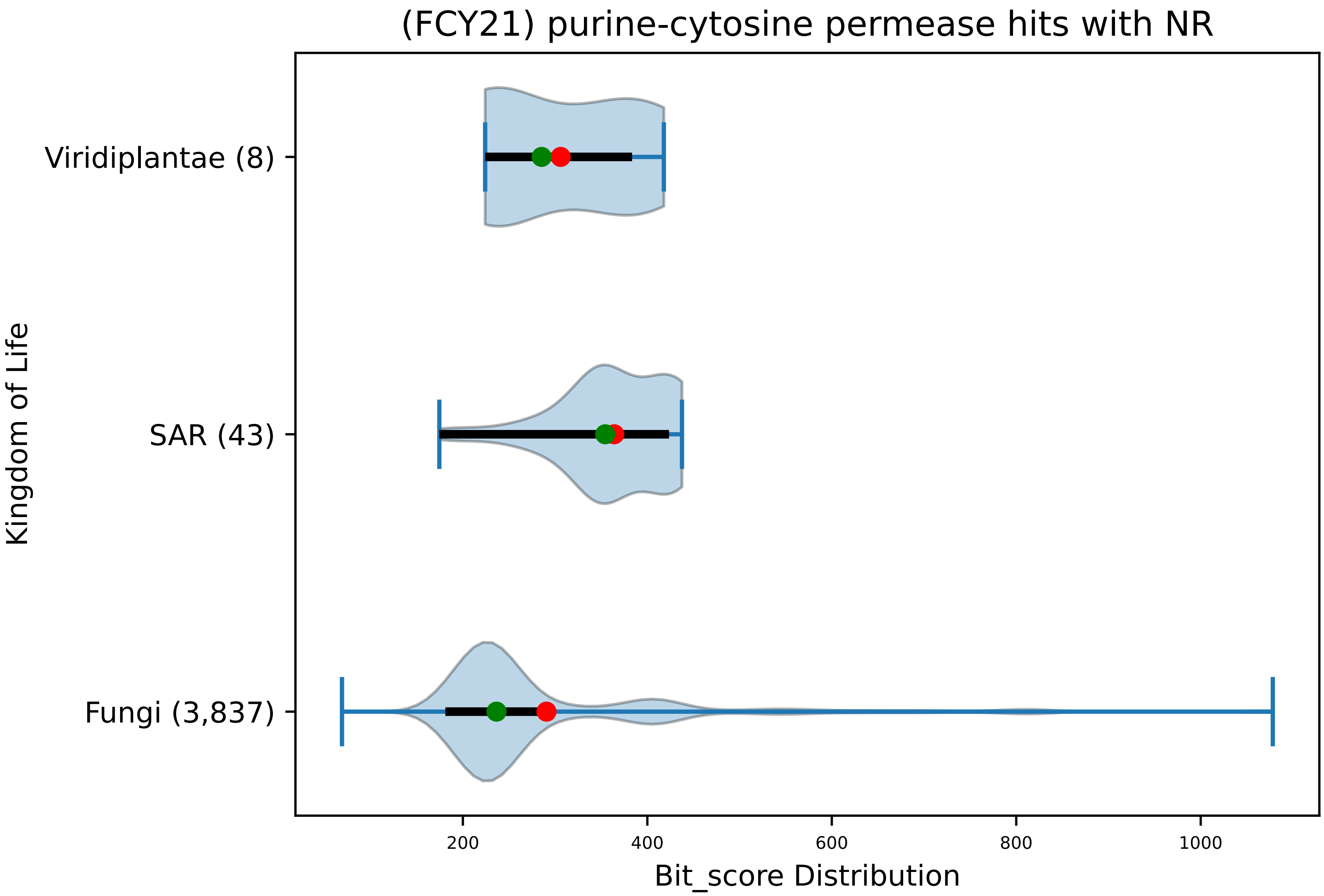


Figure S129: Non-redundant (NR) protein hits for DEG20010294/Fcy21, with expectation value of no more than 0.1. Green points are medians, and red points are arithmetic means.

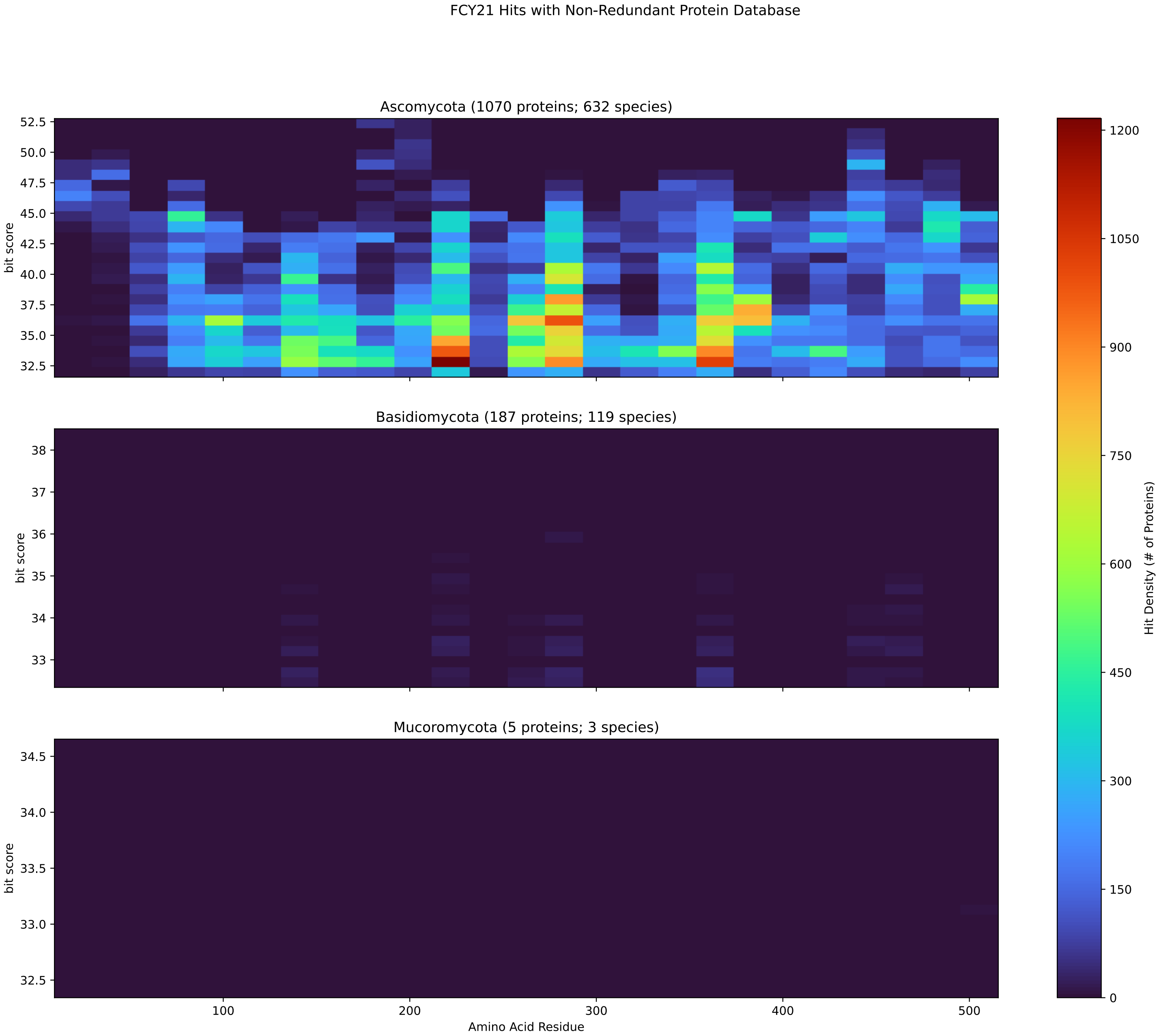


Figure S130: Non-redundant (NR) protein hits for Fcy21 in the kingdom Fungi.

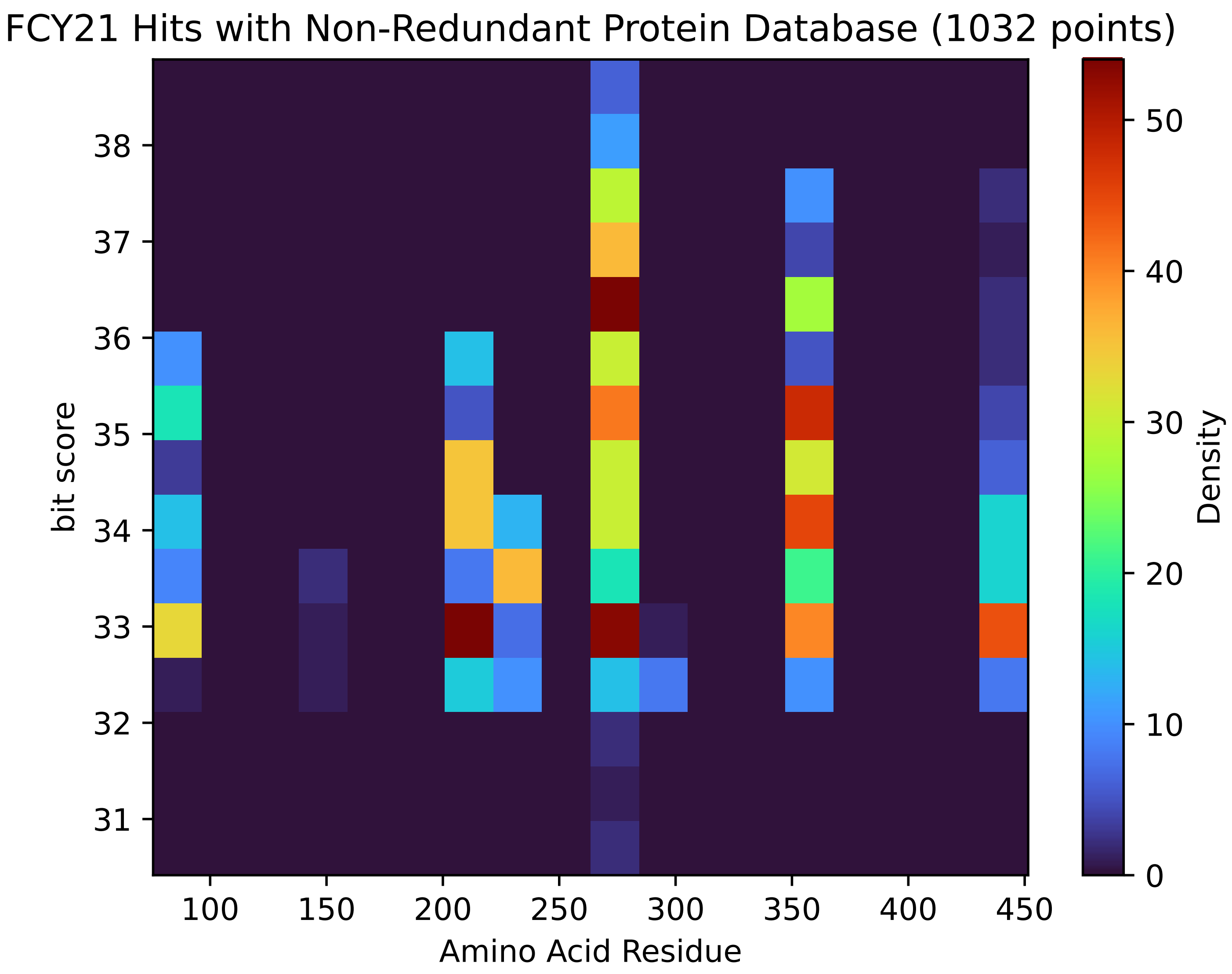


Figure S131: Non-redundant (NR) protein hits for Fcy21 in the kingdom SAR.

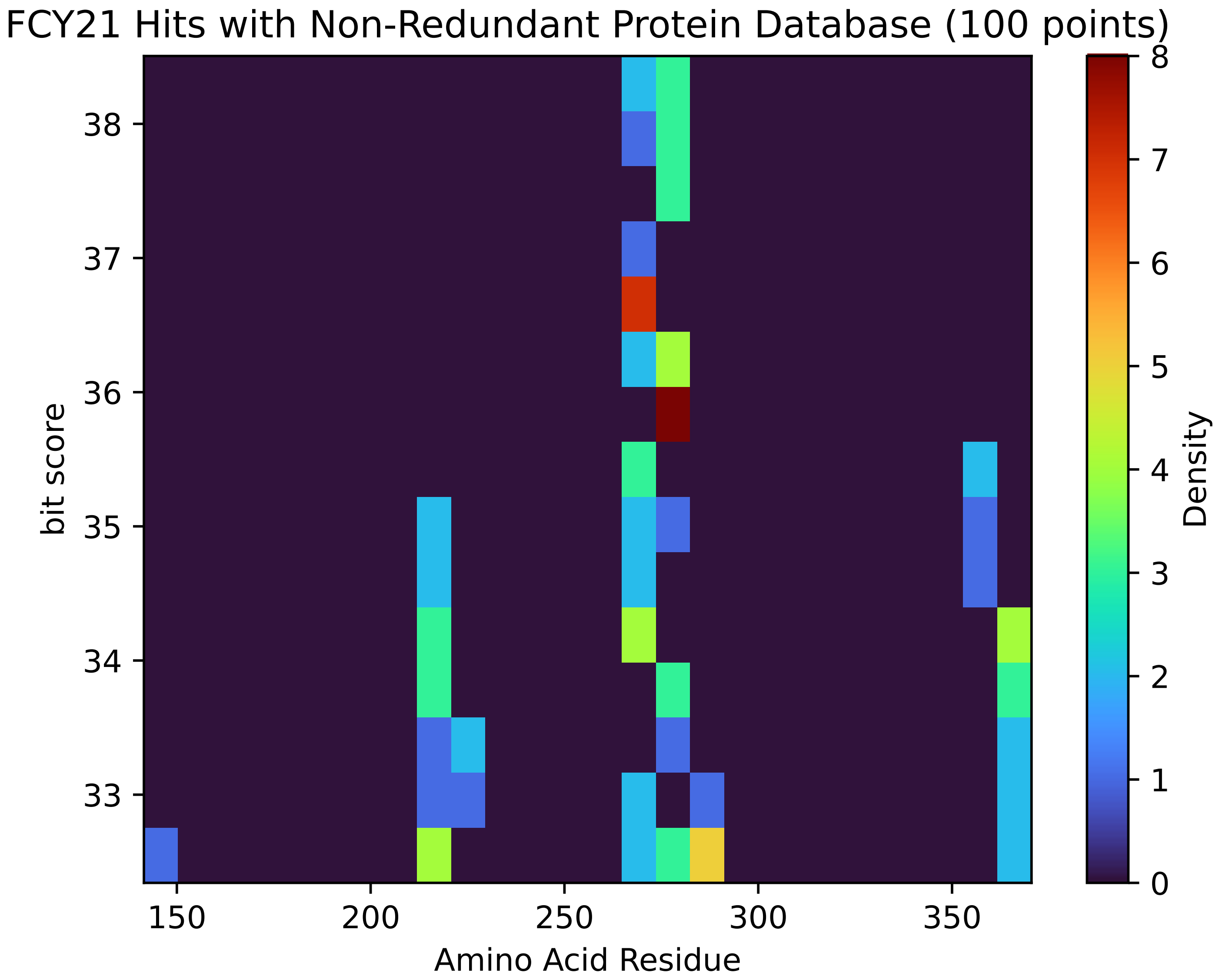


Figure S132: Non-redundant (NR) protein hits for Fcy21 in the kingdom Viridiplantae.

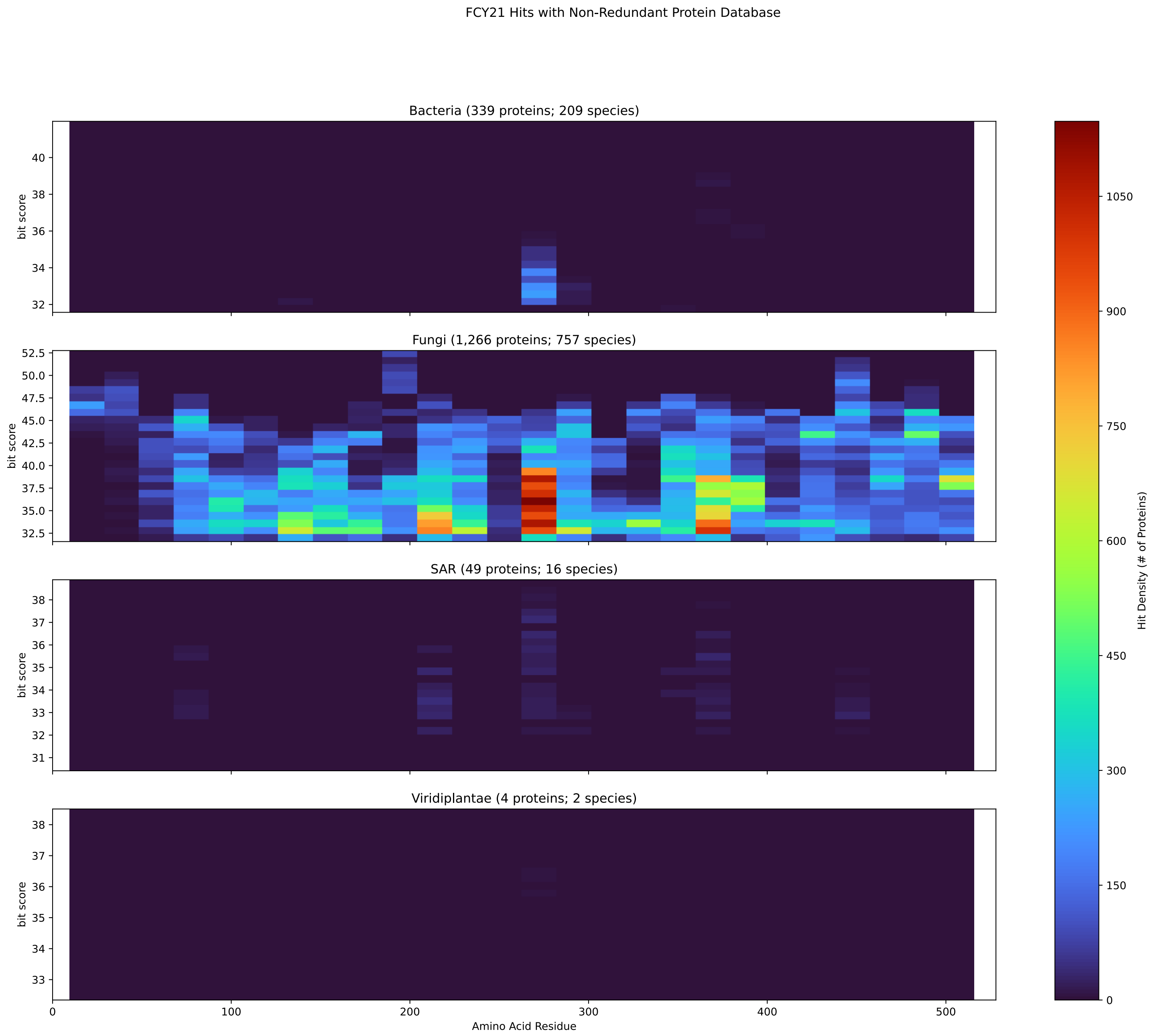


Figure S133: Non-redundant (NR) protein hits for DEG20010294/Fcy21 at 20 amino acid length queries.

#### S4.9 Fol1

##### S4.9.1 WHO Critical Pathogens

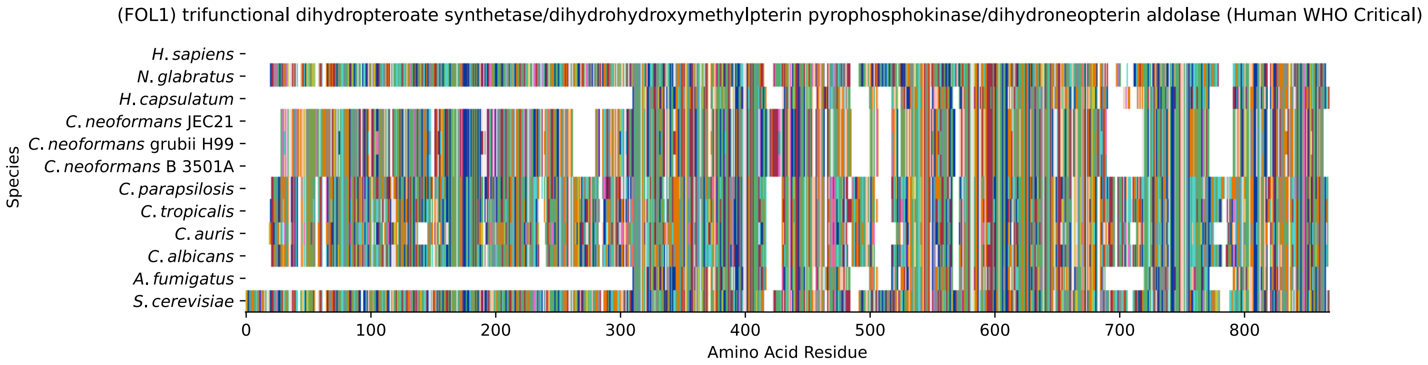


Figure S134: Multiple sequence alignment of yeast Fol1 (WHO Critical Pathogens). Cf. Figure S135 for alignment quality, and Figure S136 for Sneath similarity. Cf. Table S23 for protein names, and pairwise alignment metrics with yeast Fol1.

#### Fol1 MSA Quality

| | *A. f.* | *C. a.* | *C. a.* | *C. p.* | *C. t.* | *C. n.* | *C. n.* | *C. n.* | *H. c.* | *N. g.* | *S. c.* |
|---|---|---|---|---|---|---|---|---|---|---|---|
| *A. fumigatus* | 920.228 | 378.252 | 360.533 | 331.257 | 374.785 | 315.464 | 326.25 | 313.153 | 587.03 | 303.523 | 331.643 |
| *C. albicans* | 380.563 | 1606.65 | 754.592 | 1057.74 | 1261.9 | 310.071 | 318.546 | 313.538 | 342.428 | 497.278 | 534.258 |
| *C. auris* | 355.14 | 742.265 | 1570.44 | 723.391 | 761.911 | 288.5 | 279.641 | 282.722 | 324.324 | 496.508 | 506.138 |
| *C. parapsilosis* | 326.635 | 1041.95 | 723.391 | 1624.37 | 1060.06 | 268.085 | 265.774 | 266.544 | 305.449 | 501.13 | 522.702 |
| *C. tropicalis* | 369.392 | 1243.41 | 761.911 | 1060.06 | 1615.9 | 291.967 | 296.59 | 296.975 | 313.923 | 509.99 | 547.74 |
| *C. neoformans* B 3501A | 308.145 | 298.901 | 288.5 | 268.085 | 291.967 | 1463.74 | 1382.08 | 1454.5 | 299.671 | 315.849 | 320.857 |
| *C. neoformans* grubii H99 | 320.472 | 308.531 | 279.641 | 265.774 | 296.59 | 1382.08 | 1469.52 | 1382.85 | 301.982 | 311.227 | 313.923 |
| *C. neoformans* JEC21 | 307.375 | 302.753 | 282.722 | 266.544 | 296.975 | 1454.5 | 1382.85 | 1464.51 | 297.36 | 313.538 | 317.005 |
| *H. capsulatum* | 598.201 | 350.517 | 343.584 | 323.553 | 332.798 | 320.472 | 324.709 | 318.931 | 955.281 | 273.863 | 310.457 |
| *N. glabratus* | 298.516 | 483.411 | 496.508 | 501.13 | 509.99 | 315.849 | 311.227 | 313.538 | 256.144 | 1634.77 | 854.744 |
| *S. cerevisiae* | 327.02 | 521.931 | 506.138 | 522.702 | 547.74 | 320.857 | 313.923 | 317.005 | 292.352 | 854.744 | 1704.88 |

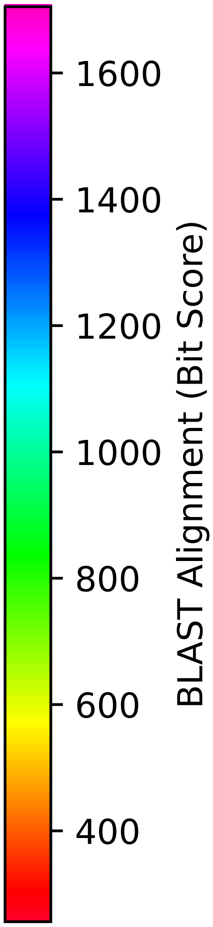


Figure S135: Multiple sequence alignment quality of Fol1 (WHO Critical Pathogens). Cf. Figures S134 and S137.

| Species | Hit Protein | Hit Length (a.a.) | evalue | align_len | bit_score | identity | positive | score | gaps | % identity | % positive |
|---|---|---|---|---|---|---|---|---|---|---|---|
| H.sapiens | - | - | - | - | - | - | - | - | - | - | - |
| N.glabratus | XP_448052.1 uncharacterized protein CAGL0J07920g Nakaseomyces glabratus | 806 | 0 | 806 | 855.129 | 417 | 561 | 2208 | 21 | 50.6 | 68.1 |
| H.capsulatum | XP_045283569.1 folic acid synthesis protein Histoplasma capsulatum G186AR | 528 | 2e-88 | 528 | 292.738 | 198 | 280 | 748 | 63 | 24.0 | 34.0 |
| C.neoformans.JEC21 | XP_570005.1 folic acid and derivative biosynthesis-related protein, putative Cryptococcus neoformans var. neoformans JEC21 | 818 | 6e-97 | 818 | 317.005 | 260 | 386 | 811 | 126 | 31.6 | 46.8 |
| C.neoformans.grubii.H99 | XP_012048132.1 dihydropteroate synthase Cryptococcus neoformans var. grubii H99 | 819 | 9.2e-96 | 819 | 313.923 | 263 | 391 | 803 | 128 | 31.9 | 47.5 |
| C.neoformans.B.3501A | XP_776823.1 hypothetical protein CNBC3140 Cryptococcus neoformans var. neoformans B-3501A | 818 | 1.5e-98 | 818 | 321.242 | 261 | 387 | 822 | 126 | 31.7 | 47.0 |
| C.parapsilosis | XP_036664526.1 uncharacterized protein CPAR2 303390 Candida parapsilosis | 828 | 4.7e-175 | 828 | 522.702 | 317 | 488 | 1345 | 65 | 38.5 | 59.2 |
| C.tropicalis | XP_002551057.1 conserved hypothetical protein Candida tropicalis MYA-3404 | 833 | 0 | 833 | 548.51 | 334 | 495 | 1412 | 73 | 40.5 | 60.1 |
| C.auris | XP_028891915.2 dihydropteroate synthase Candida auris | 823 | 4.9e-169 | 823 | 505.753 | 324 | 452 | 1301 | 83 | 39.3 | 54.9 |
| C.albicans | XP_714207.2 trifunctional dihydropteroate synthetase/dihydroxymethylpterin pyrophosphokinase/dihydroneopterin aldolase Candida albicans SC5314 | 829 | 4.5e-175 | 829 | 522.702 | 322 | 494 | 1345 | 72 | 39.1 | 60.0 |
| A.fumigatus | XP_755317.1 folic acid synthesis protein Aspergillus fumigatus Af293 | 527 | 6.1e-104 | 527 | 327.405 | 206 | 290 | 838 | 79 | 25.0 | 35.2 |

Table S23: Pairwise alignment info from yeast Fol1 (DEG20010889), cf. Figure S134.

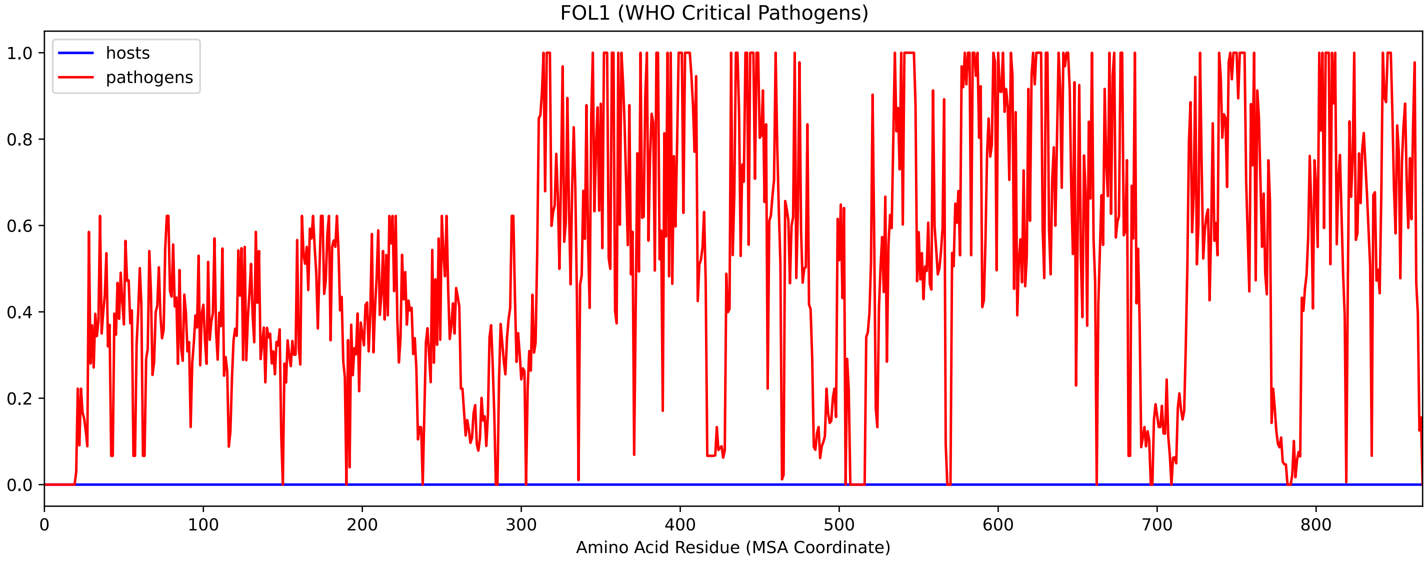


Figure S136: Sneath Similarity of Fol1 for WHO Critical Pathogens, cf. Figure S134

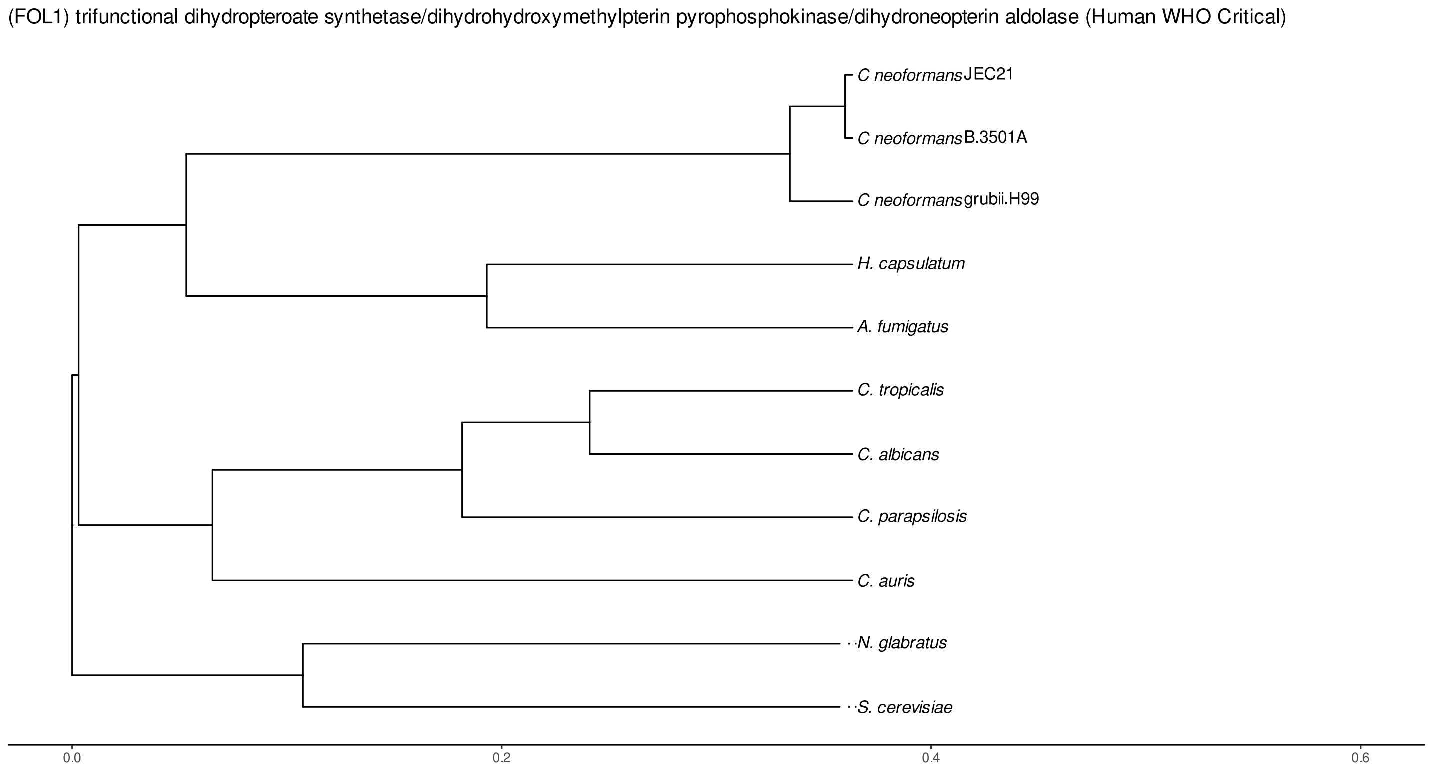


Figure S137: Phylogeny of hits for Fol1 made using clustal omega and visualized using ggtree, cf. Figures S134 and S135.

##### S4.9.2 Top 10 Agricultural Fungal Pathogens

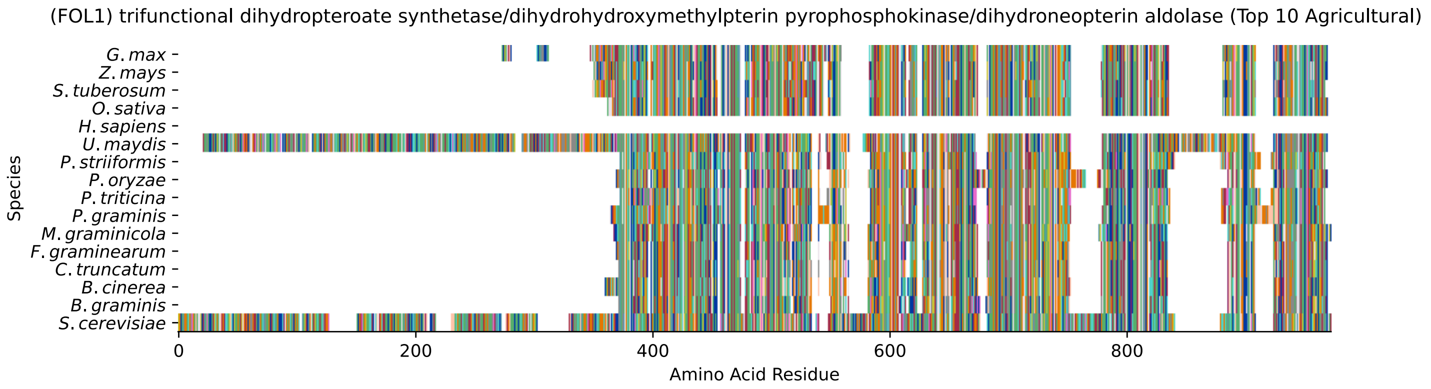


Figure S138: Multiple sequence alignment of yeast Fol1 (Top 10 Agricultural Fungal Pathogens). Cf. Figure S139 for alignment quality, and Figure S140 for Sneath similarity. Cf. Table S24 for protein names, and pairwise alignment metrics with yeast Fol1.

### Fol1 MSA Quality

| | *B. g.* | *B. c.* | *C. t.* | *F. g.* | *M. g.* | *P. g.* | *P. t.* | *P. o.* | *P. s.* | *U. m.* | *S. c.* |
|---|---|---|---|---|---|---|---|---|---|---|---|
| *B. graminis* | 919.842 | 575.859 | 521.161 | 545.814 | 441.039 | 261.922 | 271.552 | 491.5 | 277.715 | 278.1 | 283.493 |
| *B. cinerea* | 544.273 | 942.184 | 619.772 | 622.083 | 488.034 | 285.419 | 300.442 | 581.252 | 295.049 | 330.102 | 323.553 |
| *C. truncatum* | 512.301 | 640.188 | 910.983 | 653.284 | 485.337 | 271.166 | 289.656 | 580.096 | 285.419 | 313.153 | 301.212 |
| *F. graminearum* | 528.865 | 637.876 | 648.662 | 905.975 | 498.819 | 284.648 | 291.967 | 586.645 | 291.967 | 300.442 | 299.286 |
| *M. graminicola* | 414.461 | 488.034 | 463.766 | 484.567 | 946.421 | 293.508 | 286.189 | 438.728 | 316.235 | 320.857 | 311.612 |
| *P. graminis* | 279.256 | 310.071 | 279.256 | 300.442 | 318.931 | 977.622 | 657.907 | 265.388 | 677.552 | 362.073 | 256.529 |
| *P. triticina* | 280.796 | 317.005 | 289.656 | 303.138 | 304.294 | 637.106 | 923.309 | 269.24 | 620.542 | 334.724 | 248.44 |
| *P. oryzae* | 502.671 | 624.78 | 603.594 | 614.379 | 479.559 | 269.626 | 279.641 | 962.214 | 289.271 | 295.819 | 302.368 |
| *P. striiformis* | 278.1 | 308.916 | 280.411 | 295.819 | 328.561 | 654.44 | 613.609 | 272.707 | 948.347 | 367.466 | 263.462 |
| *U. maydis* | 277.33 | 350.517 | 310.071 | 306.22 | 338.191 | 357.066 | 332.028 | 280.026 | 374.4 | 1731.07 | 326.25 |
| *S. cerevisiae* | 263.077 | 323.553 | 286.96 | 286.96 | 311.612 | 241.506 | 236.498 | 273.863 | 252.677 | 303.138 | 1704.88 |

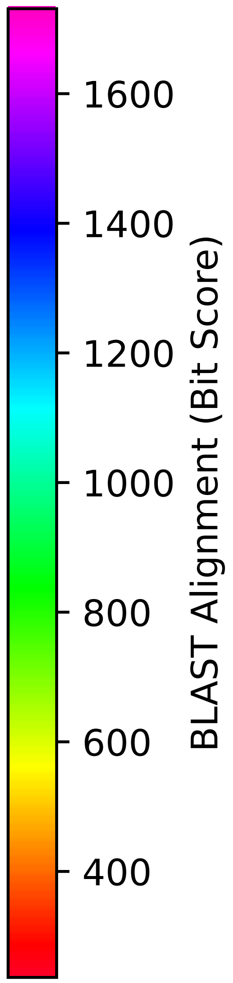


Figure S139: Multiple sequence alignment quality of Fol1 (Top 10 Agricultural Fungal Pathogens). Cf. Figures S138 and S141.

| Species | Hit Protein | Hit Length (a.a.) | evalue | align_len | bit_score | identity | positive | score | gaps | % identity | % positive |
|---|---|---|---|---|---|---|---|---|---|---|---|
| G.max | XP_003518782.1 folate synthesis bifunctional protein, mitochondrial Glycine max | 574 | 8.8e-75 | 574 | 254.218 | 180 | 290 | 648 | 74 | 21.8 | 35.2 |
| Z.mays | NP_001146490.1 Folate synthesis bifunctional protein, mitochondrial-like Zea mays | 546 | 2.9e-66 | 546 | 230.72 | 166 | 267 | 587 | 68 | 20.1 | 32.4 |
| S.tuberosum | XP_006348669.1 PREDICTED: folate synthesis bifunctional protein, mitochondrial-like Solanum tuberosum | 559 | 1.1e-72 | 559 | 247.284 | 181 | 301 | 630 | 86 | 22.0 | 36.5 |
| O.sativa | XP_015646629.1 folate synthesis bifunctional protein, mitochondrial isoform X6 Oryza sativa Japonica Group | 539 | 1.8e-72 | 539 | 247.669 | 173 | 274 | 631 | 76 | 21.0 | 33.3 |
| H.sapiens | - | - | - | - | - | - | - | - | - | - | - |
| U.maydis | XP_011388224.1 trifunctional dihydropteroate synthetase/dihydrohydroxymethylpterin pyrophosphokinase/dihydroneopterin aldolase FOL1 Ustilago maydis 521 | 923 | 1.1e-90 | 923 | 303.523 | 272 | 430 | 776 | 196 | 33.0 | 52.2 |
| P.striiformis | XP_047809565.1 hypothetical protein Pst134EA 007376 Puccinia striiformis f. sp. tritici | 529 | 1.2e-74 | 529 | 252.292 | 174 | 280 | 643 | 72 | 21.1 | 34.0 |
| P.oryzae | mRNA M BR32 EuGene 00010841-p1 — transcript=mRNA M BR32 EuGene 00010841 — gene=M BR32 EuGene 00010841 — organism=Pyricularia oryzae BR32 — gene product=unspecified product — transcript product=unspecified product — location=BR32 scaffold00001:3219749-3221562(+) — protein length=577 — sequence SO=supercontig — SO=protein coding gene — is pseudo=false | 539 | 2.2e-82 | 539 | 274.248 | 192 | 288 | 700 | 77 | 23.3 | 35.0 |
| P.triticina | XP_053018629.1 uncharacterized protein PtA15 3A441 Puccinia triticina | 530 | 2.3e-69 | 530 | 236.113 | 174 | 260 | 601 | 83 | 21.1 | 31.6 |
| P.graminis | XP_003324553.1 hypothetical protein PGTG 05359 Puccinia graminis f. sp. tritici CRL 75-36-700-3 | 546 | 5.7e-71 | 546 | 241.891 | 177 | 278 | 616 | 83 | 21.5 | 33.7 |
| M.graminicola | ZTRI 4.793.mRNA-p1 — transcript=ZTRI 4.793.mRNA — gene=ZTRI 4.793 — organism=Zymoseptoria tritici IPO323 — gene product=hypothetical protein — transcript product=hypothetical protein — location=Ztri chr 4:2544178-2550684(-) — protein length=2100 — sequence SO=chromosome — SO=protein coding gene — is pseudo=false | 533 | 2.5e-90 | 533 | 311.997 | 196 | 291 | 798 | 79 | 23.8 | 35.3 |
| F.graminearum | XP_011327992.1 hypothetical protein FGSG 09710 Fusarium graminearum PH-1 | 527 | 2.8e-87 | 527 | 286.189 | 192 | 275 | 731 | 83 | 23.3 | 33.4 |
| C.truncatum | XP_036583196.1 dihydropteroate synthase, partial Colletotrichum truncatum | 527 | 9.6e-87 | 527 | 287.345 | 195 | 271 | 734 | 86 | 23.7 | 32.9 |
| B.cinerea | XP_024553509.1 Bcfol1 Botrytis cinerea B05.10 | 542 | 2e-101 | 542 | 323.168 | 201 | 293 | 827 | 84 | 24.4 | 35.6 |
| B.graminis | VDB89524.1 — transcript=BGT96224V316 LOCUS5151 t1 — gene=BGT96224V316 LOCUS5151 — organism=Blumeria graminis f. sp. tritici 96224 — gene product=unspecified product — transcript product=unspecified product — location=LR026990:10590963-10592601(+) — protein length=513 — sequence SO=chromosome — SO=protein coding gene — is pseudo=false | 527 | 2.7e-79 | 527 | 263.077 | 191 | 275 | 671 | 85 | 23.2 | 33.4 |

Table S24: Pairwise alignment info from yeast Fol1 (DEG20010889), cf. Figure S138.

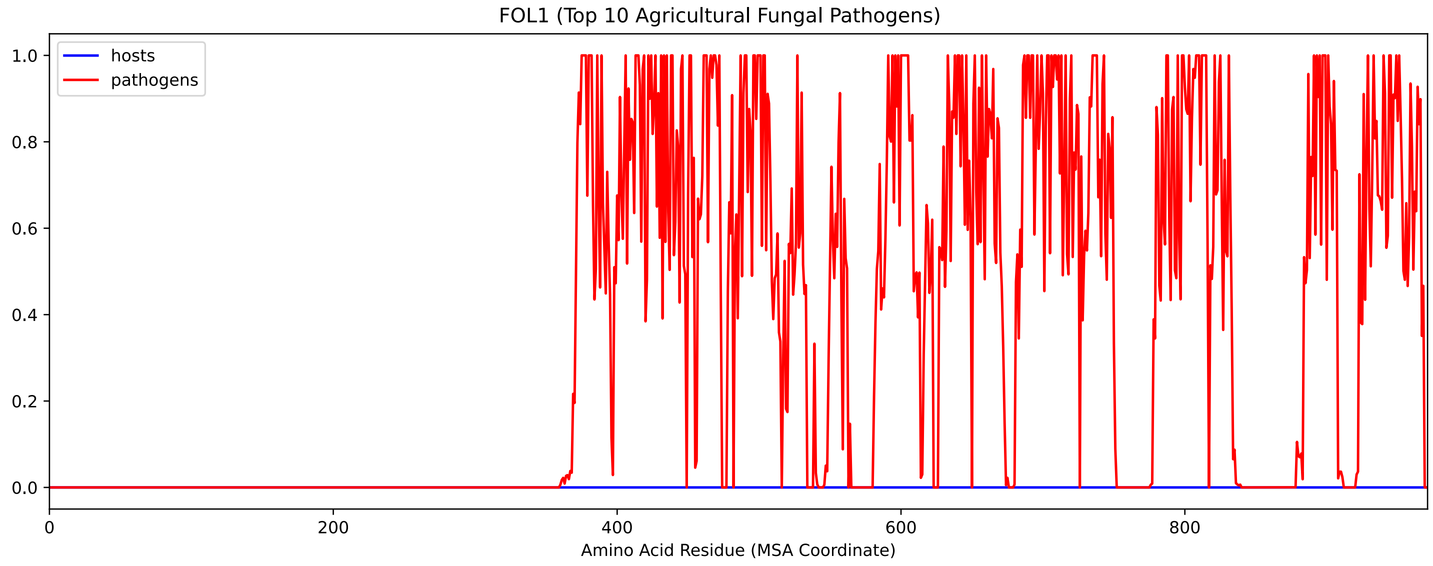


Figure S140: Sneath Similarity of Fol1 for Top 10 Agricultural Fungal Pathogens, cf. Figure S138

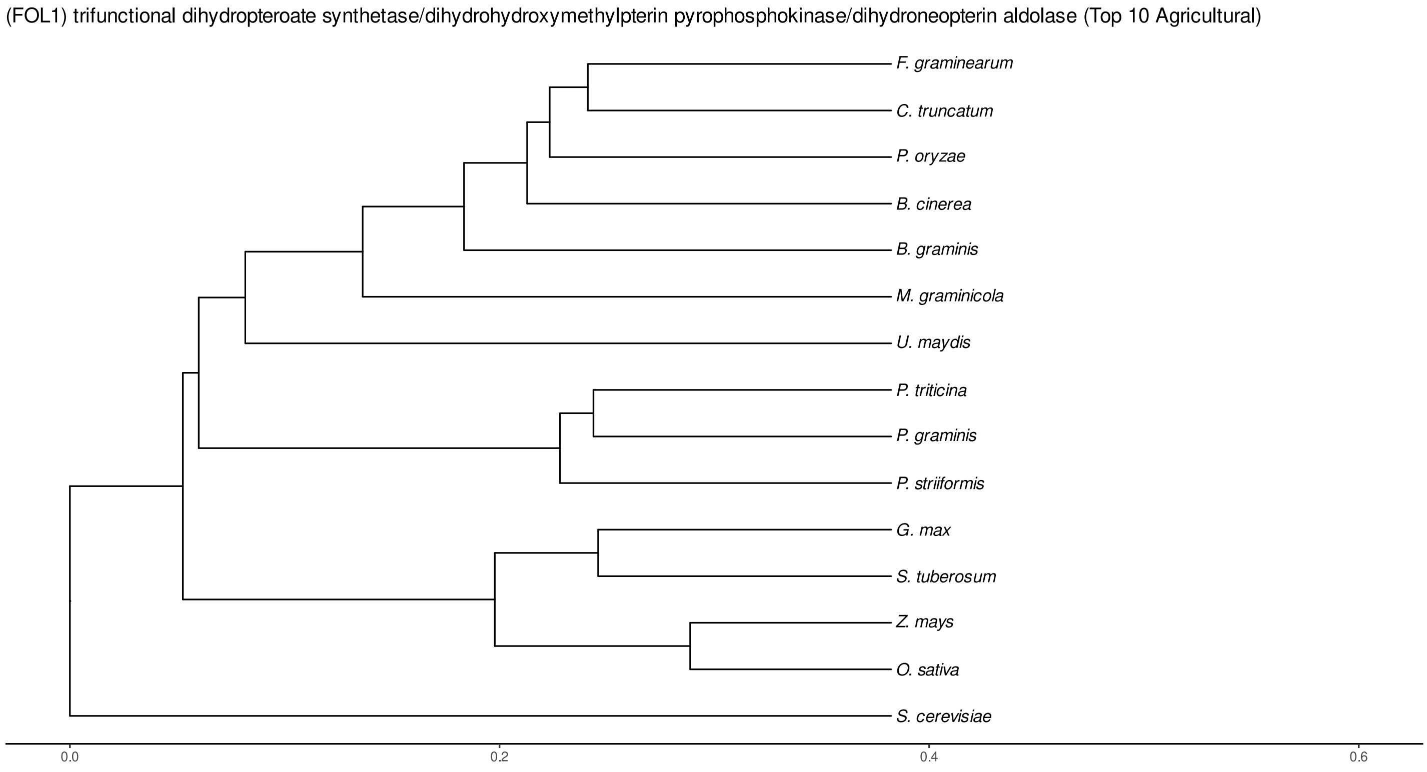


Figure S141: Phylogeny of hits for Fol1 made using clustal omega and visualized using ggtree, cf. Figures S138 and S139.

### S4.9.3 NR

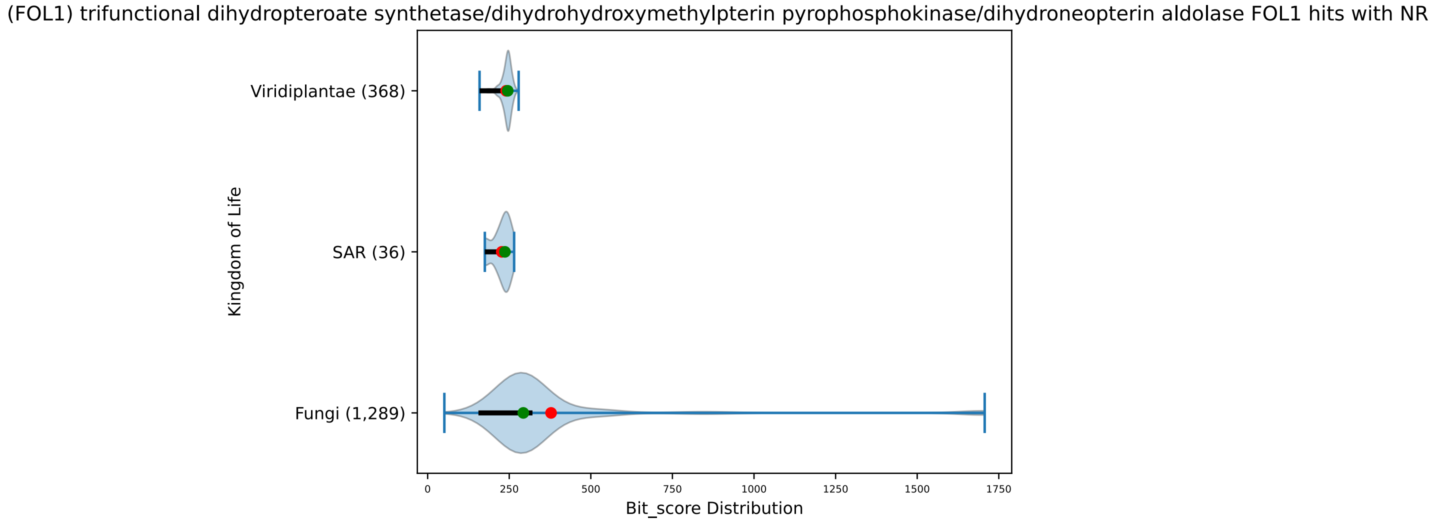


Figure S142: Non-redundant (NR) protein hits for DEG20010889/Fol1, with expectation value of no more than 0.1. Green points are medians, and red points are arithmetic means.

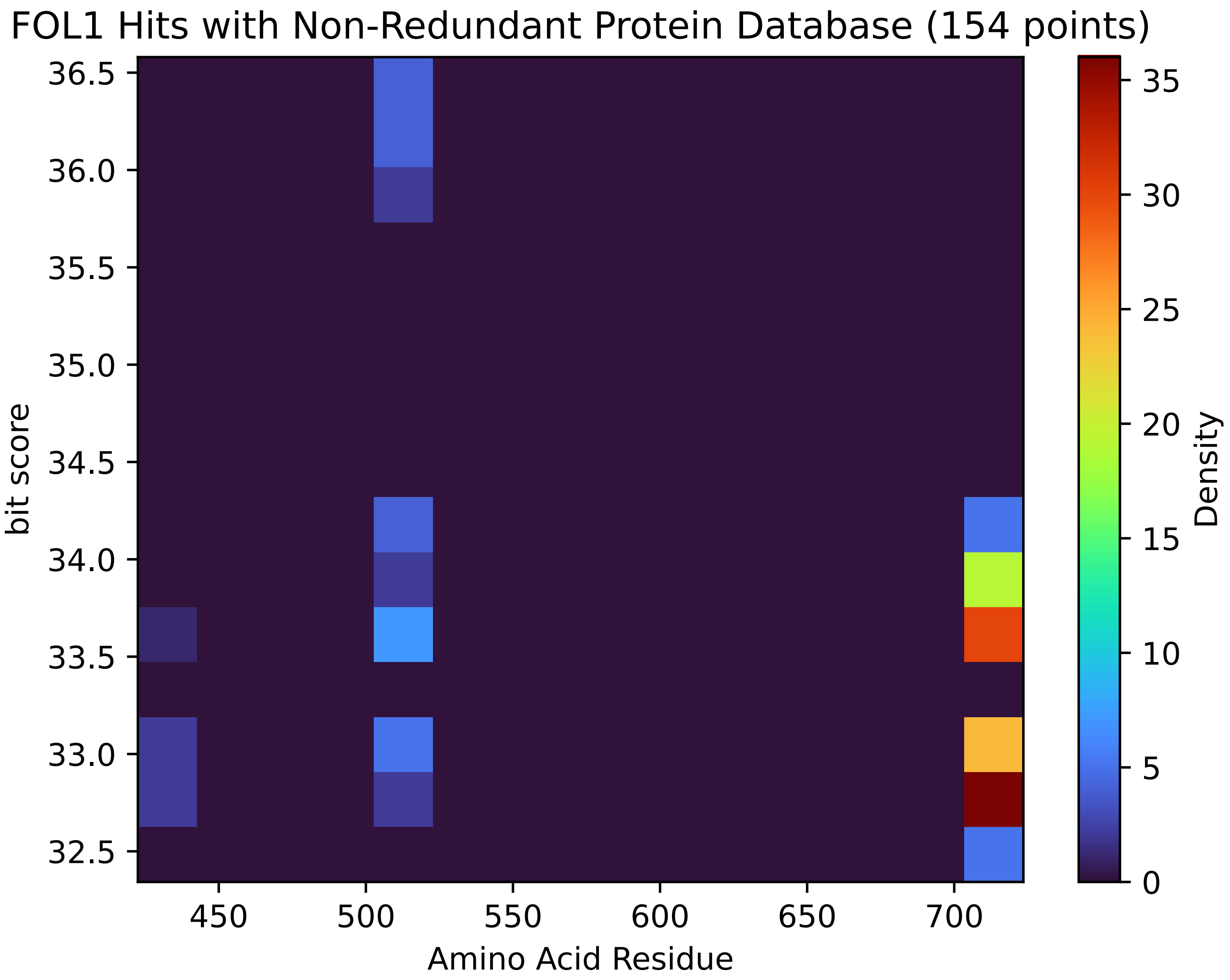


Figure S143: Non-redundant (NR) protein hits for Fol1 in the kingdom SAR.

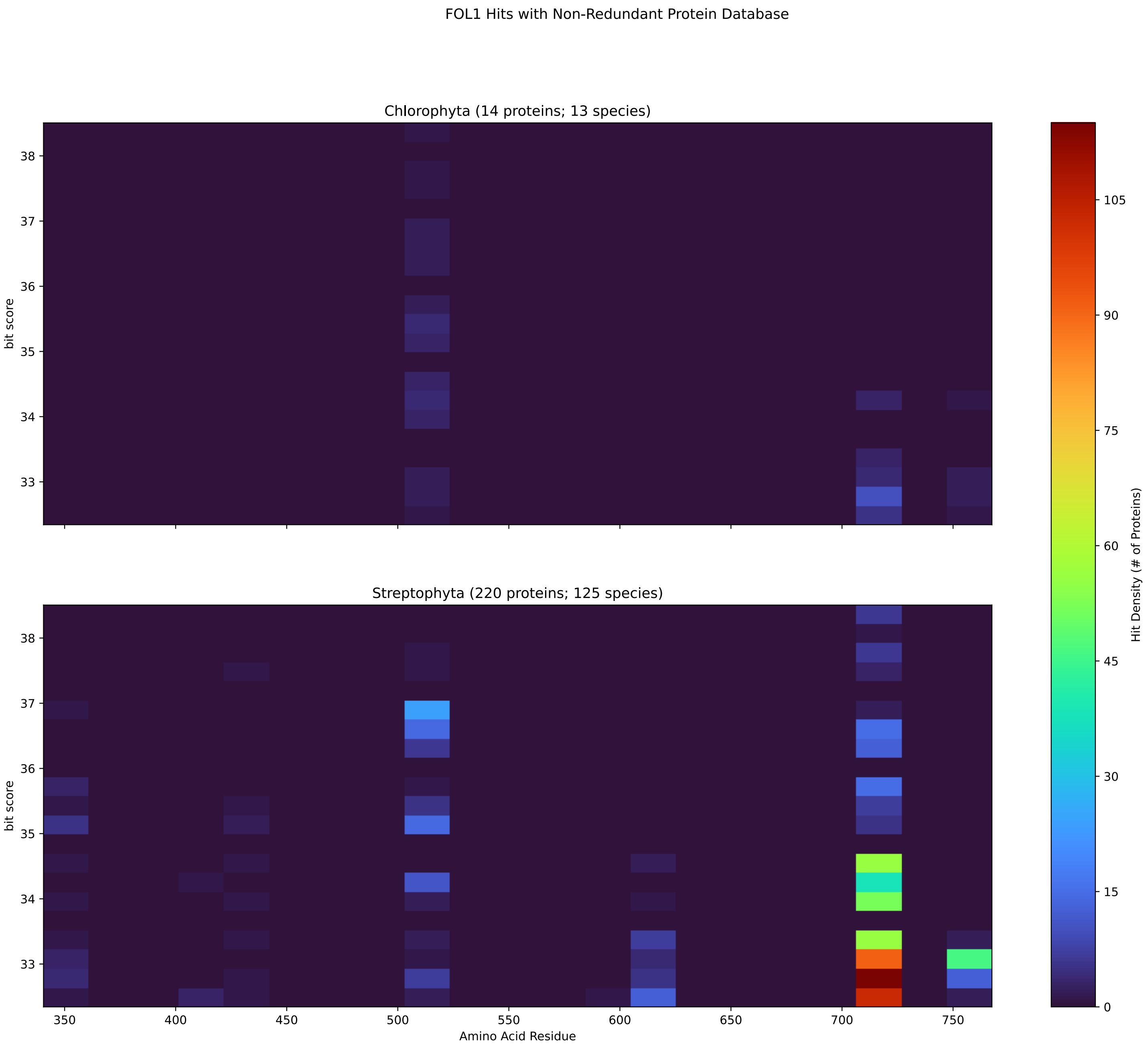


Figure S144: Non-redundant (NR) protein hits for Fol1 in the kingdom Viridiplantae.

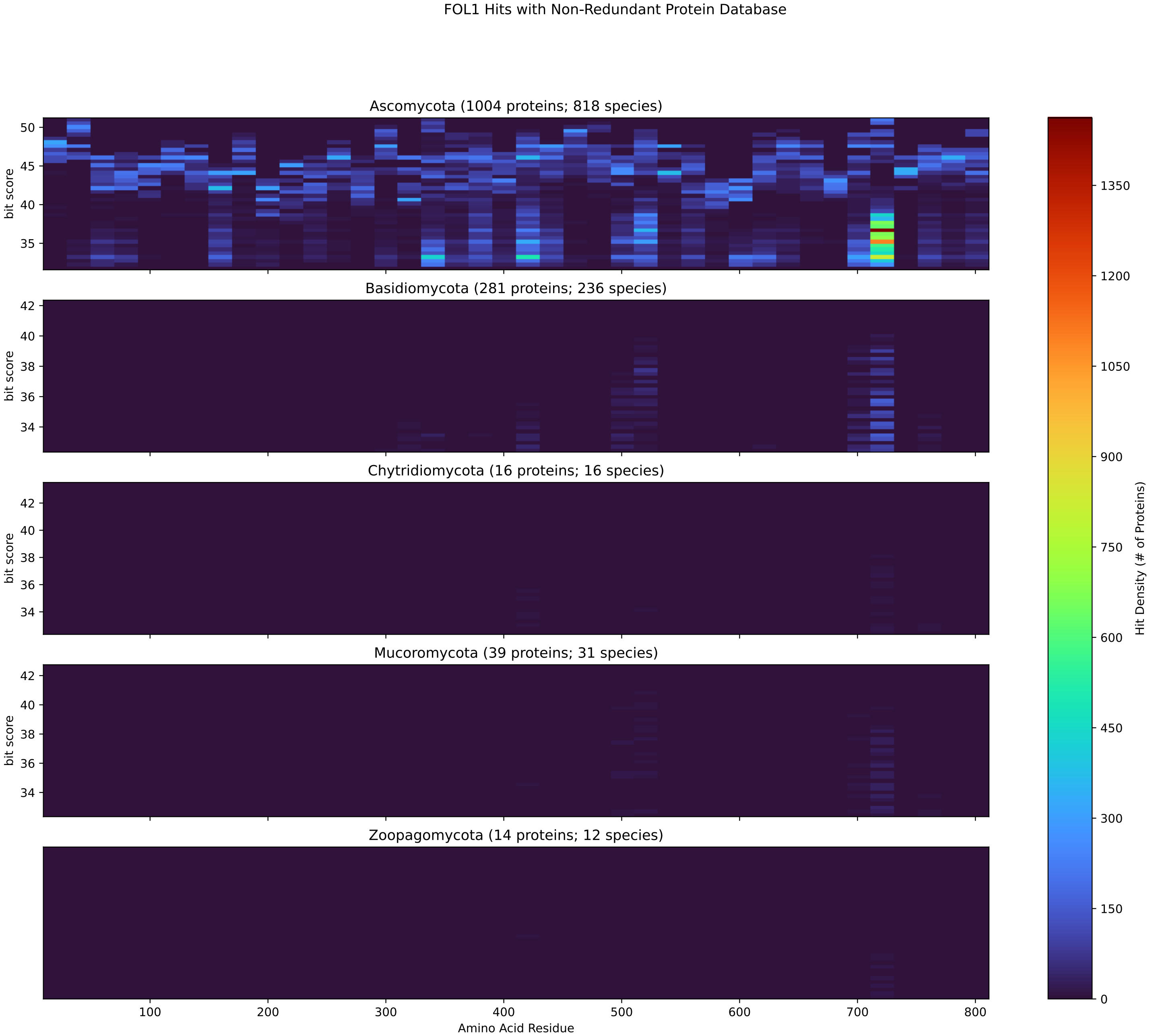


Figure S145: Non-redundant (NR) protein hits for Fol1 in the kingdom Fungi.

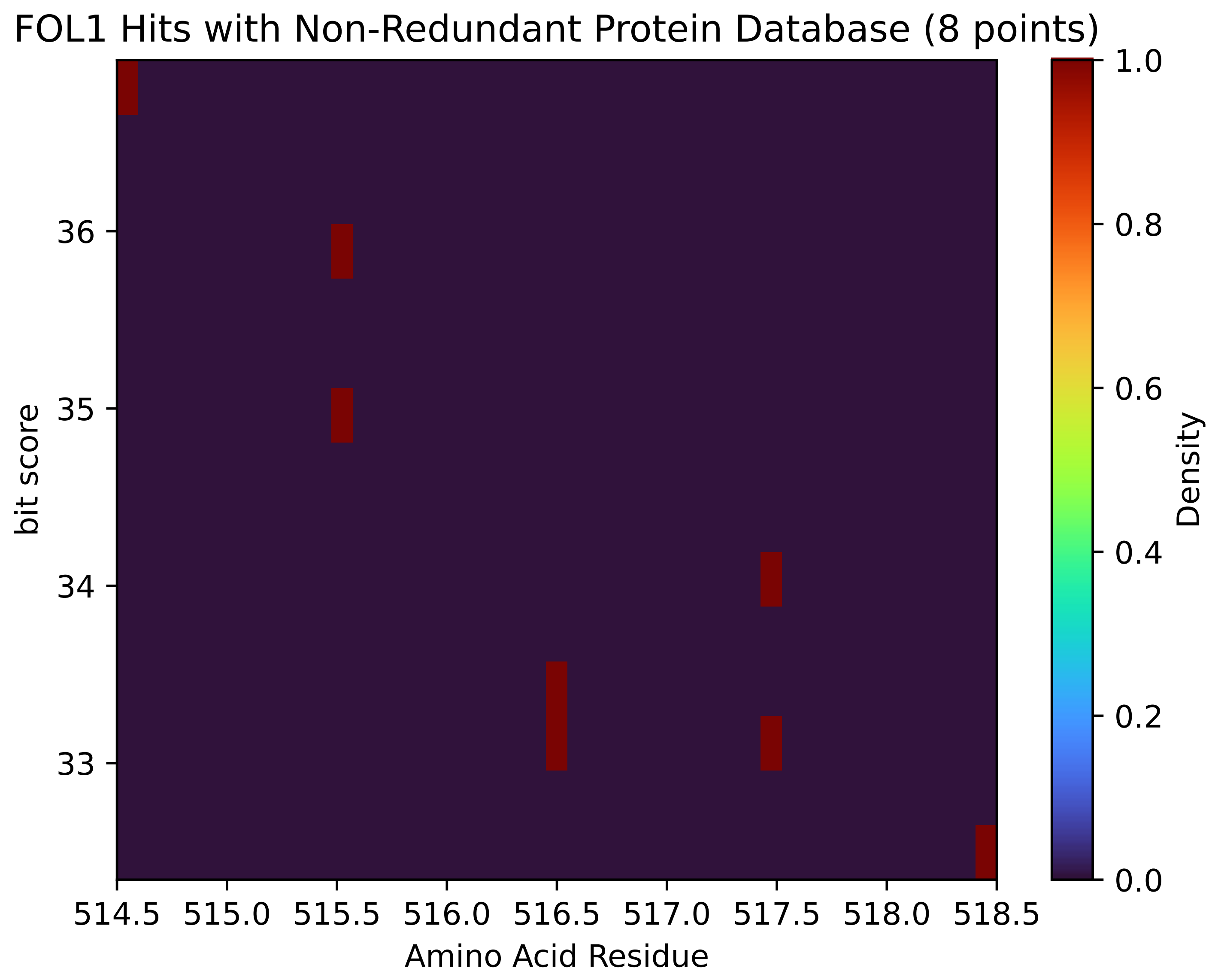


Figure S146: Non-redundant (NR) protein hits for Fol1 in the kingdom Metazoa.

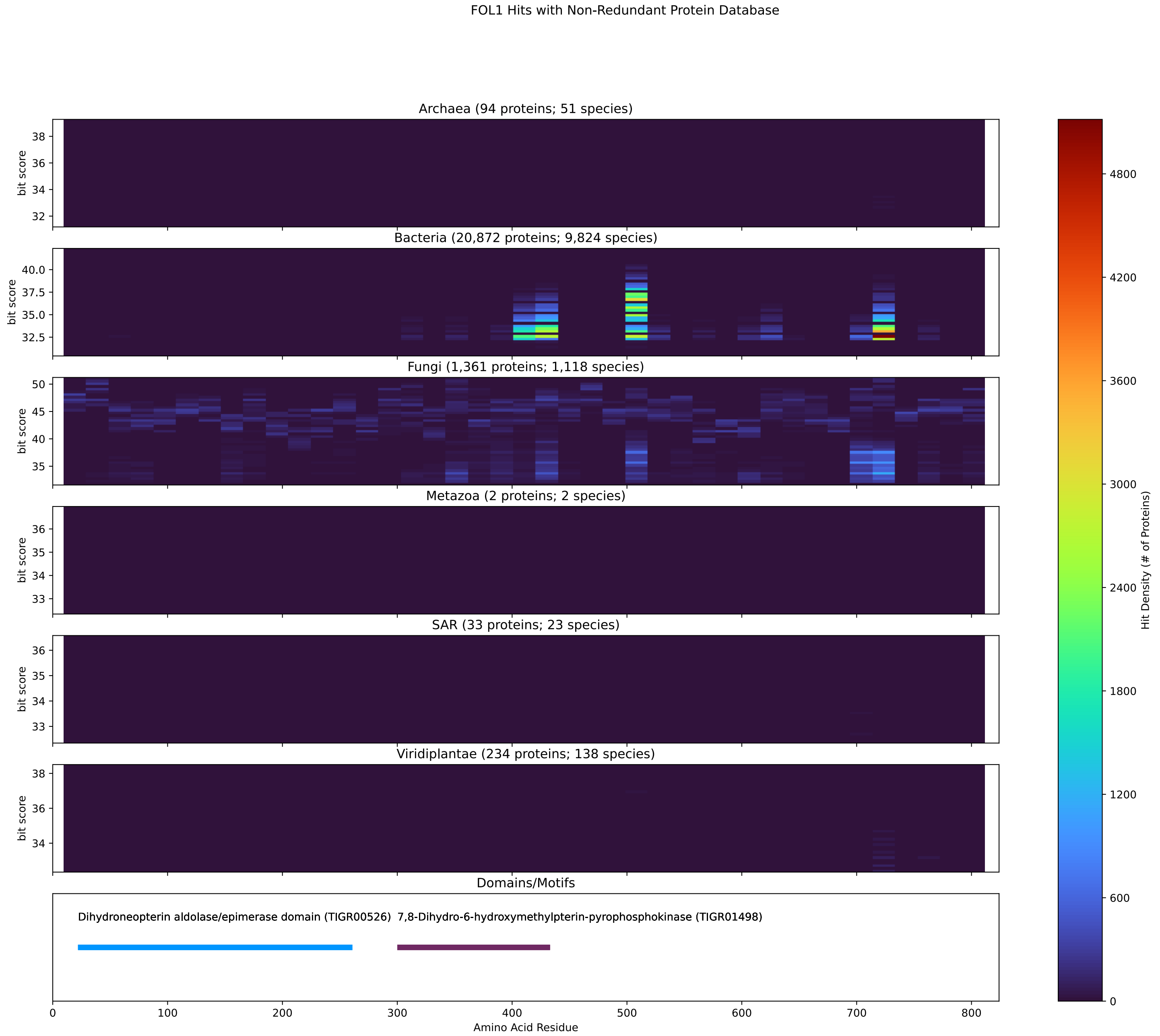


Figure S147: Non-redundant (NR) protein hits for DEG20010889/Fol1 at 20 amino acid length queries.

#### S4.10 Ilv3

##### S4.10.1 WHO Critical Pathogens

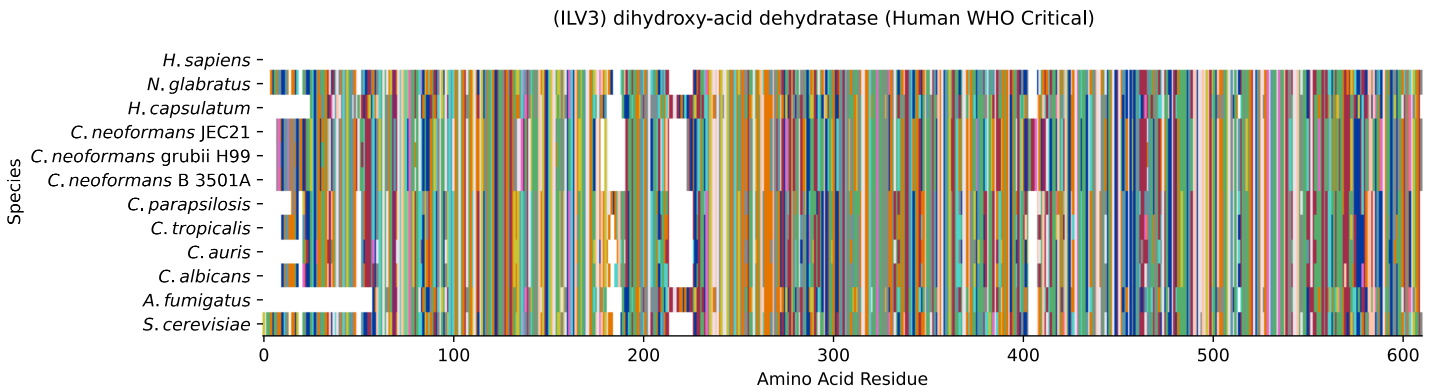


Figure S148: Multiple sequence alignment of yeast Ilv3 (WHO Critical Pathogens). Cf. Figure S149 for alignment quality, and Figure S150 for Sneath similarity. Cf. Table S25 for protein names, and pairwise alignment metrics with yeast Ilv3.

#### Ilv3 MSA Quality

| | *A. f.* | *C. a.* | *C. a.* | *C. p.* | *C. t.* | *C. n.* | *C. n.* | *C. n.* | *H. c.* | *N. g.* | *S. c.* |
|---|---|---|---|---|---|---|---|---|---|---|---|
| *A. fumigatus* | 1112.06 | 611.683 | 623.239 | 620.157 | 616.305 | 586.26 | 588.186 | 586.26 | 960.674 | 618.616 | 617.461 |
| *C. albicans* | 597.045 | 1187.17 | 989.564 | 1038.1 | 1115.52 | 681.019 | 683.33 | 681.019 | 626.32 | 897.886 | 882.863 |
| *C. auris* | 608.986 | 989.564 | 1164.06 | 987.638 | 993.801 | 673.315 | 676.011 | 673.315 | 646.351 | 893.264 | 877.47 |
| *C. parapsilosis* | 605.134 | 1038.1 | 987.638 | 1180.24 | 1039.25 | 669.078 | 673.315 | 669.078 | 640.188 | 877.856 | 879.782 |
| *C. tropicalis* | 600.897 | 1115.52 | 993.801 | 1039.25 | 1186.02 | 690.263 | 694.115 | 690.263 | 632.098 | 889.797 | 885.56 |
| *C. neoformans* B 3501A | 566.229 | 681.019 | 673.315 | 669.078 | 690.263 | 1204.5 | 1190.64 | 1204.5 | 600.127 | 681.019 | 691.034 |
| *C. neoformans* grubii H99 | 568.155 | 683.33 | 676.011 | 673.315 | 694.115 | 1190.64 | 1205.66 | 1190.64 | 602.823 | 685.641 | 692.96 |
| *C. neoformans* JEC21 | 566.229 | 681.019 | 673.315 | 669.078 | 690.263 | 1204.5 | 1190.64 | 1204.5 | 600.127 | 681.019 | 691.034 |
| *H. capsulatum* | 936.406 | 626.32 | 646.351 | 640.188 | 632.098 | 600.127 | 602.823 | 600.127 | 1181.39 | 622.468 | 634.41 |
| *N. glabratus* | 604.364 | 897.886 | 893.264 | 877.856 | 889.797 | 681.019 | 685.641 | 681.019 | 622.468 | 1202.58 | 1050.04 |
| *S. cerevisiae* | 602.823 | 882.863 | 877.47 | 879.782 | 885.56 | 691.034 | 692.96 | 691.034 | 634.41 | 1050.04 | 1210.28 |

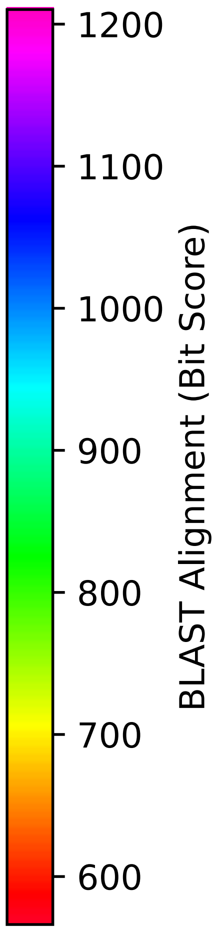


Figure S149: Multiple sequence alignment quality of Ilv3 (WHO Critical Pathogens). Cf. Figures S148 and S151.

| Species | Hit Protein | Hit Length (a.a.) | evalue | align_len | bit_score | identity | positive | score | gaps | % identity | % positive |
|---|---|---|---|---|---|---|---|---|---|---|---|
| H.sapiens | - | - | - | - | - | - | - | - | - | - | - |
| N.glabratus | XP_445144.1 uncharacterized protein CAGL0B03993g Nakaseomyces glabratus | 581 | 0 | 581 | 1051.2 | 495 | 538 | 2717 | 0 | 84.6 | 92.0 |
| H.capsulatum | XP_045291535.1 dihydroxy-acid dehydratase Histoplasma capsulatum G186AR | 576 | 0 | 576 | 634.024 | 313 | 406 | 1634 | 19 | 53.5 | 69.4 |
| C.neoformans.JEC21 | XP_572335.1 dihydroxy-acid dehydratase, putative Cryptococcus neoformans var. neoformans JEC21 | 590 | 0 | 590 | 692.189 | 341 | 433 | 1785 | 18 | 58.3 | 74.0 |
| C.neoformans.grubii.H99 | XP_012053809.1 dihydroxy-acid dehydratase Cryptococcus neoformans var. grubii H99 | 590 | 0 | 590 | 694.115 | 343 | 437 | 1790 | 18 | 58.6 | 74.7 |
| C.neoformans.B.3501A | XP_772279.1 hypothetical protein CNBL1470 Cryptococcus neoformans var. neoformans B-3501A | 590 | 0 | 590 | 692.189 | 341 | 433 | 1785 | 18 | 58.3 | 74.0 |
| C.parapsilosis | XP_036663159.1 uncharacterized protein CPAR2 100130 Candida parapsilosis | 575 | 0 | 575 | 880.167 | 420 | 486 | 2273 | 5 | 71.8 | 83.1 |
| C.tropicalis | XP_002546669.1 dihydroxy-acid dehydratase, mitochondrial precursor Candida tropicalis MYA--3404 | 577 | 0 | 577 | 884.789 | 420 | 490 | 2285 | 3 | 71.8 | 83.8 |
| C.auris | XP_028892424.1 dihydroxy-acid dehydratase, mitochondrial Candida auris | 566 | 0 | 566 | 878.241 | 410 | 486 | 2268 | 0 | 70.1 | 83.1 |
| C.albicans | XP_721948.1 dihydroxy-acid dehydratase Candida albicans SC5-314 | 577 | 0 | 577 | 882.093 | 418 | 486 | 2278 | 3 | 71.5 | 83.1 |
| A.fumigatus | XP_750105.1 mitochondrial dihydroxy acid dehydratase, putative Aspergillus fumigatus Af29-3 | 543 | 0 | 543 | 602.438 | 301 | 385 | 1552 | 17 | 51.5 | 65.8 |

Table S25: Pairwise alignment info from yeast Ilv3 (DEG20010579), cf. Figure S148.

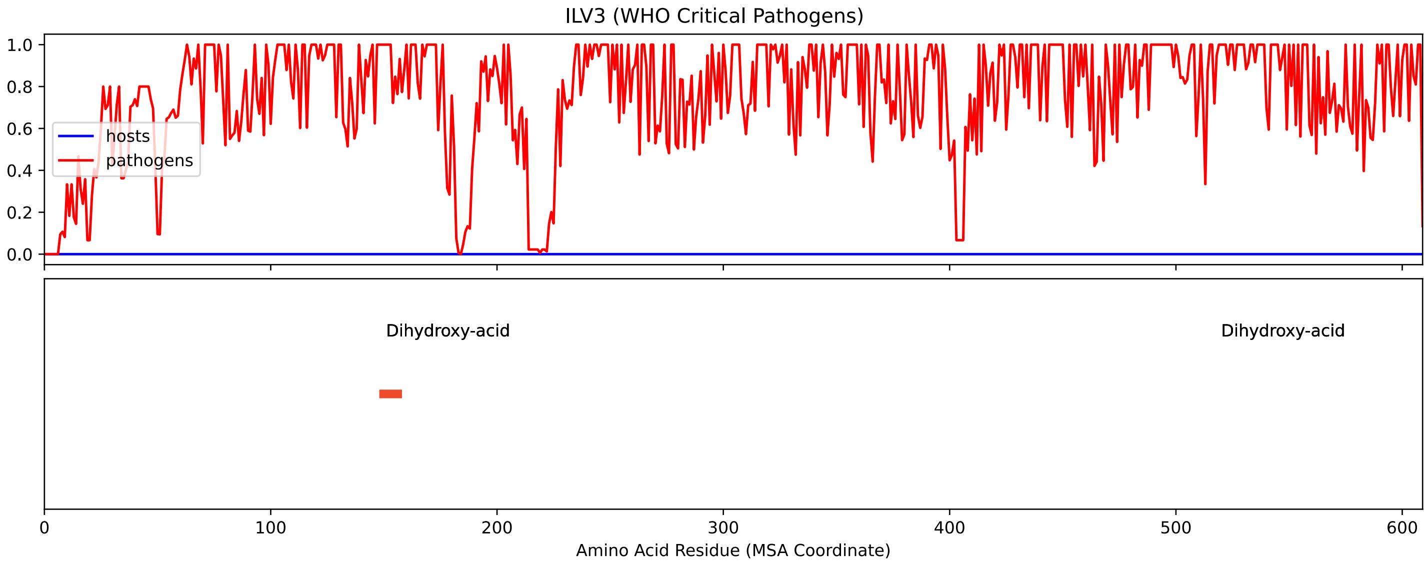


Figure S150: Sneath Similarity of Ilv3 for WHO Critical Pathogens, cf. Figure S148

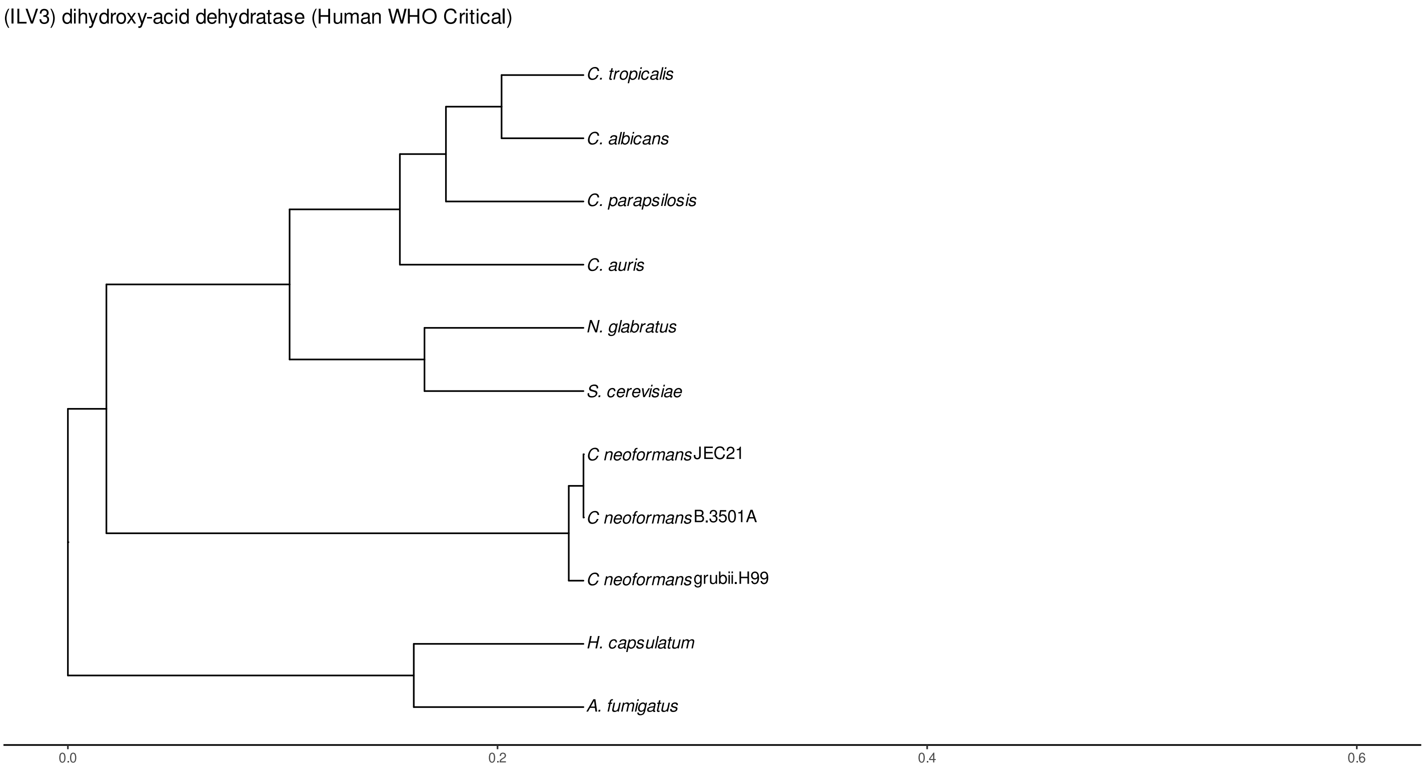


Figure S151: Phylogeny of hits for Ilv3 made using clustal omega and visualized using ggtree, cf. Figures S148 and S149.

##### S4.10.2 Top 10 Agricultural Fungal Pathogens

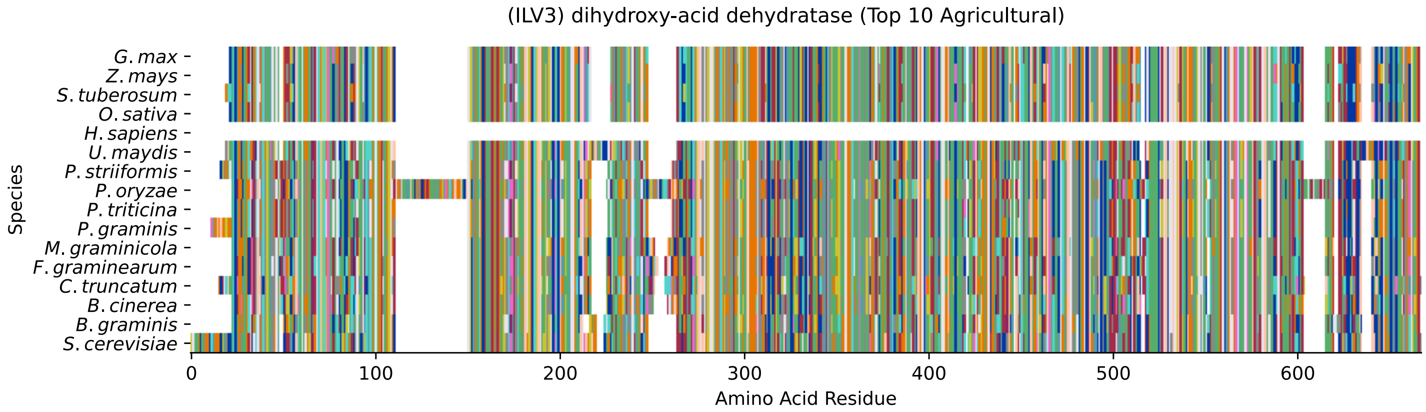


Figure S152: Multiple sequence alignment of yeast Ilv3 (Top 10 Agricultural Fungal Pathogens). Cf. Figure S153 for alignment quality, and Figure S154 for Sneath similarity. Cf. Table S26 for protein names, and pairwise alignment metrics with yeast Ilv3.

### Ilv3 MSA Quality

| | B.g. | B.c. | C.t. | F.g. | M.g. | P.g. | P.t. | P.o. | P.s. | U.m. | S.c. |
|---|---|---|---|---|---|---|---|---|---|---|---|
| *B. graminis* | 1144.8 | 629.787 | 583.948 | 639.032 | 614.764 | 682.559 | 685.256 | 609.372 | 680.248 | 631.328 | 754.977 |
| *B. cinerea* | 629.787 | 1174.07 | 759.985 | 880.937 | 867.84 | 577.4 | 577.015 | 873.233 | 572.778 | 609.372 | 620.928 |
| *C. truncatum* | 597.816 | 776.933 | 1193.33 | 765.763 | 811.216 | 561.607 | 557.37 | 717.227 | 558.91 | 584.334 | 605.134 |
| *F. graminearum* | 639.032 | 880.937 | 749.584 | 1182.16 | 824.313 | 621.698 | 632.098 | 917.531 | 620.542 | 627.091 | 639.032 |
| *M. graminicola* | 614.764 | 867.84 | 793.112 | 824.313 | 1170.61 | 595.119 | 590.497 | 792.727 | 593.578 | 584.719 | 622.854 |
| *P. graminis* | 702.975 | 598.201 | 569.311 | 643.654 | 616.305 | 1167.14 | 1076.62 | 597.816 | 1108.98 | 621.698 | 681.789 |
| *P. triticina* | 706.442 | 598.971 | 563.918 | 653.67 | 611.298 | 1076.62 | 1145.18 | 595.89 | 1106.28 | 633.254 | 692.189 |
| *P. oryzae* | 609.372 | 873.233 | 705.286 | 917.531 | 792.727 | 577.015 | 574.704 | 1287.71 | 571.622 | 579.711 | 593.578 |
| *P. striiformis* | 701.049 | 593.578 | 565.844 | 640.958 | 615.535 | 1108.98 | 1106.28 | 592.808 | 1153.66 | 622.854 | 682.174 |
| *U. maydis* | 631.328 | 609.372 | 569.311 | 627.091 | 584.719 | 599.742 | 610.912 | 579.711 | 600.512 | 1181.01 | 634.41 |
| *S. cerevisiae* | 754.977 | 620.928 | 593.193 | 639.032 | 622.854 | 663.685 | 673.315 | 593.578 | 663.3 | 634.41 | 1210.28 |

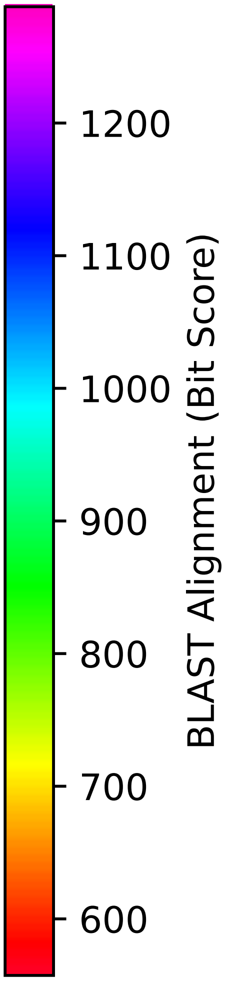


Figure S153: Multiple sequence alignment quality of Ilv3 (Top 10 Agricultural Fungal Pathogens). Cf. Figures S152 and S155.

| Species | Hit Protein | Hit Length (a.a.) | evalue | align_len | bit_score | identity | positive | score | gaps | % identity | % positive |
|---|---|---|---|---|---|---|---|---|---|---|---|
| G.max | NP_001276144.1 putative dihydroxy-acid dehydratase, mitochondrial-like Glycine max | 564 | 0 | 564 | 686.797 | 335 | 416 | 1771 | 9 | 57.3 | 71.1 |
| Z.mays | NP_001141560.1 Dihydroxy-acid dehydratase chloroplastic Zea mays | 564 | 0 | 564 | 682.945 | 335 | 415 | 1761 | 9 | 57.3 | 70.9 |
| S.tuberosum | XP_006346583.1 PREDICTED: dihydroxy-acid dehydratase, chloroplastic-like Solanum tuberosum | 566 | 0 | 566 | 679.863 | 331 | 418 | 1753 | 9 | 56.6 | 71.5 |
| O.sativa | XP_015649747.1 dihydroxy-acid dehydratase, chloroplastic Oryza sativa Japonica Group | 564 | 0 | 564 | 689.108 | 334 | 420 | 1777 | 9 | 57.1 | 71.8 |
| H.sapiens | - | - | - | - | - | - | - | - | - | - | - |
| U.maydis | XP_011389372.1 putative dihydroxy-acid dehydratase Ustilago maydis 521 | 577 | 0 | 577 | 634.41 | 315 | 401 | 1635 | 15 | 53.8 | 68.5 |
| P.striiformis | XP_047805603.1 hypothetical protein Pst134EA 015851 Puccinia striiformis f. sp. tritici | 573 | 0 | 573 | 662.529 | 336 | 414 | 1708 | 10 | 57.4 | 70.8 |
| P.oryzae | mRNA M BR32 EuGene 00030511-p1 — transcript=mRNA M BR32 EuGene 00030511 — gene=M BR32 EuGene 00030511 — organism=Pyricularia oryzae BR32 — gene product=unspecified product — transcript product=unspecified product — location=BR32 scaffold00002:4030791-4032964(+) — protein length=660 — sequence SO=supercontig — SO=protein coding gene — is pseudo=false | 631 | 0 | 631 | 592.808 | 311 | 400 | 1527 | 74 | 53.2 | 68.4 |
| P.triticina | XP_053023065.1 uncharacterized protein PtA15 8A414 Puccinia triticina | 567 | 0 | 567 | 673.315 | 341 | 415 | 1736 | 10 | 58.3 | 70.9 |
| P.graminis | XP_003327990.2 dihydroxy-acid dehydratase Puccinia graminis f. sp. tritici CRL 75-36-700–3 | 578 | 0 | 578 | 663.685 | 337 | 417 | 1711 | 10 | 57.6 | 71.3 |
| M.graminicola | ZTRI 1.138.mRNA-p1 — transcript=ZTRI 1.138.mRNA — gene=ZTRI 1.138 — organism=Zymoseptoria tritici IPO323 — gene product=similar to dihydroxy-acid dehydratase — transcript product=similar to dihydroxy-acid dehydratase — location=Ztri chr 1:550543-552333(+) — protein length=596 — sequence SO=chromosome — SO=protein coding gene — is pseudo=false | 573 | 0 | 573 | 622.468 | 310 | 398 | 1604 | 16 | 53.0 | 68.0 |
| F.graminearum | XP_011318673.1 dihydroxy-acid dehydratase Fusarium graminearum PH-1 | 577 | 0 | 577 | 638.262 | 317 | 405 | 1645 | 20 | 54.2 | 69.2 |
| C.truncatum | XP_036575211.1 dihydroxy-acid dehydratase Colletotrichum truncatum | 583 | 0 | 583 | 592.423 | 304 | 410 | 1526 | 18 | 52.0 | 70.1 |
| B.cinerea | XP_024553827.1 hypothetical protein BCIN 16g02700 Botrytis cinerea B05.10 | 572 | 0 | 572 | 619.387 | 303 | 401 | 1596 | 15 | 51.8 | 68.5 |
| B.graminis | VDB86357.1 — transcript=BGT96224V316 LOCUS3920 t1 — gene=BGT96224V316 LOCUS3920 — organism=Blumeria graminis f. sp. tritici 96224 — gene product=unspecified product — transcript product=unspecified product — location=LR026988:19021675-19023529(+) — protein length=601 — sequence SO=chromosome — SO=protein coding gene — is pseudo=false | 564 | 0 | 564 | 754.592 | 364 | 444 | 1947 | 5 | 62.2 | 75.9 |

Table S26: Pairwise alignment info from yeast Ilv3 (DEG20010579), cf. Figure S152.

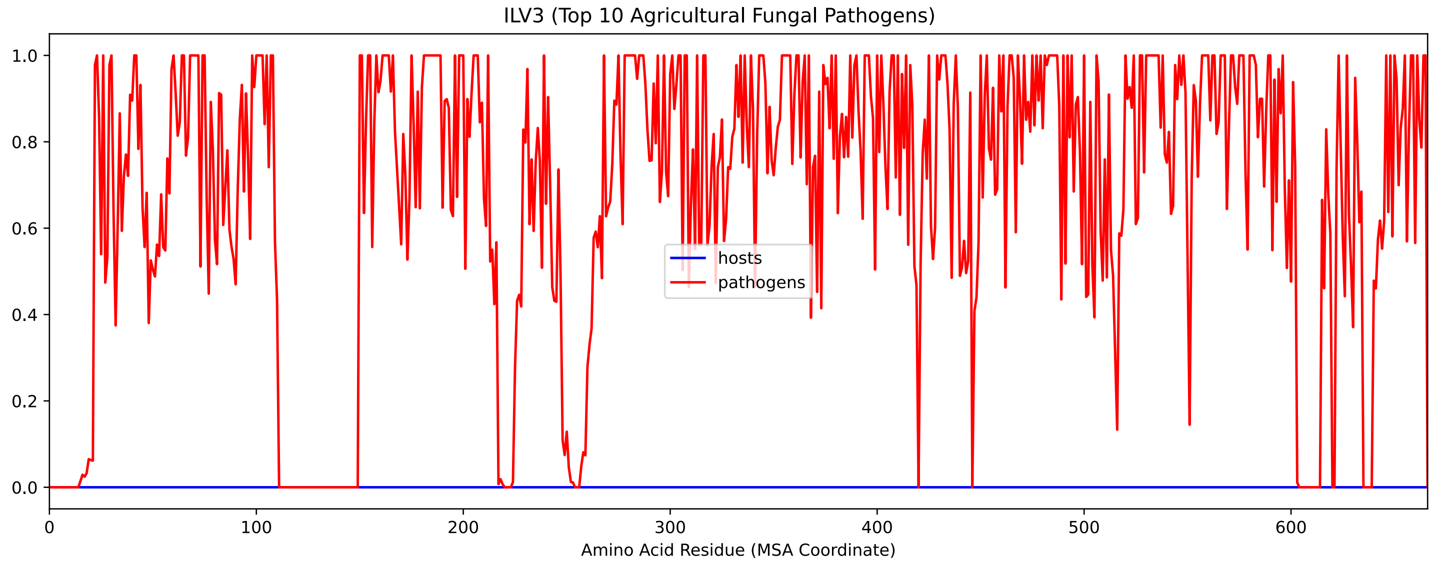


Figure S154: Sneath Similarity of Ilv3 for Top 10 Agricultural Fungal Pathogens, cf. Figure S152

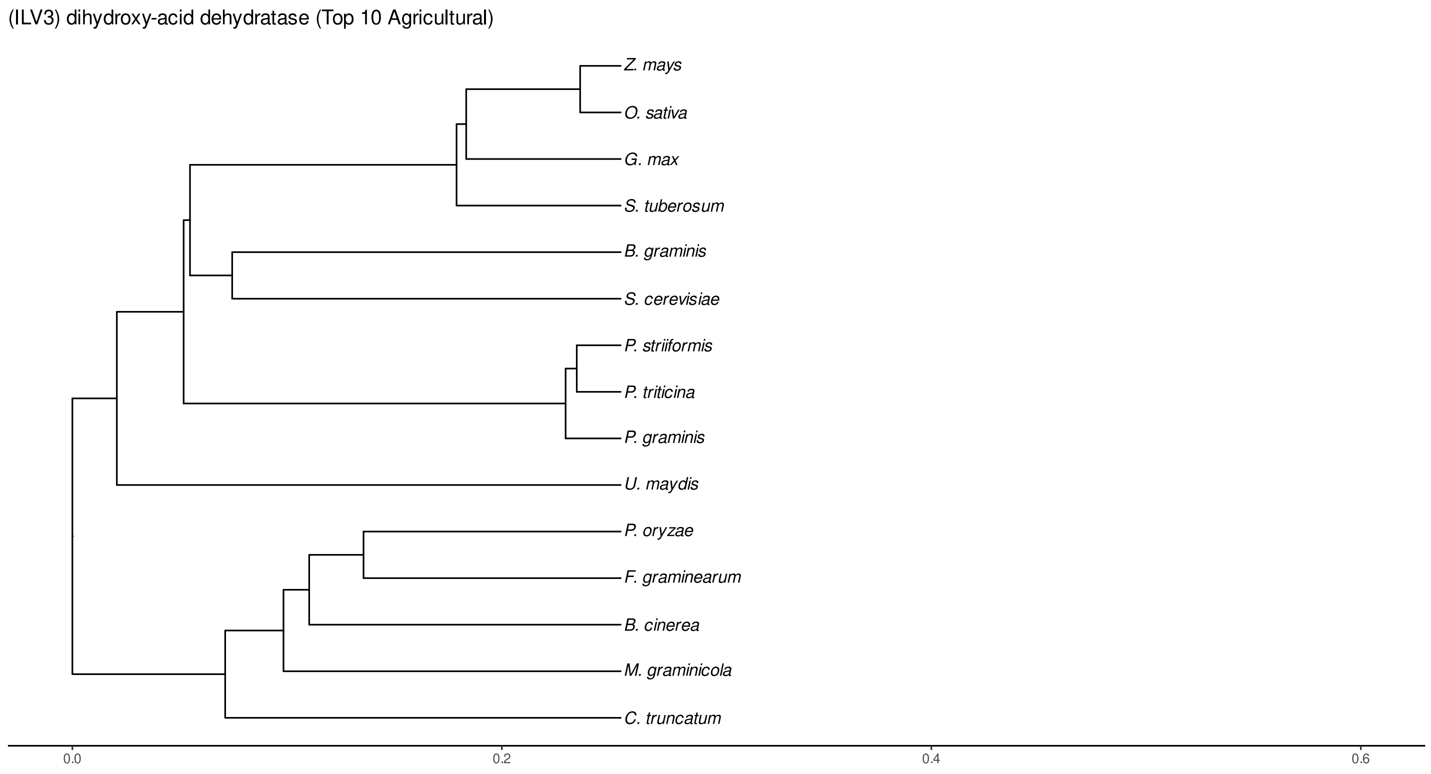


Figure S155: Phylogeny of hits for Ilv3 made using clustal omega and visualized using ggtree, cf. Figures S152 and S153.

### S4.10.3 NR

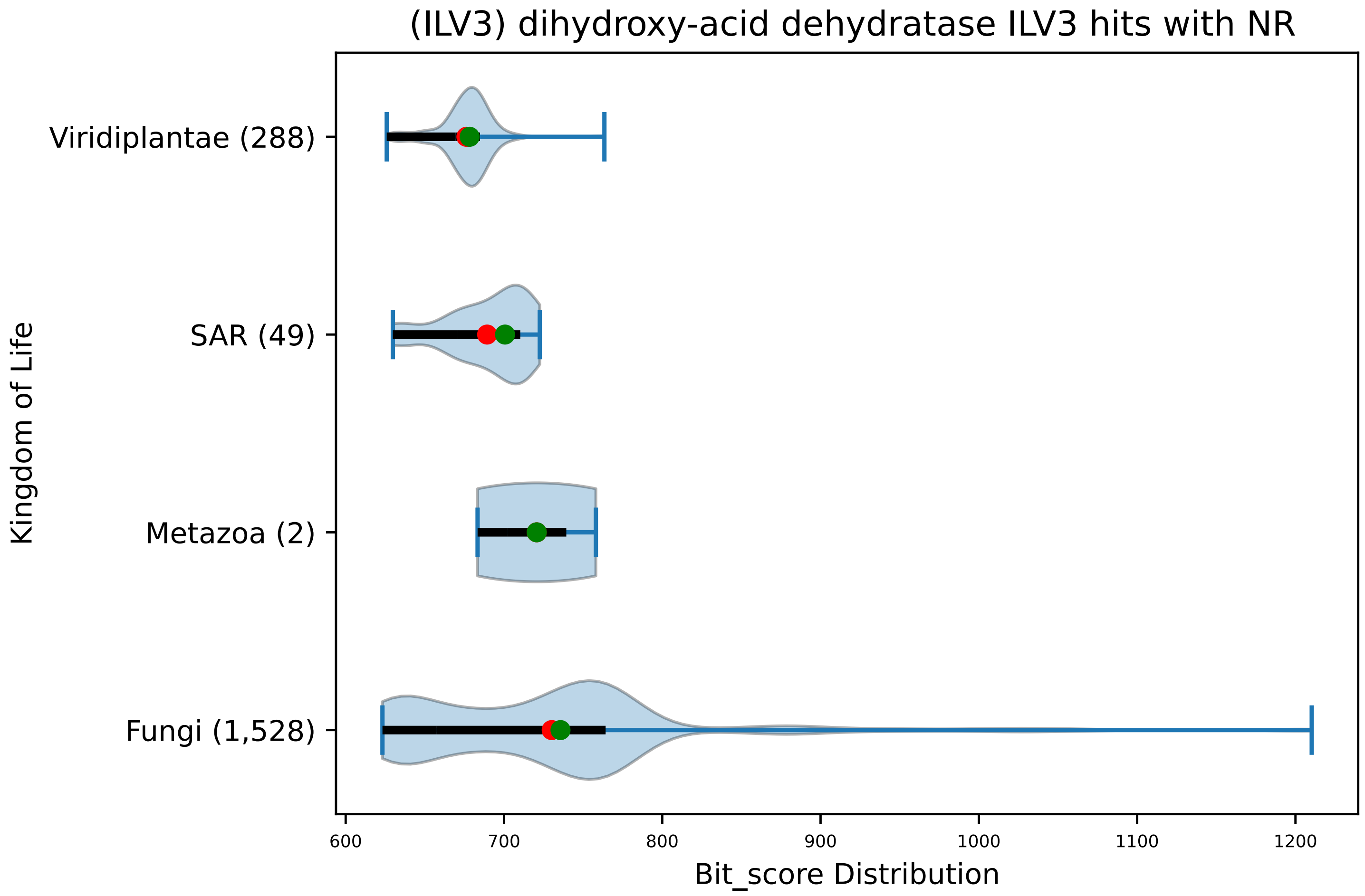


Figure S156: Non-redundant (NR) protein hits for DEG20010579/Ilv3, with expectation value of no more than 0.1. Green points are medians, and red points are arithmetic means.

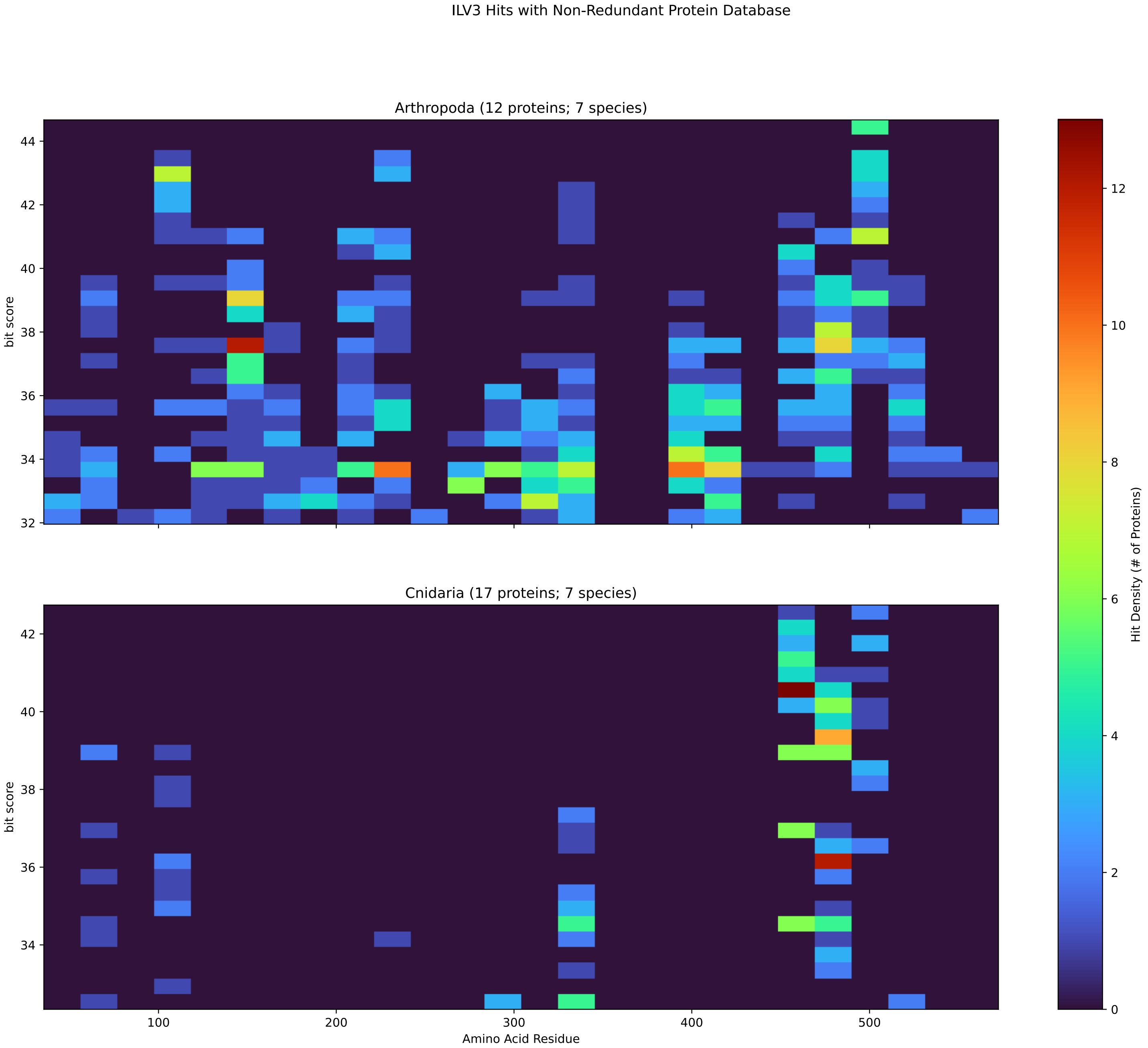


Figure S157: Non-redundant (NR) protein hits for Ilv3 in the kingdom Metazoa.

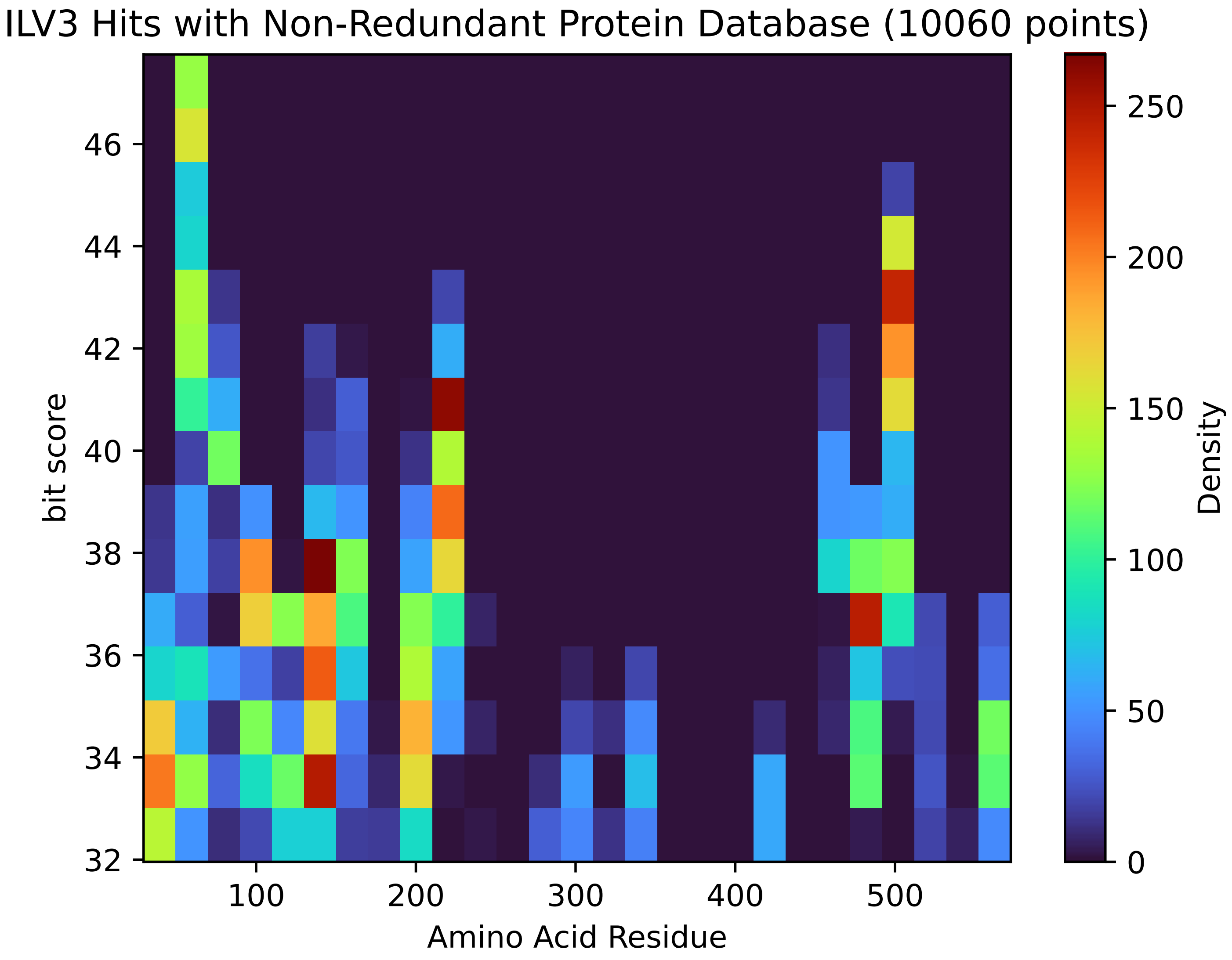


Figure S158: Non-redundant (NR) protein hits for Ilv3 in the kingdom SAR.

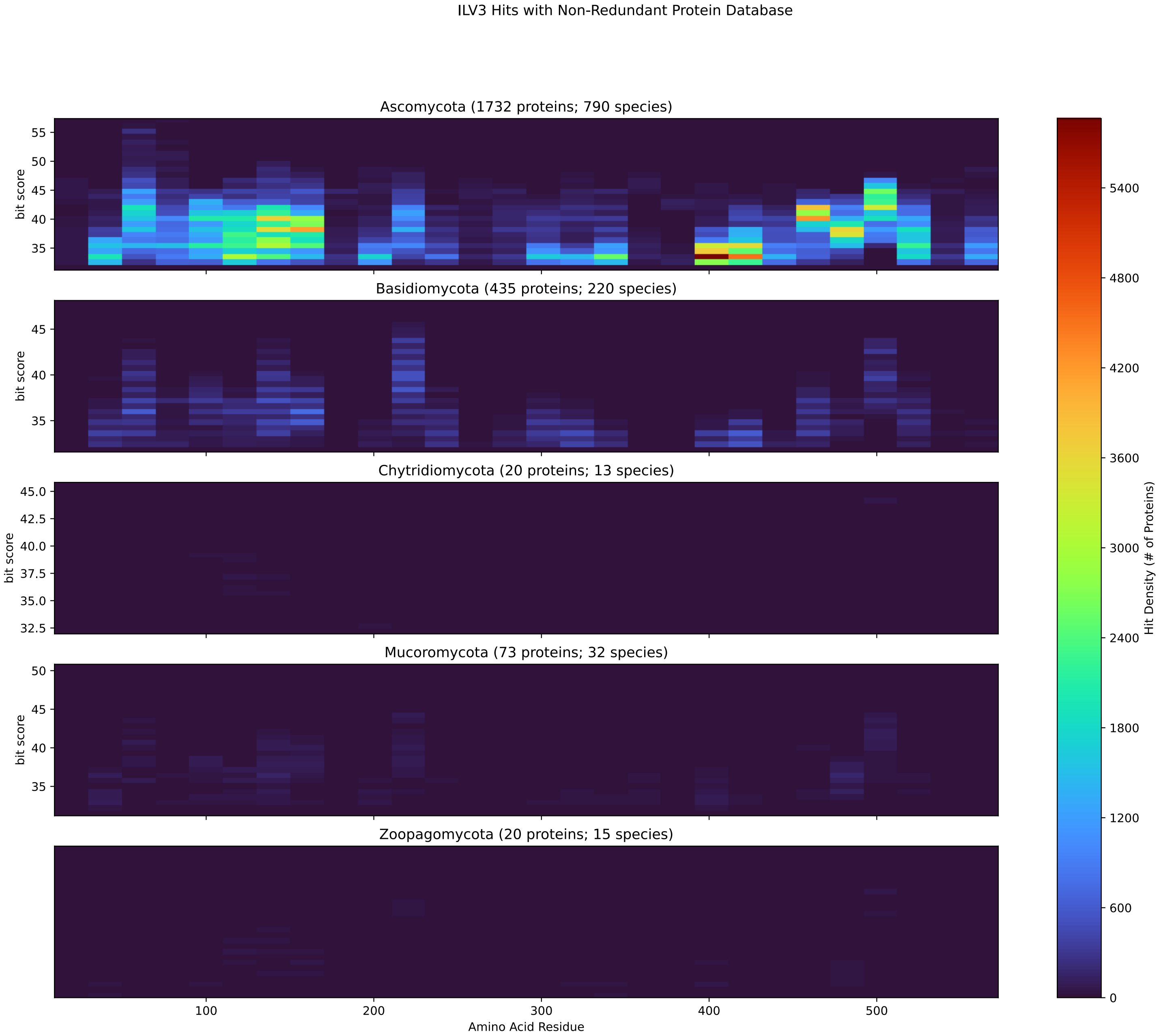


Figure S159: Non-redundant (NR) protein hits for Ilv3 in the kingdom Fungi.

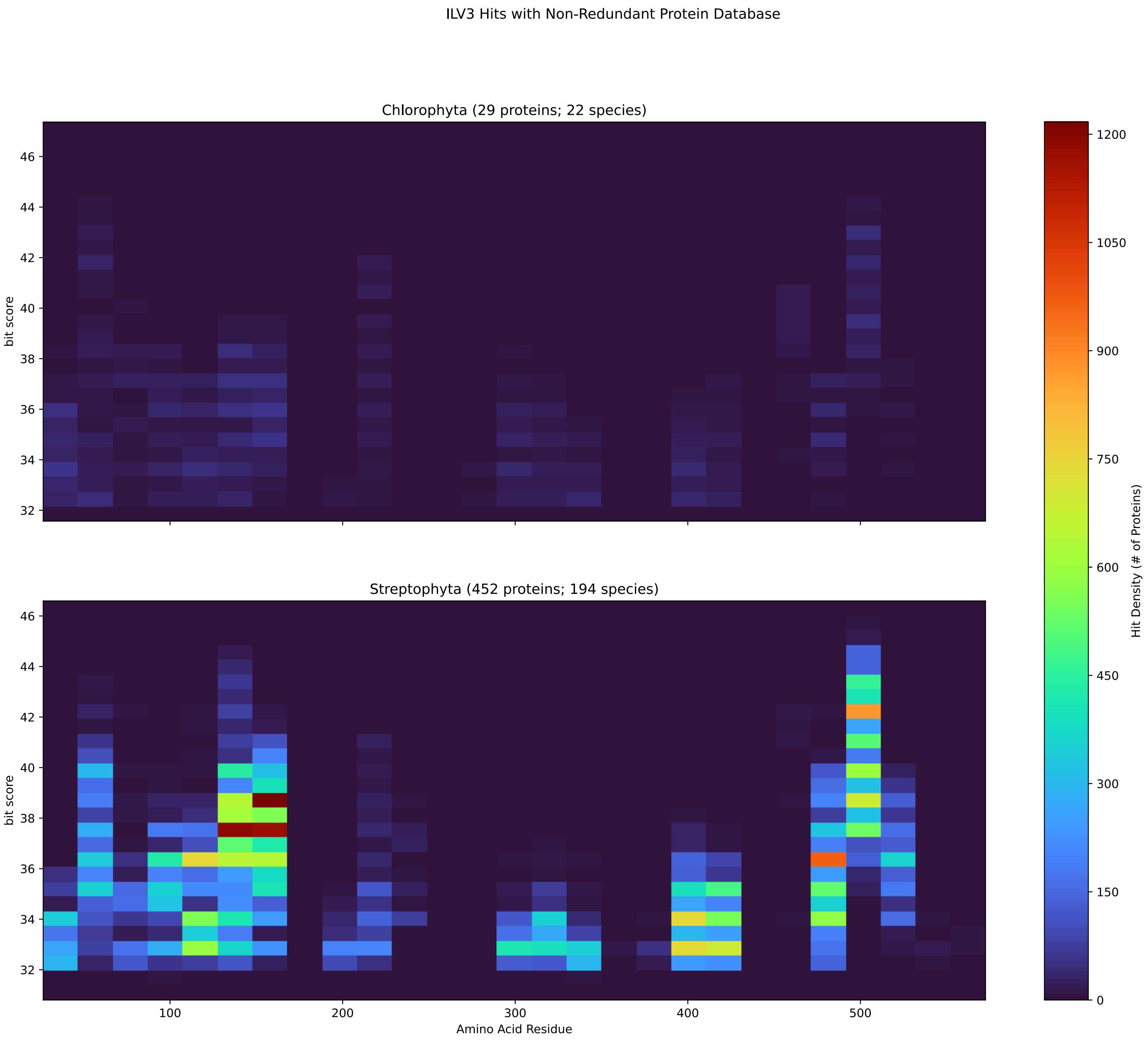


Figure S160: Non-redundant (NR) protein hits for Ilv3 in the kingdom Viridiplantae.

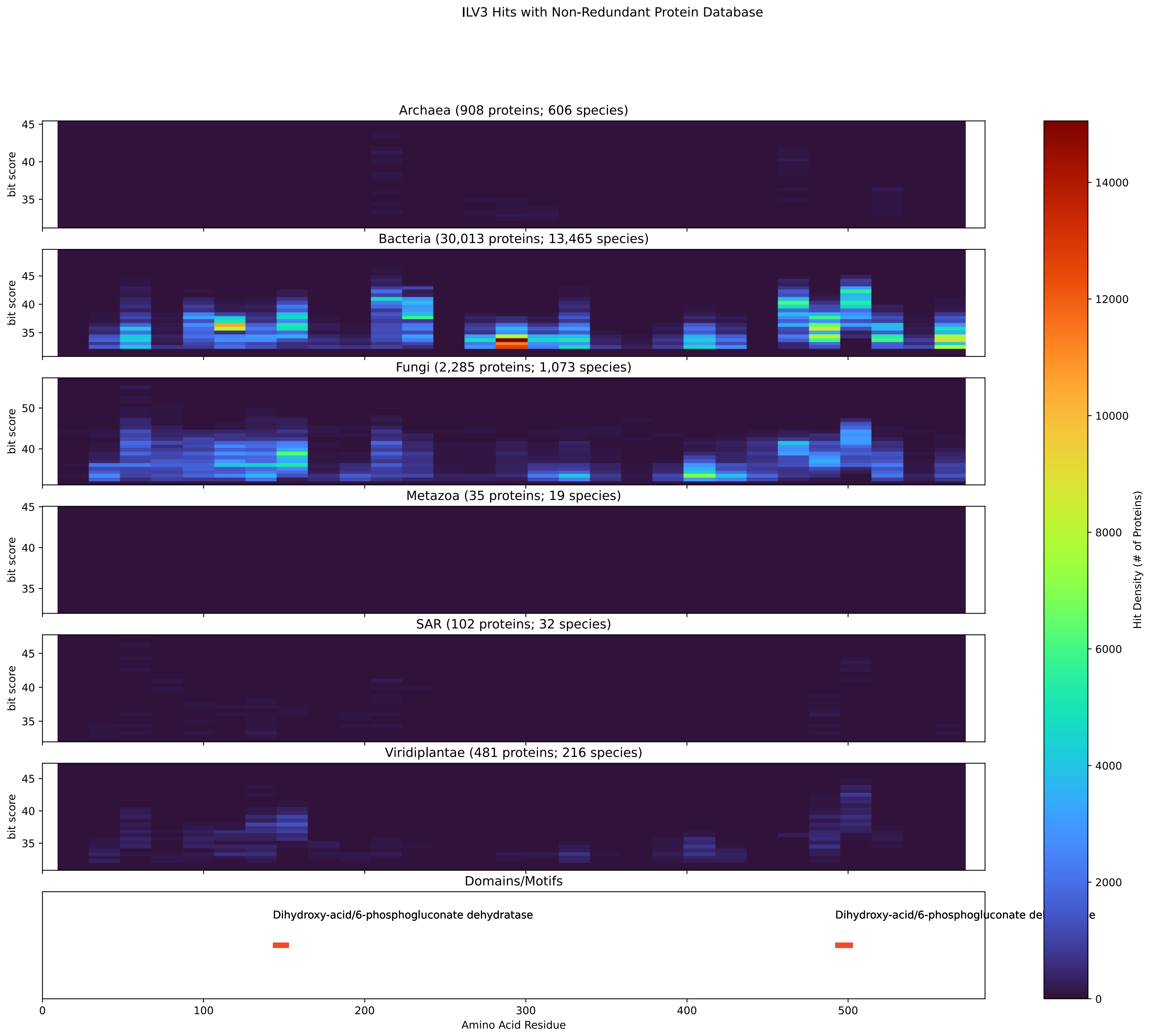


Figure S161: Non-redundant (NR) protein hits for DEG20010579/Ilv3 at 20 amino acid length queries.

#### S4.11 Ilv5

##### S4.11.1 WHO Critical Pathogens

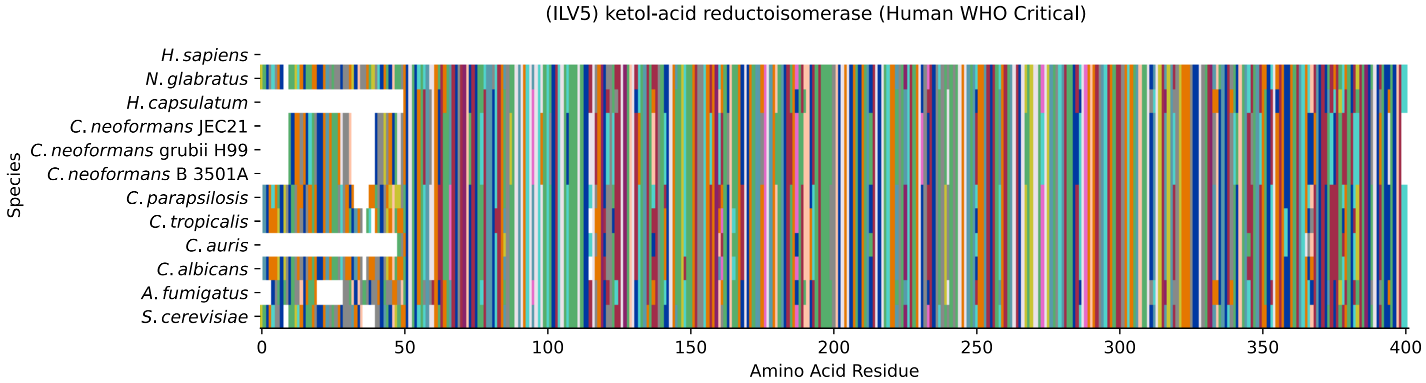


Figure S162: Multiple sequence alignment of yeast Ilv5 (WHO Critical Pathogens). Cf. Figure S163 for alignment quality, and Figure S164 for Sneath similarity. Cf. Table S27 for protein names, and pairwise alignment metrics with yeast Ilv5.

### Ilv5 MSA Quality

| | *A. f.* | *C. a.* | *C. a.* | *C. p.* | *C. t.* | *C. n.* | *C. n.* | *C. n.* | *H. c.* | *N. g.* | *S. c.* |
|---|---|---|---|---|---|---|---|---|---|---|---|
| *A. fumigatus* | 799.66 | 560.451 | 555.058 | 582.022 | 554.673 | 574.704 | 574.704 | 574.704 | 685.256 | 577.015 | 569.696 |
| *C. albicans* | 584.719 | 825.083 | 624.78 | 717.613 | 755.747 | 561.607 | 561.992 | 561.607 | 586.645 | 650.203 | 637.106 |
| *C. auris* | 591.652 | 642.499 | 728.783 | 667.537 | 638.262 | 552.362 | 552.362 | 552.362 | 600.127 | 627.476 | 628.246 |
| *C. parapsilosis* | 607.06 | 717.613 | 646.736 | 819.691 | 708.368 | 572.778 | 573.163 | 572.778 | 605.134 | 663.685 | 658.292 |
| *C. tropicalis* | 579.326 | 755.747 | 620.928 | 708.368 | 822.002 | 558.14 | 558.525 | 558.14 | 573.933 | 637.876 | 621.313 |
| *C. neoformans* B 3501A | 574.704 | 541.576 | 517.694 | 552.747 | 537.724 | 786.178 | 785.793 | 786.178 | 560.836 | 536.954 | 534.258 |
| *C. neoformans* grubii H99 | 574.704 | 541.962 | 518.079 | 553.132 | 538.11 | 785.793 | 786.178 | 785.793 | 560.451 | 537.724 | 534.258 |
| *C. neoformans* JEC21 | 574.704 | 541.576 | 517.694 | 552.747 | 537.724 | 786.178 | 785.793 | 786.178 | 560.836 | 536.954 | 534.258 |
| *H. capsulatum* | 685.256 | 563.148 | 563.533 | 581.637 | 550.821 | 560.836 | 560.451 | 560.836 | 723.391 | 585.104 | 582.022 |
| *N. glabratus* | 601.282 | 650.588 | 611.298 | 664.455 | 639.032 | 558.525 | 558.525 | 558.525 | 608.216 | 830.091 | 731.865 |
| *S. cerevisiae* | 593.964 | 637.491 | 611.298 | 659.062 | 621.698 | 554.673 | 555.058 | 554.673 | 605.905 | 731.48 | 821.231 |

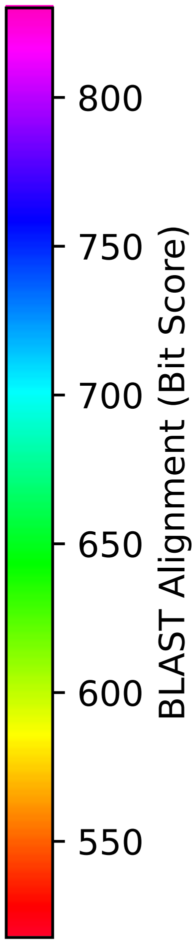


Figure S163: Multiple sequence alignment quality of Ilv5 (WHO Critical Pathogens). Cf. Figures S162 and S165.

| Species | Hit Protein | Hit Length (a.a.) | evalue | align_len | bit_score | identity | positive | score | gaps | % identity | % positive |
|---|---|---|---|---|---|---|---|---|---|---|---|
| H.sapiens | - | - | - | - | - | - | - | - | - | - | - |
| N.glabratus | XP_445105.1 uncharacterized protein CAGL0B03047g Nakaseomyces glabratus | 400 | 0 | 400 | 731.48 | 358 | 383 | 1887 | 6 | 90.6 | 97.0 |
| H.capsulatum | XP_045283741.1 ketol-acid reductoisomerase Histoplasma capsulatum G186AR | 351 | 0 | 351 | 606.675 | 287 | 315 | 1563 | 0 | 72.7 | 79.7 |
| C.neoformans.JEC21 | XP_571345.1 ketol-acid reductoisomerase, putative Cryptococcus neoformans var. neoformans JEC21 | 392 | 0 | 392 | 556.599 | 268 | 317 | 1433 | 11 | 67.8 | 80.3 |
| C.neoformans.grubii.H99 | XP_012050476.1 ketol-acid reductoisomerase, mitochondrial Cryptococcus neoformans var. grubii H99 | 392 | 0 | 392 | 556.599 | 268 | 317 | 1433 | 11 | 67.8 | 80.3 |
| C.neoformans.B.3501A | XP_774791.1 hypothetical protein CNBF2210 Cryptococcus neoformans var. neoformans B-3501A | 392 | 0 | 392 | 556.599 | 268 | 317 | 1433 | 11 | 67.8 | 80.3 |
| C.parapsilosis | XP_036663080.1 uncharacterized protein CPAR2 603140 Candida parapsilosis | 396 | 0 | 396 | 659.062 | 322 | 358 | 1699 | 3 | 81.5 | 90.6 |
| C.tropicalis | XP_002548652.1 ketol-acid reductoisomerase, mitochondrial precursor Candida tropicalis MYA-3404 | 398 | 0 | 398 | 621.698 | 307 | 354 | 1602 | 5 | 77.7 | 89.6 |
| C.auris | XP_028890742.2 ketol-acid reductoisomerase, mitochondrial Candida auris | 353 | 0 | 353 | 614.764 | 308 | 333 | 1584 | 0 | 78.0 | 84.3 |
| C.albicans | XP_714297.2 ketol-acid reductoisomerase Candida albicans SC5314 | 399 | 0 | 399 | 637.876 | 315 | 357 | 1644 | 6 | 79.7 | 90.4 |
| A.fumigatus | XP_754177.1 Ketol-acid reductoisomerase Aspergillus fumigatus Af293 | 388 | 0 | 388 | 595.89 | 279 | 325 | 1535 | 0 | 70.6 | 82.3 |

Table S27: Pairwise alignment info from yeast Ilv5 (DEG20010747), cf. Figure S162.

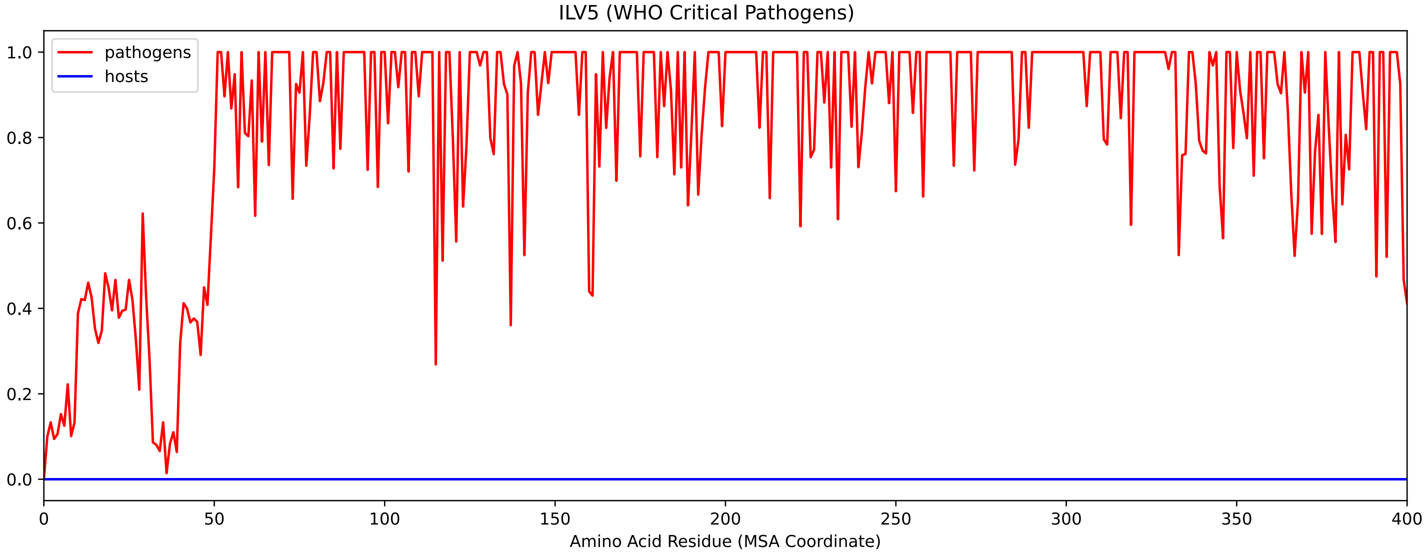


Figure S164: Sneath Similarity of Ilv5 for WHO Critical Pathogens, cf. Figure S162

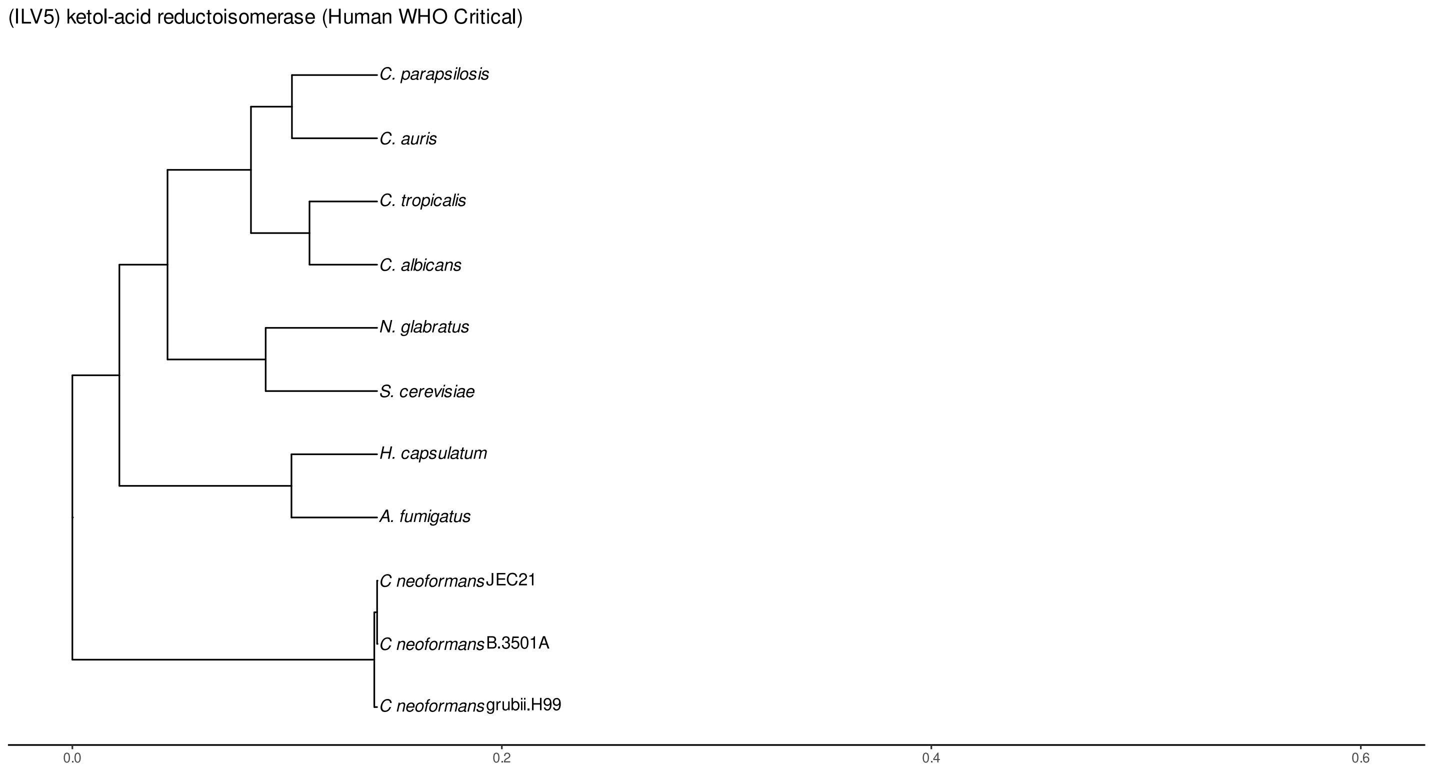


Figure S165: Phylogeny of hits for Ilv5 made using clustal omega and visualized using ggtree, cf. Figures S162 and S163.

##### S4.11.2 Top 10 Agricultural Fungal Pathogens

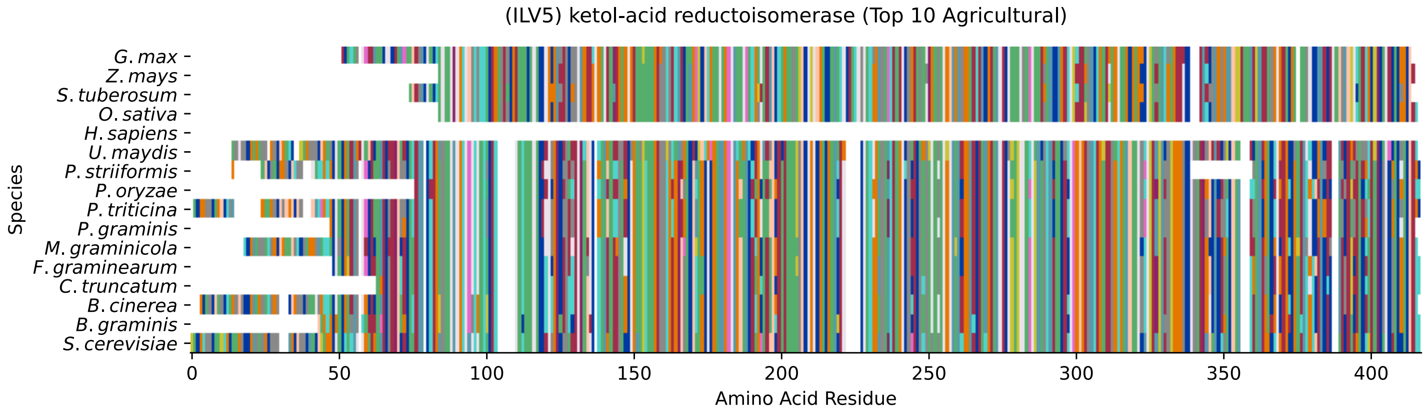


Figure S166: Multiple sequence alignment of yeast Ilv5 (Top 10 Agricultural Fungal Pathogens). Cf. Figure S167 for alignment quality, and Figure S168 for Sneath similarity. Cf. Table S28 for protein names, and pairwise alignment metrics with yeast Ilv5.

### Ilv5 MSA Quality

| | *B. g.* | *B. c.* | *C. t.* | *F. g.* | *M. g.* | *P. g.* | *P. t.* | *P. o.* | *P. s.* | *U. m.* | *S. c.* |
|---|---|---|---|---|---|---|---|---|---|---|---|
| *B. graminis* | 738.799 | 662.144 | 600.897 | 613.609 | 600.512 | 534.258 | 533.487 | 586.26 | 504.212 | 469.929 | 526.939 |
| *B. cinerea* | 662.529 | 810.446 | 631.328 | 655.596 | 652.899 | 555.444 | 555.058 | 616.305 | 533.102 | 505.753 | 567 |
| *C. truncatum* | 600.897 | 630.943 | 697.197 | 654.825 | 605.52 | 530.02 | 532.717 | 621.313 | 493.426 | 485.337 | 553.518 |
| *F. graminearum* | 613.609 | 654.44 | 654.825 | 725.317 | 632.098 | 550.821 | 553.903 | 640.573 | 514.227 | 487.649 | 563.918 |
| *M. graminicola* | 600.512 | 640.573 | 605.52 | 632.098 | 783.097 | 550.436 | 555.444 | 587.03 | 518.85 | 494.582 | 550.436 |
| *P. graminis* | 534.258 | 554.673 | 530.02 | 550.821 | 550.436 | 718.768 | 711.064 | 502.671 | 666.766 | 509.22 | 510.76 |
| *P. triticina* | 533.487 | 554.673 | 533.102 | 553.518 | 555.444 | 711.064 | 794.653 | 505.753 | 677.937 | 516.153 | 520.005 |
| *P. oryzae* | 586.26 | 615.15 | 621.313 | 640.573 | 587.03 | 502.671 | 505.753 | 670.233 | 466.463 | 461.84 | 533.487 |
| *P. striiformis* | 504.212 | 523.857 | 493.426 | 514.227 | 518.85 | 666.766 | 673.7 | 466.463 | 730.709 | 471.085 | 476.863 |
| *U. maydis* | 469.929 | 496.508 | 485.337 | 487.649 | 494.582 | 509.22 | 514.998 | 461.84 | 471.085 | 791.571 | 477.633 |
| *S. cerevisiae* | 545.814 | 579.326 | 576.63 | 587.03 | 574.318 | 533.487 | 541.191 | 556.984 | 499.59 | 501.516 | 821.231 |

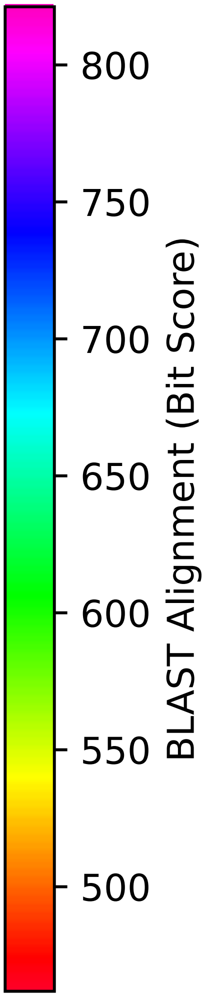


Figure S167: Multiple sequence alignment quality of Ilv5 (Top 10 Agricultural Fungal Pathogens). Cf. Figures S166 and S169.

| Species | Hit Protein | Hit Length (a.a.) | evalue | align_len | bit_score | identity | positive | score | gaps | % identity | % positive |
|---|---|---|---|---|---|---|---|---|---|---|---|
| G.max | NP_001276286.2 ketol-acid reductoisomerase, chloroplastic-like Glycine max | 363 | 4e-43 | 363 | 159.844 | 115 | 179 | 403 | 24 | 29.1 | 45.3 |
| Z.mays | NP_001169144.1 ketol-acid reductoisomerase, chloroplastic-like Zea mays | 332 | 6e-44 | 332 | 160.614 | 109 | 169 | 405 | 23 | 27.6 | 42.8 |
| S.tuberosum | XP_006348854.1 PREDICTED: ketol-acid reductoisomerase, chloroplastic Solanum tuberosum | 341 | 8.8e-43 | 341 | 157.918 | 109 | 171 | 398 | 24 | 27.6 | 43.3 |
| O.sativa | XP_015640421.1 ketol-acid reductoisomerase, chloroplastic Oryza sativa Japonica Group | 333 | 2.3e-43 | 333 | 159.458 | 108 | 168 | 402 | 23 | 27.3 | 42.5 |
| H.sapiens | - | - | - | - | - | - | - | - | - | - | - |
| U.maydis | XP_011391733.1 putative ketol--acid reductoisomerase Ustilago maydis 521 | 382 | 3.8e-179 | 382 | 503.056 | 243 | 290 | 1294 | 3 | 61.5 | 73.4 |
| P.striiformis | XP_047803961.1 hypothetical protein Pst134EA 017323 Puccinia striiformis f. sp. tritici | 375 | 2.9e-178 | 375 | 499.59 | 249 | 288 | 1285 | 21 | 63.0 | 72.9 |
| P.oryzae | mRNA M BR32 EuGene 00014811-p1 — transcript=mRNA M BR32 EuGene 00014811 — gene=M BR32 EuGene 00014811 — organism=Pyricularia oryzae BR32 — gene product=unspecified product — transcript product=unspecified product — location=BR32 scaffold000-01:4410728-4411788(+) — protein length=324 — sequence SO=supercontig — SO=protein coding gene — is pseudo=false | 323 | 0 | 323 | 556.984 | 260 | 293 | 1434 | 0 | 65.8 | 74.2 |
| P.triticina | XP_053025405.1 uncharacterized protein PtA15 11A542 Puccinia triticina | 394 | 0 | 394 | 540.421 | 268 | 309 | 1391 | 9 | 67.8 | 78.2 |
| P.graminis | XP_003323363.1 ketol-acid reductoisomerase, mitochondrial Puccinia graminis f. sp. tritici CRL 75-36-700-3 | 349 | 0 | 349 | 535.798 | 254 | 290 | 1379 | 0 | 64.3 | 73.4 |
| M.graminicola | ZTRI 1.395.mRNA-p1 — transcript=ZTRI 1.395.mRNA — gene=ZTRI 1.395 — organism=Zymoseptoria tritici IPO323 — gene product=-similar to ketol-acid reductoisomerase — transcript product=-similar to ketol-acid reductoisomerase — location=Ztri chr 1-:1345028-1346413(-) — protein length=403 — sequence SO=chromosome — SO=protein coding gene — is pseudo=false | 382 | 0 | 382 | 576.244 | 275 | 316 | 1484 | 9 | 69.6 | 80.0 |
| F.graminearum | XP_011319056.1 ketol-acid reductoisomerase Fusarium graminearum PH-1 | 350 | 0 | 350 | 588.956 | 278 | 310 | 1517 | 0 | 70.4 | 78.5 |
| C.truncatum | XP_036585043.1 ketol-acid reductoisomerase Colletotrichum truncatum | 336 | 0 | 336 | 576.63 | 269 | 305 | 1485 | 0 | 68.1 | 77.2 |
| B.cinerea | XP_001557193.1 Bcilv5 Botrytis cinerea B05.10 | 391 | 0 | 391 | 579.711 | 277 | 320 | 1493 | 0 | 70.1 | 81.0 |
| B.graminis | VCU40915.1 — transcript=BGT962-24V316 LOCUS2166 t1 — gene=BGT-96224V316 LOCUS2166 — organism=Blumeria graminis f. sp. tritici 96224 — gene product=unspecified product — transcript product=unspecified product — location=LR026987:2357263-235874-1(+) — protein length=402 — sequence SO=chromosome — SO=protein coding gene — is pseudo=false | 355 | 0 | 355 | 547.74 | 255 | 297 | 1410 | 0 | 64.6 | 75.2 |

Table S28: Pairwise alignment info from yeast Ilv5 (DEG20010747), cf. Figure S166.

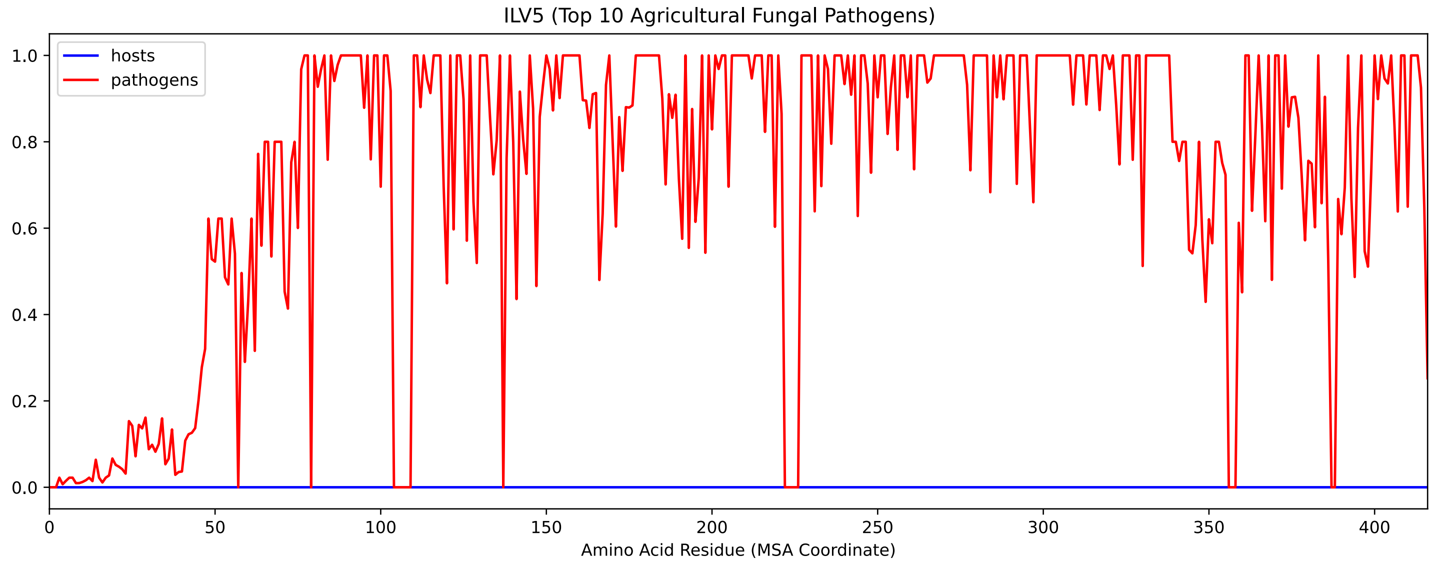


Figure S168: Sneath Similarity of Ilv5 for Top 10 Agricultural Fungal Pathogens, cf. Figure S166

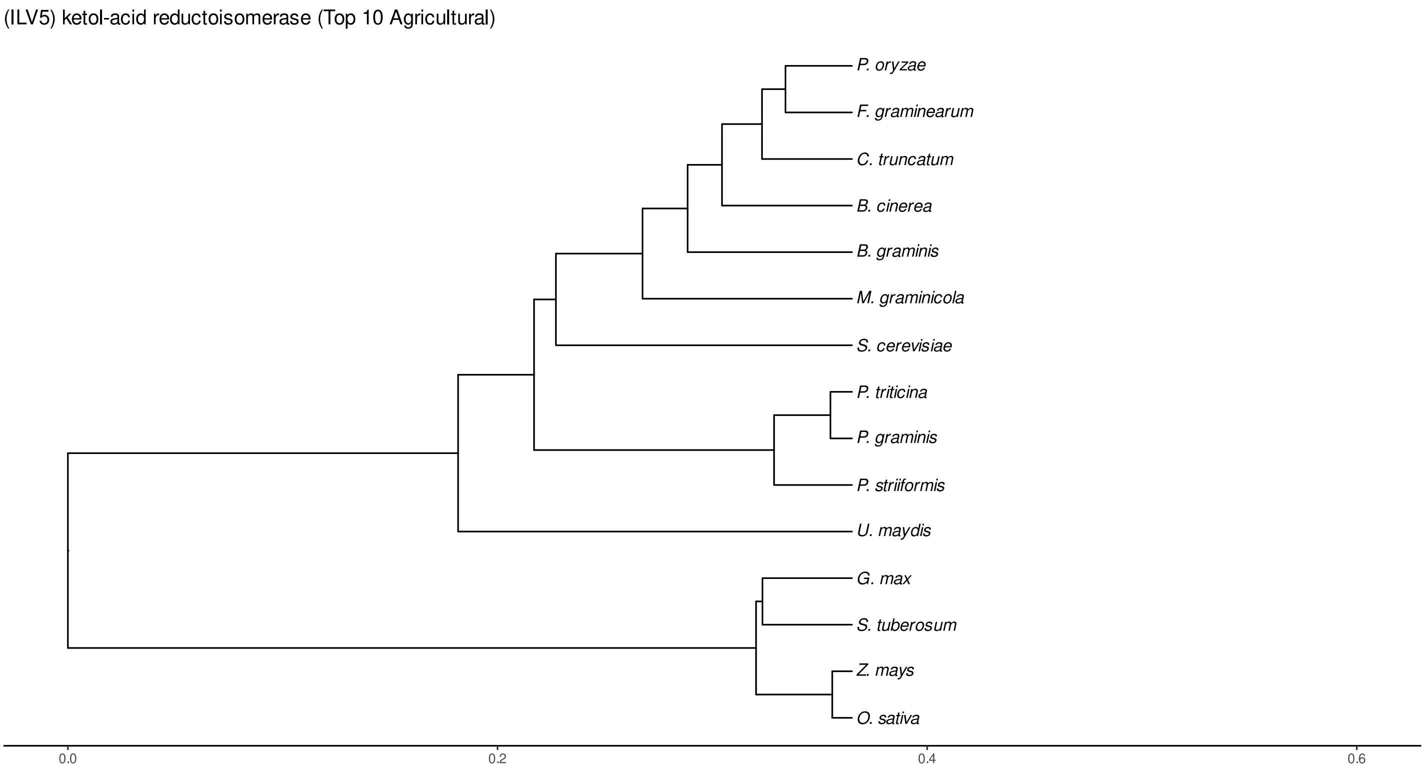


Figure S169: Phylogeny of hits for Ilv5 made using clustal omega and visualized using ggtree, cf. Figures S166 and S167.

#### S4.11.3 NR

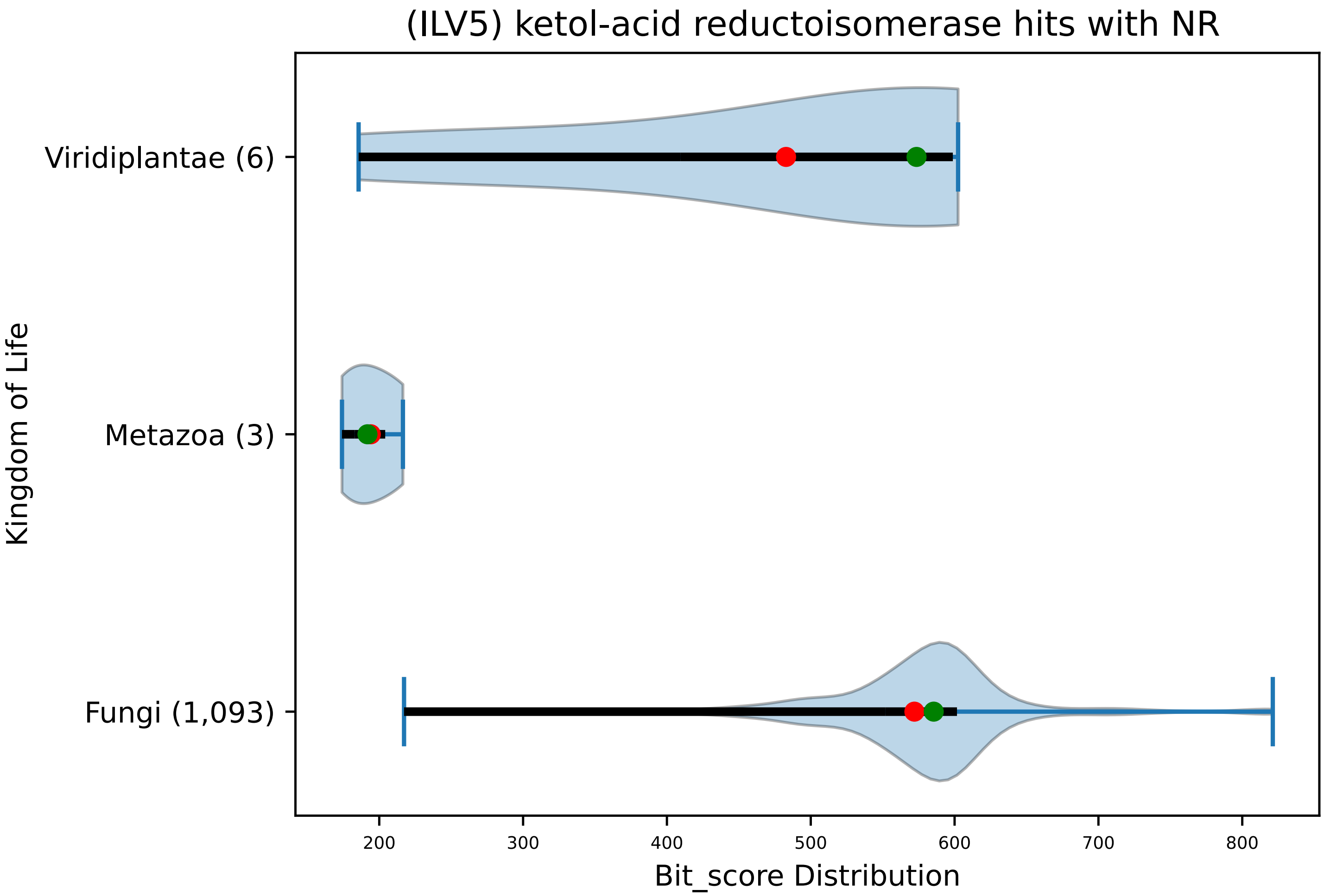


Figure S170: Non-redundant (NR) protein hits for DEG20010747/Ilv5, with expectation value of no more than 0.1. Green points are medians, and red points are arithmetic means.

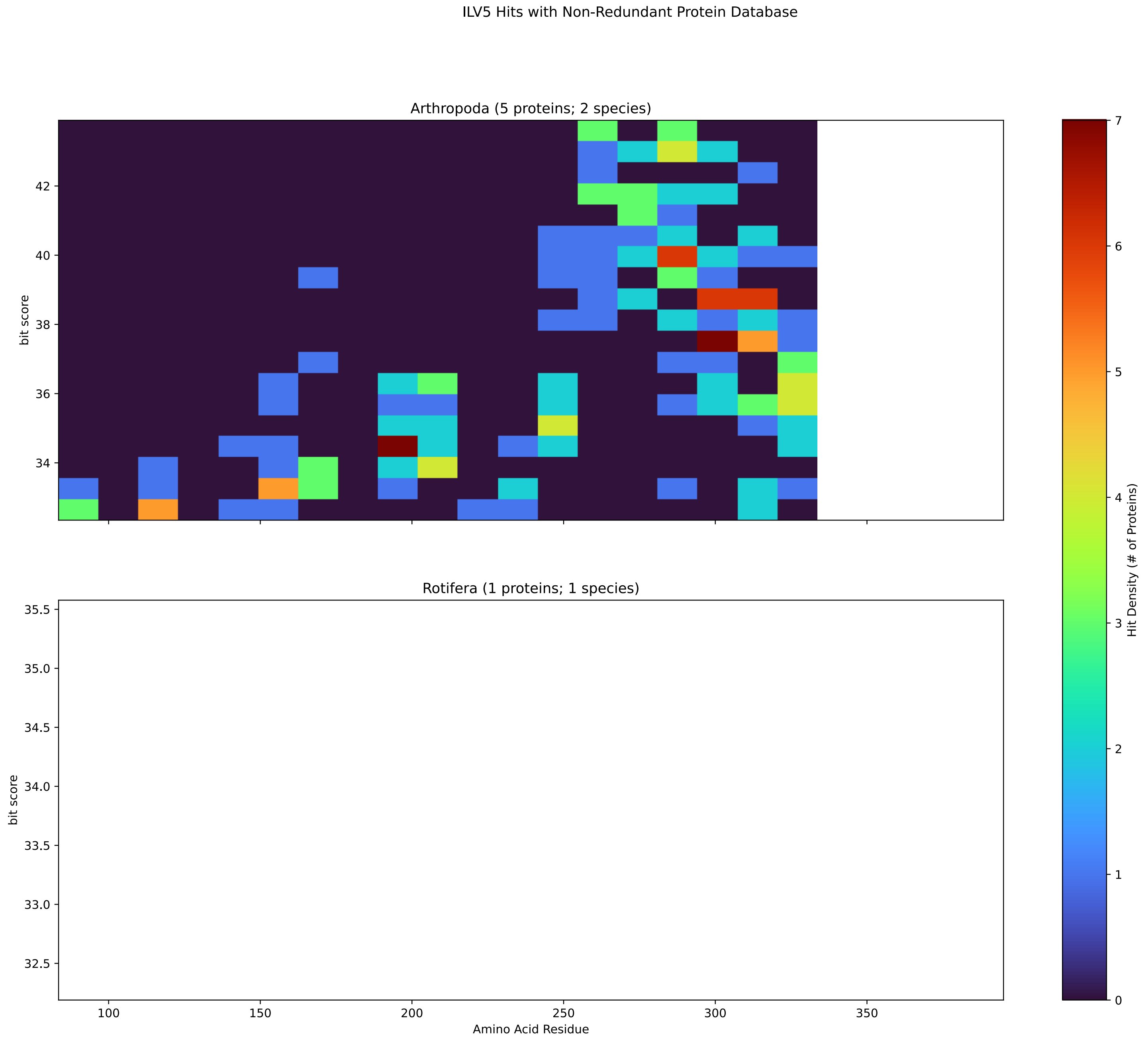


Figure S171: Non-redundant (NR) protein hits for Ilv5 in the kingdom Metazoa.

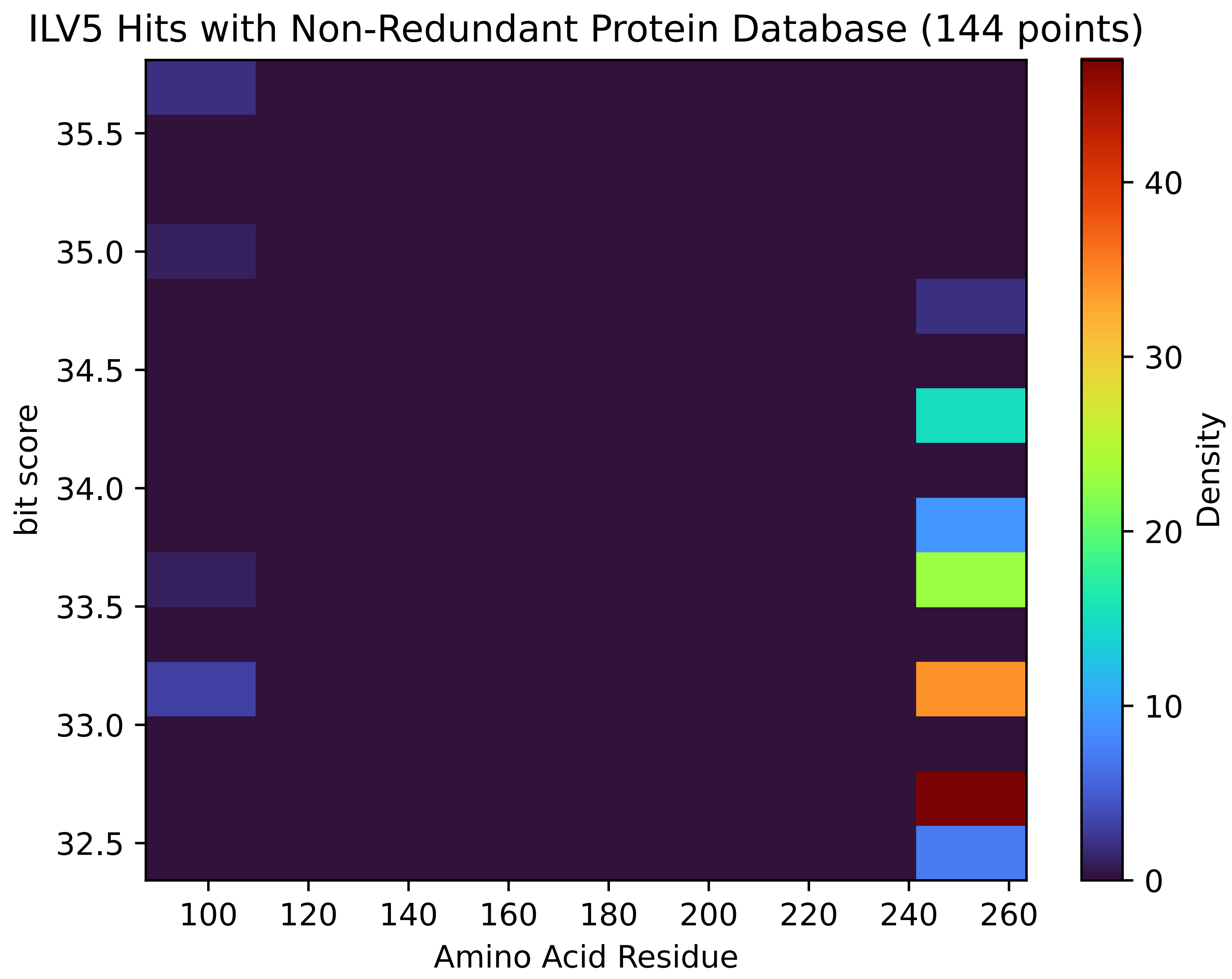


Figure S172: Non-redundant (NR) protein hits for Ilv5 in the kingdom SAR.

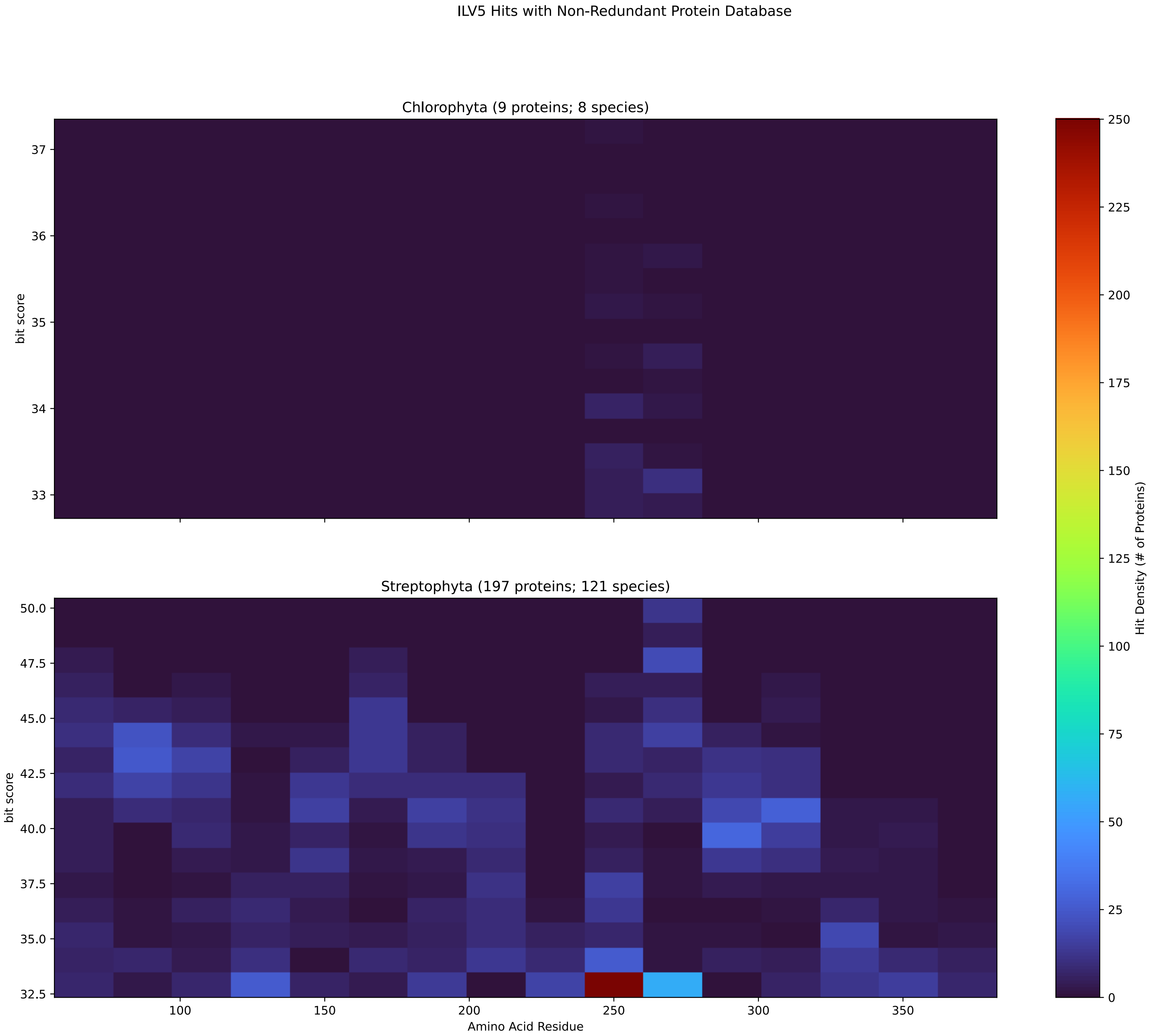


Figure S173: Non-redundant (NR) protein hits for Ilv5 in the kingdom Viridiplantae.

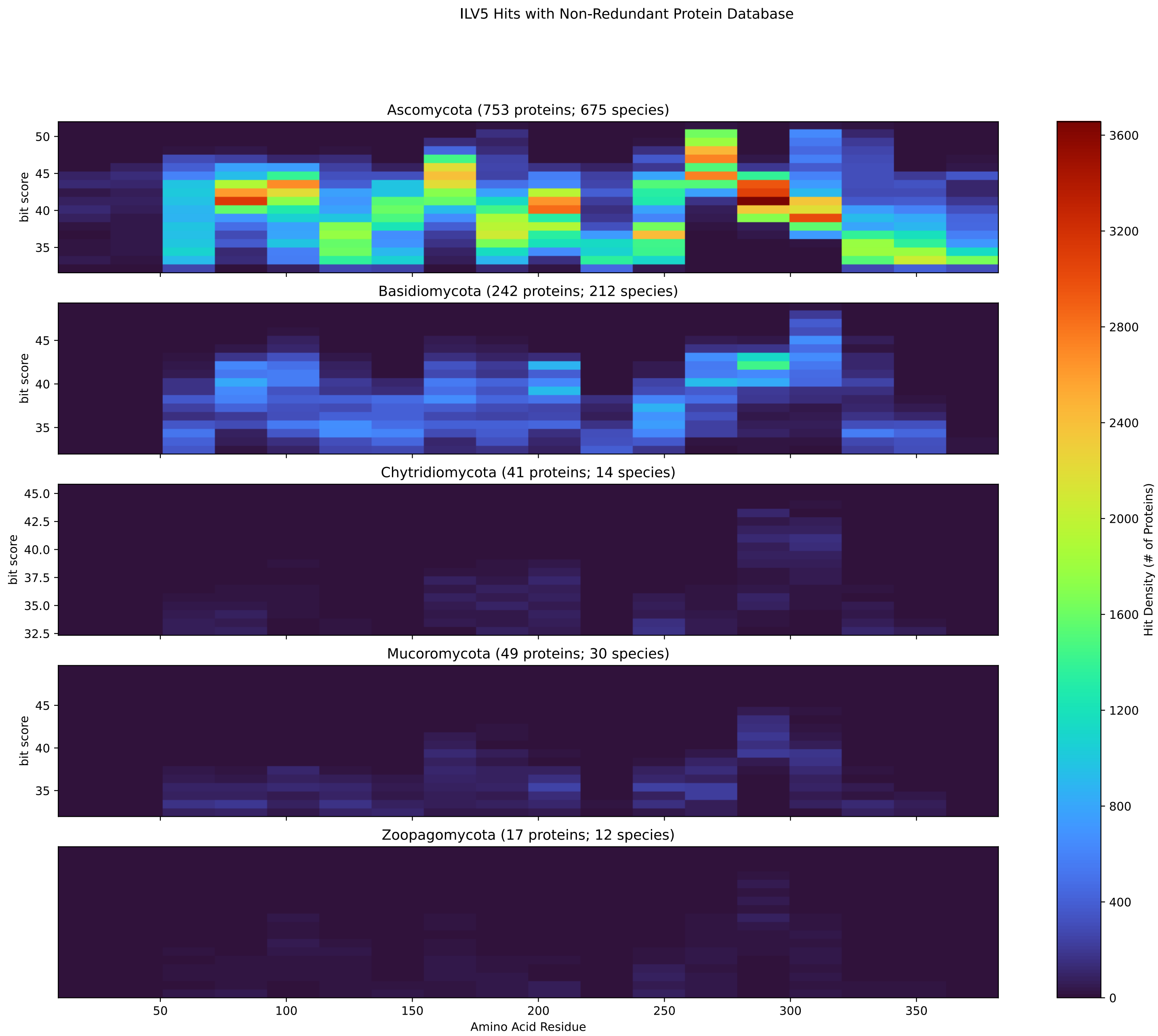


Figure S174: Non-redundant (NR) protein hits for Ilv5 in the kingdom Fungi.

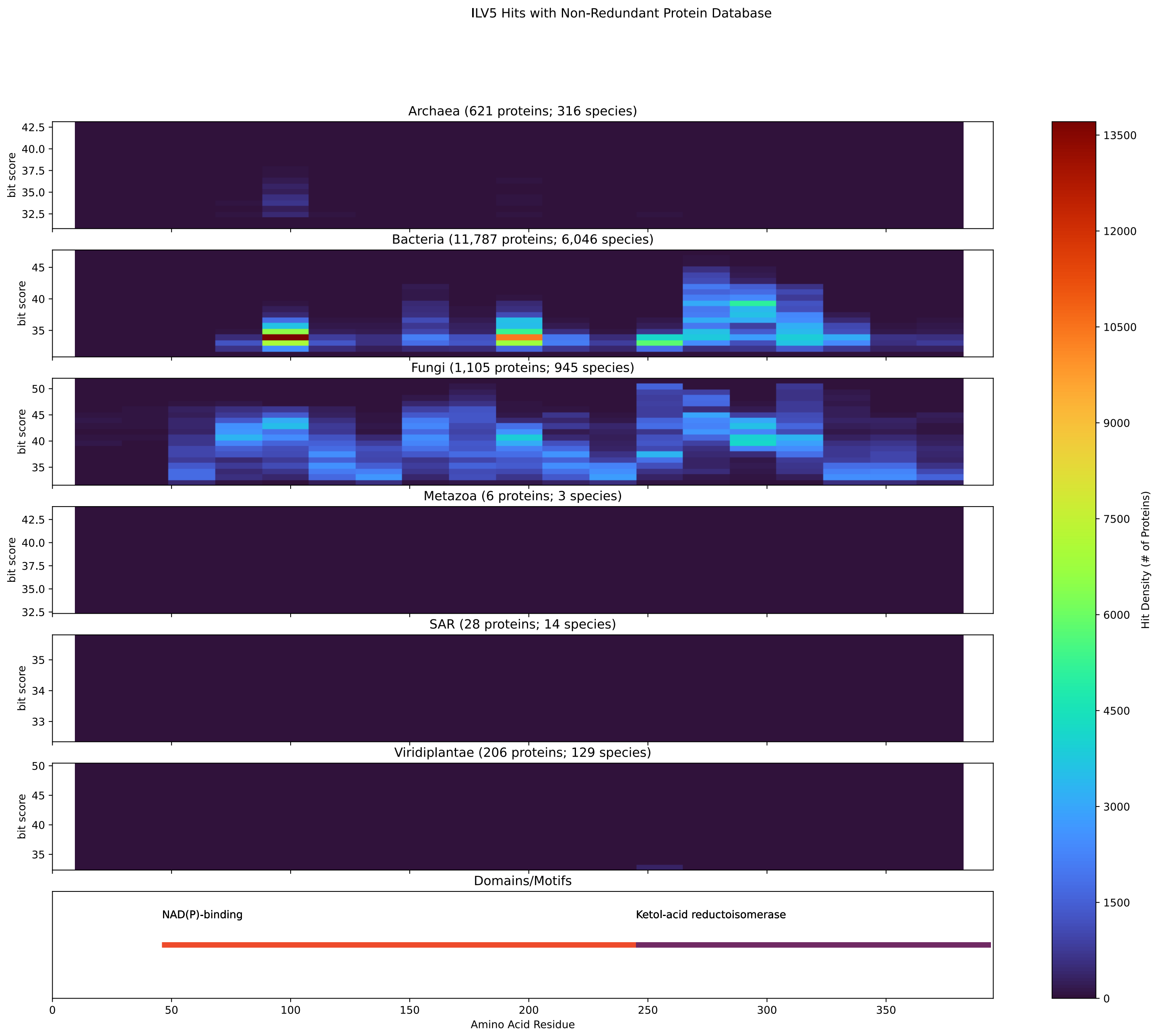


Figure S175: Non-redundant (NR) protein hits for DEG20010747/Ilv5 at 20 amino acid length queries.

#### S4.12 Rib3

##### S4.12.1 WHO Critical Pathogens

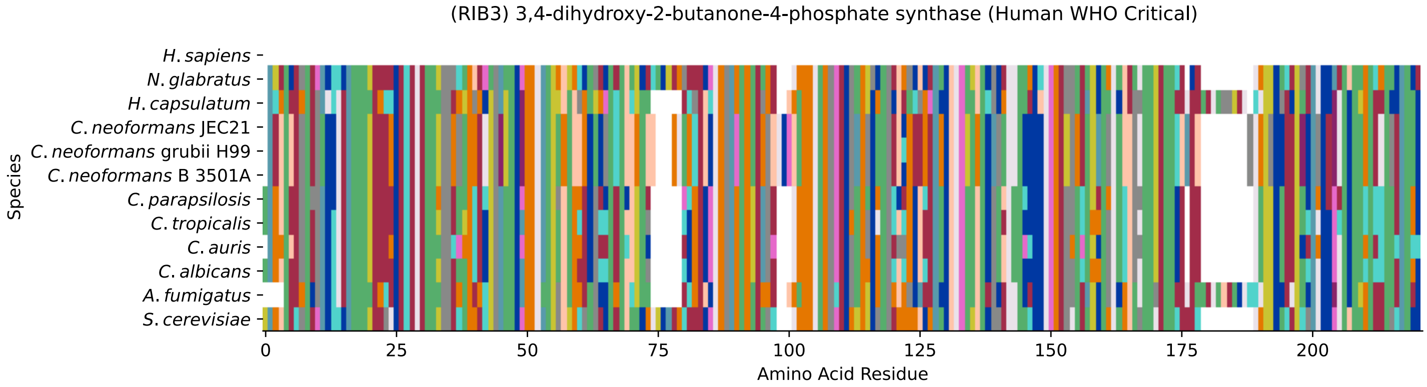


Figure S176: Multiple sequence alignment of yeast Rib3 (WHO Critical Pathogens). Cf. Figure S177 for alignment quality, and Figure S178 for Sneath similarity. Cf. Table S29 for protein names, and pairwise alignment metrics with yeast Rib3.

### Rib3 MSA Quality

| | *A. f.* | *C. a.* | *C. a.* | *C. p.* | *C. t.* | *C. n.* | *C. n.* | *C. n.* | *H. c.* | *N. g.* | *S. c.* |
|---|---|---|---|---|---|---|---|---|---|---|---|
| *A. fumigatus* | 425.246 | 226.098 | 211.846 | 211.846 | 217.238 | 185.267 | 186.808 | 185.267 | 305.449 | 212.616 | 204.912 |
| *C. albicans* | 226.098 | 416.772 | 328.176 | 351.288 | 394.815 | 212.616 | 216.083 | 212.616 | 207.994 | 253.832 | 252.292 |
| *C. auris* | 211.846 | 328.176 | 414.461 | 328.176 | 324.324 | 203.756 | 204.912 | 203.756 | 201.445 | 248.825 | 252.292 |
| *C. parapsilosis* | 211.846 | 351.288 | 328.176 | 416.001 | 344.739 | 198.749 | 200.675 | 198.749 | 193.741 | 245.358 | 251.521 |
| *C. tropicalis* | 217.238 | 394.815 | 324.324 | 344.739 | 417.542 | 202.986 | 206.068 | 202.986 | 199.134 | 250.366 | 249.595 |
| *C. neoformans* B 3501A | 185.267 | 212.616 | 203.756 | 198.749 | 202.986 | 427.172 | 424.476 | 427.172 | 183.341 | 194.897 | 193.741 |
| *C. neoformans* grubii H99 | 186.808 | 216.083 | 204.912 | 200.675 | 206.068 | 424.476 | 428.328 | 424.476 | 184.496 | 196.052 | 194.897 |
| *C. neoformans* JEC21 | 185.267 | 212.616 | 203.756 | 198.749 | 202.986 | 427.172 | 424.476 | 427.172 | 183.341 | 194.897 | 193.741 |
| *H. capsulatum* | 305.449 | 207.994 | 201.445 | 193.741 | 199.134 | 183.341 | 184.496 | 183.341 | 437.958 | 201.83 | 189.119 |
| *N. glabratus* | 212.616 | 253.832 | 248.825 | 245.358 | 250.366 | 194.897 | 196.052 | 194.897 | 201.83 | 424.861 | 315.464 |
| *S. cerevisiae* | 204.912 | 252.292 | 252.292 | 251.521 | 249.595 | 193.741 | 194.897 | 193.741 | 189.119 | 315.464 | 433.721 |

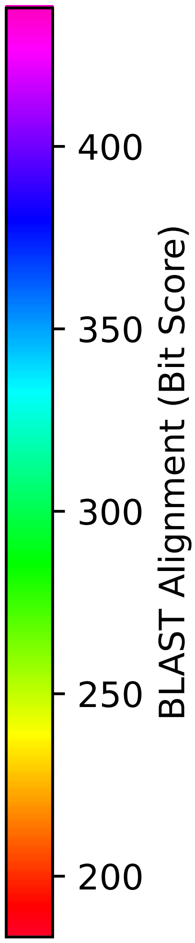


Figure S177: Multiple sequence alignment quality of Rib3 (WHO Critical Pathogens). Cf. Figures S176 and S179.

| Species | Hit Protein | Hit Length (a.a.) | evalue | align_len | bit_score | identity | positive | score | gaps | % identity | % positive |
|---|---|---|---|---|---|---|---|---|---|---|---|
| H.sapiens | - | - | - | - | - | - | - | - | - | - | - |
| N.glabratus | XP_447577.1 uncharacterized p-rotein CAGL0I07557g Nakaseomyc-es glabratus | 207 | 2.2e-111 | 207 | 315.464 | 150 | 171 | 807 | 1 | 72.1 | 82.2 |
| H.capsulatum | XP_045289685.1 3,4-dihydroxy–2-butanone 4-phosphate synthas-e Histoplasma capsulatum G186A-R | 217 | 4.5e-61 | 217 | 189.119 | 98 | 136 | 479 | 16 | 47.1 | 65.4 |
| C.neoformans.JEC21 | XP_570768.1 3,4 dihydroxy-2-b-utanone-4-phosphate synthase, putative Cryptococcus neoforma-ns var. neoformans JEC21 | 211 | 3.8e-63 | 211 | 194.512 | 99 | 133 | 493 | 7 | 47.6 | 63.9 |
| C.neoformans.grubii.H99 | XP_012050191.1 3,4-dihydroxy–2-butanone-4-phosphate synthas-e Cryptococcus neoformans var. grubii H99 | 211 | 2.1e-63 | 211 | 195.282 | 100 | 134 | 495 | 7 | 48.1 | 64.4 |
| C.neoformans.B.3501A | XP_775331.1 hypothetical prot-ein CNBE0490 Cryptococcus neof-ormans var. neoformans B-3501A | 211 | 3.6e-63 | 211 | 194.512 | 99 | 133 | 493 | 7 | 47.6 | 63.9 |
| C.parapsilosis | XP_036668231.1 uncharacterize-d protein CPAR2 700750 Candida parapsilosis | 209 | 2.8e-86 | 209 | 251.906 | 124 | 155 | 642 | 9 | 59.6 | 74.5 |
| C.tropicalis | XP_002549222.1 3,4-dihydroxy–2-butanone 4-phosphate synthas-e Candida tropicalis MYA-3404 | 208 | 2.1e-85 | 208 | 249.595 | 122 | 153 | 636 | 7 | 58.7 | 73.6 |
| C.auris | XP_028891318.1 3,4-dihydroxy–2-butanone 4-phosphate synthas-e Candida auris | 208 | 1.2e-86 | 208 | 252.677 | 123 | 156 | 644 | 9 | 59.1 | 75.0 |
| C.albicans | XP_716297.2 3,4-dihydroxy-2-b-utanone-4-phosphate synthase C-andida albicans SC5314 | 208 | 1.7e-86 | 208 | 252.292 | 122 | 157 | 643 | 7 | 58.7 | 75.5 |
| A.fumigatus | XP_751191.1 3,4-dihydroxy-2-b-utanone 4-phosphate synthase A-spergillus fumigatus Af293 | 215 | 3.3e-67 | 215 | 204.912 | 104 | 142 | 520 | 17 | 50.0 | 68.3 |

Table S29: Pairwise alignment info from yeast Rib3 (DEG20010261), cf. Figure S176.

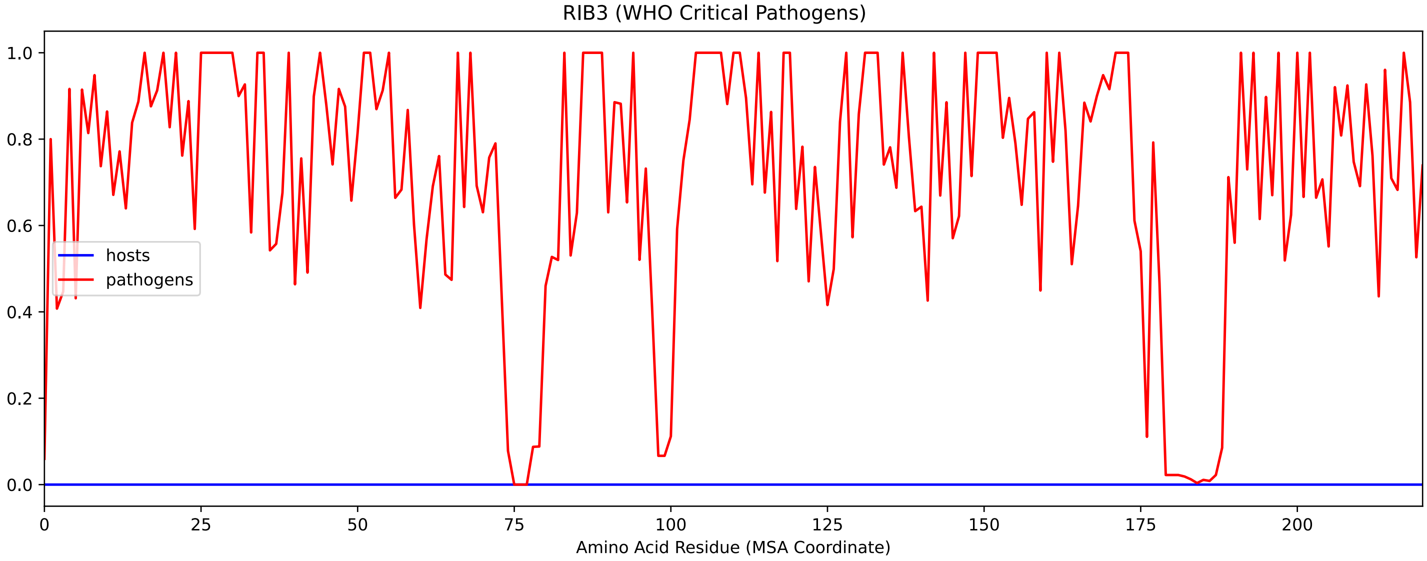


Figure S178: Sneath Similarity of Rib3 for WHO Critical Pathogens, cf. Figure S176

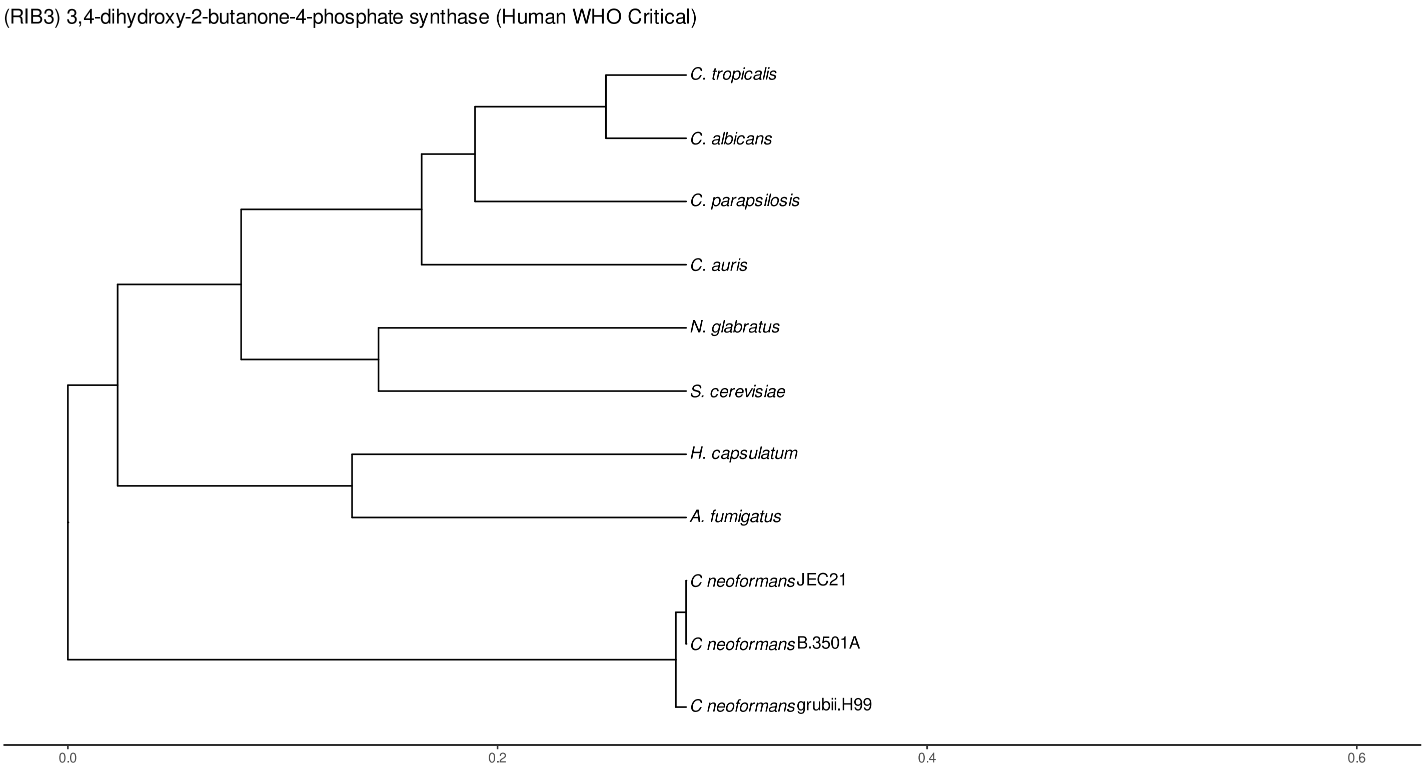


Figure S179: Phylogeny of hits for Rib3 made using clustal omega and visualized using ggtree, cf. Figures S176 and S177.

##### S4.12.2 Top 10 Agricultural Fungal Pathogens

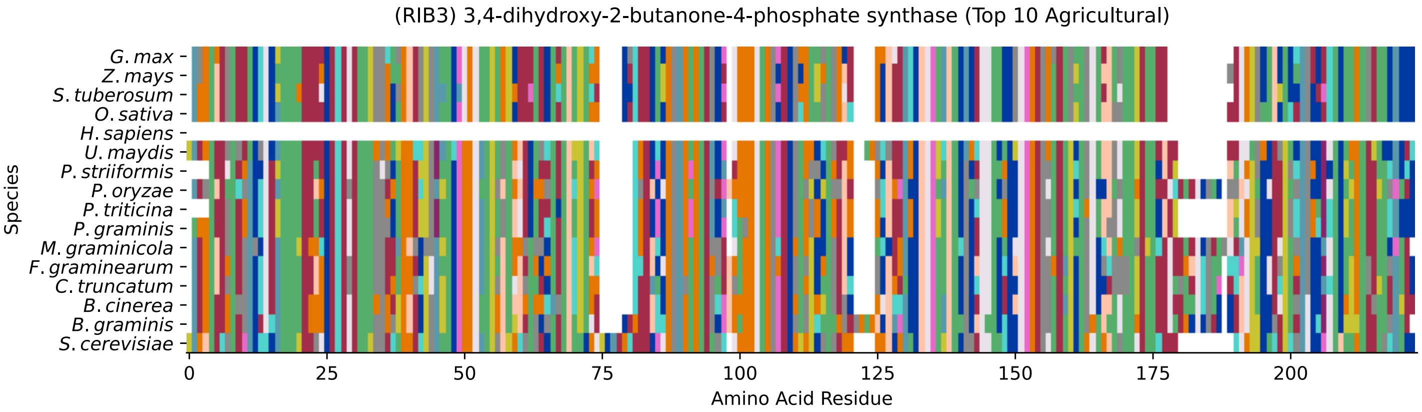


Figure S180: Multiple sequence alignment of yeast Rib3 (Top 10 Agricultural Fungal Pathogens). Cf. Figure S181 for alignment quality, and Figure S182 for Sneath similarity. Cf. Table S30 for protein names, and pairwise alignment metrics with yeast Rib3.

#### Rib3 MSA Quality

| | B. g. | B. c. | C. t. | F. g. | M. g. | P. g. | P. t. | P. o. | P. s. | U. m. | S. c. |
|---|---|---|---|---|---|---|---|---|---|---|---|
| *B. graminis* | 442.195 | 294.664 | 237.269 | 254.218 | 219.55 | 182.956 | 172.17 | 270.396 | 174.096 | 191.43 | 181.03 |
| *B. cinerea* | 294.664 | 437.573 | 269.24 | 287.345 | 270.781 | 203.756 | 201.445 | 295.819 | 201.83 | 224.557 | 200.29 |
| *C. truncatum* | 237.269 | 269.24 | 429.483 | 321.627 | 256.529 | 209.92 | 205.682 | 304.679 | 203.371 | 199.134 | 200.675 |
| *F. graminearum* | 254.218 | 287.345 | 321.627 | 434.106 | 260.381 | 198.749 | 192.971 | 312.768 | 192.971 | 203.756 | 199.519 |
| *M. graminicola* | 219.55 | 270.781 | 256.529 | 260.381 | 429.483 | 188.348 | 187.193 | 268.085 | 189.889 | 207.223 | 215.698 |
| *P. graminis* | 182.956 | 203.756 | 209.92 | 198.749 | 188.348 | 419.083 | 390.193 | 189.889 | 380.563 | 253.447 | 204.142 |
| *P. triticina* | 172.17 | 201.445 | 205.682 | 192.971 | 187.193 | 390.193 | 415.616 | 192.971 | 384.415 | 257.299 | 208.764 |
| *P. oryzae* | 270.396 | 295.819 | 304.679 | 312.768 | 268.085 | 189.889 | 192.971 | 428.713 | 191.045 | 211.46 | 210.305 |
| *P. striiformis* | 174.096 | 201.83 | 203.371 | 192.971 | 189.889 | 380.563 | 384.415 | 191.045 | 411.764 | 253.447 | 205.297 |
| *U. maydis* | 191.43 | 224.557 | 199.134 | 203.756 | 207.223 | 253.447 | 257.299 | 211.46 | 253.447 | 429.098 | 191.045 |
| *S. cerevisiae* | 181.03 | 200.29 | 200.675 | 199.519 | 215.698 | 204.142 | 208.764 | 210.305 | 205.297 | 191.045 | 433.721 |

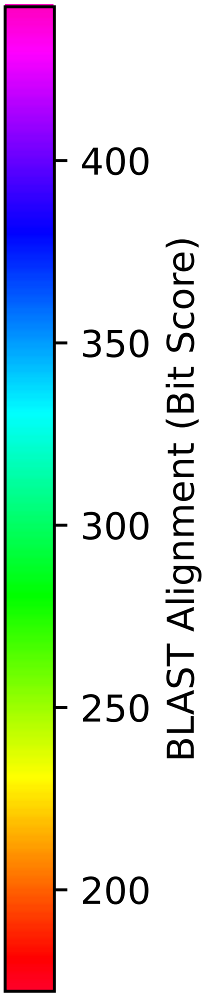


Figure S181: Multiple sequence alignment quality of Rib3 (Top 10 Agricultural Fungal Pathogens). Cf. Figures S180 and S183.

| Species | Hit Protein | Hit Length (a.a.) | evalue | align_len | bit_score | identity | positive | score | gaps | % identity | % positive |
|---|---|---|---|---|---|---|---|---|---|---|---|
| G.max | XP_003545697.1 bifunctional riboflavin biosynthesis protein RIBA 1, chloroplastic Glycine max | 207 | 3.4e-60 | 207 | 197.978 | 101 | 137 | 502 | 6 | 48.6 | 65.9 |
| Z.mays | XP_008677643.1 uncharacterized protein LOC100383821 isoform X5 Zea mays | 207 | 2.5e-59 | 207 | 193.356 | 101 | 135 | 490 | 5 | 48.6 | 64.9 |
| S.tuberosum | XP_006351756.1 PREDICTED: bifunctional riboflavin biosynthesis protein RIBA 1, chloroplastic-like Solanum tuberosum | 207 | 4.5e-60 | 207 | 197.208 | 99 | 138 | 500 | 6 | 47.6 | 66.3 |
| O.sativa | XP_015651119.1 probable bifunctional riboflavin biosynthesis protein RIBA 1, chloroplastic isoform X1 Oryza sativa Japonica Group | 207 | 6.4e-60 | 207 | 196.823 | 100 | 141 | 499 | 5 | 48.1 | 67.8 |
| H.sapiens | - | - | - | - | - | - | - | - | - | - | - |
| U.maydis | XP_011392588.1 putative 3,4-dihydroxy-2-butanone 4-phosphate synthase Ustilago maydis 521 | 211 | 9.6e-62 | 211 | 191.045 | 96 | 140 | 484 | 9 | 46.2 | 67.3 |
| P.striiformis | XP_047796296.1 uncharacterized protein Pst134EA 032002 Puccinia striiformis f. sp. tritici | 205 | 1.4e-67 | 205 | 206.068 | 101 | 140 | 523 | 7 | 48.6 | 67.3 |
| P.oryzae | mRNA M BR32 EuGene 00029391-p1 — transcript=mRNA M BR32 EuGene 00029391 — gene=M BR32 EuGene 00029391 — organism=Pyricularia oryzae BR32 — gene product=unspecified product — transcript product=unspecified product — location=BR32 scaffold000-02:3710575-3711355(+) — protein length=233 — sequence SO=supercontig — SO=protein coding gene — is pseudo=false | 217 | 2.3e-69 | 217 | 210.69 | 106 | 145 | 535 | 16 | 51.0 | 69.7 |
| P.triticina | XP_053020497.1 uncharacterized protein PtA15 5A515 Puccinia triticina | 205 | 3.7e-69 | 205 | 209.534 | 103 | 138 | 532 | 7 | 49.5 | 66.3 |
| P.graminis | XP_003338670.1 3,4-dihydroxy–2-butanone 4-phosphate synthase Puccinia graminis f. sp. tritici CRL 75-36-700-3 | 208 | 3.6e-67 | 208 | 204.527 | 98 | 141 | 519 | 7 | 47.1 | 67.8 |
| M.graminicola | ZTRI 3.823.mRNA-p1 — transcript=ZTRI 3.823.mRNA — gene=ZTRI 3.823 — organism=Zymoseptoria tritici IPO323 — gene product=similar to 3 — transcript product=similar to 3 — location=Ztri chr 3:2587765-2588507(-) — protein length=226 — sequence SO=chromosome — SO=protein coding gene — is pseudo=false | 216 | 1.4e-71 | 216 | 216.083 | 108 | 149 | 549 | 16 | 51.9 | 71.6 |
| F.graminearum | XP_011320316.1 hypothetical protein FGSG 08452 Fusarium graminearum PH-1 | 217 | 6.9e-65 | 217 | 199.134 | 100 | 138 | 505 | 16 | 48.1 | 66.3 |
| C.truncatum | XP_036587073.1 3,4-dihydroxy–2-butanone 4-phosphate synthase Colletotrichum truncatum | 217 | 2.2e-65 | 217 | 200.675 | 100 | 140 | 509 | 16 | 48.1 | 67.3 |
| B.cinerea | XP_001552345.1 Bcrib3 Botrytis cinerea B05.10 | 217 | 4.1e-65 | 217 | 200.29 | 100 | 142 | 508 | 16 | 48.1 | 68.3 |
| B.graminis | VDB94000.1 — transcript=BGT96224V316 LOCUS7590 t1 — gene=BGT96224V316 LOCUS7590 — organism=Blumeria graminis f. sp. tritici 96224 — gene product=unspecified product — transcript product=unspecified product — location=LR026992:19353413-19354355(-) — protein length=247 — sequence SO=chromosome — SO=protein coding gene — is pseudo=false | 220 | 7e-58 | 220 | 181.03 | 100 | 138 | 458 | 18 | 48.1 | 66.3 |

Table S30: Pairwise alignment info from yeast Rib3 (DEG20010261), cf. Figure S180.

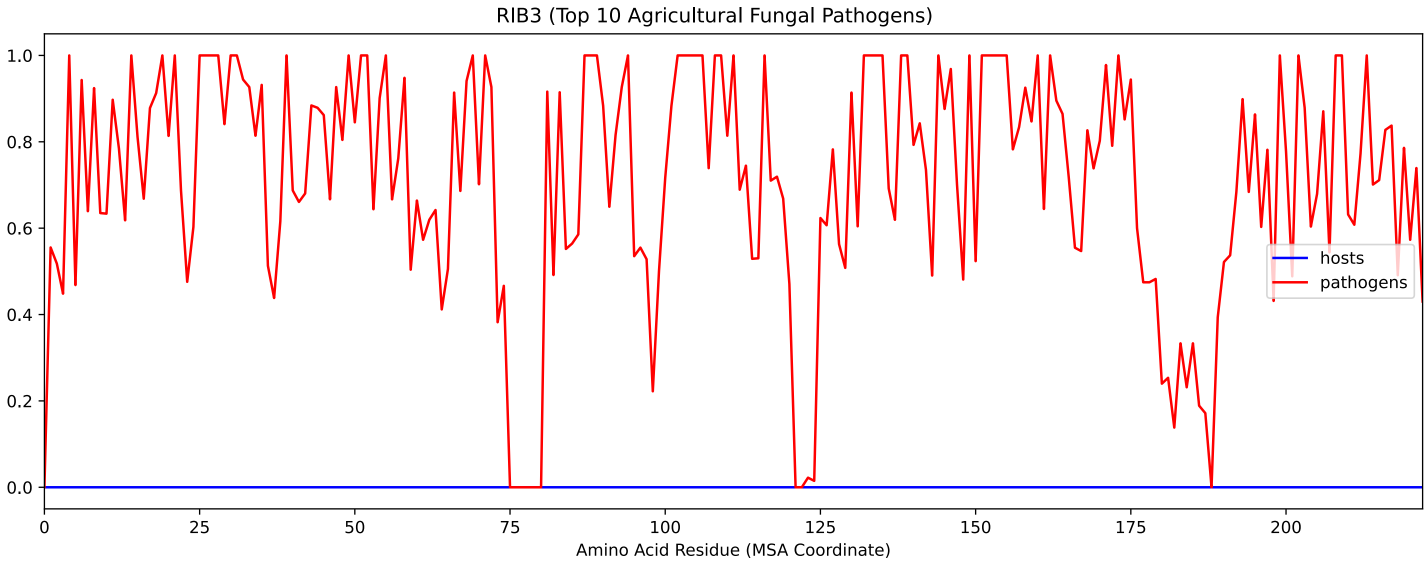


Figure S182: Sneath Similarity of Rib3 for Top 10 Agricultural Fungal Pathogens, cf. Figure S180

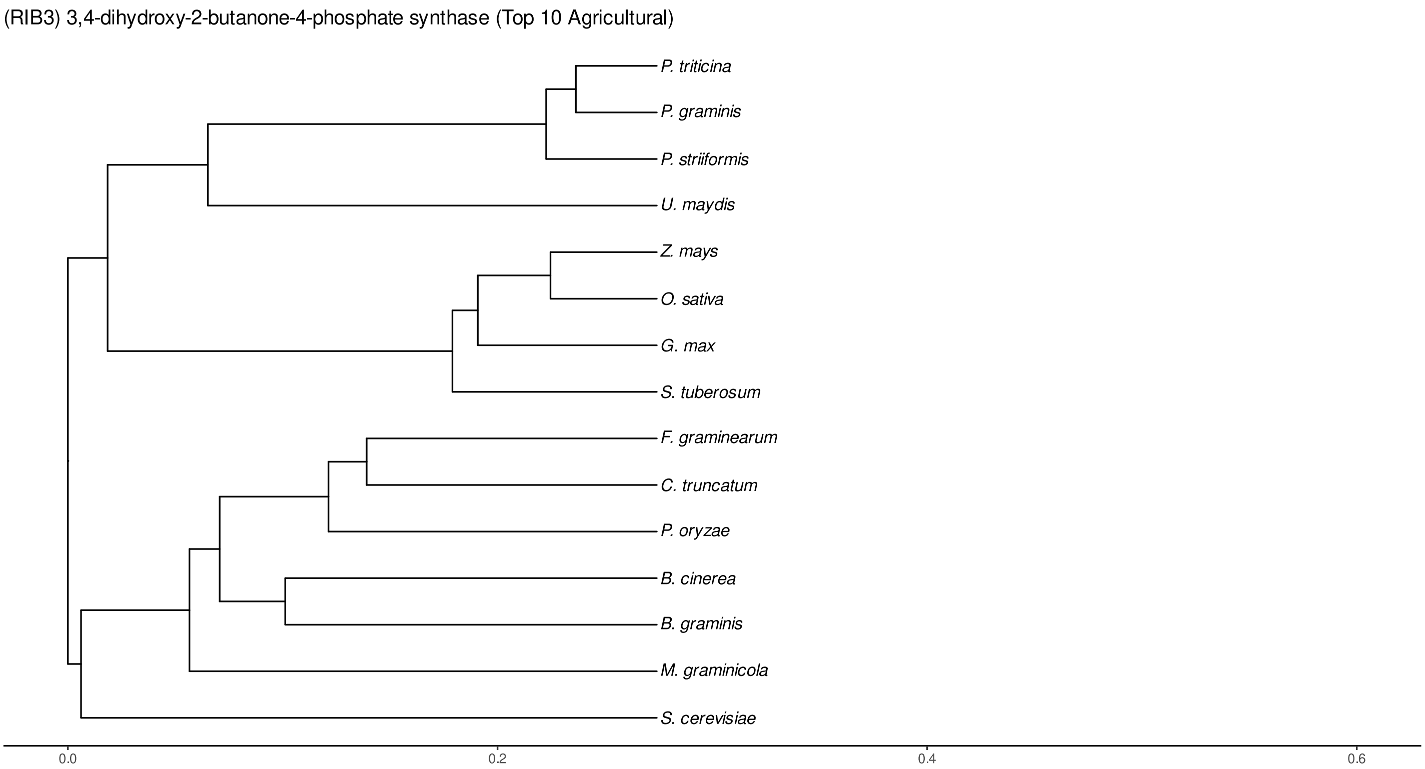


Figure S183: Phylogeny of hits for Rib3 made using clustal omega and visualized using ggtree, cf. Figures S180 and S181.

#### S4.12.3 NR

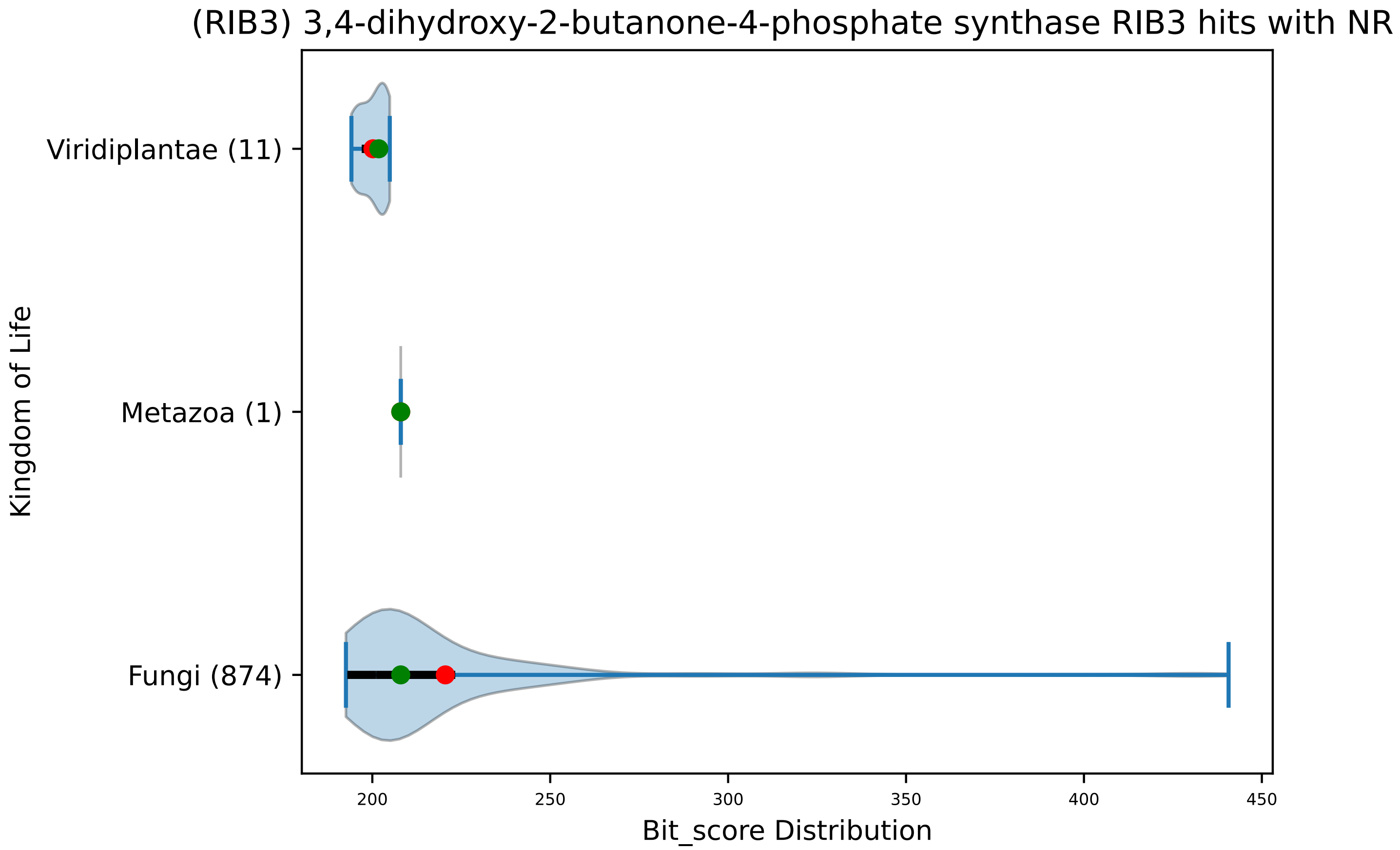


Figure S184: Non-redundant (NR) protein hits for DEG20010261/Rib3, with expectation value of no more than 0.1. Green points are medians, and red points are arithmetic means.

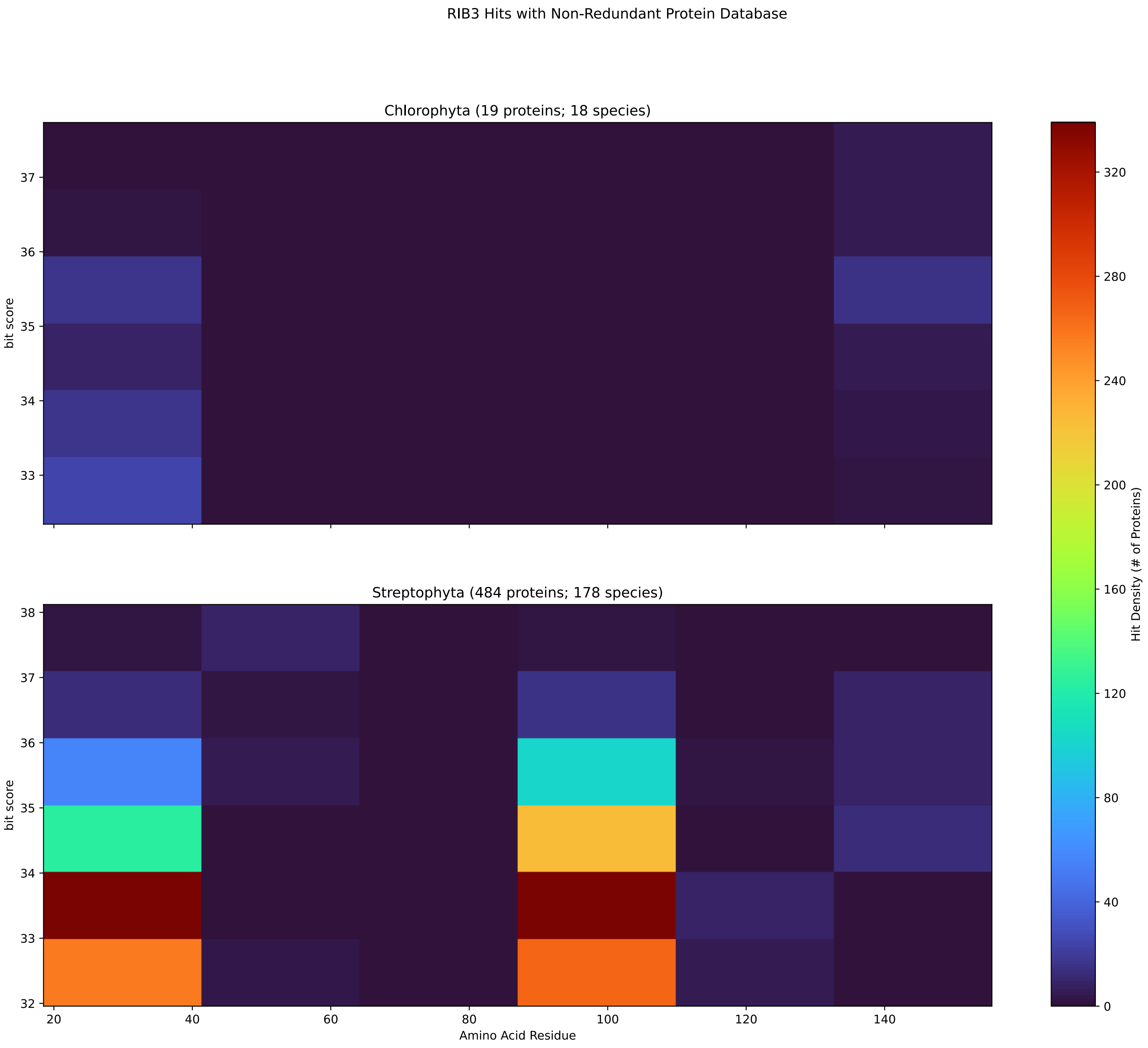


Figure S185: Non-redundant (NR) protein hits for Rib3 in the kingdom Viridiplantae.

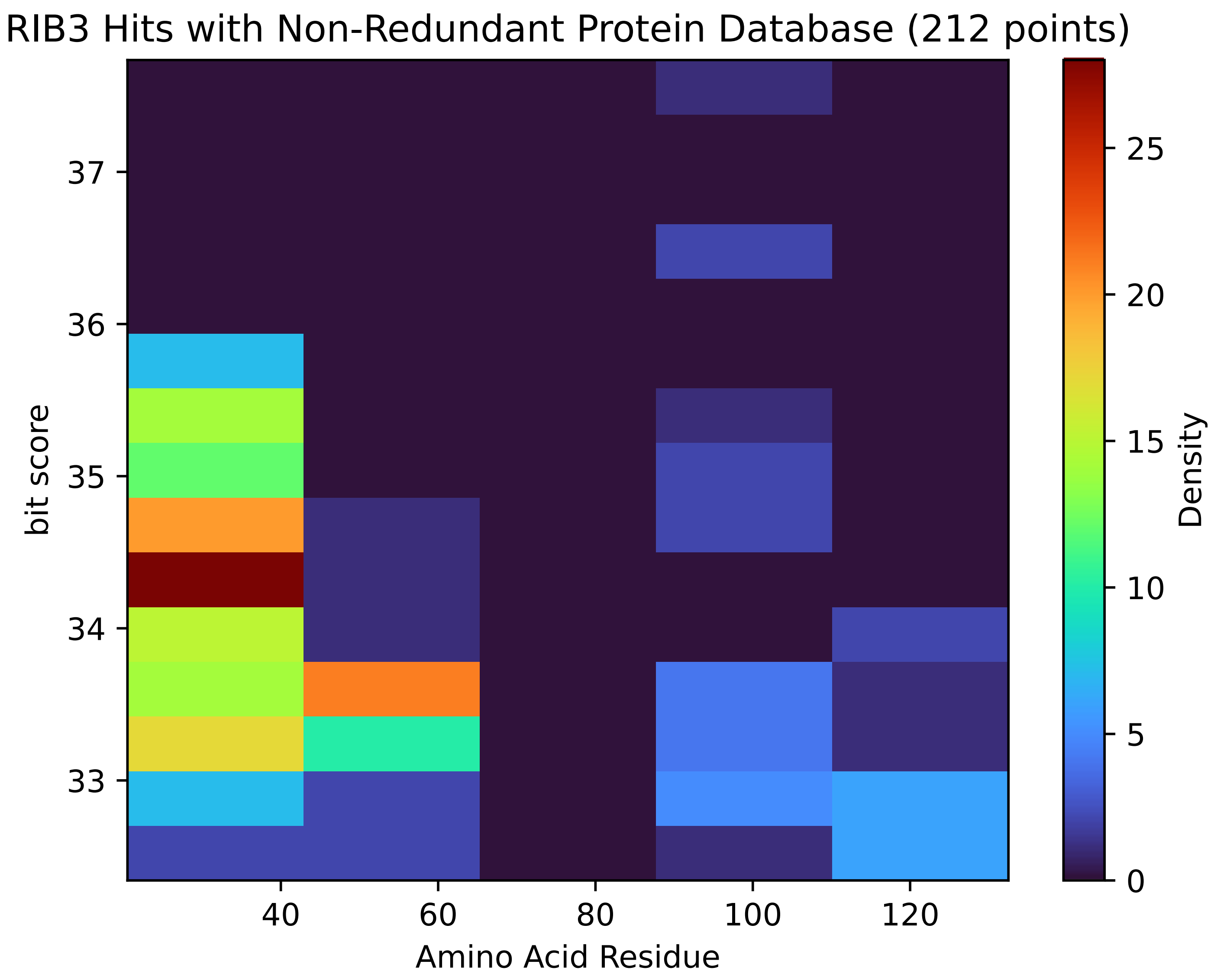


Figure S186: Non-redundant (NR) protein hits for Rib3 in the kingdom SAR.

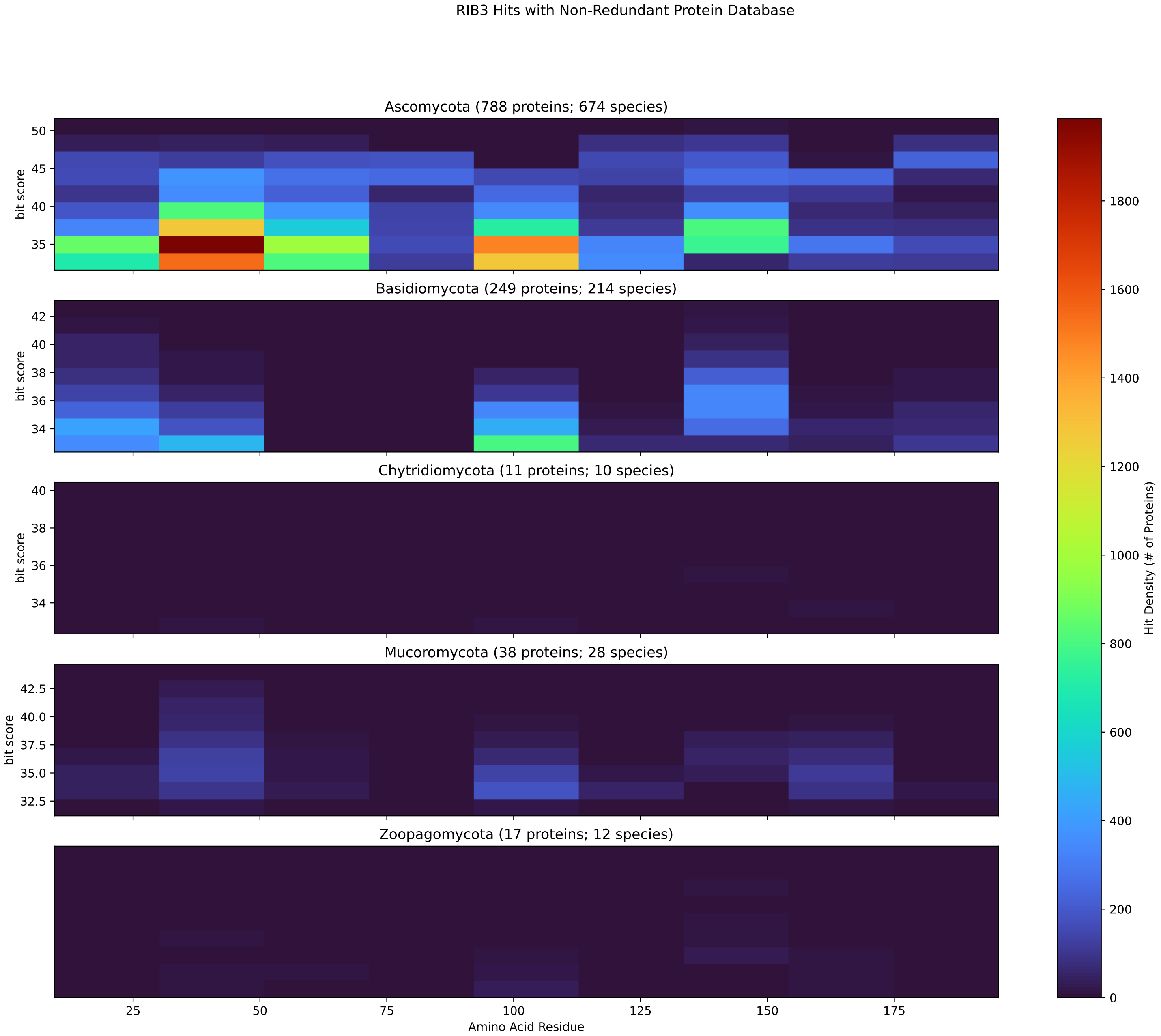


Figure S187: Non-redundant (NR) protein hits for Rib3 in the kingdom Fungi.

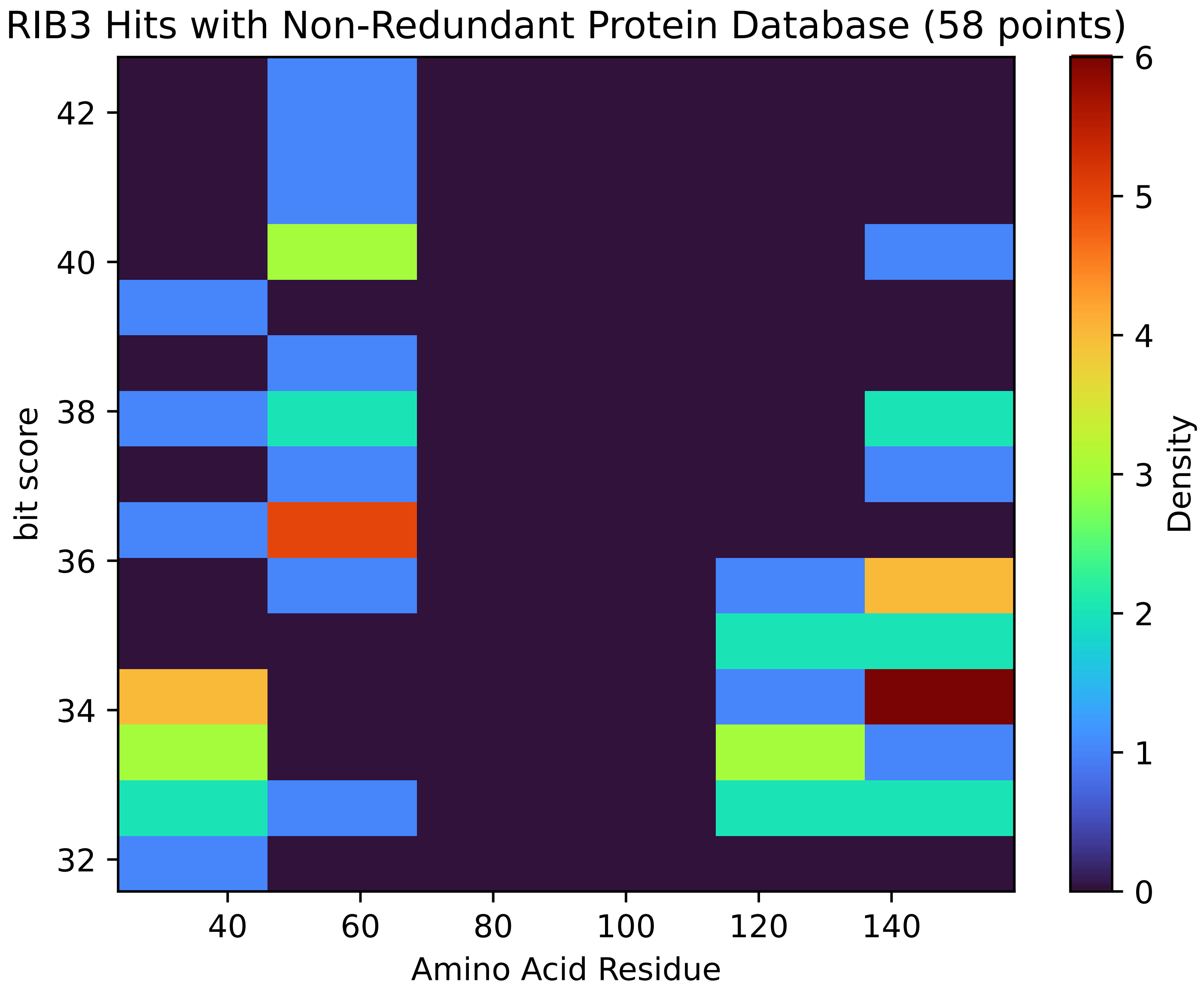


Figure S188: Non-redundant (NR) protein hits for Rib3 in the kingdom Metazoa.

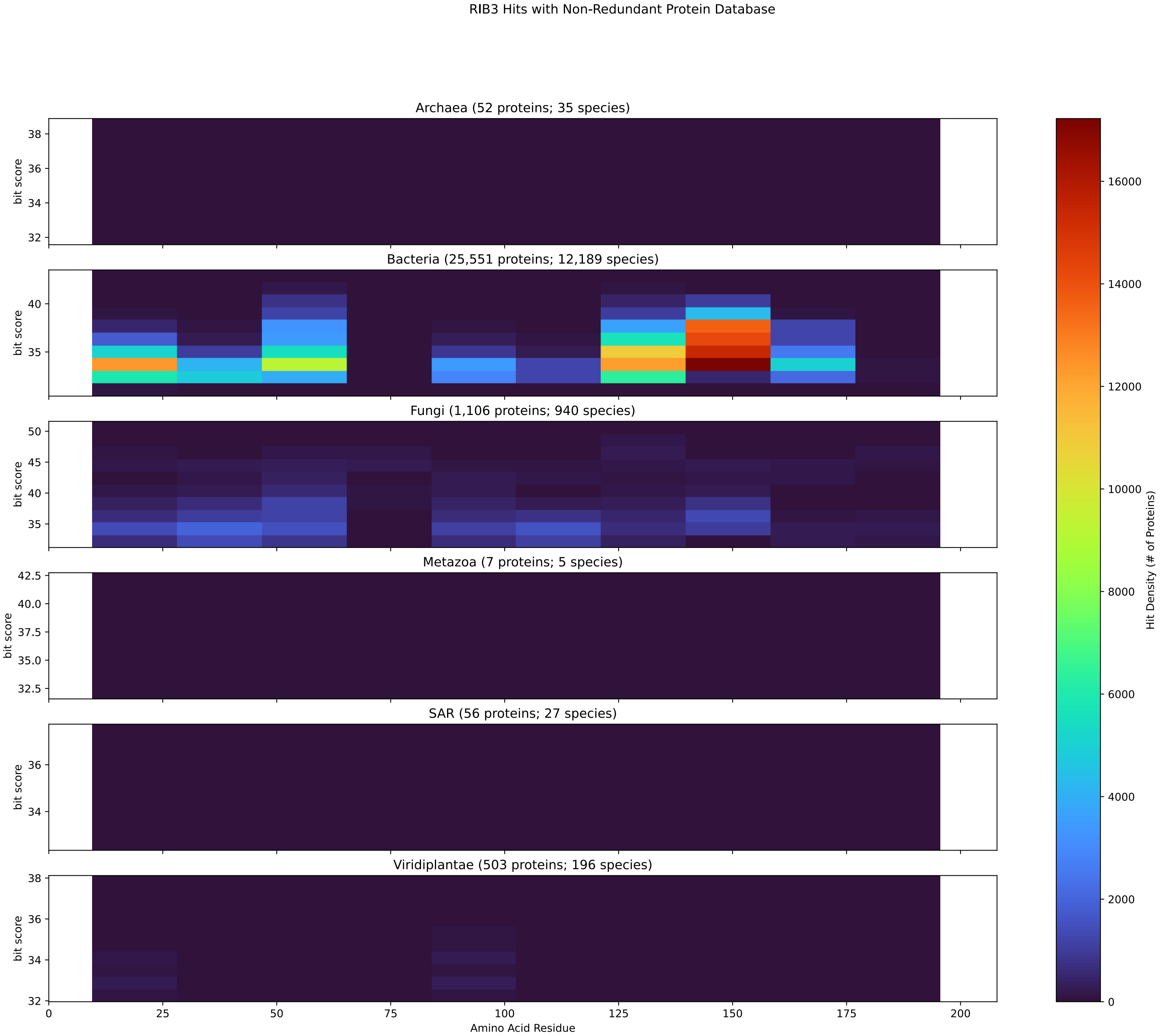


Figure S189: Non-redundant (NR) protein hits for DEG20010261/Rib3 at 20 amino acid length queries.

#### S4.13 Rib5

##### S4.13.1 WHO Critical Pathogens

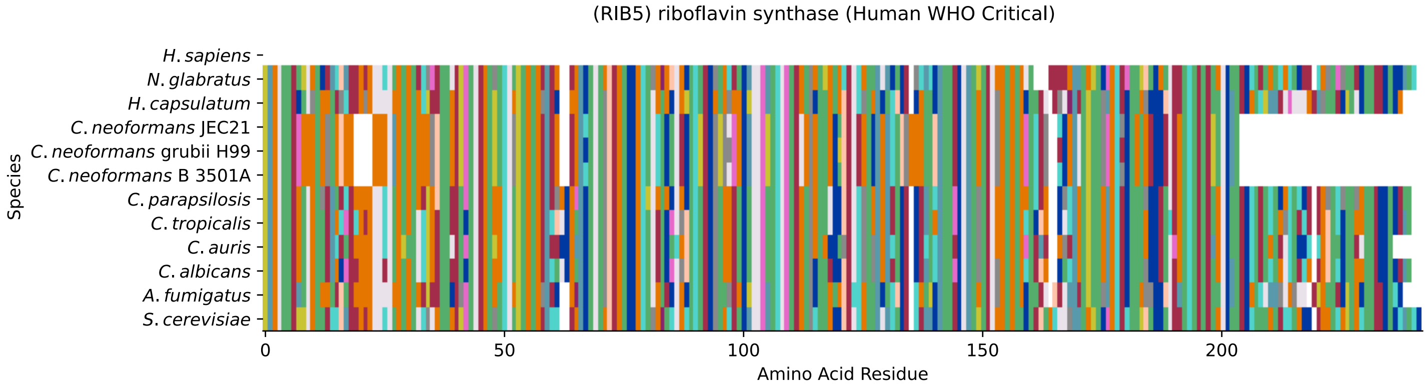


Figure S190: Multiple sequence alignment of yeast Rib5 (WHO Critical Pathogens). Cf. Figure S191 for alignment quality, and Figure S192 for Sneath similarity. Cf. Table S31 for protein names, and pairwise alignment metrics with yeast Rib5.

#### Rib5 MSA Quality

| | *A. f.* | *C. a.* | *C. a.* | *C. p.* | *C. t.* | *C. n.* | *C. n.* | *C. n.* | *H. c.* | *N. g.* | *S. c.* |
|---|---|---|---|---|---|---|---|---|---|---|---|
| *A. fumigatus* | 473.396 | 258.84 | 244.588 | 261.922 | 245.358 | 211.46 | 216.468 | 211.46 | 380.563 | 239.195 | 233.032 |
| *C. albicans* | 243.432 | 471.855 | 343.969 | 375.555 | 427.172 | 199.904 | 202.986 | 199.904 | 261.536 | 259.225 | 274.248 |
| *C. auris* | 228.409 | 343.969 | 464.922 | 338.576 | 340.502 | 202.986 | 204.912 | 202.986 | 249.98 | 261.536 | 255.758 |
| *C. parapsilosis* | 246.128 | 375.555 | 338.576 | 470.7 | 379.022 | 211.075 | 214.542 | 211.075 | 263.848 | 272.707 | 273.092 |
| *C. tropicalis* | 232.261 | 427.172 | 340.502 | 379.022 | 470.7 | 193.741 | 197.208 | 193.741 | 246.128 | 259.225 | 268.085 |
| *C. neoformans* B 3501A | 207.608 | 199.904 | 202.986 | 211.075 | 193.741 | 399.438 | 390.578 | 399.438 | 216.083 | 190.274 | 189.119 |
| *C. neoformans* grubii H99 | 211.846 | 202.986 | 204.912 | 214.542 | 197.208 | 390.578 | 400.208 | 390.578 | 221.09 | 193.741 | 194.512 |
| *C. neoformans* JEC21 | 207.608 | 199.904 | 202.986 | 211.075 | 193.741 | 399.438 | 390.578 | 399.438 | 216.083 | 190.274 | 189.119 |
| *H. capsulatum* | 363.614 | 261.536 | 249.98 | 263.848 | 246.128 | 216.083 | 221.09 | 216.083 | 474.937 | 236.498 | 238.039 |
| *N. glabratus* | 226.098 | 259.225 | 261.536 | 272.707 | 259.225 | 190.274 | 193.741 | 190.274 | 236.498 | 477.633 | 355.525 |
| *S. cerevisiae* | 219.164 | 274.248 | 255.758 | 273.092 | 268.085 | 189.119 | 194.512 | 189.119 | 238.039 | 355.525 | 484.182 |

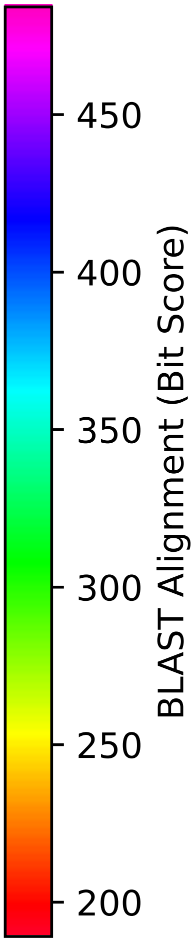


Figure S191: Multiple sequence alignment quality of Rib5 (WHO Critical Pathogens). Cf. Figures S190 and S193.

| Species | Hit Protein | Hit Length (a.a.) | evalue | align_len | bit_score | identity | positive | score | gaps | % identity | % positive |
|---|---|---|---|---|---|---|---|---|---|---|---|
| H.sapiens | - | - | - | - | - | - | - | - | - | - | - |
| N.glabratus | XP_445640.1 uncharacterized p-rotein CAGL0D05302g Nakaseomyc-es glabratus | 237 | 1.4e-126 | 237 | 356.295 | 172 | 205 | 913 | 3 | 72.3 | 86.1 |
| H.capsulatum | XP_045285331.1 riboflavin syn-thase Histoplasma capsulatum G-186AR | 235 | 8.5e-80 | 235 | 238.039 | 114 | 160 | 606 | 2 | 47.9 | 67.2 |
| C.neoformans.JEC21 | XP_569876.1 riboflavin syntha-se, putative Cryptococcus neof-ormans var. neoformans JEC21 | 202 | 9.8e-61 | 202 | 188.734 | 96 | 135 | 478 | 7 | 40.3 | 56.7 |
| C.neoformans.grubii.H99 | XP_012048272.1 riboflavin syn-thase, alpha subunit Cryptococ-cus neoformans var. grubii H99 | 202 | 8.4e-63 | 202 | 194.126 | 98 | 136 | 492 | 7 | 41.2 | 57.1 |
| C.neoformans.B.3501A | XP_776613.1 hypothetical prot-ein CNBC1060 Cryptococcus neof-ormans var. neoformans B-3501A | 202 | 9.4e-61 | 202 | 188.734 | 96 | 135 | 478 | 7 | 40.3 | 56.7 |
| C.parapsilosis | XP_036663168.1 uncharacterize-d protein CPAR2 100220 Candida parapsilosis | 238 | 1.1e-93 | 238 | 273.092 | 132 | 181 | 697 | 4 | 55.5 | 76.1 |
| C.tropicalis | XP_002546658.1 riboflavin syn-thase alpha chain Candida trop-icalis MYA-3404 | 238 | 7.9e-92 | 238 | 268.47 | 130 | 178 | 685 | 4 | 54.6 | 74.8 |
| C.auris | XP_028891770.1 riboflavin syn-thase, alpha subunit Candida a-uris | 234 | 3.4e-87 | 234 | 256.144 | 130 | 168 | 653 | 4 | 54.6 | 70.6 |
| C.albicans | XP_721932.2 riboflavin syntha-se Candida albicans SC5314 | 238 | 3.4e-94 | 238 | 274.248 | 134 | 176 | 700 | 4 | 56.3 | 73.9 |
| A.fumigatus | XP_750373.1 riboflavin syntha-se, alpha subunit Aspergillus fumigatus Af293 | 234 | 2.4e-72 | 234 | 219.164 | 115 | 157 | 557 | 1 | 48.3 | 66.0 |

Table S31: Pairwise alignment info from yeast Rib5 (DEG20010082), cf. Figure S190.

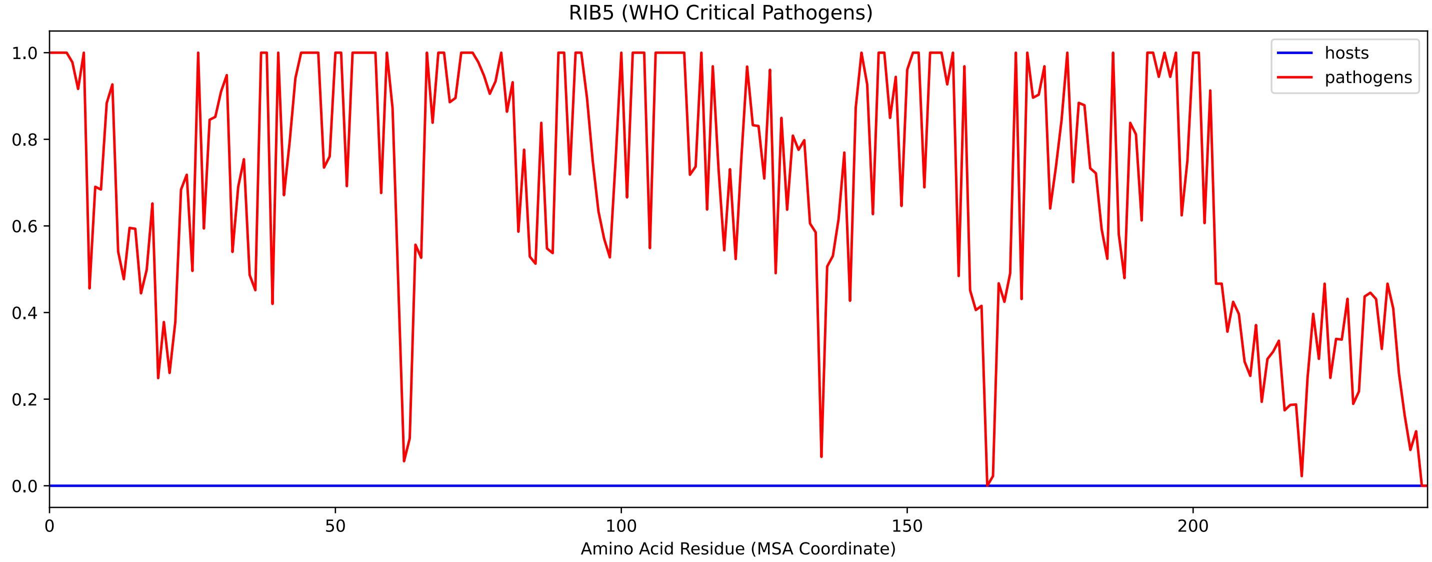


Figure S192: Sneath Similarity of Rib5 for WHO Critical Pathogens, cf. Figure S190

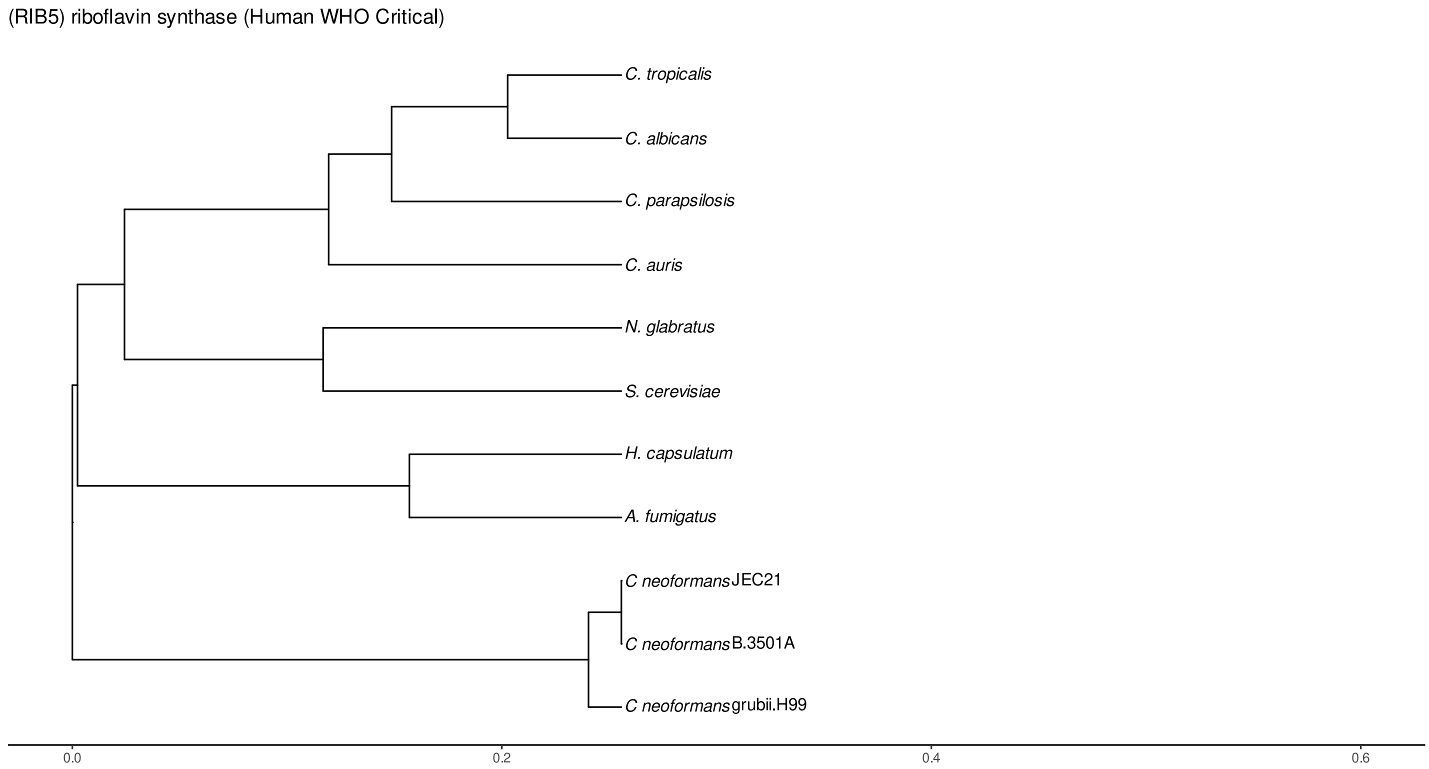


Figure S193: Phylogeny of hits for Rib5 made using clustal omega and visualized using ggtree, cf. Figures S190 and S191.

##### S4.13.2 Top 10 Agricultural Fungal Pathogens

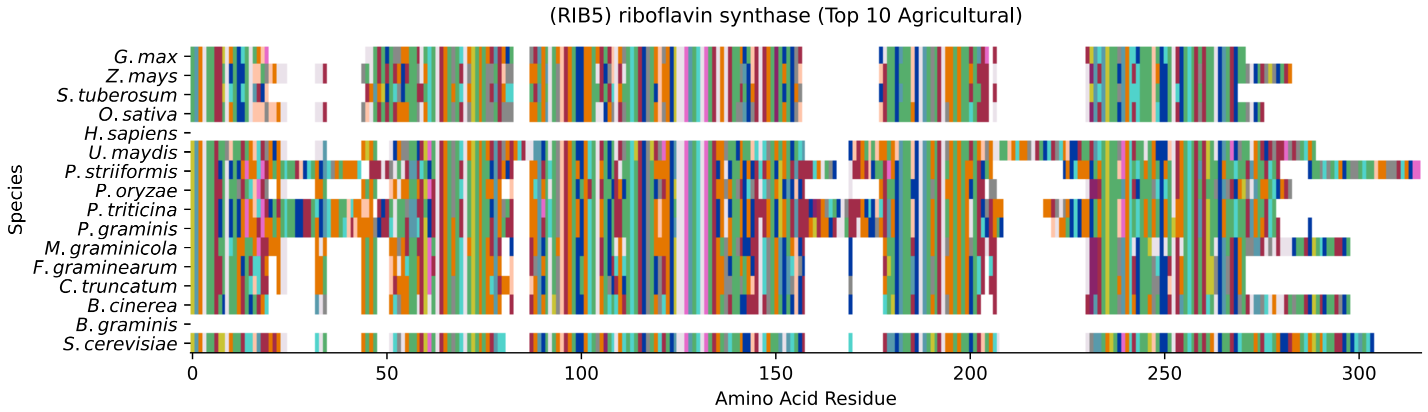


Figure S194: Multiple sequence alignment of yeast Rib5 (Top 10 Agricultural Fungal Pathogens). Cf. Figure S195 for alignment quality, and Figure S196 for Sneath similarity. Cf. Table S32 for protein names, and pairwise alignment metrics with yeast Rib5.

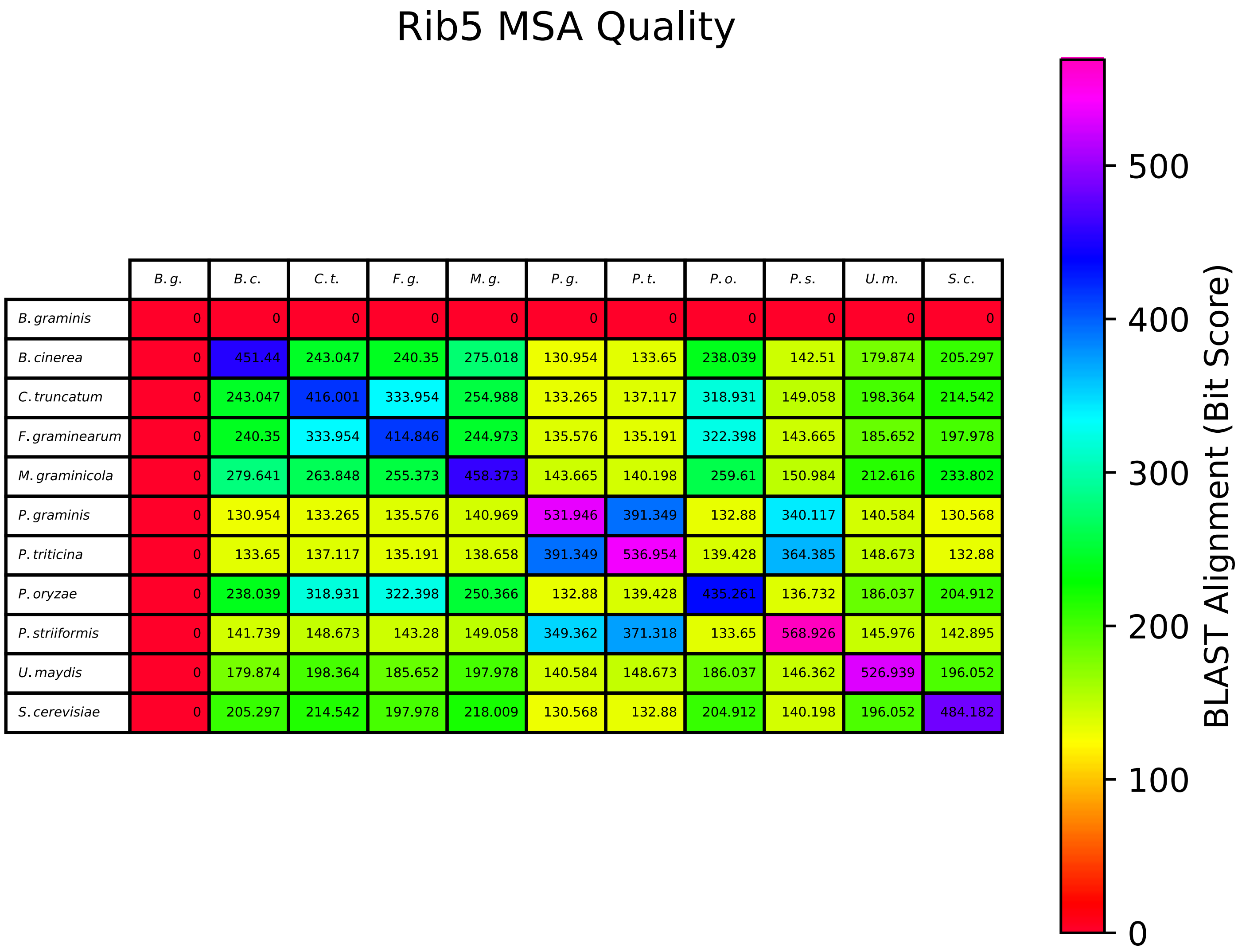


| | B. g. | B. c. | C. t. | F. g. | M. g. | P. g. | P. t. | P. o. | P. s. | U. m. | S. c. |
|---|---|---|---|---|---|---|---|---|---|---|---|
| B. graminis | 0 | 0 | 0 | 0 | 0 | 0 | 0 | 0 | 0 | 0 | 0 |
| B. cinerea | 0 | 451.44 | 243.047 | 240.35 | 275.018 | 130.954 | 133.65 | 238.039 | 142.51 | 179.874 | 205.297 |
| C. truncatum | 0 | 243.047 | 416.001 | 333.954 | 254.988 | 133.265 | 137.117 | 318.931 | 149.058 | 198.364 | 214.542 |
| F. graminearum | 0 | 240.35 | 333.954 | 414.846 | 244.973 | 135.576 | 135.191 | 322.398 | 143.665 | 185.652 | 197.978 |
| M. graminicola | 0 | 279.641 | 263.848 | 255.373 | 458.373 | 143.665 | 140.198 | 259.61 | 150.984 | 212.616 | 233.802 |
| P. graminis | 0 | 130.954 | 133.265 | 135.576 | 140.969 | 531.946 | 391.349 | 132.88 | 340.117 | 140.584 | 130.568 |
| P. triticina | 0 | 133.65 | 137.117 | 135.191 | 138.658 | 391.349 | 536.954 | 139.428 | 364.385 | 148.673 | 132.88 |
| P. oryzae | 0 | 238.039 | 318.931 | 322.398 | 250.366 | 132.88 | 139.428 | 435.261 | 136.732 | 186.037 | 204.912 |
| P. striiformis | 0 | 141.739 | 148.673 | 143.28 | 149.058 | 349.362 | 371.318 | 133.65 | 568.926 | 145.976 | 142.895 |
| U. maydis | 0 | 179.874 | 198.364 | 185.652 | 197.978 | 140.584 | 148.673 | 186.037 | 146.362 | 526.939 | 196.052 |
| S. cerevisiae | 0 | 205.297 | 214.542 | 197.978 | 218.009 | 130.568 | 132.88 | 204.912 | 140.198 | 196.052 | 484.182 |

Figure S195: Multiple sequence alignment quality of Rib5 (Top 10 Agricultural Fungal Pathogens). Cf. Figures S194 and S197.

| Species | Hit Protein | Hit Length (a.a.) | evalue | align_len | bit_score | identity | positive | score | gaps | % identity | % positive |
|---|---|---|---|---|---|---|---|---|---|---|---|
| G.max | XP_003539920.1 riboflavin synthase Glycine max | 207 | 1.5e-53 | 207 | 174.866 | 96 | 135 | 442 | 11 | 40.3 | 56.7 |
| Z.mays | NP_001149354.1 uncharacterized protein LOC100282978 Zea mays | 226 | 3.4e-49 | 226 | 164.081 | 87 | 137 | 414 | 15 | 36.6 | 57.6 |
| S.tuberosum | XP_006357493.1 PREDICTED: riboflavin synthase Solanum tuberosum | 205 | 2.4e-52 | 205 | 171.014 | 92 | 134 | 432 | 11 | 38.7 | 56.3 |
| O.sativa | XP_015618419.1 riboflavin synthase isoform X1 Oryza sativa Japonica Group | 216 | 4.8e-47 | 216 | 158.303 | 84 | 129 | 399 | 10 | 35.3 | 54.2 |
| H.sapiens | - | - | - | - | - | - | - | - | - | - | - |
| U.maydis | XP_011388024.1 riboflavin synthase Ustilago maydis 521 | 259 | 1e-62 | 259 | 196.438 | 104 | 152 | 498 | 36 | 43.7 | 63.9 |
| P.striiformis | XP_047801472.1 hypothetical protein Pst134EA 022744 Puccinia striiformis f. sp. tritici | 282 | 1.4e-40 | 282 | 139.813 | 97 | 145 | 351 | 47 | 40.8 | 60.9 |
| P.oryzae | mRNA M BR32 EuGene 00002111-p1 — transcript=mRNA M BR32 EuGene 00002111 — gene=M BR32 EuGene 00002111 — organism=Pyricularia oryzae BR32 — gene product=unspecified product — transcript product=unspecified product — location=BR32 scaffold00001:606531-607319(-) — protein length=214 — sequence SO=supercontig — SO=protein coding gene — is pseudo=false | 219 | 5.3e-67 | 219 | 204.912 | 101 | 149 | 520 | 6 | 42.4 | 62.6 |
| P.triticina | XP_053025945.1 uncharacterized protein PtA15 12A379 Puccinia triticina | 264 | 6.4e-38 | 264 | 132.88 | 91 | 133 | 333 | 51 | 38.2 | 55.9 |
| P.graminis | XP_003330596.1 riboflavin synthase, alpha subunit Puccinia graminis f. sp. tritici CRL 75-36-700-3 | 272 | 6.6e-37 | 272 | 130.568 | 92 | 128 | 327 | 64 | 38.7 | 53.8 |
| M.graminicola | ZTRI 2.263.mRNA-p1 — transcript=ZTRI 2.263.mRNA — gene=ZTRI 2.263 — organism=Zymoseptoria tritici IPO323 — gene product=similar to riboflavin synthase — transcript product=similar to riboflavin synthase — location=Ztri chr 2:855688-856441(−) — protein length=234 — sequence SO=chromosome — SO=protein coding gene — is pseudo=false | 232 | 5.2e-72 | 232 | 218.394 | 110 | 157 | 555 | 1 | 46.2 | 66.0 |
| F.graminearum | XP_011316898.1 riboflavin synthase alpha chain Fusarium graminearum PH-1 | 207 | 2.2e-63 | 207 | 196.823 | 100 | 142 | 499 | 8 | 42.0 | 59.7 |
| C.truncatum | XP_036583443.1 riboflavin synthase Colletotrichum truncatum | 207 | 4e-70 | 207 | 214.157 | 103 | 147 | 544 | 6 | 43.3 | 61.8 |
| B.cinerea | XP_024551623.1 Bcrib5 Botrytis cinerea B05.10 | 232 | 1.1e-66 | 232 | 204.912 | 108 | 149 | 520 | 9 | 45.4 | 62.6 |
| B.graminis | - | - | - | - | - | - | - | - | - | - | - |

Table S32: Pairwise alignment info from yeast Rib5 (DEG20010082), cf. Figure S194.

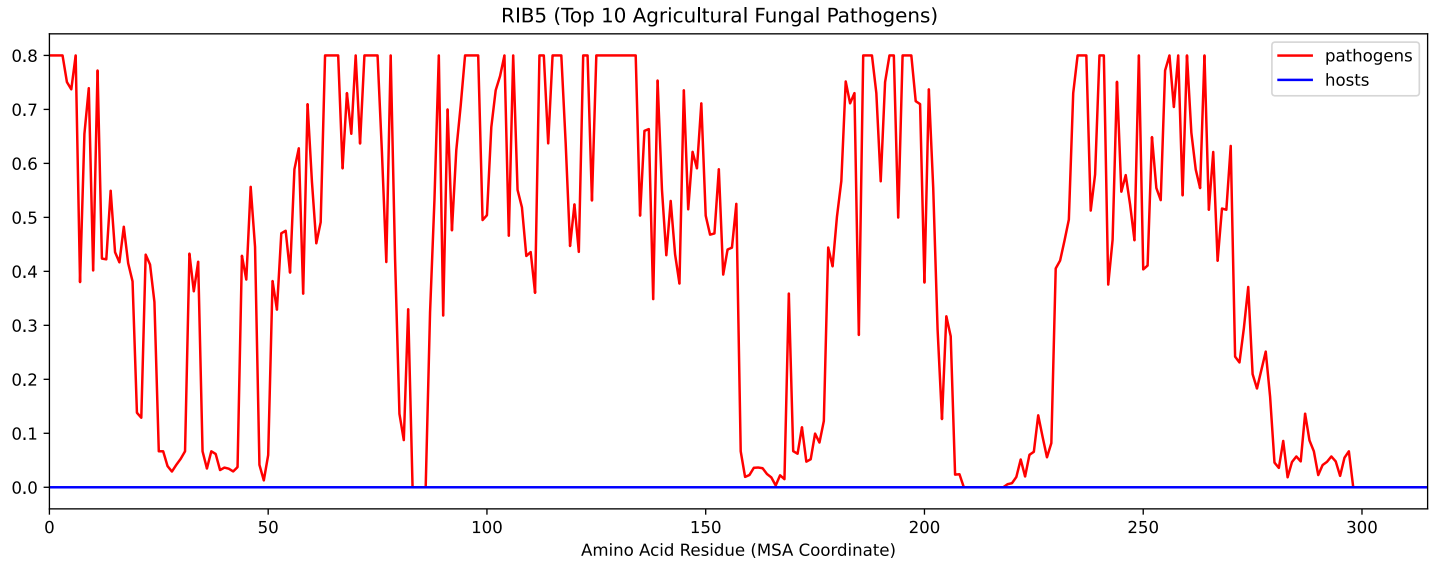


Figure S196: Sneath Similarity of Rib5 for Top 10 Agricultural Fungal Pathogens, cf. Figure S194

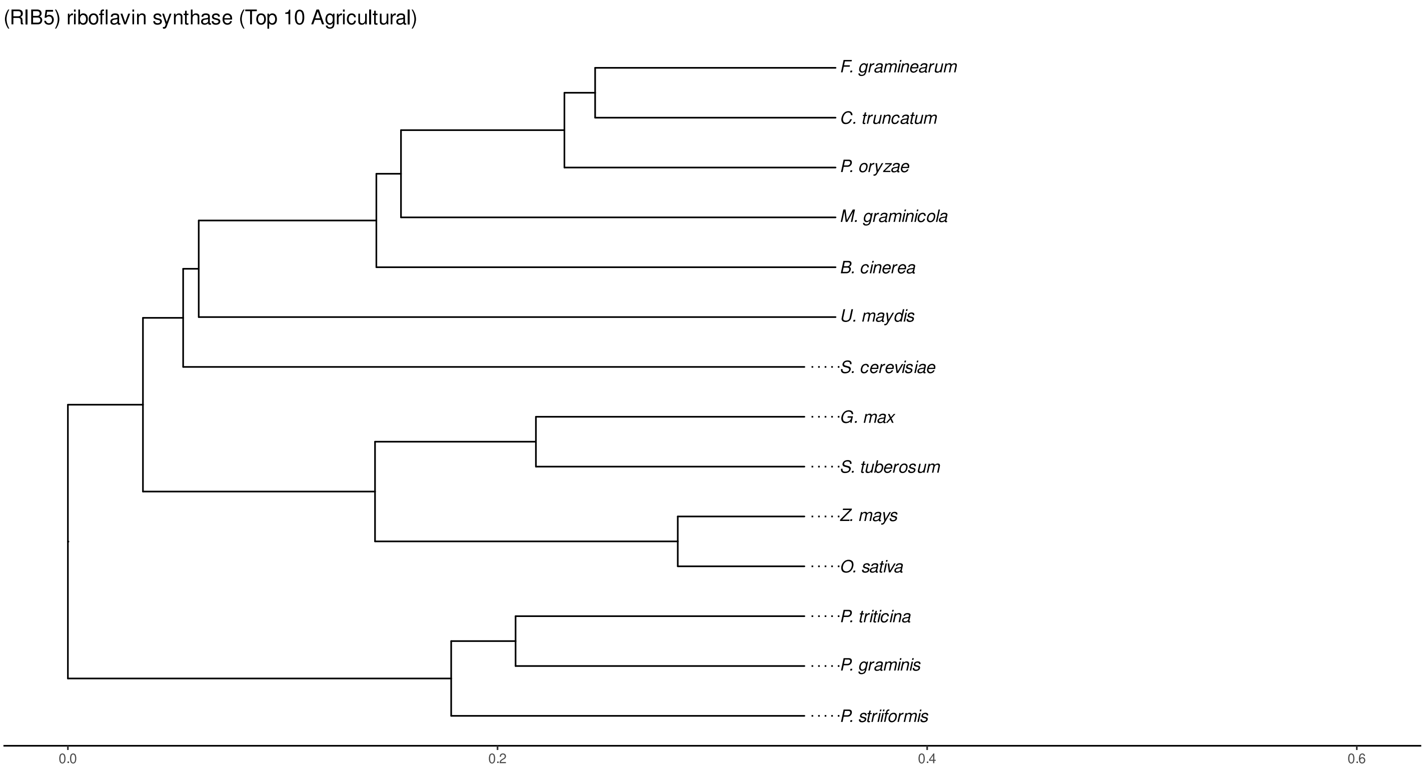


Figure S197: Phylogeny of hits for Rib5 made using clustal omega and visualized using ggtree, cf. Figures S194 and S195.

#### S4.13.3 NR

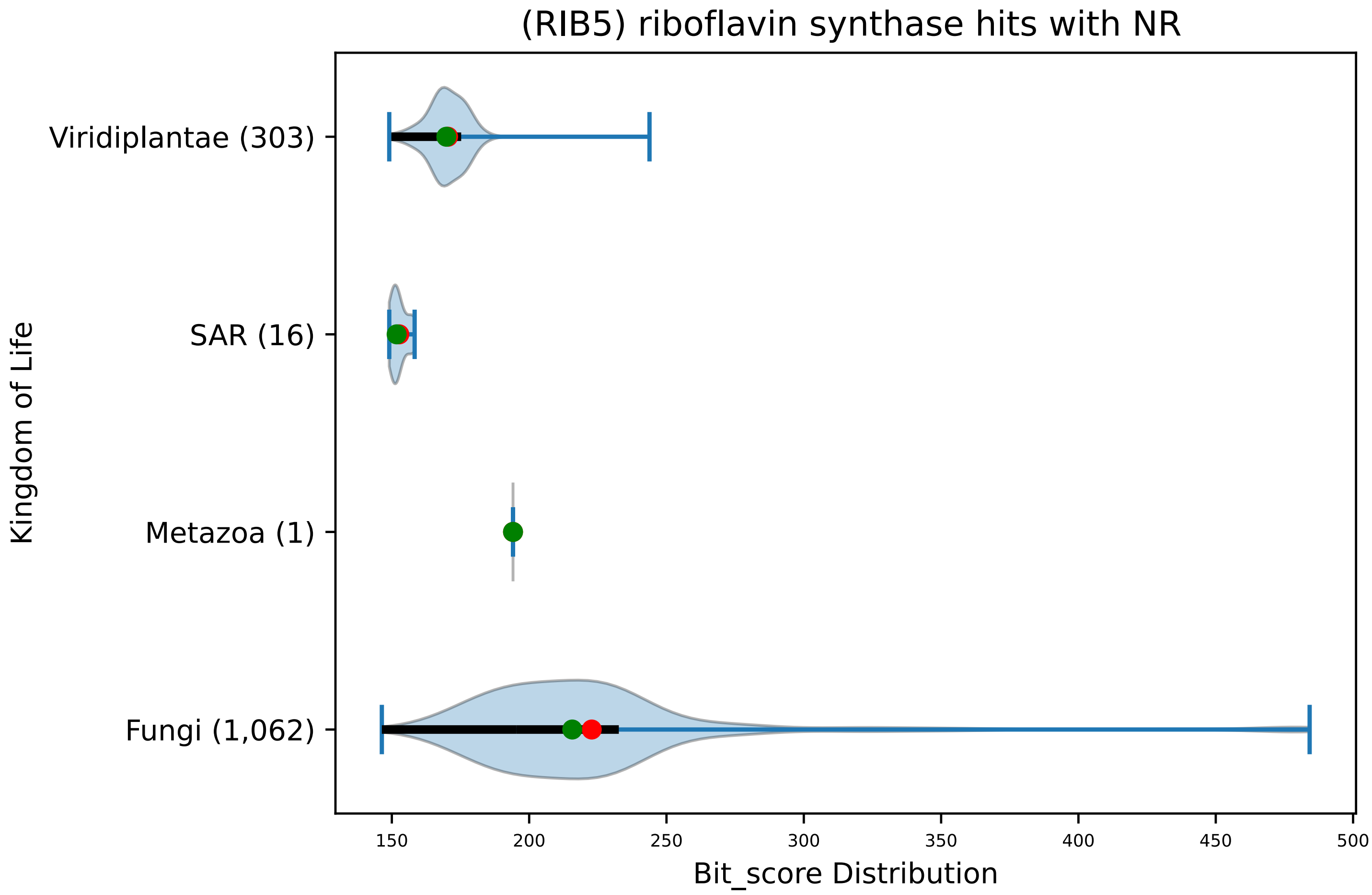


Figure S198: Non-redundant (NR) protein hits for DEG20010082/Rib5, with expectation value of no more than 0.1. Green points are medians, and red points are arithmetic means.

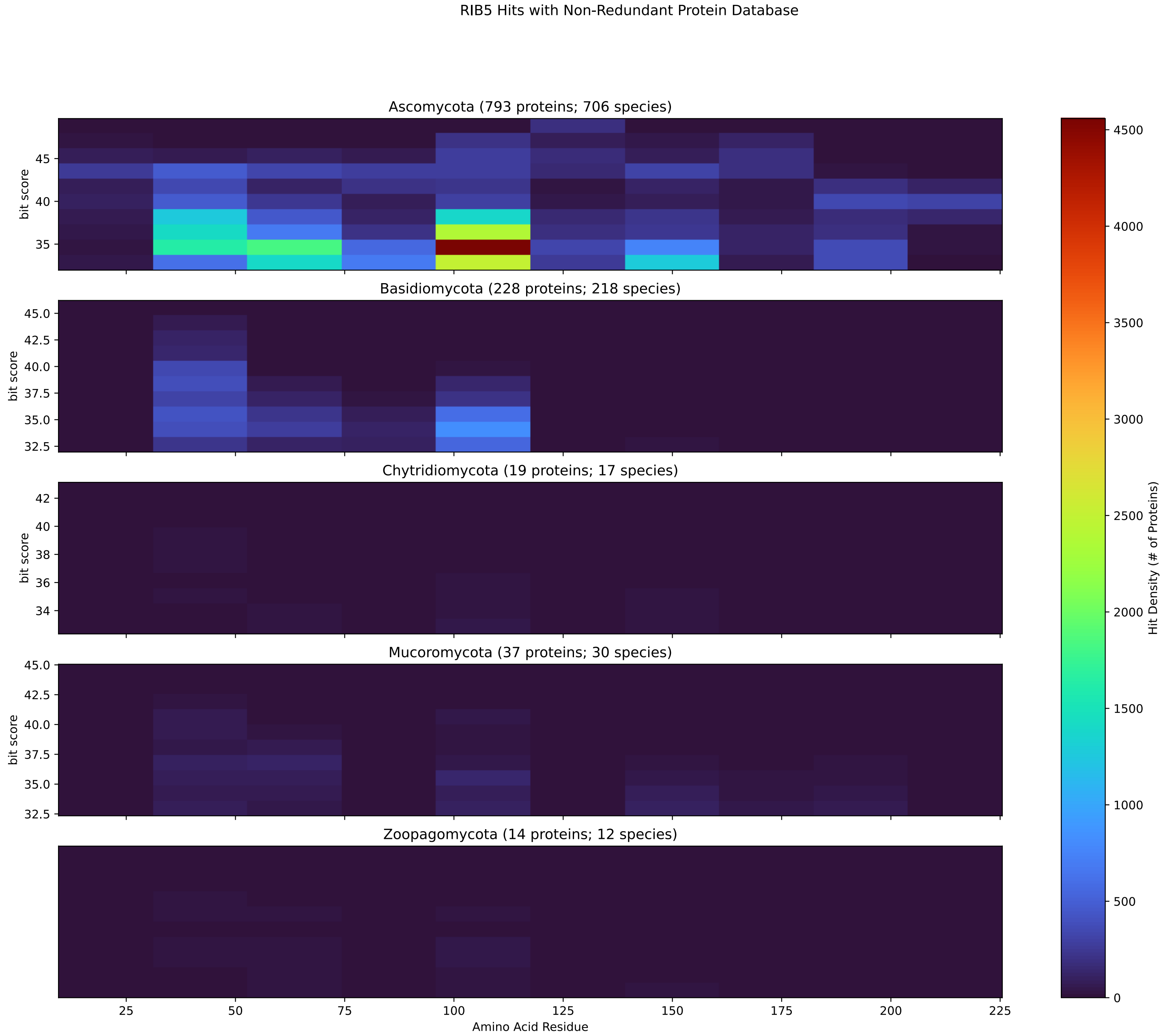


Figure S199: Non-redundant (NR) protein hits for Rib5 in the kingdom Fungi.

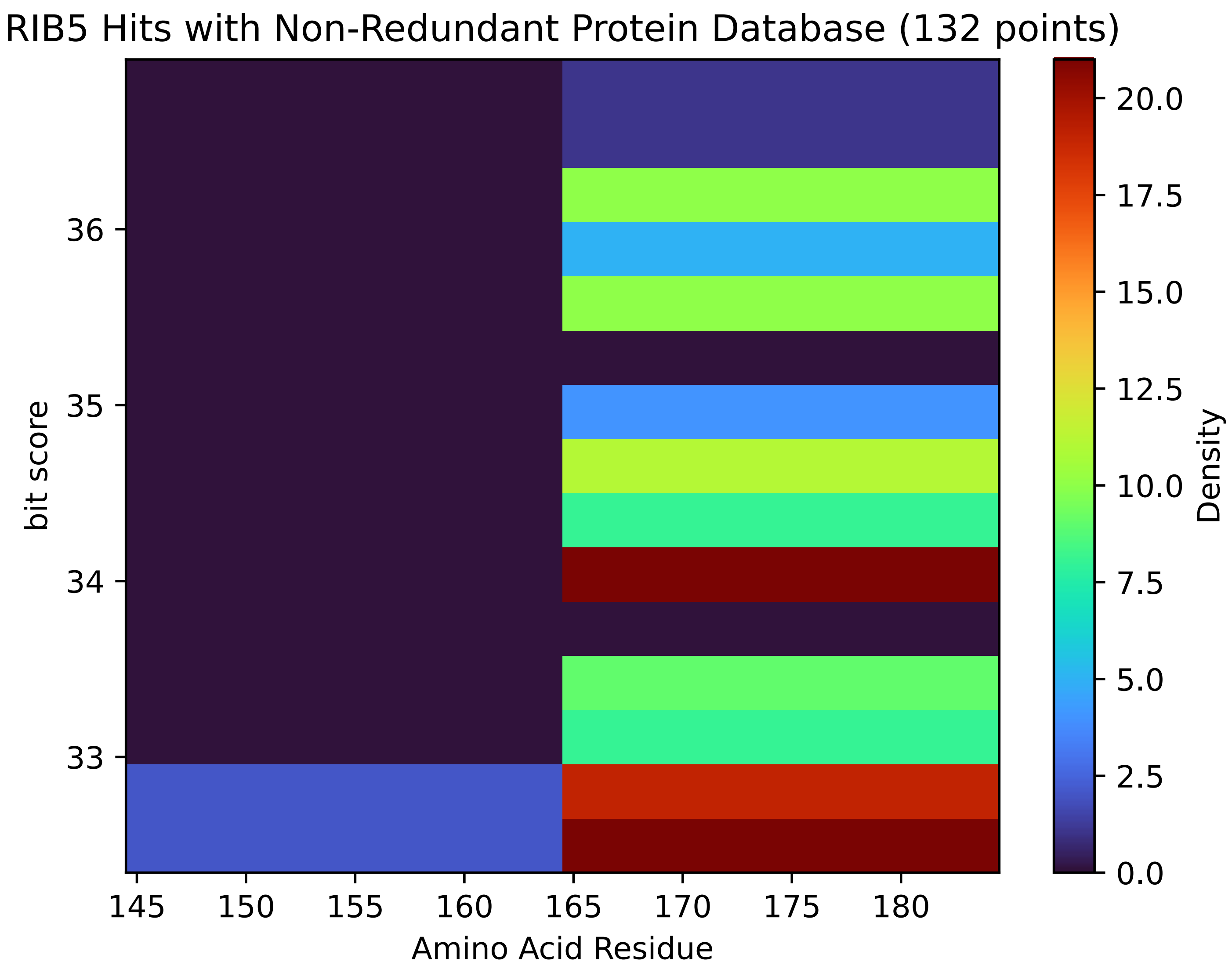


Figure S200: Non-redundant (NR) protein hits for Rib5 in the kingdom SAR.

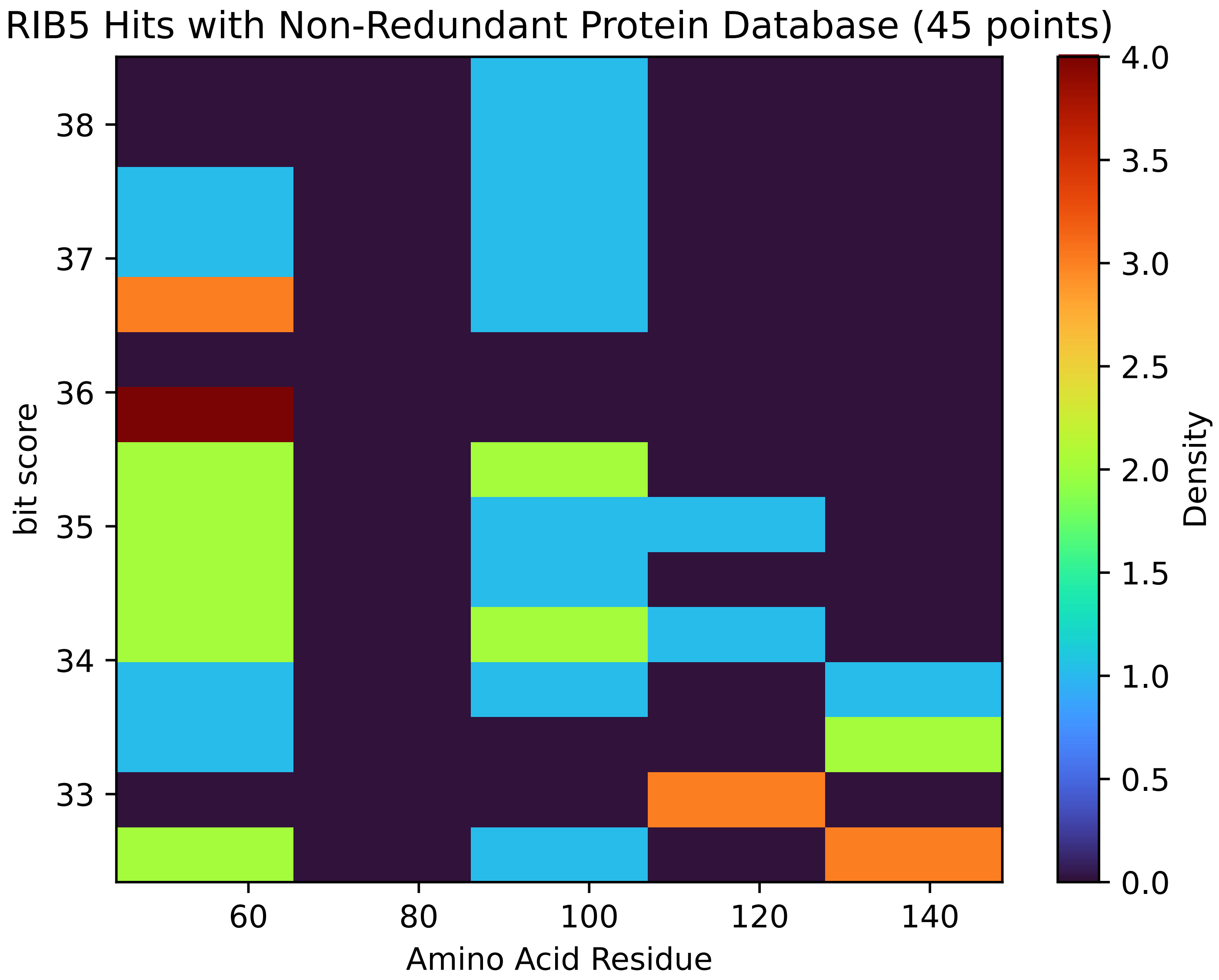


Figure S201: Non-redundant (NR) protein hits for Rib5 in the kingdom Metazoa.

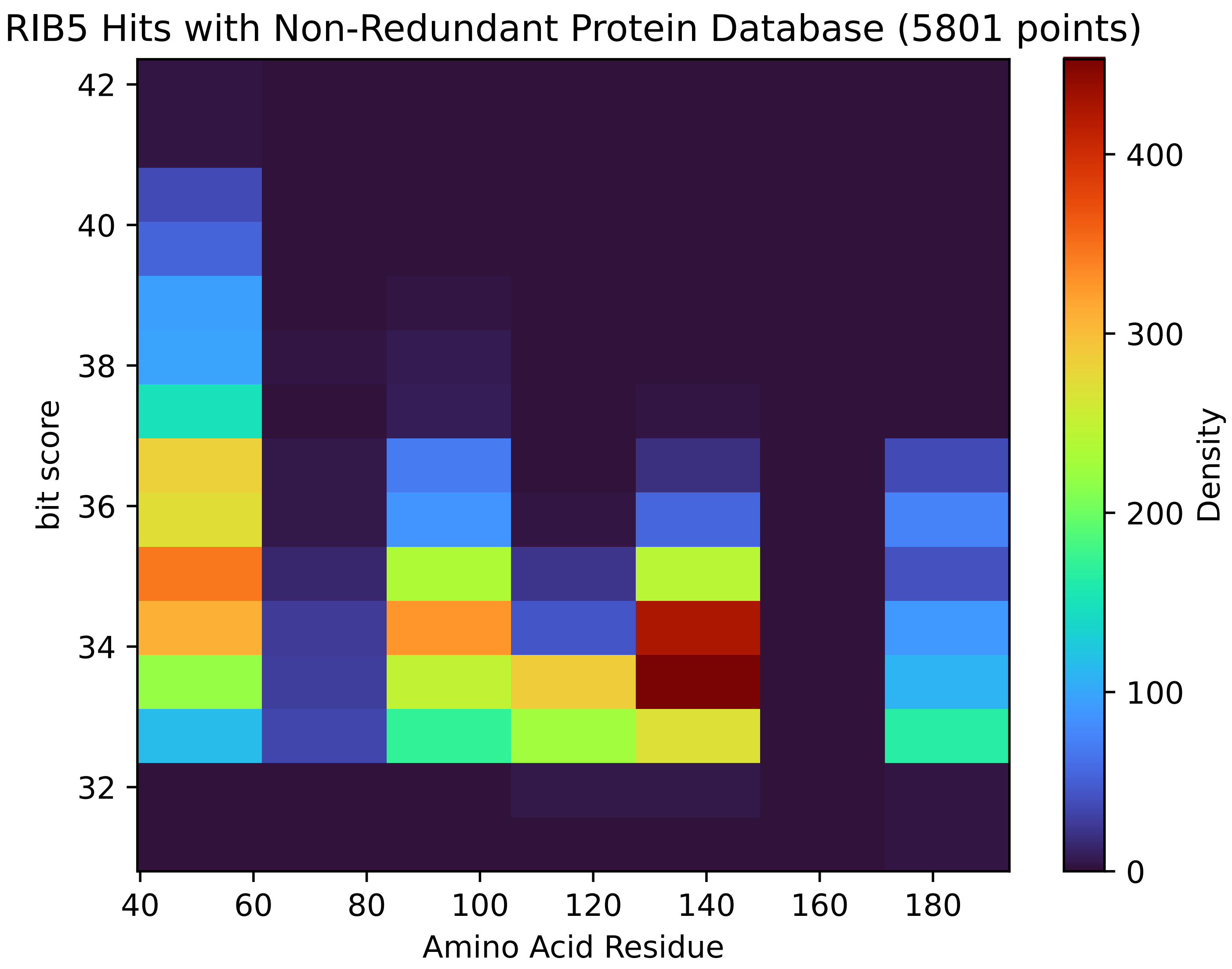


Figure S202: Non-redundant (NR) protein hits for Rib5 in the kingdom Viridiplantae.

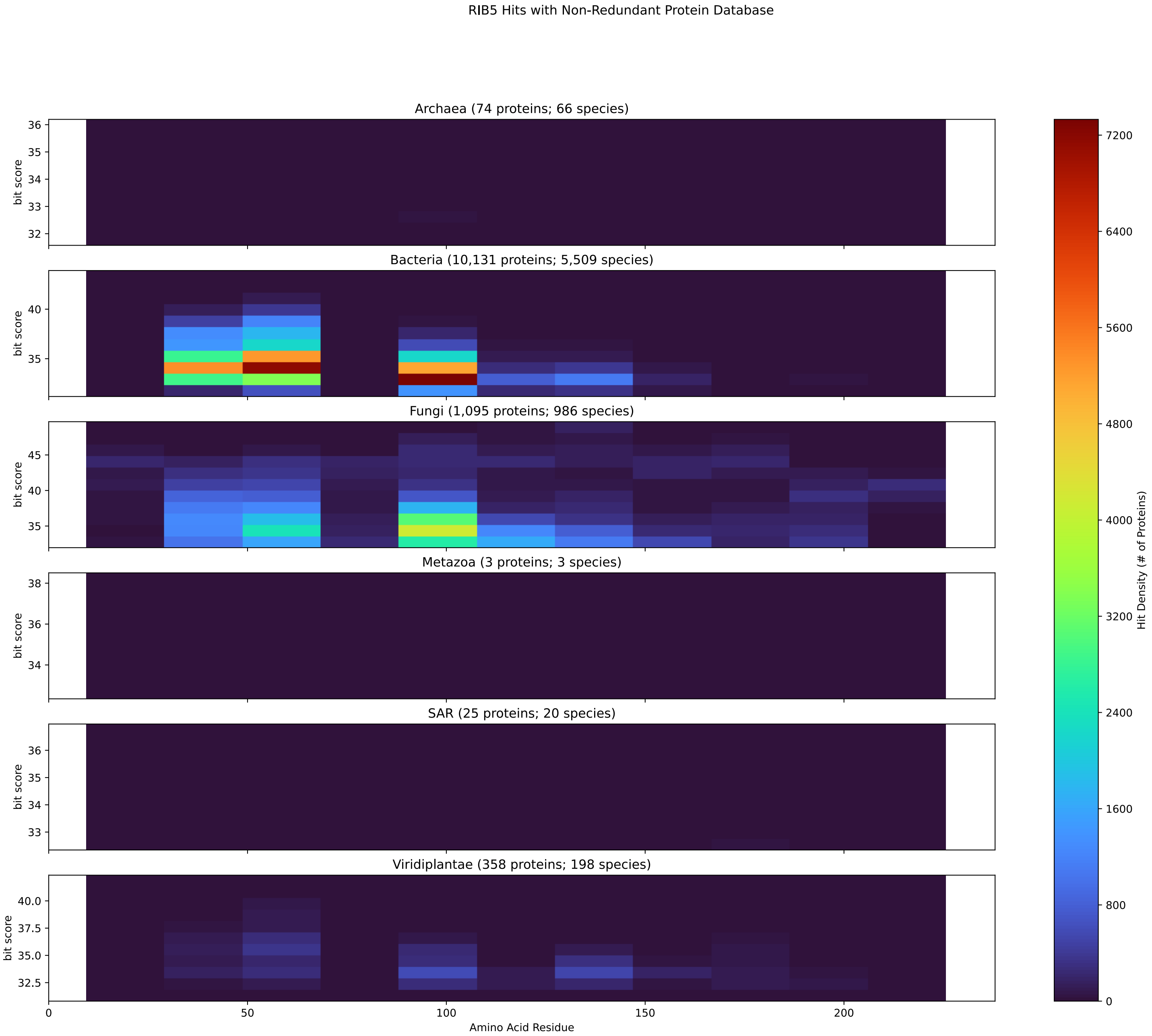


Figure S203: Non-redundant (NR) protein hits for DEG20010082/Rib5 at 20 amino acid length queries.

#### S4.14 Ssy1

##### S4.14.1 WHO Critical Pathogens

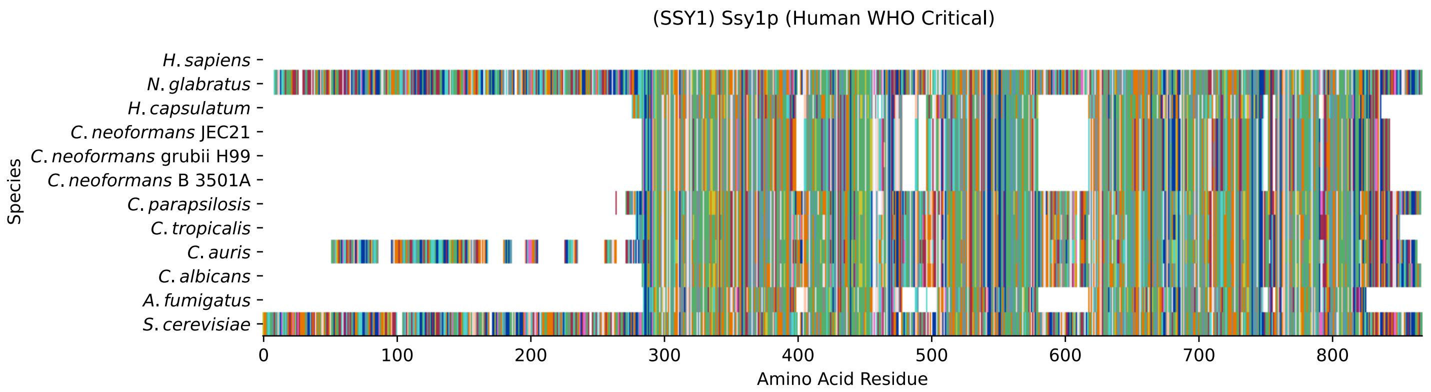


Figure S204: Multiple sequence alignment of yeast Ssy1 (WHO Critical Pathogens). Cf. Figure S205 for alignment quality, and Figure S206 for Sneath similarity. Cf. Table S33 for protein names, and pairwise alignment metrics with yeast Ssy1.

#### SSY1 MSA Quality

| | *A. f.* | *C. a.* | *C. a.* | *C. p.* | *C. t.* | *C. n.* | *C. n.* | *C. n.* | *H. c.* | *N. g.* | *S. c.* |
|---|---|---|---|---|---|---|---|---|---|---|---|
| *A. fumigatus* | 937.176 | 269.626 | 234.187 | 231.106 | 264.618 | 449.129 | 446.817 | 449.129 | 357.836 | 231.491 | 239.58 |
| *C. albicans* | 269.626 | 1174.84 | 806.209 | 907.516 | 1003.43 | 310.842 | 311.997 | 310.842 | 258.07 | 453.366 | 472.241 |
| *C. auris* | 244.588 | 822.002 | 1503.03 | 770.77 | 818.92 | 316.235 | 315.464 | 316.235 | 264.618 | 480.715 | 485.337 |
| *C. parapsilosis* | 240.35 | 926.006 | 771.926 | 1199.11 | 913.294 | 302.753 | 303.138 | 302.753 | 262.692 | 451.825 | 462.611 |
| *C. tropicalis* | 264.618 | 1003.43 | 803.897 | 899.042 | 1139.41 | 317.775 | 320.087 | 317.775 | 254.603 | 466.077 | 475.322 |
| *C. neoformans* B 3501A | 449.129 | 310.842 | 306.99 | 291.967 | 317.775 | 998.423 | 984.941 | 998.423 | 444.891 | 304.679 | 304.679 |
| *C. neoformans* grubii H99 | 446.817 | 311.997 | 306.22 | 291.967 | 320.087 | 984.941 | 998.038 | 984.941 | 444.121 | 308.916 | 303.138 |
| *C. neoformans* JEC21 | 449.129 | 310.842 | 306.99 | 291.967 | 317.775 | 998.423 | 984.941 | 998.423 | 444.891 | 304.679 | 304.679 |
| *H. capsulatum* | 357.836 | 258.07 | 259.225 | 253.447 | 254.603 | 444.891 | 444.121 | 444.891 | 984.941 | 247.284 | 257.299 |
| *N. glabratus* | 237.654 | 461.455 | 478.019 | 446.432 | 473.781 | 317.005 | 320.857 | 317.005 | 256.914 | 1739.16 | 922.924 |
| *S. cerevisiae* | 239.58 | 472.241 | 472.626 | 450.284 | 475.322 | 304.679 | 303.138 | 304.679 | 257.299 | 901.353 | 1758.42 |

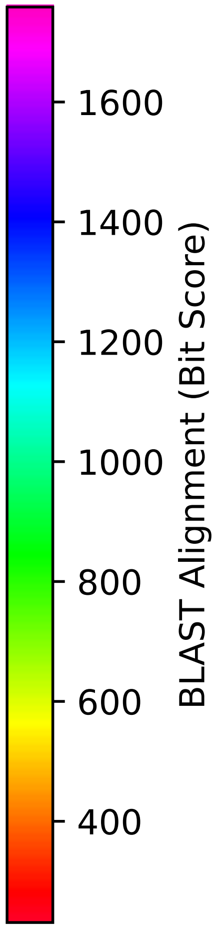


Figure S205: Multiple sequence alignment quality of Ssy1 (WHO Critical Pathogens). Cf. Figures S204 and S207.

| Species | Hit Protein | Hit Length (a.a.) | evalue | align_len | bit_score | identity | positive | score | gaps | % identity | % positive |
|---|---|---|---|---|---|---|---|---|---|---|---|
| H.sapiens | - | - | - | - | - | - | - | - | - | - | - |
| N.glabratus | XP_445735.1 uncharacterized protein CAGL0E01089g Nakaseomyces glabratus | 857 | 0 | 857 | 901.353 | 457 | 606 | 2328 | 23 | 53.6 | 71.1 |
| H.capsulatum | XP_045289907.1 arginine permease Histoplasma capsulatum G18-6AR | 553 | 1.9e-77 | 553 | 260.381 | 174 | 280 | 664 | 62 | 20.4 | 32.9 |
| C.neoformans.JEC21 | XP_568394.1 amino acid transporter, putative Cryptococcus neoformans var. neoformans JEC21 | 555 | 6.9e-95 | 555 | 307.76 | 181 | 300 | 787 | 61 | 21.2 | 35.2 |
| C.neoformans.grubii.H99 | XP_012052990.1 AAT family amino acid transporter Cryptococcus neoformans var. grubii H99 | 555 | 1.9e-94 | 555 | 306.605 | 183 | 298 | 784 | 61 | 21.5 | 35.0 |
| C.neoformans.B.3501A | XP_772149.1 hypothetical protein CNBM0690 Cryptococcus neoformans var. neoformans B-3501A | 555 | 6.6e-95 | 555 | 307.76 | 181 | 300 | 787 | 61 | 21.2 | 35.2 |
| C.parapsilosis | XP_036666456.1 uncharacterized protein CPAR2 209340 Candida parapsilosis | 603 | 7e-150 | 603 | 457.988 | 249 | 365 | 1177 | 24 | 29.2 | 42.8 |
| C.tropicalis | XP_002547148.1 hypothetical protein CTRG 01454 Candida tropicalis MYA-3404 | 575 | 3.8e-159 | 575 | 485.337 | 251 | 359 | 1248 | 19 | 29.5 | 42.1 |
| C.auris | XP_028890914.2 hypothetical protein Candida auris | 817 | 8.1e-156 | 817 | 473.396 | 290 | 439 | 1217 | 104 | 34.0 | 51.5 |
| C.albicans | XP_720938.2 Ssy1p Candida albicans SC5314 | 589 | 2.9e-157 | 589 | 480.33 | 254 | 374 | 1235 | 24 | 29.8 | 43.9 |
| A.fumigatus | XP_748191.1 amino acid permease, putative Aspergillus fumigatus Af293 | 535 | 7.6e-72 | 535 | 245.743 | 153 | 267 | 626 | 66 | 18.0 | 31.3 |

Table S33: Pairwise alignment info from yeast Ssy1 (DEG20010185), cf. Figure S204.

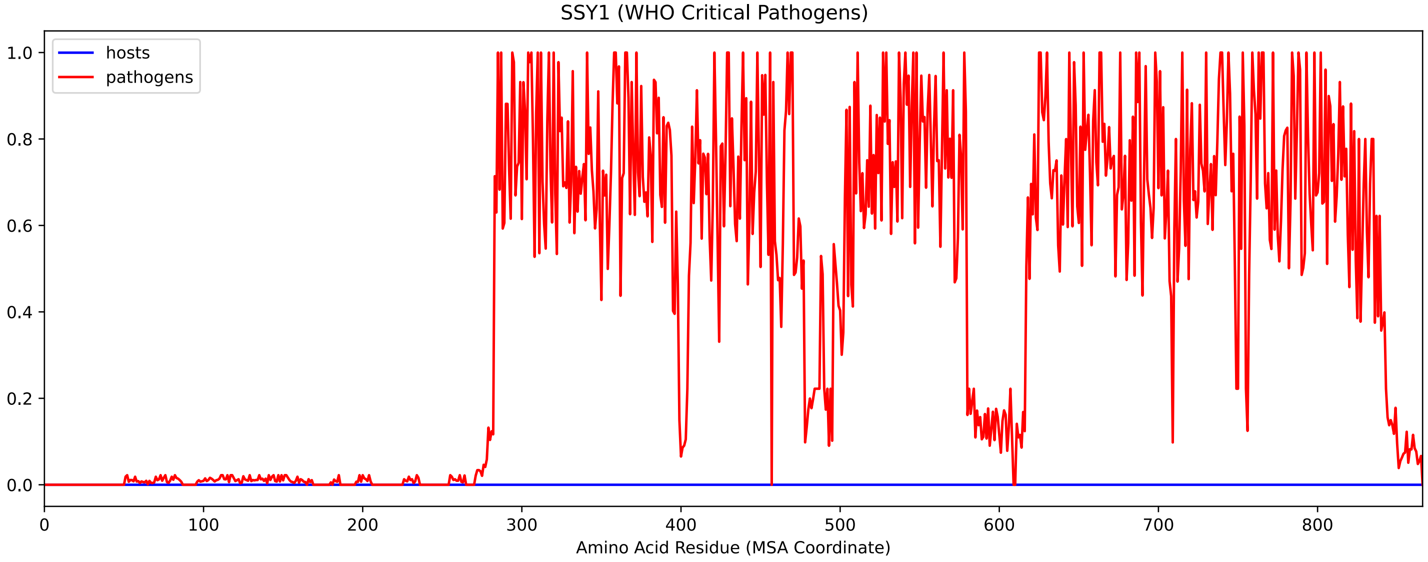


Figure S206: Sneath Similarity of Ssy1 for WHO Critical Pathogens, cf. Figure S204

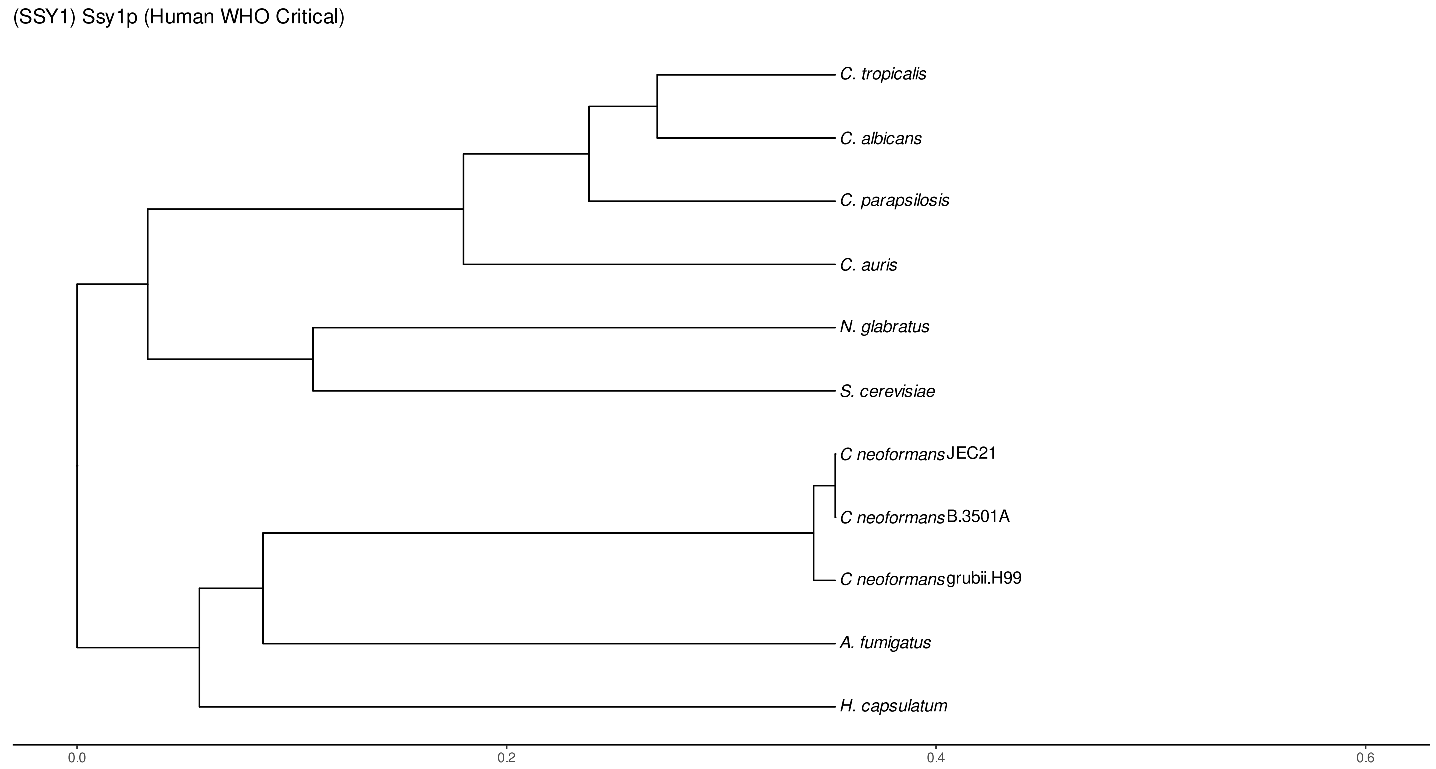


Figure S207: Phylogeny of hits for Ssy1 made using clustal omega and visualized using ggtree, cf. Figures S204 and S205.

##### S4.14.2 Top 10 Agricultural Fungal Pathogens

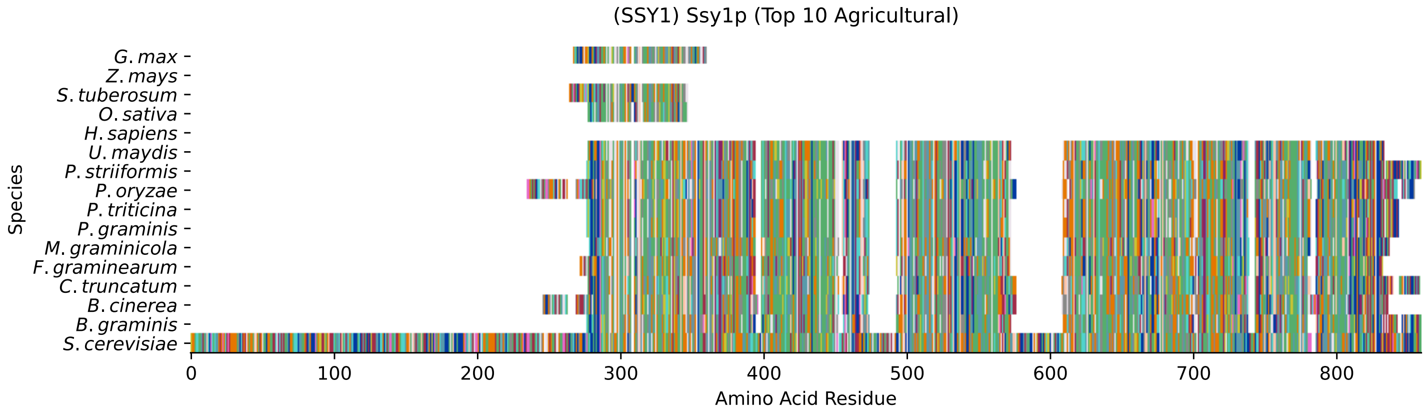


Figure S208: Multiple sequence alignment of yeast Ssy1 (Top 10 Agricultural Fungal Pathogens). Cf. Figure S209 for alignment quality, and Figure S210 for Sneath similarity. Cf. Table S34 for protein names, and pairwise alignment metrics with yeast Ssy1.

#### Ssy1 MSA Quality

| | *B.g.* | *B.c.* | *C.t.* | *F.g.* | *M.g.* | *P.g.* | *P.t.* | *P.o.* | *P.s.* | *U.m.* | *S.c.* |
|---|---|---|---|---|---|---|---|---|---|---|---|
| *B.graminis* | 1013.06 | 364.77 | 390.193 | 539.265 | 299.671 | 444.506 | 432.18 | 374.4 | 452.21 | 357.066 | 218.779 |
| *B.cinerea* | 369.007 | 1056.2 | 457.218 | 362.459 | 355.525 | 472.626 | 449.514 | 350.132 | 481.871 | 430.639 | 256.529 |
| *C.truncatum* | 402.905 | 458.373 | 1027.7 | 385.185 | 355.525 | 469.929 | 447.973 | 359.762 | 480.715 | 410.223 | 268.085 |
| *F.graminearum* | 539.265 | 351.673 | 373.629 | 980.319 | 308.145 | 434.106 | 424.476 | 354.369 | 436.802 | 373.629 | 250.366 |
| *M.graminicola* | 307.375 | 363.999 | 355.525 | 319.701 | 970.689 | 420.624 | 411.764 | 288.5 | 434.876 | 444.891 | 250.751 |
| *P.graminis* | 444.506 | 465.692 | 455.292 | 434.106 | 407.142 | 999.194 | 884.019 | 399.053 | 904.049 | 473.011 | 289.656 |
| *P.triticina* | 451.44 | 468.389 | 457.218 | 444.121 | 415.616 | 919.072 | 1001.5 | 400.979 | 930.243 | 481.1 | 303.908 |
| *P.oryzae* | 374.4 | 340.502 | 344.739 | 354.369 | 278.1 | 399.053 | 387.882 | 1103.2 | 391.349 | 353.599 | 236.498 |
| *P.striiformis* | 452.21 | 473.781 | 466.848 | 436.802 | 422.165 | 904.049 | 895.575 | 391.349 | 1031.55 | 479.174 | 295.434 |
| *U.maydis* | 375.941 | 444.506 | 410.223 | 392.504 | 447.588 | 491.115 | 473.011 | 364.77 | 496.123 | 968.378 | 273.092 |
| *S.cerevisiae* | 218.779 | 253.447 | 261.151 | 250.366 | 240.736 | 289.656 | 290.426 | 236.498 | 295.434 | 263.462 | 1758.42 |

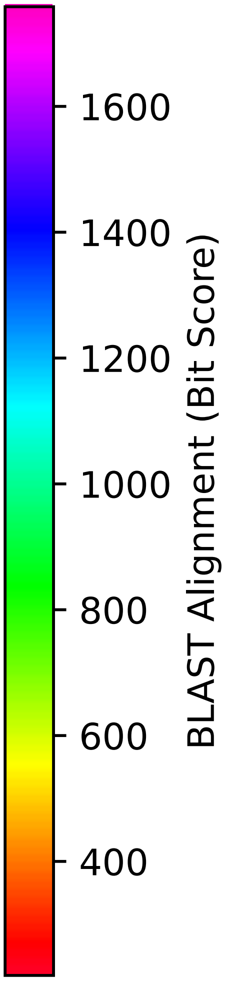


Figure S209: Multiple sequence alignment quality of Ssy1 (Top 10 Agricultural Fungal Pathogens). Cf. Figures S208 and S211.

| Species | Hit Protein | Hit Length (a.a.) | evalue | align_len | bit_score | identity | positive | score | gaps | % identity | % positive |
|---|---|---|---|---|---|---|---|---|---|---|---|
| G.max | XP_003555221.1 cationic amino acid transporter 7, chloroplastic Glycine max | 92 | 0.019 | 92 | 39.6614 | 29 | 46 | 91 | 2 | 3.4 | 5.4 |
| Z.mays | - | - | - | - | - | - | - | - | - | - | - |
| S.tuberosum | XP_006353075.1 PREDICTED: cationic amino acid transporter 1--like Solanum tuberosum | 90 | 0.00083 | 90 | 43.1282 | 29 | 49 | 100 | 9 | 3.4 | 5.8 |
| O.sativa | XP_015627412.1 cationic amino acid transporter 9, chloroplastic Oryza sativa Japonica Group | 69 | 0.05 | 69 | 37.7354 | 22 | 35 | 86 | 1 | 2.6 | 4.1 |
| H.sapiens | - | - | - | - | - | - | - | - | - | - | - |
| U.maydis | XP_011389724.1 putative general amino acid permease Ustilago maydis 521 | 552 | 2e-79 | 552 | 266.159 | 171 | 276 | 679 | 68 | 20.1 | 32.4 |
| P.striiformis | XP_047811281.1 hypothetical protein Pst134EA 002462 Puccinia striiformis f. sp. tritici | 578 | 1.1e-91 | 578 | 299.286 | 180 | 295 | 765 | 67 | 21.1 | 34.6 |
| P.oryzae | mRNA M BR32 EuGene 00049831-p1 — transcript=mRNA M BR32 EuGene 00049831 — gene=M BR32 EuGene 00049831 — organism=Pyricularia oryzae BR32 — gene product=unspecified product — transcript product=unspecified product — location=BR32 scaffold000-04:1622099-1624325(-) — protein length=549 — sequence SO=supercontig — SO=protein coding gene — is pseudo=false | 610 | 8.1e-69 | 610 | 236.884 | 168 | 287 | 603 | 71 | 19.7 | 33.7 |
| P.triticina | XP_053023344.1 uncharacterized protein PtA15 8A695 Puccinia triticina | 561 | 5.9e-90 | 561 | 294.278 | 180 | 288 | 752 | 66 | 21.1 | 33.8 |
| P.graminis | XP_003329009.2 AAT family amino acid transporter Puccinia graminis f. sp. tritici CRL 75–36-700-3 | 561 | 4.1e-89 | 561 | 292.352 | 170 | 292 | 747 | 66 | 20.0 | 34.3 |
| M.graminicola | ZTRI 4.872.mRNA-p1 — transcript=ZTRI 4.872.mRNA — gene=ZTRI 4.872 — organism=Zymoseptoria tritici IPO323 — gene product=-similar to amino acid permease — transcript product=similar to amino acid permease — location=Ztri chr 4:2785263-2787024-(-) — protein length=550 — sequence SO=chromosome — SO=protein coding gene — is pseudo=false | 555 | 2.5e-71 | 555 | 243.817 | 171 | 281 | 621 | 69 | 20.1 | 33.0 |
| F.graminearum | XP_011326118.1 hypothetical protein FGSG 06508 Fusarium graminearum PH-1 | 554 | 3.5e-75 | 554 | 253.447 | 173 | 272 | 646 | 67 | 20.3 | 31.9 |
| C.truncatum | XP_036576509.1 amino acid permease Colletotrichum truncatum | 575 | 9.6e-79 | 575 | 265.003 | 172 | 285 | 676 | 65 | 20.2 | 33.5 |
| B.cinerea | XP_001552845.1 hypothetical protein BCIN 14g01410 Botrytis cinerea B05.10 | 589 | 9.5e-76 | 589 | 256.914 | 172 | 289 | 655 | 71 | 20.2 | 33.9 |
| B.graminis | VDB92912.1 — transcript=BGT962-24V316 LOCUS6671 t1 — gene=BGT-96224V316 LOCUS6671 — organism-=Blumeria graminis f. sp. tritici 96224 — gene product=unspecified product — transcript product=unspecified product — location=LR026992:5342891-534455-2(+) — protein length=536 — sequence SO=chromosome — SO=protein coding gene — is pseudo=false | 584 | 3.1e-64 | 584 | 223.402 | 162 | 281 | 568 | 83 | 19.0 | 33.0 |

Table S34: Pairwise alignment info from yeast Ssy1 (DEG20010185), cf. Figure S208.

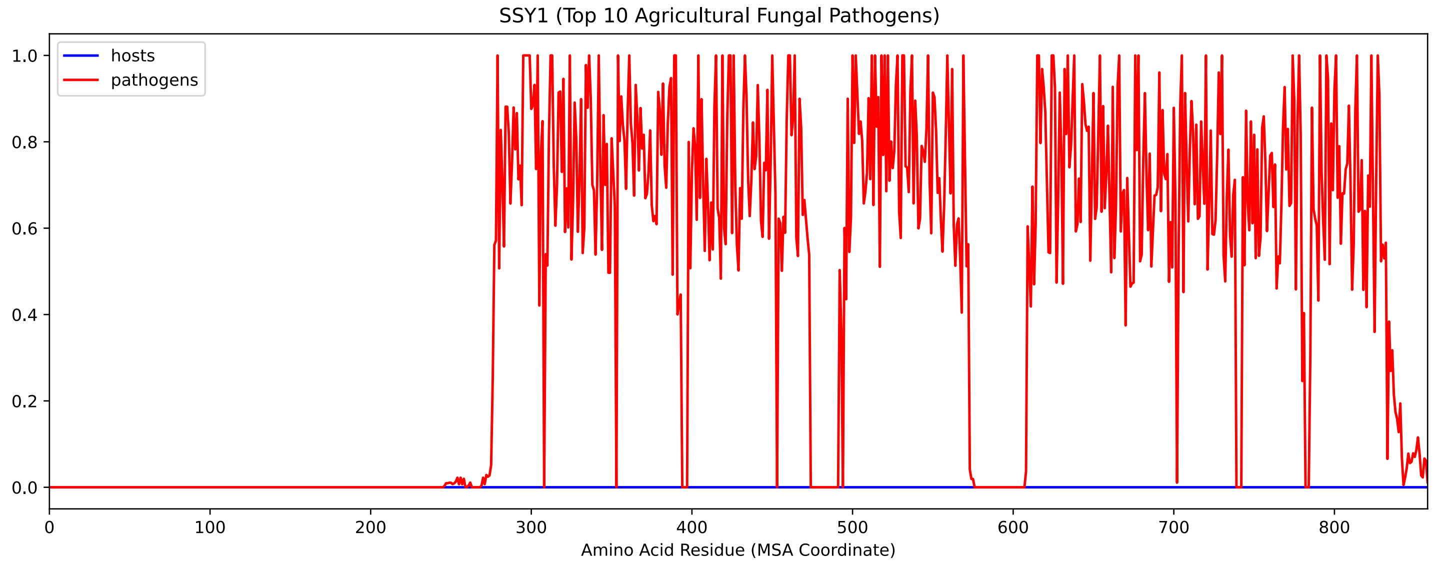


Figure S210: Sneath Similarity of Ssy1 for Top 10 Agricultural Fungal Pathogens, cf. Figure S208

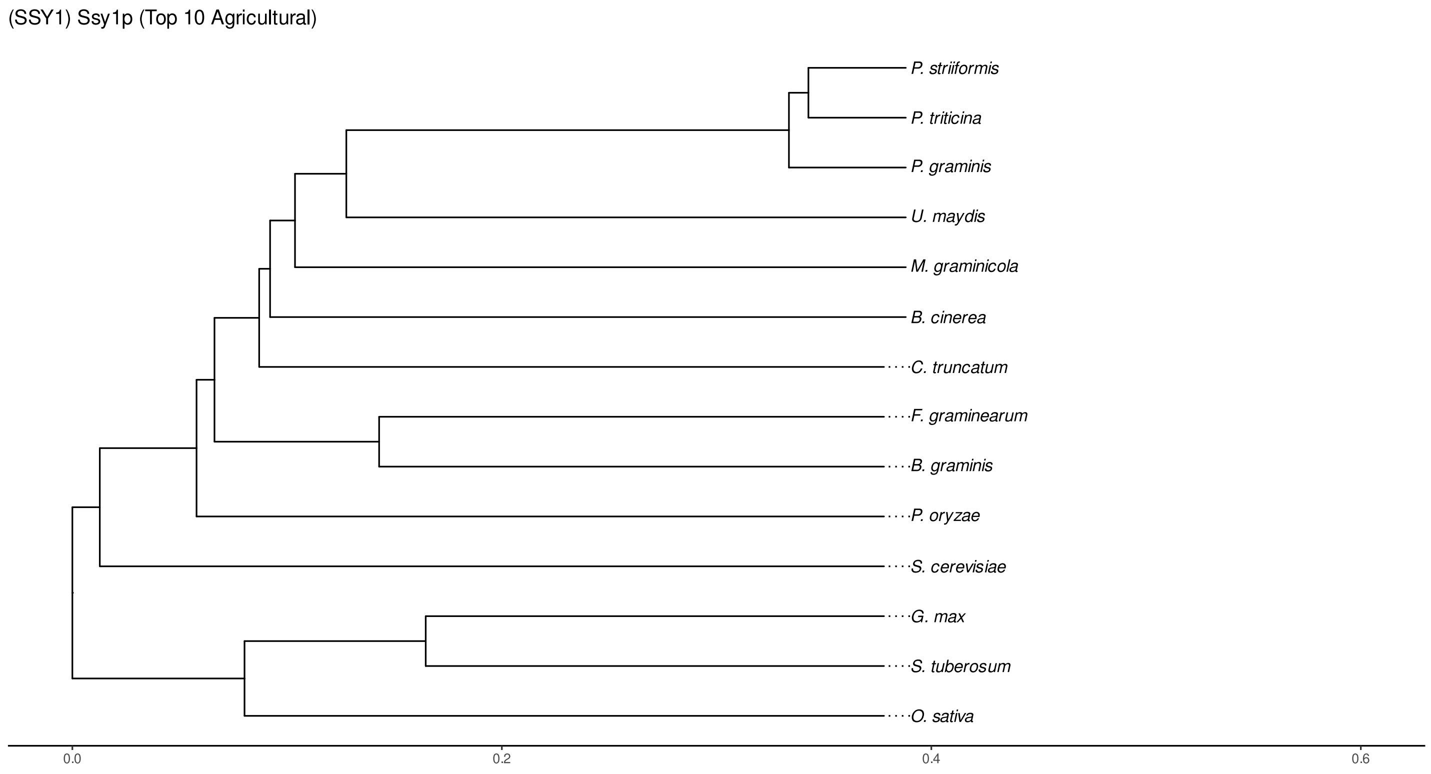


Figure S211: Phylogeny of hits for Ssy1 made using clustal omega and visualized using ggtree, cf. Figures S208 and S209.

### S4.14.3 NR

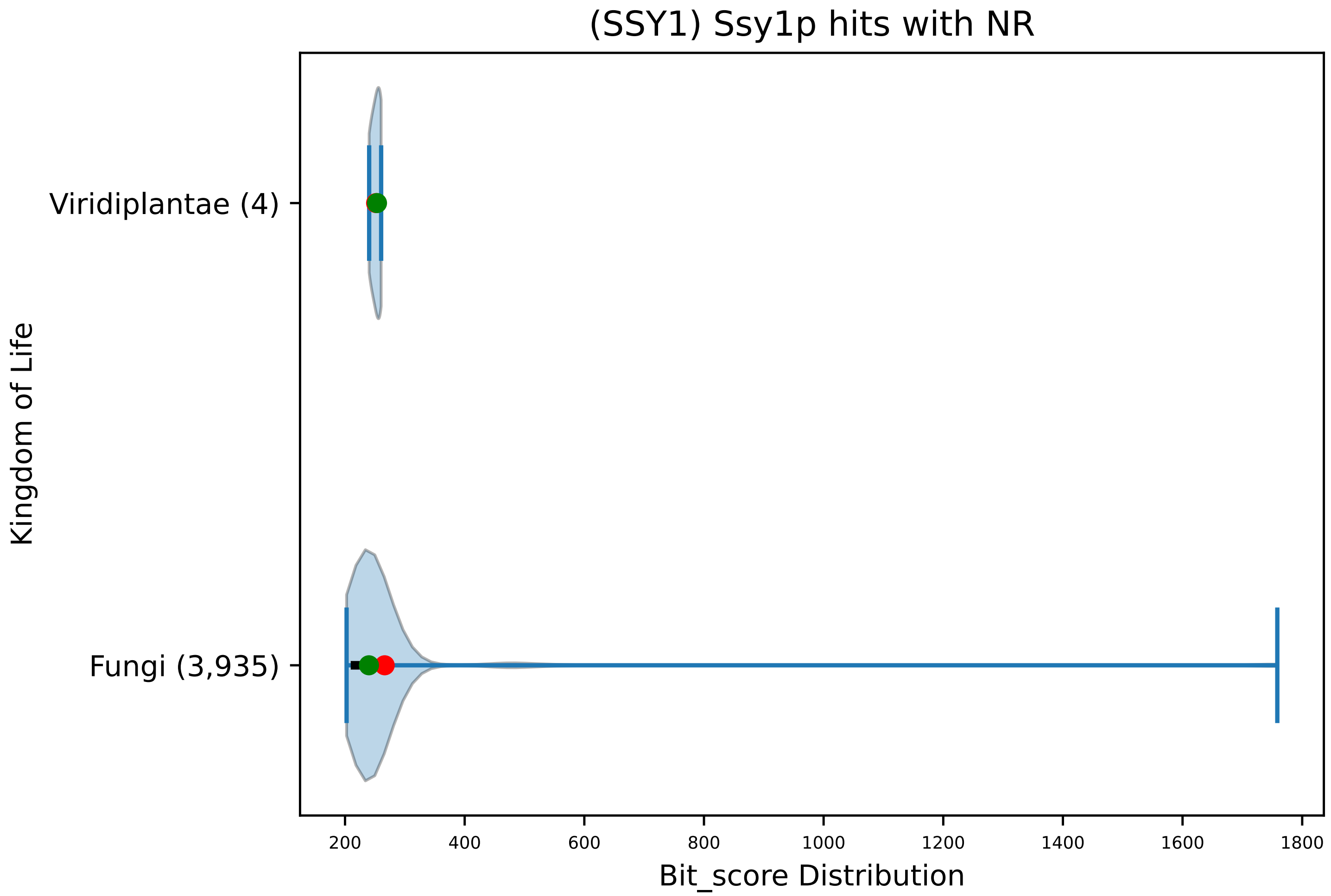


Figure S212: Non-redundant (NR) protein hits for DEG20010185/Ssy1, with expectation value of no more than 0.1. Green points are medians, and red points are arithmetic means.

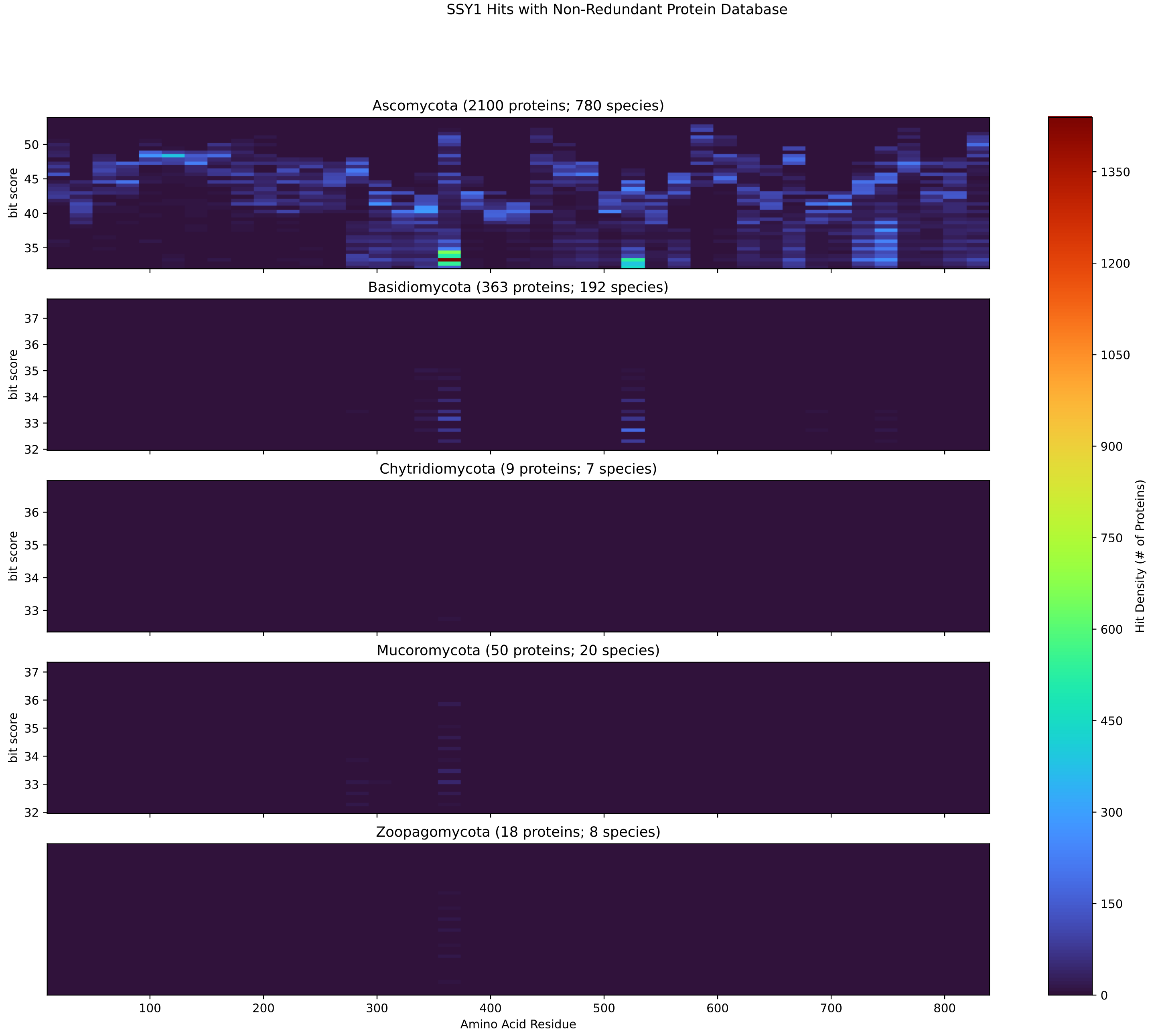


Figure S213: Non-redundant (NR) protein hits for Ssy1 in the kingdom Fungi.

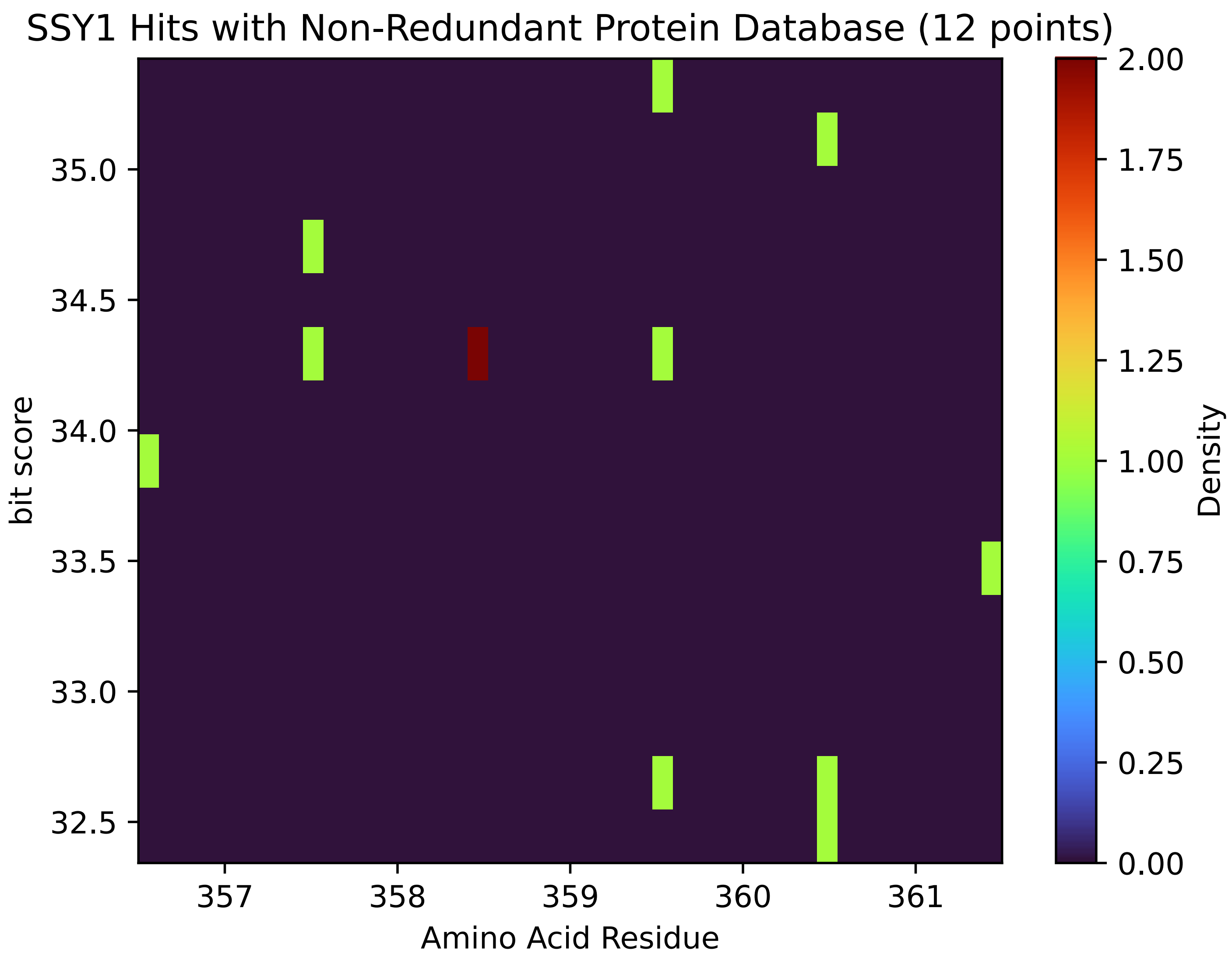


Figure S214: Non-redundant (NR) protein hits for Ssy1 in the kingdom Metazoa.

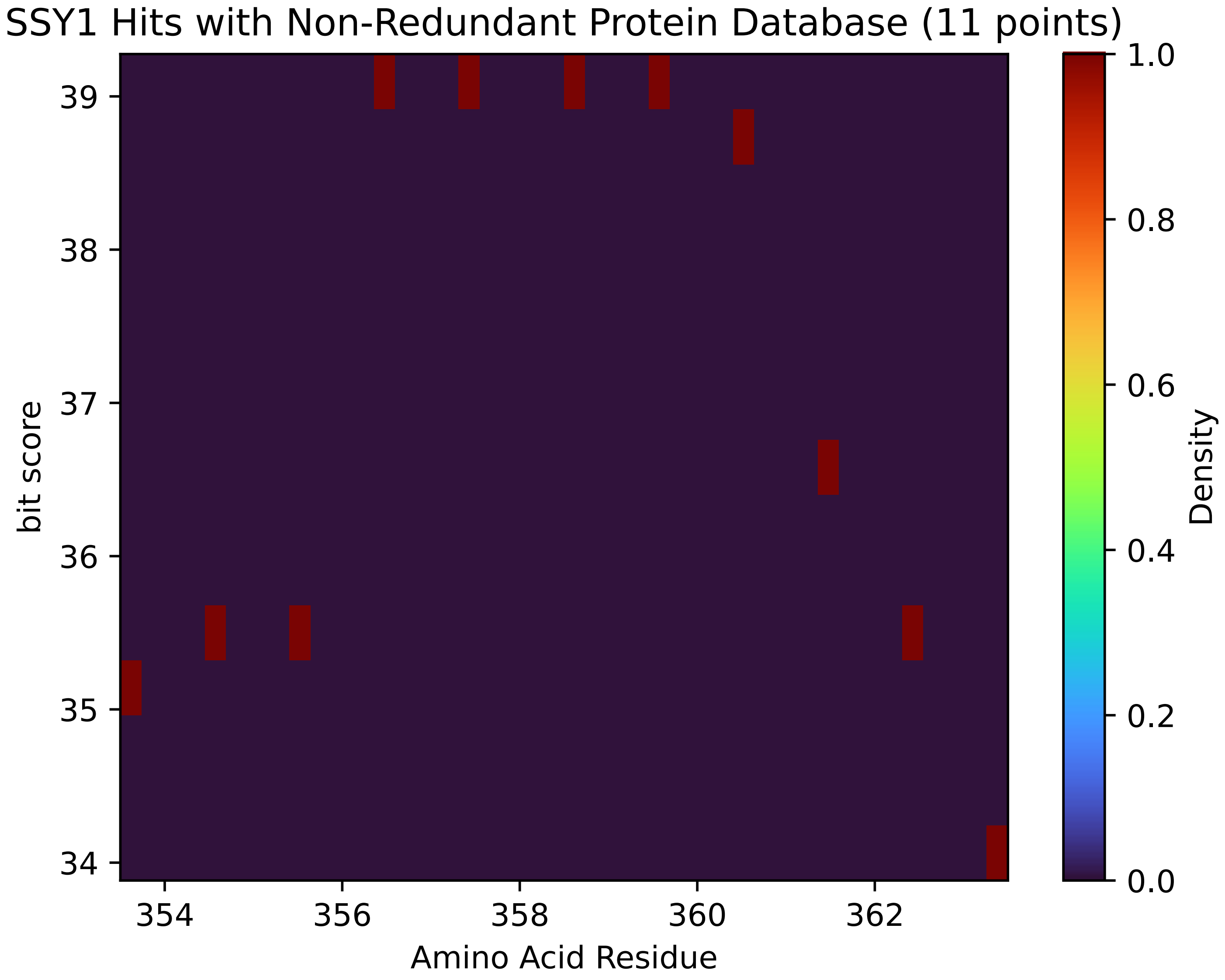


Figure S215: Non-redundant (NR) protein hits for Ssy1 in the kingdom SAR.

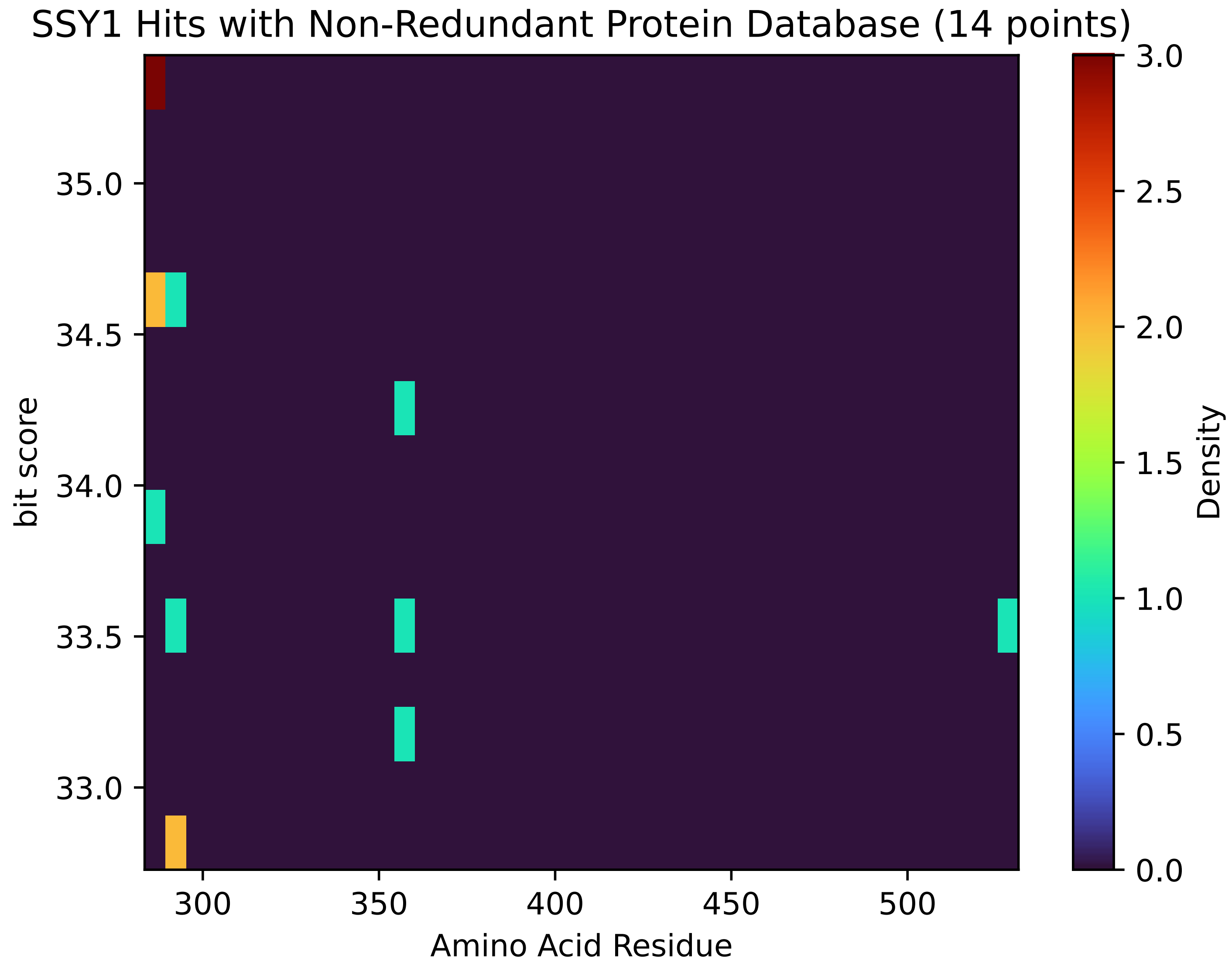


Figure S216: Non-redundant (NR) protein hits for Ssy1 in the kingdom Viridiplantae.

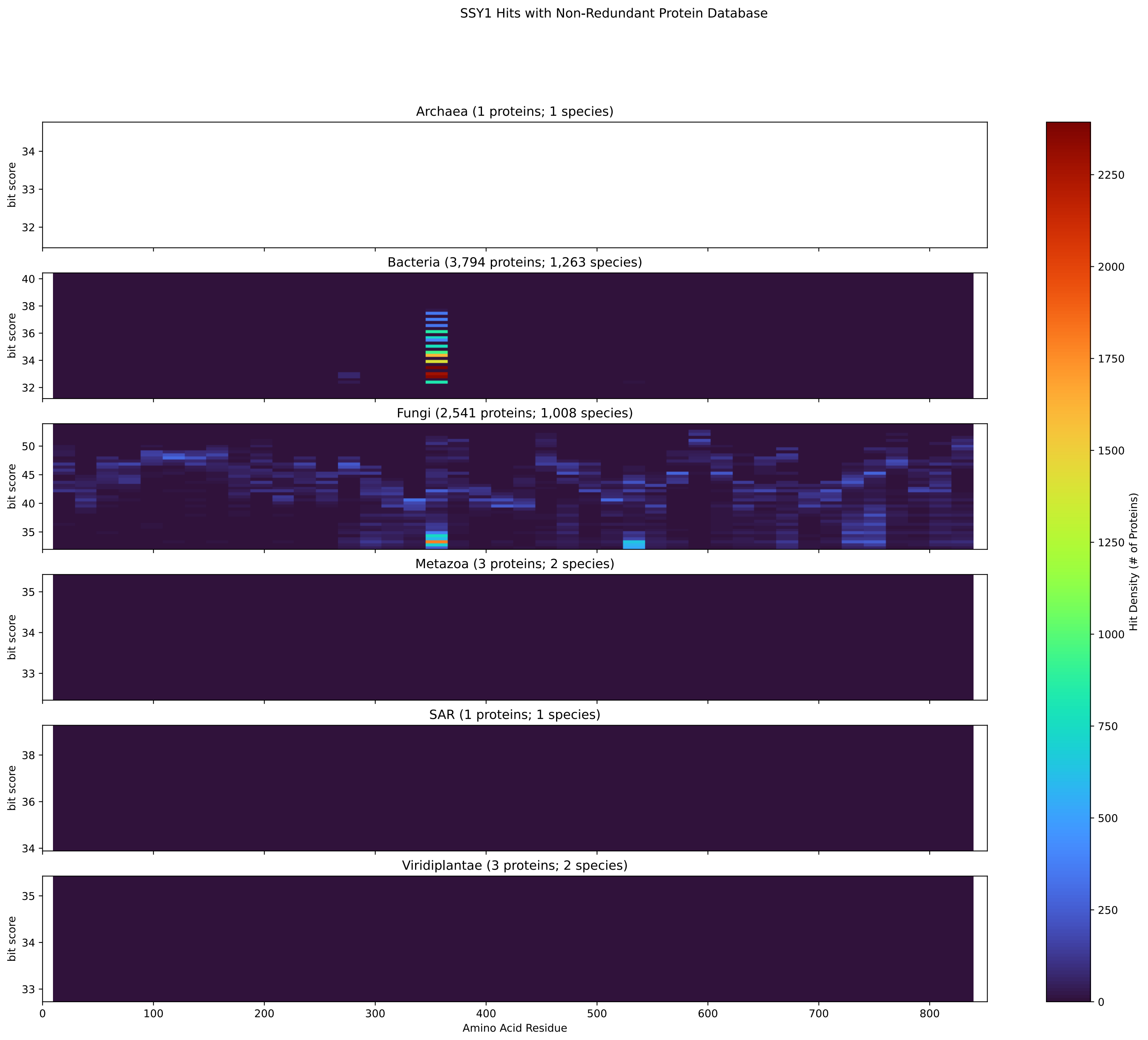


Figure S217: Non-redundant (NR) protein hits for DEG20010185/Ssy1 at 20 amino acid length queries.

#### S4.15 Ste12

##### S4.15.1 WHO Critical Pathogens

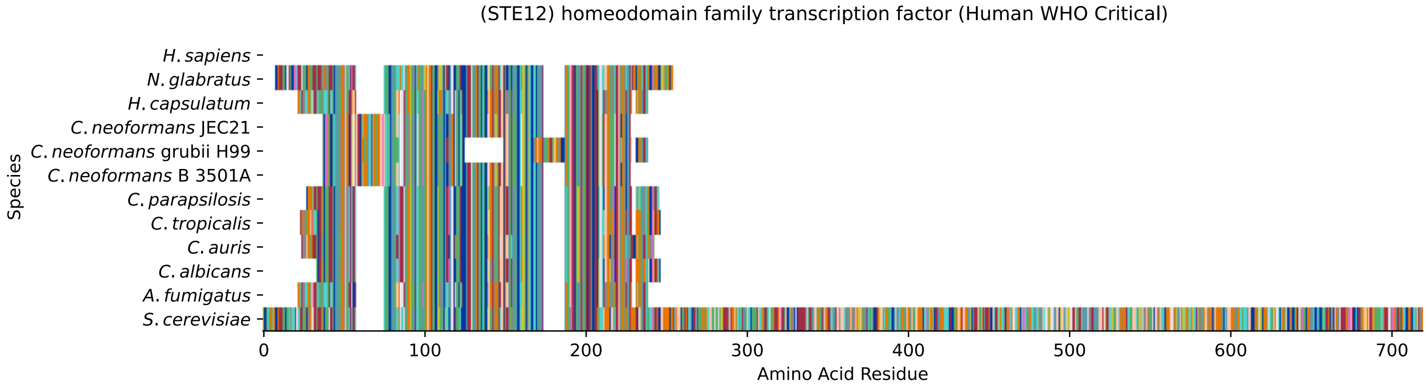


Figure S218: Multiple sequence alignment of yeast Ste12 (WHO Critical Pathogens). Cf. Figure S219 for alignment quality, and Figure S220 for Sneath similarity. Cf. Table S35 for protein names, and pairwise alignment metrics with yeast Ste12.

#### Ste12 MSA Quality

| | *A. f.* | *C. a.* | *C. a.* | *C. p.* | *C. t.* | *C. n.* | *C. n.* | *C. n.* | *H. c.* | *N. g.* | *S. c.* |
|---|---|---|---|---|---|---|---|---|---|---|---|
| *A. fumigatus* | 382.104 | 234.958 | 244.588 | 232.261 | 238.424 | 235.343 | 164.081 | 235.343 | 358.992 | 222.246 | 236.113 |
| *C. albicans* | 234.958 | 382.874 | 309.301 | 352.058 | 367.081 | 197.978 | 135.191 | 197.978 | 231.106 | 276.174 | 280.796 |
| *C. auris* | 244.588 | 309.301 | 392.889 | 305.449 | 314.694 | 198.364 | 132.109 | 198.364 | 239.58 | 272.322 | 277.715 |
| *C. parapsilosis* | 232.261 | 352.058 | 305.449 | 389.037 | 358.221 | 199.134 | 137.117 | 199.134 | 233.032 | 275.789 | 276.559 |
| *C. tropicalis* | 238.424 | 367.081 | 314.694 | 358.221 | 401.749 | 199.134 | 135.191 | 199.134 | 233.032 | 279.256 | 283.878 |
| *C. neoformans* B 3501A | 235.343 | 197.978 | 198.364 | 199.134 | 199.134 | 368.237 | 273.863 | 368.237 | 238.81 | 197.593 | 192.971 |
| *C. neoformans* grubii H99 | 164.081 | 135.191 | 132.109 | 137.117 | 135.191 | 273.863 | 362.073 | 273.863 | 167.548 | 131.339 | 121.709 |
| *C. neoformans* JEC21 | 235.343 | 197.978 | 198.364 | 199.134 | 199.134 | 368.237 | 273.863 | 368.237 | 238.81 | 197.593 | 192.971 |
| *H. capsulatum* | 358.992 | 231.106 | 239.58 | 233.032 | 233.032 | 238.81 | 167.548 | 238.81 | 380.563 | 218.009 | 233.802 |
| *N. glabratus* | 222.246 | 276.174 | 272.322 | 275.789 | 279.256 | 197.593 | 131.339 | 197.593 | 218.009 | 448.358 | 310.457 |
| *S. cerevisiae* | 236.113 | 280.796 | 277.715 | 276.559 | 283.878 | 192.971 | 121.709 | 192.971 | 233.802 | 310.457 | 1425.22 |

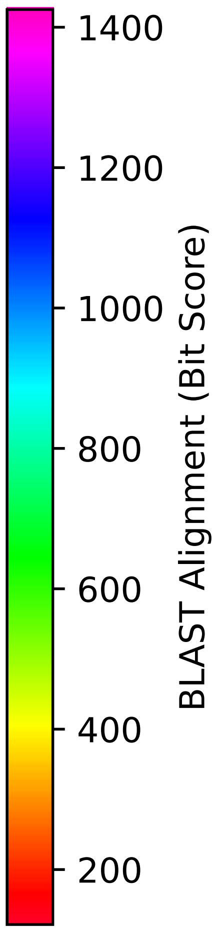


Figure S219: Multiple sequence alignment quality of Ste12 (WHO Critical Pathogens). Cf. Figures S218 and S221.

| Species | Hit Protein | Hit Length (a.a.) | evalue | align_len | bit_score | identity | positive | score | gaps | % identity | % positive |
|---|---|---|---|---|---|---|---|---|---|---|---|
| H.sapiens | - | - | - | - | - | - | - | - | - | - | - |
| N.glabratus | XP_449399.1 uncharacterized p-rotein CAGL0M01254g Nakaseomyc-es glabratus | 216 | 4.4e-101 | 216 | 320.087 | 144 | 180 | 819 | 1 | 20.9 | 26.2 |
| H.capsulatum | XP_045286576.1 transcription factor steA Histoplasma capsul-atum G186AR | 187 | 2.3e-71 | 187 | 244.588 | 109 | 147 | 623 | 3 | 15.8 | 21.4 |
| C.neoformans.JEC21 | XP_024512781.1 conserved hypo-thetical protein Cryptococcus neoformans var. neoformans JEC-21 | 178 | 8.1e-56 | 178 | 202.986 | 97 | 122 | 515 | 18 | 14.1 | 17.7 |
| C.neoformans.grubii.H99 | XP_012049551.1 transcription factor STE12 Cryptococcus neof-ormans var. grubii H99 | 202 | 9.3e-32 | 202 | 130.568 | 78 | 104 | 327 | 57 | 11.3 | 15.1 |
| C.neoformans.B.3501A | XP_776009.1 hypothetical prot-ein CNBD0580 Cryptococcus neof-ormans var. neoformans B-3501A | 178 | 9.2e-56 | 178 | 202.601 | 97 | 122 | 514 | 18 | 14.1 | 17.7 |
| C.parapsilosis | XP_036666384.1 uncharacterize-d protein CPAR2 208600 Candida parapsilosis | 189 | 2.6e-88 | 189 | 287.345 | 129 | 157 | 734 | 2 | 18.8 | 22.8 |
| C.tropicalis | XP_002549862.1 transcription factor CPH1 Candida tropicalis MYA-3404 | 194 | 2.9e-89 | 194 | 290.041 | 132 | 156 | 741 | 2 | 19.2 | 22.7 |
| C.auris | XP_028890469.2 protein STE12 Candida auris | 189 | 1.6e-87 | 189 | 281.952 | 126 | 152 | 720 | 0 | 18.3 | 22.1 |
| C.albicans | XP_713877.1 homeodomain famil-y transcription factor Candida albicans SC5314 | 184 | 2.5e-89 | 184 | 290.812 | 129 | 154 | 743 | 2 | 18.8 | 22.4 |
| A.fumigatus | XP_754013.1 sexual developmen-t transcription factor SteA As-pergillus fumigatus Af293 | 187 | 3.9e-72 | 187 | 246.128 | 110 | 147 | 627 | 3 | 16.0 | 21.4 |

Table S35: Pairwise alignment info from yeast Ste12 (DEG20010472), cf. Figure S218.

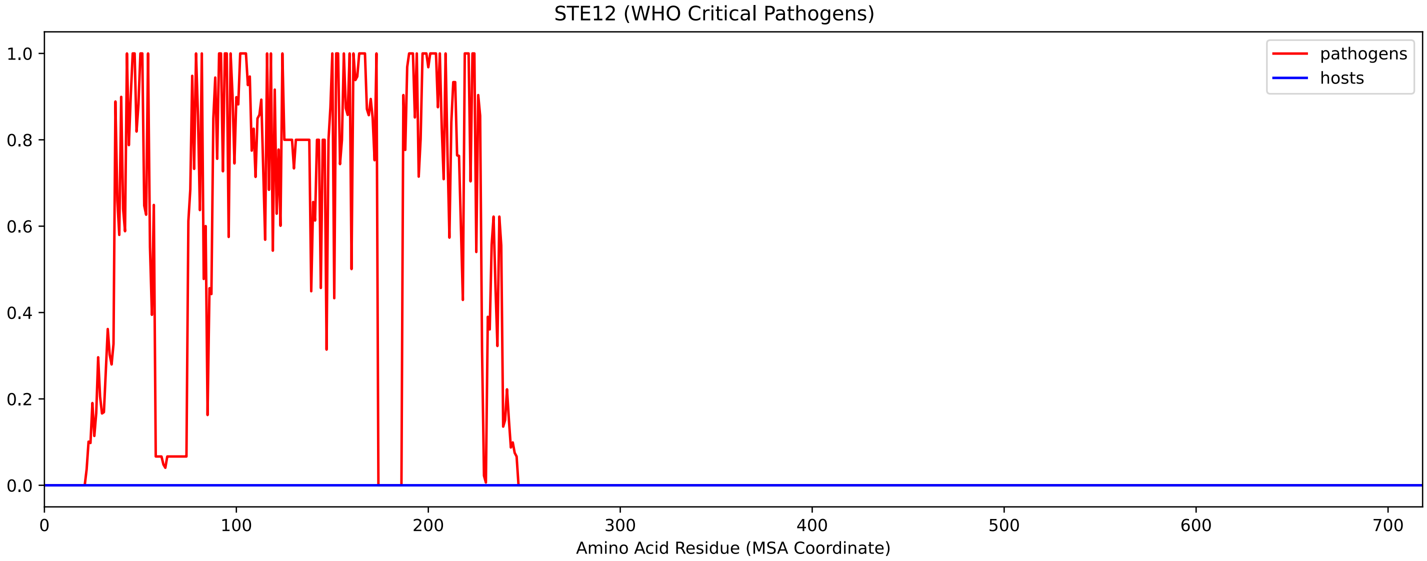


Figure S220: Sneath Similarity of Ste12 for WHO Critical Pathogens, cf. Figure S218

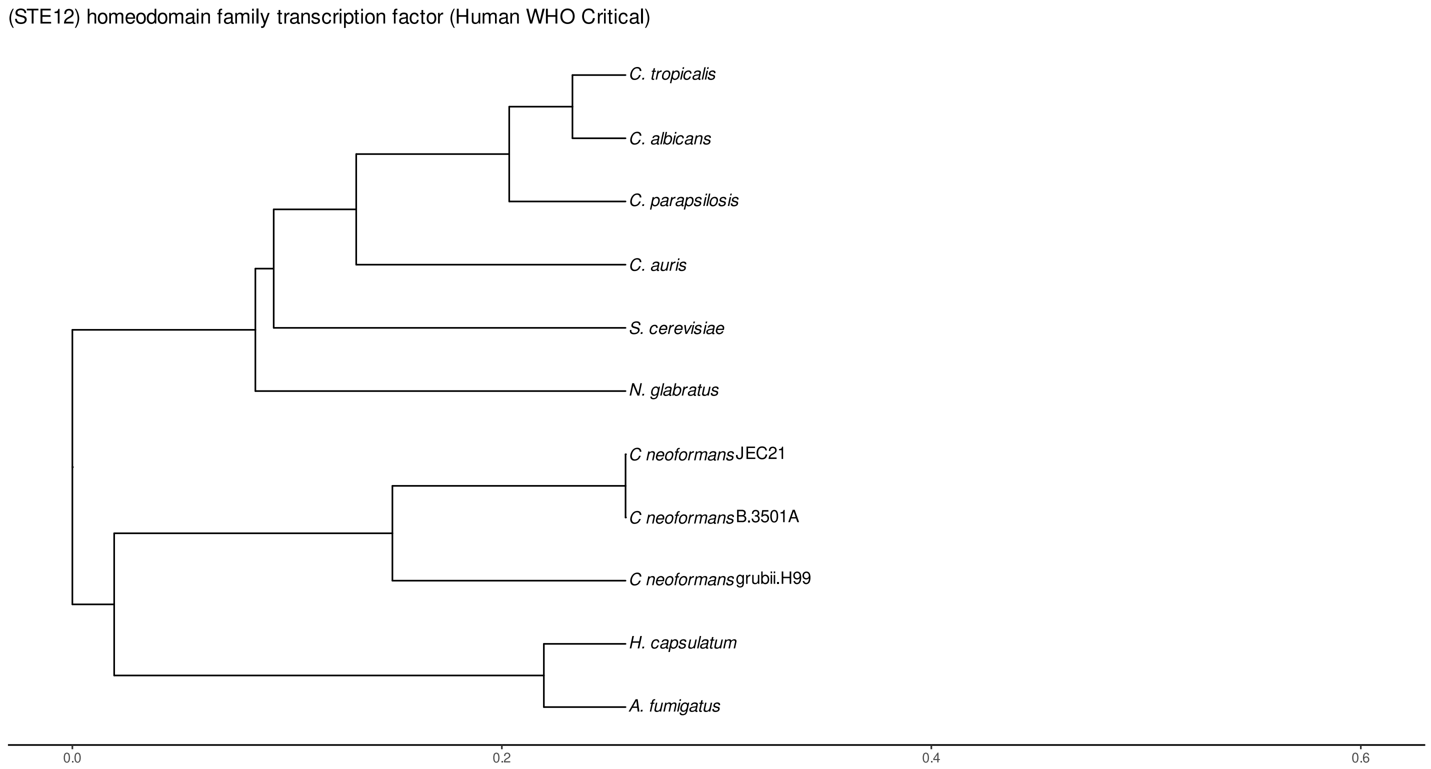


Figure S221: Phylogeny of hits for Ste12 made using clustal omega and visualized using ggtree, cf. Figures S218 and S219.

##### S4.15.2 Top 10 Agricultural Fungal Pathogens

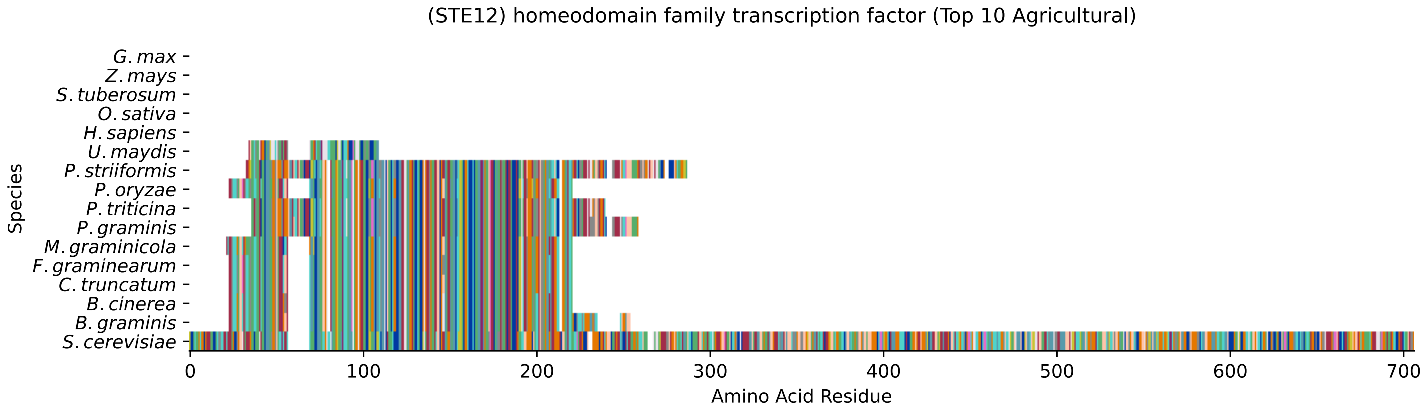


Figure S222: Multiple sequence alignment of yeast Ste12 (Top 10 Agricultural Fungal Pathogens). Cf. Figure S223 for alignment quality, and Figure S224 for Sneath similarity. Cf. Table S36 for protein names, and pairwise alignment metrics with yeast Ste12.

#### Ste12 MSA Quality

| | B.g. | B.c. | C.t. | F.g. | M.g. | P.g. | P.t. | P.o. | P.s. | U.m. | S.c. |
|---|---|---|---|---|---|---|---|---|---|---|---|
| B. graminis | 418.313 | 375.17 | 364.385 | 365.925 | 357.451 | 254.218 | 254.603 | 363.614 | 259.225 | 0 | 234.958 |
| B. cinerea | 375.17 | 376.711 | 361.688 | 362.459 | 357.066 | 253.062 | 253.832 | 360.533 | 258.455 | 0 | 233.417 |
| C. truncatum | 364.385 | 361.688 | 378.252 | 375.941 | 355.525 | 252.677 | 253.832 | 377.096 | 257.299 | 0 | 237.269 |
| F. graminearum | 365.925 | 362.459 | 375.941 | 379.022 | 352.443 | 253.062 | 253.832 | 374.785 | 257.684 | 0 | 237.654 |
| M. graminicola | 357.451 | 357.066 | 355.525 | 352.443 | 379.793 | 251.906 | 253.062 | 354.755 | 256.914 | 0 | 224.557 |
| P. graminis | 254.218 | 253.062 | 252.677 | 253.062 | 251.906 | 444.121 | 403.675 | 253.062 | 433.721 | 0 | 199.519 |
| P. triticina | 254.603 | 253.832 | 253.832 | 253.832 | 253.062 | 403.675 | 413.305 | 254.218 | 402.905 | 0 | 196.438 |
| P. oryzae | 363.614 | 360.533 | 377.096 | 374.785 | 354.755 | 253.062 | 254.218 | 379.407 | 257.684 | 0 | 237.269 |
| P. striiformis | 259.225 | 258.455 | 257.299 | 257.684 | 256.914 | 433.721 | 402.905 | 257.684 | 507.294 | 0 | 204.142 |
| U. maydis | 0 | 15.779 | 16.5494 | 16.1642 | 17.705 | 0 | 0 | 16.5494 | 0 | 127.102 | 27.7202 |
| S. cerevisiae | 234.958 | 233.417 | 237.269 | 237.654 | 224.557 | 199.519 | 196.438 | 237.269 | 204.142 | 27.7202 | 1425.22 |

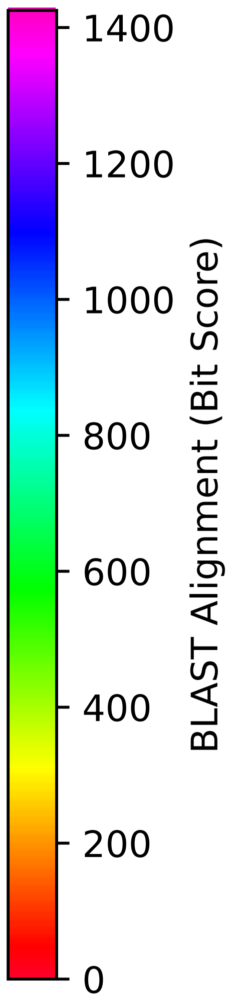


Figure S223: Multiple sequence alignment quality of Ste12 (Top 10 Agricultural Fungal Pathogens). Cf. Figures S222 and S225.

| Species | Hit Protein | Hit Length (a.a.) | evalue | align_len | bit_score | identity | positive | score | gaps | % identity | % positive |
|---|---|---|---|---|---|---|---|---|---|---|---|
| G.max | - | - | - | - | - | - | - | - | - | - | - |
| Z.mays | - | - | - | - | - | - | - | - | - | - | - |
| S.tuberosum | - | - | - | - | - | - | - | - | - | - | - |
| O.sativa | - | - | - | - | - | - | - | - | - | - | - |
| H.sapiens | - | - | - | - | - | - | - | - | - | - | - |
| U.maydis | XP_011389356.1 uncharacterize-d protein UMAG 02954 Ustilago maydis 521 | 65 | 0.034 | 65 | 35.8094 | 16 | 26 | 81 | 2 | 2.3 | 3.8 |
| P.striiformis | XP_047804633.1 hypothetical p-rotein Pst134EA 017974 Puccini-a striiformis f. sp. tritici | 253 | 6.8e-59 | 253 | 212.616 | 117 | 155 | 540 | 25 | 17.0 | 22.5 |
| P.oryzae | mRNA M BR32 EuGene 00048151-p1 — transcript=mRNA M BR32 EuGe-ne 00048151 — gene=M BR32 EuGe-ne 00048151 — organism=Pyricul-aria oryzae BR32 — gene produc-t=unspecified product — transc-ript product=unspecified produ-ct — location=BR32 scaffold000-04:1144110-1146702(-) — protei-n length=764 — sequence SO=sup-ercontig — SO=protein coding g-ene — is pseudo=false | 185 | 7.8e-72 | 185 | 246.899 | 112 | 146 | 629 | 3 | 16.3 | 21.2 |
| P.triticina | XP_053024960.1 uncharacterize-d protein PtA15 11A92 Puccinia triticina | 203 | 1.4e-56 | 203 | 205.682 | 102 | 137 | 522 | 17 | 14.8 | 19.9 |
| P.graminis | XP_003325390.2 transcription factor STE12 Puccinia graminis f. sp. tritici CRL 75-36-700–3 | 222 | 8.8e-58 | 222 | 209.149 | 109 | 144 | 531 | 20 | 15.8 | 20.9 |
| M.graminicola | ZTRI 8.206.mRNA-p1 — transcrip-t=ZTRI 8.206.mRNA — gene=ZTRI 8.206 — organism=Zymoseptoria tritici IPO323 — gene product=-similar to transcription facto-r steA — transcript product=si-milar to transcription factor steA — location=Ztri chr 8:759-315-761612(+) — protein length-=726 — sequence SO=chromosome — SO=protein coding gene — is pseudo=false | 187 | 6.8e-68 | 187 | 235.343 | 105 | 145 | 599 | 3 | 15.3 | 21.1 |
| F.graminearum | XP_011327056.1 transcription factor steA Fusarium graminear-um PH-1 | 185 | 8.5e-73 | 185 | 248.44 | 111 | 146 | 633 | 3 | 16.1 | 21.2 |
| C.truncatum | XP_036586582.1 ste12-like tra-nscription factor Colletotrich-um truncatum | 185 | 3.3e-72 | 185 | 247.284 | 112 | 146 | 630 | 3 | 16.3 | 21.2 |
| B.cinerea | XP_024551490.1 Bcste12 Botryt-is cinerea B05.10 | 185 | 4.5e-71 | 185 | 243.047 | 109 | 145 | 619 | 3 | 15.8 | 21.1 |
| B.graminis | VCU39448.1 — transcript=BGT962-24V316 LOCUS707 t1 — gene=BGT9-6224V316 LOCUS707 — organism=B-lumeria graminis f. sp. tritic-i 96224 — gene product=unspeci-fied product — transcript prod-uct=unspecified product — loca-tion=LR026985:958957-961327(+) — protein length=692 — sequen-ce SO=chromosome — SO=protein coding gene — is pseudo=false | 205 | 4e-71 | 205 | 243.047 | 112 | 152 | 619 | 3 | 16.3 | 22.1 |

Table S36: Pairwise alignment info from yeast Ste12 (DEG20010472), cf. Figure S222.

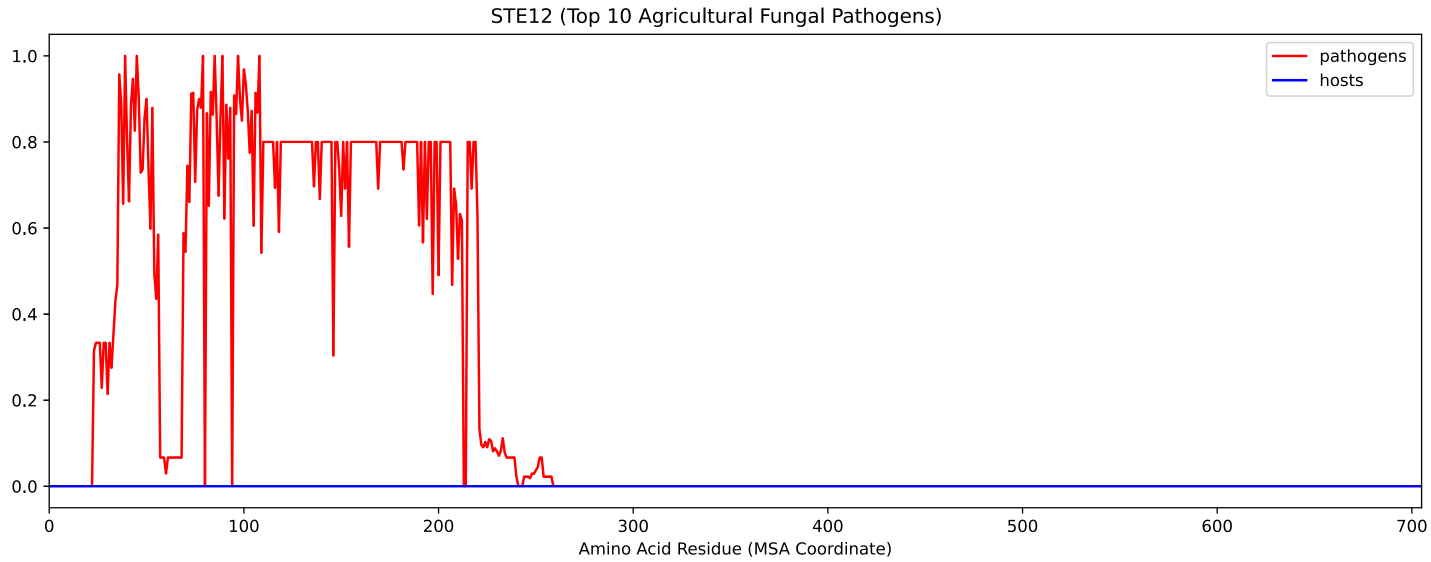


Figure S224: Sneath Similarity of Ste12 for Top 10 Agricultural Fungal Pathogens, cf. Figure S222

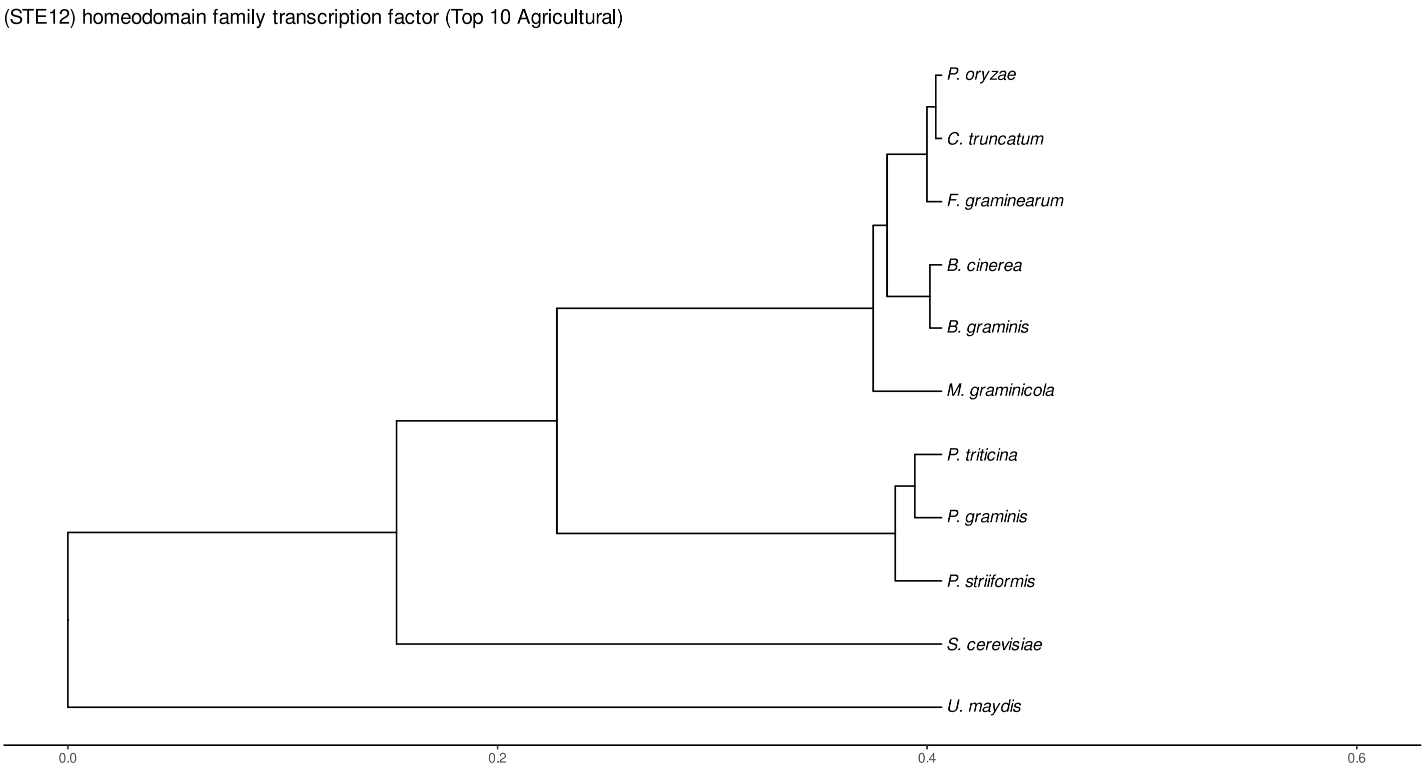


Figure S225: Phylogeny of hits for Ste12 made using clustal omega and visualized using ggtree, cf. Figures S222 and S223.

#### S4.15.3 NR

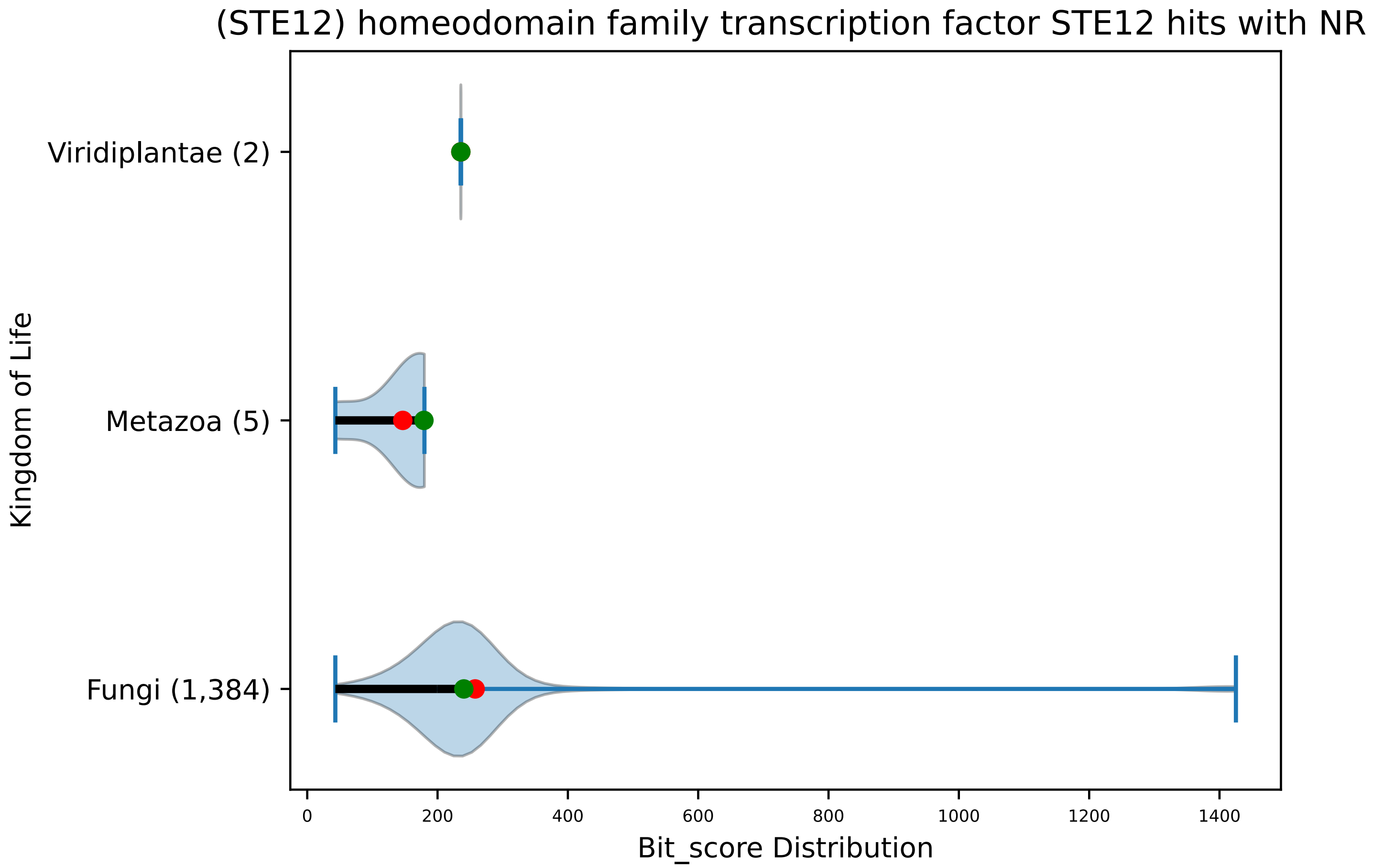


Figure S226: Non-redundant (NR) protein hits for DEG20010472/Ste12, with expectation value of no more than 0.1. Green points are medians, and red points are arithmetic means.

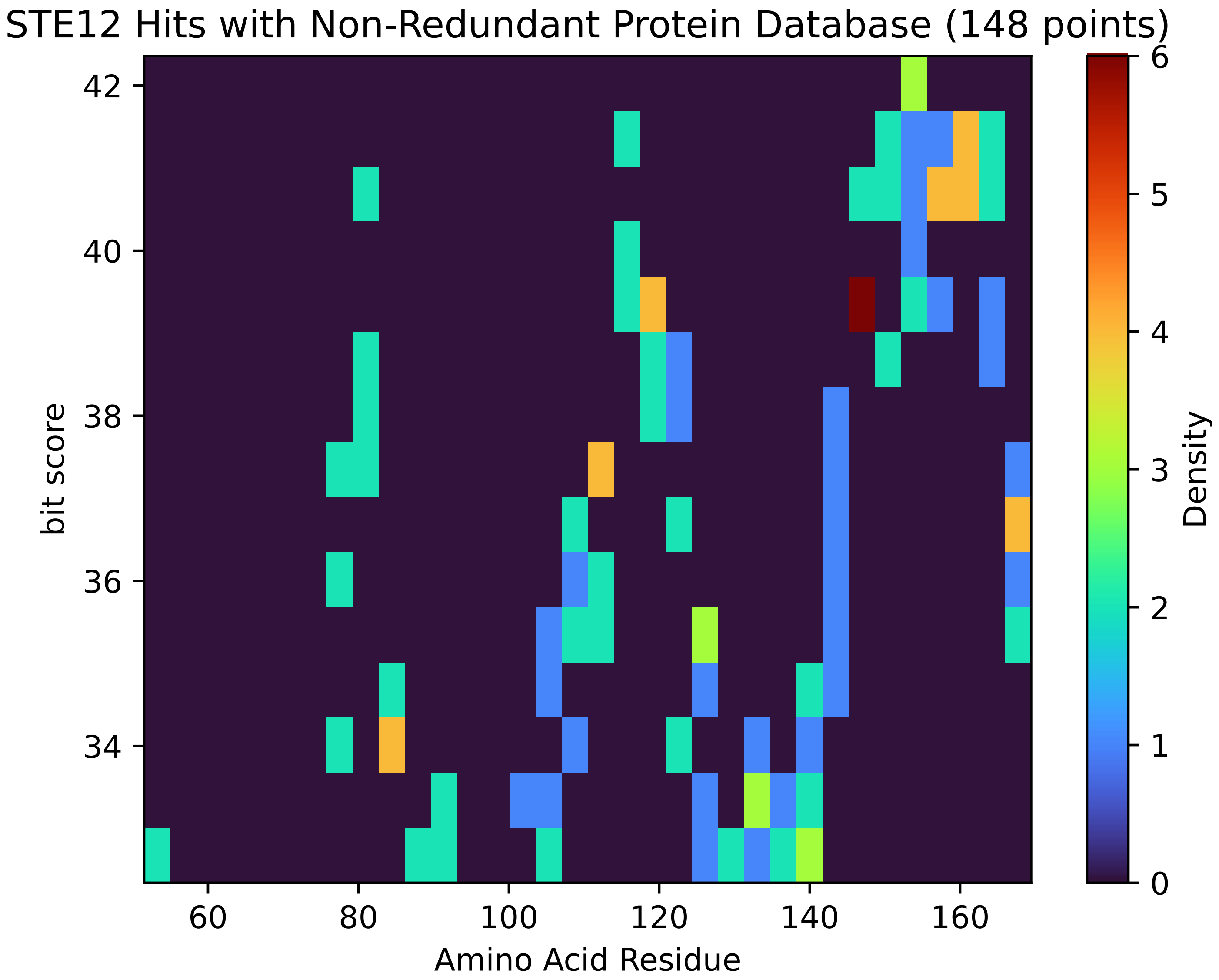


Figure S227: Non-redundant (NR) protein hits for Ste12 in the kingdom Viridiplantae.

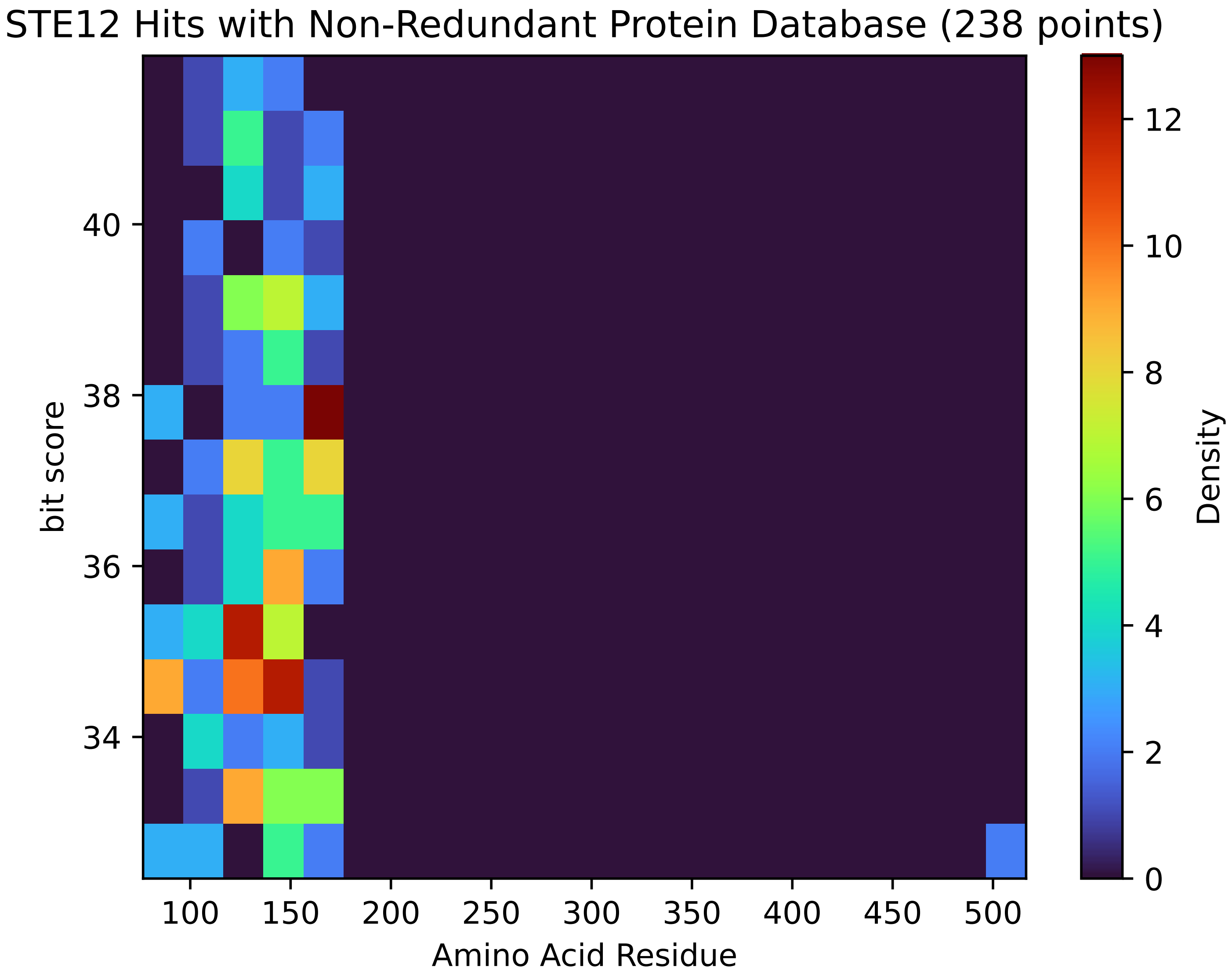


Figure S228: Non-redundant (NR) protein hits for Ste12 in the kingdom Metazoa.

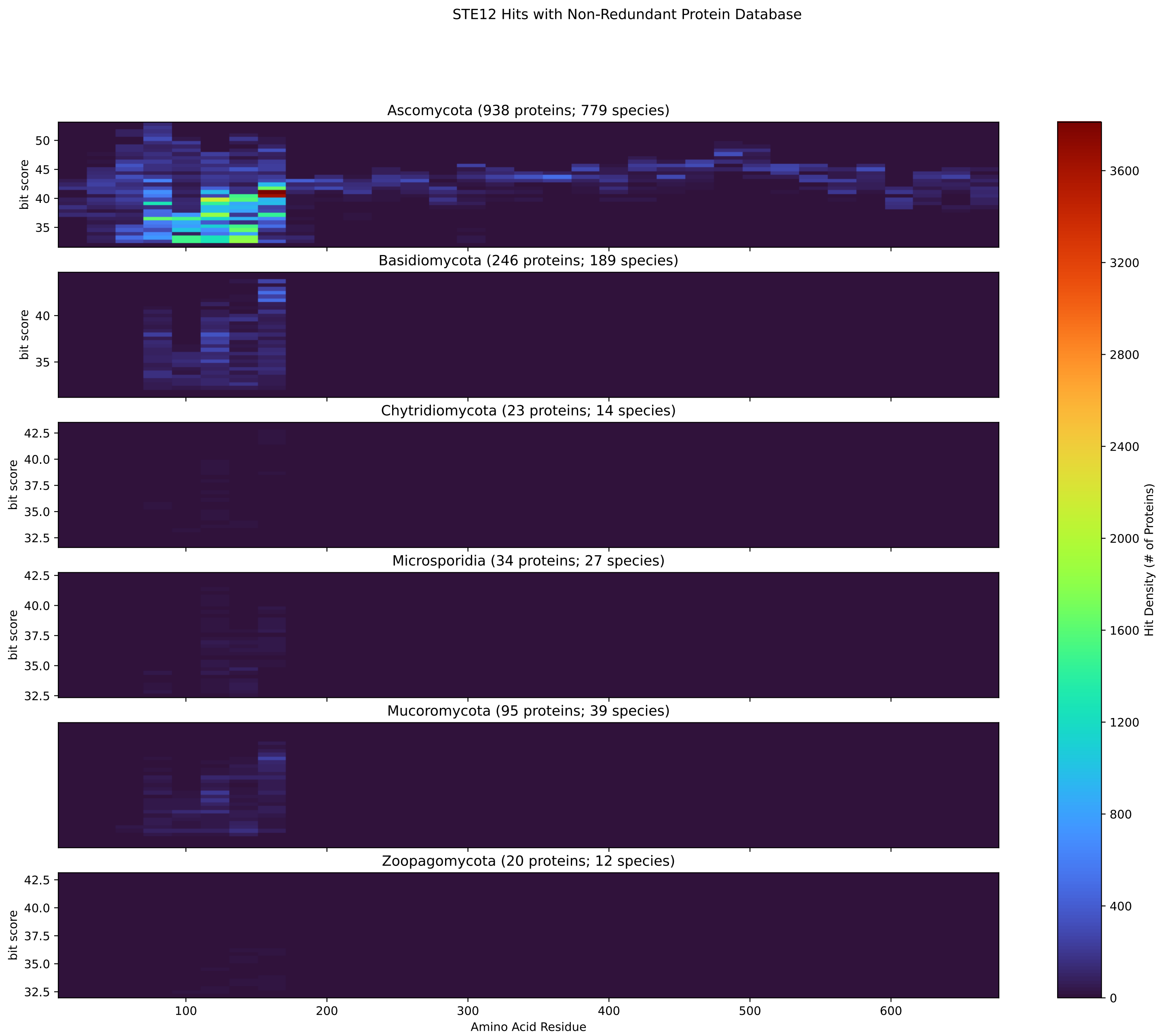


Figure S229: Non-redundant (NR) protein hits for Ste12 in the kingdom Fungi.

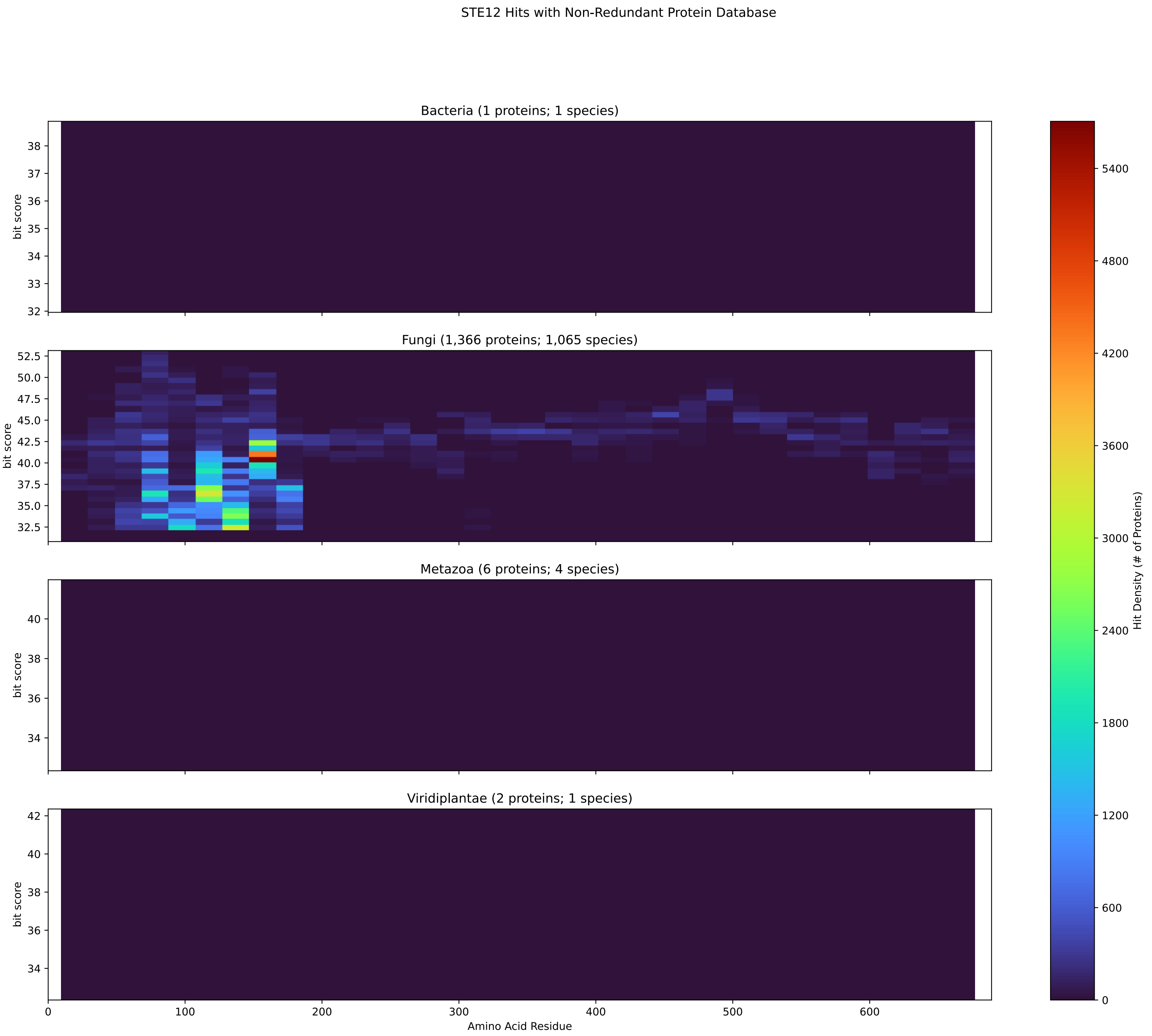


Figure S230: Non-redundant (NR) protein hits for DEG20010472/Ste12 at 20 amino acid length queries.

#### S4.16 Trl1

##### S4.16.1 WHO Critical Pathogens

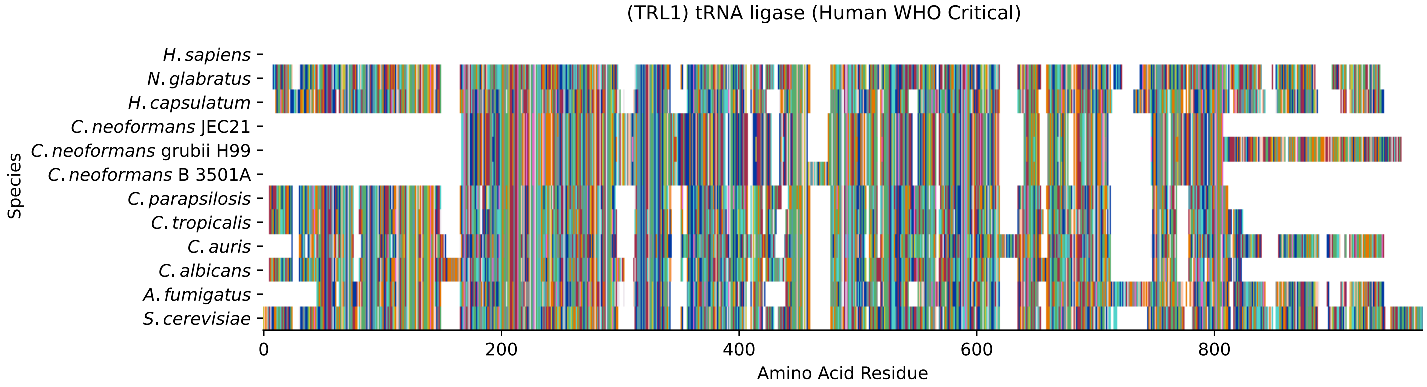


Figure S231: Multiple sequence alignment of yeast Trl1 (WHO Critical Pathogens). Cf. Figure S232 for alignment quality, and Figure S233 for Sneath similarity. Cf. Table S37 for protein names, and pairwise alignment metrics with yeast Trl1.

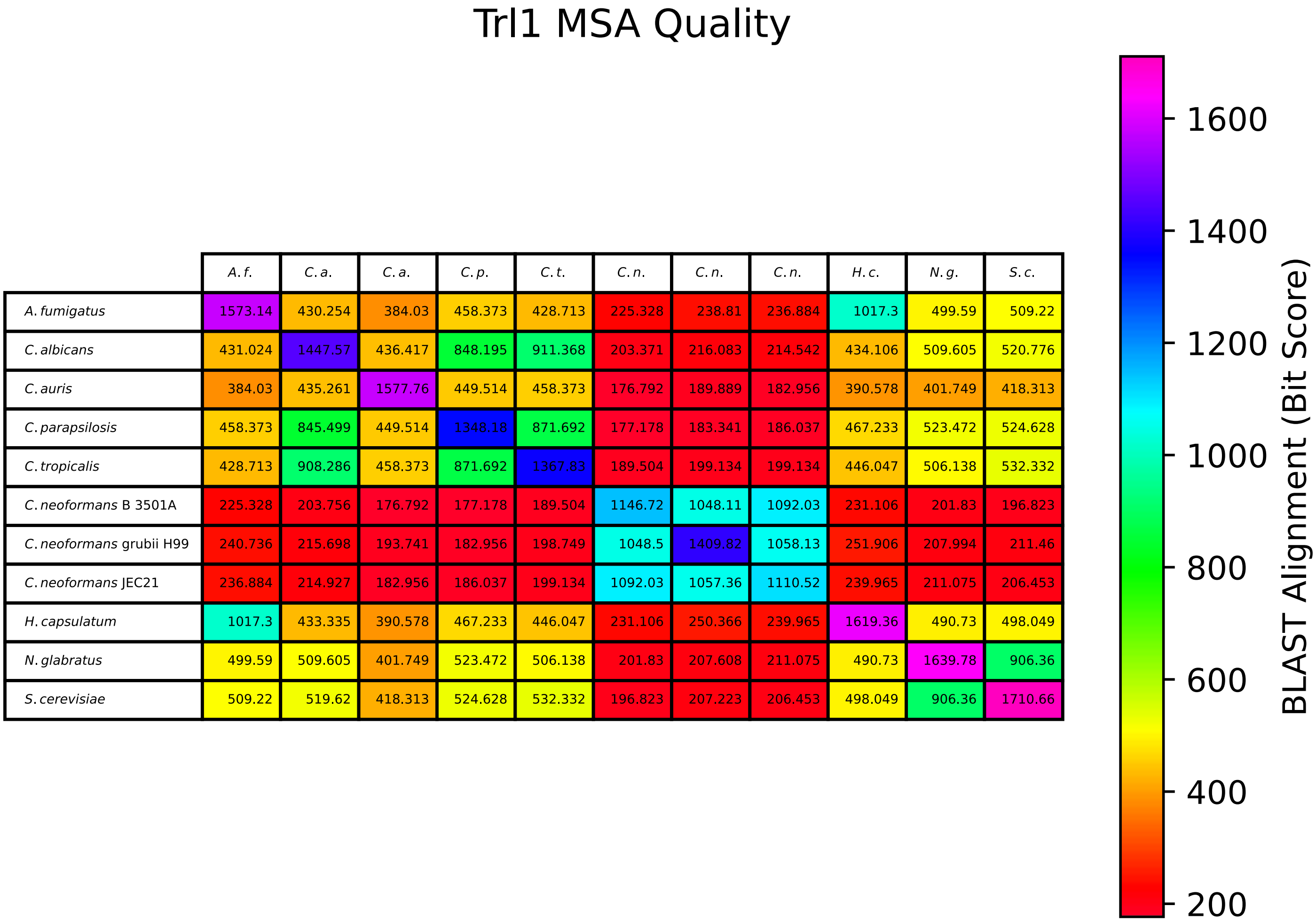


| | *A. f.* | *C. a.* | *C. a.* | *C. p.* | *C. t.* | *C. n.* | *C. n.* | *C. n.* | *H. c.* | *N. g.* | *S. c.* |
|---|---|---|---|---|---|---|---|---|---|---|---|
| *A. fumigatus* | 1573.14 | 430.254 | 384.03 | 458.373 | 428.713 | 225.328 | 238.81 | 236.884 | 1017.3 | 499.59 | 509.22 |
| *C. albicans* | 431.024 | 1447.57 | 436.417 | 848.195 | 911.368 | 203.371 | 216.083 | 214.542 | 434.106 | 509.605 | 520.776 |
| *C. auris* | 384.03 | 435.261 | 1577.76 | 449.514 | 458.373 | 176.792 | 189.889 | 182.956 | 390.578 | 401.749 | 418.313 |
| *C. parapsilosis* | 458.373 | 845.499 | 449.514 | 1348.18 | 871.692 | 177.178 | 183.341 | 186.037 | 467.233 | 523.472 | 524.628 |
| *C. tropicalis* | 428.713 | 908.286 | 458.373 | 871.692 | 1367.83 | 189.504 | 199.134 | 199.134 | 446.047 | 506.138 | 532.332 |
| *C. neoformans* B 3501A | 225.328 | 203.756 | 176.792 | 177.178 | 189.504 | 1146.72 | 1048.11 | 1092.03 | 231.106 | 201.83 | 196.823 |
| *C. neoformans* grubii H99 | 240.736 | 215.698 | 193.741 | 182.956 | 198.749 | 1048.5 | 1409.82 | 1058.13 | 251.906 | 207.994 | 211.46 |
| *C. neoformans* JEC21 | 236.884 | 214.927 | 182.956 | 186.037 | 199.134 | 1092.03 | 1057.36 | 1110.52 | 239.965 | 211.075 | 206.453 |
| *H. capsulatum* | 1017.3 | 433.335 | 390.578 | 467.233 | 446.047 | 231.106 | 250.366 | 239.965 | 1619.36 | 490.73 | 498.049 |
| *N. glabratus* | 499.59 | 509.605 | 401.749 | 523.472 | 506.138 | 201.83 | 207.608 | 211.075 | 490.73 | 1639.78 | 906.36 |
| *S. cerevisiae* | 509.22 | 519.62 | 418.313 | 524.628 | 532.332 | 196.823 | 207.223 | 206.453 | 498.049 | 906.36 | 1710.66 |

Figure S232: Multiple sequence alignment quality of Trl1 (WHO Critical Pathogens). Cf. Figures S231 and S234.

| Species | Hit Protein | Hit Length (a.a.) | evalue | align_len | bit_score | identity | positive | score | gaps | % identity | % positive |
|---|---|---|---|---|---|---|---|---|---|---|---|
| H.sapiens | - | - | - | - | - | - | - | - | - | - | - |
| N.glabratus | XP_449481.1 uncharacterized p-rotein CAGL0M03091g Nakaseomyc-es glabratus | 804 | 0 | 804 | 906.746 | 450 | 572 | 2342 | 30 | 54.4 | 69.2 |
| H.capsulatum | XP_045288514.1 tRNA ligase Hi-stoplasma capsulatum G186AR | 819 | 1.2e-164 | 819 | 496.893 | 308 | 457 | 1278 | 76 | 37.2 | 55.3 |
| C.neoformans.JEC21 | XP_024512786.1 RNA ligase (AT-P), putative Cryptococcus neof-ormans var. neoformans JEC21 | 567 | 5.4e-56 | 567 | 206.838 | 157 | 280 | 525 | 67 | 19.0 | 33.9 |
| C.neoformans.grubii.H99 | XP_012049418.1 tRNA ligase Cr-yptococcus neoformans var. gru-bii H99 | 713 | 5.8e-56 | 713 | 207.223 | 183 | 326 | 526 | 81 | 22.1 | 39.4 |
| C.neoformans.B.3501A | XP_775856.1 hypothetical prot-ein CNBD2650 Cryptococcus neof-ormans var. neoformans B-3501A | 582 | 6.9e-53 | 582 | 197.593 | 157 | 283 | 501 | 80 | 19.0 | 34.2 |
| C.parapsilosis | XP_036664316.1 uncharacterize-d protein CPAR2 301270 Candida parapsilosis | 680 | 7.9e-176 | 680 | 524.628 | 299 | 415 | 1350 | 31 | 36.2 | 50.2 |
| C.tropicalis | XP_002550758.1 tRNA ligase Ca-ndida tropicalis MYA-3404 | 693 | 5.2e-179 | 693 | 533.102 | 295 | 427 | 1372 | 33 | 35.7 | 51.6 |
| C.auris | XP_028888635.2 tRNA ligase Ca-ndida auris | 806 | 2.7e-134 | 806 | 417.927 | 282 | 431 | 1073 | 79 | 34.1 | 52.1 |
| C.albicans | XP_721349.1 tRNA ligase Candi-da albicans SC5314 | 722 | 2.3e-173 | 722 | 519.62 | 302 | 426 | 1337 | 57 | 36.5 | 51.5 |
| A.fumigatus | XP_752336.1 tRNA ligase Asper-gillus fumigatus Af293 | 802 | 3.4e-169 | 802 | 509.605 | 312 | 441 | 1311 | 94 | 37.7 | 53.3 |

Table S37: Pairwise alignment info from yeast Trl1 (DEG20010555), cf. Figure S231.

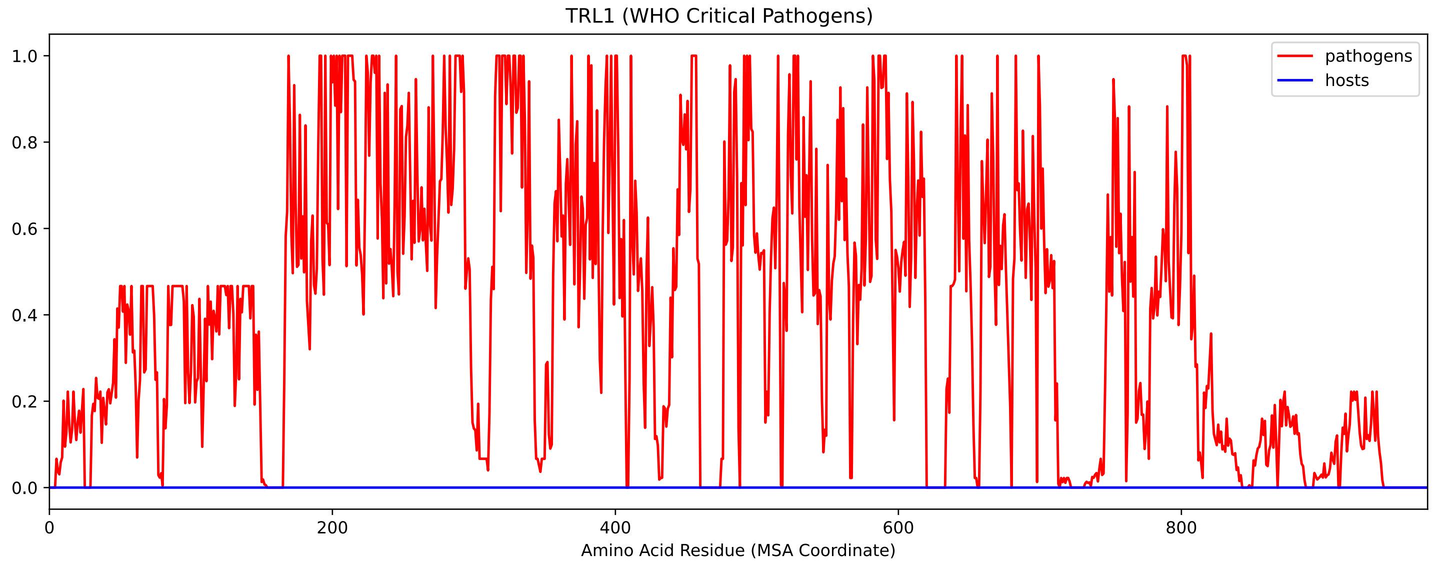


Figure S233: Sneath Similarity of Trl1 for WHO Critical Pathogens, cf. Figure S231

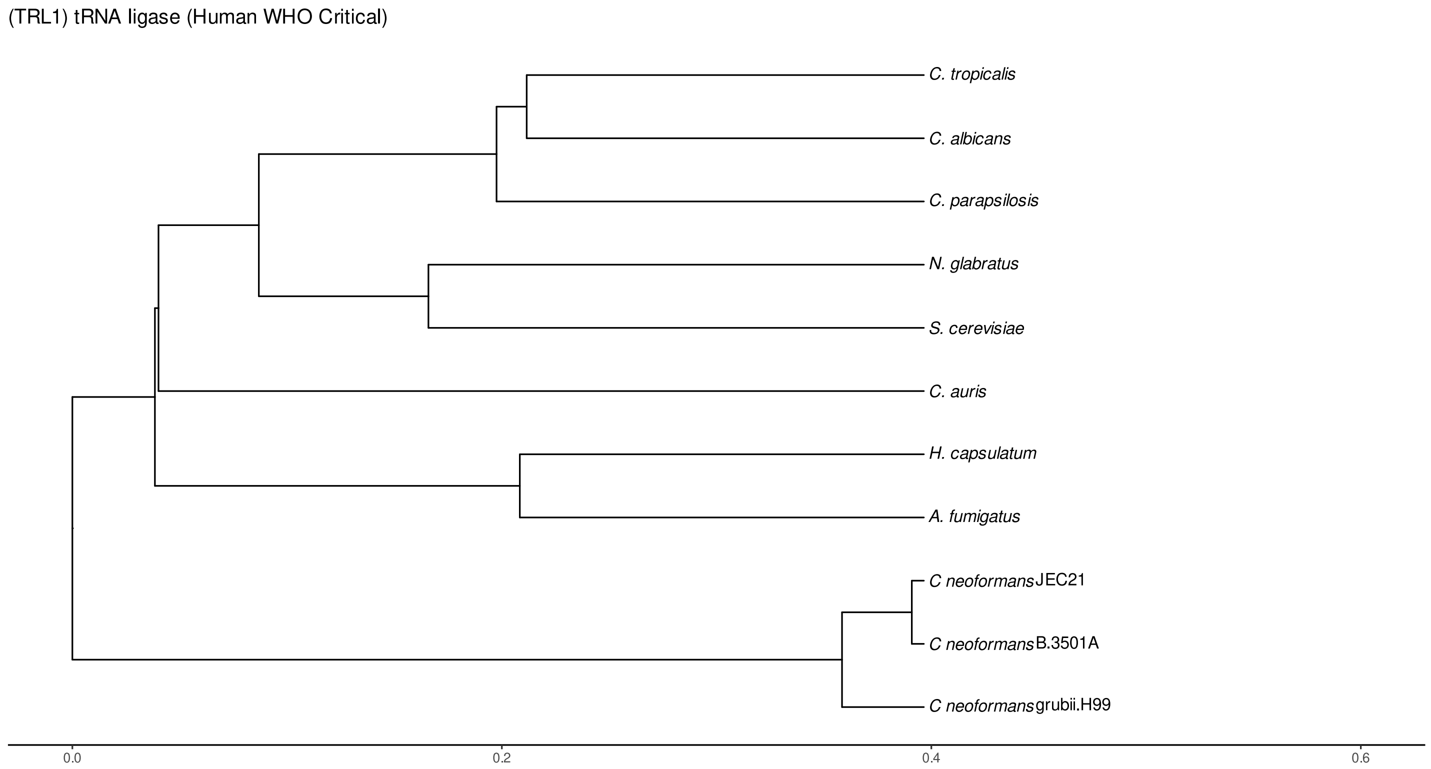


Figure S234: Phylogeny of hits for Trl1 made using clustal omega and visualized using ggtree, cf. Figures S231 and S232.

##### S4.16.2 Top 10 Agricultural Fungal Pathogens

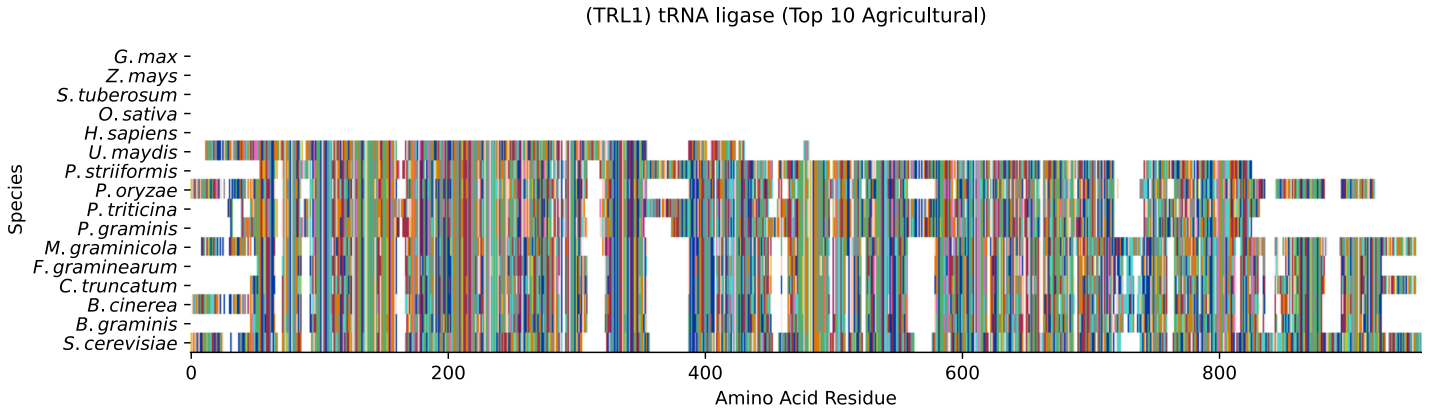


Figure S235: Multiple sequence alignment of yeast Trl1 (Top 10 Agricultural Fungal Pathogens). Cf. Figure S236 for alignment quality, and Figure S237 for Sneath similarity. Cf. Table S38 for protein names, and pairwise alignment metrics with yeast Trl1.

#### Trl1 MSA Quality

| | B. g. | B. c. | C. t. | F. g. | M. g. | P. g. | P. t. | P. o. | P. s. | U. m. | S. c. |
|---|---|---|---|---|---|---|---|---|---|---|---|
| *B. graminis* | 1508.43 | 880.167 | 793.112 | 783.867 | 737.643 | 303.138 | 283.878 | 687.567 | 291.197 | 231.491 | 0 |
| *B. cinerea* | 864.759 | 1602.42 | 901.353 | 900.197 | 845.884 | 329.717 | 306.99 | 802.357 | 320.472 | 235.728 | 0 |
| *C. truncatum* | 793.882 | 918.302 | 1580.46 | 1061.98 | 850.892 | 333.569 | 335.109 | 894.419 | 305.449 | 246.514 | 0 |
| *F. graminearum* | 774.237 | 910.598 | 1062.75 | 1498.8 | 802.742 | 327.791 | 317.775 | 865.914 | 294.278 | 251.906 | 0 |
| *M. graminicola* | 718.768 | 845.884 | 841.262 | 792.727 | 1679.84 | 336.65 | 330.102 | 751.125 | 309.301 | 251.521 | 0 |
| *P. graminis* | 294.664 | 329.717 | 330.102 | 325.479 | 336.65 | 1498.03 | 983.786 | 346.665 | 1139.79 | 223.016 | 0 |
| *P. triticina* | 283.878 | 315.849 | 344.739 | 323.553 | 340.502 | 1003.05 | 1494.56 | 338.191 | 991.104 | 236.498 | 0 |
| *P. oryzae* | 669.848 | 802.357 | 889.026 | 862.448 | 751.125 | 346.665 | 327.405 | 1595.48 | 319.701 | 252.677 | 0 |
| *P. striiformis* | 283.108 | 320.472 | 302.368 | 291.967 | 309.301 | 1139.79 | 962.6 | 319.701 | 1501.11 | 231.876 | 0 |
| *U. maydis* | 230.72 | 235.728 | 246.128 | 251.521 | 251.521 | 223.016 | 236.113 | 252.677 | 231.876 | 789.26 | 0 |
| *S. cerevisiae* | 0 | 0 | 0 | 0 | 0 | 0 | 0 | 0 | 0 | 0 | 0 |

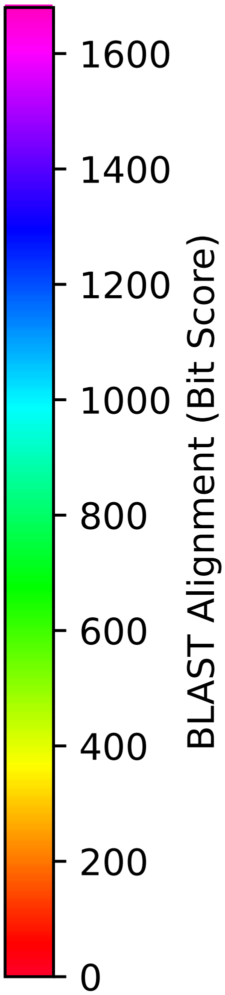


Figure S236: Multiple sequence alignment quality of Trl1 (Top 10 Agricultural Fungal Pathogens). Cf. Figures S235 and S238.

| Species | Hit Protein | Hit Length (a.a.) | evalue | align_len | bit_score | identity | positive | score | gaps | % identity | % positive |
|---|---|---|---|---|---|---|---|---|---|---|---|
| G.max | - | - | - | - | - | - | - | - | - | - | - |
| Z.mays | - | - | - | - | - | - | - | - | - | - | - |
| S.tuberosum | - | - | - | - | - | - | - | - | - | - | - |
| O.sativa | - | - | - | - | - | - | - | - | - | - | - |
| H.sapiens | - | - | - | - | - | - | - | - | - | - | - |
| U.maydis | XP_011386892.1 tRNA ligase Ustilago maydis 521 | 380 | 6e-62 | 380 | 224.942 | 140 | 192 | 572 | 45 | 16.9 | 23.2 |
| P.striiformis | XP_047800696.1 hypothetical protein Pst134EA 024359 Puccinia striiformis f. sp. tritici | 736 | 2.6e-80 | 736 | 277.33 | 225 | 340 | 708 | 98 | 27.2 | 41.1 |
| P.oryzae | mRNA M BR32 EuGene 00050301-p1 — transcript=mRNA M BR32 EuGene 00050301 — gene=M BR32 EuGene 00050301 — organism=Pyricularia oryzae BR32 — gene product=unspecified product — transcript product=unspecified product — location=BR32 scaffold00004:1761131-1763962(-) — protein length=943 — sequence SO=supercontig — SO=protein coding gene — is pseudo=false | 815 | 2.2e-160 | 815 | 490.345 | 304 | 450 | 1261 | 71 | 36.8 | 54.4 |
| P.triticina | XP_053026401.1 uncharacterized protein PtA15 13A245 Puccinia triticina | 760 | 1.6e-80 | 760 | 276.559 | 221 | 336 | 706 | 108 | 26.7 | 40.6 |
| P.graminis | XP_003326541.2 hypothetical protein PGTG 07519 Puccinia graminis f. sp. tritici CRL 75-36--700-3 | 736 | 4.8e-85 | 736 | 290.041 | 228 | 362 | 741 | 85 | 27.6 | 43.8 |
| M.graminicola | ZTRI 1.1976.mRNA-p1 — transcript=ZTRI 1.1976.mRNA — gene=ZTRI 1.1976 — organism=Zymoseptoria tritici IPO323 — gene product=similar to trna ligase — transcript product=similar to trna ligase — location=Ztri chr 1:5571847-5574309(-) — protein length=820 — sequence SO=chromosome — SO=protein coding gene — is pseudo=false | 848 | 4.6e-158 | 848 | 480.33 | 311 | 457 | 1235 | 81 | 37.6 | 55.3 |
| F.graminearum | XP_011318754.1 hypothetical protein FGSG 11594 Fusarium graminearum PH-1 | 773 | 3.9e-157 | 773 | 477.248 | 293 | 429 | 1227 | 75 | 35.4 | 51.9 |
| C.truncatum | XP_036575045.1 tRNA ligase Colletotrichum truncatum | 807 | 4.3e-171 | 807 | 515.383 | 317 | 448 | 1326 | 70 | 38.3 | 54.2 |
| B.cinerea | XP_001558448.2 Bctrl1 Botrytis cinerea B05.10 | 825 | 3.9e-163 | 825 | 494.967 | 311 | 461 | 1273 | 88 | 37.6 | 55.7 |
| B.graminis | VDB94992.1 — transcript=BGT96224V316 LOCUS8047 t1 — gene=BGT96224V316 LOCUS8047 — organism=Blumeria graminis f. sp. tritici 96224 — gene product=unspecified product — transcript product=unspecified product — location=LR026993:6965370-6967882(+) — protein length=803 — sequence SO=chromosome — SO=protein coding gene — is pseudo=false | 780 | 3.9e-142 | 780 | 437.958 | 294 | 427 | 1125 | 75 | 35.6 | 51.6 |

Table S38: Pairwise alignment info from yeast Trl1 (DEG20010555), cf. Figure S235.

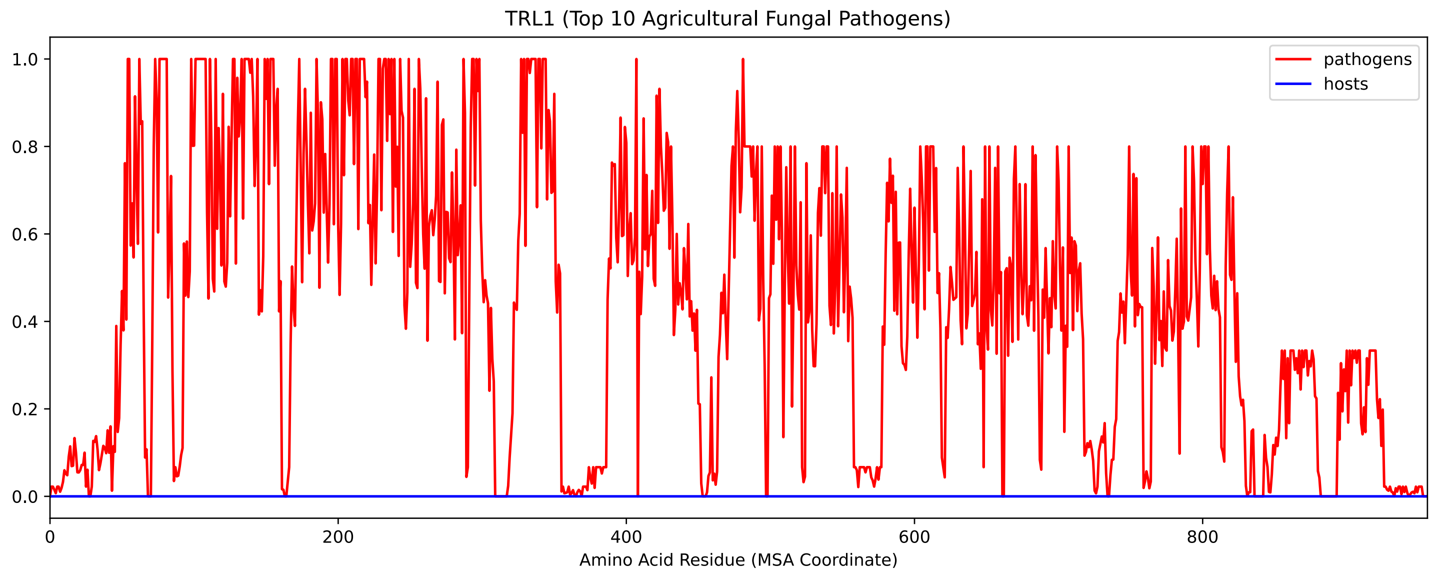


Figure S237: Sneath Similarity of Trl1 for Top 10 Agricultural Fungal Pathogens, cf. Figure S235

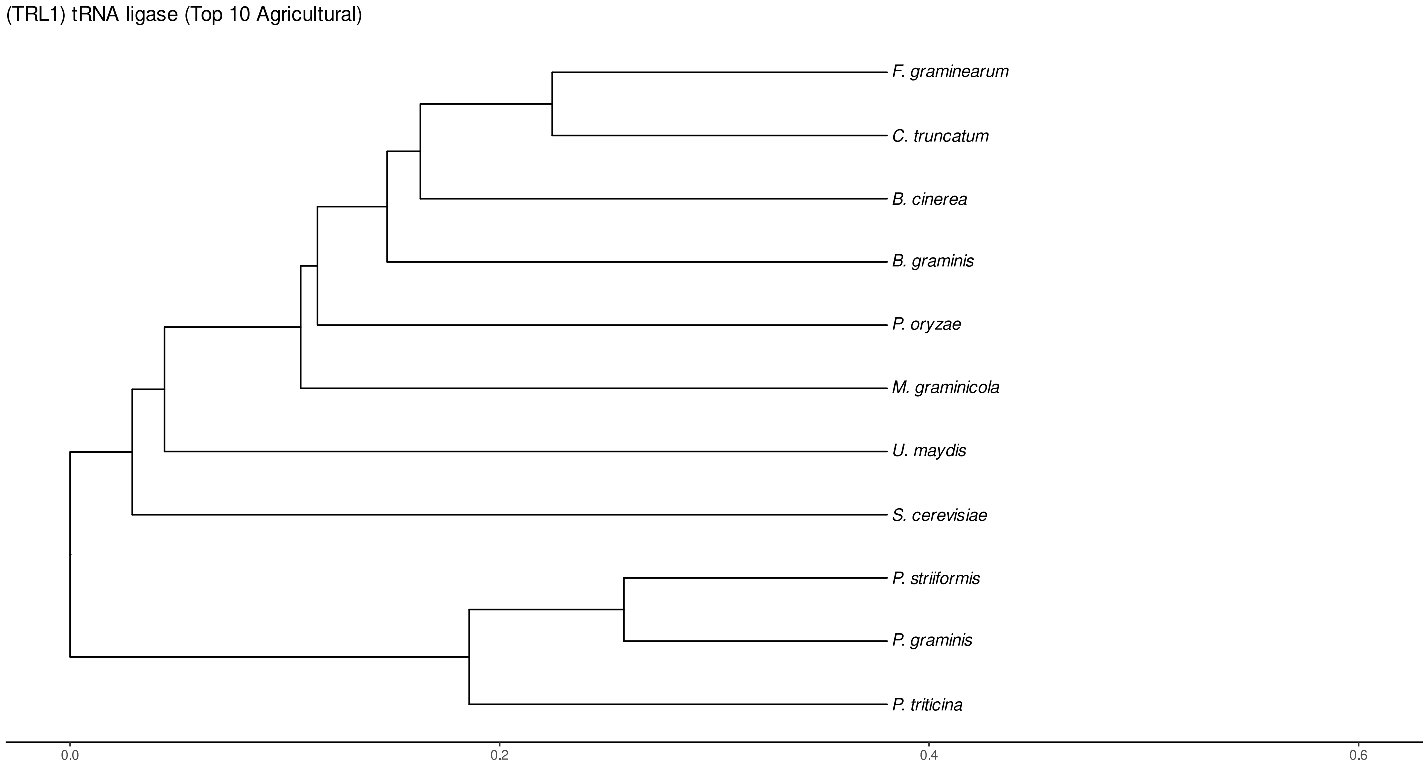


Figure S238: Phylogeny of hits for Trl1 made using clustal omega and visualized using ggtree, cf. Figures S235 and S236.

### S4.16.3 NR

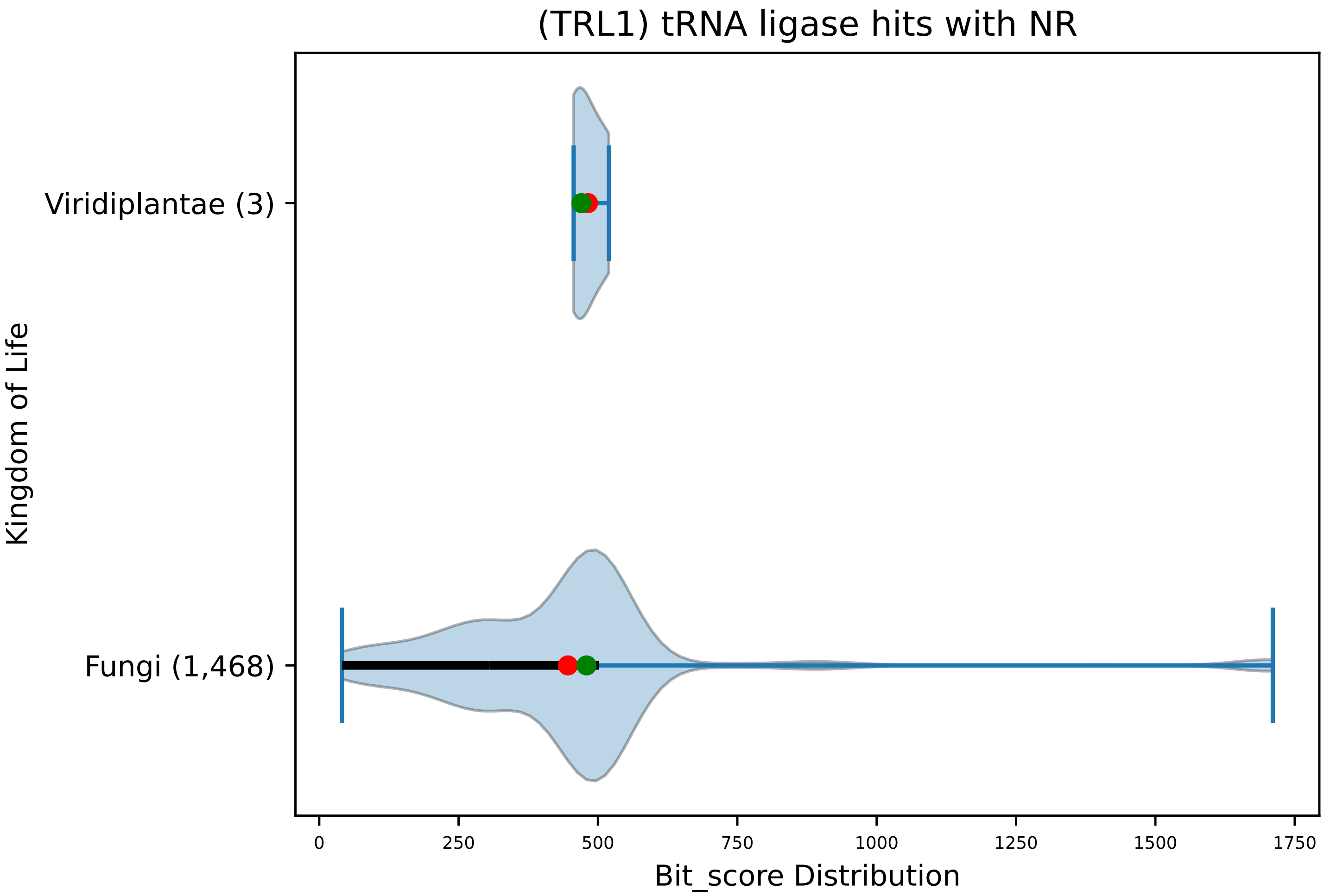


Figure S239: Non-redundant (NR) protein hits for DEG20010555/Trl1, with expectation value of no more than 0.1. Green points are medians, and red points are arithmetic means.

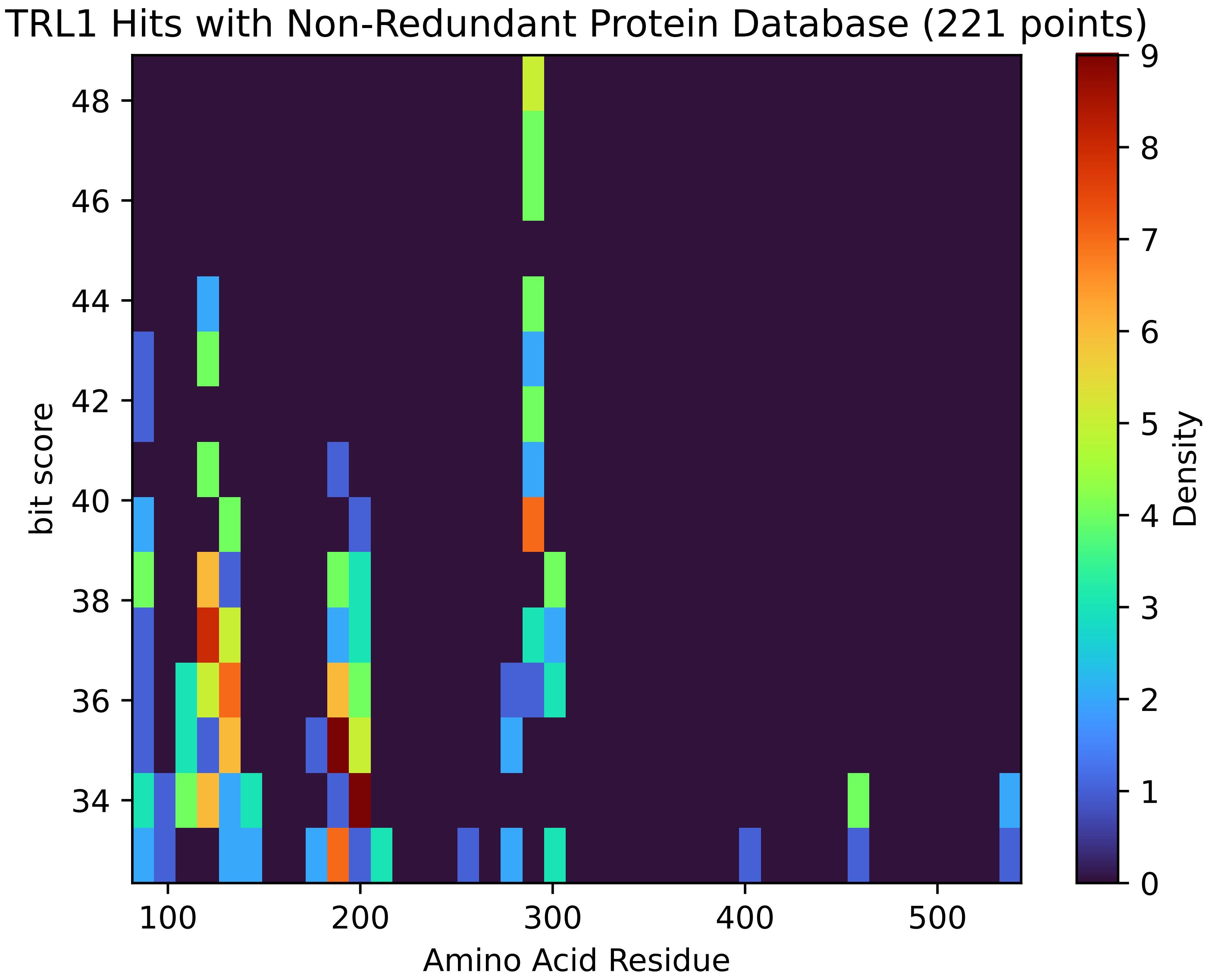


Figure S240: Non-redundant (NR) protein hits for Trl1 in the kingdom Viridiplantae.

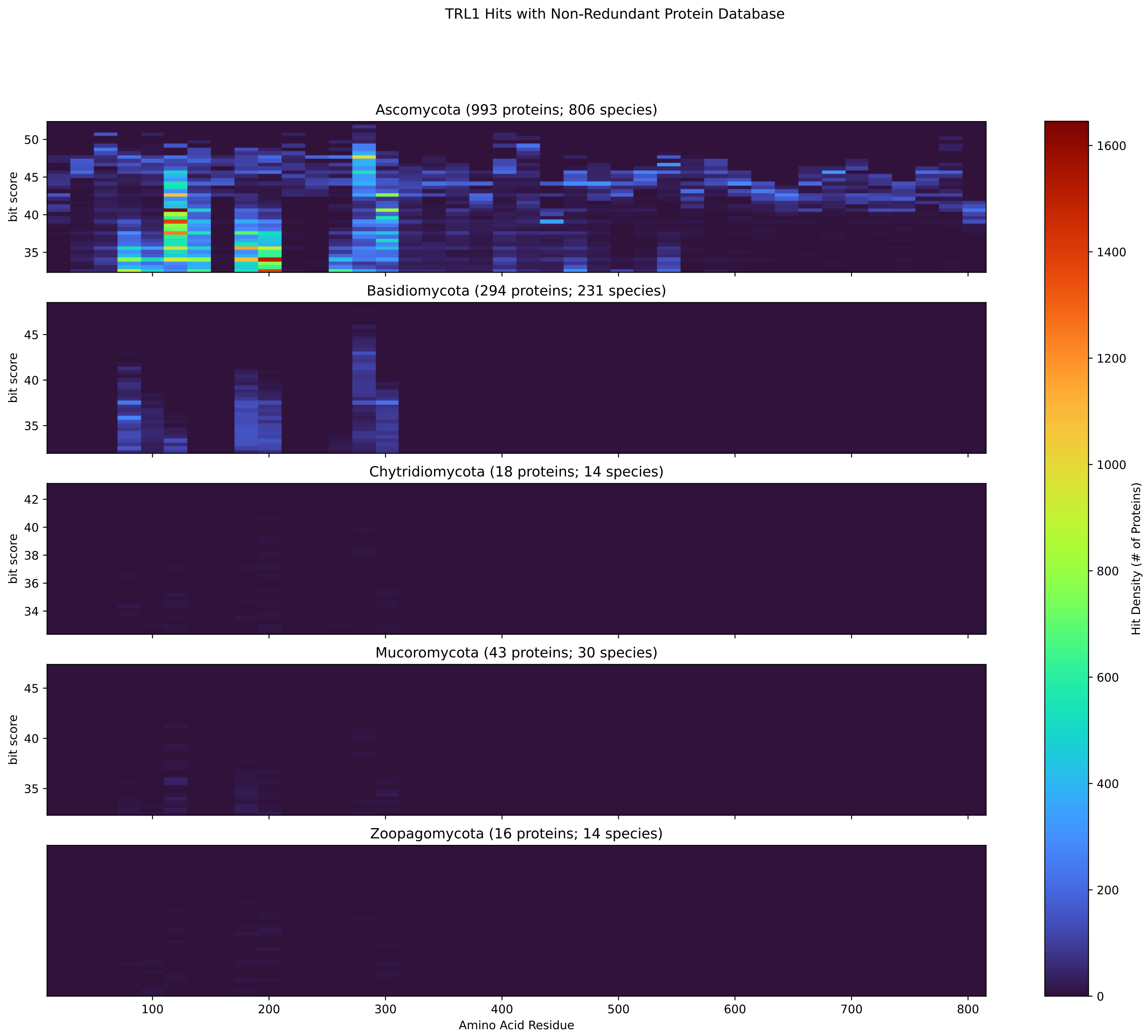


Figure S241: Non-redundant (NR) protein hits for Trl1 in the kingdom Fungi.

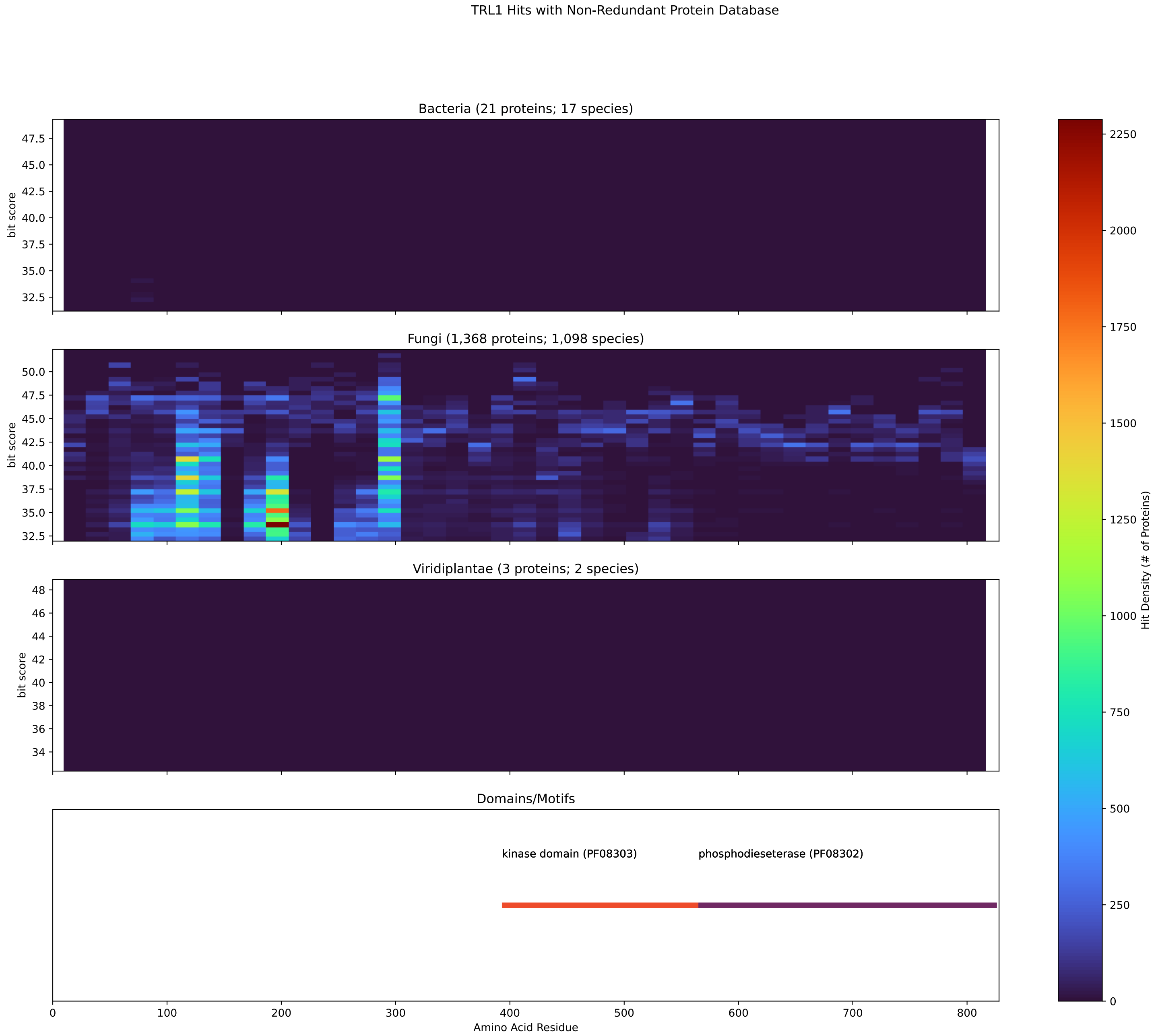


Figure S242: Non-redundant (NR) protein hits for DEG20010555/Trl1 at 20 amino acid length queries.

#### S4.17 Yef3

##### S4.17.1 WHO Critical Pathogens

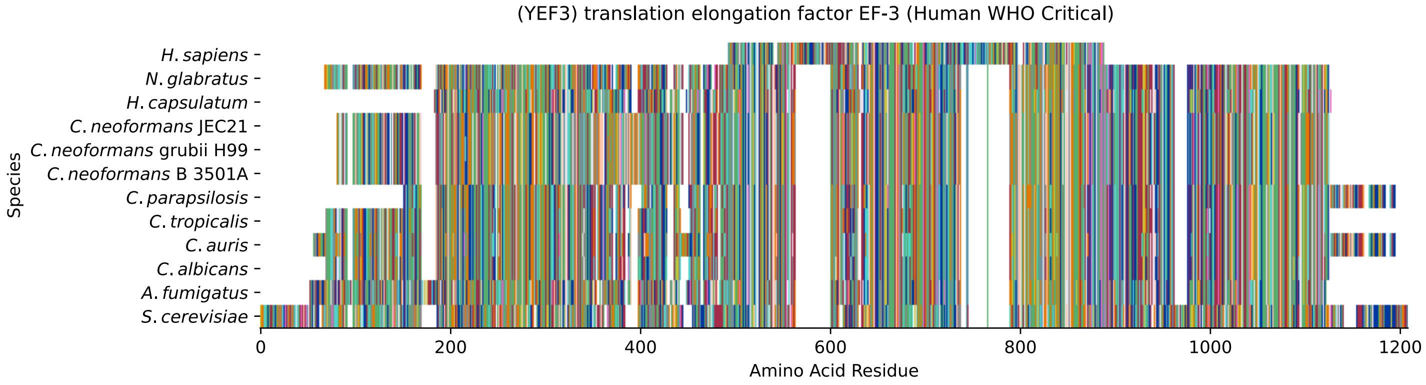


Figure S243: Multiple sequence alignment of yeast Yef3 (WHO Critical Pathogens). Cf. Figure S244 for alignment quality, and Figure S245 for Sneath similarity. Cf. Table S39 for protein names, and pairwise alignment metrics with yeast Yef3.

#### Yef3 MSA Quality

| | *A. f.* | *C. a.* | *C. a.* | *C. p.* | *C. t.* | *C. n.* | *C. n.* | *C. n.* | *H. c.* | *N. g.* | *S. c.* |
|---|---|---|---|---|---|---|---|---|---|---|---|
| *A. fumigatus* | 1957.96 | 922.924 | 951.429 | 900.582 | 918.687 | 801.586 | 801.201 | 801.586 | 1395.18 | 955.666 | 631.328 |
| *C. albicans* | 909.057 | 1857.03 | 1520.75 | 1466.44 | 1715.66 | 794.653 | 795.423 | 794.653 | 949.503 | 1277.69 | 597.43 |
| *C. auris* | 939.488 | 1520.75 | 2024.59 | 1506.12 | 1501.88 | 805.823 | 801.971 | 805.823 | 962.6 | 1270.37 | 616.305 |
| *C. parapsilosis* | 897.501 | 1484.93 | 1521.14 | 1818.13 | 1468.75 | 806.209 | 801.971 | 806.209 | 962.214 | 1221.07 | 604.749 |
| *C. tropicalis* | 908.672 | 1726.07 | 1511.89 | 1472.6 | 1847.4 | 803.512 | 803.512 | 803.512 | 950.658 | 1290.4 | 600.897 |
| *C. neoformans* B 3501A | 787.334 | 794.653 | 805.823 | 788.104 | 793.882 | 1877.83 | 1866.66 | 1877.83 | 803.897 | 797.349 | 658.292 |
| *C. neoformans* grubii H99 | 787.719 | 795.423 | 801.971 | 784.252 | 793.497 | 1866.66 | 1875.91 | 1866.66 | 804.668 | 793.497 | 660.603 |
| *C. neoformans* JEC21 | 787.334 | 794.653 | 805.823 | 788.104 | 793.882 | 1877.83 | 1866.66 | 1877.83 | 803.897 | 797.349 | 658.292 |
| *H. capsulatum* | 1383.24 | 949.503 | 962.6 | 952.199 | 946.806 | 803.897 | 804.668 | 803.897 | 1702.18 | 951.044 | 629.787 |
| *N. glabratus* | 956.822 | 1295.8 | 1287.32 | 1219.53 | 1305.81 | 807.364 | 804.283 | 807.364 | 964.911 | 1869.74 | 620.928 |
| *S. cerevisiae* | 633.254 | 609.757 | 629.402 | 609.372 | 606.29 | 676.011 | 678.707 | 676.011 | 647.121 | 621.698 | 2163.27 |

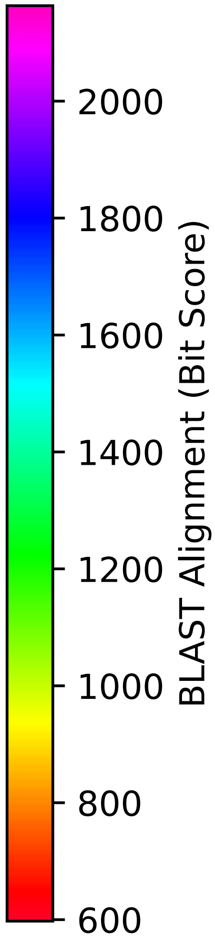


Figure S244: Multiple sequence alignment quality of Yef3 (WHO Critical Pathogens). Cf. Figures S243 and S246.

| Species | Hit Protein | Hit Length (a.a.) | evalue | align_len | bit_score | identity | positive | score | gaps | % identity | % positive |
|---|---|---|---|---|---|---|---|---|---|---|---|
| H.sapiens | NP_001338227.1 ATP-binding cassette sub-family F member 3 isoform 2 Homo sapiens | 396 | 3.2e-39 | 396 | 159.073 | 117 | 172 | 401 | 98 | 11.2 | 16.5 |
| N.glabratus | XP_445575.1 uncharacterized protein CAGL0D03674g Nakaseomyces glabratus | 939 | 0 | 939 | 620.928 | 364 | 545 | 1600 | 60 | 34.9 | 52.2 |
| H.capsulatum | XP_045289814.1 elongation factor 3 Histoplasma capsulatum G-186AR | 851 | 0 | 851 | 647.892 | 368 | 511 | 1670 | 58 | 35.2 | 48.9 |
| C.neoformans.JEC21 | XP_566522.1 mRNA export factor elf1, putative Cryptococcus neoformans var. neoformans JEC-21 | 922 | 0 | 922 | 676.011 | 377 | 527 | 1743 | 38 | 36.1 | 50.5 |
| C.neoformans.grubii.H99 | XP_012046531.1 elongation factor 3 Cryptococcus neoformans var. grubii H99 | 922 | 0 | 922 | 678.322 | 379 | 527 | 1749 | 38 | 36.3 | 50.5 |
| C.neoformans.B.3501A | XP_778089.1 hypothetical protein CNBA0920 Cryptococcus neoformans var. neoformans B-3501A | 922 | 0 | 922 | 676.011 | 377 | 527 | 1743 | 38 | 36.1 | 50.5 |
| C.parapsilosis | XP_036668290.1 uncharacterized protein CPAR2 701340 Candida parapsilosis | 916 | 0 | 916 | 610.527 | 349 | 538 | 1573 | 62 | 33.4 | 51.5 |
| C.tropicalis | XP_002546443.1 mRNA export factor elf1 Candida tropicalis MYA-3404 | 932 | 0 | 932 | 606.675 | 343 | 537 | 1563 | 55 | 32.9 | 51.4 |
| C.auris | XP_028888995.2 hypothetical protein Candida auris | 1010 | 0 | 1010 | 632.098 | 369 | 573 | 1629 | 61 | 35.3 | 54.9 |
| C.albicans | XP_716466.1 Elf1p Candida albicans SC5314 | 944 | 0 | 944 | 609.372 | 349 | 541 | 1570 | 76 | 33.4 | 51.8 |
| A.fumigatus | XP_747719.1 mRNA-nucleus export ATPase (Elf1), putative Aspergillus fumigatus Af293 | 966 | 0 | 966 | 634.024 | 385 | 553 | 1634 | 64 | 36.9 | 53.0 |

Table S39: Pairwise alignment info from yeast Yef3 (DEG20010729), cf. Figure S243.

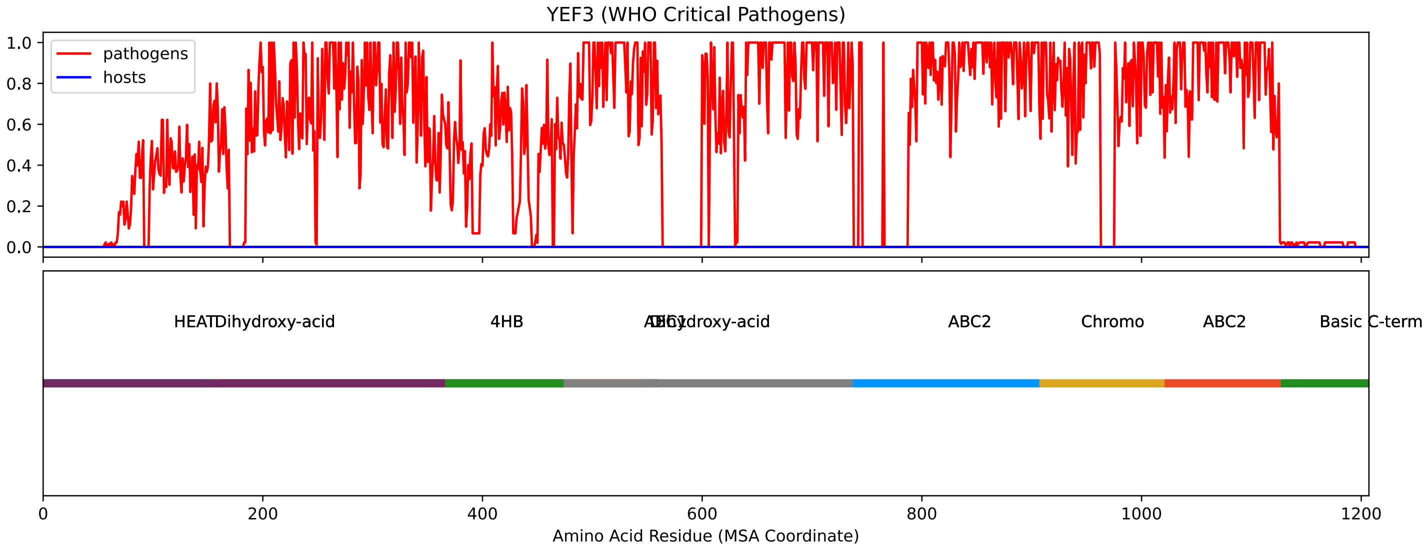


Figure S245: Sneath Similarity of Yef3 for WHO Critical Pathogens, cf. Figure S243

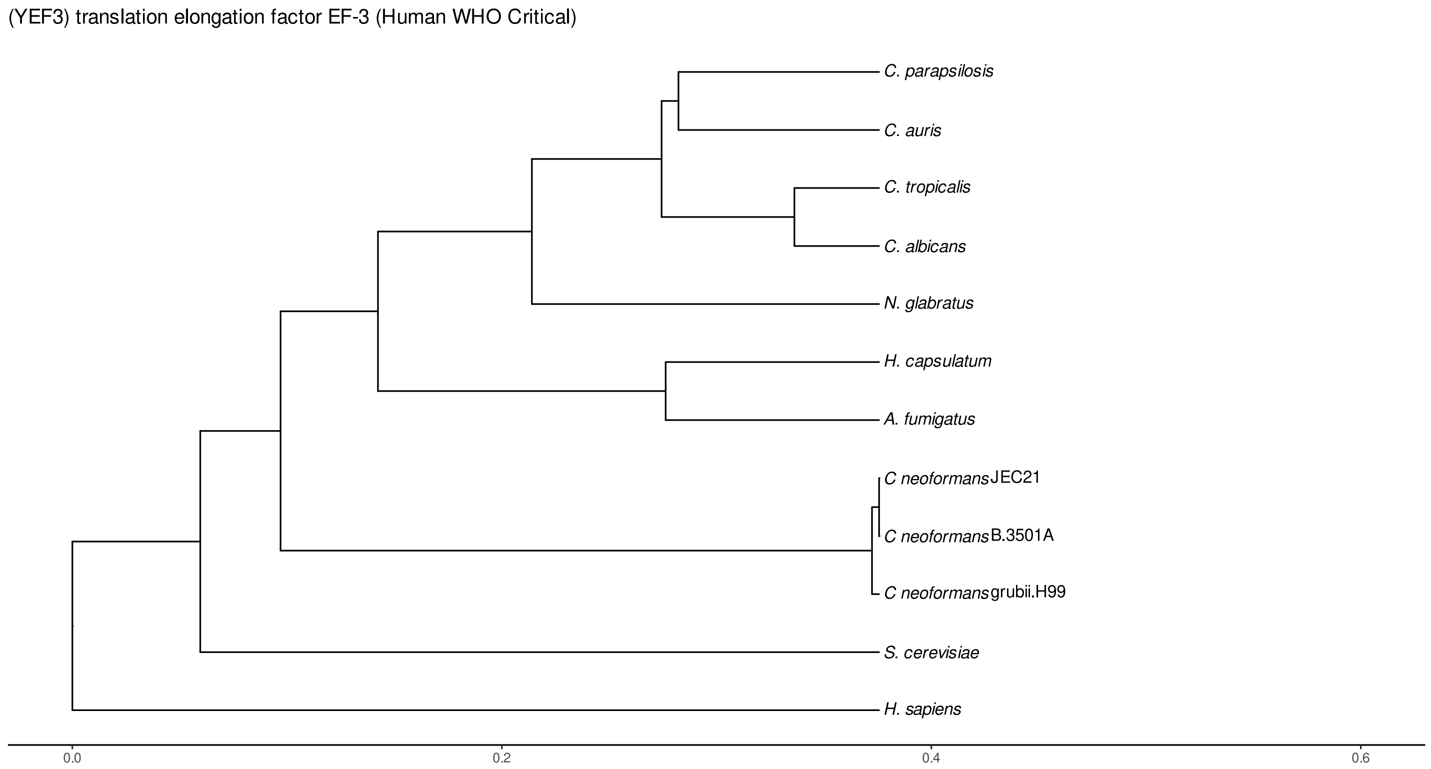


Figure S246: Phylogeny of hits for Yef3 made using clustal omega and visualized using ggtree, cf. Figures S243 and S244.

##### S4.17.2 Top 10 Agricultural Fungal Pathogens

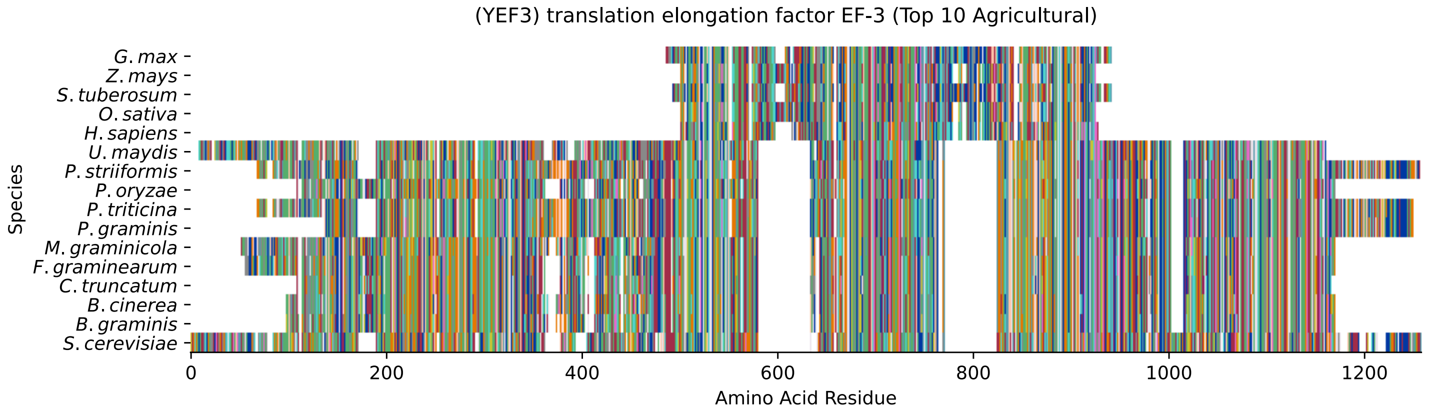


Figure S247: Multiple sequence alignment of yeast Yef3 (Top 10 Agricultural Fungal Pathogens). Cf. Figure S248 for alignment quality, and Figure S249 for Sneath similarity. Cf. Table S40 for protein names, and pairwise alignment metrics with yeast Yef3.

#### Yef3 MSA Quality

| | *B.g.* | *B.c.* | *C.t.* | *F.g.* | *M.g.* | *P.g.* | *P.t.* | *P.o.* | *P.s.* | *U.m.* | *S.c.* |
|---|---|---|---|---|---|---|---|---|---|---|---|
| *B. graminis* | 1890.16 | 1580.84 | 1466.06 | 1485.7 | 1377.07 | 778.859 | 780.015 | 1443.33 | 778.859 | 821.617 | 0 |
| *B. cinerea* | 1580.84 | 1894.78 | 1455.66 | 1471.83 | 1380.16 | 784.637 | 785.408 | 1457.58 | 780.015 | 845.114 | 0 |
| *C. truncatum* | 1466.06 | 1455.66 | 1849.33 | 1588.93 | 1352.42 | 770.77 | 776.548 | 1536.55 | 764.992 | 823.157 | 0 |
| *F. graminearum* | 1485.7 | 1471.83 | 1588.93 | 1956.8 | 1406.35 | 793.112 | 788.875 | 1566.59 | 781.941 | 838.565 | 0 |
| *M. graminicola* | 1377.07 | 1380.16 | 1352.42 | 1406.35 | 1993.01 | 780.4 | 782.711 | 1339.71 | 776.548 | 844.343 | 0 |
| *P. graminis* | 778.859 | 784.637 | 770.77 | 793.112 | 780.4 | 1975.67 | 1924.06 | 781.941 | 1919.05 | 951.429 | 0 |
| *P. triticina* | 780.015 | 785.408 | 776.548 | 788.875 | 782.711 | 1924.06 | 2100.86 | 784.637 | 2032.3 | 976.082 | 0 |
| *P. oryzae* | 1443.33 | 1457.58 | 1536.55 | 1566.59 | 1339.71 | 781.941 | 784.637 | 1854.72 | 776.548 | 806.979 | 0 |
| *P. striiformis* | 779.63 | 785.793 | 771.926 | 787.719 | 782.326 | 1919.05 | 2032.3 | 782.711 | 2116.27 | 990.334 | 0 |
| *U. maydis* | 838.565 | 861.677 | 844.729 | 864.759 | 871.307 | 975.696 | 1002.28 | 829.706 | 1004.59 | 2025.75 | 0 |
| *S. cerevisiae* | 0 | 0 | 0 | 0 | 0 | 0 | 0 | 0 | 0 | 0 | 0 |

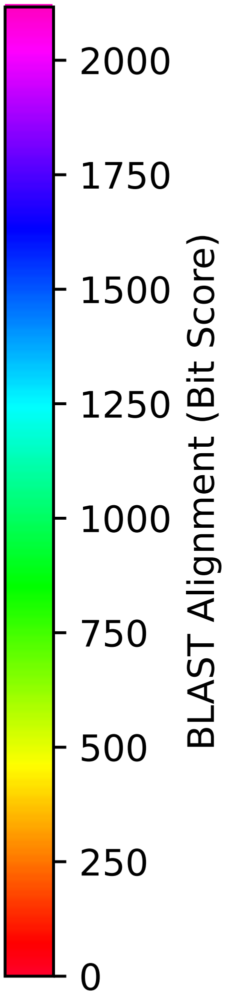


Figure S248: Multiple sequence alignment quality of Yef3 (Top 10 Agricultural Fungal Pathogens). Cf. Figures S247 and S250.

| Species | Hit Protein | Hit Length (a.a.) | evalue | align_len | bit_score | identity | positive | score | gaps | % identity | % positive |
|---|---|---|---|---|---|---|---|---|---|---|---|
| G.max | XP_003542630.1 ABC transporter F family member 4 Glycine max | 439 | 3.6e-47 | 439 | 182.185 | 124 | 195 | 461 | 117 | 11.9 | 18.7 |
| Z.mays | NP_001349290.1 ABC transporter F family member 3-like Zea mays | 408 | 6.8e-42 | 408 | 165.622 | 120 | 174 | 418 | 114 | 11.5 | 16.7 |
| S.tuberosum | XP_006362455.1 PREDICTED: ABC transporter F family member 4-like Solanum tuberosum | 431 | 1.8e-48 | 431 | 185.267 | 127 | 192 | 469 | 114 | 12.2 | 18.4 |
| O.sativa | XP_015626086.1 ABC transporter F family member 3 Oryza sativa Japonica Group | 408 | 2e-42 | 408 | 167.162 | 120 | 174 | 422 | 114 | 11.5 | 16.7 |
| H.sapiens | NP_001338227.1 ATP-binding cassette sub-family F member 3 isoform 2 Homo sapiens | 396 | 3.2e-39 | 396 | 159.073 | 117 | 172 | 401 | 98 | 11.2 | 16.5 |
| U.maydis | XP_011390724.1 putative mrna export factor elf1 Ustilago maydis 521 | 1001 | 0 | 1001 | 689.878 | 406 | 580 | 1779 | 59 | 38.9 | 55.6 |
| P.striiformis | XP_047803383.1 hypothetical protein Pst134EA 019464 Puccinia striiformis f. sp. tritici | 1043 | 0 | 1043 | 644.425 | 374 | 581 | 1661 | 83 | 35.8 | 55.7 |
| P.oryzae | mRNA M BR32 EuGene 00061071-p1 — transcript=mRNA M BR32 EuGene 00061071 — gene=M BR32 EuGene 00061071 — organism=Pyricularia oryzae BR32 — gene product=unspecified product — transcript product=unspecified product — location=BR32 scaffold00005:2291252-2294682(-) — protein length=1118 — sequence SO=supercontig — SO=protein coding gene — is pseudo=false | 922 | 0 | 922 | 669.848 | 381 | 548 | 1727 | 64 | 36.5 | 52.5 |
| P.triticina | XP_053024635.1 uncharacterized protein PtA15 10A503 Puccinia triticina | 1040 | 0 | 1040 | 643.269 | 380 | 578 | 1658 | 89 | 36.4 | 55.4 |
| P.graminis | XP_003319416.1 hypothetical protein PGTG 01590 Puccinia graminis f. sp. tritici CRL 75-36-700-3 | 977 | 0 | 977 | 645.966 | 369 | 559 | 1665 | 83 | 35.3 | 53.5 |
| M.graminicola | ZTRI 5.109.mRNA-p1 — transcript=ZTRI 5.109.mRNA — gene=ZTRI 5.109 — organism=Zymoseptoria tritici IPO323 — gene product=similar to mRNA-nucleus export ATPase (Elf1)/ABC Transporter — transcript product=similar to mRNA-nucleus export ATPase (Elf1)/ABC Transporter — location=Ztri chr 5:449343-452784(+-) — protein length=1128 — sequence SO=chromosome — SO=protein coding gene — is pseudo=false | 984 | 0 | 984 | 625.935 | 376 | 547 | 1613 | 76 | 36.0 | 52.4 |
| F.graminearum | XP_011320226.1 prion formation protein 1 Fusarium graminearum PH-1 | 1001 | 0 | 1001 | 664.07 | 392 | 573 | 1712 | 90 | 37.5 | 54.9 |
| C.truncatum | XP_036575201.1 ABC transporter Colletotrichum truncatum | 916 | 0 | 916 | 672.544 | 383 | 545 | 1734 | 60 | 36.7 | 52.2 |
| B.cinerea | XP_001545837.2 Bcnew1 Botrytis cinerea B05.10 | 935 | 0 | 935 | 667.537 | 388 | 550 | 1721 | 63 | 37.2 | 52.7 |
| B.graminis | VDB89801.1 — transcript=BGT96224V316 LOCUS5359 t1 — gene=BGT96224V316 LOCUS5359 — organism=Blumeria graminis f. sp. tritici 96224 — gene product=unspecified product — transcript product=unspecified product — location=LR026990:14052689-14056061(+) — protein length=1104 — sequence SO=chromosome — SO=protein coding gene — is pseudo=false | 935 | 0 | 935 | 647.121 | 385 | 545 | 1668 | 66 | 36.9 | 52.2 |

Table S40: Pairwise alignment info from yeast Yef3 (DEG20010729), cf. Figure S247.

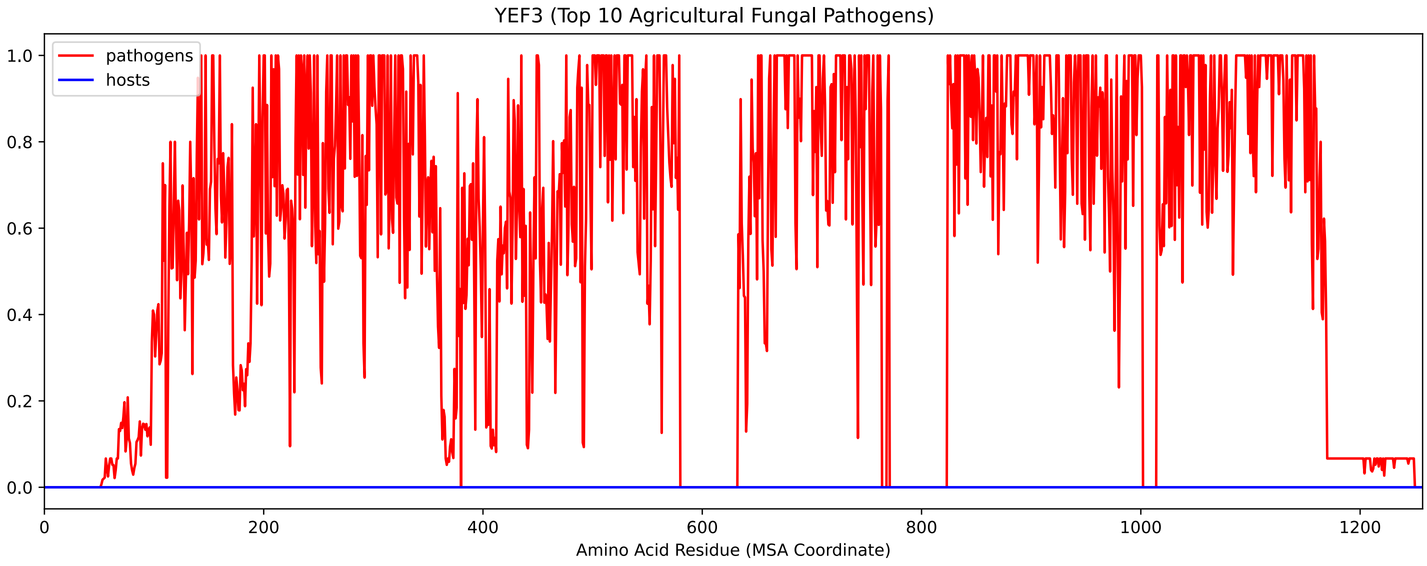


Figure S249: Sneath Similarity of Yef3 for Top 10 Agricultural Fungal Pathogens, cf. Figure S247

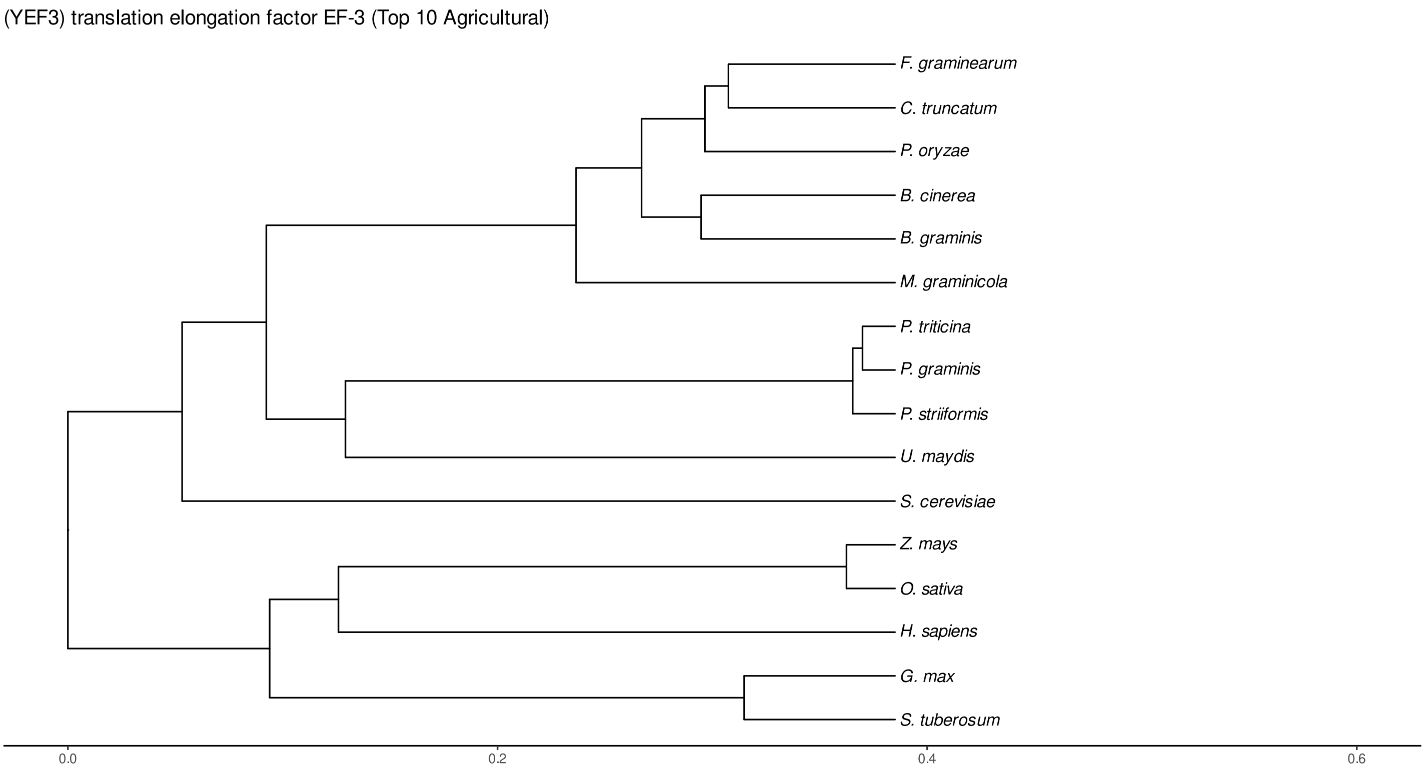


Figure S250: Phylogeny of hits for Yef3 made using clustal omega and visualized using ggtree, cf. Figures S247 and S248.

#### S4.17.3 NR

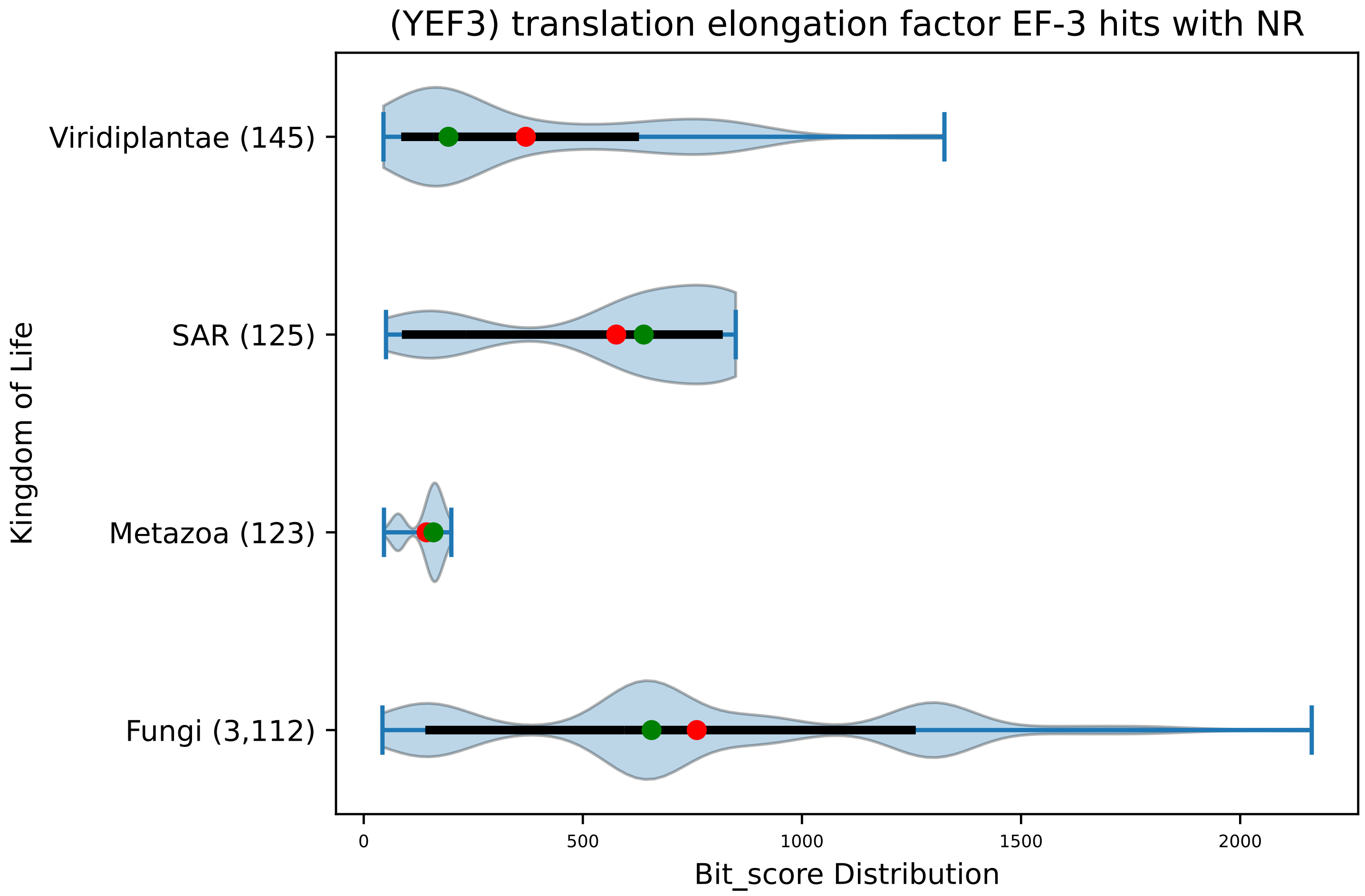


Figure S251: Non-redundant (NR) protein hits for DEG20010729/Yef3, with expectation value of no more than 0.1. Green points are medians, and red points are arithmetic means.

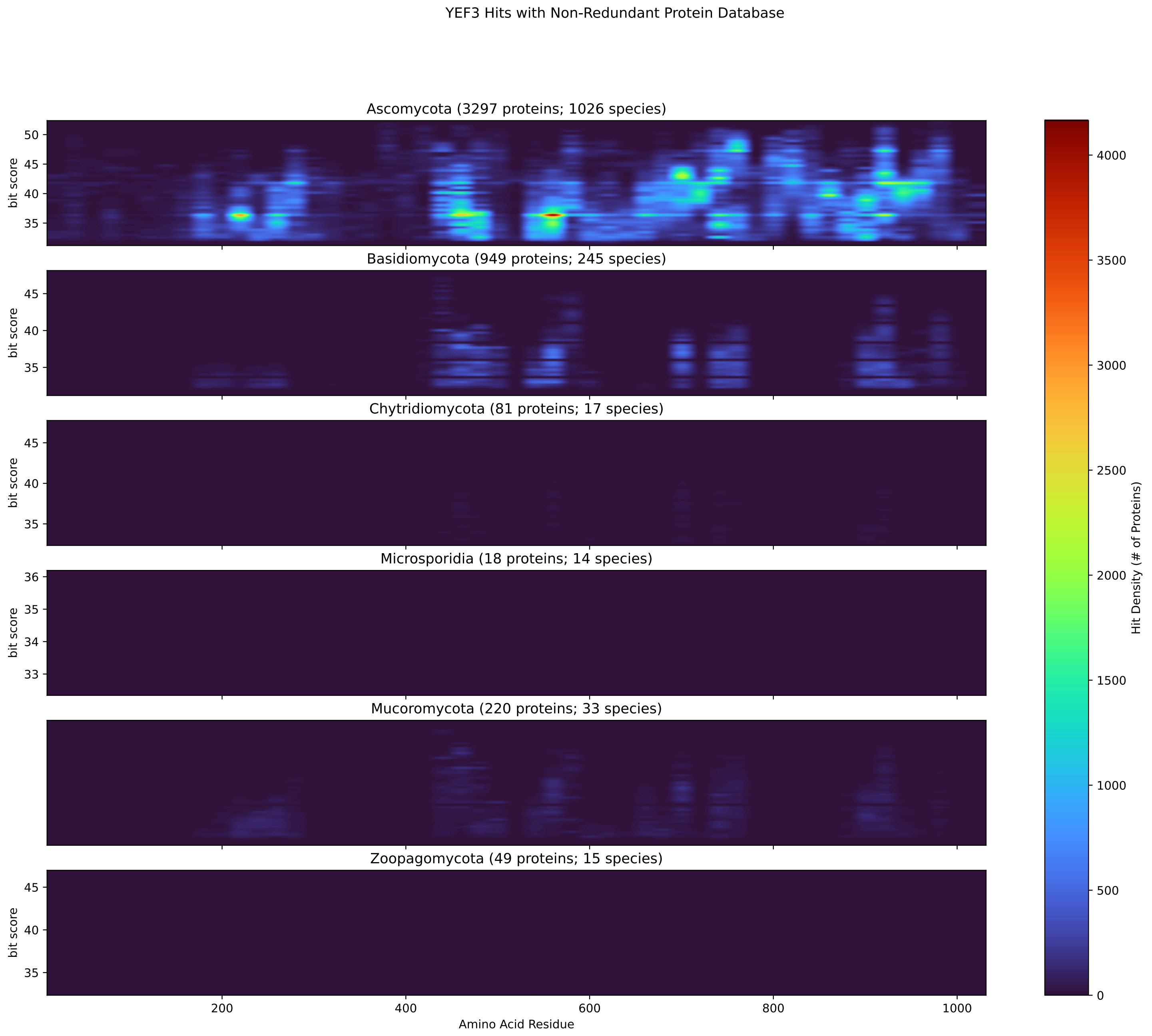


Figure S252: Non-redundant (NR) protein hits for Yef3 in the kingdom Fungi.

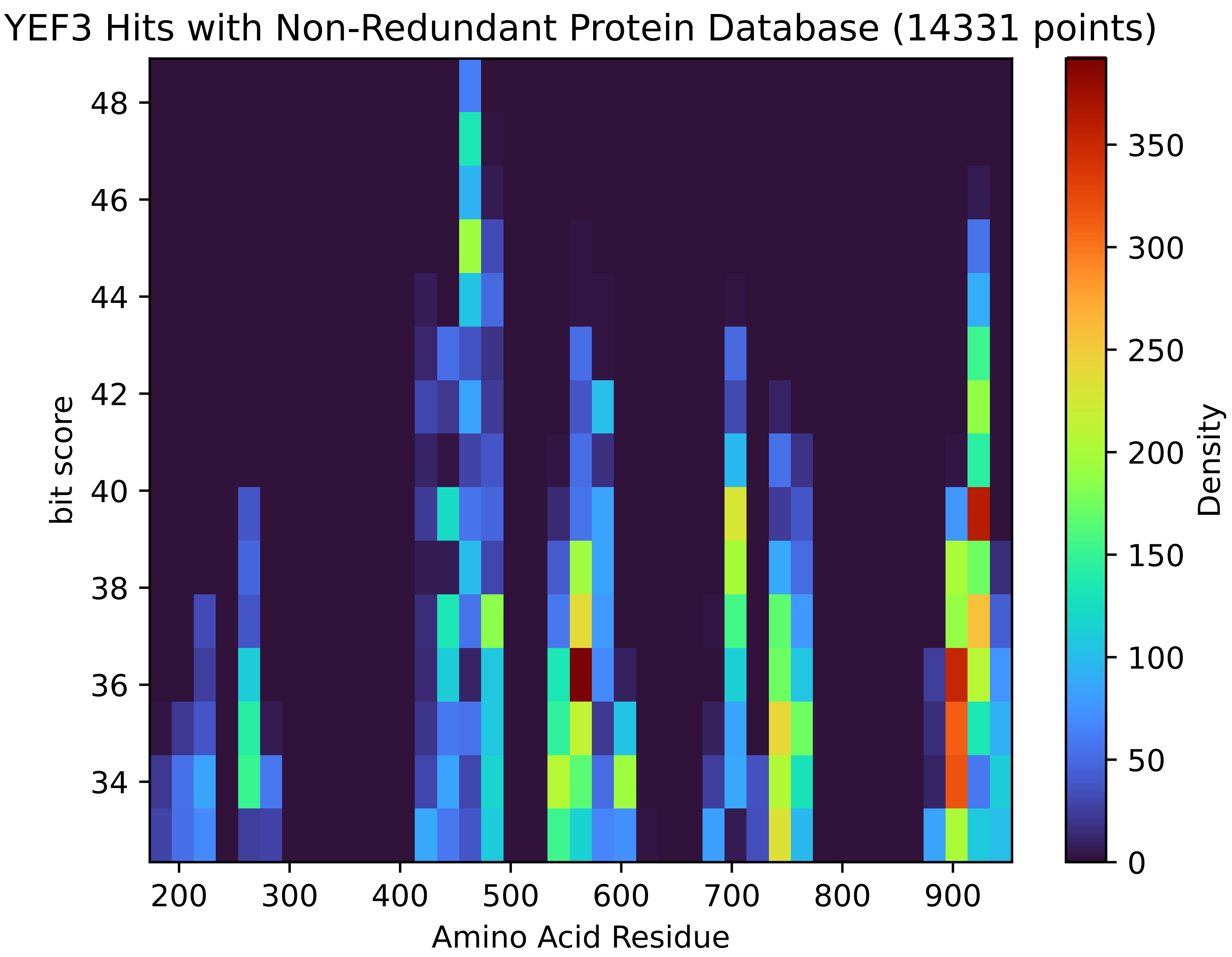


Figure S253: Non-redundant (NR) protein hits for Yef3 in the kingdom SAR.

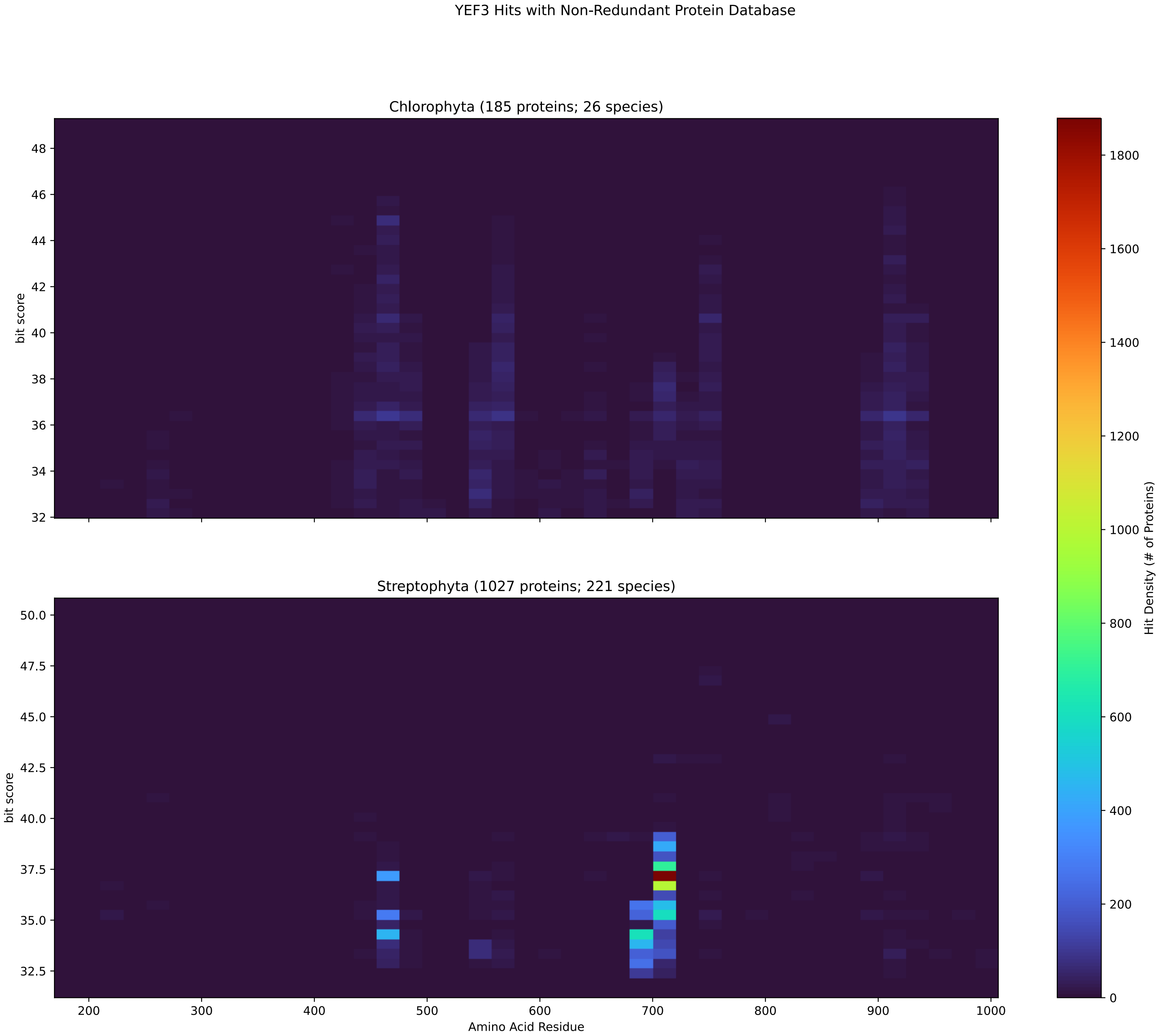


Figure S254: Non-redundant (NR) protein hits for Yef3 in the kingdom Viridiplantae.

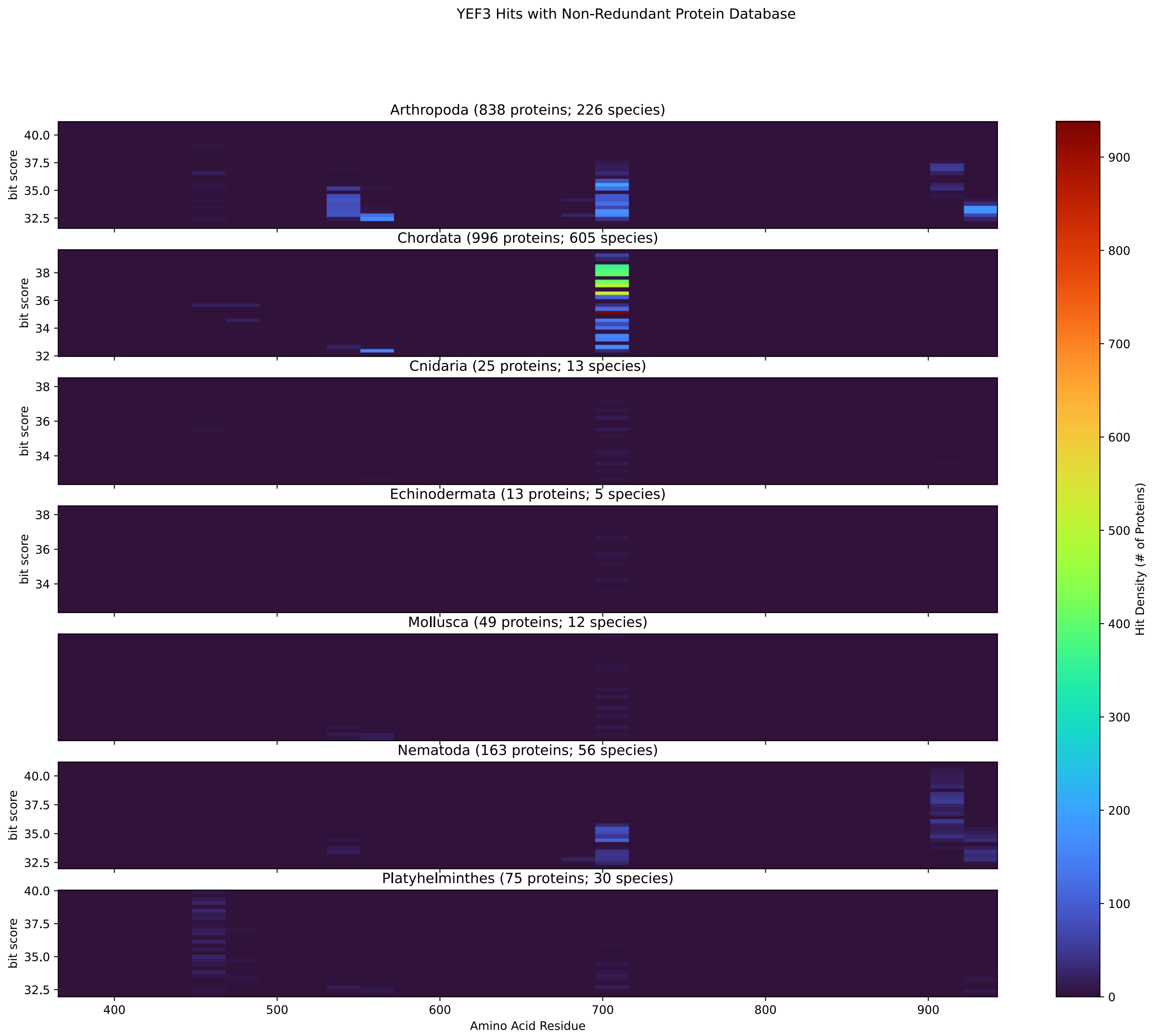


Figure S255: Non-redundant (NR) protein hits for Yef3 in the kingdom Metazoa.

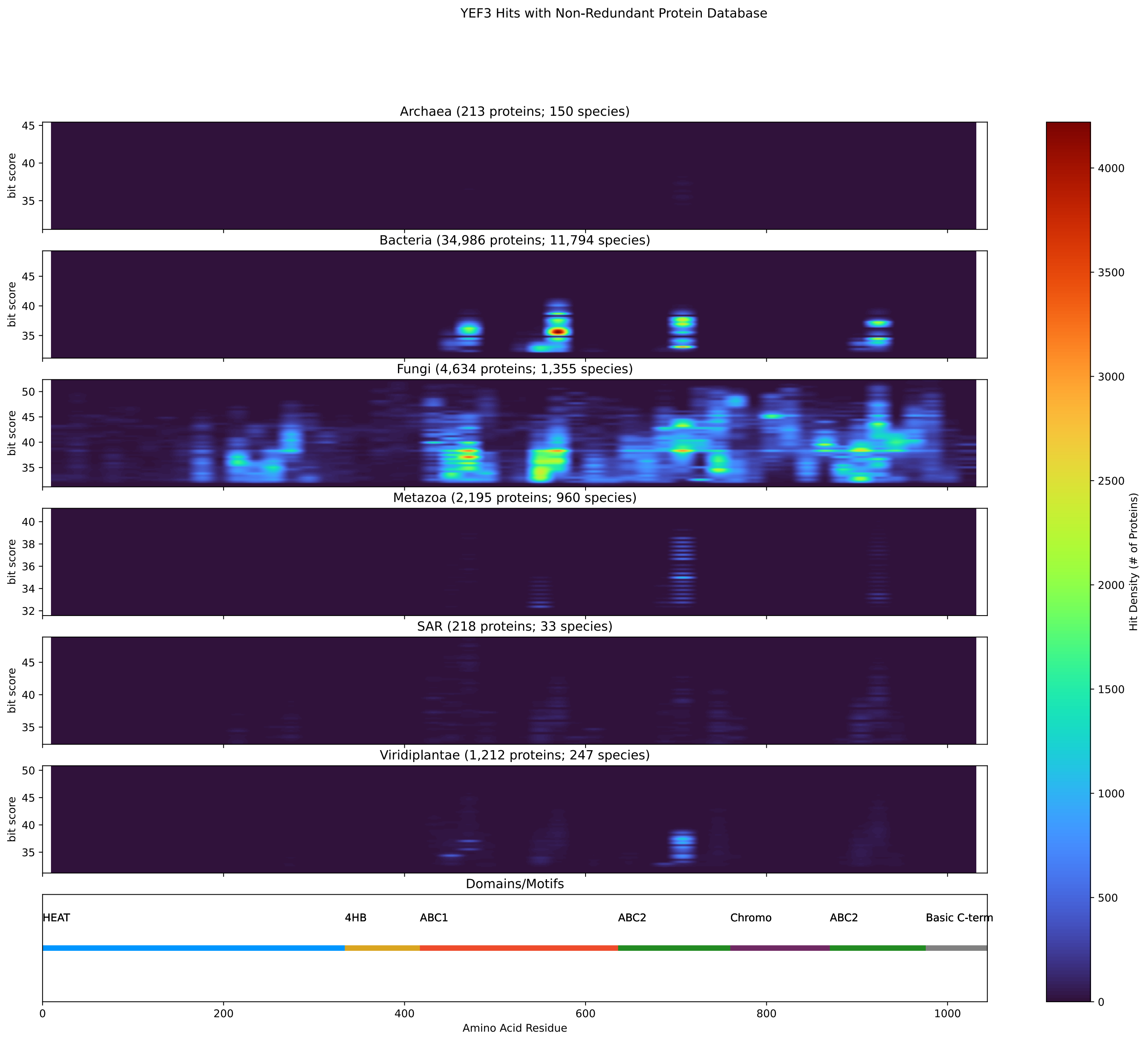


Figure S256: Non-redundant (NR) protein hits for DEG20010729/Yef3 at 20 amino acid length queries.

### S5 Previously identified protein targets

#### S5.1 Erg11

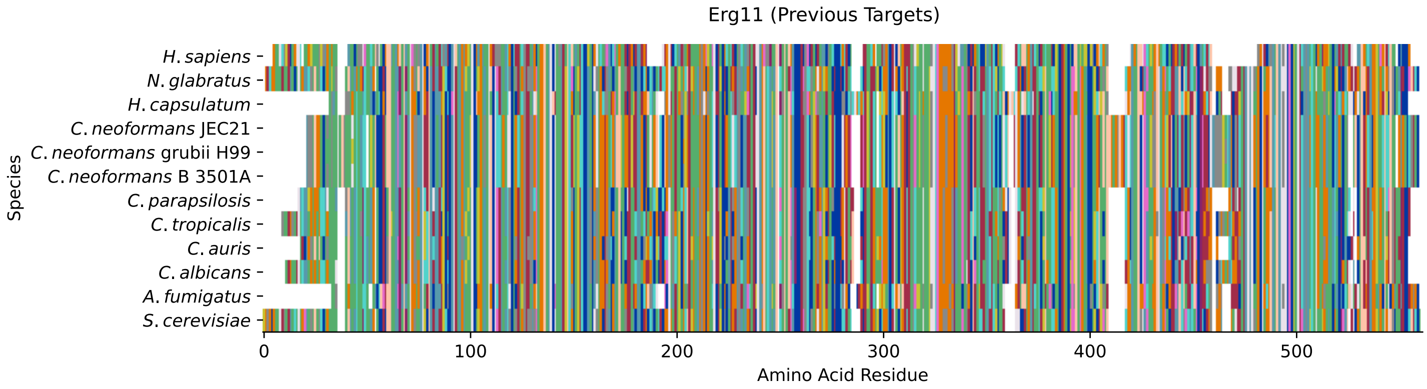


Figure S257: MSA for Erg11; cf. Table S41 for gene names and exact alignment values.

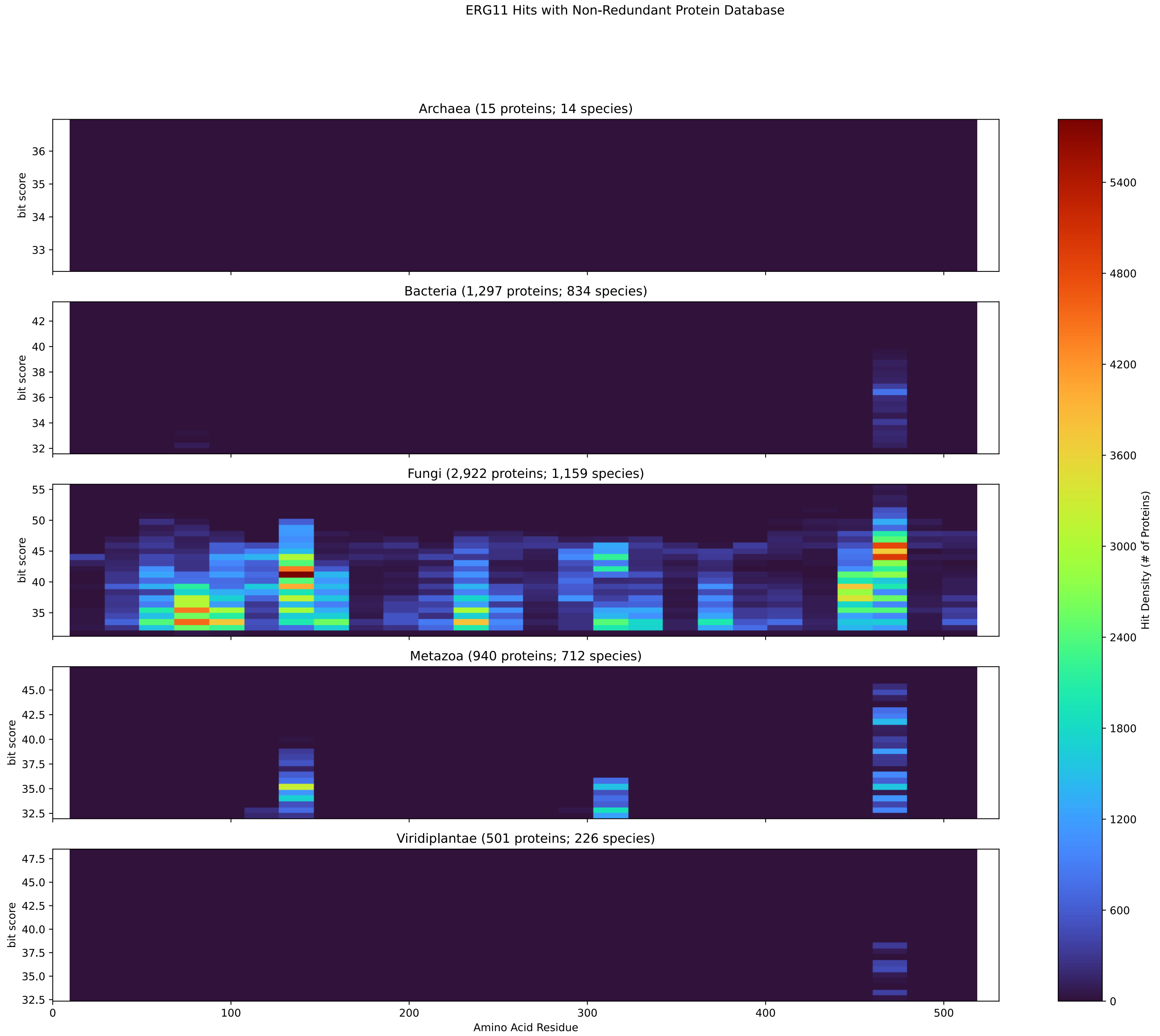


Figure S258: Histogram of Erg11 hits against NR in 20-aa windows

| Species | Hit Protein | Hit Length (a.a.) | evalue | align_len | bit_score | identity | positive | score | gaps | % identity | % positive |
|---|---|---|---|---|---|---|---|---|---|---|---|
| H.sapiens | lanosterol 14-alpha demethylas- e isoform 1 precursor Homo sap- iens | 522 | 9e-104 | 522 | 323.168 | 191 | 290 | 827 | 33 | 36.0 | 54.6 |
| N.glabratus | uncharacterized protein CAGL0E- 04334g Nakaseomyces glabratus | 530 | 0 | 530 | 938.717 | 440 | 486 | 2425 | 2 | 82.9 | 91.5 |
| H.capsulatum | cytochrome P450 sterol 14 alph- a-demethylase Histoplasma caps- ulatum G186AR | 503 | 6.1e-175 | 503 | 501.901 | 253 | 339 | 1291 | 15 | 47.6 | 63.8 |
| C.neoformans.JEC21 | sterol 14-demethylase, putativ- e Cryptococcus neoformans var. neoformans JEC21 | 531 | 2.8e-169 | 531 | 488.804 | 238 | 351 | 1257 | 29 | 44.8 | 66.1 |
| C.neoformans.grubii.H99 | cytochrome P450, family 51 (st- erol 14-demethylase) Cryptococ- cus neoformans var. grubii H99 | 531 | 5.1e-170 | 531 | 490.73 | 240 | 351 | 1262 | 29 | 45.2 | 66.1 |
| C.neoformans.B.3501A | hypothetical protein CNBA0300 Cryptococcus neoformans var. n- eoformans B-3501A | 531 | 2.7e-169 | 531 | 488.804 | 238 | 351 | 1257 | 29 | 44.8 | 66.1 |
| C.parapsilosis | uncharacterized protein CPAR2 303740 Candida parapsilosis | 512 | 0 | 512 | 717.227 | 341 | 410 | 1850 | 7 | 64.2 | 77.2 |
| C.tropicalis | cytochrome P450 51 Candida tro- picalis MYA-3404 | 528 | 0 | 528 | 714.146 | 346 | 409 | 1842 | 13 | 65.2 | 77.0 |
| C.auris | lanosterol 14-alpha demethylas- e Candida auris | 510 | 0 | 510 | 742.651 | 343 | 420 | 1916 | 5 | 64.6 | 79.1 |
| C.albicans | sterol 14-demethylase Candida albicans SC5314 | 521 | 0 | 521 | 711.064 | 339 | 411 | 1834 | 9 | 63.8 | 77.4 |
| A.fumigatus | 14-alpha sterol demethylase Cy- p51B Aspergillus fumigatus Af2- 93 | 498 | 1.6e-180 | 498 | 516.924 | 249 | 352 | 1330 | 7 | 46.9 | 66.3 |

Table S41: Pairwise alignment info from yeast Erg11

#### S5.2 Erg24

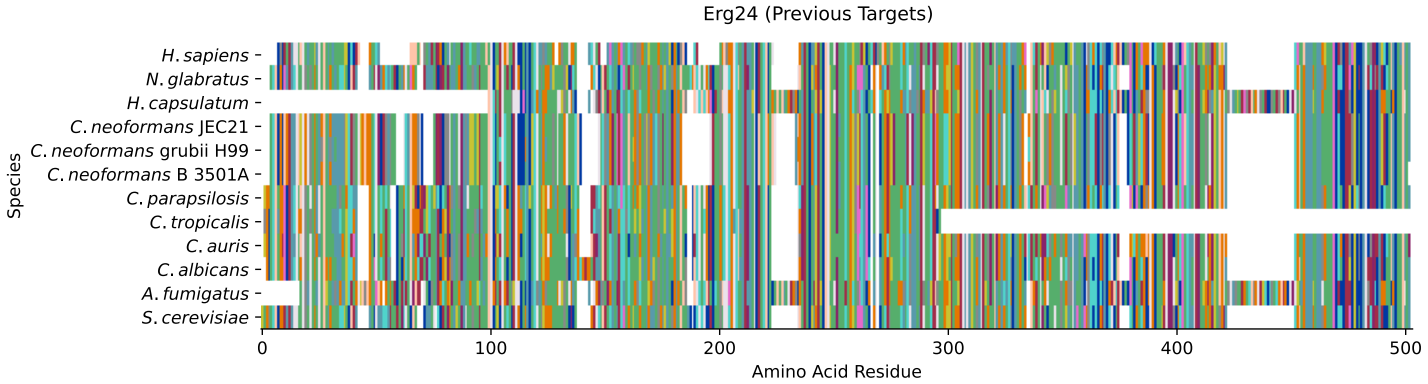


Figure S259: MSA for Erg24; cf. Table S42 for gene names and exact alignment values.

| Species | Hit Protein | Hit Length (a.a.) | evalue | align_len | bit_score | identity | positive | score | gaps | % identity | % positive |
|---|---|---|---|---|---|---|---|---|---|---|---|
| H.sapiens | delta(14)-sterol reductase LBR Homo sapiens | 442 | 1.3e-90 | 442 | 289.271 | 176 | 242 | 739 | 40 | 40.1 | 55.1 |
| N.glabratus | uncharacterized protein CAGL0I-02970g Nakaseomyces glabratus | 436 | 0 | 436 | 705.671 | 331 | 388 | 1820 | 2 | 75.4 | 88.4 |
| H.capsulatum | c-14 sterol reductase Histopla-sma capsulatum G186AR | 394 | 8.7e-88 | 394 | 275.018 | 146 | 228 | 702 | 49 | 33.3 | 51.9 |
| C.neoformans.JEC21 | C-14 sterol reductase, putativ-e Cryptococcus neoformans var. neoformans JEC21 | 448 | 6.3e-122 | 448 | 361.303 | 198 | 274 | 926 | 34 | 45.1 | 62.4 |
| C.neoformans.grubii.H99 | delta14-sterol reductase Crypt-ococcus neoformans var. grubii H99 | 448 | 6.1e-126 | 448 | 371.318 | 201 | 278 | 952 | 34 | 45.8 | 63.3 |
| C.neoformans.B.3501A | hypothetical protein CNBA1040 Cryptococcus neoformans var. n-eoformans B-3501A | 447 | 5e-121 | 447 | 358.992 | 197 | 273 | 920 | 34 | 44.9 | 62.2 |
| C.parapsilosis | uncharacterized protein CPAR2 405900 Candida parapsilosis | 444 | 5.3e-175 | 444 | 495.352 | 250 | 324 | 1274 | 10 | 56.9 | 73.8 |
| C.tropicalis | hypothetical protein CTRG 0186-9 Candida tropicalis MYA-3404 | 270 | 6.6e-88 | 270 | 267.314 | 139 | 177 | 682 | 4 | 31.7 | 40.3 |
| C.auris | delta(14)-sterol reductase Can-dida auris | 445 | 1.3e-173 | 445 | 491.886 | 256 | 326 | 1265 | 12 | 58.3 | 74.3 |
| C.albicans | delta(14)-sterol reductase Can-dida albicans SC5314 | 453 | 0 | 453 | 514.998 | 264 | 331 | 1325 | 25 | 60.1 | 75.4 |
| A.fumigatus | c-14 sterol reductase Aspergil-lus fumigatus Af293 | 479 | 1.3e-89 | 479 | 280.411 | 174 | 256 | 716 | 67 | 39.6 | 58.3 |

Table S42: Pairwise alignment info from yeast Erg24

#### S5.3 Erg2

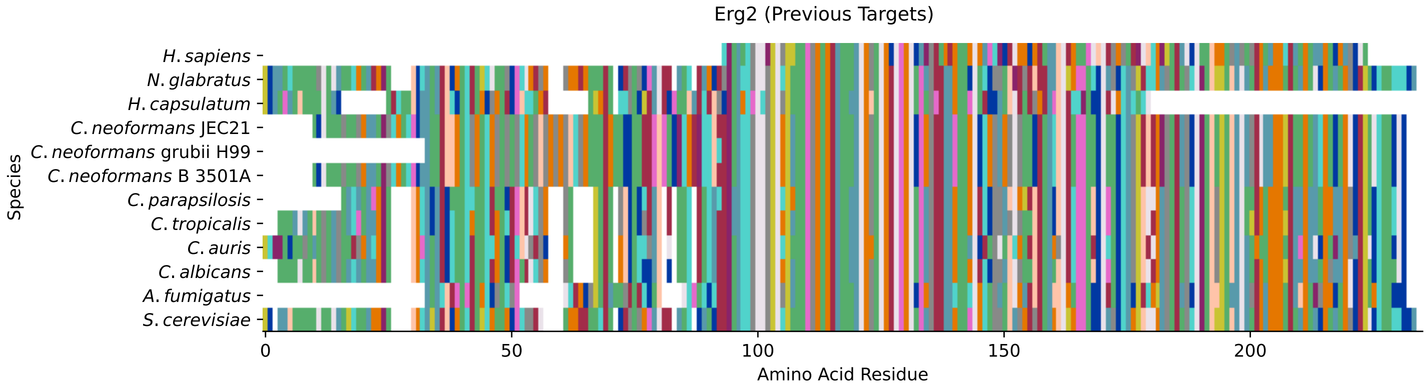


Figure S260: MSA for Erg2; cf. Table S43 for gene names and exact alignment values.

| Species | Hit Protein | Hit Length (a.a.) | evalue | align_len | bit_score | identity | positive | score | gaps | % identity | % positive |
|---|---|---|---|---|---|---|---|---|---|---|---|
| H.sapiens | sigma non-opioid intracellular receptor 1 isoform 8 Homo sapiens | 131 | 3.6e-23 | 131 | 94.7449 | 55 | 77 | 234 | 0 | 24.7 | 34.5 |
| N.glabratus | uncharacterized protein CAGL0L-10714g Nakaseomyces glabratus | 224 | 4.8e-127 | 224 | 356.295 | 170 | 193 | 913 | 2 | 76.2 | 86.5 |
| H.capsulatum | C-8 sterol isomerase Histoplasma capsulatum G186AR | 170 | 1.5e-39 | 170 | 134.42 | 73 | 104 | 337 | 10 | 32.7 | 46.6 |
| C.neoformans.JEC21 | C-8 sterol isomerase, putative Cryptococcus neoformans var. neoformans JEC21 | 225 | 9.2e-59 | 225 | 184.496 | 104 | 137 | 467 | 19 | 46.6 | 61.4 |
| C.neoformans.grubii.H99 | C-8 sterol isomerase Cryptococcus neoformans var. grubii H99 | 202 | 1.3e-58 | 202 | 184.111 | 99 | 126 | 466 | 13 | 44.4 | 56.5 |
| C.neoformans.B.3501A | hypothetical protein CNBA8120 Cryptococcus neoformans var. neoformans B-3501A | 225 | 8.9e-59 | 225 | 184.496 | 104 | 137 | 467 | 19 | 46.6 | 61.4 |
| C.parapsilosis | uncharacterized protein CPAR2 109890 Candida parapsilosis | 205 | 4e-85 | 205 | 249.98 | 129 | 144 | 637 | 4 | 57.8 | 64.6 |
| C.tropicalis | C-8 sterol isomerase Candida tropicalis MYA-3404 | 218 | 5.7e-81 | 218 | 239.58 | 127 | 147 | 610 | 8 | 57.0 | 65.9 |
| C.auris | C-8 sterol isomerase ERG2 Candida auris | 221 | 6.7e-79 | 221 | 234.187 | 116 | 157 | 596 | 5 | 52.0 | 70.4 |
| C.albicans | C-8 sterol isomerase Candida albicans SC5314 | 218 | 1.4e-83 | 218 | 246.128 | 128 | 152 | 627 | 8 | 57.4 | 68.2 |
| A.fumigatus | C-8 sterol isomerase (Erg-1), putative Aspergillus fumigatus Af293 | 192 | 1.7e-64 | 192 | 198.749 | 99 | 128 | 504 | 6 | 44.4 | 57.4 |

Table S43: Pairwise alignment info from yeast Erg2

#### S5.4 Fks1

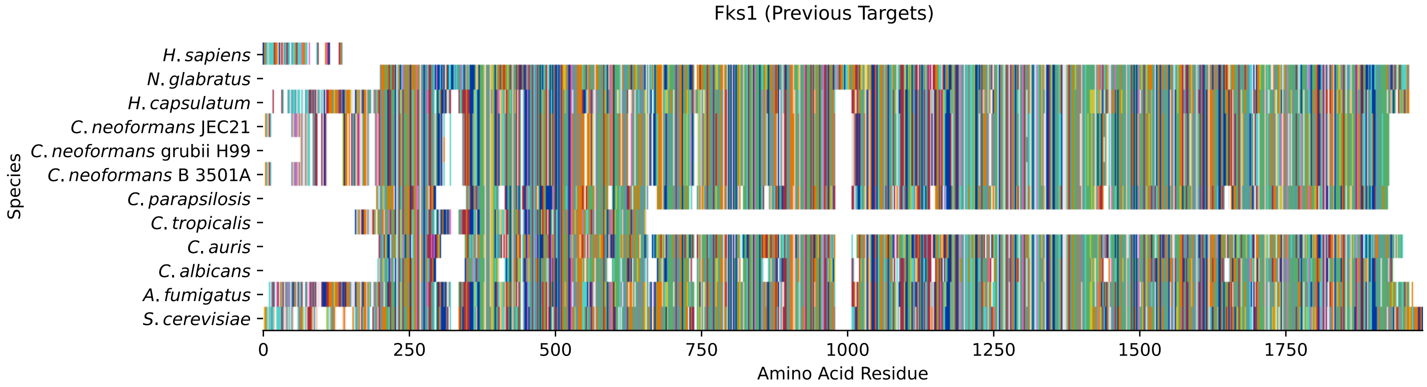


Figure S261: MSA for Fks1; cf. Table S44 for gene names and exact alignment values.

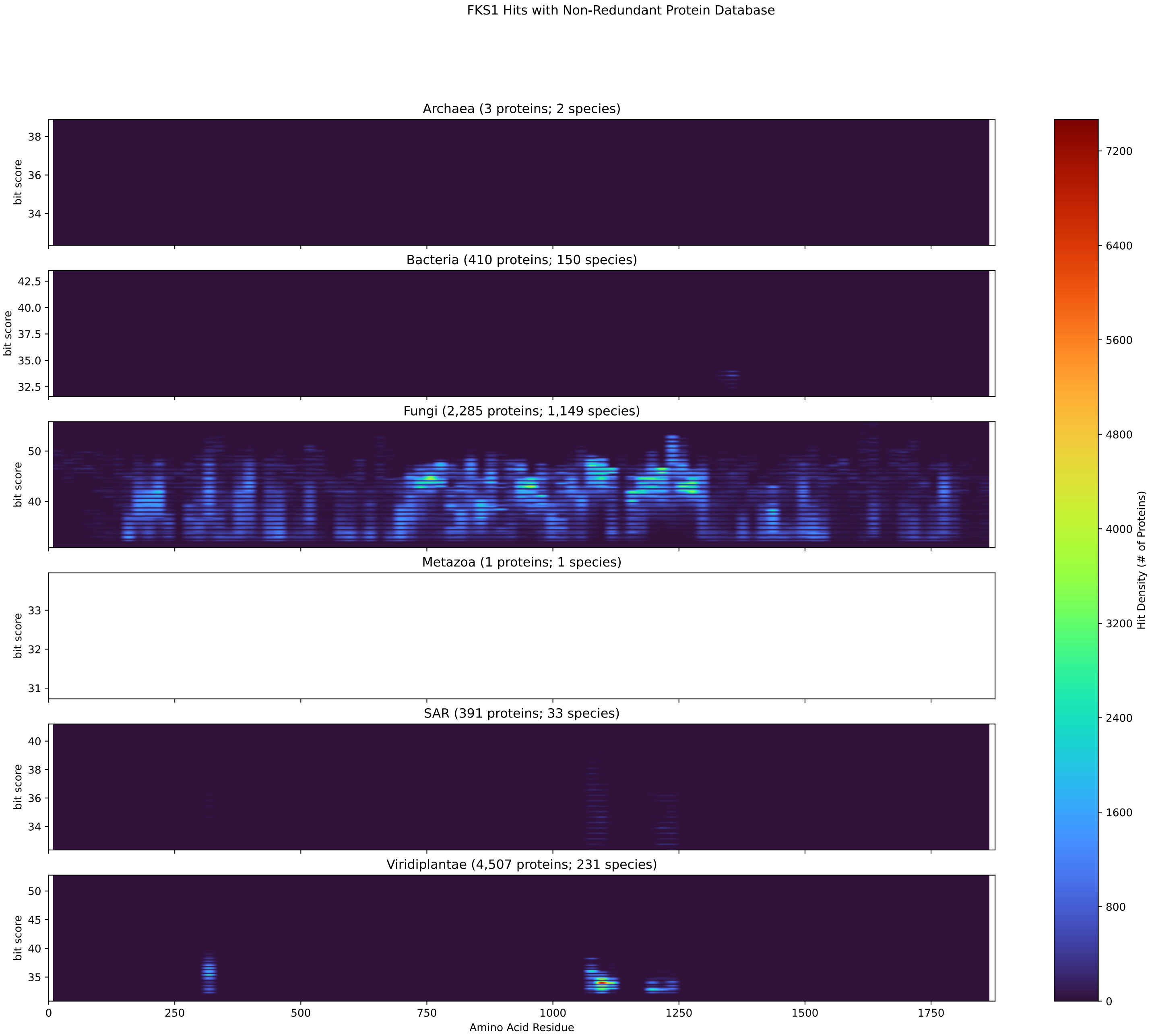


Figure S262: Histogram of Fks1 hits against NR in 20-aa windows

| Species | Hit Protein | Hit Length (a.a.) | evalue | align_len | bit_score | identity | positive | score | gaps | % identity | % positive |
|---|---|---|---|---|---|---|---|---|---|---|---|
| H.sapiens | TATA-binding protein-associated factor 2N isoform 2 Homo sapiens | 104 | 0.025 | 104 | 41.9726 | 32 | 47 | 97 | 20 | 1.7 | 2.5 |
| N.glabratus | uncharacterized protein CAGL0M-13827g Nakaseomyces glabratus | 1765 | 0 | 1765 | 1888.62 | 953 | 1257 | 4891 | 73 | 50.8 | 67.0 |
| H.capsulatum | glucan synthase Histoplasma capsulatum G186AR | 1879 | 0 | 1879 | 2468.34 | 1208 | 1449 | 6396 | 56 | 64.4 | 77.2 |
| C.neoformans.JEC21 | 1,3-beta-glucan synthase, putative Cryptococcus neoformans var. neoformans JEC21 | 1808 | 0 | 1808 | 2004.56 | 997 | 1271 | 5192 | 59 | 53.1 | 67.7 |
| C.neoformans.grubii.H99 | 1,3-beta-glucan synthase component FKS1 Cryptococcus neoformans var. grubii H99 | 1782 | 0 | 1782 | 1998.79 | 991 | 1260 | 5177 | 55 | 52.8 | 67.1 |
| C.neoformans.B.3501A | hypothetical protein CNBN2360 Cryptococcus neoformans var. neoformans B-3501A | 1808 | 0 | 1808 | 2005.33 | 997 | 1271 | 5194 | 59 | 53.1 | 67.7 |
| C.parapsilosis | uncharacterized protein CPAR2 109680 Candida parapsilosis | 1690 | 0 | 1690 | 1472.99 | 763 | 1085 | 3812 | 136 | 40.6 | 57.8 |
| C.tropicalis | 1,3-beta-glucan synthase component GLS1, partial Candida tropicalis MYA-3404 | 485 | 0 | 485 | 684.1 | 328 | 398 | 1764 | 8 | 17.5 | 21.2 |
| C.auris | hypothetical protein Candida auris | 1702 | 0 | 1702 | 1627.45 | 810 | 1137 | 4213 | 50 | 43.2 | 60.6 |
| C.albicans | Gsl1p Candida albicans SC5314 | 1689 | 0 | 1689 | 1486.09 | 783 | 1068 | 3846 | 139 | 41.7 | 56.9 |
| A.fumigatus | 1,3-beta-glucan synthase catalytic subunit FksP Aspergillus fumigatus Af293 | 1898 | 0 | 1898 | 2426.36 | 1212 | 1463 | 6287 | 64 | 64.6 | 77.9 |

Table S44: Pairwise alignment info from yeast Fks1

#### S5.5 Fks3

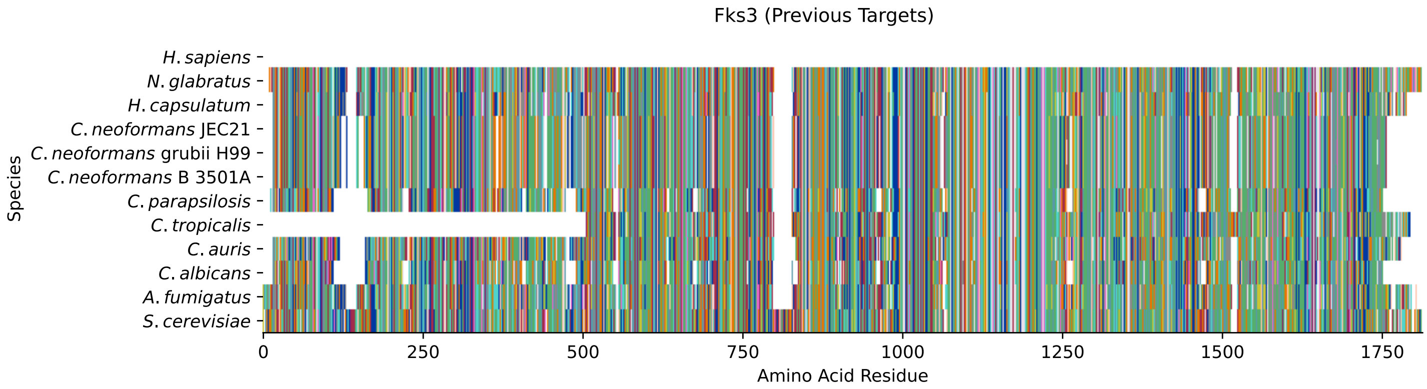


Figure S263: MSA for Fks3; cf. Table S45 for gene names and exact alignment values.

| Species | Hit Protein | Hit Length (a.a.) | evalue | align_len | bit_score | identity | positive | score | gaps | % identity | % positive |
|---|---|---|---|---|---|---|---|---|---|---|---|
| H.sapiens | - | - | - | - | - | - | - | - | - | - | - |
| N.glabratus | uncharacterized protein CAGL0K-04037g Nakaseomyces glabratus | 1787 | 0 | 1787 | 2006.88 | 980 | 1278 | 5198 | 63 | 54.9 | 71.6 |
| H.capsulatum | glucan synthase Histoplasma ca-psulatum G186AR | 1758 | 0 | 1758 | 2002.64 | 975 | 1254 | 5187 | 73 | 54.6 | 70.2 |
| C.neoformans.JEC21 | 1,3-beta-glucan synthase, puta-tive Cryptococcus neoformans v-ar. neoformans JEC21 | 1723 | 0 | 1723 | 1706.42 | 863 | 1149 | 4418 | 77 | 48.3 | 64.3 |
| C.neoformans.grubii.H99 | 1,3-beta-glucan synthase compo-nent FKS1 Cryptococcus neoform-ans var. grubii H99 | 1723 | 0 | 1723 | 1702.18 | 861 | 1143 | 4407 | 77 | 48.2 | 64.0 |
| C.neoformans.B.3501A | hypothetical protein CNBN2360 Cryptococcus neoformans var. n-eoformans B-3501A | 1723 | 0 | 1723 | 1706.03 | 864 | 1149 | 4417 | 77 | 48.4 | 64.3 |
| C.parapsilosis | uncharacterized protein CPAR2 109680 Candida parapsilosis | 1725 | 0 | 1725 | 1459.12 | 771 | 1083 | 3776 | 167 | 43.2 | 60.6 |
| C.tropicalis | 1,3-beta-glucan synthase compo-nent GLS1 Candida tropicalis M-YA-3404 | 1287 | 0 | 1287 | 1488.01 | 751 | 953 | 3851 | 51 | 42.0 | 53.4 |
| C.auris | hypothetical protein Candida a-uris | 1749 | 0 | 1749 | 1512.28 | 787 | 1111 | 3914 | 103 | 44.1 | 62.2 |
| C.albicans | Gsl1p Candida albicans SC5314 | 1720 | 0 | 1720 | 1500.34 | 792 | 1090 | 3883 | 175 | 44.3 | 61.0 |
| A.fumigatus | 1,3-beta-glucan synthase catal-ytic subunit FksP Aspergillus fumigatus Af293 | 1782 | 0 | 1782 | 1965.27 | 994 | 1271 | 5090 | 74 | 55.7 | 71.2 |

Table S45: Pairwise alignment info from yeast Fks3

#### S5.6 Gsc2

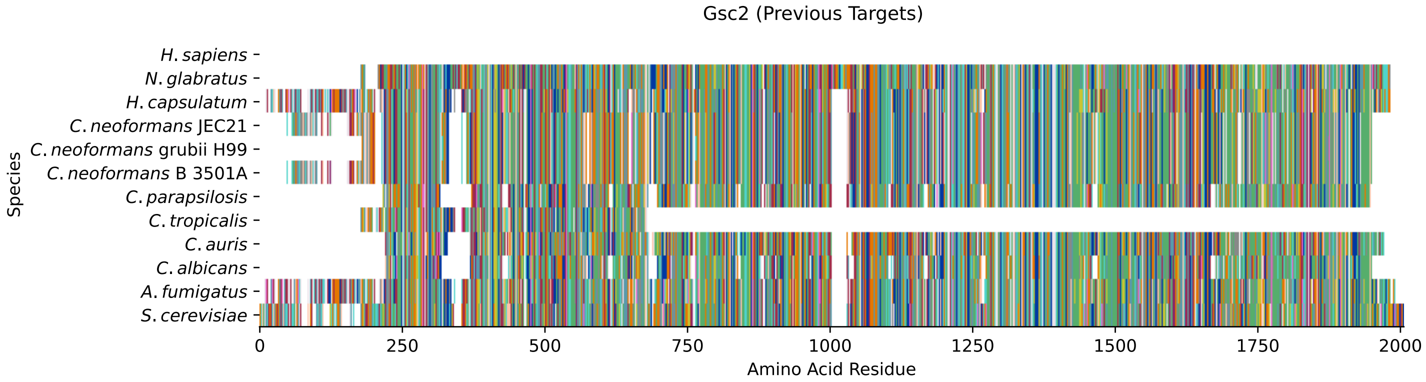


Figure S264: MSA for Gsc2; cf. Table S46 for gene names and exact alignment values.

| Species | Hit Protein | Hit Length (a.a.) | evalue | align_len | bit_score | identity | positive | score | gaps | % identity | % positive |
|---|---|---|---|---|---|---|---|---|---|---|---|
| H.sapiens | - | - | - | - | - | - | - | - | - | - | - |
| N.glabratus | uncharacterized protein CAGL0M-13827g Nakaseomyces glabratus | 1787 | 0 | 1787 | 1897.09 | 959 | 1266 | 4913 | 66 | 50.6 | 66.8 |
| H.capsulatum | glucan synthase Histoplasma ca-psulatum G186AR | 1905 | 0 | 1905 | 2468.73 | 1207 | 1450 | 6397 | 82 | 63.7 | 76.5 |
| C.neoformans.JEC21 | 1,3-beta-glucan synthase, puta-tive Cryptococcus neoformans v-ar. neoformans JEC21 | 1813 | 0 | 1813 | 2006.11 | 997 | 1266 | 5196 | 74 | 52.6 | 66.8 |
| C.neoformans.grubii.H99 | 1,3-beta-glucan synthase compo-nent FKS1 Cryptococcus neoform-ans var. grubii H99 | 1733 | 0 | 1733 | 2009.96 | 986 | 1243 | 5206 | 59 | 52.0 | 65.6 |
| C.neoformans.B.3501A | hypothetical protein CNBN2360 Cryptococcus neoformans var. n-eoformans B-3501A | 1813 | 0 | 1813 | 2006.11 | 996 | 1264 | 5196 | 74 | 52.5 | 66.7 |
| C.parapsilosis | uncharacterized protein CPAR2 109680 Candida parapsilosis | 1687 | 0 | 1687 | 1488.01 | 775 | 1094 | 3851 | 130 | 40.9 | 57.7 |
| C.tropicalis | 1,3-beta-glucan synthase compo-nent GLS1, partial Candida tro-picalis MYA-3404 | 489 | 0 | 489 | 678.322 | 331 | 391 | 1749 | 7 | 17.5 | 20.6 |
| C.auris | hypothetical protein Candida a-uris | 1704 | 0 | 1704 | 1632.08 | 812 | 1139 | 4225 | 54 | 42.8 | 60.1 |
| C.albicans | Gsl1p Candida albicans SC5314 | 1684 | 0 | 1684 | 1478.77 | 786 | 1069 | 3827 | 139 | 41.5 | 56.4 |
| A.fumigatus | 1,3-beta-glucan synthase catal-ytic subunit FksP Aspergillus fumigatus Af293 | 1914 | 0 | 1914 | 2419.81 | 1217 | 1461 | 6270 | 80 | 64.2 | 77.1 |

Table S46: Pairwise alignment info from yeast Gsc2

### S6 Genus-specific Good Targets

#### S6.1 Ccc1

##### S6.1.1 WHO Critical Pathogens

| Species | Hit Protein | Hit Length (a.a.) | evalue | align_len | bit_score | identity | positive | score | gaps | % identity | % positive |
|---|---|---|---|---|---|---|---|---|---|---|---|
| H.sapiens | - | - | - | - | - | - | - | - | - | - | - |
| N.glabratus | XP_445212.1 uncharacterized p-rotein CAGL0C00693g Nakaseomyc-es glabratus | 322 | 2.5e-133 | 322 | 379.793 | 224 | 256 | 974 | 23 | 69.6 | 79.5 |
| H.capsulatum | XP_045287029.1 CCC1 Histoplas-ma capsulatum G186AR | 246 | 1.4e-50 | 246 | 168.318 | 94 | 139 | 425 | 30 | 29.2 | 43.2 |
| C.neoformans.JEC21 | XP_568012.1 membrane fraction protein, putative Cryptococcu-s neoformans var. neoformans J-EC21 | 178 | 5.4e-22 | 178 | 93.5893 | 72 | 100 | 231 | 27 | 22.4 | 31.1 |
| C.neoformans.grubii.H99 | XP_012048745.1 membrane fract-ion protein Cryptococcus neofo-rmans var. grubii H99 | 177 | 4.8e-23 | 177 | 96.6709 | 73 | 100 | 239 | 25 | 22.7 | 31.1 |
| C.neoformans.B.3501A | XP_773602.1 hypothetical prot-ein CNBI2160 Cryptococcus neof-ormans var. neoformans B-3501A | 178 | 5.2e-22 | 178 | 93.5893 | 72 | 100 | 231 | 27 | 22.4 | 31.1 |
| C.parapsilosis | XP_036667882.1 uncharacterize-d protein CPAR2 405240 Candida parapsilosis | 325 | 3.1e-96 | 325 | 285.804 | 165 | 218 | 730 | 14 | 51.2 | 67.7 |
| C.tropicalis | XP_002547990.1 conserved hypo-thetical protein Candida tropi-calis MYA-3404 | 325 | 1.1e-99 | 325 | 294.278 | 169 | 216 | 752 | 18 | 52.5 | 67.1 |
| C.auris | XP_028891400.2 hypothetical p-rotein Candida auris | 264 | 1.2e-104 | 264 | 306.605 | 152 | 196 | 784 | 3 | 47.2 | 60.9 |
| C.albicans | XP_712749.1 Ccc1p Candida alb-icans SC5314 | 325 | 1.2e-95 | 325 | 283.878 | 165 | 212 | 725 | 18 | 51.2 | 65.8 |
| A.fumigatus | XP_754622.1 vacuolar iron tra-nsporter Ccc1, putative Asperg-illus fumigatus Af293 | 265 | 1.6e-60 | 265 | 194.512 | 116 | 156 | 493 | 13 | 36.0 | 48.4 |

Table S47: Pairwise alignment info from *S. cerevisiae* Ccc1 with all pathogens.

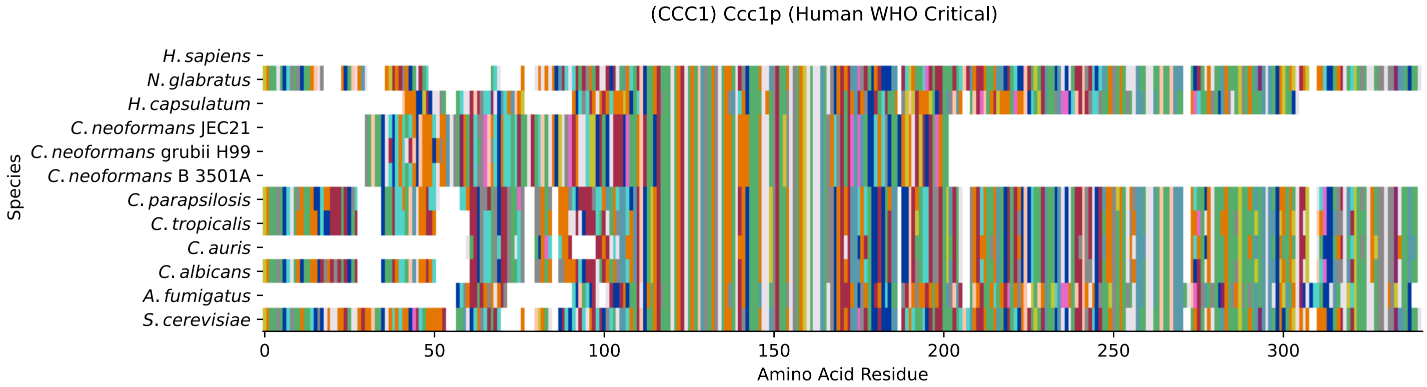


Figure S265: MSA of Ccc1, with all pathogens in the WHO Critical Pathogens group. Cf. Table S47 and Fig. S266.

#### Ccc1 MSA Quality

| | *A. f.* | *C. a.* | *C. a.* | *C. p.* | *C. t.* | *C. n.* | *C. n.* | *C. n.* | *H. c.* | *N. g.* | *S. c.* |
|---|---|---|---|---|---|---|---|---|---|---|---|
| *A. fumigatus* | 516.924 | 182.956 | 174.096 | 162.54 | 184.882 | 80.8777 | 81.2629 | 80.8777 | 223.016 | 157.918 | 166.777 |
| *C. albicans* | 189.504 | 620.928 | 377.481 | 452.981 | 542.732 | 85.1149 | 83.5741 | 85.1149 | 165.236 | 270.781 | 263.462 |
| *C. auris* | 174.096 | 341.658 | 521.161 | 379.793 | 357.836 | 85.1149 | 85.1149 | 85.1149 | 166.777 | 275.404 | 264.618 |
| *C. parapsilosis* | 178.718 | 442.965 | 405.986 | 630.172 | 473.011 | 82.0333 | 82.4185 | 82.0333 | 164.081 | 273.092 | 266.544 |
| *C. tropicalis* | 189.504 | 517.694 | 381.719 | 457.603 | 626.32 | 82.8037 | 83.1889 | 82.8037 | 158.303 | 269.626 | 267.7 |
| *C. neoformans* B 3501A | 93.9745 | 98.9821 | 102.449 | 98.9821 | 99.3673 | 349.362 | 311.612 | 349.362 | 95.5153 | 106.686 | 108.997 |
| *C. neoformans* grubii H99 | 94.3597 | 98.9821 | 102.834 | 99.3673 | 99.3673 | 311.227 | 348.591 | 311.227 | 95.9005 | 107.071 | 111.694 |
| *C. neoformans* JEC21 | 93.9745 | 98.9821 | 102.449 | 98.9821 | 99.3673 | 349.362 | 311.612 | 349.362 | 95.5153 | 106.686 | 108.997 |
| *H. capsulatum* | 223.016 | 160.999 | 166.777 | 164.081 | 157.147 | 82.8037 | 82.8037 | 82.8037 | 498.819 | 157.532 | 155.606 |
| *N. glabratus* | 186.037 | 285.034 | 309.686 | 285.419 | 289.271 | 90.5077 | 91.2781 | 90.5077 | 171.4 | 597.045 | 372.859 |
| *S. cerevisiae* | 194.512 | 284.263 | 306.22 | 286.189 | 294.278 | 92.8189 | 95.9005 | 92.8189 | 169.088 | 379.793 | 638.647 |

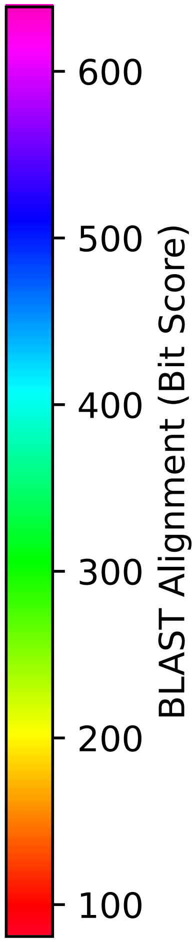


Figure S266: 2-D MSA of Ccc1, with all pathogens in the group. Cf. Table S47.

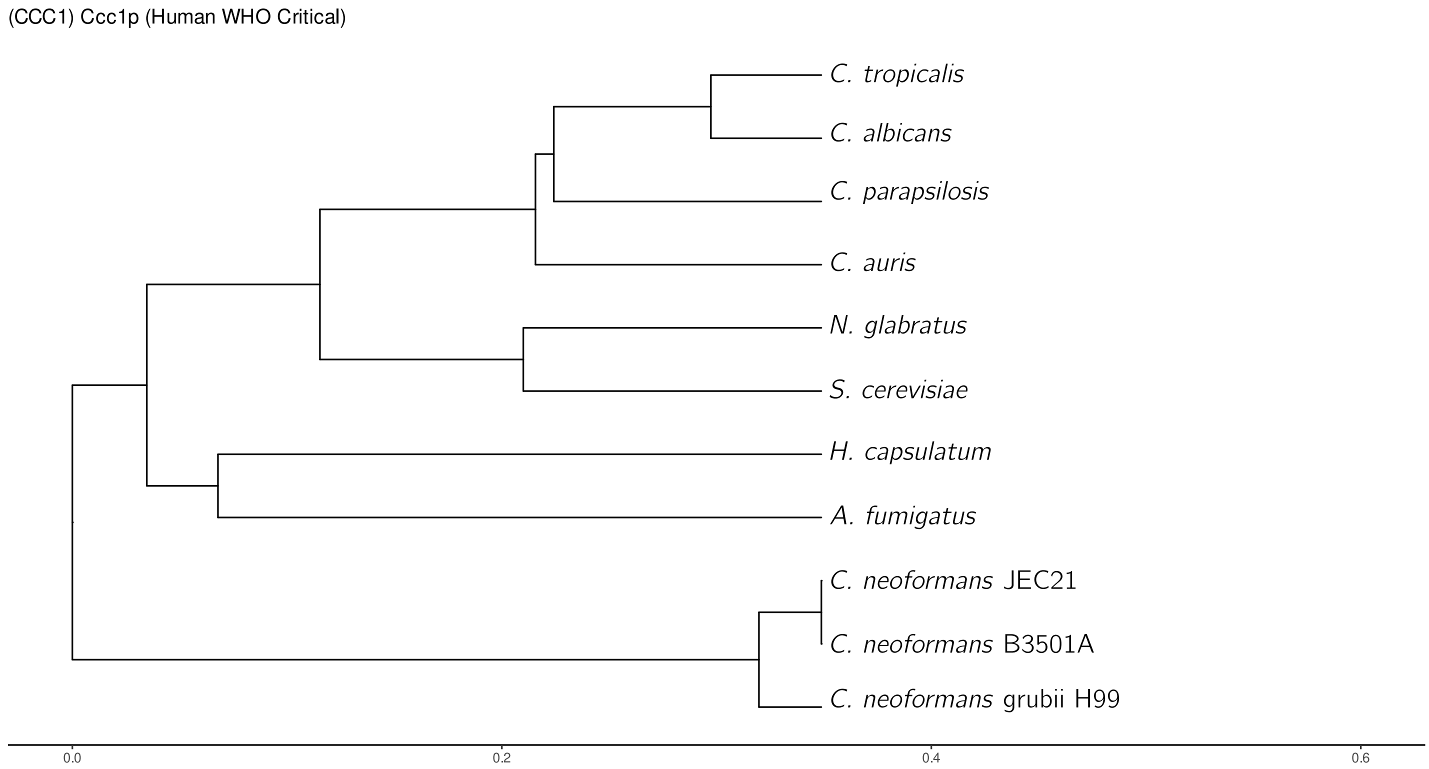


Figure S267: Phylogeny of Ccc1, with all pathogens in the group. Cf. Table S47, Figures S265 and S266. Plot was made using ggtree and ggplot2.

##### S6.1.2 Aspergillus

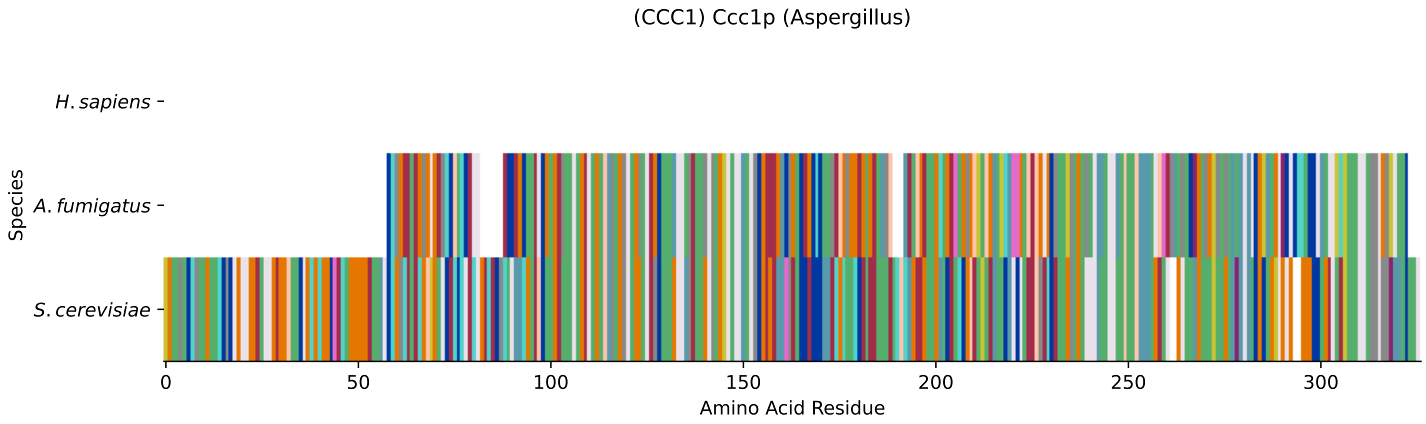


Figure S268: MSA of Ccc1 with only *Aspergillus*, with all genes in the group. Cf. Table S47 and Fig. S265.

##### S6.1.3 Candida

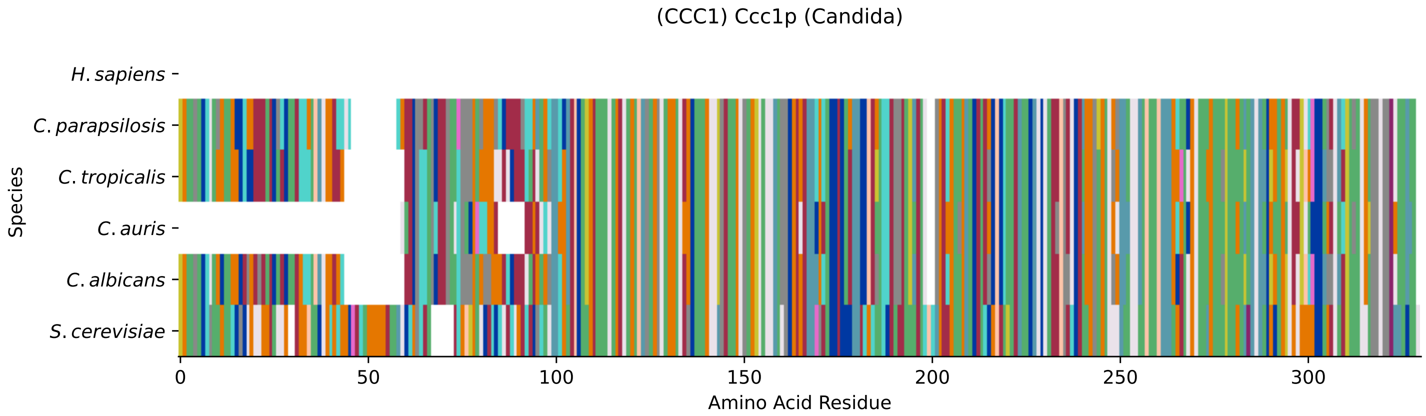


Figure S269: MSA of Ccc1 with only *Candida*, with all genes in the group. Cf. Table S47 and Fig. S265.

##### S6.1.4 Cryptococcus

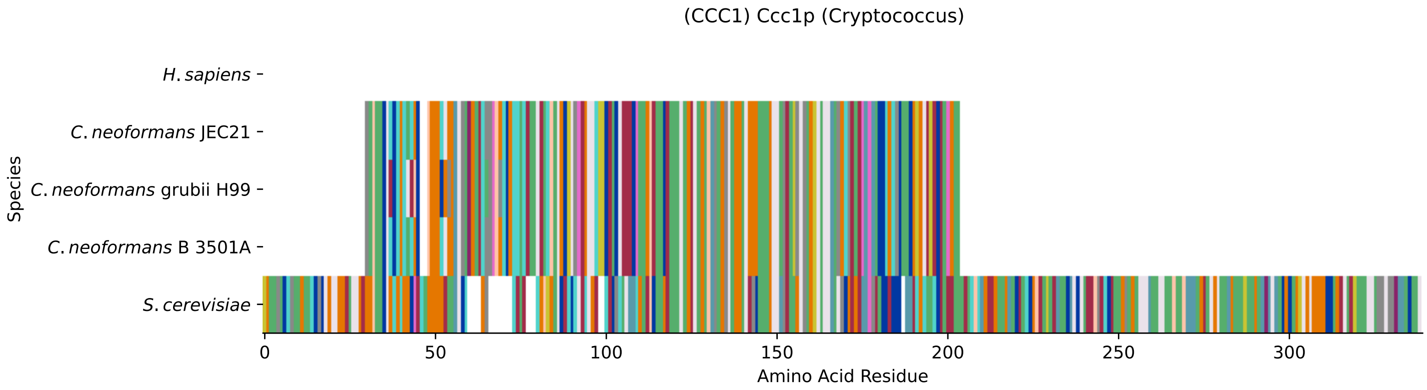


Figure S270: MSA of Ccc1 with only *Cryptococcus*, with all genes in the group. Cf. Table S47 and Fig. S265.

##### S6.1.5 Top 10 Agricultural Fungal Pathogens

| Species | Hit Protein | Hit Length (a.a.) | evalue | align_len | bit_score | identity | positive | score | gaps | % identity | % positive |
|---|---|---|---|---|---|---|---|---|---|---|---|
| G.max | XP_003530927.1 vacuolar iron transporter 1 Glycine max | 222 | 2.2e-48 | 222 | 163.31 | 85 | 136 | 412 | 9 | 26.4 | 42.2 |
| Z.mays | XP_008670169.1 vacuolar iron transporter 2 Zea mays | 211 | 6.6e-46 | 211 | 156.762 | 87 | 129 | 395 | 12 | 27.0 | 40.1 |
| S.tuberosum | XP_006355092.2 PREDICTED: vac- uolar iron transporter 1, part- ial Solanum tuberosum | 223 | 8.1e-45 | 223 | 154.836 | 86 | 132 | 390 | 13 | 26.7 | 41.0 |
| O.sativa | XP_015636127.1 vacuolar iron transporter 1.1 Oryza sativa J- aponica Group | 221 | 6.6e-48 | 221 | 161.77 | 86 | 136 | 408 | 9 | 26.7 | 42.2 |
| H.sapiens | - | - | - | - | - | - | - | - | - | - | - |
| U.maydis | XP_011388301.1 uncharacterize- d protein UMAG 06298 Ustilago maydis 521 | 287 | 1e-44 | 287 | 155.606 | 102 | 146 | 392 | 63 | 31.7 | 45.3 |
| P.striiformis | XP_047807231.1 hypothetical p- rotein Pst134EA 011404 Puccini- a striiformis f. sp. tritici | 231 | 9.1e-42 | 231 | 148.673 | 89 | 127 | 374 | 44 | 27.6 | 39.4 |
| P.oryzae | mRNA M BR32 EuGene 00028701-p1 — transcript=mRNA M BR32 EuGe- ne 00028701 — gene=M BR32 EuGe- ne 00028701 — organism=Pyricul- aria oryzae BR32 — gene produc- t=unspecified product — transc- ript product=unspecified produ- ct — location=BR32 scaffold000- 02:3526435-3527539(+) — protei- n length=338 — sequence SO=sup- ercontig — SO=protein coding g- ene — is pseudo=false | 196 | 4.7e-32 | 196 | 120.939 | 88 | 114 | 302 | 17 | 27.3 | 35.4 |
| P.triticina | XP_053019341.1 uncharacterize- d protein PtA15 4A235 Puccinia triticina | 273 | 4.7e-48 | 273 | 164.851 | 104 | 147 | 416 | 48 | 32.3 | 45.7 |
| P.graminis | XP_003334675.1 hypothetical p- rotein PGTG 16534 Puccinia gra- minis f. sp. tritici CRL 75-36- -700-3 | 301 | 9.7e-43 | 301 | 149.443 | 98 | 153 | 376 | 68 | 30.4 | 47.5 |
| M.graminicola | ZTRI 5.300.mRNA-p1 — transcrip- t=ZTRI 5.300.mRNA — gene=ZTRI 5.300 — organism=Zymoseptoria tritici IPO323 — gene product=- similar to vacuolar iron trans- porter Ccc1 — transcript produ- ct=similar to vacuolar iron tr- ansporter Ccc1 — location=Ztri chr 5:1094678-1095529(-) — pr- otein length=283 — sequence SO- =chromosome — SO=protein codin- g gene — is pseudo=false | 229 | 2.9e-57 | 229 | 185.652 | 104 | 138 | 470 | 6 | 32.3 | 42.9 |
| F.graminearum | XP_011327657.1 hypothetical p- rotein FGSG 07832 Fusarium gra- minearum PH-1 | 220 | 3.1e-49 | 220 | 164.466 | 91 | 135 | 415 | 5 | 28.3 | 41.9 |
| C.truncatum | XP_036587805.1 vacuolar iron transporter Colletotrichum tru- ncatum | 213 | 2.6e-58 | 213 | 189.889 | 108 | 136 | 481 | 9 | 33.5 | 42.2 |
| B.cinerea | XP_001546942.1 hypothetical p- rotein BCIN 11g00770 Botrytis cinerea B05.10 | 218 | 8.6e-47 | 218 | 159.458 | 112 | 139 | 402 | 8 | 34.8 | 43.2 |
| B.graminis | - | - | - | - | - | - | - | - | - | - | - |

Table S48: Pairwise alignment info from *S. cerevisiae* Ccc1 with all pathogens.

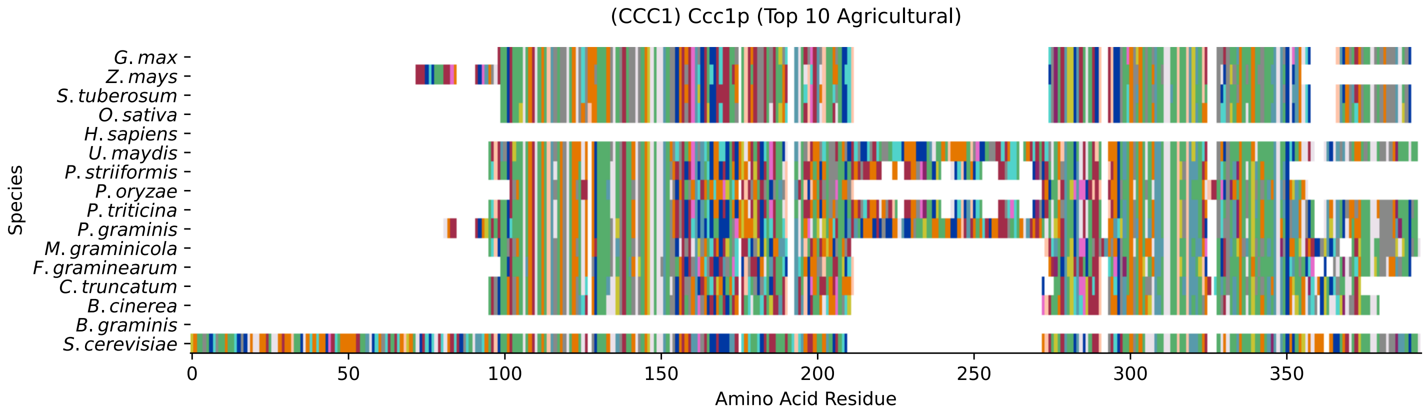


Figure S271: MSA of Ccc1, with all pathogens in the Top 10 Agricultural Fungal Pathogens group. Cf. Table S48 and Fig. S272.

#### Ccc1 MSA Quality

| | B.g. | B.c. | C.t. | F.g. | M.g. | P.g. | P.t. | P.o. | P.s. | U.m. | S.c. |
|---|---|---|---|---|---|---|---|---|---|---|---|
| *B. graminis* | 0 | 0 | 0 | 0 | 0 | 0 | 0 | 0 | 0 | 0 | 0 |
| *B. cinerea* | 0 | 433.335 | 228.024 | 145.976 | 180.644 | 116.316 | 133.65 | 167.933 | 128.642 | 128.257 | 171.4 |
| *C. truncatum* | 0 | 197.978 | 427.172 | 150.599 | 192.971 | 90.8929 | 132.88 | 207.608 | 127.487 | 87.8113 | 167.548 |
| *F. graminearum* | 0 | 117.087 | 150.599 | 437.187 | 149.828 | 138.658 | 141.354 | 97.4413 | 134.42 | 140.584 | 141.354 |
| *M. graminicola* | 0 | 145.976 | 192.971 | 149.828 | 453.751 | 108.997 | 124.02 | 154.066 | 117.857 | 125.176 | 158.303 |
| *P. graminis* | 0 | 85.1149 | 90.8929 | 96.2857 | 108.997 | 588.956 | 301.982 | 74.7146 | 239.195 | 232.646 | 124.79 |
| *P. triticina* | 0 | 103.605 | 132.88 | 141.354 | 124.02 | 301.982 | 546.584 | 82.0333 | 411.764 | 244.202 | 136.732 |
| *P. oryzae* | 0 | 171.4 | 235.728 | 128.257 | 185.267 | 91.2781 | 105.531 | 377.481 | 103.605 | 110.153 | 132.109 |
| *P. striiformis* | 0 | 102.064 | 134.42 | 137.887 | 120.168 | 248.054 | 439.499 | 82.0333 | 463.381 | 212.231 | 135.576 |
| *U. maydis* | 0 | 92.8189 | 87.8113 | 140.584 | 125.176 | 232.646 | 244.202 | 85.5001 | 204.912 | 565.074 | 124.79 |
| *S. cerevisiae* | 0 | 160.229 | 187.578 | 162.54 | 184.496 | 149.443 | 163.31 | 119.398 | 147.902 | 152.91 | 638.647 |

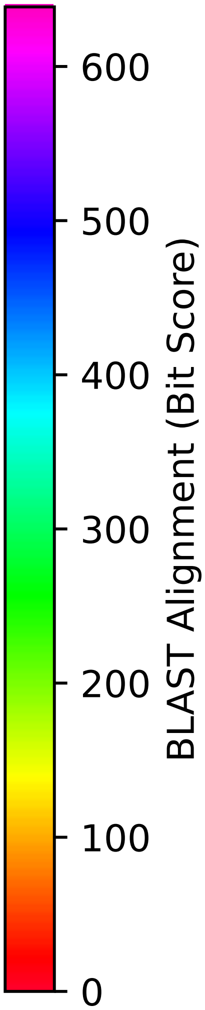


Figure S272: 2-D MSA of Ccc1, with all pathogens in the group. Cf. Table S48.
